# Pangenomics provides insights into the role of synanthropy in barn swallow evolution

**DOI:** 10.1101/2022.03.28.486082

**Authors:** Simona Secomandi, Guido Roberto Gallo, Marcella Sozzoni, Alessio Iannucci, Elena Galati, Linelle Abueg, Jennifer Balacco, Manuela Caprioli, William Chow, Claudio Ciofi, Joanna Collins, Olivier Fedrigo, Luca Ferretti, Arkarachai Fungtammasan, Bettina Haase, Kerstin Howe, Woori Kwak, Gianluca Lombardo, Patrick Masterson, Graziella Messina, Anders Pape Møller, Jacquelyn Mountcastle, Timothy A. Mousseau, Joan Ferrer-Obiol, Anna Olivieri, Arang Rhie, Diego Rubolini, Marielle Saclier, Roscoe Stanyon, David Stucki, Françoise Thibaud-Nissen, James Torrance, Antonio Torroni, Kristina Weber, Roberto Ambrosini, Andrea Bonisoli-Alquati, Erich D. Jarvis, Luca Gianfranceschi, Giulio Formenti

**Affiliations:** Department of Biosciences, University of Milan, Milan, Italy; Department of Biology, University of Florence, Sesto Fiorentino (FI), Italy; Vertebrate Genome Lab, The Rockefeller University, New York City, USA; Department of Environmental Sciences and Policy, University of Milan, Milan, Italy; Wellcome Sanger Institute, Cambridge, UK; Department of Biology and Biotechnology “L. Spallanzani”, University of Pavia, Pavia, Italy; DNAnexus Inc, USA; Hoonygen, Seoul, Republic of Korea; Hoonygen, Seoul, Korea; National Center for Biotechnology Information, National Library of Medicine, National Institutes of Health, Bethesda, MD 20894, USA; Ecologie Systématique Evolution, Université Paris-Sud, CNRS, AgroParisTech, Université Paris-Saclay, Orsay Cedex, France; Department of Biological Sciences, University of South Carolina, Columbia, SC, 29208, USA; Genome Informatics Section, Computational and Statistical Genomics Branch, National Human Genome, National Human Genome Research Institute, National Institutes of Health (Bethesda, Maryland, USA); Department of Developmental and Stem Cell Biology, Institut Pasteur/CNRS UMR3738, Cellular Plasticity and Disease Modelling, 25 Rue du Docteur Roux, 75015 Paris; Pacific Biosciences, Menlo Park, CA, USA; Department of Biological Sciences, California State Polytechnic University - Pomona, Pomona, CA, USA; The Rockefeller University (New York, NY, USA), Vertebrate Genome Laboratory, and HHMI

**Keywords:** Genome assembly, comparative genomics, pangenomics, genetic markers, positive selection, synanthropy

## Abstract

Insights into the evolution of non-model organisms are often limited by the lack of reference genomes. As part of the Vertebrate Genomes Project, we present a new reference genome and a pangenome produced with High-Fidelity long reads for the barn swallow *Hirundo rustica*. We then generated a reference-free multialignment with other bird genomes to identify genes under selection. Conservation analyses pointed at genes enriched for transcriptional regulation and neurodevelopment. The most conserved gene is *CAMK2N2*, with a potential role in fear memory formation. In addition, using all publicly available data, we generated a comprehensive catalogue of genetic markers. Genome-wide linkage disequilibrium scans identified potential selection signatures at multiple loci. The top candidate region comprises several genes and includes *BDNF*, a gene involved in stress response, fear memory formation, and tameness. We propose that the strict association with humans in this species is linked with the evolution of pathways typically under selection in domesticated taxa.

## Introduction

The association with anthropogenic environments includes different degrees of dependence, starting with synanthropy, when a species continues to live in areas occupied and altered by humans (1,2), and ending with domestication, when humans directly control selective pressures. Domestication has been extensively studied in birds (3) and mammals (4,5), where it has been linked to modifications of behavioural mechanisms, particularly a reduction in fear and reactive aggression responses and increased tameness, presumably related to alterations of specific physiological and developmental processes. Among these, neural crest cells development (6,7), corticosteroid hormones release (8) and other stress tolerance-related pathways, such as the glutamatergic signaling (9), are well-documented. Synanthropic species have adapted to exploit human environments without the need of an obligate dependency on anthropogenic resources (10). Typical adaptations are related to immune system response (11), resistance to pollutants (12–14), dietary (15), and behavioural changes (16). Because of its strong association with humans, the barn swallow (*Hirundo rustica*) is a well-suited model to investigate the evolution and genetic bases of behaviours correlated to synanthropy. The barn swallow is a well-studied (17–22) migratory passerine bird with six recognised subspecies in Europe, Asia, Africa and the Americas (23). While still poorly understood, recent studies have started to shed light on its genetics (18,24–29). The barn swallow demographic history reconstructed from genomic data suggests that the current barn swallow distribution derives from a relatively recent expansion, probably driven by the spread of human settlements providing more nesting opportunities (28,30). In the European subspecies, no evidence of population structure was observed, likely due to extensive gene flow between breeding populations (25). Studies on the barn swallow genomic architecture and adaptations have been limited by the lack of a highly contiguous, complete, and well-annotated reference genome for the species. The first reference genome, released in 2019 by our research group (31), was a scaffold-level assembly for the Eurasian subspecies generated combining PacBio long-read sequencing (32) and Bionano Direct Label and Stain (DLS) optical mapping (33). The second was a fragmented assembly for the same subspecies based on Illumina short reads and released in 2020 by the B10K Consortium (34,35). Here we present the first chromosome-level assembly for the Eurasian barn swallow *H. r. rustica*, generated using the Vertebrate Genomes Project (VGP) assembly pipeline (36), and the first pangenome for the species to expand the characterization of its intraspecific variation (37,38). With this assembly we identified conserved and accelerated genomic regions in the barn swallow genome, and generated a catalogue of genetic markers to detect high-linkage disequilibrium (LD) regions. Both approaches pointed at candidate genes known to be implicated in stress response, fear memory formation and vocal learning in songbirds (39). These processes are associated with tameness and domestication in birds (9), suggesting that the synanthropic habits of barn swallows could have evolved through similar selective pressures and pathways as those shaping the evolution of domestic taxa.

## Results and discussion

### A reference genome for the barn swallow

Using the VGP genome assembly pipeline v1.6 (36) (Additional file 1: Figure S1), we generated the first chromosome-level reference genome assembly (‘bHirRus1’ hereafter) and an alternative-haplotype assembly for the barn swallow. This included generating contigs with PacBio CLR long reads and scaffolding them with 10x linked reads, Bionano optical maps, and Hi-C reads. We also generated a mitochondrial genome for the species (Additional file 1: Figure S2, Supplementary Note). We sequenced a female, to obtain both Z and W sex chromosomes. After our manual curation (Supplementary Note) the primary assembly was 1.11 Giga base pairs (Gbp) long, with a scaffold NG50 of 73 Mega base pairs (Mbp) and a per-base consensus accuracy of Q43.7 (∼0.42 base errors/10 kilo base pairs, kbp; Additional file 1: Tables S1-2, see Supplementary Note for the extended evaluation). We assigned 98.2% of the assembled sequence to 39 autosomes and the Z and W sex chromosomes (Additional file 1: Figure S3a, Table S3), which are usually challenging to assemble due to their highly repetitive nature (40). The assembly exceeds the VGP standard metrics (6.7.Q40.C90) (36). The chromosome reconstruction (2n = 80) matches our cytogenetic analysis (Fig. 1a; Additional file 1: Supplementary Note), and is in line with the current literature on pachytene karyotypes of the barn swallow (41). Based on the original chicken chromosome classification (42) and our chromosome sizes (Additional file 1: Table S3, Supplementary Note), we define chromosomes 1-6 and Z as macrochromosomes, 7-13 and W as intermediate-size chromosomes, and 14-39 as microchromosomes. The size of the assembled chromosomes tightly correlates with the size of the chromosomes estimated from karyotype images (Spearman’s ρ = 0.99, n = 40, P < 2.2 × 10^−16^, Fig. 1b, Additional file 1: Table S4). As expected (36), PacBio long-reads coverage shows haploid coverage for Z and W (Fig. 1c track A). The total repeat content of the assembly is 271 Mbp (22.9%, Fig. 1c track B, Additional file 1: Table S3), in line with Genomescope2.0 (43) predictions (Additional file 1: Figure S4a, Table S5). The GC content is 42.5% (Fig. 1c track C, Additional file 1: Table S3). Functional gene completeness, measured with BUSCO (44), is 96% (Additional file 1: Figure S5a, Table S6).

**Fig. 1.**
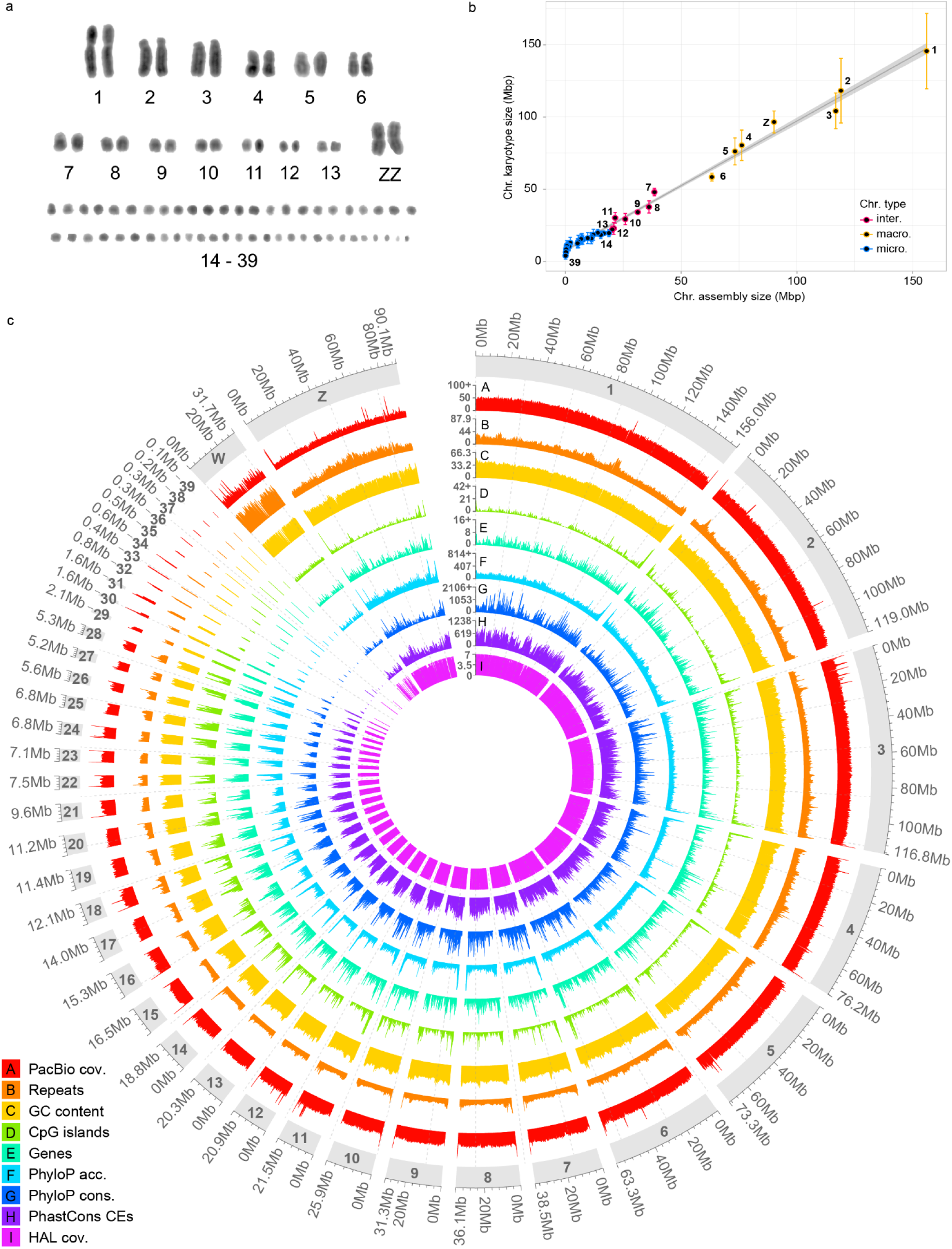
Karyotype reconstruction and chromosome-level assembly for the barn swallow. **a** DAPI-stained karyotype of a male Eurasian barn swallow (inverted colours). **b** Correlation between assembled chromosome length (x) and the estimated chromosome length from karyotype images (y). W sex chromosome is absent due to the sex of the karyotyped sample. **c** Circular representation of bHirRus1 chromosomes. All data are plotted using 200 kbp windows and the highest values were capped for visualisation whenever necessary (marked with +). PacBio long-read coverage (A); % repeat density (B); % GC (C); CpG islands density (D); gene density from NCBI annotation (E); accelerated sites density computed with phyloP (F); conserved sites density computed with phyloP (G); conserved elements (CEs) density computed from phastCons analysis (H); coverage of bHirRus1 in the Cactus HAL alignment, i.e. number of species aligned (I).

### Functional annotation

Newly-generated IsoSeq and RNAseq data, RNAseq data from other individuals (45) (Table S7), and protein alignments were used to guide the gene prediction process to generate the first NCBI RefSeq annotation for the species (NCBI *Hirundo rustica* Annotation Release 100). The NCBI Eukaryotic genome annotation pipeline (36,46) identified 18,578 genes and pseudogenes, of which 15,516 were protein coding. Among these, 15,130 (97.5%) aligned to UniProtKB/Swiss-Prot curated proteins, covering ≥ 50% of the query sequence, while 10,797 (69.6%) coding sequences aligned for ≥ 95%. In line with other birds (47), ∼52% of the total bp is annotated as genes, of which ∼90% are annotated as introns and ∼5% as coding DNA sequences (Additional file 1: Table S8).

### Chromosome size and genomic content

Differences in GC, CpG islands, gene and repeat content between bird macro-, intermediates and microchromosomes are likely the product of the evolutionary process that led to stable chromosome types in birds (48). Similarly to the zebra finch *Taeniopygia guttata (49)* genome, bHirRus1 chromosome size negatively correlates with GC content (Spearman’s ρ = -0.972, n = 40, P < 2.2 × 10^−16^), CpG island density (Spearman’s ρ = -0.925, n = 40, P < 2.2 × 10^−16^), gene density (Spearman’s ρ = -0.364, P < 2.5 × 10^−2^) and repeat density (Spearman’s ρ = -0.51, n = 40, P = 1.2 × 10^−3^; Fig 1c; Additional file 1: Figure S6). Indeed, microchromosomes are GC-rich (Wilcoxon test, W = 0, P = 1.4 × 10^−7^), CpG-rich (Wilcoxon test, W = 3, P = 2.2 × 10^−7^), gene-rich (Wilcoxon test, W = 94, P = 9.9 × 10^−3^) and repeat-rich (Wilcoxon test, W = 103, P = 2.0 × 10^−2^) with respect to the other types of chromosomes.

### Comparison between bHirRus1 and the previous scaffold-level assembly

Compared to the previous assembly (here after ‘Chelidonia’, scaffold NG50 26 Mbp; Additional file 1: Table S1), the VGP assembly pipeline and our subsequent manual curation increased the assembly contiguity to the chromosome level (see Supplementary Note for an extended comparison). Assembly QV is also considerably increased (43.7 vs 34; Additional file 1: Table S2). The repeat content decreased from 315 Mb to 271 Mb (Additional file 1: Figure S5c). BUSCO completeness increased in bHirRus1 (96% vs 95.9%), and BUSCO genes were less duplicated (0.8% vs 1.3%) and less fragmented (1.1% vs 1.2%; Additional file 1: Figure S5c, Table S6). We reconciled the larger size of Chelidonia (1.2 Gbp; Table S1) with the size of bHirRus1 (1.11 Gbp) by identifying 55 Mbp of repeats, sequence overlaps, low-coverage regions and haplotigs in Chelidonia (Additional file 1: Table S9, Supplementary Note).

### Reference-free whole-genome multiple species alignment and selection analysis

To identify regions under positive and negative (purifying) selection, we generated a reference-free, whole-genome multiple alignment using Cactus (50,51). The alignment included bHirRus1, six publicly-available chromosome-level Passeriformes genomes, and the chicken GRCg7b genome (Additional file 1: Figure S7, Table S10). Most of the species are synanthropic, domesticated or live partially in contact with humans. Overall, the coverage of the alignment with bHirRus1 was uniform in macrochromosomes, intermediate chromosomes, with the exception of chromosome W, and the largest microchromosomes (Fig. 1 track I; Additional file 1: Table S11). The mean alignability between all the species and the barn swallow was ∼76% (Additional file 1: Table S10). Using a 4-fold-degenerate sites neutral model and the Cactus alignment, we found that 0.96% of bHirRus1 bases are accelerated (i.e, evolve at higher rate than that under neutral evolution) and 2.71% are conserved (i.e, evolve at a lower rate) using phyloP with false-discovery rate (FDR) correction (52) (Fig. 1c track F-G; Additional file 1: Figure S8, Table S12). Approximately 52% and 63% of accelerated and conserved nucleotides, respectively, fell within genes (Additional file 1: Figure S8e, Table S12). Only ∼0.9% and ∼17% of accelerated and conserved bases overlapped with coding sequences (CDS), in line with previous studies (53,54). Using phastCons (55) and an *ad hoc* parameters set (coverage and smoothing), we identified ∼3 million conserved elements (CEs) covering 12.3% of the barn swallow genome (133 Mbp; Fig. 1c track H; Additional file 1: Table S12). Similarly to the phyloP analysis, significant overlaps were observed between CEs and genes (∼61%), with ∼14% of CEs overlapping CDS (Additional file 1: Figure S8e, Table S12), as expected (53,54). While conserved sites density was weakly positively correlated with chromosome sizes (Spearman’s ρ = 0.35, n = 40, P < 3.4 × 10^−2^), without significant differences between chromosome classes (Wilcoxon test, W = 244, P = 0.189), accelerated sites density was strongly negatively correlated with chromosome size (Spearman’s ρ = -0.80, n = 40, P < 9.5 × 10^−8^), with microchromosomes richer in accelerated sites than the other chromosomes (Wilcoxon test, W = 50, P = 4.6 × 10^−5^), as already observed in other birds (56). The Gene Ontology (GO) analysis on the top 5% genes with highest overlapping with phyloP accelerated sites (Additional file 1: Table S13) did not disclose any enriched GO term (Additional file 1: Table S14, Supplementary Note). PhyloP conserved sites showed a highly significant positive correlation with CEs detected with phastCons (Spearman’s ρ = 0.83, n = 108010, P < 2.2 × 10^−16^; Fig. 1c). Since phyloP sites can be considered a higher confidence subset within the larger phastCons set, we focussed our subsequent analyses on phyloP sites. As expected, CDS were the most conserved (57) (Additional file 1: Figure S8c, Table S12). The GO analysis on the top 5% genes with highest overlapping between CDS and phyloP conserved bases (Additional file 1: Table S15) revealed an enrichment for genes involved in DNA-binding, transcriptional regulation and nervous system development (Additional file 1: Table S16). The top 20 genes were largely involved in neural development and differentiation (Additional file 1: Table S15, Supplementary Notes). The top candidate was *CAMK2N2* (89% CDS bases conserved; Additional file 1: Table S15), located on chromosome 10. In the Cactus alignment, in correspondence with its CDS coordinates, all the species have the same base composition, with the exception of the chicken, which has a few SNPs (Additional file 1: Figure S9). PhyloP conserved bases were located only in regions without SNPs, while phastCons CEs comprise also regions which are not fully conserved between all species. *CAMK2N2* encodes a protein that acts as an inhibitor of calcium/calmodulin-dependent protein kinase II (*CAMKII*). *CAMKII* has a vital role in long-term potentiation of synaptic strength (LTP) and learning, via regulation of glutamate receptors (AMPA) (58–62). *CAMKII* is also one of the main calcium/calmodulin targets after the activation of NMDA (N-methyl-d-aspartate) glutamate receptors, which are involved in memory formation (63). Moreover, a peptide derived from *CAMK2N2 (*tatCN21) impairs fear memory formation by blocking *CAMKII* activity (64), and overexpression of *CAMK2N2* in the hippocampus was found involved in memory formation (65). In the Bengalese finch *Lonchura striata domestica (9)*, one of the species included in the Cactus alignment, the glutamatergic system contributed to the attenuation of stress response and aggressive behaviour under domestication. Finally, in high stress lines of the domesticated Japanese quail *Coturnix japonica, CAMK2N2* and *CAMKII* have been detected as deleted, together with other genes in the same networks, compared with low stress lines (66,67). Loss of genes in this network may be responsible for the reduced growth rate and low basal weight of the high stress quails (67). Since *CAMK2N2* is likely involved in behavioural and physiological changes under domestication in birds, we evaluated it in relation to the onset of synanthropic habits in the barn swallow. We generated an alignment of transcripts from 38 species (17 domesticated or synanthropic, 21 wild; Additional file 1: Table S17). However, we did not observe any pattern specific to domesticated or synanthropic species, and the single-gene phylogenetic tree substantially matched the known phylogeny. Thus, any role of *CAMK2N2* in synanthropic habits or domestication would have to be ascribed to non-coding regulatory elements. In vocal learning bird species, domestication was also found involved in the control of dopaminergic signalling in neural circuits that are crucial for vocal learning (9). Among the top 20 genes with the most overlap between CDS and phyloP conserved bases (Additional file 1: Table S15), *FOXP2* has 74% of its CDS bases conserved. This gene received great attention for its role in language and speech, since mutations in its sequence cause, among others, speech impairments (68–73). In the zebra finch, a vocal learner like the barn swallow, this gene has a marked expression in brain regions involved in song learning (74–77). Another candidate gene detected and previously associated with song learning is *UBE2D3* (75% CDS conserved; Additional file 1: Table S15), a gene located in a region of the human genome associated with musical abilities (78–80), which include recognizing, reproducing and memorising sounds. *CAMK2N2, FOXP2* and *UBE2D3* were also in the top 5% genes with the most overlaps between CDS and CEs bases detected with phastCons (Additional file 1: Table S18).

### Towards a pangenome for the barn swallow

Despite the high resolution achieved with chromosome-level assemblies, population genomic studies based on traditional linear reference genomes face limitations when aiming to describe complete variation among individuals (81). To reduce bias towards a single reference genome, we generated high-coverage (∼15-30x) HiFi whole-genome sequencing (WGS) data for five additional *H. r. rustica* individuals (Additional file 1: Tables S19-S20), assembled them with Hifiasm (82), and used both primary and alternate haplotypes (Additional file 1: Table S21) to generate the first pangenome variation graph (37,38) for the species (Fig. 2, Additional file 1: Figure S10). The HiFi-based primary assemblies had a contig NG50 between 2-8.6 Mbp, while the alternate between 0.2-1.7 Mbp, proportional to sequencing coverage (Additional file 1: Tables S20; S21). All primary assemblies shared 90.5% of their sequence (core genome), while all the HiFi individuals, considering both primary and alternate, shared 92.6% of bHirRus1 genes (Fig. 2c-d; Additional file 1: Table S22). 1.36% (236) of bHirRus1 annotated genes were not found in the HiFi assemblies (Fig. 2d; Additional file 1: Tables S22-23). Of those genes, 79 were found in the raw-reads of at least one individual for > 80% of their sequence with > 99% identity (Additional file 1: Table S23). The absence of the remaining 157 genes (0.87%) from both HiFi raw reads and HiFi-based assemblies, may either be due to the known GA dropout in HiFi reads (83), or to real gene losses in those individuals.

**Fig. 2.**
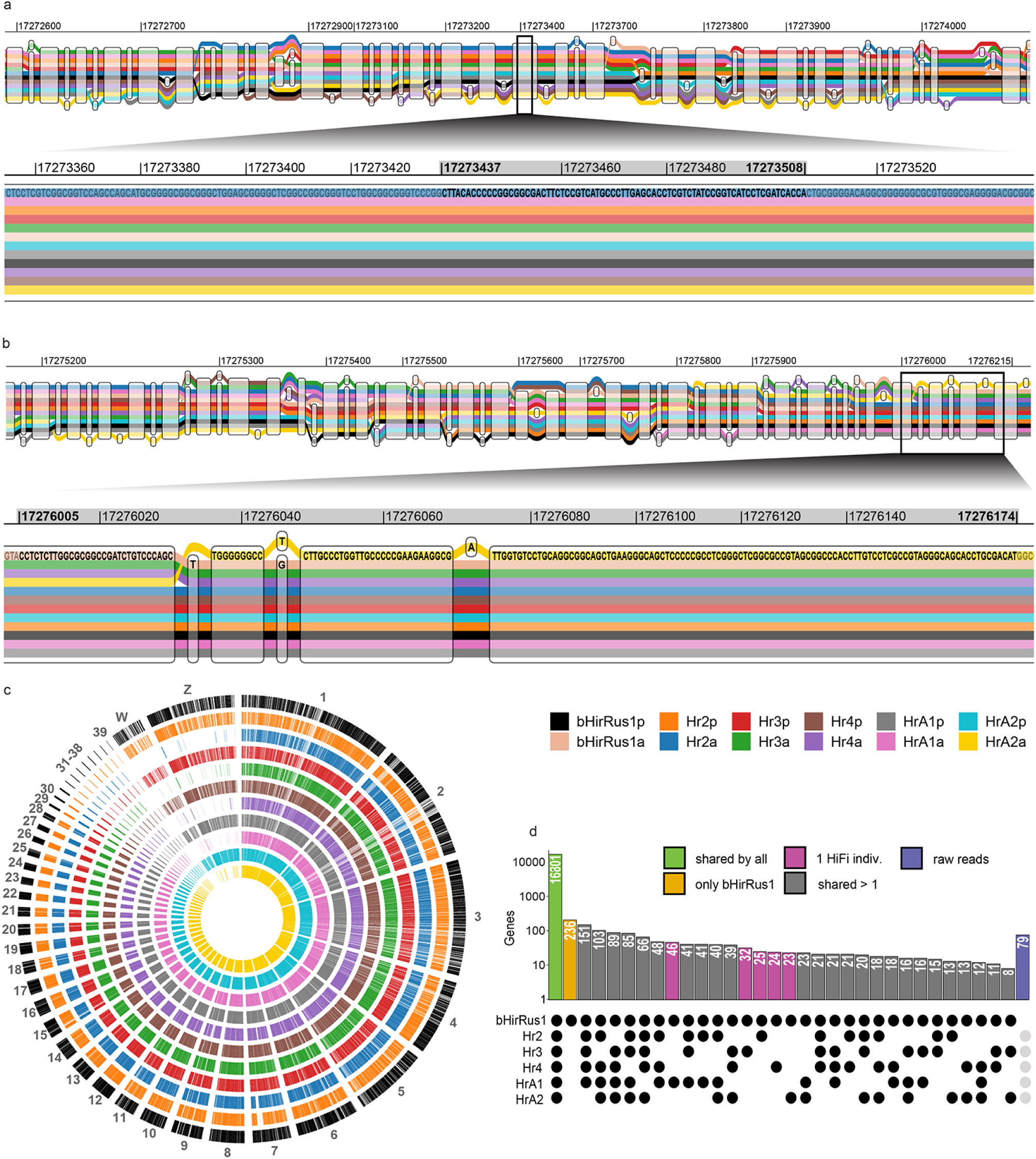
The first pangenome for the barn swallow. **a** *CAMK2N2* initial region in the barn swallow pangenome. bHirRus1 Chr10 (‘bHirRus1p’) is shown together with the alternate assembly ‘bHirRus1a’, the five HiFi-based primary (Hr2p, Hr3p, Hr4p, HrA1p, HrA2p), and their alternate assemblies (Hr2a, Hr3a, Hr4a, HrA1a, HrA2a). The zoomed part shows the first CDS (grey rectangle, 17,273,437-17,273,508). **b** *CAMK2N2* terminal region. The zoomed part shows the details of the second CDS (grey rectangle, 17,276,005-17,276,174). **c** Circos plot showing the annotated genes of bHirRus1 (primary, black) and orthologs in the other individuals (primary and alternate combined). Tracks follow the same colour legend as a and b. **d** Presence or absence of bHirRus1 genes in the other individuals included in the pangenome. The histogram reports the number of genes shared between bHirRus1 (primary) and each of the other individuals or groups of individuals (primary and alternate assemblies combined). The majority of the genes are shared between all individuals (green), while only 236 genes are exclusive of bHirRus1 (yellow). Genes shared only between bHirRus1 and another individual are shown in purple. The remaining bHirRus1 genes were found in 2 or more individuals (grey). Seventy-nine out of the 236 genes exclusive of bHirRus1 were found with BLAST (84) in at least one individual HiFi reads (violet), and therefore not properly assembled in the HiFi-based assemblies.

### Marker catalogue and genome-wide density

In parallel to our phylogenomic analyses, we used bHirRus1 as reference and our high coverage HiFi WGS dataset (ds1, ∼20x coverage, N = 5) to generate a comprehensive catalogue of single-nucleotide polymorphisms (SNPs; Additional file 1: Supplementary Note). We complemented this information with all the publicly available genomic data for the species (Additional file 1: Figure S11, Table S24), including two Illumina WGS datasets (28,29) (ds2 and ds3.1, ∼6.8x, N = 159) and four ddRAD datasets (24,25,27,28) (ds3.2 through ds6, ∼0.07x; N = 1,162). Despite the fewer individuals in HiFi WGS, the average SNP density and distribution (Fig. 3, light blue track; 142.37 SNPs/10 kbp; Additional file 1: Table S25) was comparable to the one computed for Illumina WGS (Fig. 3, dark blue track; 160.34 SNPs/10 kbp; Additional file 1: Table S25), suggesting that this sequencing method yields a high and accurate reads mappability even when only small datasets are available. We also performed a coverage titration experiment (Additional file 1: Supplementary Note) and found that SNP distribution was still uniform across chromosomes even when HiFi WGS were downsampled to 5x (96.33 SNPs/10 kbp; Additional file 1: Figure S12, Table 25). Chromosome W showed the lowest SNP density among all chromosomes (HiFi WGS 3.16 SNPs/10 kbp; HiFi WGS 5x 1.01 SNPs/10 kbp; Illumina WGS 1.38 SNPs/10 kbp), in line with the fact that chromosome W is present as single copy only in females (the heterogametic sex), and it has the highest content of heterochromatin and repeat elements, hindering variant calling (85). In contrast, we identified a higher number of SNP markers on chromosome Z (HiFi WGS 31.8 SNPs/10 kbp; HiFi WGS 5x 2.34 SNPs/10 kbp; Illumina WGS 53.3 SNPs/10 kbp). As expected, ddRAD exhibited very localised peaks of SNP density (0.8 SNPs/10 kbp; Fig. 3, red track). Particularly, ddRAD identified an extremely low number of SNPs on chromosome Z (0.27 SNPs/10 kbp) and no SNPs on microchromosome 33 (Additional file 1: Figure S13). Opposed to previous findings in humans (86,87), we detected a positive correlation between chromosome GC content and SNP density in all datasets (Additional file 1: Supplementary Note).

**Fig. 3.**
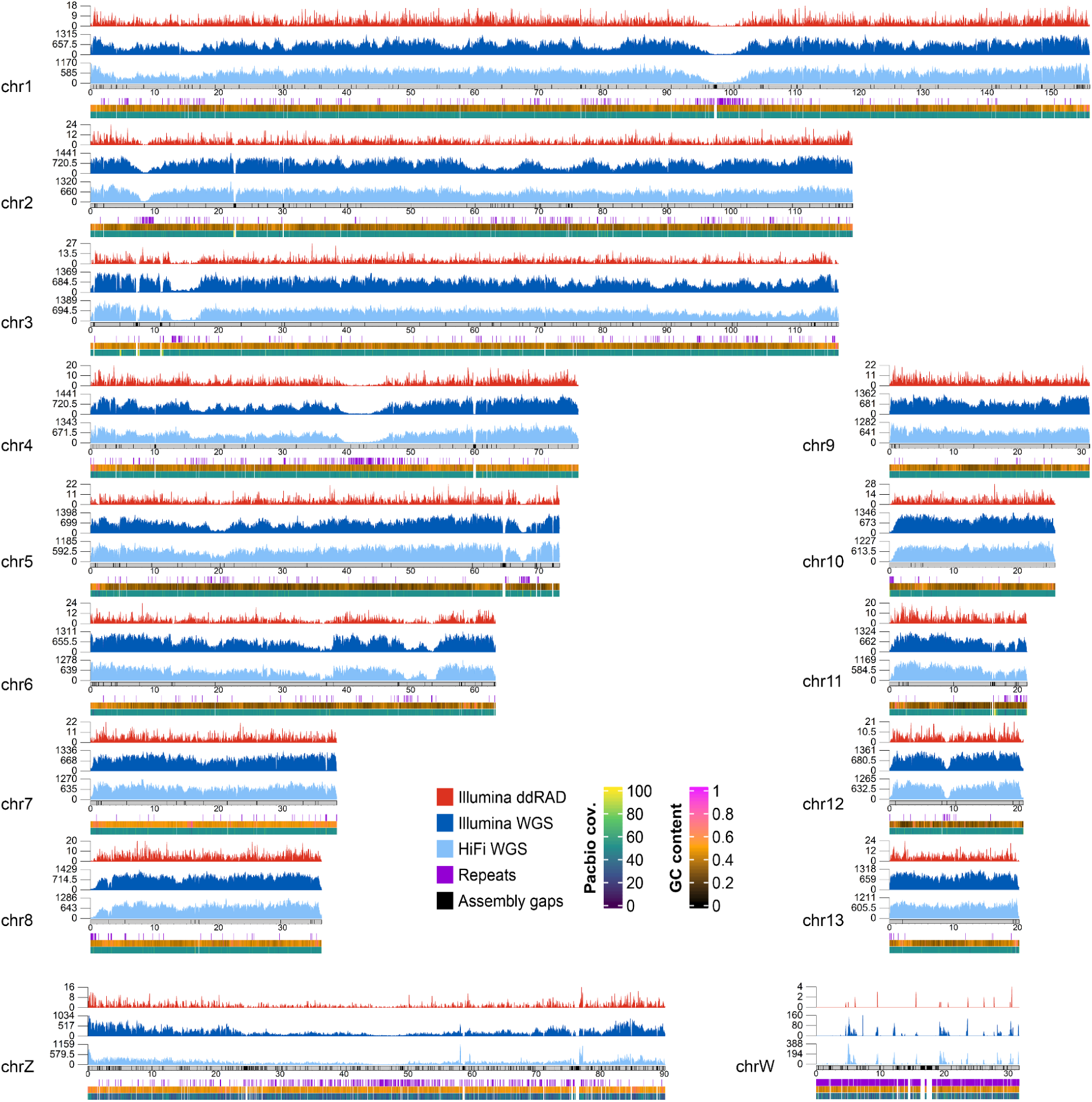
SNPs density per chromosome. Macrochromosomes and intermediate chromosomes are shown here, while microchromosomes are shown in Extended Data Fig. 7. SNP density, coloured according to the different types of genomic data used, was computed over 40 kbp windows. The numbers on the y axis of each density track indicate the maximum and average values of SNP density for each track. Light blue: HiFi WGS data (ds1). Dark blue: Illumina WGS data from ds2 and ds3.1. Red: Illumina ddRAD data from ds3.2 through ds6.8. All available samples from the same sequencing technology were considered together. Additional tracks in the lower panel show repetitive regions of the genome (violet bars; only regions larger than 3 kbp are plotted), GC content and PacBio reads coverage. Grey ideograms represent chromosomes in scale, with assembly gaps highlighted as black bars.

### Genome-wide linkage disequilibrium

LD reflects the evolutionary history of populations as it can be influenced by selective pressures (88–90), recombination rate (91,92), migration (93), genetic drift (94) and population admixture (95,96). Assessing its decay is pivotal to the success of genome-wide association studies (GWAS) (97,98) because it provides an estimate of the number of molecular markers required to detect significant associations between markers and causative loci. Since WGS is usually very useful to describe LD patterns (92), and no previous study estimated genome-wide LD decay in the barn swallow, we evaluated LD using the SNPs in our catalogue derived from WGS (ds2 and ds3.1). Genome-wide average r^2^ varied between *H. rustica* subspecies (Fig. 4a, Additional file 1: Table S26). As expected (99), absolute r^2^ decreased with increasing sample size and marker distance (*H. r. savignii, H. r. erythrogaster, H. r. transitiva*; Fig. 4a, Additional file 1: Table S26). Overall, our results indicate that the genetic association between loci in the barn swallow is extremely low and decreases rapidly within the first 10 kbp, as expected in large panmictic populations (25,100). Average r^2^ at increasing distance varied also across chromosome types, confirming that avian microchromosomes are characterised by higher rates of meiotic recombination, and thus lower LD, than macrochromosomes (Fig. 4b; Additional file 1: Table S27) (48,101,102).

**Fig. 4.**
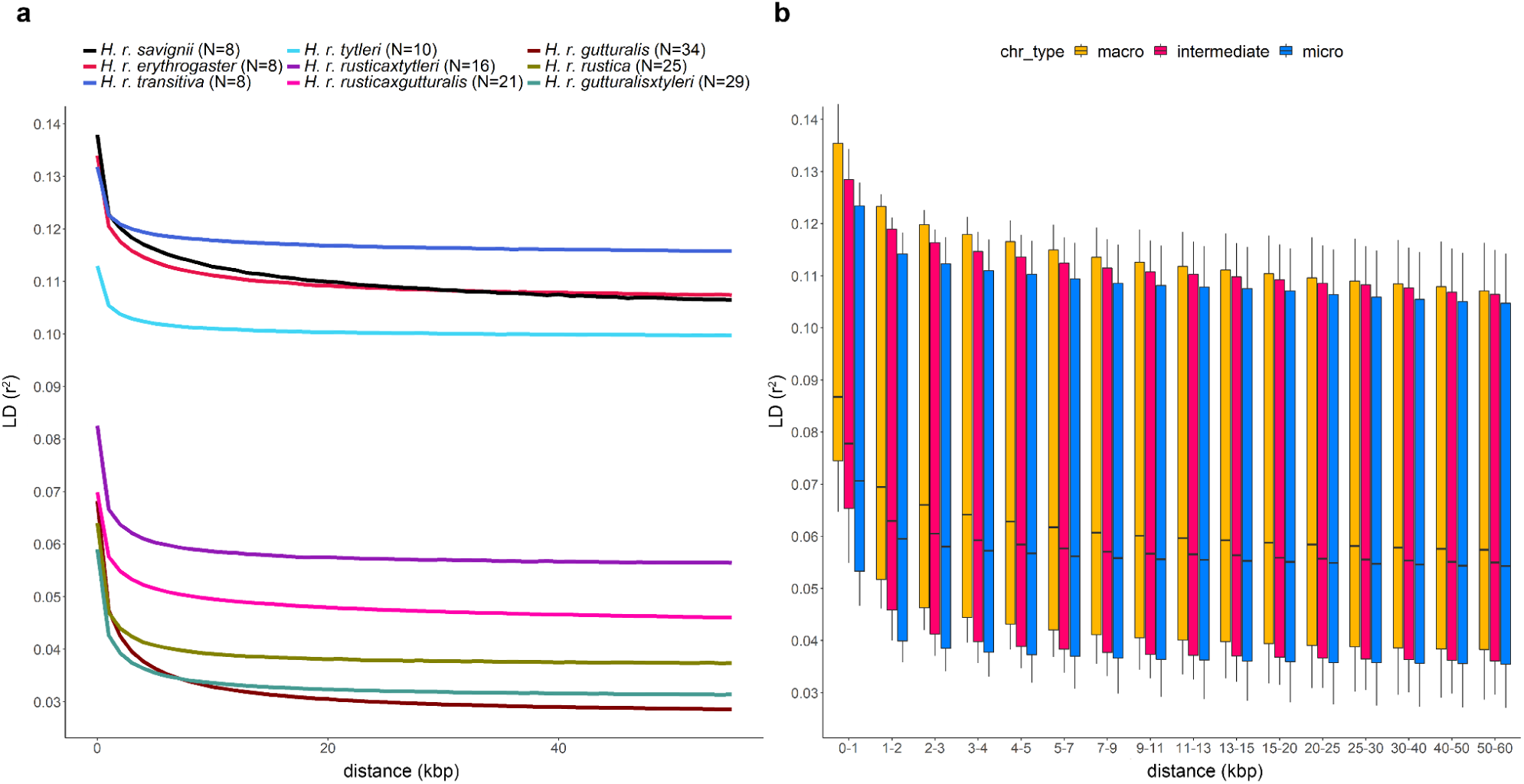
Linkage disequilibrium decay in the barn swallow genome. **a** Average r^2^ values plotted against physical distance (kbp) for the different populations belonging to ds2 and ds3.1 (Illumina WGS data). **b** Average r^2^ values in macrochromosomes, intermediate chromosomes, and microchromosomes according to pairwise distance (kbp) between SNPs. LD median estimates were obtained averaging values from all Illumina WGS data populations (ds2 and ds3.1).

### Candidate genes in high LD blocks

We performed an initial chromosome scan using Illumina WGS data from the *H. r. erythrogaster* and *H. r. savignii* subspecies (28) (ds3.1) to identify potential regions of interest (ROIs) exhibiting high LD values (average r^2^ > 0.3). Despite the small sample size and the rapid genome-wide LD decay, our analyses revealed the presence of 78 ROIs, many of which (N = 57/78) spanned at least one annotated protein coding gene (N = 83; Table S28). Excluding ROIs containing sequences potentially collapsed in the reference or not overlapping with annotated genes, the locus showing the highest r^2^ values was on chr6 (ROI 45) and harboured four genes (*CCDC34, LGR4, LIN7C* and *BDNF*; Fig. 5a; Additional file 1: Table S28). Among these genes, *BDNF* is particularly interesting because it encodes a major neurotrophin involved in neuronal plasticity and differentiation (103,104). In zebra finch males, its transcript is upregulated to high levels in the high vocal centre (HVC) by singing activity (105), particularly when juveniles start to emit vocalisations, and its tissue-specific overexpression significantly increases during sensorimotor song learning (39,106,107). BDNF is also implicated in neural crest cells development (108), and studies in multiple domesticated mammalian species suggest a role for the modification of neural crest development in driving the concerted evolution of tame phenotypes during domestication (i.e., ‘domestication syndrome’) (6,7). It is also extensively implicated in the response to stress, fear, and fear memory consolidation (109). Similarly to other species (110), barn swallow *BDNF* presents alternative transcripts (Fig. 5b), three of which (transcript variants X2, X3, X4) lead to the same amino acid sequence, suggesting the presence of important regulatory elements. In other bird species, temperature (chicken (111,112)) and prolonged social isolation (zebra finch (112)) affect the expression of *BDNF* through a methylation-mediated mechanism associated with CpG sites located within CpG islands upstream of the translation start site, as well as in the coding region. Initially, using WGS data from American and Egyptian samples (28) (ds3.1), we detected 6 LD blocks comprising 104 SNPs within the *BDNF* gene region. Of these SNPs, 30 directly alter CpG sites, either in the reference or in the alternate allele sequence (Additional file 1: Table S29). The highest LD values were identified within *H. r. savignii* population (Additional file 1: Figure S14a), where we also detected an average homozygosity (i.e. the average proportion of homozygous genotypes) of ∼88.8% across all samples for the genotyped SNPs within the gene (Additional file 1: Table S29). The strong LD detected at CpG sites may indicate that certain alleles have been favoured by selection (97,113). In the specific case of the Egyptian barn swallow, where there is evidence of a past bottleneck event (28), we cannot exclude that genetic drift may have also played a role. However, the same genomic region in all other available WGS populations (ds2) had similar LD patterns (Additional file 1: Figure S14). For instance, *H. r. transitiva* showed very high pairwise LD values within *BDNF* gene coordinates (Additional file 1: Figure S14c). We further confirmed the presence of a potential selective signature within this genomic region by computing population haplotype homozygosity statistics (iHS, the integrated haplotype homozygosity score) on chr6 in WGS ds3.1, ds2.1 and ds2.2. The ROI harbouring *BDNF* identified with genome-wide LD scans was associated with significant outlier peaks also in this analysis (Additional file 1: Figures S15-16). Four CpG islands are present within the sequence of *BDNF* in the barn swallow (Fig. 5b, blue blocks). The first CpG island corresponds to one of the two genomic regions containing methylated sites previously described in zebra finch (112). We found that four of the seven CpG sites reported in zebra finch are conserved in the barn swallow (Fig. 5c, highlighted in yellow). One SNP present in our barn swallow markers catalogue (chr6:53,908,036) directly affects a CpG site adjacent to a zebra finch methylation site (112) (Fig. 5c, SNP adjacent to the first highlighted CpG site). We also analysed this region in the Cactus multialignment and found that all of the zebra finch CpG sites are conserved in all other bird species, except for the chicken, where only two sites are conserved as CpG (Fig. 5c). The presence and conservation of CpG sites in the barn swallow, together with the identified selection signatures associated with this genomic region, reinforce the importance of these sites. CpG islands are known to directly affect the transcription of genes by altering local chromatin structure, mostly through methylation of CpG dinucleotides (111). For *BDNF* methylation-dependent transcriptional regulation involving CpG islands has been shown to affect fear memory consolidation (114), a process strictly involved in domestication. Methylation state assays could potentially help to further investigate the role played by epigenetic modifications of *BDNF* in the barn swallow, providing additional insights on the evolution of tameness-related habits in this species.

**Fig. 5.**
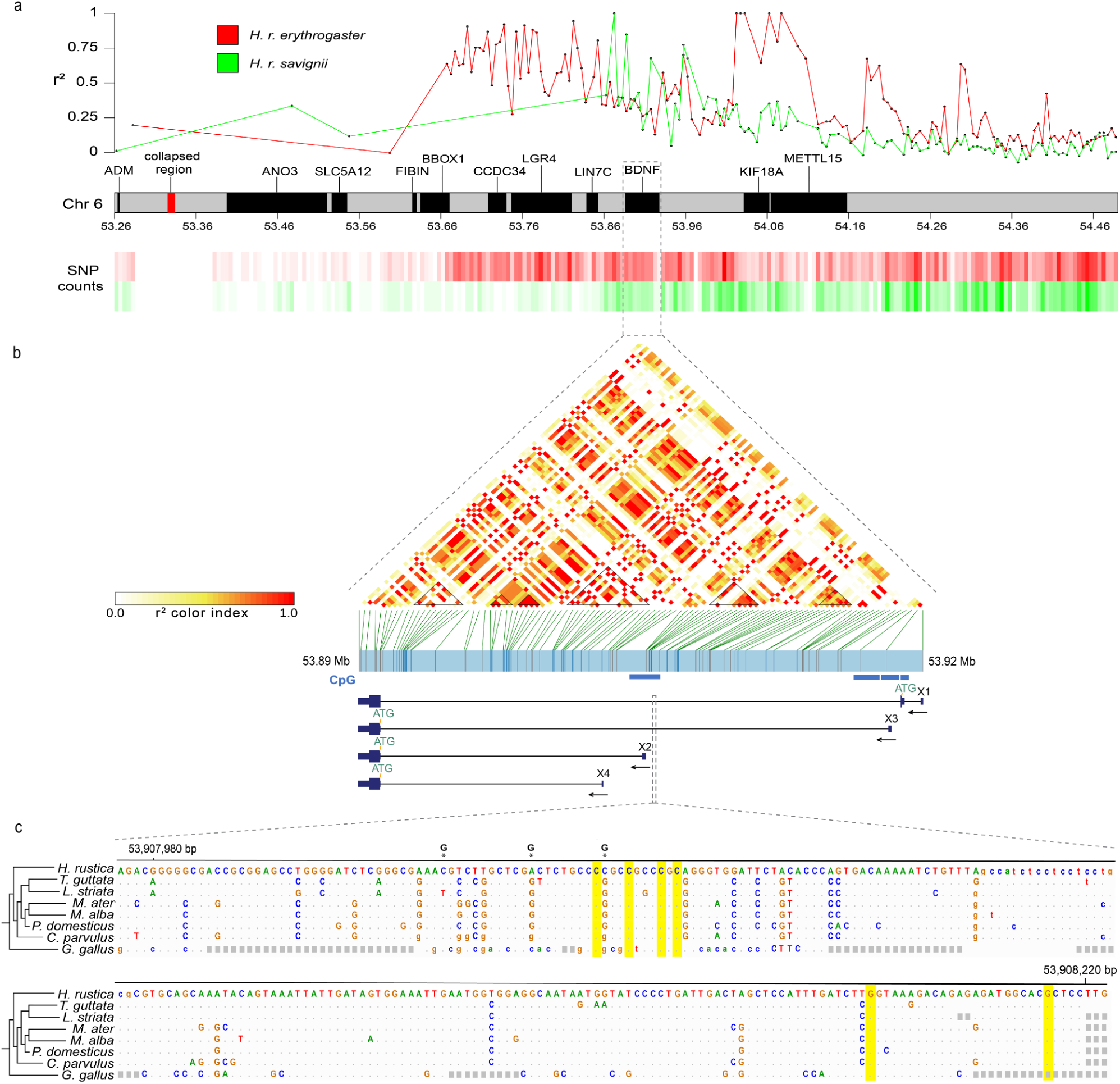
Patterns of LD blocks in genomic regions on chr6. **a** Average r^2^ values computed over 5 kbp windows on chr. 6 (upper panel; from 53.26 Mb to 54.49 Mb) for the *H. r. savignii* (green) and *H. r. erythrogaster* (red) populations (ds3.1). The region shown in the plot extends beyond ROI 45. Each point represents the average r^2^ value per window and was placed at the midpoint of the genomic region. The heatmap in the lower panel represents SNP counts for the two populations analysed. **b** Upper panel: LD heatmap within *BDNF* gene coordinates considering the two populations combined. Black triangles indicate LD blocks. Blue horizontal blocks mark the presence of CpG islands. Lower panel: barn swallow *BDNF* four transcript isoforms X1, X2, X3 and X4 (big rectangles: coding exons; small rectangles: non coding exons; horizontal line: introns; arrows indicate the direction of transcription). **c** Cactus multiple alignment of the zebra finch (second line) region containing CpG sites important for methylation-dependent regulation (112). Asterisks: SNPs present in barn swallow marker catalogue. Alternate base is shown on top of the barn swallow reference sequence. Yellow: zebra finch methylated sites (112). The second, third and sixth CpG sites are conserved in the barn swallow. The first one (at position 53,908,035) is not fixed in the barn swallow but the transition of the adjacent polymorphic site from reference (C) to alternate (G) allele leads to the formation of a CpG site.

## Conclusion

Using our high-quality, karyotype-validated and fully annotated chromosome-level reference genome for the barn swallow in combination with comparative and population genomics, we detected genes involved in domestication and song learning. Particularly, *CAMK2N2* has a role in fear memory formation and is likely involved in the glutamatergic system, which in turn plays a key role in domestication through the attenuation of stress response and aggressive behaviour. Similarly, *BDNF* is also involved in stress response and fear memory consolidation, as well as tameness during domestication, through its role in neural crest development. Based on these results, we propose that the strict association with humans in this species is linked with the evolution of pathways suppressing fear response and promoting tameness that are typically under selection in domesticated taxa.

## Methods

### Genome sequencing, assembly and annotation

HMW (High Molecular Weight) DNA was extracted from muscle tissue of a female barn swallow captured in a farm near Milan (Italy) and sequenced using 10x Genomics and Arima Hi-C technologies (Additional file 1: Supplementary Methods). Genomescope2.0 (43) was run online (http://qb.cshl.edu/genomescope/genomescope2.0/) starting from the *k*-mer (31 bp) histogram resulting from Meryl (115) (Additional file 1: Supplementary Methods). Newly generated data were combined with PacBio CLR long reads and Bionano optical maps already available for the same individual (31), using the VGP standard genome assembly pipeline 1.6 (36) (Additional file 1: Figure S1, Supplementary Methods). Briefly, Pacbio CLR long reads were assembled using FALCON (116), contigs were phased with FALCON-unzip (117) and polished with Arrow (smrtanalysis 5.1.0.26412). Two sets of contigs were generated, primary, representing one of the haplotypes, and alternate, representing the secondary haplotype. The primary contigs were purged (118), generating purged contigs and alternate haplotigs. The latter were merged with the alternate contigs and purged again. The primary purged contigs were then subjected to three steps of scaffolding with 10x linked reads, Bionano optical maps and Hi-C reads, generating chromosome-level scaffolds. Final scaffolds were merged with the alternate contigs and the mitogenome, generated with NOVOplasty (119) (Additional file 1: Supplementary Methods), polished with Arrow (smrtanalysis 5.1.0.26412) and Freebayes (120), and separated again in the two haplotypes, which then went through two steps of manual curation (121) (Additional file 1: Supplementary Methods). The primary curated assembly was annotated with IsoSeq and RNAseq data (Additional file 1: Table S7, Supplementary Methods).

### Karyotype reconstruction

To confirm the chromosomal structure of our assembly, a karyotype for the barn swallow was generated using a cultured cell protocol (Additional file 1: Supplementary Methods). Chromosome sizes were predicted from the karyotype images and correlated with the assembly chromosome sizes (Additional file 1: Supplementary Methods).

### Assembly evaluation and comparison with Chelidonia

Summary assembly statistics were computed with a script included in the VGP assembly pipeline GitHub repository (https://github.com/VGP/vgp-assembly/blob/master/pipeline/stats/asm_stats.sh). The assembly was further evaluated using BUSCO (44,122), Merqury (115), and Hi-C contact heatmaps (Additional file 1: Supplementary Methods). PacBio CLR long reads were aligned to the assembly and repeats were masked with a combination of Windowmasker (123) and RepeatMasker (124,125) (Additional file 1: Supplementary Methods). The same procedure was applied to Chelidonia (31). A purge_dups (118) run was performed on the latter with default parameters. Correlations between bHirRus1 chromosome size and genomic content were performed with Spearman nonparametric rank tests (126) and Wilcoxon signed-rank tests (127) (Additional file 1: Supplementary Methods).

### Cactus alignment

Progressive Cactus (50) v1.3.0 with default parameters was used initially to align bHirRus1 with 8 chromosome-level annotated Passeriformes genomes available on NCBI and the Chicken genome (Table S10). A maximum of 10 species were chosen for computational limits using Cactus. Despite different runs with the same parameters, two species failed to align (*Parus major* and *Ficedula albicollis*) and were excluded from the subsequent analyses. The guide tree was taken from TimeTree (128) (Additional file 1: Figure S7, Supplementary Methods). The genomes were soft-masked with WindowMasker (123) and RepeatMasker (124) (http://www.repeatmasker.org) (50) and then aligned (Additional file 1: Supplementary Methods). The alignment coverage was calculated with halAlignmentDepth (129) with the --noAncestors option and bHirRus1 as target species.

### Neutral model estimation

PHAST v1.5 (130) was used in combination with the HAL toolkit (129) for the selection analyses. An alignment in the MAF format was extracted for each bHirRus1 chromosome from the Cactus HAL output using hal2maf (129) with the --noAncestors and --onlyOrthologs options. The MAFs were post-processed with maf_stream (https://github.com/joelarmstrong/maf_stream), as previously described (57). The non-conserved neutral model was trained from fourfold degenerate (4d) sites in the coding regions of the barn swallow annotation (55,131). Briefly, CDS that fall within bHirRus1 chromosomes were extracted from the NCBI gff3 annotation file. msa_view (130) was used to extract 4d codons and 4d sites from each MAF separately, using the correspondent CDS coordinates. The combined 4d sites were used with phyloFit (--subst-mod REV --EM) to generate the neutral model.

### PhyloP analysis

To detect conserved and accelerated bases, phyloP (130) was run on each chromosome separately using the neutral model with LRT method and in the CONACC mode. Due to the low total branch length between the aligned species (57), no significant calls were found after the false discovery rate (FDR) (52) correction with 0.05 as significance level. We increased the statistical power of the constraint analysis by running phyloP on 10bp windows. Briefly, the aligned coordinates of bHirRus1 in the Cactus alignment were obtained and divided into 10bp windows. PhyloP was run again on the windows (LRT method and CONACC mode), and the FDR correction at 5% was applied. Windows smaller than 10bp were discarded and windows overlapping with assembly gaps were removed. Spearman nonparametric rank test (126) was used to correlate chromosome size and the fraction covered by phyloP sites (Additional file 1: Table S3). Wilcoxon signed-rank test (127) was used to compare differences between microchromosomes and the other chromosomes.

### PhastCons analysis

An additional conservation analysis was performed using PhastCons (130) with the same neutral model as phyloP analysis, to predict discrete conserved elements (CEs). PhastCons requires parameter tuning to reach the desired levels of smoothing and coverage (130). Each chromosome MAF file was split in chunks and 200 of them were randomly selected. phastCons was run on each chunk with initial parameters (Additional file 1: Supplementary Methods) generating tuned conserved and non-conserved models. These models were then used with phastCons to predict conservation scores and discrete conserved elements. Levels of smoothing and coverage were checked and the analysis was repeated again until the desired tuning was reached (Additional file 1: Supplementary Methods). Following Craig et al. (54), windows that overlapped for more than 20% with an assembly gap were removed, and all bases that fell into gaps were filtered out. Correlations between phyloP conserved elements and phastcons CEs as the number of elements per 10kb windows were computed with the Spearman correlation rank test (126).

### Candidate genes detection and *CAMK2N2* tree construction

To detect candidate genes, we intersected the conserved and accelerated bases detected with each annotated class extracted with GenomicFeatures (Additional file 1: Supplementary Methods). Bases overlapping with more than one feature were assigned hierarchically based on the first appearance (54,132) in this order: CDS, 5’ UTR, 3’ UTR, intronic, intergenic. Genes without identified orthologs (“LOC” genes) were discarded. To look at differences in *CAMK2N2* transcript between species with different levels of association with humans, the transcript sequences of 38 species were downloaded from NCBI and aligned with Muscle on MEGA (133). The tree was then generated using the Maximum likelihood method, a generalised time reversible (GTR) model and a gamma distribution (G) with 5 categories.

### Gene ontology enrichment analysis

The gene ontology analysis was performed on the top 500 genes with the most overlaps with phyloP accelerated and conserved sites using *gage (134)* R package (Additional file 1: Supplementary Methods).

### Generation of the pangenome and orthologs analysis

For the generation of the pangenome, additional 5 Italian barn swallow individuals were sampled (Additional file 1: Supplementary Methods). HMW DNA was extracted from the blood samples and sequenced with the PacBio HiFi technology (Additional file 1: Supplementary Methods). HiFi reads were checked for adaptor contamination and trimmed accordingly with cutadapt v3.2 (135) (Additional file 1: Supplementary Methods). Hifiasm (82) was used to assemble both primary and alternate assemblies which were then purged with purge_dups (118) using custom cutoffs (83) (Table 31). Both primary and alternate HiFi-based assemblies were masked with Windowmasker (123) and RepeatMasker (124). The pangenome was generated with the Cactus (50) v1.3.0 Pangenome Pipeline (https://github.com/ComparativeGenomicsToolkit/cactus/blob/master/doc/pangenome.md, Additional file 1: Supplementary Methods) using HiFi-based assemblies and bHirRus1 primary and alternate assemblies. Orthologous genes were found running HALPER (136) following the steps described on GitHub (https://github.com/pfenninglab/halLiftover-postprocessing). Briefly, from the HAL alignment, the coverage of bHirRus1 was calculated with halAlignmentDepth (129). Then, a file for the ortholog extension was generated from the coverage file and halLiftover (129) and used to lift bHirRus1 gene coordinates on the aligned HiFi assemblies. Orthologs were then found using the lifted genes. The resulting lists of orthologs were manually evaluated to find genes shared between individuals. The 236 genes that were found only in the bHirRus1 assembly were searched in the HiFi raw reads with BLAST 2.10.1+ (84). The alignments were checked to find genes present for more than 80% of their sequence in the reads and 99% identity with the query sequence.

### SNP catalogue

To generate the catalogue of genetic variants, all publicly available datasets were combined with our newly generated Hifi reads set (see “HiFi reads processing for genetic variants identification” Methods section). For each publicly available dataset, sequencing adapters and low quality bases were trimmed when present, and reads were aligned to the bHirRus1 reference genome. Freebayes v1.3 (120) was used on the alignments to call variants, parallelizing the process with a script from the VGP assembly pipeline (Additional file 1: Supplementary Methods). Variants were then split by population and markers were filtered for quality, read depth supporting each variant call, average fraction of missing sites among individuals and minor allele frequency (maf). Samples showing > 70% of missing genotypes were removed. Variants within repetitive regions were excluded, and only SNPs were extracted for downstream analysis. Details relative to the filters and threshold values used can be found in the Additional file 1: Supplementary Methods section.

For SNP density plotting and its correlation with genomic features, all data using the same sequencing technology were merged (HiFi WGS; Illumina WGS; Illumina ddRAD). SNP density was computed across all chromosomes (excluding unlocalized/unplaced scaffolds) over 10 kbp windows and these values were correlated with the GC content per window using the Spearman nonparametric rank test (126). SNPs falling in genic, intergenic, exonic and intronic regions (as determined from NCBI annotation) for each chromosome in the different datasets were counted. To plot SNP density across all chromosomes, the KaryoploteR package (137) was used, computing its value over 40 kbp intervals (Additional file 1: Supplementary Methods).

### Linkage disequilibrium analysis

Genome-wide LD decay was evaluated in all Illumina WGS datasets by calculating the r^2^ coefficient using Plink v1.9 (138), considering all marker pairs within a 55 kbp distance. To calculate average r^2^, SNP pairs were grouped according to their distance in bins of 1 kbp (range 1-55 kbp; Additional file 1: Supplementary Methods). The same approach was used to calculate average r^2^ values per chromosome group (macrochromosomes, intermediate and microchromosomes), except that values were then averaged across specific distance bins (Additional file 1: Supplementary Methods).

### LD scans and extended haplotype homozygosity statistics

To scan chromosomes for regions containing alleles exhibiting high local LD values, Plink v1.9 (138) was used, considering marker pairs within a 15 kbp distance maximum. For the first LD scan, Illumina WGS data from ds3.1 were used. Each chromosome was divided into non-overlapping 5 kbp sliding windows to compute average LD (Additional file 1: Supplementary Methods). Next, only genomic windows with average r^2^ > 0.3 were extracted and intersected with annotated features to generate a list of top candidate genes carrying alleles with high LD. Windows were excluded if in proximity (within ∼5 kbp) with potentially collapsed or low-confidence assembly regions (considering a PacBio reads coverage value higher than twice the average genome-wide coverage or lower than 10, respectively). Before computing within population haplotype homozygosity statistics (iHS) in ds3.1, ds2.1 and ds2.2, variants present on chr6 were phased and specifically filtered according to genotype missingness and maf parameters (Additional file 1: Supplementary Methods).

### HiFi reads processing for genetic variants identification

HiFi reads from ds1 were aligned to bHirRus1 and small variants were called using deepvariant v1.0.0 (139). Only biallelic SNPs were kept, and variants falling within repetitive regions were removed. Next, variants were filtered according to genotype quality (quality > 20) and variant site depth (5% and 95% quantiles of the read depth values distribution were used to set the minimum and maximum site coverage). Joint variant calling of single-nucleotide variants (SNVs) and small insertions-deletions (indels) was performed using gVCF files from DeepVariant v1.1.0 per-sample calls, jointly called with GLNexus (140) pipeline (Additional file 1: Supplementary Methods). For structural variants (SVs), *pbsv* v2.6.0 (141) (commit v2.4.1-155-g281bd17) was used for per-sample and joint variant calling.

### Titration and phasing experiments with HiFi reads

HiFi reads were first randomly downsampled and two titration experiments were conducted, the first one using variants obtained with individual variant calling and the second one with joint variant calling (N = 5). Estimation of haplotype-phased blocks length was also performed (Additional file 1: Supplementary Methods).

## Data availability

Scripts used in this paper are available on GitHub (https://github.com/SwallowGenomics/BarnSwallow). Primary and alternate assemblies used in this study are available on NCBI under accession numbers GCF_015227805.1 and GCA_015227805.3. All raw data supporting the genome assembly are available on GenomeArk (https://vgp.github.io/genomeark/Hirundo_rustica/), and will also be available upon publication in SRA. The HiFi data will be made available upon publication. IsoSeq and RNAseq data are available on NCBI under the accession numbers SRR13516425, SRR13516426, SRR13516427, SRR9184408 and SRR9184409. The SNPs catalogue will be available upon publication on Dryad.

## Acknowledgments

This work would have not been possible without the dedication of Prof. Nicola Saino. We received support from: the Italian Ministry of Education, University and Research (MIUR) for the project PRIN2017 2017CWHLHY (L.G. and A.T.); Dipartimenti di Eccellenza Program (2018–2022) -Department of Biology and Biotechnology “L. Spallanzani’’ University of Pavia (to A.O., L.F. and A.T.); the CSU Program for Education & Research in Biotechnology (CSUPERB) (to A.B.-A.); Howard Hughes Medical Institute (to E.D.J.); Samuel Freeman Charitable Trust (to T.A.M. and A.P.M.). The work of F.T.-N. and P.M. was supported by the Intramural Research Program of the National Library of Medicine, National Institutes of Health. We thank the INDACO Platform team (a project of High Performance Computing at the University of Milan, http://www.unimi.it), in particular Dr. Alessio Alessi, as well as Prof. Aureliano Bombarely for providing computational resources and technical assistance. We thank Prof. Guido Grilli (Department of Veterinary, University of Milan, Milan, Italy) for euthanising and dissecting the barn swallow individual used for the assembly and annotation of bHirRus1 reference genome. We thank Dr. Alessandra Costanzo for her help in obtaining barn swallows blood samples.

## Author contributions

S.S., G.R.G., A.I, E.G, J.B, M.C., J.M, M.Sa., R.S. and G.F. performed the wet lab experiments.

S.S., G.R.G, A.T., A.B.-A., L.G and G.F. planned the experiments.

S.S., G.R.G., M.So., A.I., J.F.O., R.S., P.M., K.W., L.G. and G.F. analysed the data.

S.S., G.R.G., M.So., A.B.-A., L.G. and G.F. drafted the manuscript.

C.C., A.P.M, T.M, A.T., A.B.-A., E.D.J. and L.G. provided computational resources or funding.

S.S., W.C., J.C., K.H. and J.T. performed manual curation.

S.S., P.M. and F.T.-N. performed assembly annotation.

J.B., O.F., B.H. and J.M., generated the raw sequencing data.

S.S. generated the genome assembly with support from A.F. and A.R. S.S., A.I., M.C., D.R., R.A. and G.F. contributed to sampling.

S.S., L.A., W.K., E.D.J. and G.F. handled data submission.

L.F., G.L., A.O., J.F.-O., D.R., A.T., R.A., A.B.-A. and E.D.J contributed to the general discussion. All authors reviewed the final manuscript and approved it.

