## Supplementary Material for "Pangenomics provides insights into the role of synanthropy in barn swallow evolution"

### Additional file 1

**Supplementary Figures** 3

**Supplementary Methods** 24

1 - bHirRus1 genome assembly 24
1 - 1 Sampling 24
1 - 2 DNA extraction 24
1 - 3 Library preparation and sequencing 24
1 - 4 Genomescope2.0 predictions 25
1 - 5 Mitogenome assembly 25
1 - 6 Assembly pipeline 25
1 - 6 - 1 Contigging, phasing and duplication removal 26
1 - 6 - 2 Scaffolding 26
1 - 6 - 3 Polishing 27
1 - 6 - 4 Manual curation 27
1 - 6 - 5 Annotation 28
2 - Karyotype description and chromosome classification 29
2 - 1 Karyotype construction 29
2 - 2 Karyotype chromosome size estimations 29
2 - 3 Chromosome classification 30
3 - bHirRus1 evaluation 30
3 - 1 Raw read alignments 30
3 - 2 Assembly statistics 30
3 - 3 Hi-C contact heatmaps 30
3 - 4 Masking of repetitive regions 31
3 - 5 QV, k-mer completeness and duplication content 31
3 - 6 Chromosome size and genomic content 31
4 - Haplotig purging in Chelidonia 31
5 - Selection analysis on multiple whole-genome alignments 32
5 - 1 Cactus alignment 32
5 - 2 Neutral model estimation and phyloP analysis 32
5 - 3 phastCons parameters tuning 32
5 - 4 Gene ontology (GO) analysis 33
6 - Generation of the pangenome 33
6 - 1 Sampling, DNA extraction, and PacBio Hifi sequencing 33
6 - 2 Pangenome assembly 34
7 - SNP catalogue generation 35
7 - 1 Publicly available datasets 35
7 - 2 Variant calling and filtering 35
7 - 3 SNP density statistics and correlations with genomic features 36
7 - 4 SNP density plotting 37
8 - LD and haplotype statistics analysis 37
8 - 1 Genome-wide LD decay 37
8 - 2 LD scans 38

8 - 3 iHS computation 38
9 - HiFi reads processing for SNP catalogue, titration and phasing experiment 39
9 - 1 HiFi reads alignment, variant calling and filtering 39
9 - 2 Titration of HiFi reads 40
9 - 3 Phasing of HiFi read sets 40
10 - Graphical representations 41

**Supplementary Note 43**
1 - bHirRus1 manual curation 43
1 - 1 First curation step 43
1 - 2 BUSCO genes recovery 43
2 - Barn swallow mitogenome 44
3 - Karyotype reconstruction 44
4 - bHirRus1 evaluation 45
5 - Comparison between bHirRus1 and Chelidonia 45
6 - PhyloP analysis 46
7 - SNP catalogue description 47
7 - 1 - SNP counts 47
7 - 2 - Correlation between SNP density and genomic features 47
8 - HiFi read mapping, variant calling, titration and phasing experiments 48
8 - 1 Sequencing and read mapping 48
8 - 2 Variant calling and titration experiment 48
8 - 3 Haplotype phasing with HiFi reads 49

**References 50**

**Supplementary Tables 57**

### Supplementary Figures

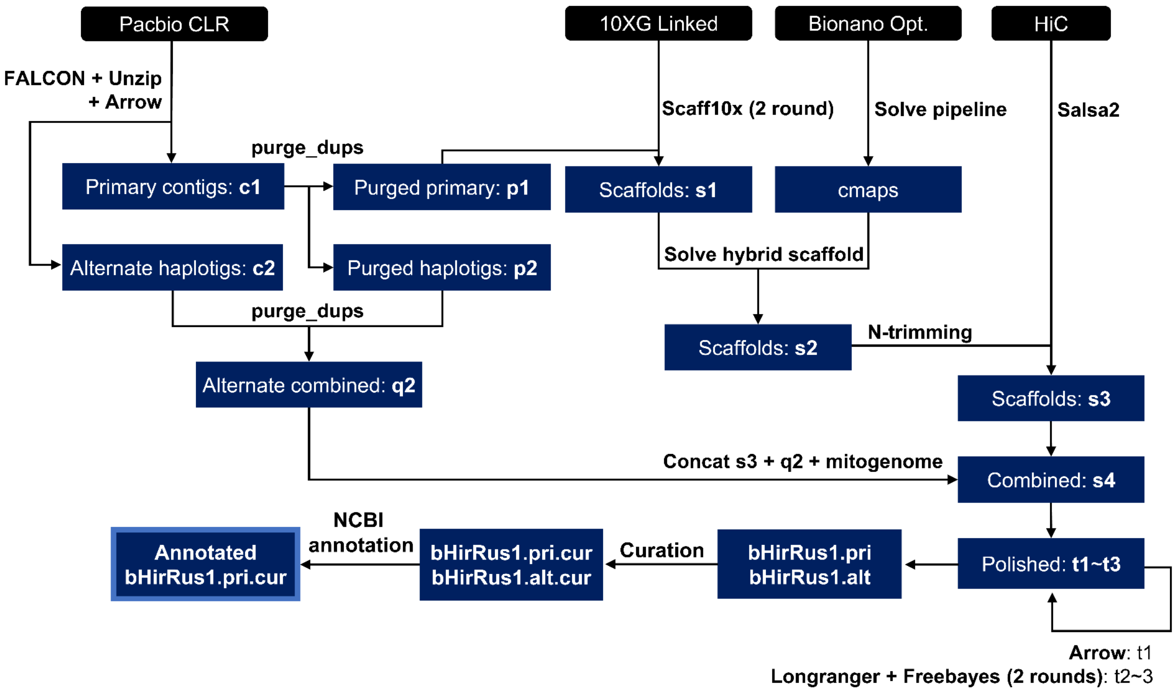

**Figure S1** Flowchart of the VGP assembly pipeline 1.6 (redrawn from Rhie et al. (1)). Primary and alternate pseudo-haplotypes are generated with FALCON-Unzip (2) using Pacbio CLR reads; haplotigs are purged with purge_dups (3); a first step of scaffolding is performed with 10X linked-reads using Scaff10X (https://github.com/wtsi-hpag/Scaff10X); additional scaffolding and mis-joins breaking are performed with Bionano Solve (https://bionanogenomics.com/support/software-downloads/) and Bionano optical maps; chromosome-level scaffolds are generated with Hi-C reads using Salsa2 (4); gaps filling and initial base-error polishing are performed using Arrow (Pacific BioSciences); two additional rounds of polishing with 10X linked-reads were performed with FreeBayes (5); expert manual curation is performed at the end to correct structural assembly errors introduced by the automated tools.

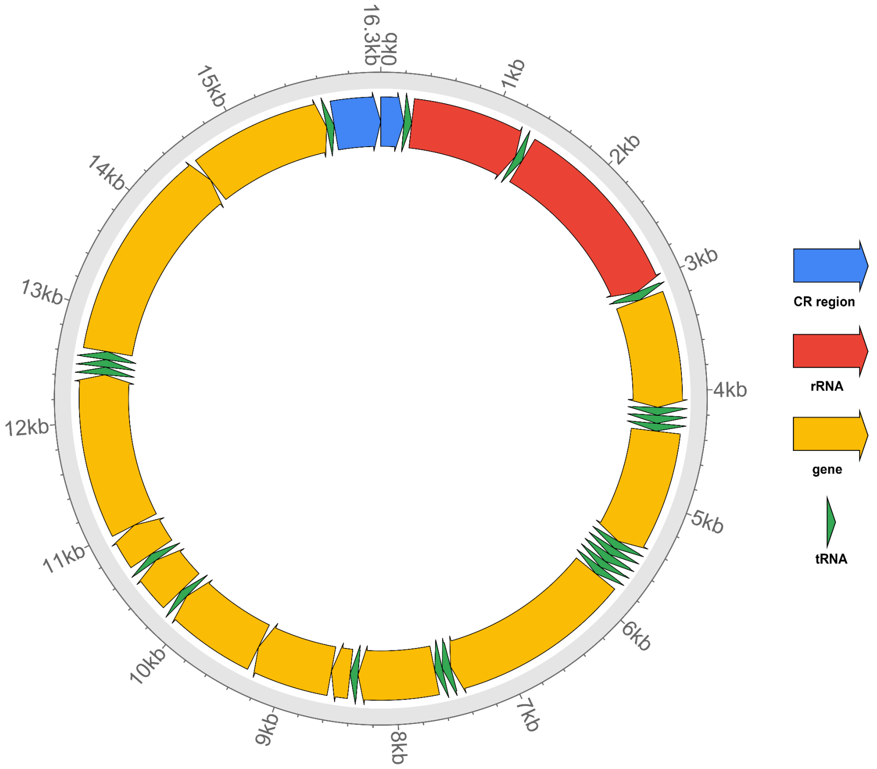

**Figure S2** Circular representation of the barn swallow mitogenome assembled with NOVOplasty (6). The control regions (CR) are coloured in blue, rRNA genes in red, tRNA genes in green, other genes in yellow.

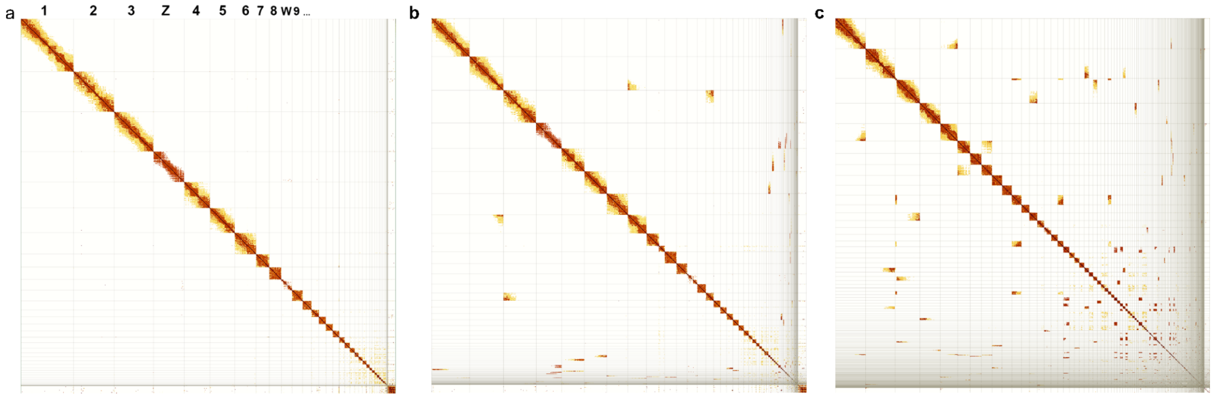

**Figure S3**. Hi-C interaction heatmaps. The linear sequence of the reference genome assembly is represented on both axes and the diagonal shows 3D proximity of interacting pairs. The strength of the interaction is given by colour intensity.A scaffold is considered a full chromosome when the number of interchromosomal interactions is negligible. **a** Curated bHirRus1 assembly. No off-diagonal interactions are visible. Scaffolds are labelled by their chromosome number. **b** bHirRus1 assembly before curation. A number of off-diagonal interactions are still visible, which can either result from missing links between scaffolds of the same chromosome or from misassembly. **c** Chelidonia assembly. The assembly is still substantially fragmented, with several off-diagonal Hi-C interactions.

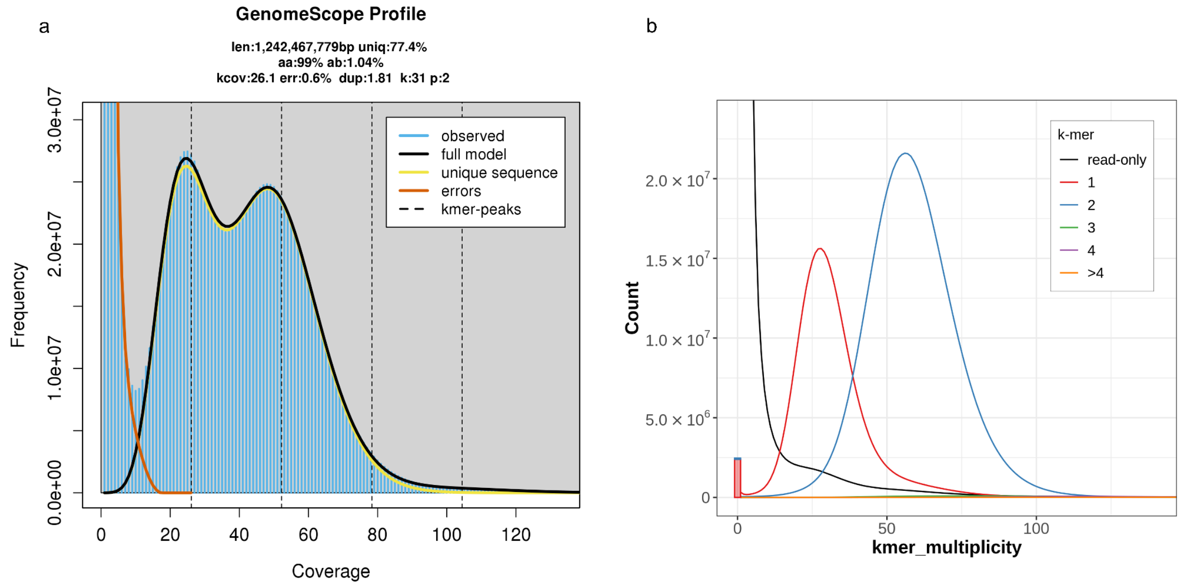

**Figure S4.** *K*-mers based statistics for bHirRus1. **a** Genomescope2.0 (7) *k*-mer profile for bHirRus1 generated from trimmed 10X linked reads, used to estimate genome size, repetitiveness and heterozygosity (top). The x-axis represents multiplicity in the read set, while the y-axis represents their cumulative frequency. **b** Merqury spectra-cn plots for bHirRus1. with k-mer multiplicity in the 10X linked reads (x-axis) versus their frequency (y-axis). Coloured curves discriminate *k*-mer occurrences in the assembly. The bar at the origin of the graph represents *k*-mers found only in the assembly (assembly errors). Two frequency peaks are visible: a haploid peak at ~25× coverage (half average coverage, red), representing *k*-mers found once in the assembly (haplotype specific), and a diploid peak at ~50× (average coverage, blue) representing *k*-mers found twice in the assembly (shared between haplotypes). No *k*-mers resulting from artificial duplications (green, purple, yellow) are visible.

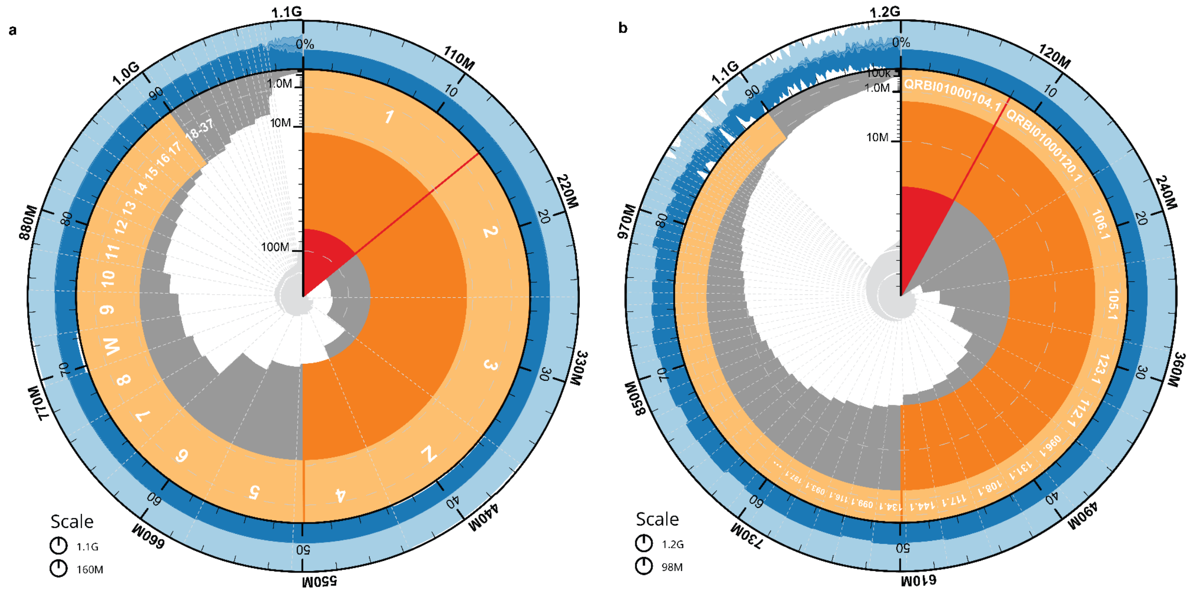

c

| Assembly | bHirRus1 | Chelidonia |
|---|---|---|
| Genome size (Gbp) | 1.1 | 1.2 |
| Number scaffolds | 617 | 364 |
| Longest scaffold (Mbp) | 156 | 98 |
| N50 (Mbp) | 76 | 26 |
| NG50 (Mbp) | 73 | 26 |
| N90 (Mbp) | 12 | 2 |
| BUSCO complete genes (%) | 96 | 95.9 |
| BUSCO fragmented genes (%) | 1.1 | 1.2 |
| BUSCO duplicated genes (%) | 0.8 | 1.3 |
| BUSCO missing genes (%) | 2.9 | 3 |
| Repeat content (Mbp) | 271 | 315 |

Composition
GC
AT
N

Scaffold statistics
Log10 scaffold count
Scaffold length
Longest scaffold
N50 length
N90 length

**Figure S5.** Snail plot and assembly summary statistics. The main plot is divided into 1,000 size-ordered bins around the circumference. Scaffold length distribution is shown in dark grey with the plot radius scaled to the longest scaffold (red). Orange and pale-orange arcs show scaffold N50 and N90, respectively. The pale grey spiral shows the cumulative scaffold count on a log scale, with white scale lines showing successive orders of magnitude. The blue and pale-blue areas around the plot show the GC, AT and N content in the same bins as the inner plot. **a** bHirRus1 snail plot. **b** Chelidonia snail plot. **c** Assembly summary statistics and BUSCO results (vertebrata_odb10) of Chelidonia and bHirRus1.

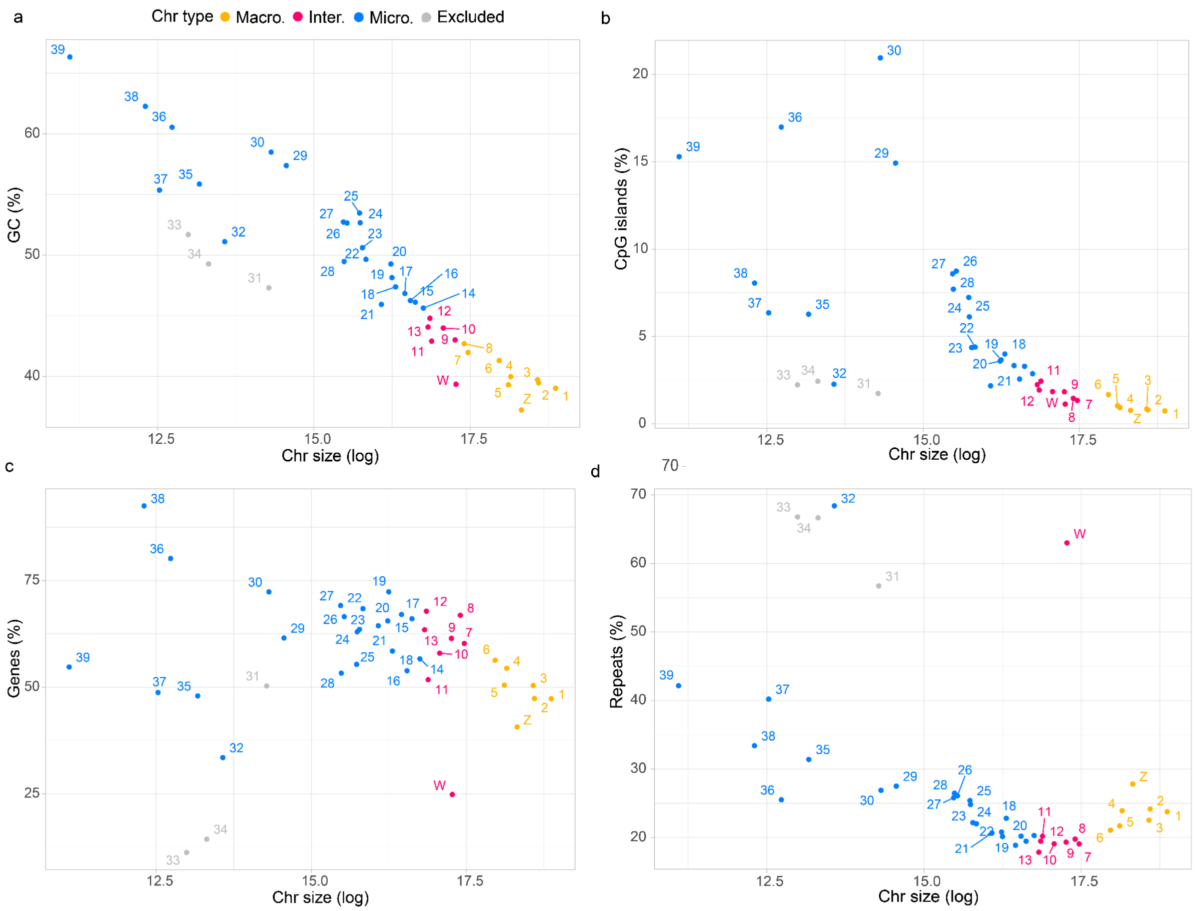

**Figure S6.** Correlation between chromosome length (log) and genomic features. Chromosome lengths tend to correlate inversely with **a** GC content, **b** CpG islands, **c** gene content, **d** repeat content. Chr 31, 33, 34 were excluded based on their higer PacBio raw-reads coverage with respect to the other chromosomes.

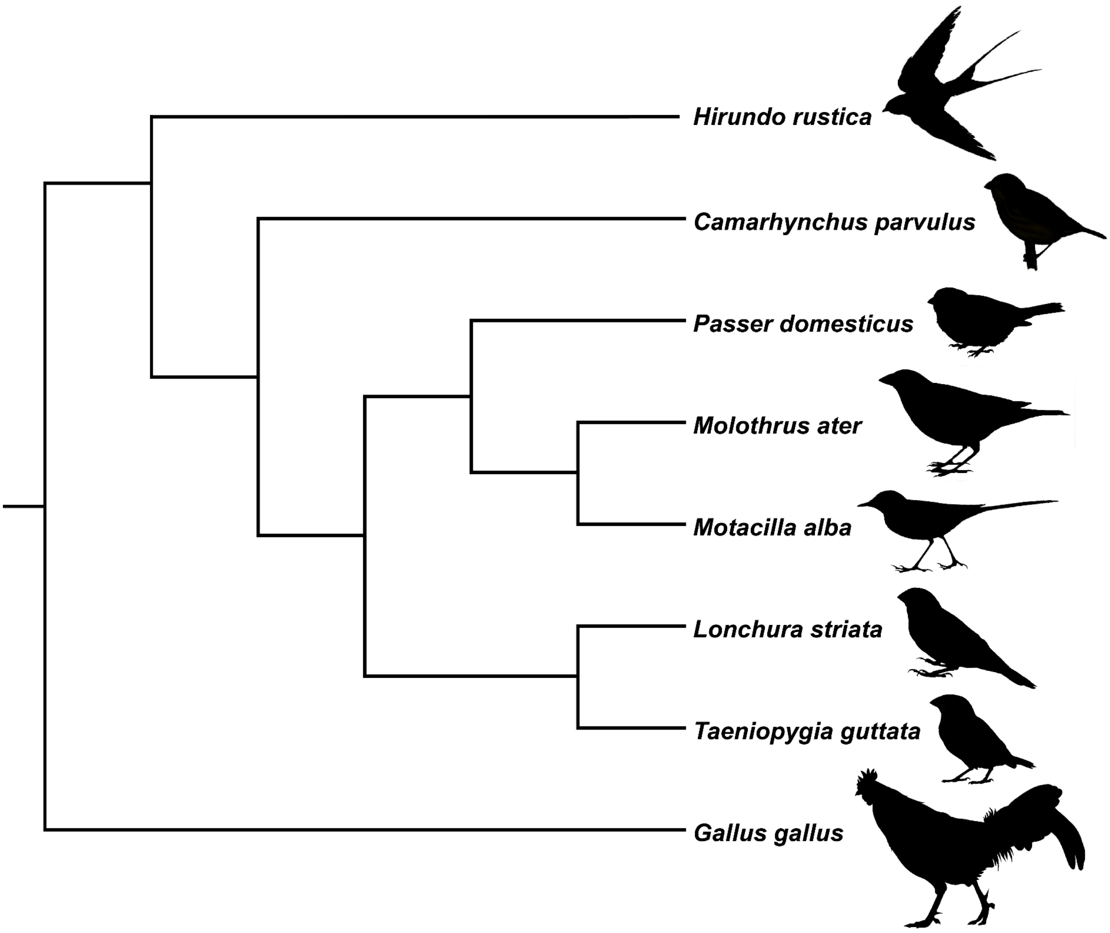

**Figure S7** Cactus aligned species tree topology. Cactus guide tree: (Gallus_gallus:98.04286929, ( (Hirundo_rustica:43.70000000, (Ficedula_albicollis:42.77829604,Parus_major:42.77829604):0.92170396):0.00000000, (Camarhynchus_parvulus:38.00000000, ( (Passer_domesticus:34.80000000, (Molothrus_ater:24.06782500,Motacilla_alba:24.06782500):10.73217500):0.00000000, (Lonchura_striata:10.12359500,Taeniopygia_guttata:10.12359500):24.67640500):3.20000000):5.70000000):54.34286929);

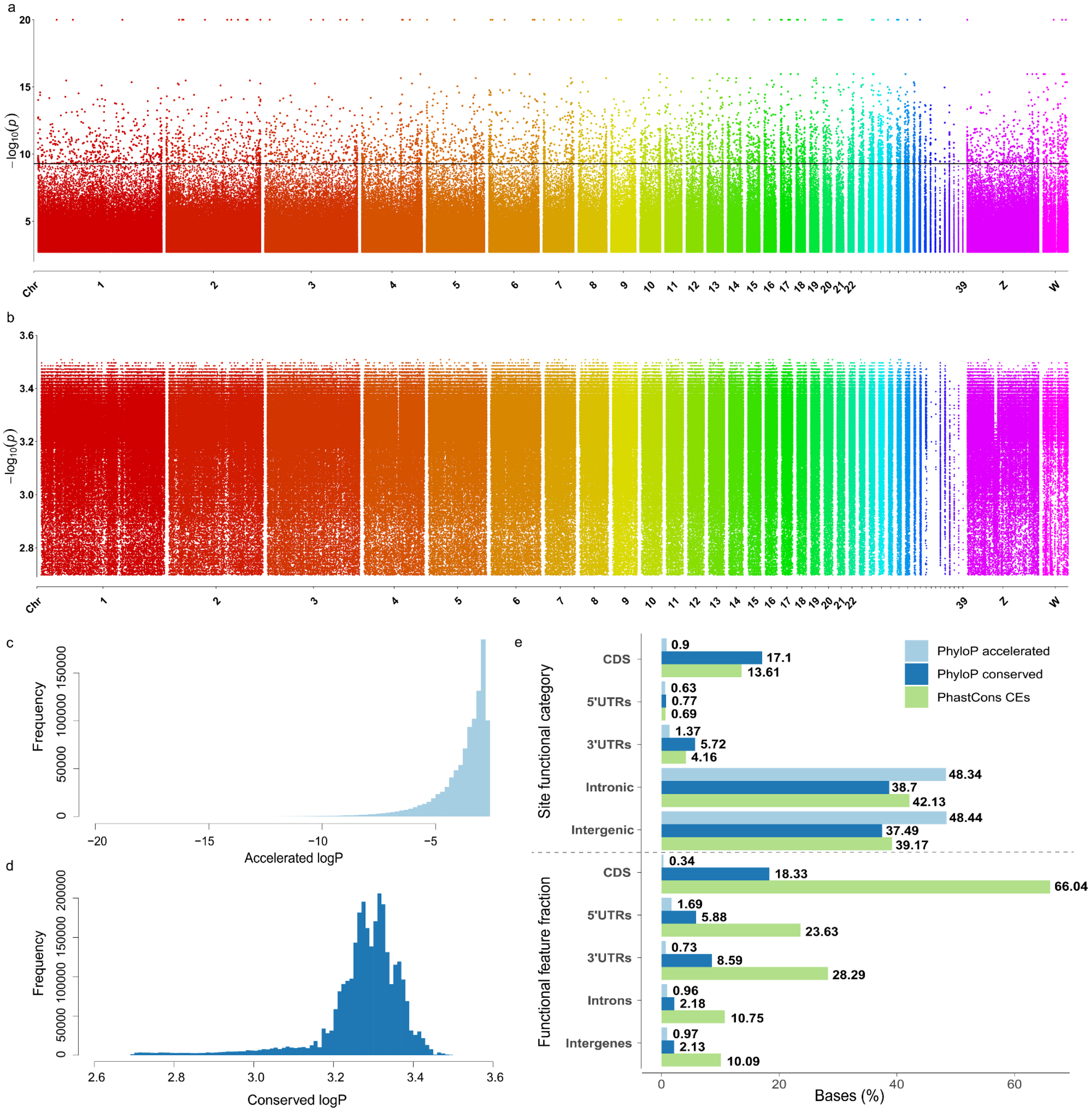

**Figure S8** Results from selection analysis. **a** Manhattan plot of the FDR-corrected accelerated sites identified with PhyloP. Each point represents a 10 bp conserved or accelerated site. LogP values are on the x axis. The horizontal line defines the Bonferroni threshold (9.3). **b** Manhattan plot of the FDR-corrected conserved sites found with PhyloP. No significant bases were found after Bonferroni correction. **c**, Frequency distribution of accelerated sites lod values. **d** Frequency distribution of conserved sites lod values. **e** Intersection between PhyloP conserved and accelerated, and PhastCons CEs bases with bHirRus1 genomic features. Top panel represents the percentage of PhyloP/PhastCons bases that overlap with genomic features, while the bottom panel represents the fraction of genomic features covered by accelerated or conserved bases. The CDS were the most conserved according to both phyloP and phastCons.

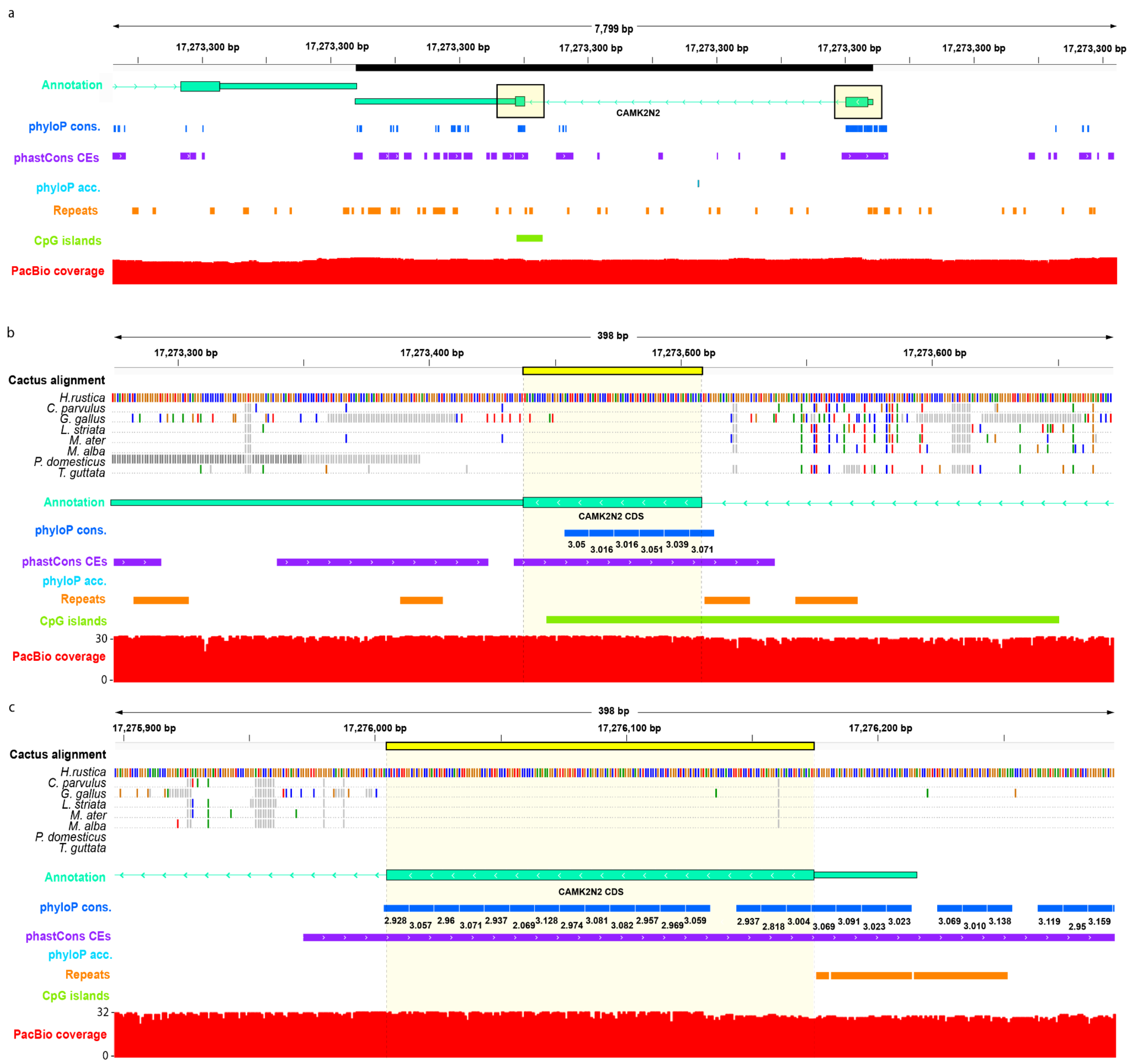

**Figure S9** CDS of *CAMK2N2* on chr10. **A** Whole gene. The black bar represents the gene coordinates. The yellow rectangles highlight the CDS regions. **B** First CDS (17,273,437-17,273,508). **c** second CDS (17,276,005-17,276,174). In b and c the yellow horizontal bar represents the CDS coordinates. Tracks represent Cactus MAF alignment referenced to the barn swallow, gff3 bHirRus1 gene annotation, phyloP conserved 10 bp windows with their logPvalue, phastCons CEs, PhyloP accelerated 10 bp windows, repeats, CpG islands and PacBio coverage. In the Cactus alignment track, all the barn swallow bases and divergent sites in the alignment are shown (green A, red T, blue C, orange G). Dots represent identical sites. SNPs in the other species are represented with bases with different colours than the barn swallow, while the points represent sites with the same base calling.

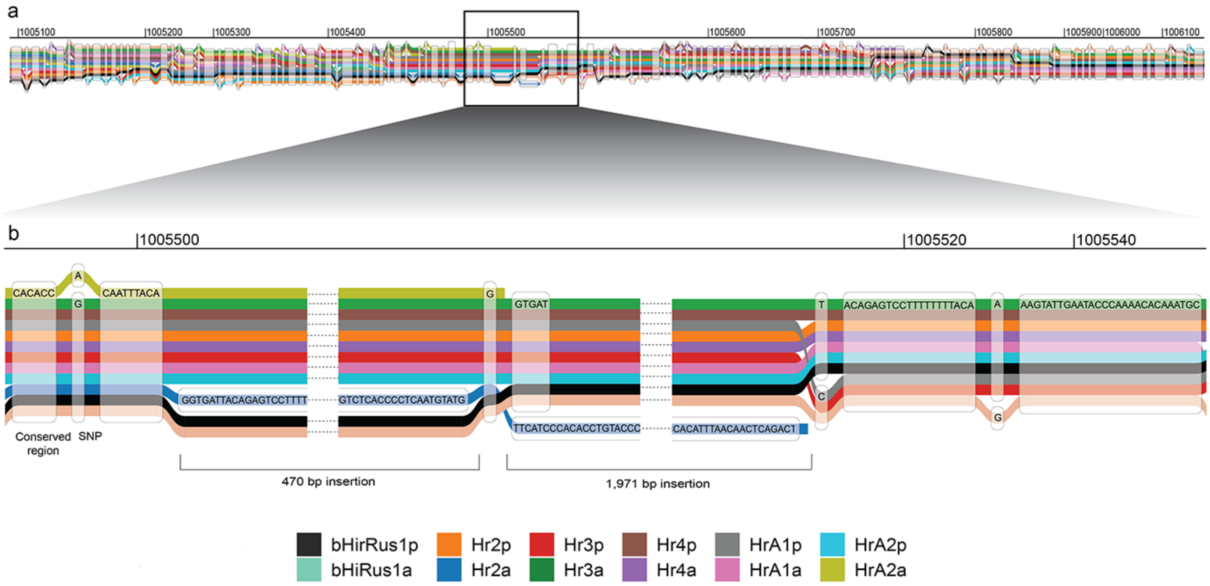

**Figure S10** Graphic representation of a portion of the barn swallow pangenome. **a** bHirRus1 Chr1:1,005,100-1,006,100 (1 kbp) in the barn swallow pangenome aligned to the alternate assembly bHirRus1a, the five HiFi-based primary (Hr2p, Hr3p, Hr4p, HrA1p, HrA2p) and their alternate assemblies (Hr2a, Hr3a, Hr4a, HrA1a, HrA2a). **b** Detail of the region showing two insertions in the alternate assembly of Hr2 (blue), as well as SNPs and conserved regions between assemblies.

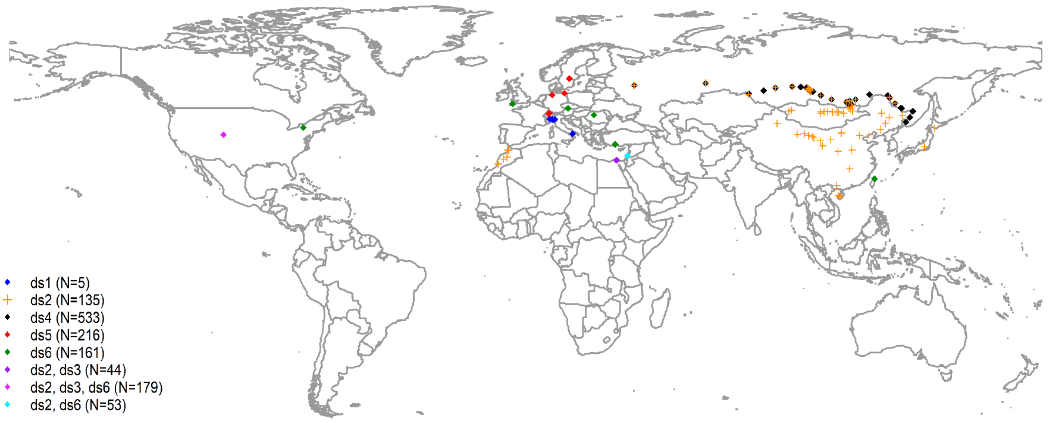

**Figure S11** Sampling locations of all individuals used to generate the SNP catalogue. Purple, fuchsia and light blue colours indicate sampling locations in common between datasets indicated in the legend. Sampling locations from ds2 are plotted with a different shape (cross) to distinguish them from black points (ds4), as some sampling locations partially overlap on the map.

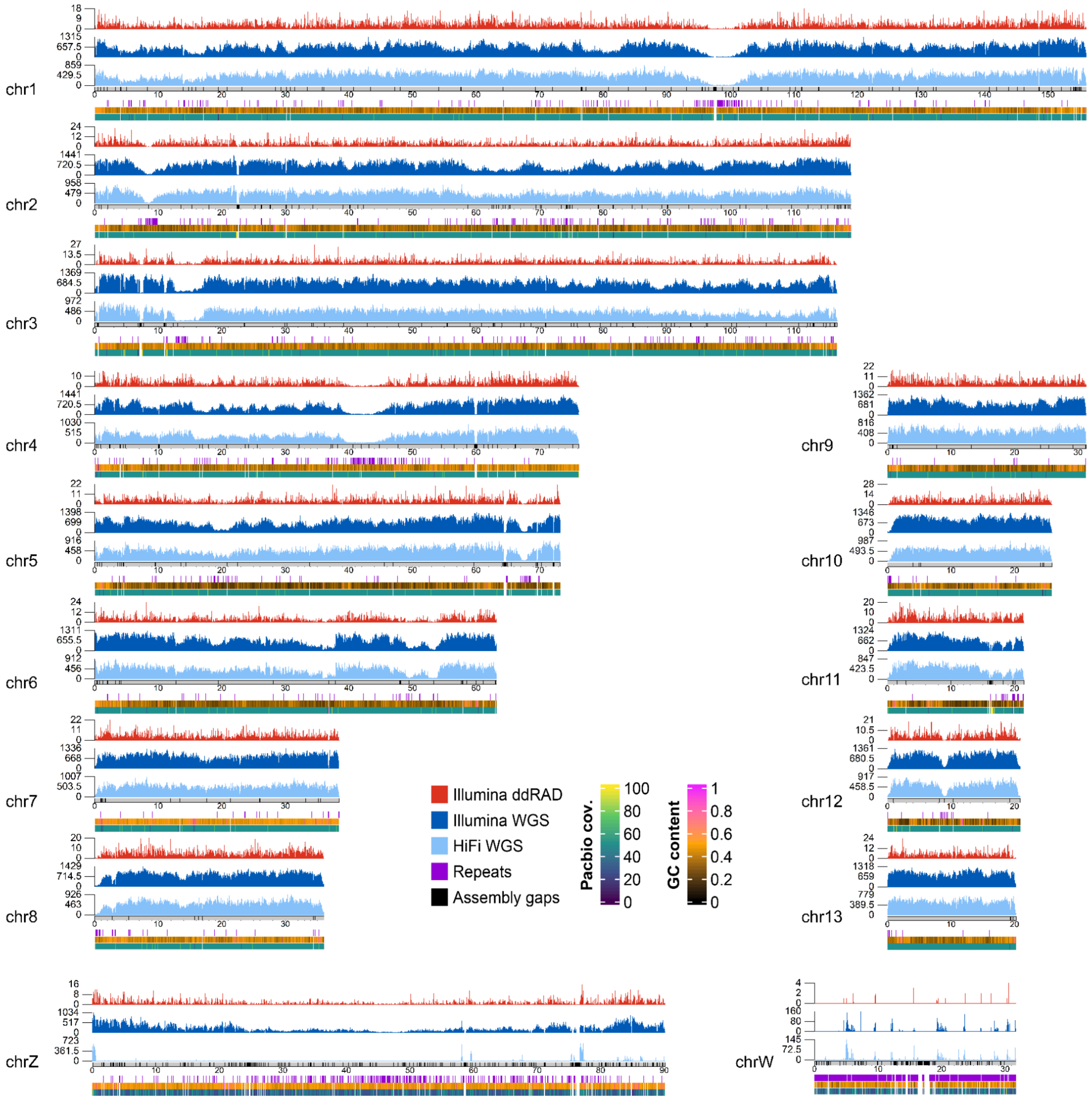

**Figure S12.** SNPs density per chromosome using a downsampled (5x) set of HiFi reads. Only macrochromosomes and intermediate chromosomes are shown. Sex chromosomes are shown at the bottom. SNP density, coloured according to the different types of genomic data used, was computed over 40 kbp windows. Maximum and average SNP density are indicated on the y axis. Light blue: HiFi WGS data (ds1). Dark blue: Illumina WGS data from ds2 and ds3.1. Red: Illumina ddRAD data from ds3.2 through ds6.8. All available samples from the same sequencing technology were considered together. Additional tracks in the lower panel show repetitive regions of the genome (violet bars; only regions larger than 3 kbp are plotted), GC content and PacBio reads coverage. Grey ideograms represent chromosomes in scale, with assembly gaps highlighted as black bars. The distribution of SNPs in the HiFi dataset after titration appears comparable to the full coverage distribution, as expected from a random subsampling of genomic data. The only chromosomes where downsampling led to a visible reduction in SNP density are sex chromosomes, consistent with their haploid coverage.

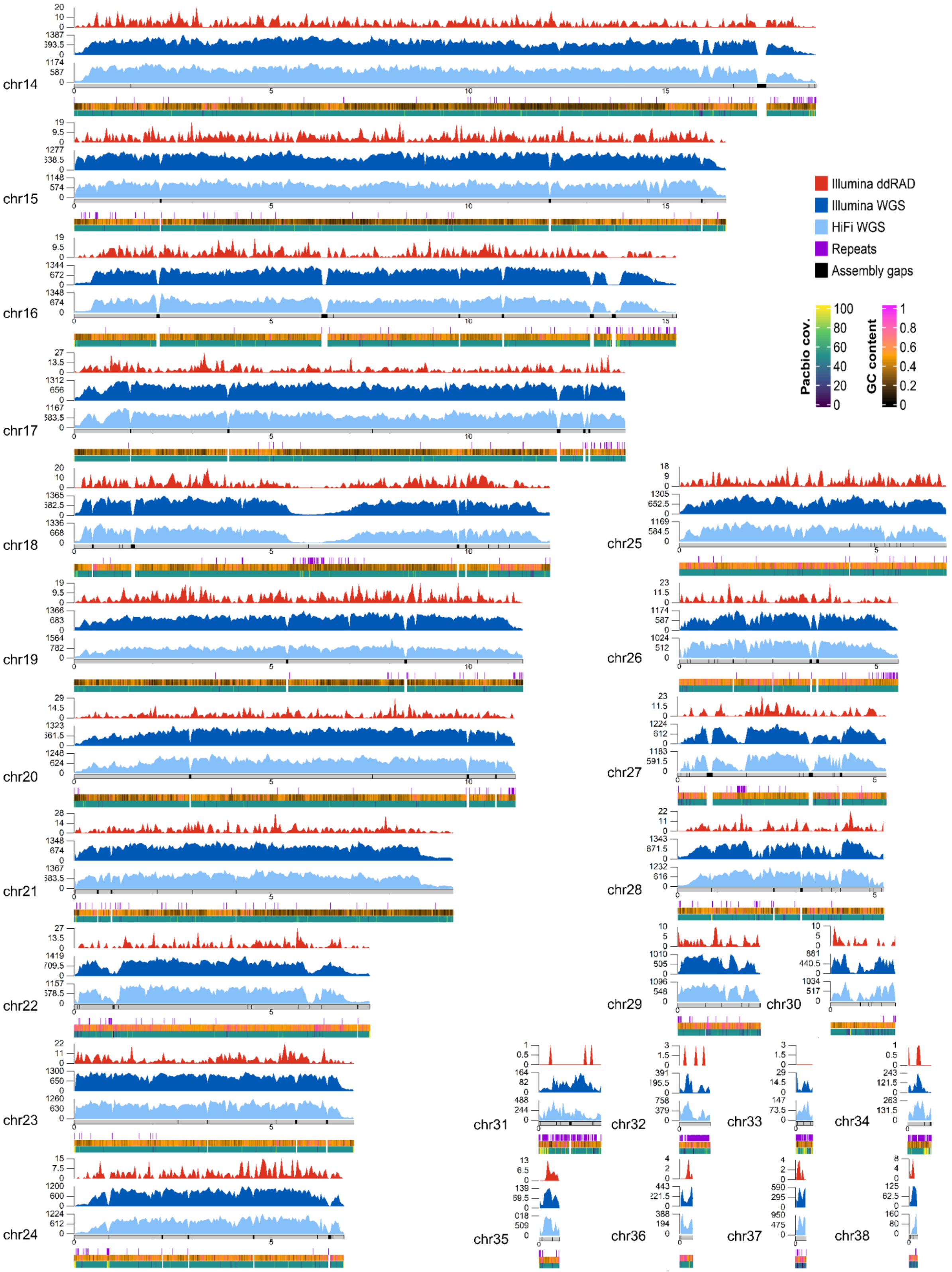

**Figure S13** SNPs density across microchromosomes. SNP density, coloured according to the different types of genomic data used, was computed over 40 kbp windows. The numbers on the y axis of each density track indicate the maximum and average values of SNP density for each track. Light blue: HiFi WGS data (ds1). Dark blue: Illumina WGS data from ds2 and ds3.1. Red: Illumina ddRAD data from ds3.2 through ds6.8. All available samples from the same sequencing technology were considered together. Additional tracks in the lower panel show repetitive regions of the genome (violet bars; only regions larger than 1 kbp are plotted), GC content and PacBio reads coverage. Grey ideograms represent chromosomes in scale, with assembly gaps highlighted as black bars.

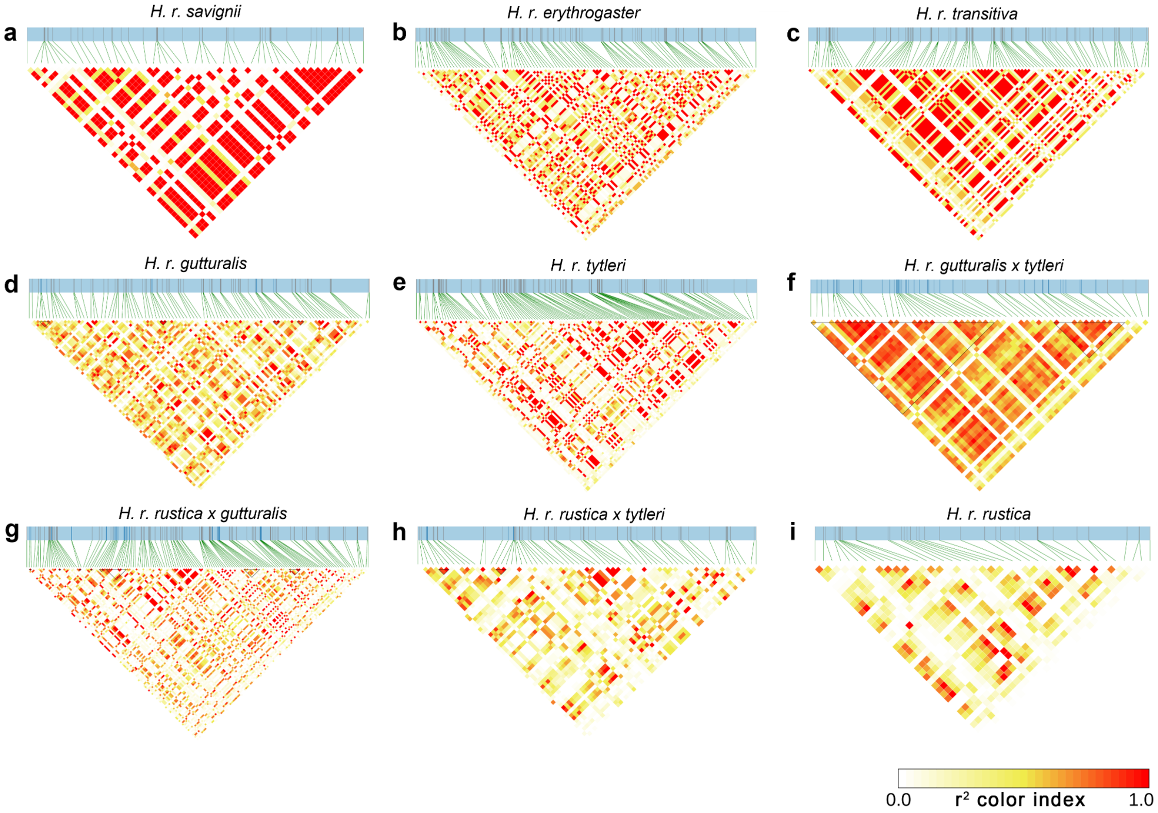

**Figure S14** LD heatmap within *BDNF* gene coordinates (chr6: 53886627-53927580) in Illumina WGS from dataset2 and dataset3.1.1. **a** *H. r. savignii* population (ds3.1.1). **b** *H. r. erythrogaster* population (ds3.1.2). **c** *H. r. transitiva* population (ds2.3). **d** *H. r. gutturalis* population (ds2.1). **e** *H. r. tytleri* population (ds2.4). **f** *H. r. gutturalis x tytleri* population (ds2.5). **g** *H. r. rustica x gutturalis* population (ds2.6). **h** *H. r. rustica x tytleri* population (ds2.7). **i** *H. r. rustica* population (ds2.2).

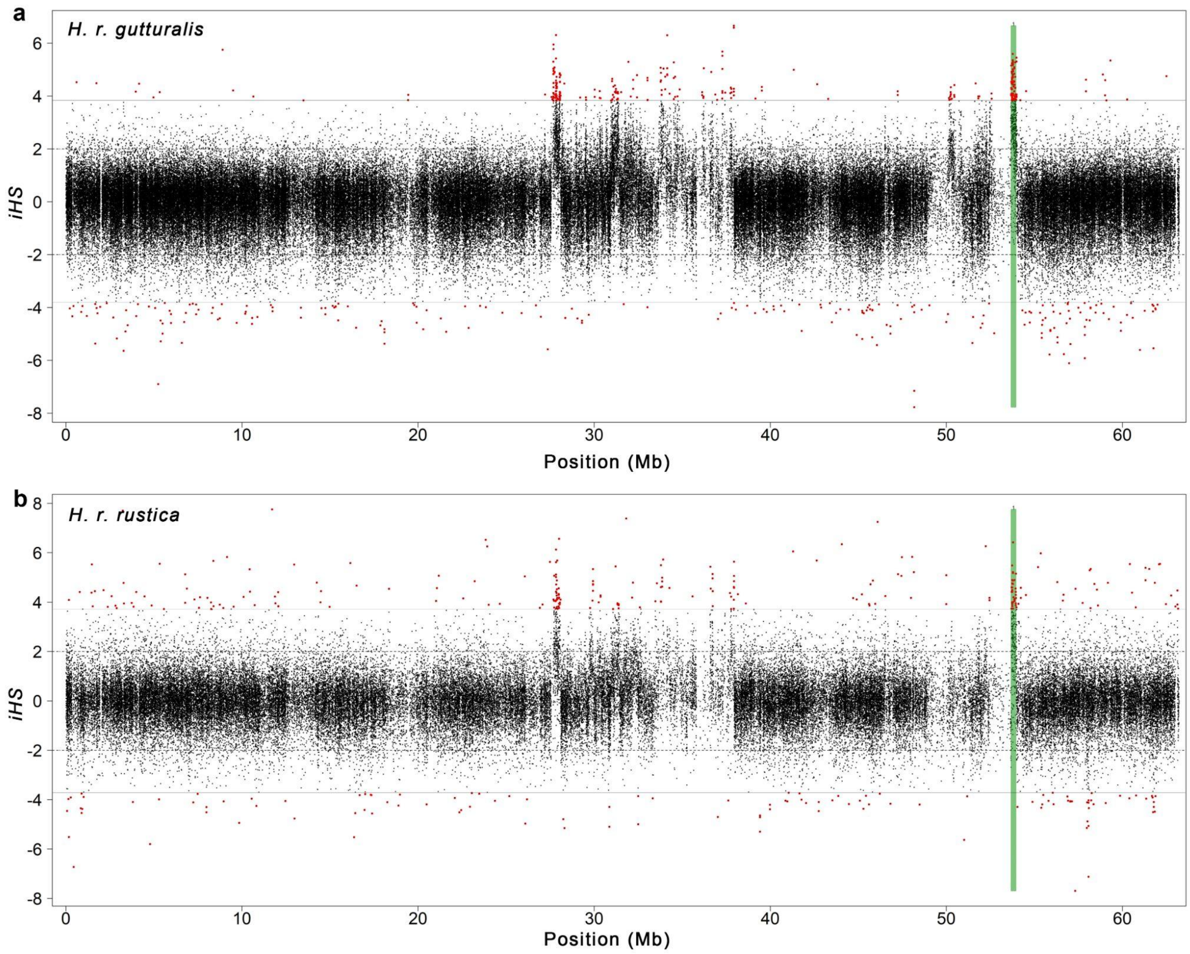

**Figure S15** Integrated haplotype homozygosity score (iHS) computed on chr6. Horizontal dashed lines (at +2, -2 values) represent the threshold identifying statistically significant iHS scores. Horizontal solid lines represent the threshold calculated after FDR correction. Statistically significant values after FDR correction are highlighted in red. The green vertical bars indicate the ROI with high LD values harbouring the *BDNF* gene (chr6:53680000-53954999). **a** *H. r. gutturalis* population (ds2.1). **b** *H. r. rustica* population (ds2.2).

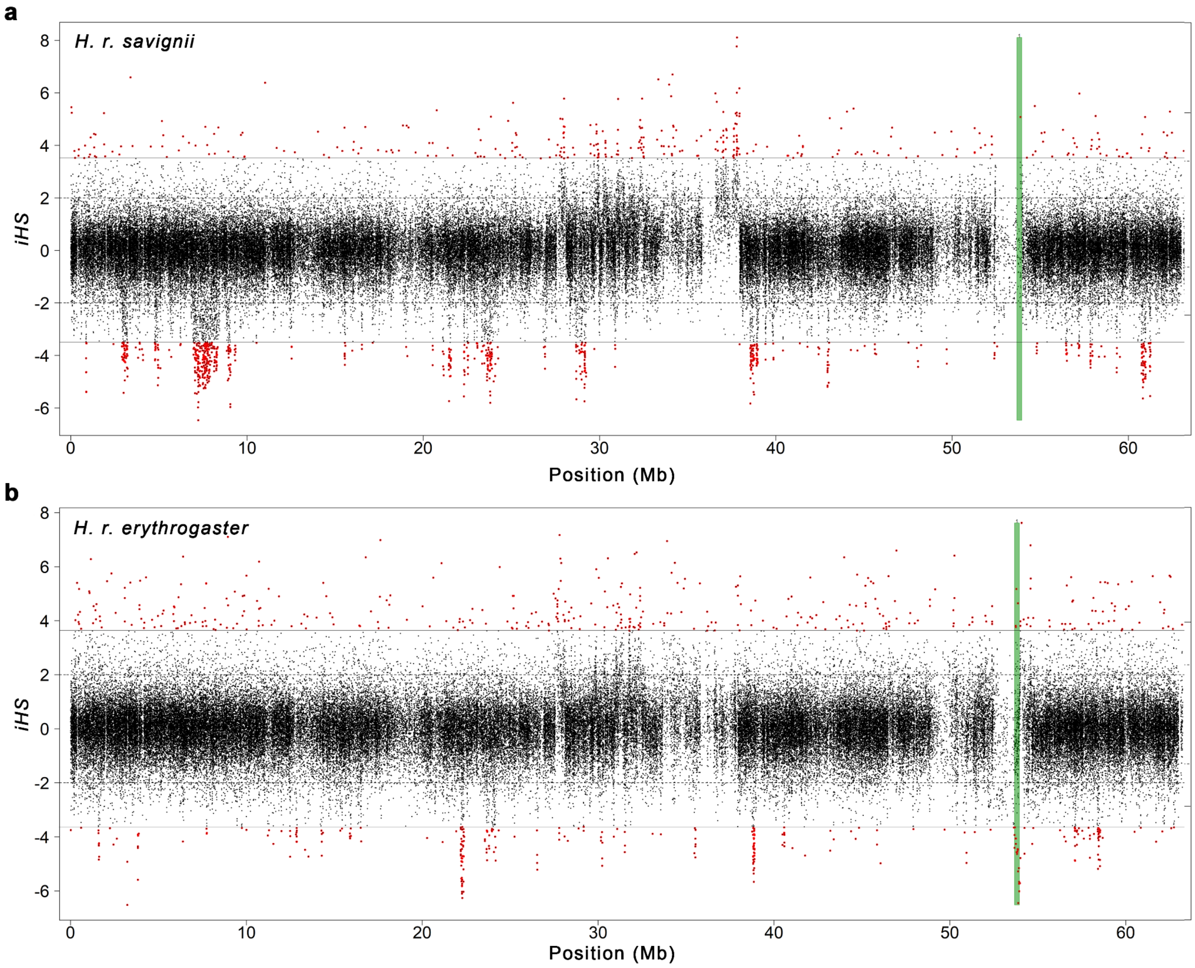

**Figure S16.** Integrated haplotype homozygosity score (iHS) computed on chr6. Horizontal dashed lines ( +2, -2) represent the thresholds for statistically significant iHS scores. Horizontal solid lines represent the threshold after FDR correction. Statistically significant values after FDR correction are highlighted in red. The green vertical bars indicate the ROI with high LD values harbouring the *BDNF* gene (chr6:53680000-53954999). a) *H. r. savignii* population (ds3.1.1). b) *H. r. erythrogaster* population (ds3.1.2).

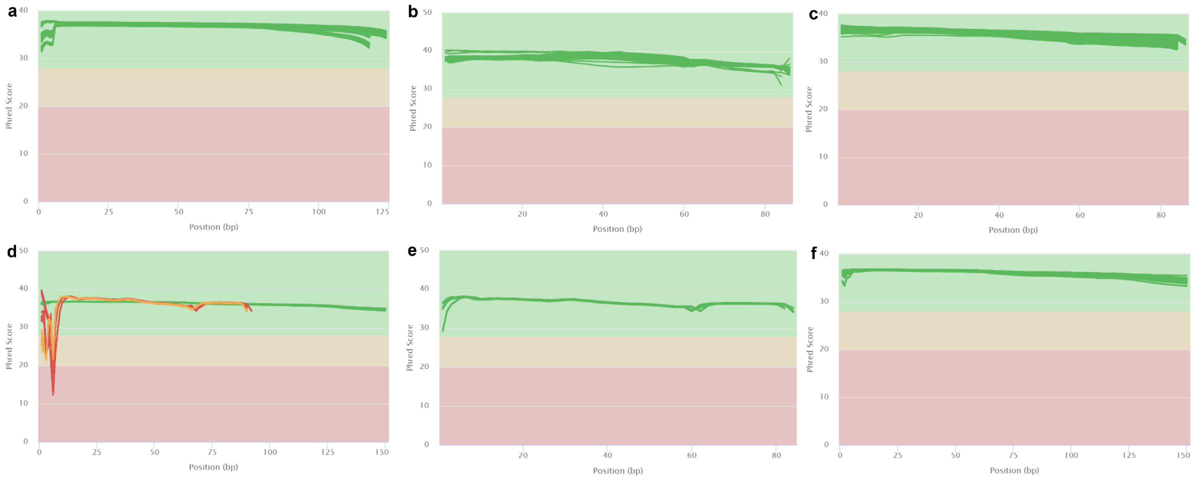

**Figure S17**. Per base quality scores (y-axis) from fastqc output performed on genomic data of the different datasets after adapter sequences trimming. **a** ds5; **b** ds6; **c** ds4; **d** ds3. American and Egyptian samples (ds3.2.1 and ds3.2.2) showed low quality scores at the beginning of the reads and were further processed to trim low quality bases; **e)** ds3.2.1 and ds3.2.2 after removal of low quality bases; **f** ds2.

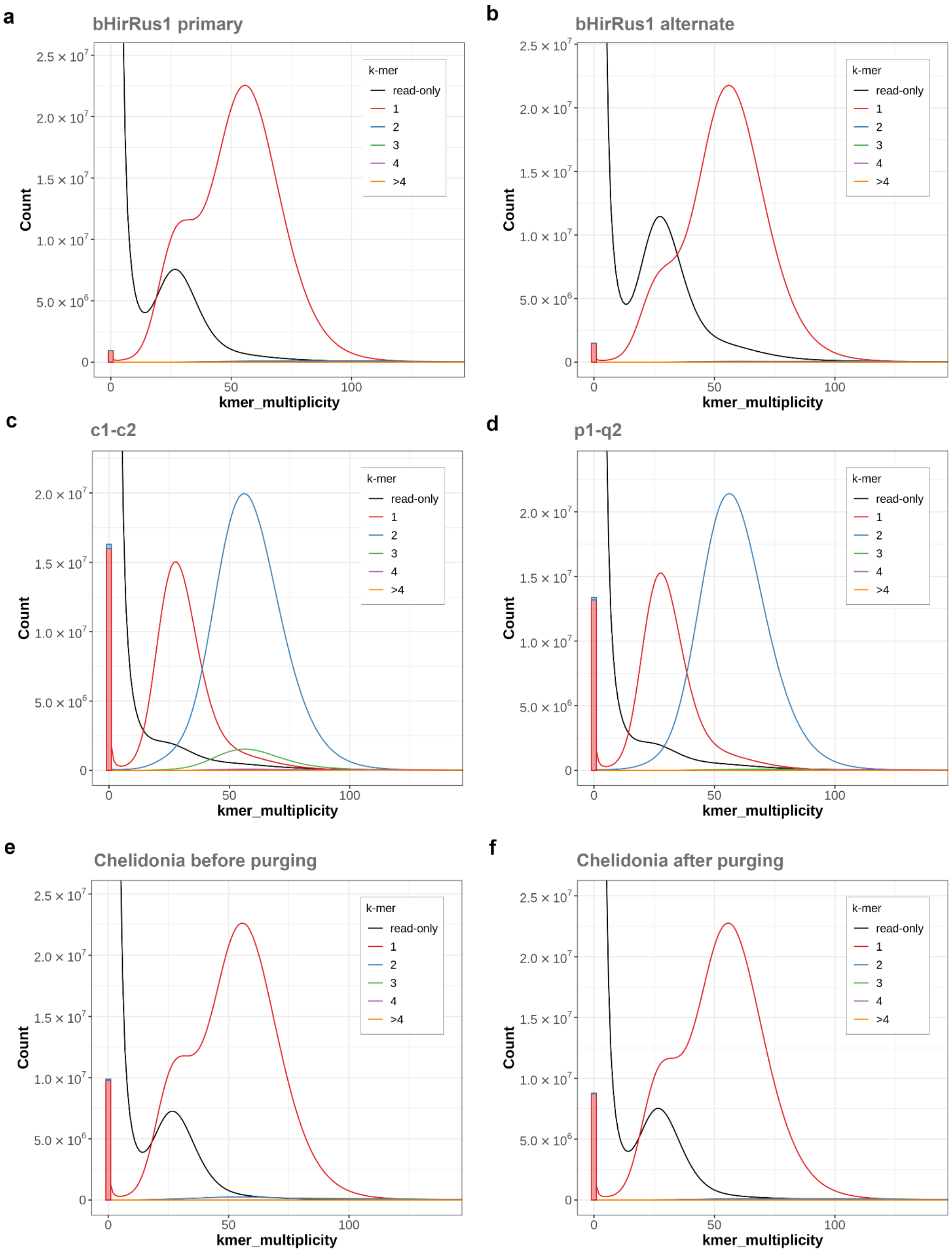

**Figure S18**. Merqury unstacked spectra-cn plots. **a** bHirRus1 primary assembly only. **b** bHirRus1 alternate assembly only; **c** Initial pseudo-haplotype assembly including primary contigs (c1) and alternate haplotigs (c2) generated by FALCON-unzip before purging. Retained haplotigs in the primary assembly show up as 3-copy k-mers (green curve) at about 50x (diploid coverage); **d** Pseudo-haplotype assembly after purge_dups (3) (p1 and q2 intermediates). Purging effectively removed 3-copy k-mers; **e** Chelidonia haploid assembly shows a higher number of false *k*-mers (bars at the origin) compared to bHirRus1 (a,b), as well as the presence of *k*-mers with diploid coverage in the read set (~50×) found twice in the assembly (blue curve); **f** Chelidonia after purge_dups (3). Duplicated k-mers were removed (blue curve reduced).

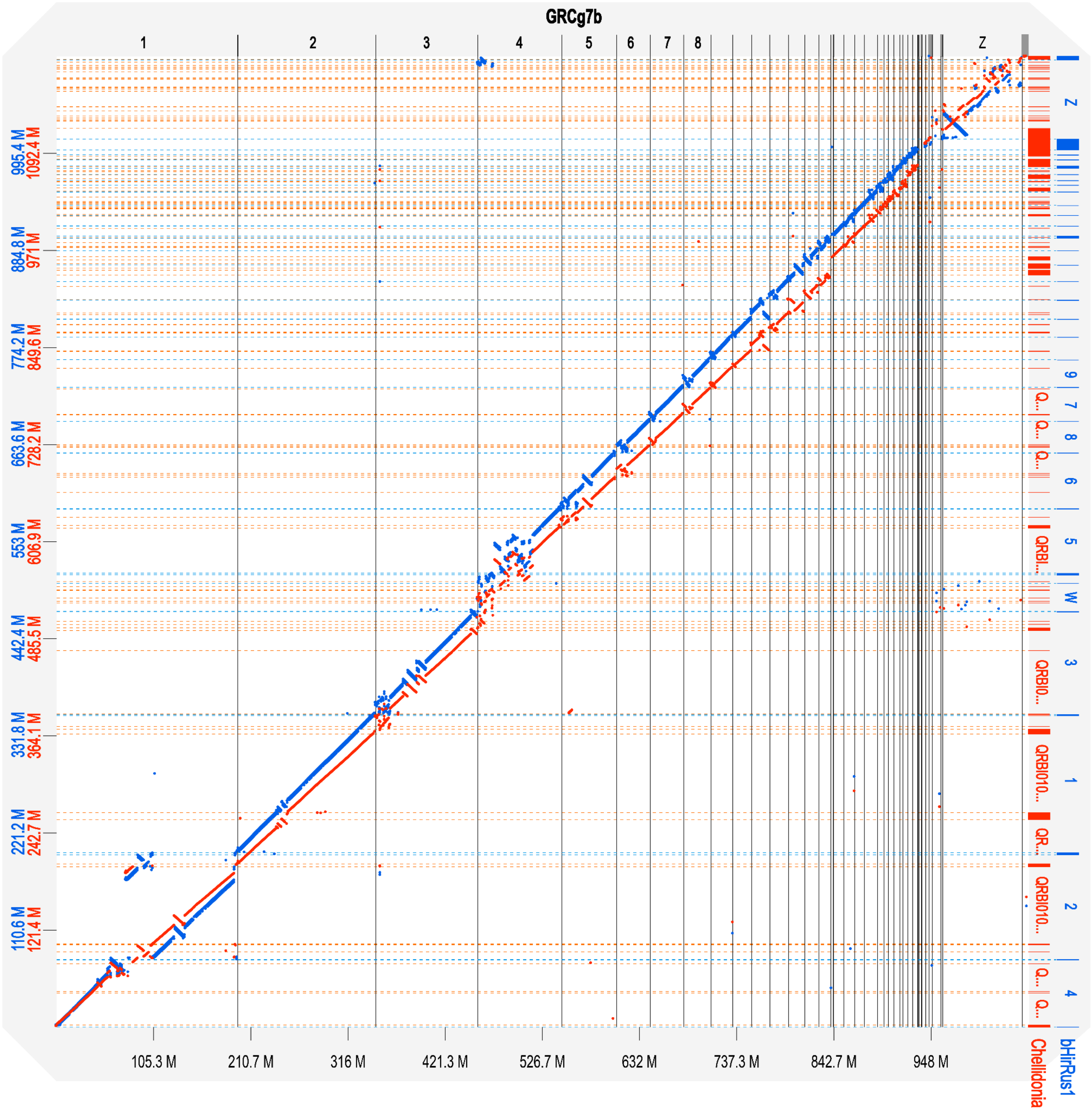

**Figure S19** Dotplot alignment of bHirRus1 (blue) and Chelidonia (red) with the latest VGP chicken assembly GRCg7b. Chromosome numbers and coordinates are reported for GRCg7b (x-axis), Chelidonia (y-axis, red) and bHirRus1 (y-axis, blue). Black vertical lines, red horizontal lines and blue dashed horizontal lines define chromosome and scaffold boundaries in the chicken assembly, in Chelidonia, and in bHirRus1, respectively.

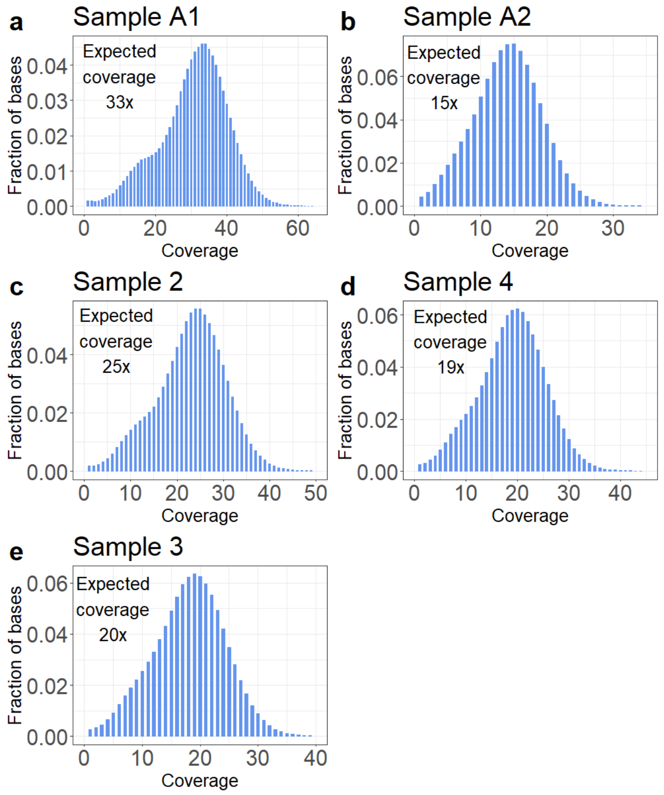

**Figure S20.** Histogram of alignment coverage for all samples of ds1. Per-base coverage was computed with bedtools genomecov.

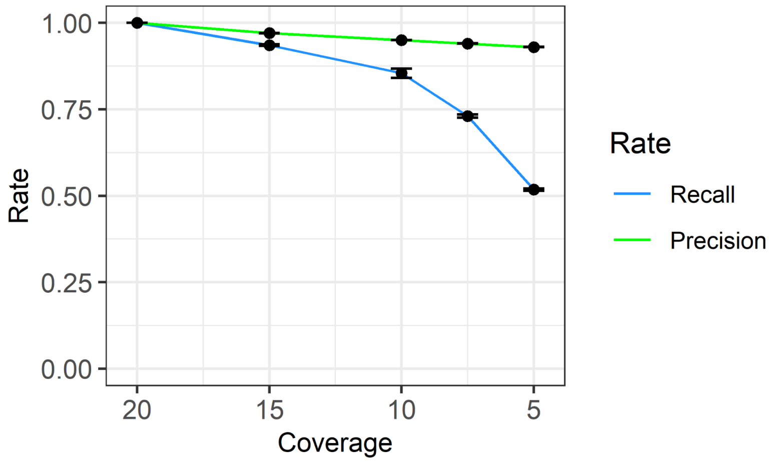

**Figure S21.** Recall and precision rates in individual titration experiments at varying HiFi coverage (x-axis). Each observation represents the average rate with standard deviation computed across all five samples (ds1) considering the full read set as truth (Supplementary Methods). Recall rate appears to sharply decrease between 10x and 5x, while precision rate does not decrease significantly at low coverage values.

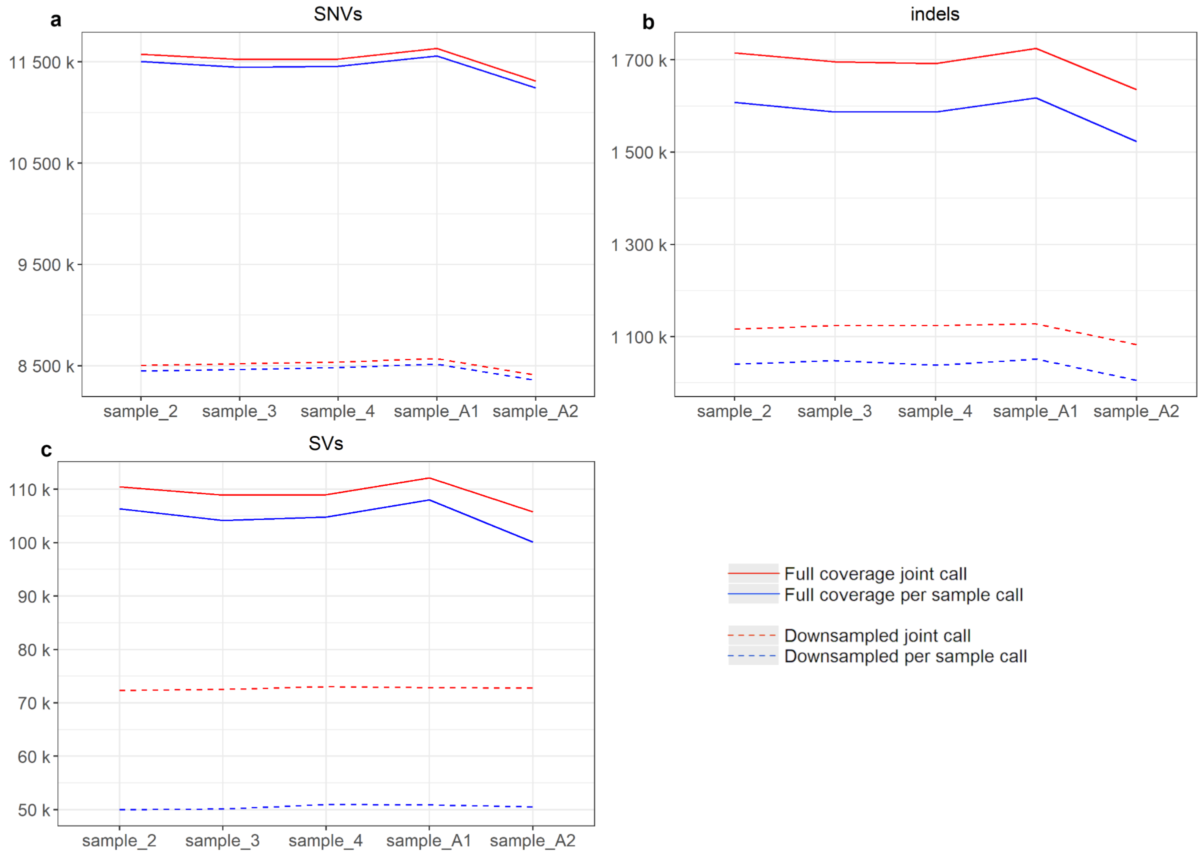

**Figure S22.** Absolute counts of joint and solo variant calls for the five HiFi read sets using the full or downsampled (5x) data sets. Variant calls shown for **a** SNVs, **b** indels (per-sample calls from DeepVariant, joint calls from DeepVariant and GLNexus), and **c** SVs (per-sample and joint calling with *pbsv)*.

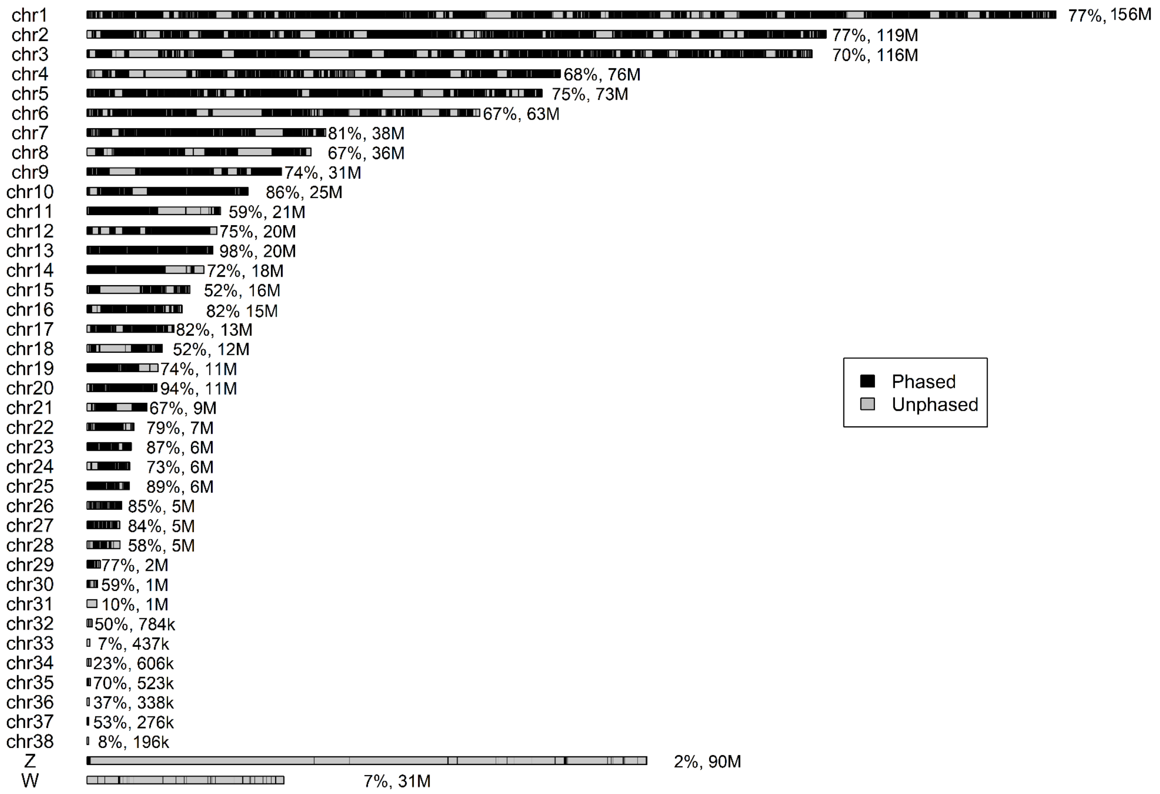

**Figure S23.** Haplotype phasing per chromosome with HiFi technology. Length of phased blocks (black) for sample A1 reported as example. Percentage of phasing and chromosome size is reported alongside each chromosome.

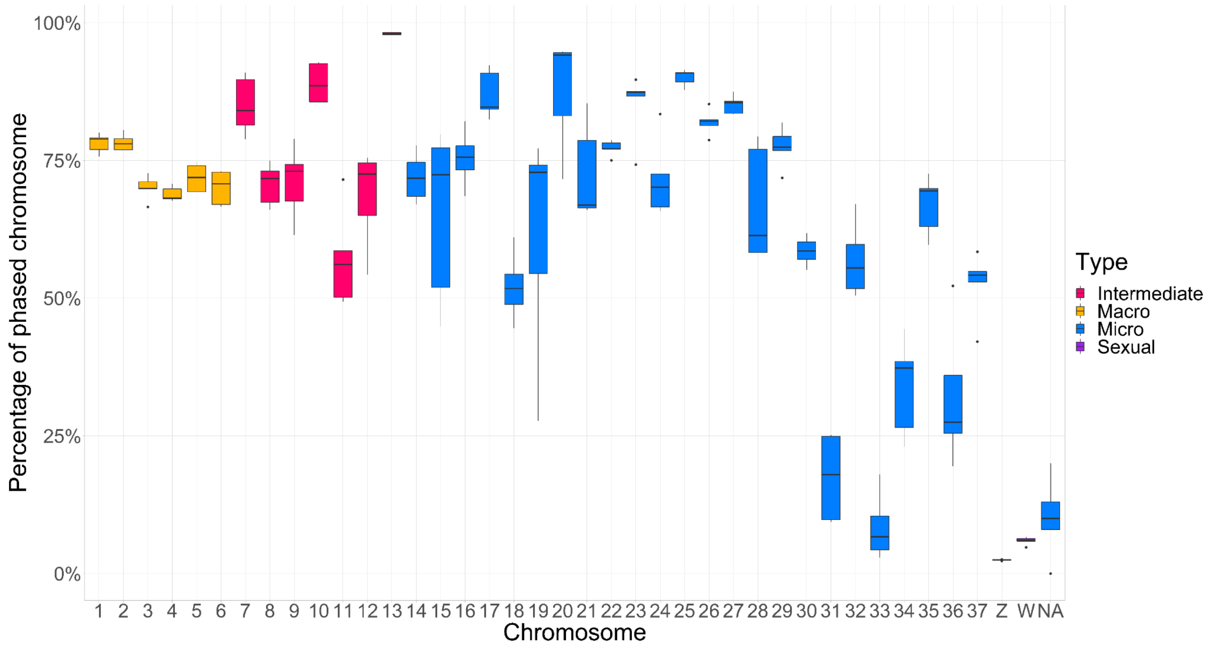

**Figure S24.** Haplotype phasing per chromosome with HiFi technology. The boxplot shows the percentage of phased chromosomes, coloured by type, across all samples.

### Supplementary Methods

#### 1 - bHirRus1 genome assembly

##### 1 - 1 Sampling

Tissues were collected from the same ringed barn swallow female whose blood was used for producing the previous barn swallow 'Chelidonia' assembly (8). The individual was recaptured in June 2018 in the same farm near Milan (45.4N 9.3E) and euthanized under permission N. 5104 issued on 11.04.2018 by Regione Lombardia. Tissues were dissected by an experienced avian veterinary, flash frozen immediately after dissection, and stored at -80 °C. The identity of the individual was further corroborated by manual inspection of read alignments of the newly generated reads to the Chelidonia assembly.

##### 1 - 2 DNA extraction

High molecular weight (HMW) DNA extraction was performed with the Bionano animal tissue DNA isolation fibrous tissue protocol (cat# RE-013-10;document number 30071). Approximately 55 mg of frozen muscle tissue was fixed in formaldehyde (2%) and homogenised with the Qiagen TissueRuptor. The lysate was included in agarose plugs, which were then treated with Proteinase K and RNase A. The DNA was recovered and purified from the plugs through a drop dialysis with 1x TE. Pulsed Field Gel Electrophoresis (PFGE; Pippin Pulse, SAGE Science, Beverly, MA) and Qubit were used for DNA quality control. According to the PFGE run, a large fraction of the isolated DNA was >250kbp.

##### 1 - 3 Library preparation and sequencing

Genomic data from four different sequencing technologies were used for the assembly: Pacific Biosciences (PacBio) CLR long-reads, 10x Genomics linked reads (short-reads), Bionano optical maps with one restriction enzyme (DLS) labelling, and Hi-C reads from Arima Genomics. PacBio long-reads and Bionano optical maps were reused from Chelidonia assembly (8). Linked-reads libraries were generated using the 10x Genomics Chromium platform (Genome Library Kit & Gel Bead Kit v2 PN-120258, Genome Chip Kit v2 PN-120257, i7 Multiplex Kit PN-120262) and sequenced on an Illumina NovaSeq S4 150bp PE lane at ~60X coverage. Hi-C libraries were generated by Arima Genomics (https://arimagenomics.com/) using muscle *in-vivo* cross-linking with the Arima-HiC kit (P/N: A510008) with 2-enzymes proximity ligation. Proximally-ligated DNA was subjected to shearing, size-selection (~200-600bp) with SPRI beads, and enrichment with streptavidin beads for the biotin-labelled DNA. KAPA Hyper Prep kit (P/N: KK8504) was employed to generate libraries compatible with Illumina technologies. The libraries were amplified through PCR and purified with SPRI beads. Libraries were sequenced on a Illumina HiSeq X (~60X coverage) after a quality check with Bioanalyzer and qPCR.

A quality control for each sequencing data type was performed with Mash (9) to detect potential outlier sequencing runs or species contamination. Mash was run with 21-mers to generate sketches of size 10,000. No contamination was detected.

##### 1 - 4 Genomescope2.0 predictions

Genomescope2.0 (7) (http://qb.cshl.edu/genomescope/genomescope2.0/) was used to estimate genome size, heterozygosity and repeat content through statistical analyses of *k*-mer profiles in unassembled sequencing data. A 31-mer database was generated with Meryl (10) using the 10x linked reads while trimming off the barcodes (i.e. the first 23 bp of the forward read).

##### 1 - 5 Mitogenome assembly

For the *de novo* assembly of the barn swallow mitogenome used in the polishing step, 10X reads were trimmed with the process_10xReads.py script from proc10xG (https://github.com/ucdavis-bioinformatics/proc10xG) with -a and -b 16 parameters. Trimmed reads were aligned to the Chelidonia assembly (8) with bowtie2 (11) and unmapped reads were extracted. NOVOplasty (6) was run with default parameters (read length = 151, insert size = 300) to assemble the mitogenome *de novo* from the unmapped reads. The mitogenome annotation was performed with MITOS2 (12). As sanity check, we aligned and mapped our complete mitochondrial sequence to the *Hirundo r. rustica* mitochondrial Reference Sequence (HrrRS, GenBank accession number MZ905359), which is included in a companion study on barn swallow mitogenome relationships (Lombardo et al., submitted).

##### 1 - 6 Assembly pipeline

The assembly was performed on the DNAnexus cloud-based informatic platform for genomic data analyses (https://www.dnanexus.com/) using the VGP standard genome assembly pipeline 1.6 (1) (https://github.com/VGP/vgp-assembly; Figure S1). The assembly was named 'bHirRus1' after the individual used for sequencing, which in turn is based on VGP guidelines for genome identifiers (1).

*1 - 6 - 1 Contigging, phasing and duplication removal*

PacBio subreads from Formenti et al. 2019 (8) were used in the first FALCON (13) contigging step. A genome size estimate of 1.31 Gbp (http://www.genomesize.com/) was used for read coverage calculation. Pre-assembled contigs underwent a phasing step with FALCON-unzip (2) (smrtanalysis 3.0.0) and a first round of Arrow (smrtanalysis 5.1.0.26412) polishing. FALCON and FALCON-unzip were run with default parameters, with the exception of parameters related to the identification of read overlaps. Raw reads overlaps were computed with DALIGNER options -k14 -e0.75 -s100 -l2500 -h240 -w8, and pre-assembled reads (preads) overlaps with DALIGNER options -k24 -e.90 -s100 -l1000 -h600. FALCON-unzip generated a set of primary contigs (labelled c1) representing the primary pseudo-haplotype, and a set of alternate haplotigs (c2), representing the secondary haplotypes (Figure S1). Purge_dups (3) was run on c1 primary contigs to remove any retained haplotig from the primary assembly, particularly in highly divergent regions, and to remove overlaps, collapsed repeats and low- and high-coverage contigs. Purged primary contigs (p1) were scaffolded, whilst all the alternate sequences were included into the p2 intermediate. The latter was merged with c2 alternate haplotigs and subjected to another round of purge_dups to remove additional haplotigs and overlaps. Purged alternate haplotigs (q2) were employed during the polishing step (Figure S1). To confirm the removal of haplotigs and overlaps, the evaluation tool Merqury (10) was run on primary and alternate contigs before and after purging.

*1 - 6 - 2 Scaffolding*

After purge_dups, a three-steps scaffolding strategy was performed on the p1 purged primary contigs using Illumina short-reads (10x Genomics), Bionano optical maps and Hi-C reads (Figure S1). To join proximal contigs, 10x linked reads were aligned to the p1 intermediate in two rounds and an adjacency matrix was produced from the barcodes using scaff10X v2.0-2.1 (https://github.com/wtsi-hpag/Scaff10X). Two scaffolding rounds were performed with options -matrix 2000 -reads 12 -link 10 and then -matrix 2000 -reads 8 -link 10. Contigs were then joined with 100 bp gaps (‘N’s). The resulting s1 intermediate was then scaffolded with Bionano DLS optical maps (8) using Bionano Solve v3.2.1 in non-haplotype assembly mode with a DLE-1 one enzyme non-nicking approach, obtaining s2. Finally, Hi-C reads from Arima were aligned to the s2 intermediate with the Arima Genomics mapping pipeline (https://github.com/ArimaGenomics/mapping_pipeline). Forward and reverse reads were aligned independently with BWA-MEM (14) with the -B8 parameter and filtered with a minimum mapping quality of 10. Reads containing a restriction enzyme site were trimmed at the 3’ end, and the aligned single reads were paired again. Processed alignments were employed for scaffolding with Salsa v2.2 (4) with -m yes -i 5 -p yes parameters and -e GATC, GANTC to indicate restriction enzymes used for library generation.

*1 - 6 - 3 Polishing*

Polishing was performed to improve the assembly per-base accuracy (QV) (1). We targeted Q40 (99.99% accuracy or 1 error/10 kbp) (1). To prevent haplotype switches and overpolishing of NUMTs (1,15), s3 scaffolded primary assembly was merged with q2 alternate combined haplotigs and the barn swallow mitogenome from NOVOplasty (6) (Figures S1-2). The s4 combined intermediate was polished with Arrow (pacific Biosciences; smrtanalysis 5.1.0.26412) with the command '`pbalign --minAccuracy=0.75 --minLength=50--minAnchorSize=12--maxDivergence=30--concordant--algorithm=blasr --algorithmOptions=--useQuality --maxHits=1 --hitPolicy=random --seed=1`' for read alignment, and with '`variantCaller --skipUnrecognizedContigs haploid -x 5 -q 20 -X120 -v --algorithm=arrow`' for consensus polishing, using PacBio CLR (t1). Two additional rounds of polishing with linked-reads were performed on t1, generating the t2 intermediate, and the final t3 polished assembly. In this step, raw-reads were aligned with Longranger align 2.2.2 and variants were called with Freebayes v1.2.0 (5) with default parameters. Finally, bcftools consensus (16) with options i'QUAL>1 && (GT="AA" || GT="Aa")' -Hla was used to generate the consensus.

*1 - 6 - 4 Manual curation*

Manual assembly curation entails the removal of contaminants and false duplications, the correction of structural assembly errors and the identification and assignment of chromosomal units. For bHirRus1, a dedicated decontamination pipeline, the genome evaluation browser gEVAL (geval.org.uk) and HiGlass Hi-C 2D maps were used (17). Since no reference for chromosome assignment was already established for the barn swallow, chromosomes were numbered in decreasing size order. A second curation step was performed using the results from BUSCO 4.1.4 (18–20), which indirectly assessed functional completeness through the prediction of highly conserved BUSCO vertebrate genes (complete, complete and single-copy, complete and duplicated, fragmented and missing). The absence, duplication or fragmentation of BUSCO genes can be evidence of assembly errors or missing sequences. BUSCO was run with the vertebrata_odb10 database and 'chicken' as training species for gene prediction on bHirRus1 and Chelidonia to assess differences in functional completeness, but also on the alternate assembly and the assembly pipeline intermediates c1, p1 and p2, to assess whether purge_dups (3) removed unintended sequences from the primary assembly. The BUSCO results were manually evaluated to detect missing genes in bHirRus1 that were found in the other assemblies, and could, therefore, be recovered. Nucleotide-nucleotide BLAST 2.10.1+ (21) was used to search in bHirRus1 the sequence of the missing genes retrieved from the corresponding assembly. These genes were erroneously not detected by BUSCO in bHirRus1. To confirm the presence of the genes found with BLAST and rescue the remaining bHirRus1 missing genes from the other assemblies, the scaffold or contig sequences containing the predicted BUSCO genes were aligned to bHirRus1 with MUMMer NUCmer (22). The alignment files were filtered maintaining only query alignment >1 kbp with an identity >98% with the reference sequence. Alignment coordinates were then manually evaluated. If the gene coordinates in the scaffolds failed to align to bHirRus1, the missing scaffold fragments were extracted from Chelidonia and the alternate assembly and added to bHirRus1. The rescued sequences were trimmed accordingly to avoid the insertion of duplicates and gaps. BUSCO and BLAST analysis were repeated on the new assembly version to confirm the addition of the rescued genes.

*1 - 6 - 5 Annotation*

Total RNA was extracted and purified using the QIAGEN RNAeasy kit (Cat. No. 74104). For each tissue type (brain and ovary), ~30mg was used, kept on dry ice and cut into 2mm pieces before being disrupted and homogenised with the Qiagen TissueRuptor II (Cat No./ID: 9002755). The RNA quality of all samples was measured using a Fragment Analyzer (Agilent Technologies, Santa Clara, CA) and quantified with a Qubit 2 Fluorometer (Qubit™ RNA BR Assay Kit - Catalog number: Q10210). PacBio Iso-seq libraries were prepared according to the "Procedure & Checklist – Iso-Seq™ Express Template Preparation for Sequel® and Sequel II Systems (PN 101-763-800 Version 01)". Briefly, cDNA was reverse transcribed using the NEBNext® Single Cell/Low Input cDNA Synthesis & Amplification Module (New England BioLabs, cat. no. E6421S) and Iso-Seq Express Oligo Kit (PacBio PN 10 1-737-500) from 300ng of total RNA for both brain and ovary. Amplified cDNA was cleaned with ProNex® Beads (Promega - Catalog numbers: NG2001). For each sample, a PacBio library was prepared using the Pacific Biosciences SMRTbell Express Template Prep Kit 2.0 (PN 101-685-400) following the manufacturer protocol. PacBio Iso-seq libraries were sequenced on a PacBio Sequel using sequencing chemistry 3.0 and with 20 hours movie time, 4 hour pre-extension and PacBio 1M v3 (#101-531-000) smrtcells. We sequenced one smrtcell for each Iso-seq library using sequencing kit 3.0 (#101-597-800). We then used the Iso-seq application in the Pacbio smrtlink package to generate Circular

Consensus Sequences (CCSs), re-move cDNA primers and concatemers, identified strandedness, trim polyA tails, and perform de novo clustering and consensus call to output high-quality full-length consensus isoforms. Truseq stranded mRNA libraries (TruSeq Stranded mRNA LT Sample Prep Kit/TruSeq Stranded mRNA Sample Preparation Guide, Part # 15031047 Rev. E) were generated and sequenced on a Novaseq6000 S4 lane (150bp PE) at Psomagen, Inc. A total of 6 libraries were sequenced: 2 for brain, 2 for ovary and 2 for muscle RNA samples. To obtain the coordinates of the different functional features of bHirRus1 (genes, exons, introns, CDS, 5' UTR, 3' UTR) for the following analysis, we parsed the NCBI annotation GFF3 file with GenomicFeatures (23) using a modified R script, excluding tRNAs, pseudogenes and C/V_gene_segments. Scripts used for this analysis can be found on GitHub (https://github.com/SwallowGenomics/BarnSwallow/tree/main/Analyses/GenomicFeatures)

#### 2 - Karyotype description and chromosome classification

##### 2 - 1 Karyotype construction

Tissue biopsies were obtained from a male *Hirundo r. rustica* sampled under permit N. 3268 issued on 12.03.2019 by Regione Lombardia. The sex of the individual was confirmed by PCR amplification of sex-specific genomic regions as described in Griffith et al., 1996 (24). Cells were cultured in a medium composed of 50% RPMI1640 and 50% Iscove's Modified Dulbecco's Medium, supplemented with 10% foetal bovine serum, 1% penicillin (10,000 units/ml) - streptomycin (10 mg/ml), 1% gentamycin sulphate (10 mg/ml), 0.5% amphotericin B (250 ug/ml) and 1% L-glutamine (200 mM) and incubated at 41°C with 5% $CO_2$. Chromosome preparations were made following standard procedures (25). Diploid number and chromosome morphology were determined from the analyses of 20 mitotic cells stained with 4 ′,6-diamidino-2-phenylindole (DAPI).

##### 2 - 2 Karyotype chromosome size estimations

Chromosomes sizes were estimated from four karyotype images using the chromosome_size software (https://git.mpi-cbg.de/dibrov/chromosome_size#example). The average size value was calculated for each chromosome. Sizes were correlated with the assembly chromosome sizes using Spearman nonparametric rank test (26).

##### 2 - 3 Chromosome classification

We assigned bHirRus1 chromosomes to the three typical avian chromosomal groups (macrochromosomes, intermediate chromosomes, microchromosomes), adapting the classification described by the chicken genome consortium (27). Here the authors assigned chromosomes ranging from 188 to 56.6 Mb to macrochromosomes, chromosomes from 33 Mb to 20 Mb to intermediates and chromosomes smaller than 20 Mb to microchromosomes. For the barn swallow genome, we designated chr7 (38.46 Mb) and chr8 (36.08 Mb) to the intermediate group, given their divergence in size with the larger macrochromosomes.

#### 3 - bHirRus1 evaluation

The commands used for the assembly evaluation can be found on the project GitHub page (https://github.com/SwallowGenomics/BarnSwallow/blob/main/Analyses/assembly_evaluation/assembly_evaluation.txt)

##### 3 - 1 Raw read alignments

Raw PacBio subreads were converted to fastq files with samtools (16) bam2fq 1.10. Each read set was aligned to both assemblies with bwa-mem (14) 0.7.17-r1188 and then converted to bam with samtools sort 1.10 with the -o option. The coverage was calculated from the bam file with mosdepth (28).

##### 3 - 2 Assembly statistics

Assembly metrics for all the assemblies were obtained with asm_stats.sh (https://github.com/VGP/vgp-assembly/blob/master/pipeline/stats/asm_stats.sh) with the mean predicted haploid genome size from Genomescope2.0 (1,241,727,742 bp; Table S5).

##### 3 - 3 Hi-C contact heatmaps

The three-dimensional conformation of chromosomes can be visualised as Hi-C interaction heatmaps through the alignment of the read set against the assembly. Contact maps were created from bwa-mem (14) alignments with PretextMap (https://github.com/wtsi-hpag/PretextMap) and visualised with PretextView (https://github.com/wtsi-hpag/PretextView).

##### 3 - 4 Masking of repetitive regions

The assemblies were soft-masked with WindowMasker 1.0.0 (29) and RepeatMasker 4.1.0 (30,31) (http://www.repeatmasker.org). RepeatMasker was run with NCBI/RMBLAST 2.10.0+ with Dfam_3.1 (profile HMM library) and Repbase (32) version 20170127 as repeat databases with the 'aves' repeat library. First, the genomes were processed separately with both tools. Then, 1-base repeat coordinates from RepeatMasker were used to further mask the Windowmasker-masked genome with bedtools maskfasta.

##### 3 - 5 QV, k-mer completeness and duplication content

Meryl (10) was used to count 21-mers from 10x linked reads that was then used in Merqury (10), a reference-free tool that computes per-base assembly accuracy (QV), completeness and *k*-mer multiplicity.

##### 3 - 6 Chromosome size and genomic content

Spearman nonparametric rank test (26) was used for the correlation between features and chromosome sizes, while Wilcoxon signed-rank test (33) was used to compare differences between microchromosomes and the other chromosomes. GC content was calculated with bedtools (34) nuc. CpG islands for bHirRus1 were downloaded from the UCSC browser (https://genome.ucsc.edu/cgi-bin/hgGateway). The fraction of the chromosomes covered by CG, CpG islands, genes and repeats (in percentage), was correlated with chromosome sizes (Table S3). Based on their high PacBio long-reads coverage (Table S3), microchromosomes 31, 33 and 34, representing approximately 0.2% of the assembly sequence (2.7 Mbp), were excluded from all correlation analysis.

#### 4 - Haplotig purging in Chelidonia

To confirm the presence of alternate haplotigs in Chelidonia and to investigate whether they affected *k*-mer and BUSCO (18,19) completeness, and increased the size of the assembly, we ran purge_dups (3) on Chelidonia with default parameters. The removal of retained haplotigs was evaluated with BUSCO (18,19), Merqury (10) and asm_stats.

#### 5 - Selection analysis on multiple whole-genome alignments

##### 5 - 1 Cactus alignment

Progressive Cactus (35) v1.3.0 was run with this command:

```
cactus --logInfo --logError --binariesMode local --workDir=/data/workDir jobStore SeqFile3.txt alignment.hal
```

The SeqFile.txt file contained the paths to the masked assembly files of the 10 bird species (Table S10) and the guide tree taken from TimeTree (36) (Figure S7) in Newick format. The alignment coverage was calculated with halAlignmentDepth (37) with the --noAncestors option and the barn swallow as the target species, excluding also the two species that did not align. Coverage was computed for each chromosome separately and then combined. The parameter --step 200000 was added to the command to generate track I of Figure 1. More details on the commands can be found on the project GitHub page (https://github.com/SwallowGenomics/BarnSwallow/tree/main/Analyses/Cactus%20alignment).

##### 5 - 2 Neutral model estimation and phyloP analysis

The command used to estimate the neutral model can be found on GitHub (https://github.com/SwallowGenomics/BarnSwallow/blob/main/Analyses/Selection%20analysis/neutral_model_estimation.txt), together with commands used to perform the phyloP analysis (https://github.com/SwallowGenomics/BarnSwallow/blob/main/Analyses/Selection%20analysis/phyloP_analysis.txt).

##### 5 - 3 phastCons parameters tuning

The same neutral model used for the phyloP analysis was used for phastCons (38) analysis. Given the low number of species and the high number of sites in our alignment, point 4.1 of PhastCons HOW TO guide (39) was followed. The initial length expected for phastCons was guessed at 20 bp, while the target coverage, which is the fraction of bases expected to be conserved, was set at 0.174. This value was calculated as the ratio between the expected conservation fraction (13.2% (40)) and the mean mappability between the barn swallow and the aligned genomes (76%; Table S10). The parameters were tuned such that around 65-70% of the CDS bases were covered by phastCons conserved elements (CEs) (41,42) and the smoothing PIT was around 10

(39,42). Briefly, each chromosome MAF file extracted for phyloP analysis was split into 1 kbp chunks and 200 chunks were randomly selected from the set. PhastCons was run on each sampled chunk with the --no-post-probs and --gc 0.425 tuning options, using the initial expected length and coverage, as well as the previously generated 4d non-conserved neutral model. The parameters, initially estimated separately, were averaged with phyloBoot (38), obtaining tuned conserved and non-conserved neutral models, which were then used by phastCons to predict conserved elements and conservation scores on each chunk. The smoothing level was checked with conEntropy (38) and coverage between CDS and the predicted CEs was manually verified. The analysis was repeated until the desired smoothing and coverage were reached (--target-coverage 0.22 --expected-length 8). The commands used for this analysis can be found on GitHub (https://github.com/SwallowGenomics/BarnSwallow/blob/main/Analyses/Selection%20analysis/phastCons_analysis.txt).

##### 5 - 4 Gene ontology (GO) analysis

The GO analysis was performed with the *gage (43)* R package. Previous to *gage* analysis, *bioMart (44)* R package was used to retrieve correspondence between the zebra finch and human Ensembl IDs and associate the latter with GO terms. The zebra finch annotation was used since the barn swallow could not be found on Ensemble yet at the time of the analysis. Human genes were used since annotation with GO terms would be more accurate. The script used can be found on the project GitHub page (https://github.com/SwallowGenomics/BarnSwallow/tree/main/Analyses/Gene_ontology).

#### 6 - Generation of the pangenome

##### 6 - 1 Sampling, DNA extraction, and PacBio Hifi sequencing

~100 µl of blood from five Italian barn swallows (*H. r. rustica*), were collected in heparinized capillary tubes through a minimally invasive sampling procedure in June 2019 (sample A1 and A2), July 2020 (sample 2), April 2019 (sample 3) and May 2019 (sample 4). Sampling was performed under permission 3268 of 12.03.2019 by Regione Lombardia. Sampling locations are reported in Table S19. High Molecular Weight (HMW) DNA was extracted from whole blood for samples A1 and A2, while for the other samples the starting material was centrifuged blood. Circulomics Nanobind Tissue Big DNA kit (SKU NB-900-701-01) was used to extract HMW DNA, following manufacturer's instructions. DNA absorbance was checked as quality and purity control by Nanodrop and average fragments length was verified with a Pulsed Field Gel Electrophoresis (PFGE). To perform PFGE, the Pulsaphor system with a hexagonal electrode array (Amersham Pharmacia Biotech) was employed. Genomic DNA was loaded on a 1% agarose gel in 0.5X TBE (running conditions: 165V, 60 sec pulses for the first 12 hours, 90 sec pulses for the last 12 hours; 8°C). Gel was stained with Ethidium Bromide 2 µg/ml in TBE 0.5X for 30 minutes; to acquire images, Geldoc (Bio-Rad) was used. To perform a second round of sequencing and achieve a higher coverage, DNA was re-extracted from samples A1,2,3,4 using the Qiagen Genomic tip columns and protocol. HMW DNA was sequenced by a PacBio sequencing service provider at Brigham Young University, Provo, UT (USA), where it was sheared using a Megaruptor 3. Libraries were prepared using the PacBio "SMRTbell express template Prep kit 2.0". Final size selection was performed using the Blue Pippin.

##### 6 - 2 Pangenome assembly

Hifi reads were screened for PacBio adapter contamination and then processed with cutadapt v3.2 (45). Reads were assembled with Hifiasm (46). Genomescope2.0 (7) was used to predict assembly statistics from HiFi raw data (Table S30). Both primary and alternate assemblies were purged using purge_dups (3) with the minimap2 option -xasm20 and custom cutoffs (Table S31) (47). The two cutoffs were calculated starting from the *k*-mer coverage (kcov) computed by Genomescope2.0 (7) (value1 = kcov*1.5, value2 = value1*3). The assemblies were masked with WindowMasker 1.0.0 (29) and RepeatMasker 4.1.0 (30) to reduce the alignment computational time (35). The Cactus Pangenome Pipeline included in Cactus (35) v1.3.0 was run as described in the software documentation (https://github.com/ComparativeGenomicsToolkit/cactus/blob/master/doc/pangenome.md). Briefly, Minigraph (48) v0.14-r415 was used to generate a GFA graph starting from the HiFi primary and alternate assemblies and bHirRus1 primary and alternate assemblies with the -xggs preset. Then, cactus-graphmap was used to align the input fasta sequences to the minigraph. Cactus-align was then used to run Cactus in pangenome mode to generate both a HAL alignment and a vg graph starting from the previous alignment. The commands used for the assembly of the pangenome and subsequent ortholog analysis can be found on the project GitHub page (https://github.com/SwallowGenomics/BarnSwallow/tree/main/Analyses/Pangenome)

#### 7 - SNP catalogue generation

##### 7 - 1 Publicly available datasets

To generate the catalogue of genetic variants, all publicly available datasets were used (Table S24). Raw reads were downloaded using fasterq-dump v2.9.1 from SRA Toolkit (https://github.com/ncbi/sra-tools). Data was single-end, except WGS data in ds2 and ds3.1 and ddRAD data in ds5. Quality control was performed on all raw reads using Fastqc v0.11.9 (https://www.bioinformatics.babraham.ac.uk/projects/fastqc/) and Multiqc v1.9 (49) (https://github.com/ewels/MultiQC). Low quality bases were trimmed using Cutadapt v2.10 (45) (Figure S17). BBDuk, from BBMap v38.18 (50) was used to remove Illumina adapters (k = 23, max mismatches = 1). Fastq files were aligned to bHirRus1 reference genome using bowtie2 v2.4.1 (11). The unmasked genome was used as reference. For WGS data, duplicated reads were removed using the Picard MarkDuplicates tool v2.23.4 (http://broadinstitute.github.io/picard/). Samtools v1.9 (51) (https://github.com/samtools/samtools) was used to sort and index alignments. Alignment files generated from paired-end genomic data were further processed with Bam clipOverlap software v1.0.14 (https://genome.sph.umich.edu/wiki/BamUtil:_clipOverlap) to trim overlaps between paired reads. The complete pipeline used to download and align reads is available on the project github page (https://github.com/SwallowGenomics/BarnSwallow/blob/main/Analyses/popgen_data_download_alignment/popgen_pipeline.bash).

##### 7 - 2 Variant calling and filtering

Freebayes v1.3.1 (5) (https://github.com/freebayes/freebayes) was used to call variants. To reduce computational time, a script adapted from the VGP assembly pipeline (https://github.com/VGP/vgp-assembly/blob/master/pipeline/freebayes-polish/freebayes_v1.3.sh) was used to parallelize the process by subsetting the reference genome by scaffold. Variants were called with the options --min-mapping-quality 10 --min-base-quality 20 --populations (all other parameters were left to default). Due to the lower sequencing coverage, --min-alternate-count 0 was used for ds6. The coordinates of the repetitive regions were extracted from the masked reference genome with a python script (https://gist.github.com/danielecook/cfaa5c359d99bcad3200) and the unmasked regions identified with bedtools v2.29.2 (34) using the complement command. All vcf files were first filtered to remove variants falling within repetitive regions, multiallelic SNPs and indels. Further filtering steps and thresholds are detailed in Table S33.

We removed sites showing more than twice the mean read depth across samples (INFO/DP field). In the vcf generated by Freebayes, genotype quality is expressed as QR (quality reference) and QA (quality alternate). We marked as missing all genotypes in which both values were below the threshold reported in Table S33. For FMT/DP filtering, we used as maximum value twice the average DP value and we approximated the 5% quantile of the distribution to set the minimum value. Individuals presenting a high amount of missing data (> 70%) were discarded (Table S33). All filters were applied using bcftools v1.1 (51) (https://github.com/samtools/bcftools) with the view and filter commands, except the removal of variants falling within repetitive regions, performed with bedtools v2.29.2 (34) using the intersect command and the coordinates of the unmasked regions previously identified. Standard statistics from the VCF files (in particular average site depth and average individual depth) were calculated using VCFtools v.0.1.16 (52) (https://github.com/vcftools/vcftools). An example of the complete set of commands used to filter variants (from ds2.2) can be found here (https://github.com/SwallowGenomics/BarnSwallow/blob/main/Analyses/variants%20filtering/filtering_commands.txt).

**7 - 3 SNP density statistics and correlations with genomic features**

For all the analyses described in Supplementary Methods sections 7-3 and 7-4, all datasets generated with the same sequencing technology were combined (HiFi WGS; Illumina WGS; Illumina ddRAD). SNP density for each chromosome was computed on 10 kbp windows and SNPs were counted using bedtools v2.29.2 (34) with the coverage -counts option. Average SNP density for each chromosome was then calculated in R using the weighted mean function. Mean value was weighted for the window size to take into account truncated windows potentially present at chromosome ends. The average SNP density values across all chromosomes for each sequencing technology was calculated in R using the weighted mean function (in this case the mean was weighted for the chromosome size). For the HiFi dataset (ds1) also a 5x downsampled HiFi dataset was generated (Supplementary Methods Section 9-2, first titration experiment) considering the 20x read coverage of each sample (except for the A2 sample, starting from 15x) as the truth set (variants from the 5x reads set were intersected with variants from the 20x reads set using bedtools v2.29.2 (34) with the intersect command). For each chromosome and dataset, SNPs falling in intervals corresponding to genic, intergenic, exonic and intronic regions as determined from NCBI annotation were counted using bedtools v2.29.2 (34) with the coverage -counts option. To analyse correlations between SNP density and GC content in our catalogue, the GC content was calculated using bedtools v2.29.2 (34) with the -nuc option on 10 kbp windows and SNPs were counted every 10 kbp window. Correlation was tested in R computing the Spearman nonparametric rank test (26) with the R function cor.test. Unlocalized/unplaced scaffolds were excluded from the analysis. bedtools v2.29.2 (34) was used to divide the genome in 10 kbp windows, using the makewindows command with the -w 10000 flag.

##### 7 - 4 SNP density plotting

To plot SNP distribution across chromosomes, SNP density was computed with the R (53) package *karyoploteR (54)*. Additional tracks included repetitive regions, GC content, raw reference reads coverage and assembly gaps. Repeats were annotated by Windowmasker 1.0.0 (29) and Repeatmasker 4.1.0 (30,31). GC content was calculated using bedtools v2.29.2 (34) with the -nuc option on 1 kbp windows. Per base coverage of raw reference reads was calculated by aligning reads back to the bHirRus1 assembly and using bedtools v2.29.2 (34) with the genomecov -d option. Values were then averaged every 500 bp (https://github.com/SwallowGenomics/BarnSwallow/blob/main/Analyses/coverage_analysis/avg_coverage.bash). Standardised values were attributed to specific coverage intervals: 0 for low coverage (between 0 and 10), 100 for regions showing double the average coverage value (95), or higher, and intermediate fixed values for coverage between 10 and 95. Assembly gaps were removed from computation of GC content, repeat content and PacBio reads coverage.

#### 8 - LD and haplotype statistics analysis

##### 8 - 1 Genome-wide LD decay

LD decay was evaluated using $r^2$ from Freebayes v1.3.1 (5) variant calls. $r^2$ values were calculated using Plink v1.9 (55). To estimate LD decay trend across the whole genome in filtered ds2 and ds3.1, we considered marker pairs within a 55 kbp distance with the option --bcf file.bcf --r2 dprime yes-really --ld-window 999999 --ld-window-kb 55 --ld-window-r2 0 --allow-extra-chr --out LD55kb. Option –ld-window 999999 is required to consider variant pairs more than 9 lines apart from each other (56). To calculate average $r^2$, SNP pairs were grouped according to their distance in bins of 1 kbp (range 1-55 kbp) using a custom perl script (https://github.com/SwallowGenomics/BarnSwallow/blob/main/Analyses/LD-scripts/LDaverage.pl), that was run on Plink output. The same approach was used to calculate average $r^2$ values per chromosome group (macrochromosomes, intermediate and microchromosomes), except that values were then averaged across specific distance bins. Sex chromosomes were excluded from the chromosome group LD analysis.

**8 - 2 LD scans**

Before performing the LD scans, variants were filtered with bedtools v2.29.2 (34) using as maximum coverage (95x) twice the average PacBio reads coverage genome wide (47.7x) and 10x as the minimum, so to ensure the exclusion of SNPs falling within collapsed or ambiguous regions of the genome. We used Plink v1.9 (55) with options --bcf file.bcf --r2 dprime yes-really --ld-window 10000 --ld-window-kb 15 --ld-window-r2 0 --allow-extra-chr --out LD15kb on Illumina WGS data from American and Egyptian samples (ds3.1). To scan for genes showing high LD values, $r^2$ was chosen as it is generally more informative for small datasets and also more consistent with allele frequency variation (57), whereas D′ can be more prone to inflation. To compute the average LD, each scaffold was divided in sliding non-overlapping 5 kbp windows with a custom perl script (https://github.com/SwallowGenomics/BarnSwallow/blob/main/Analyses/LD-scripts/chr_ld.pl), requiring a minimum of 100 markers per window. Only genomic windows with average $r^2 > 0.3$ were extracted (Table S28). The threshold was chosen based on similar studies (57,58). Coordinates were intersected with the NCBI annotation to find genes potentially carrying alleles in LD using bedtools v2.29.2 (34). For further analysis, two 5 kbp intervals were joined into the same ROI if the distance between them was lower than 100 kbp. Intervals showing high LD values were excluded if in proximity (within ~5 kbp) with potentially collapsed or low-confidence assembly regions (considering a PacBio reads coverage value higher than twice the average genome-wide coverage or lower than 10, respectively) or if not carrying any annotated gene. For Fig. 5a, we computed the average LD in correspondence with ROI 45 on chr.6 for the *H. r. savignii* and *H. r. erythrogaster* populations separately, requiring a minimum of 10 markers per window. For *BDNF*, LD heatmaps were generated using LDBlockShow v1.36 (59) (https://github.com/BGI-shenzhen/LDBlockShow) with the options -InVCF file.vcf -OutPut Scaffold_name -Region Scaffold:start-end -OutPng -SeleVar 2. CpG islands along the *BDNF* sequence were identified with cpgiscan v1.0 (60) (https://github.com/jzuoyi/cpgiscan), combining neighbouring CpG islands when their distance was lower than 100 bp.

**8 - 3 iHS computation**

To calculate iHS, namely the standardised log-ratio of the iHH (integrated haplotype homozygosity) values for the two alleles, variants present on chr6 were phased with WhatsHap v0.18 (61) (https://github.com/whatshap/whatshap) and the Rehh (62) R package was used. Before iHS computation, variants were filtered to remove sites showing a fraction of missing genotypes across samples higher than 0.1 and sites with maf < 5%, using Rehh filtering options min_perc_geno.mrk = 90 and min_maf = 0.05. Extended haplotype statistics were then calculated using the scan_hh (with the polarised = FALSE option) and the ihh2ihs (setting freqbin = 1) functions. To perform FDR correction, the qvalue R package was used (https://github.com/StoreyLab/qvalue). This analysis was performed on populations relative to ds3.1, ds2.1 and ds2.2. The complete list of commands used for iHS computation can be found here (https://github.com/SwallowGenomics/BarnSwallow/blob/main/Analyses/iHS%20analysis/iHS_analysis_script.R).

#### 9 - HiFi reads processing for SNP catalogue, titration and phasing experiment

##### 9 - 1 HiFi reads alignment, variant calling and filtering

HiFi reads from ds1 samples were aligned to bHirRus1 with pbmm2 v1.3.0 (https://github.com/PacificBiosciences/pbmm2) using default parameters for PacBio CCS reads with the options align --preset CCS --sort -j 32 --log-level INFO reference.mmi reads.ccs.bam file.aligned.ccs.bam. The genome-wide coverage of mapped reads was computed with bedtools v2.29.2 (34) using the genomecov command. At first, alignments were used to call small variants using DeepVariant v1.0.0 (63) (https://github.com/google/deepvariant) with default parameters for PacBio reads individually for each sample. Variants were first filtered to remove multiallelic SNPs and indels. SNPs falling within repetitive regions were removed as described for the publicly available datasets. Next, only SNPs with a genotype quality value higher than 20 were kept, and 5% and 95% quantiles of the read depth values distribution were used to set the minimum and maximum site coverage. Filters were applied using bcftools v1.1 (51), and filtered variants from each sample were merged with the same tool to estimate and plot SNP density across chromosomes as described for Illumina WGS and ddRAD data. For the comparison between Illumina and HiFi technology, Samtools v1.9 (51) was used with the view command and the -q flag to exclude reads with a mapping quality value lower than 30 (for Illumina data) and 60 (for HiFi data), based on Hon et al (64). The proportion of the genome covered by the alignment was computed with bedtools v2.29.2 (34) with the genomecov -bg option. All bases with read depth ≥ 1 were extracted from bedtools output. HiFi joint variant calling of SNVs and indels was performed using gVCF files from DeepVariant v1.1.0 (63) per-sample calls, jointly called with GLNexus (65) pipeline (https://github.com/PacificBiosciences/pb-human-wgs-workflow-snakemake). For joint calling of SNVs and indels, DeepVariant v1.1.0 (63) was run twice, the second time after an intermediate variants phasing step performed with WhatsHap v1.0 (61). For SVs, *pbsv* v2.6.0 (66) (commit v2.4.1-155-g281bd17) (https://github.com/PacificBiosciences/pbsv) was used for per-sample and joint variant calling. The minimum SV length was set to 20 bp.

**9 - 2 Titration of HiFi reads**

Two downsampling experiments were conducted, the first one after individual variant calling and the second one after joint variant calling (N = 5). For the individual titration experiment, all HiFi reads were first downsampled to 20x coverage using Rasusa v0.3.0 (67) (https://github.com/mbhall88/rasusa), except for the A2 sample where the sequencing coverage was 15x. Three different truth sets were generated, first (truth set 1) using the vcf file derived from the 20x coverage alignment of each sample; second (truth set 2) by intersecting this 20x file with a set of publicly available barn swallow variants (dst3.1); third (truth set 3) from the intersection of all variants from the 5 samples at full sequencing coverage. Each read set was further downsampled at 15x, 10x and 5x, in triplicate for each condition. Reads were aligned to bHirRus1 and variants were called as described in section 9-1 for per-sample variant calling. Specific filters were applied as described in section 9-1. The three different truth sets were then intersected with the variants recovered after every titration using bcftools v1.1 (51) with the isec command and the -w1 flag. Recall rate, precision and F1 score were estimated for each titration experiment. The recall rate at the different coverage values was estimated as the number of shared variants after intersection divided by the total number of variants in the truth set for each sample, while the precision rate was estimated as the number of shared variants after intersection divided by the total number of variants identified in each particular titration replicate. The F1 score, the harmonic mean between recall rate and precision rate, was estimated as $F1 = 2 \times \frac{precision \times recall}{precision + recall}$. For the second titration experiment, reads were randomly downsampled using Rasusa v0.3.0 (67) tool as described above for the first experiment. Reads were aligned to bHirRus1 using pbmm2 v1.4.0, variants were called as described in 9-1 for joint variant calling and recall rate was estimated considering the full-coverage joint calling as truth set.

**9 - 3 Phasing of HiFi read sets**

Variants obtained with HiFi reads (ds1) were filtered to remove multiallelic SNPs and indels. Only SNPs with a genotype quality value higher than 20 were kept, and 5% and 95% quantiles of the read depth values distribution were used to set the minimum and maximum site coverage. Next, to estimate and plot haplotype-phased blocks length across chromosomes, variants were phased using WhatsHap development version v.1.2.dev2+g3dffe4a (61) with the options stats --chr_lengths --tsv.

#### 10 - Graphical representations

The R (53) package *ggplot2 (68)* was used to generate correlation plots (Fig.1b; Fig.2c), histograms (Fig.2d; Figure S8) and the gene presence-absence matrix (Fig. 2d). The R package *circlize* was used to generate Circos plots (Fig.1c; Fig.2c; Figure S2). SequenceTubeMap (69) was used to graphically represent pangenome regions (Fig.2a,b; Figure S10). MEGA X software (70) was used to generate the phylogenetic trees. The Hi-C contact heatmaps were visualised with PretextView (https://github.com/wtsi-hpag/PretextView). The *k*-mer profiles were generated with Genomescope2.0 (7) (http://qb.cshl.edu/genomescope/genomescope2.0/) and Merqury (10) (Figure S4b, S18). Snail plots were generated with BloobToolKit (71) (Figure S5a,b). Alignment dot plot was generated with D-genies (72) (Figure S19). Manhattan plots were generated with the R package *CMplot* (Figure S8a,b). IGV (73) was used to visualise aligned features to the genome (Figure S9).

R (53) package *karyoploteR (54)* was used to plot SNP density visualisation across all chromosomes (Fig. 3). SNP density was computed using the internal function kpPlotDensity using 40 kbp as window size, for the three types of sequencing technologies considered. To plot SNPs distribution across all chromosomes for the 5x downsampled HiFi dataset (Figure S12), the 20x read coverage of each sample (except for the A2 sample, starting from 15x) was used as the truth set (variants from the 5x reads set were intersected with variants from the 20x reads set before plotting). Both coverage and GC content were plotted with the kpHeatmap function. The heatmap relative to Pacbio coverage was generated using the *viridis* package. Repeats and assembly gaps were plotted using the kpPlotRegions function. Only repeats larger than 3 kbp (larger than 1 kbp for Extended Data Fig. 7, relative to microchromosomes) were plotted. The figure legend was generated using the *ComplexHeatmap (74)* package. Unlocalized/unplaced scaffolds were excluded. The R package *ggplot2* was used to plot genome-wide LD decay (geom_line function) and LD per chromosome group (geom_boxplot function) (Fig 4). After LD scans, LD values were plotted with the KaryoploteR (54) package using the kpPoints and kpLines functions. SNP counts for the two populations were plotted with the kpHeatmap function. The *BDNF* transcript isoforms structure was drawn using the *ggplot2* package. IGV (73) was used to visualise *BDNF* region containing previously annotated methylation sites from the Cactus multialignment (Fig. 5).

The map showing sampling locations from all datasets was generated in R using the packages *ggplot2 (68)*, *rnaturalearth*, *sf* and *rnaturalearthdata* (Figure S11). iHS values were plotted using the manhattanplot function of the *Rehh (62)* package (Figures S15; S16). Histograms of the HiFi reads coverage were generated with the *ggplot2 (68)* package using the geom_bar function (Figure S20). To plot recall rate values after HiFi titration experiments, the functions geom_line and geom_point of the *ggplot2* package were used. For the second titration experiment, the legend was generated using the *ComplexHeatmap (74)* package and plots were arranged together with the packages *grid* and *gridExtra* (Figures S21; S22). Before plotting phased blocks length, the WhatsHap development version v.1.2.dev2+g3dffe4a (61) command stats --gtf was used to generate a .gtf file with the size and position of the phased blocks. Phased blocks computed from HiFi reads were plotted with the *KaryoploteR (54)* package using the kpRegions function (Figure S23). The percentage of phased chromosomes, coloured by type, averaged across samples, was plotted with the *ggplot2 (68)* function geom_boxplot (Figure S24). See this github section (https://github.com/SwallowGenomics/BarnSwallow/tree/main/Plots%20and%20figures) to retrieve the lists of commands used for all figures and plots.

### Supplementary Note

#### 1 - bHirRus1 manual curation

##### 1 - 1 First curation step

A first round of manual curation performed at the end of the assembly pipeline (Figure S1) introduced 174 rearrangements by breaking and joining scaffolds and resulted in the removal of 8 false duplications. This reduced the genome length by 3.2 Mb, decreased the scaffold count by 14% to 578 and increased the scaffold N50 by 23% to 76.2 Mb (Table S1). Overall, 98.27% assembly sequence could be assigned to chromosomes (Table S3).

##### 1 - 2 BUSCO genes recovery

The comparison between curated bHirRus1 and Chelidonia (8) revealed a discrepancy in BUSCO (18–20) gene completeness. Chelidonia overall completeness was 95.9%, while curated bHirRus1 was 94.8%, with 139 missing genes (Table S6). A slight decrease in BUSCO scores in highly contiguous assemblies generated with Hi-C scaffolding has been previously observed (e.g. in the yellow perch (75)). The difference is not necessarily due to a higher gene fragmentation, but to BUSCO mode of operation in relation to the identification of the best candidate genomic regions across different scaffolds. In a second step of manual curation, BUSCO results were carefully analysed in order to improve the functional completeness of our assembly. In bHirRus1, we found 13 of the BUSCO genes missing in Chelidonia (Table S34). However, of the 139 missing BUSCO genes in bHirRus1, 53 were identified in Chelidonia, 21 in the alternate assembly (14 in common with those in Chelidonia) and one in the VGP assembly pipeline intermediate p1 (Table S35-36). According to the BLAST search of the gene sequences extracted from Chelidonia, the alternate assembly, and p1, 19 of those genes were indeed present in bHirRus1 (Table S35, S37), implying that BUSCO results were partially inaccurate, and several genes that should have been identified as present were missed possibly because of automated gene prediction issues. In addition, 7 of the 9 missing alternate genes in bHirRus1 (g20-25, g27; Table S35) were found by BUSCO in the primary contigs (c1) at the beginning of the VGP assembly pipeline (Figure S1), and in the alternate haplotig (p2) after haplotig purging with purge_dups (3), but not in the primary purged contigs (p1), showing that while purge_dups removes false duplications and repeats, it can also remove some coding genomic regions. In particular, 6 of the removed regions containing the genes, were flagged as haplotigs and one as repetitive by purge_dups (Table S38). A manual evaluation of the alignments between bHirRus1 and Chelidonia, alternate assembly and p1 contigs or scaffolds containing the missing genes, further confirmed the presence of the genes and the absence of the missing ones in our VGP assembly (Table S39). One gene (g1, Table S39) extracted from Chelidonia partially aligned to bHirRus1 and therefore was not recovered to avoid introducing duplications. The sequences of the missing genes were extracted from the alternate assembly and from Chelidonia together with the entire scaffold fragment that did not contain significant alignments to bHirRus1. In total, 42 genes were recovered. Thirty-two sequences containing 35 of the missing gene sequences were extracted from Chelidonia, and 7 sequences containing one gene each were extracted from the alternate assembly (Table S40). Two of the 9 alternate assembly genes that were missing from bHirRus1 (g27-28; Table S35) were recovered from Chelidonia after a careful evaluation, and some scaffolds were trimmed to avoid introducing duplicated sequences (See Notes column in Table S40). The recovered sequences were renamed accordingly and added to bHirRus1 to generate a more complete assembly. The BLAST search was repeated on the updated assembly and confirmed the presence of the recovered genes (Table S41). BUSCO completeness of the updated assembly was 96%, also slightly higher than that of Chelidonia (95.9%; Table S6). After the gene recovery, the missing genes decreased from 139 to 97 in bHirRus1 (-1.3%; Table S6). *K*-mer completeness increased from 33.2% to 33.3% in the new version with respect to the original, while QV slightly decreased from 44% to 43.7%, due to the introduction of relatively lower quality sequences in the recovered genes (Table S2). Duplicated genes minimally increased as a result of the process (from 0.46% to 0.49%; Table S42).

#### 2 - Barn swallow mitogenome

The mitogenome sequence, included in the VGP assembly pipeline for polishing, was 16,277 bp (Figure S2). Our annotation included 2 rRNA genes, 20 tRNA genes and other 12 genes, in line with closely related birds (15) (Table S43). We could not resolve the CR region due to the presence of repetitive elements, but this is presented for another individual in our companion paper (Lombardo et al., submitted).

#### 3 - Karyotype reconstruction

The karyotype of the barn swallow is composed of 2n = 80 chromosomes, which corroborate previous studies (76,77) (Fig. 1). The karyotype is made of 7 pairs of macrochromosomes (pairs 1-6 and the Z chromosome), 7 pairs of chromosomes of intermediate size (pairs 7-13) and 26 pairs of microchromosomes (pairs 14-39). The morphology of the macrochromosomes is similar to that of other swallow species (78). In particular, pairs 1, 4,

10 and the Z chromosomes are metacentric, pairs 2, 3 and 5 are acrocentric, while pair 6 is submetacentric. Microchromosome morphology is not identifiable due to their small size.

##### 4 - bHirRus1 evaluation

Genomescope v2.0 (7), starting from unassembled 10X linked-reads, predicted a genome size of 1.24 Gbp, a repeat content of 281 Mbp and a heterozygosity of 1.04% (Figure S4a, Table S5). The final assembly size was 1.11 Gbp (Table S1), slightly smaller than expected (1.24 Gbp) (Figure S4a, Table S5). However, likely due to the high heterozygosity and possibly due the bias often introduced during 10x library preparation, the model fit from Genomescope v2.0 was only 88.8%, and therefore the estimations may be inaccurate (Figure S4a, Table S5). The Hi-C contact heatmap confirmed the chromosomal structure of the assembly (2n = 78 + ZW), showing a strong correlation between read pairs and the corresponding scaffold sequence (Figure S3a). The final scaffold N50 and NG50 are 76 Mbp and 73 Mbp, respectively, with a maximum scaffold length of 156 Mbp (Table S1, Figure S5c). The primary assembly also included 1,719 contigs with a NG50 of 2.8 Mbp and a NG50 of 2.3 Mbp, 1,103 gaps with a max gap length of 859 Mbp and a N50 of 67 Mbp (Table S1). According to *k*-mer-based metrics (10), bHirRus1 has a per-base consensus accuracy of Q43.7 and a *k*-mer completeness of 83.3%, which reaches 93.9% when combined to the alternate assembly (74.2%) (Table S2). Duplications in the original assembly after FALCON+unzip (c1) were 4.0% and decreased to 0.50% after purging with purge_dups (3) (p1) during the assembly pipeline, and to 0.49% in the final assembly after curation (Figure S1, Figure S18, Table S42). BUSCO (18–20) completeness was 96% in bHirRus1, comprising 95.2% of complete single-copy *Vertebrata* orthologous genes and a 0.8% of duplicated complete genes, while 1.1% of identified BUSCO genes were fragmented and 2.9% were missing (Figure S5, Table S6). Repeat masking of the new assembly revealed 271 Mbp of repeats (Table S2), close to the Genomescope v2.0 prediction of 280 Mbp (Table S5).

##### 5 - Comparison between bHirRus1 and Chelidonia

Compared to Chelidonia (8), bHirRus1 is 108 Mbp smaller (1.11 Gbp vs. 1.21 Gbp) and slightly more fragmented, containing a larger number of scaffolds (617 vs. 364) and contigs (1,677 vs. 1,355; Figure S5b,c, Table S1). Since the same data was used to generate the contigs, this is likely due to the different assembler employed (Canu (79) in Chelidonia, FALCON-Unzip (2) in bHirRus1). However, the scaffold contiguity is significantly higher in bHirRus1 (scaffold N50 76 Mbp vs. 26 Mbp, scaffold NG50 73 Mbp vs. 26 Mbp; Figure S5c, Table S1). The lower contiguity of Chelidonia was also confirmed by the Hi-C contact heatmap (Figure

S3c), a data type previously unavailable (8), and by the alignment with the Chicken genome GRCg7b (Figure S19). While the scaffold NG50 increased in bHirRus1, the contig NG50 reduced from 5.6 Mbp in Chelidonia to 2.7 Mbp, in line with the increase in gap number (1,103 vs 971) (Table S1). However, the total gap size was lower in bHirRus1 than Chelidonia (25.5 Mbp vs 40 Mbp, Table S1). Merqury (10) computed a QV of 34 for Chelidonia, an order of magnitude less than that of bHirRus1 (Table S2). Conversely, *k*-mer completeness in bHirRus1 was lower than that of Chelidonia (83.2% vs. 84.9%), but only when the alternate assembly was left out (Table S2). In this case, the higher *k*-mer completeness is likely due to the presence of false duplications, i.e. duplicated copies of highly divergent heterozygous regions. Indeed, the haplotig purging in Chelidonia only consisted in discarding small contigs identified as haplotigs by BLAST searches (8). Merqury spectra-cn plot generated for Chelidonia, revealed that the duplication content of Chelidonia was three times higher than that of bHirRus1 (1.3% vs. 0.49%; Figure S4b, S18e, Table S42), and contained more duplicated BUSCO genes (1.2% vs 0.8%; Figure S5c, Table S6). The haplotig purging performed on Chelidonia, reduced its size by 55 Mbp (1.16 Gbp vs. 1.21 Gbp), leading to an assembly size closer to that of bHirRus1 (1.11 Gbp; Table S1). Twenty-seven Mbp were flagged by purge_dups as haplotigs, 16 Mbp as repeats, 10 Mbp as overlaps and 1.7 Mbp as high-coverage regions (Table S9). These results led us to conclude that the larger assembly size and completeness of Chelidonia are largely explained by retained alternate haplotigs. Haplotigs removal was confirmed by the reduction of the duplication content in the purged assembly (from 1.3% to 0.55%; Figure S18e,f, Table S42), which remained higher than that of bHirRus1 (0.49%). BUSCO identified 13 duplicated genes less (28 vs. 42), while minimally affecting the overall completeness (95.8% vs. 95.9%; Table S6). The *k*-mer completeness of Chelidonia after purging decreased from 84.9% to 84.2%, whilst the QV increased from 34 to 34.2 (Table S2). Moreover, Chelidonia has a higher repeat content (315 Mbp) than bHirRus1 (271 Mbp), according to Windowmasker (29) and RepeatMasker (30), also higher than the Genomescope2.0 (7) prediction (280 Mbp; Figure S4a, Table S5).

#### 6 - PhyloP analysis

The rate of accelerated and conserved sites was lower than that detected in other bird species (40,41,80), most likely because of the lower number of species and the shorter total branch length between the aligned species (40). Applying a more conservative Bonferroni correction (81) on the FDR corrected bases, we detected ~64 kbp (~0.01%) of significantly accelerated sites and no significantly conserved ones (Table S12). Bonferroni-corrected accelerated bases, being a subset of FDR-corrected ones, follow the same pattern of overlaps with genomic features (Table S12). The 20 genes with the most overlaps with phyloP accelerated sites (Table S13) comprise genes encoding for mitochondrial (*MRPL55 (82)*, *COQ10A (83)*) and ribosomal proteins (*RPS23 (84)*), genes involved in hair growth and morphogenesis (*HOXC12 (85)*, *HOXC11 (86), SNRPE (87)*), neuron survival and migration (*ZNHIT3 (88)*), oxidative stress response (*BFR2 (89)*), inflammatory response (IL23A (90–92)), neural development (*RBM8A (93)*), cognitive deficits (ZNF653 (94)), photoreception (*GNGT2 (95)*), angiogenesis and vascular remodelling (*ACVRL1 (96–99)*), embryonic stem cell differentiation (*POLR3GL (100)*) and stem cell self-renewal (MINDY1 (101)), metabolic processes (*SCLY (102,103)*), ion transport (*KCNE3 (104,105)*), and pain transmission and emotional processing (*KCNIP3 (106)*). The 20 genes with the most overlaps with phyloP conserved sites (Table S15) were largely involved in neural development and differentiation (*NFIA (107)*, *SOX2 (108)*, *CNOT2 (109)*, *UBE2D2 (110), UBE2D3 (110)*, *GJD2 (111)*, *FOXP2 (112)*, *AMD1 (113)*, *FGF12 (114)*, *HMGN3 (115)*, *POU2F1 (116)*, *UBE2N (110)*, *LMO4 (117)*, *MOSMO (118)*).

#### 7 - SNP catalogue description

##### 7 - 1 - SNP counts

A comprehensive SNP catalogue was generated using all publicly available genomic data for the barn swallow, including individuals from all extant subspecies as well as five high coverage samples sequenced with HiFi technology (Figure S11). Genetic marker counts obtained after variant filtering (Table S33 and Methods) are summarised in Table S44, while average site depth and average individual depth for each dataset are reported in Table S45. Results relative to the HiFi dataset (both full coverage and downsampled dataset) are discussed in more detail in Supplementary Note section 8-2. SNP counts and density for all chromosomes are summarised in Table S25. No SNP was detected on chromosome 39. For each sequencing technology, we also computed the number of SNPs by annotation category (genic, intergenic, exonic, intronic; Tables S46-S49).

##### 7 - 2 - Correlation between SNP density and genomic features

For all datasets, SNP counts across all chromosomes were correlated with GC content by computing Spearman nonparametric rank test (26). We found a positive correlation between chromosome GC content and SNP density (GC content was calculated as the fraction of GC bases every 10 kbp; SNP density was calculated over 10 kbp windows across all genome as described in Supplementary Methods) in all datasets (HiFi WGS rho =

0.25, S = $1.59 \times 10^{14}$, p-value < $2.2 \times 10^{-16}$; downsampled 5x HiFi WGS rho = 0.17, S =$1.75 \times 10^{14}$, p-value < $2.2 \times 10^{-16}$; Illumina WGS rho = 0.41, S = $1.25 \times 10^{14}$, p-value < $2.2 \times 10^{-16}$; Illumina ddRAD rho = 0.016, S = $2.08 \times 10^{14}$, p-value = $2.4 \times 10^{-7}$).

#### 8 - HiFi read mapping, variant calling, titration and phasing experiments

##### 8 - 1 Sequencing and read mapping

HiFi WGS for the five samples from ds1 generated a mean read length across samples of 15609.7 bp (≥ Q20). About 25.2 Gbp of ≥ Q20 data were generated on average for each sample, and median read ≥ Q20 quality averaged across samples was 31 (Table S20). Read alignment had a mean mapped concordance of ~97% across all samples, with an average of 1.64 million reads mapped to bHirRus1 (Table S50). We verified read alignment coverage for each sample using bedtools genomecov and verified that the distribution of bases at a certain coverage value was consistent with the expectation based on sequencing coverage (Figure S20). The secondary peak observed at half average coverage, particularly in samples sequenced at higher coverage (A1, 2, 4), is presumably due to the fact that all five samples belonged to the heterogametic sex. We also compared the proportion of the genome covered by uniquely mapped reads between Illumina WGS reads (96.35%) and HiFi reads (97.25%).

##### 8 - 2 Variant calling and titration experiment

Given the high accuracy of the Hifi reads, we performed a titration experiment to ascertain how sequencing depth affects variant calling. This aimed at identifying the minimum coverage required to detect reliable variants, with the goal of a recall rate > 60%. Recall rates from individual variant calling are reported in Table S51. We found that the steepest decrease in the number of variants detected occurred between 7.5x and 5x (Figure S21). At 5x, for all truth sets considered, ~50% of the variants called at 20x were not recalled (Table S51). These preliminary results suggest sequencing coverage of about 7.5x to recover an adequate number of SNPs for subsequent analysis. Precision remained high even at 5x (average precision rate across all samples of 0.93 for truth set 1). Truth sets 2 and 3 were excluded from further evaluation because they resulted in very low values of precision, even at high coverage (Table S51; this is presumably due to the fact that both these truth sets include only a limited subset of the variants present in our datasets). In addition, we performed joint variant calling to assess whether an increased number of variants can be recovered from the five combined HiFi read sets, by combining reads for variants shared between samples for a joint call, with subsequent genotyping for each sample for the variant. While joint calling for SNVs and indels was comparable to performance with individually called variants for full and downsampled coverage, the recovery of structural variants (including deletions, insertions, inversions, duplications, translocations, copy number variations) is significantly improved by joint calling compared to per-sample variant calling (72,691 structural variants on average per sample with joint calling, vs. 50,502 with per-sample calling, compared to 109,248 for full HiFi read set and joint calling; recall rate 0.665 vs 0.462; Figure S22, Table S52). These data suggest that, thanks to the significantly higher variants recall achieved with a joint call approach, lowering the sequencing coverage to 5x can be considered as an alternative approach when working with HiFi data, particularly when interested in recovering structural variants.

**8 - 3 Haplotype phasing with HiFi reads**

We took advantage of the high accuracy and length of HiFi reads to reconstruct phased haplotype blocks across chromosomes for all 5 samples (Table S53). The percentage of phased blocks across chromosomes for sample A1 is shown in Figure S23. We obtained 73% of average phased sequence for macrochromosomes, 77% for intermediate and 64% for microchromosomes (Figure S24). Phasing in sex chromosomes is lower than 6%, due to all samples being females, which can be regarded the average error of the method. This can be caused by regions collapsed in the assembly or by regions with high repeat content.

58. Bejarano D, Martínez R, Manrique C, Parra LM, Rocha JF, Gómez Y, et al. Linkage disequilibrium levels

and allele frequency distribution in Blanco Orejinegro and Romosinuano Creole cattle using medium density SNP chip data. Genet Mol Biol. 2018;41 (2):426–33.

67. Hall MB. Rasusa: Randomly subsample sequencing reads to a specified coverage. Zenodo; 2019.

77. Malinovskaya LP, Tishakova K, Shnaider EP, Borodin PM, Torgasheva AA. Heterochiasmy and Sexual Dimorphism: The Case of the Barn Swallow (Hirundo rustica, Hirundinidae, Aves). Genes [Internet]. 2020

Sep 24;11 (10). Available from: http://dx.doi.org/10.3390/genes11101119

95. Balcueva EA, Wang Q, Hughes H, Kunsch C, Yu Z, Robishaw JD. Human G Protein γ11 and γ14 Subtypes

Define a New Functional Subclass. Exp Cell Res. 2000 Jun 15;257 (2):310–9.

#### Supplementary Tables

| Supplementary Tables Summary | | |
|---|---|---|
| Table | S1 | Assembly statistics |
| Table | S2 | K-mer completeness and QV |
| Table | S3 | Chromosome sizes, GC content, gene density and CpG island density. |
| Table | S4 | Karyotype chromosomes size estimations |
| Table | S5 | Genomescope2.0: Genome size, heterozygosity and repeat content prediction |
| Table | S6 | BUSCO results |
| Table | S7 | RNAseq and IsoSeq public data |
| Table | S8 | Genomic features from annotation |
| Table | S9 | Chelidonia haplotig purging results |
| Table | S10 | Species used in Cactus alignment |
| Table | S11 | bHirRus1 Cactus alignment coverage per chromosome |
| Table | S12 | Selection analysis |
| Table | S13 | PhyloP accelerated genes |
| Table | S14 | Gene ontology analysis on PhyloP accelerated genes. |
| Table | S15 | PhyloP conserved genes |
| Table | S16 | Gene ontology analysis on PhyloP conserved genes |
| Table | S17 | 38 species with CAMK2N2 transcripts downloaded from NCBI |
| Table | S18 | PhastCons conserved genes |
| Table | S19 | Barn swallow individuals sequenced with Hifi technology |
| Table | S20 | HiFi sequencing statistics |
| Table | S21 | HiFi assembly statistics |
| Table | S22 | Pangenome ortholog analysis |
| Table | S23 | 236 genes found only in bHirRus1 |
| Table | S24 | Geographical origin, type of genomic data, population sizes and sequencing coverage for all genomic data analysed |
| Table | S25 | SNP counts and density across chromosomes for all datasets analysed. |
| Table | S26 | Whole genome LD values in Illumina WGS data from ds2 and ds3.1 |
| Table | S27 | LD values according to chromosome type in Illumina WGS data from ds2 and ds3.1 |
| Table | S28 | ROIs with highest average LD values (>= 0.3) computed over 5 kbps windows using Illumina WGS data from ds3.1 |
| Table | S29 | Genotypes of *H. r. savignii* samples (ds3.1.1) within the *BDNF* gene region |

| | | |
|---|---|---|
| Table | S30 | Genomescope2.0 predictions based on HiFi data |
| Table | S31 | Custom purge_dups cutoffs for HiFi |
| Table | S32 | Haplotig purging on HiFi assemblies |
| Table | S33 | Parameters used to filter vcf files of different publicly avalable datasets analyzed |
| Table | S34 | Missing Chelidonia BUSCOs that were found in bHirRus1 |
| Table | S35 | Summary of missing BUSCO in bHirRus1 that were found in Chelidonia, alternate assembly and pipeline intermediates |
| Table | S36 | Information of BUSCO genes missing from bHirRus1, but found in the other assemblies or intermediates |
| Table | S37 | BLAST search of the missing genes in bHirRus1 |
| Table | S38 | Purged c1 sequences that contained BUSCO genes |
| Table | S39 | Alignment coordinates for BUSCO genes recovery |
| Table | S40 | Recovered sequences from Chelidonia and the alternate assembly |
| Table | S41 | BLAST search of the missing genes in bHirRus1 after gene recovery |
| Table | S42 | False duplication content |
| Table | S43 | Mitogenome annotation |
| Table | S44 | Number of SNPs identified in the different datasets |
| Table | S45 | Summary of the values relative to variant depth in the different datasets after our filtering pipeline |
| Table | S46 | SNP counts from ds1 (HiFi 20x) falling in the different genomic regions |
| Table | S47 | SNP counts from downsampled ds1 (HiFi 5x intersected with HiFi 20x as truth set) falling in the different genomic regions |
| Table | S48 | SNP counts from all Illumina WGS datasets (ds2 and ds3.1) combined falling in the different genomic regions |
| Table | S49 | SNP counts from all ddRAD datasets (ds3.2 through ds6) combined falling in the different genomic regions |
| Table | S50 | HiFi data alignment statistics |
| Table | S51 | SNP recall rate after downsampling for all HiFi samples (ds1). |
| Table | S52 | SVs, indels and SVs number for all HiFi samples (ds1) after individual variant calling (per-sample calling) compared to joint variant calling. |
| Table | S53 | Statistics of phased blocks across chromosomes for all HiFi samples (ds1) |

| Table S1 - Assembly statistics. | | | |
|---|---|---|---|
| **a)** | | | |
| **bHirRus1** | **Scaffolds** | **Contigs** | **Gaps** |
| **Total bp** | 1,105,955,550 | 1,080,421,138 | 25,534,411 |
| **Number** | 617 | 1,719 | 1,102 |
| **Max length (bp)** | 156,035,725 | 18,822,688 | 859,144 |
| **N50 (bp)** | 76,187,387 | 2,794,774 | 67,235 |
| **N90 (bp)** | 12,073,725 | 410,547 | 25,050 |
| **NG50 (bp)** | 73,257,097 | 2,317,408 | - |
| **NG90 (bp)** | - | - | - |
| **b)** | | | |
| **bHirRus1 non-curated** | **Scaffolds** | **Contigs** | **Gaps** |
| **Total bp** | 1,108,599,991 | 1,079,323,631 | 29,276,367 |
| **Number** | 670 | 1,690 | 1,022 |
| **Max length (bp)** | 112,528,062 | 18,822,688 | 859,144 |
| **N50 (bp)** | 61,988,788 | 2,794,774 | 87,847 |
| **N90 (bp)** | 5,759,618 | 416,910 | 20,407 |
| **NG50 (bp)** | 54,964,849 | 2,317,408 | - |
| **NG90 (bp)** | - | - | - |
| **c)** | | | |
| **bHirRus1 after first curation step** | **Scaffolds** | **Contigs** | **Gaps** |
| **Total bp** | 1,104,070,652 | 1,078,538,515 | 25,532,137 |
| **Number** | 578 | 1,677 | 1,099 |
| **Max length (bp)** | 156,035,725 | 18,822,688 | 859,144 |
| **N50 (bp)** | 76,187,387 | 2,794,774 | 67,235 |
| **N90 (bp)** | 12,073,725 | 421,471 | 25,050 |
| **NG50 (bp)** | 73,257,097 | 2,317,408 | - |
| **NG90 (bp)** | - | - | - |
| **d)** | | | |
| **Chelidonia** | **Scaffolds** | **Contigs** | **Gaps** |

| | | | |
|---|---|---|---|
| **Total bp** | 1,213,743,879 | 1,173,760,729 | 39,983,150 |
| **Number** | 364 | 1.335 | 971 |
| **Max length (bp)** | 98,053,015 | 33,289,027 | 1,519,005 |
| **N50 (bp)** | 25,954,216 | 6,124,600 | 190,808 |
| **N90 (bp)** | 2,002,624 | 363.863 | 36,955 |
| **NG50 (bp)** | 25,954,216 | 5,557,227 | - |
| **NG90 (bp)** | 1,397,752 | 139.41 | - |
| **e)** | | | |
| **Chelidonia after haplotig p** | **Scaffolds** | **Contigs** | **Gaps** |
| **Total bp** | 1,158,593,414 | 1,118,668,438 | 39,925,031 |
| **Number** | 301 | 995 | 747 |
| **Max length (bp)** | 97,796,368 | 33,289,027 | 1,721,071 |
| **N50** | 30,885,980 | 6,435,469 | 281,095 |
| **N90** | 2,436,503 | 591,306 | 50,550 |
| **NG50** | 25,929,213 | 5,538,306 | - |
| **NG90** | 734,124 | 34,521 | - |

**Table S2 - K-mer completeness and QV.** Metrics were computed by Merqury.

| | bHirRus1 | Alternate assembly | bHirRus1+alternat | bHirRus1 after first curation step | bHirRus1 after first curation s | Chelidonia | Chelidonia after haplotig purging |
|---|---|---|---|---|---|---|---|
| **QV** | 43.7271 | 41.1616 | 42.3478 | 44.0223 | 42.4594 | 33.9565 | 34.2602 |
| **Completeness** | 83.3386 | 74.181 | 93.8648 | 83.2454 | 93.7927 | 84.8622 | 84.2029 |

**Table S3 - Genome-wide densities.** The table report the counts, bps and % of each feature for the entire genome, the assembled chromosomes combined and the single chromosomes. PacBio coverage computed with mosdepth; GC content computed with bedtools nuc; CpG islands downloaded from UCSC browser; gene density computed from GenomicFeatures file; Repeat densities computed with RepeatMasker and Windowmasker. Total number, total bps and % of conserved and accelerated sites in the whole genomes are not reported since the analysis was performed only on the chromosomes. At the bottom of the tables are also reported the sizes of the unlocalized scaffolds and the total chromosome sizes (assembled chromosomes+unlocalized scaffolds)

| CHR | Type | Size (bp) | % of the genome | PacBio long-reads co | GC (%) | CpG islands ( | CpG islands ( | CpG island (%) | Genes (n) | Genes (bp) | Genes (%) | Repeats (n) | Repeats (bp) | Repeats (%) | phyloP cons. (n) | phyloP cons. (bp) | phyloP cons. (% | phyloP acc. (n) | phyloP acc. (bp) | phyloP acc. site | phastCons CEs | phastCons CEs | phastCons CEs (%) |
|---|---|---|---|---|---|---|---|---|---|---|---|---|---|---|---|---|---|---|---|---|---|---|---|
| **whole genome** | whole genome | 1105955551 | 100 | 57.40 | 42.50 | 29760 | 18392197 | 1.66 | 18136 | 573692145 | 51.9 | 5435749 | 271184266 | 24.5 | - | - | - | - | - | - | - | - | - |
| **assembled chromos** | assembled chromoso | 1082536200 | 97.88 | 53.74 | 48.07 | 27977 | 17328813 | 1.60 | 17328 | 567449897 | 52.4 | 5437882 | 258831469 | 23.9 | 2930332 | 29303320 | 2.71 | 1036417 | 10364170 | 0.96 | 2961683 | 132672359 | 12.26 |
| **1** | macrochromosome | 156035725 | 14.11 | 50.17 | 39.01 | 1817 | 1152644 | 0.74 | 1571 | 73751238 | 47.3 | 792065 | 37089578 | 23.8 | 404956 | 4049560 | 2.60 | 120554 | 1205540 | 0.77 | 418143 | 17878588 | 11.46 |
| **2** | macrochromosome | 119023421 | 10.76 | 50.51 | 39.46 | 1556 | 476947 | 1.84 | 1377 | 56311777 | 47.3 | 606828 | 28789135 | 24.2 | 253770 | 2537700 | 2.13 | 101676 | 1016760 | 0.85 | 299875 | 12177931 | 10.23 |
| **3** | macrochromosome | 116801625 | 10.56 | 51.96 | 39.72 | 1635 | 954856 | 0.80 | 1451 | 58852296 | 50.4 | 588993 | 26328924 | 22.5 | 303143 | 3031430 | 2.60 | 87870 | 878700 | 0.75 | 320348 | 13537507 | 11.59 |
| **4** | macrochromosome | 76187387 | 6.89 | 50.80 | 39.96 | 1143 | 978963 | 0.84 | 1013 | 41464053 | 54.4 | 380754 | 18226983 | 23.9 | 174329 | 1743290 | 2.29 | 60697 | 606970 | 0.80 | 203395 | 8274516 | 10.86 |
| **5** | macrochromosome | 73257097 | 6.62 | 57.45 | 39.29 | 1169 | 702977 | 0.92 | 969 | 36954170 | 50.4 | 377368 | 15908291 | 21.7 | 160990 | 1609900 | 2.20 | 67393 | 673930 | 0.92 | 186908 | 7692850 | 10.50 |
| **6** | macrochromosome | 63258489 | 5.72 | 62.31 | 41.30 | 1506 | 753219 | 1.03 | 1100 | 35603438 | 56.3 | 318822 | 13338924 | 21.1 | 194714 | 1947140 | 3.08 | 53118 | 531180 | 0.84 | 188769 | 8427264 | 13.32 |
| **7** | macrochromosome | 38459648 | 3.48 | 50.91 | 41.96 | 916 | 1057873 | 1.67 | 595 | 23164754 | 60.2 | 198246 | 7337756 | 19.1 | 141070 | 1410700 | 3.67 | 36285 | 362850 | 0.94 | 126387 | 5874037 | 15.27 |
| **8** | macrochromosome | 36085389 | 3.26 | 52.09 | 42.69 | 904 | 513359 | 1.33 | 622 | 24121741 | 66.9 | 183829 | 7139026 | 19.8 | 123520 | 1235200 | 3.42 | 33498 | 334980 | 0.93 | 111320 | 5090875 | 14.11 |
| **9** | intermediate | 31262510 | 2.83 | 50.62 | 43.00 | 985 | 524497 | 1.45 | 606 | 19199400 | 61.4 | 163527 | 6044835 | 19.3 | 124986 | 1249860 | 4.00 | 28539 | 285390 | 0.91 | 108380 | 5149355 | 16.47 |
| **10** | intermediate | 25880253 | 2.34 | 54.44 | 43.98 | 828 | 573090 | 1.83 | 550 | 14992756 | 57.9 | 134649 | 4942530 | 19.1 | 84166 | 841660 | 3.25 | 26544 | 265440 | 1.03 | 84141 | 3760197 | 14.53 |
| **11** | intermediate | 21491857 | 1.94 | 55.62 | 42.90 | 771 | 524657 | 2.44 | 437 | 11117722 | 51.7 | 111564 | 4341902 | 20.2 | 100227 | 1002270 | 4.66 | 21390 | 213900 | 1.00 | 75418 | 3853997 | 17.93 |
| **12** | intermediate | 20890524 | 1.89 | 54.48 | 44.78 | 687 | 404388 | 1.94 | 414 | 14162264 | 67.8 | 108870 | 4068062 | 19.5 | 76561 | 765610 | 3.66 | 21751 | 217510 | 1.04 | 72567 | 3376097 | 16.16 |
| **13** | intermediate | 20272128 | 1.83 | 50.73 | 44.06 | 767 | 453684 | 2.24 | 441 | 12857672 | 63.4 | 105290 | 3620896 | 17.9 | 92210 | 922100 | 4.55 | 18701 | 187010 | 0.92 | 78364 | 3783495 | 18.66 |
| **14** | microchromosome | 18810845 | 1.70 | 49.66 | 45.63 | 832 | 539222 | 2.87 | 390 | 10650232 | 56.6 | 99754 | 3820192 | 20.3 | 66974 | 669740 | 3.56 | 22440 | 224400 | 1.19 | 65350 | 2976387 | 15.82 |
| **15** | microchromosome | 16541138 | 1.50 | 58.10 | 46.11 | 918 | 544360 | 3.29 | 425 | 10921192 | 66.0 | 87604 | 3219983 | 19.5 | 57329 | 573290 | 3.47 | 19344 | 193440 | 1.17 | 59175 | 2674122 | 16.17 |
| **16** | microchromosome | 15277844 | 1.38 | 47.05 | 46.24 | 689 | 391323 | 2.56 | 369 | 8223585 | 53.8 | 79244 | 3087558 | 20.2 | 57475 | 574750 | 3.76 | 17818 | 178180 | 1.17 | 53631 | 2525730 | 16.53 |
| **17** | microchromosome | 13985943 | 1.26 | 47.32 | 46.83 | 812 | 466364 | 3.33 | 388 | 9371747 | 67.0 | 72741 | 2639636 | 18.9 | 56744 | 567440 | 4.06 | 17212 | 172120 | 1.23 | 53091 | 2536552 | 18.14 |
| **18** | microchromosome | 12073725 | 1.09 | 80.86 | 47.38 | 704 | 482386 | 4.00 | 344 | 7056116 | 58.4 | 63892 | 2755203 | 22.8 | 37533 | 375330 | 3.11 | 16938 | 169380 | 1.40 | 38866 | 1755026 | 14.54 |
| **19** | microchromosome | 11382101 | 1.03 | 47.91 | 48.13 | 701 | 415512 | 3.65 | 364 | 8233132 | 72.3 | 60738 | 2294165 | 20.2 | 49924 | 499240 | 4.39 | 14103 | 141030 | 1.24 | 43767 | 2132495 | 18.74 |
| **20** | microchromosome | 11194341 | 1.01 | 45.67 | 49.26 | 716 | 401986 | 3.59 | 322 | 7333529 | 65.5 | 61230 | 2329230 | 20.8 | 52239 | 522390 | 4.67 | 14246 | 142460 | 1.27 | 44100 | 2118521 | 18.92 |
| **21** | microchromosome | 9617204 | 0.87 | 56.14 | 45.93 | 374 | 209080 | 2.17 | 226 | 6191303 | 64.4 | 51494 | 1989546 | 20.7 | 29307 | 293070 | 3.05 | 9877 | 98770 | 1.03 | 32153 | 1426211 | 14.83 |
| **22** | microchromosome | 7507825 | 0.68 | 67.45 | 49.64 | 616 | 329424 | 4.39 | 255 | 5133824 | 68.4 | 40927 | 1653185 | 22.0 | 23998 | 239980 | 3.20 | 13952 | 139520 | 1.86 | 22785 | 1116153 | 14.87 |
| **23** | microchromosome | 7098401 | 0.64 | 54.90 | 50.60 | 570 | 309878 | 4.37 | 202 | 4508772 | 63.5 | 39724 | 1574390 | 22.2 | 26140 | 261400 | 3.68 | 13778 | 137780 | 1.94 | 24385 | 1149720 | 16.20 |
| **24** | microchromosome | 6843954 | 0.62 | 50.83 | 52.67 | 806 | 419332 | 6.13 | 278 | 4309473 | 63.0 | 38812 | 1698044 | 24.8 | 16952 | 169520 | 2.48 | 16422 | 164220 | 2.40 | 18699 | 838995 | 12.26 |
| **25** | microchromosome | 6778862 | 0.61 | 43.99 | 53.47 | 719 | 490341 | 7.23 | 261 | 3749222 | 55.3 | 39871 | 1721968 | 25.4 | 19747 | 197470 | 2.91 | 15240 | 152400 | 2.25 | 20030 | 962553 | 14.20 |
| **26** | microchromosome | 5553549 | 0.50 | 43.46 | 52.64 | 699 | 485267 | 8.74 | 274 | 3692809 | 66.5 | 30872 | 1448967 | 26.1 | 16868 | 168680 | 3.04 | 11891 | 118910 | 2.14 | 16550 | 825479 | 14.86 |
| **27** | microchromosome | 5236451 | 0.47 | 43.33 | 52.73 | 632 | 449798 | 8.59 | 288 | 3619008 | 69.1 | 29723 | 1351663 | 25.8 | 11563 | 115630 | 2.21 | 12973 | 129730 | 2.48 | 13687 | 749182 | 14.31 |
| **28** | microchromosome | 5297670 | 0.48 | 41.78 | 49.47 | 584 | 408389 | 7.71 | 180 | 2821564 | 53.3 | 28949 | 1403408 | 26.5 | 10615 | 106150 | 2.00 | 13624 | 136240 | 2.57 | 10133 | 590469 | 11.15 |
| **29** | microchromosome | 2102120 | 0.19 | 39.33 | 57.38 | 466 | 313738 | 14.92 | 210 | 1292667 | 61.5 | 12025 | 578216 | 27.5 | 4197 | 41970 | 2.00 | 7708 | 77080 | 3.67 | 5288 | 271999 | 12.94 |
| **30** | microchromosome | 1648998 | 0.15 | 47.77 | 58.49 | 374 | 345438 | 20.95 | 128 | 1192418 | 72.3 | 10474 | 443550 | 26.9 | 4509 | 45090 | 2.73 | 6155 | 61550 | 3.73 | 4418 | 244996 | 14.86 |
| **31** | microchromosome | 1590086 | 0.14 | 101.28 | 47.30 | 56 | 27608 | 1.74 | 43 | 799486 | 50.3 | 6078 | 901772 | 56.7 | 571 | 5710 | 0.36 | 1218 | 12180 | 0.77 | 714 | 136484 | 8.58 |
| **32** | microchromosome | 784579 | 0.07 | 48.44 | 51.10 | 45 | 17753 | 2.26 | 20 | 262551 | 33.5 | 2892 | 536670 | 68.4 | 3 | 30 | 0.00 | 809 | 8090 | 1.03 | 147 | 93928 | 11.97 |
| **33** | microchromosome | 437724 | 0.04 | 107.05 | 51.69 | 19 | 9753 | 2.23 | 19 | 49143 | 11.2 | 1870 | 292340 | 66.8 | 9 | 90 | 0.02 | 307 | 3070 | 0.70 | 126 | 37296 | 8.52 |
| **34** | microchromosome | 606149 | 0.05 | 136.62 | 49.27 | 33 | 14768 | 2.44 | 14 | 86962 | 14.4 | 2636 | 403912 | 66.6 | 673 | 6730 | 1.11 | 83 | 830 | 0.14 | 73 | 22231 | 3.67 |
| **35** | microchromosome | 523230 | 0.05 | 40.04 | 55.86 | 74 | 32811 | 6.27 | 33 | 250821 | 47.9 | 2779 | 164279 | 31.4 | 832 | 8320 | 1.59 | 990 | 9900 | 1.89 | 1319 | 71325 | 13.63 |
| **36** | microchromosome | 338027 | 0.03 | 47.78 | 60.55 | 82 | 57414 | 16.99 | 47 | 270902 | 80.1 | 2192 | 86242 | 25.5 | 332 | 3320 | 0.98 | 1510 | 15100 | 4.47 | 432 | 28024 | 8.29 |
| **37** | microchromosome | 276370 | 0.02 | 38.22 | 55.36 | 43 | 17558 | 6.35 | 22 | 134656 | 48.7 | 1408 | 111092 | 40.2 | 64 | 640 | 0.23 | 371 | 3710 | 1.34 | 119 | 33845 | 12.25 |
| **38** | microchromosome | 220485 | 0.02 | 33.25 | 62.27 | 35 | 17761 | 8.06 | 20 | 203909 | 92.5 | 1633 | 73634 | 33.4 | 119 | 1190 | 0.54 | 213 | 2130 | 0.97 | 65 | 9819 | 4.45 |
| **39** | microchromosome | 65965 | 0.01 | 30.76 | 66.34 | 16 | 10090 | 15.30 | 5 | 36088 | 54.7 | 500 | 27806 | 42.2 | 11 | 110 | 0.17 | 167 | 1670 | 2.53 | 13 | 544 | 0.82 |
| **W** | intermediate | 31704074 | 2.87 | 37.08 | 39.34 | 583 | 356740 | 1.13 | 161 | 7866828 | 24.8 | 81834 | 25072792 | 79.1 | 17645 | 176450 | 0.56 | 11739 | 117390 | 0.37 | 17863 | 1944170 | 6.13 |
| **Z** | macrochromosome | 90132487 | 8.15 | 25.03 | 37.22 | 1095 | 693565 | 0.77 | 894 | 36634677 | 40.7 | 415181 | 19975184 | 22.2 | 133317 | 1333170 | 1.48 | 97273 | 972730 | 1.08 | 140689 | 6523394 | 7.24 |
| **Unlocalized chromos** | Unlocalized | 3831338 | 0.35 | - | - | - | - | - | - | - | - | - | - | - | - | - | - | - | - | - | | - | - |
| **Tot chr sizes** | Tot chr sizes | 1086367538 | 98.23 | - | - | - | - | - | - | - | - | - | - | - | - | - | - | - | - | - | | - | - |

**Table S4 - Karyotype chromosomes size estimations.** Chromosome sizes (bps) were estimated from 4 different karyotype images

| chr | size | est_1.1 | est_1.2 | est_2.1 | est_2.2 | est_3.1 | est_3.2 | est_4.1 | est_4.2 | mean | sd |
|---|---|---|---|---|---|---|---|---|---|---|---|
| **1** | 156035725 | 124716232 | 132023269 | 132064072 | 140197050 | 189644645 | 184393699 | 134626955 | 126998453 | 145583046.88 | 26033432.93 |
| **2** | 119023421 | 98396708 | 96623156 | 141281448 | 147400545 | 142691415 | 119245329 | 100567303 | 99170535 | 118172054.88 | 22415250.09 |
| **3** | 116801625 | 104143019 | 97829171 | 95117115 | 86519395 | 121260227 | 122053975 | 106154375 | 100137528 | 104151850.63 | 12339331.91 |
| **Z** | 90132487 | 96339387 | 94565834 | 82569092 | 93180692 | 107339113 | 105079985 | 97343992 | 95839780 | 96532234.38 | 7569650.04 |
| **4** | 76187387 | 68601022 | 74347333 | 81639608 | 85202627 | 92929540 | 96409818 | 69086299 | 75103146 | 80414924.13 | 10486800.34 |
| **5** | 73257097 | 78674801 | 65692395 | 83266204 | 67464989 | 92257907 | 77665161 | 78433901 | 66185319 | 76205084.63 | 9320124.73 |
| **6** | 63258489 | 58598184 | 54909194 | 56698476 | 59177098 | 59836366 | 63621932 | 59416366 | 55548393 | 58475751.13 | 2781945.52 |
| **7** | 38459648 | 46679909 | 49375709 | 44925022 | 48410584 | 45426792 | 52570521 | 47060340 | 50068764 | 48064705.13 | 2560680.64 |
| **8** | 36085389 | 37599319 | 41146425 | 37798984 | 42523857 | 29857125 | 34314324 | 37390407 | 41473268 | 37762963.63 | 4182319.08 |
| **9** | 31262510 | 33059024 | 34548808 | 34778163 | 36792044 | 32421541 | 32055196 | 34704315 | 34919202 | 34159786.63 | 1558305.24 |
| **10** | 25880253 | 24829739 | 26106697 | 30750403 | 30285661 | 28208572 | 35840762 | 25679044 | 33200103 | 29362622.63 | 3882546.1 |
| **11** | 21491857 | 34619750 | 33059024 | 30827860 | 32222085 | 25460984 | 28635975 | 32662885 | 24497163 | 30248215.75 | 3696818.04 |
| **12** | 20890524 | 19438138 | 19509081 | 26257901 | 29278721 | 22224936 | 27597997 | 19447310 | 19877084 | 22953896.00 | 4123263.95 |
| **13** | 20272128 | 23056186 | 24475028 | 22927252 | 22539968 | 18683600 | 19049945 | 23315283 | 23745058 | 22224040.00 | 2153655.8 |
| **14** | 18810845 | 22985244 | 20289444 | 19209320 | 20526087 | 14898034 | 18195140 | 22025958 | 19554753 | 19710497.50 | 2473206.01 |
| **15** | 16541138 | 20005675 | 19083428 | 20371174 | 18279836 | 18256197 | 20820613 | 20199416 | 19124979 | 19517664.75 | 970668.07 |
| **16** | 15277844 | 19438138 | 22488649 | 15568844 | 18047466 | 16973989 | 15508609 | 21381296 | 18587760 | 18499343.88 | 2540320.03 |
| **17** | 13985943 | 22062997 | 21424518 | 19209320 | 18822035 | 16180242 | 17706680 | 21381296 | 22348289 | 19891922.13 | 2251248.72 |
| **18** | 12073725 | 22488649 | 21992054 | 16885612 | 17272896 | 12822078 | 14287459 | 20091972 | 22778064 | 18577348.00 | 3835705.18 |
| **19** | 11382101 | 16529512 | 20147560 | 16033585 | 18899492 | 12394676 | 12150446 | 19124979 | 20091972 | 16921527.75 | 3238700.52 |
| **20** | 11194341 | 19296254 | 16387628 | 15259016 | 15878671 | 10257663 | 11234583 | 19662197 | 19984528 | 15995067.50 | 3713891.16 |
| **21** | 9617204 | 19296254 | 20005675 | 18202379 | 14794274 | 12089388 | 9830260 | 15257005 | 20414303 | 16236192.25 | 3894416.65 |
| **22** | 7507825 | 15536322 | 20502270 | 14019705 | 15104102 | 10562950 | 11967273 | 15471893 | 17620767 | 15098160.25 | 3103760.32 |
| **23** | 7098401 | 19580023 | 18232123 | 17970009 | 13167679 | 10501893 | 12211503 | 19554753 | 15471893 | 15836234.50 | 3525102.19 |
| **24** | 6843954 | 18303065 | 16600454 | 15104102 | 18434750 | 10318720 | 9402857 | 17943098 | 16653773 | 15345102.38 | 3565037.65 |
| **25** | 6778862 | 17593644 | 19225312 | 15104102 | 14794274 | 10807180 | 10562950 | 17405879 | 19339866 | 15604150.88 | 3455911.45 |
| **26** | 5553549 | 16103859 | 19296254 | 10689057 | 12238196 | 7632190 | 9586030 | 15901668 | 18695204 | 13767807.25 | 4339271.42 |
| **28** | 5297670 | 12202043 | 15891033 | 13632420 | 11076342 | 8914397 | 9463915 | 12356025 | 15471893 | 12376008.50 | 2555143.47 |
| **27** | 5236451 | 17522701 | 12627696 | 13167679 | 13322593 | 8120650 | 7815362 | 15579336 | 12785800 | 12617727.13 | 3315417.24 |
| **29** | 2102120 | 12982406 | 16813280 | 12160739 | 12702937 | 7449017 | 8364880 | 16653773 | 16868661 | 12999461.63 | 3702284.27 |
| **30** | 1648998 | 13479001 | 13124291 | 12005825 | 10843971 | 7326902 | 7143729 | 11711363 | 12033694 | 10958597.00 | 2438307.64 |
| **31** | 1590086 | 12485812 | 12343927 | 6971124 | 11541083 | 5861522 | 8059592 | 13645350 | 13323019 | 10528928.63 | 3075418.04 |
| **32** | 784579 | 14968785 | 14755959 | 10998885 | 12315652 | 9097570 | 6594212 | 12033694 | 11496476 | 11532654.13 | 2768828.53 |
| **34** | 606149 | 14046538 | 14046538 | 6816210 | 8907547 | 5983637 | 8059592 | 13645350 | 14504899 | 10751288.88 | 3645789.72 |
| **35** | 523230 | 10570375 | 11918275 | 6119098 | 6816210 | 4945659 | 5312004 | 9669933 | 11496476 | 8356003.75 | 2864893.18 |

| | | | | | | | | | | | |
|---|---|---|---|---|---|---|---|---|---|---|---|
| **33** | 437724 | 11350738 | 9577185 | 9139918 | 9527203 | 6227867 | 5983637 | 12893244 | 9240158 | 9242493.75 | 2320233.91 |
| **36** | 338027 | 9009648 | 12982406 | 8442806 | 8597720 | 4274026 | 5312004 | 8380609 | 10744370 | 8467948.63 | 2762760.84 |
| **37** | 276370 | 10570375 | 6526674 | 6816210 | 7048581 | 3419221 | 3480278 | 6016847 | 6446622 | 6290601.00 | 2250873.38 |
| **38** | 220485 | 7803632 | 6384790 | 9062461 | 5809270 | 2991818 | 2625473 | 7735946 | 9562489 | 6496984.88 | 2590688.63 |
| **39** | 65965 | 3192395 | 4965948 | 5731813 | 8442806 | 1099035 | 2137013 | 2363761 | 4620079 | 4069106.25 | 2367423.91 |

**Table S5 - Genomescope2.0: Genome size, heterozygosity and repeat content prediction.**

| property | min | max | mean |
|---|---|---|---|
| **Homozygous (aa) (%)** | 98.9601 | 98.9682 | 98.96415 |
| **Heterozygous (ab) (%)** | 1.03185 | 1.03988 | 1.035865 |
| **Genome Haploid Length (bp)** | 1,240,987,705 | 1,242,467,779 | 1,241,727,742 |
| **Genome Repeat Length (bp)** | 280,515,586 | 280,850,146 | 280,682,866 |
| **Genome Unique Length (bp)** | 960,472,119 | 961,617,634 | 961,044,877 |
| **Model Fit (%)** | 79.4413 | 98.3039 | 88.8726 |
| **Read Error Rate (%)** | 0.599615 | 0.599615 | 0.599615 |

GenomeScope version 2.0, p = 2, k = 31.

| **Table S6 - BUSCO results.** BUSCO was ran with the vertebrata_odb10 database and chicken as Augustus training species. | | | | | |
|---|---|---|---|---|---|
| | **bHirRus1** | **Alternate assembly** | **bHirRus1 before BUSCO gene recovery** | **Chelidonia** | **Chelidonia after haplotig purging** |
| **Complete BUSCOs (C)** | 3220 | 2926 | 3181 | 3215 | 3213 |
| **Complete and single-copy BUSCOs (** | 3192 | 2908 | 3153 | 3173 | 3185 |
| **Complete and duplicated BUSCOs (D** | 28 | 18 | 28 | 42 | 28 |
| **Fragmented BUSCOs (F)** | 37 | 70 | 34 | 40 | 40 |
| **Missing BUSCOs (M)** | 97 | 358 | 139 | 99 | 101 |
| **Total BUSCO groups searched** | 3354 | 3354 | 3354 | 3354 | 3354 |
| | C:96.0%[S:95.2%,D:0.8%],F:1.1%,M | C:87.2%[S:86.7%,D:0.5%],F:2.1%,M:10.7%,n:3354 | C:94.8%[S:94.0%,D:0.8%],F:1.0%,M:4.2%,n:3354 | C:95.9%[S:94.6%,D:1.3%],F:1.2% | C:95.8%[S:95.0%,D:0.8%],F:1.2%,M:3.0%,n:3354 |

**Table S7 - RNAseq and IsoSeq public data**

| Strategy | Tissue | Instrument | Experiment acc | Run accession | Spots (Mb) | Bases (Gb) | Link |
|---|---|---|---|---|---|---|---|
| **RNAseq** | Ovary | Illumina NovaSeq 6000 | SRX9927592 | SRR13516425 | 56.9 | 17.2 | https://www.ncbi.nlm.nih.gov/sra/SRX9927592[accn] |
| **RNAseq** | Brain | Illumina NovaSeq 6000 | SRX9927591 | SRR13516426 | 48.8 | 14.7 | https://www.ncbi.nlm.nih.gov/sra/SRX9927591%5baccn |
| **RNAseq** | Muscle | Illumina NovaSeq 6000 | SRX9927590 | SRR13516427 | 54 | 16.3 | https://www.ncbi.nlm.nih.gov/sra/12987859 |
| **RNAseq** | Brain | Illumina HiSeq 2500 | SRX7523274 | SRR10853074 | 81 | 8.1 | https://www.ncbi.nlm.nih.gov/sra/SRX7523274[accn |
| **IsoSeq** | Ovary | PacBio SMRT Sequel | SRX5956896 | SRR9184408 | 0.29 | 0.097 | https://www.ncbi.nlm.nih.gov/sra/SRX5956896%5baccn |
| **IsoSeq** | Brain | PacBio SMRT Sequel | SRX5956895 | SRR9184409 | 0.24 | 0.077 | https://www.ncbi.nlm.nih.gov/sra/SRX5956895%5baccn |

**Table S8 - Genomic features from annotation.** The different features were obtained with GenomicFeature R package from the gff annotation file. Overlapping coordinates were merged for the calculations.

| Feature | total bp | % of the assembly | % of genes |
|---|---|---|---|
| **Genes** | 573692145 | 51.9 | - |
| **Introns** | 524783002 | 47.5 | 91.5 |
| **Exons** | 55560589 | 5.0 | 9.7 |
| **CDS** | 28324328 | 2.6 | 4.9 |
| **5'UTRs** | 3952868 | 0.4 | 0.7 |
| **3'UTRs** | 19748013 | 1.8 | 3.4 |
| **Intergenic regions** | 532263405 | 48.1 | 92.8 |

**Table S9 - Chelidonia haplotig purging results.**

| Type | n° | length (bp) |
|---|---|---|
| **HIGHCOV** | 14 | 1661729 |
| **HAPLOTIG** | 268 | 27114574 |
| **OVLP** | 159 | 10465028 |
| **REPEAT** | 153 | 15814511 |
| **tot** | 594 | 55055842 |

**Table S10 - Species used in Cactus alignment.** The scientific name, the common name, the NCBI assembly name and acession number are shown for each species. Genome lenght and N50 were taken from NCBI. The table also report the number of masked bases and percentage of the genome that was masked, and also the bases aligned to the barn swallow and the percentage of the genome that aligned.

| Scientific name | Common name | NCBI name | Accession number | Tot bp | N50 (bp) | Masked bp | % masked | Aligned bp | % aligned |
|---|---|---|---|---|---|---|---|---|---|
| ***Molothrus ater*** | Brown-headed cowb | BPBGC_Mater_1.0 | GCF_012460135.1 | 1,087,312,585 | 52,124,711 | 269,471,518 | 24.78 | 881,341,738 | 79.69 |
| ***Motacilla alba*** | White wagtail | Motacilla_alba_V1.0 | GCF_015832195.1 | 1,072,670,728 | 72,386,170 | 272,868,451 | 25.44 | 880,594,475 | 79.62 |
| ***Camarhynchus parvulus*** | Small tree finch | STF_HiC | GCF_901933205.1 | 1,051,609,828 | 70,356,807 | 261,698,129 | 24.89 | 874,775,347 | 79.09 |
| ***Passer domesticus*** | House sparrow | Passer_domesticus- | GCA_001700915.1 | 1,042,720,703 | 6,373,860 | 228,033,610 | 21.87 | 871,458,883 | 78.79 |
| ***Lonchura striata domestica*** | Bengalese finch | lonStrDom2 | GCF_005870125.1 | 1,060,269,806 | 71,975,342 | 245,415,318 | 23.15 | 863,868,784 | 78.11 |
| ***Taenopygia guttata*** | Zebra finch | bTaeGut2.pat.W.v2 | GCF_008822105.2 | 1,068,988,109 | 70,879,221 | 256,101,562 | 23.96 | 860,556,598 | 77.81 |
| ***Gallus gallus*** | Chicken | bGalGal1.mat.broile | GCF_016699485.2 | 1,053,332,251 | 90,861,225 | 264,817,877 | 25.14 | 662,165,156 | 59.87 |
| | | | | | | | | | **Mean coverage** |
| | | | | | | | | | 76.14 |

**Table S11 - bHirRus1 Cactus alignment coverage per chromosome**

| Chr | 1 | 2 | 3 | Z | 4 | 5 | 6 | 7 | 8 | W |
|---|---|---|---|---|---|---|---|---|---|---|
| Size (bp) | 156035725 | 119023421 | 116801625 | 90132487 | 76187387 | 73257097 | 63258489 | 38459648 | 36085389 | 31704074 |
| 0 genomes | 22532775 | 17120388 | 16367836 | 1759600 | 12969937 | 10014716 | 7934165 | 3870322 | 4473612 | 26864875 |
| 1 genome | 94404 | 86564 | 70252 | 12899 | 56265 | 78449 | 53327 | 36235 | 30249 | 24743 |
| 2 genomes | 109679 | 114723 | 90631 | 16001 | 77688 | 70938 | 58950 | 43369 | 44714 | 95090 |
| 3 genomes | 274008 | 298724 | 191620 | 44761 | 189633 | 166941 | 143002 | 106206 | 115394 | 234938 |
| 4 genomes | 1559855 | 1425231 | 1086574 | 200355 | 875693 | 810028 | 757292 | 491506 | 465077 | 166864 |
| 5 genomes | 5332627 | 4657566 | 3781877 | 681347 | 3194956 | 2720606 | 2322007 | 1440963 | 1451433 | 303379 |
| 6 genomes | 35242269 | 27027733 | 25048151 | 2674231 | 15721209 | 16710628 | 14350140 | 8797534 | 8003464 | 1344969 |
| 7 genomes | 90890108 | 68292492 | 70164684 | 4841346 | 43102006 | 42684791 | 37639606 | 23673513 | 21501446 | 2669216 |
| % aligned | 85.56 | 85.62 | 85.99 | 98.05 | 82.98 | 86.33 | 87.46 | 89.94 | 87.60 | 15.26 |
| % not aligned | 14.44 | 14.38 | 14.01 | 1.95 | 17.02 | 13.67 | 12.54 | 10.06 | 12.40 | 84.74 |

| Chr | 9 | 10 | 11 | 12 | 13 | 14 | 15 | 16 | 17 | 18 |
|---|---|---|---|---|---|---|---|---|---|---|
| Size (bp) | 31262510 | 25880253 | 21491857 | 20890524 | 20272128 | 18810845 | 16541138 | 15277844 | 13985943 | 12073725 |
| 0 genomes | 2994007 | 2829335 | 2491859 | 2788498 | 2023772 | 2452724 | 1579808 | 2336924 | 1721483 | 2267587 |
| 1 genome | 27026 | 26193 | 19301 | 32787 | 16771 | 24623 | 24375 | 16522 | 16317 | 27237 |
| 2 genomes | 38765 | 42310 | 25876 | 30390 | 21061 | 45298 | 21016 | 26656 | 21378 | 34654 |
| 3 genomes | 107071 | 102602 | 63611 | 76214 | 53703 | 73997 | 60815 | 63252 | 53879 | 99275 |
| 4 genomes | 431152 | 435300 | 261302 | 263131 | 219746 | 283376 | 245029 | 223732 | 210481 | 301281 |
| 5 genomes | 1252595 | 1239054 | 795016 | 789694 | 768595 | 818648 | 747715 | 646004 | 650277 | 772153 |
| 6 genomes | 7358268 | 6419206 | 4764344 | 4803906 | 4427105 | 4282907 | 4346476 | 3526897 | 3460215 | 2840829 |
| 7 genomes | 19053626 | 14786253 | 13070548 | 12105904 | 12741375 | 10829272 | 9515904 | 8437857 | 7851913 | 5730709 |
| % aligned | 90.42 | 89.07 | 88.41 | 86.65 | 90.02 | 86.96 | 90.45 | 84.70 | 87.69 | 81.22 |
| % not aligned | 9.58 | 10.93 | 11.59 | 13.35 | 9.98 | 13.04 | 9.55 | 15.30 | 12.31 | 18.78 |

| Chr | 19 | 20 | 21 | 22 | 23 | 24 | 25 | 26 | 27 | 28 | 29 |
|---|---|---|---|---|---|---|---|---|---|---|---|
| Size (bp) | 11382101 | 11194341 | 9617204 | 7507825 | 7098401 | 6843954 | 6778862 | 5553549 | 5236451 | 5297670 | 2102120 |
| 0 genomes | 1328753 | 1345733 | 1179314 | 1531678 | 1076232 | 1590607 | 1353172 | 1252173 | 1363306 | 2172676 | 636352 |
| 1 genome | 17196 | 13281 | 9355 | 17867 | 15562 | 17142 | 19566 | 18536 | 16945 | 28661 | 14911 |
| 2 genomes | 24082 | 17046 | 11652 | 30822 | 17156 | 25826 | 26353 | 20175 | 30470 | 34797 | 16723 |
| 3 genomes | 51016 | 42646 | 23613 | 79264 | 34376 | 64027 | 66240 | 54283 | 96822 | 107405 | 43901 |
| 4 genomes | 205288 | 158865 | 100480 | 203219 | 131589 | 196198 | 211551 | 163204 | 236583 | 214801 | 100617 |
| 5 genomes | 547100 | 464075 | 353390 | 453555 | 392658 | 498113 | 515989 | 463395 | 484255 | 409558 | 238219 |
| 6 genomes | 2798493 | 2592367 | 2300716 | 1961827 | 1913710 | 1893733 | 1911842 | 1571580 | 1337925 | 976611 | 494811 |
| 7 genomes | 6410173 | 6560328 | 5638684 | 3229593 | 3517118 | 2558308 | 2674149 | 2010203 | 1670145 | 1353161 | 556586 |
| % aligned | 88.33 | 87.98 | 87.74 | 79.60 | 84.84 | 76.76 | 80.04 | 77.45 | 73.97 | 58.99 | 69.73 |
| % not aligned | 11.67 | 12.02 | 12.26 | 20.40 | 15.16 | 23.24 | 19.96 | 22.55 | 26.03 | 41.01 | 30.27 |

| Chr | 30 | 31 | 32 | 33 | 34 | 35 | 36 | 37 | 38 | 39 |
|---|---|---|---|---|---|---|---|---|---|---|
| Size (bp) | 1648998 | 1590086 | 784579 | 437724 | 606149 | 523230 | 338027 | 276370 | 220485 | 65965 |
| 0 genomes | 626826 | 1264350 | 716285 | 407203 | 583775 | 154497 | 178933 | 201909 | 192908 | 54934 |
| 1 genome | 11081 | 10791 | 12234 | 729 | 147 | 1361 | 5768 | 665 | 1194 | 129 |
| 2 genomes | 14520 | 4317 | 1494 | 2599 | 498 | 3700 | 12342 | 4820 | 1565 | 1010 |
| 3 genomes | 40326 | 15874 | 12182 | 2293 | 3153 | 4307 | 32148 | 7982 | 5353 | 1580 |
| 4 genomes | 94972 | 23373 | 12594 | 2507 | 1192 | 17733 | 51294 | 20097 | 10409 | 5189 |
| 5 genomes | 174771 | 39040 | 12903 | 6662 | 1490 | 44006 | 30589 | 11709 | 7359 | 2569 |
| 6 genomes | 312494 | 107858 | 15680 | 11940 | 11481 | 112568 | 18316 | 14798 | 508 | 25 |
| 7 genomes | 374008 | 124483 | 1207 | 3791 | 4413 | 185058 | 8637 | 14390 | 1189 | 529 |
| % aligned | 61.99 | 20.49 | 8.70 | 6.97 | 3.69 | 70.47 | 47.07 | 26.94 | 12.51 | 16.72 |
| % not aligned | 38.01 | 79.51 | 91.30 | 93.03 | 96.31 | 29.53 | 52.93 | 73.06 | 87.49 | 83.28 |

**Table S12 - Selection analysis.** a) Total conserved and accelerated elements and bases computed with PhyloP and PhastCons and corrected with a 5% FDR and Bonferroni correction. No significant calls were retrieved in the PhyloP conservation analysis with FDR+Bonferroni. b) Number and percentage of FDR corrected conserved and accelerated bases that fall into different genomic features (extractet with GenomicFeatures R package). c) Base pair number of genomic feature in the chromosomes and percentage of genomic features bases that are coverered by conserved or accelerated elements.

| a) | total (bp) | n° elements | Chr fraction (%) |
|---|---|---|---|
| PhyloP acc. (FDR) | 10364130 | 1036413 | 0.96 |
| PhyloP acc. (FDR + Bonferroni) | 63710 | 6371 | 0.01 |
| PhyloP cons. (FDR) | 29303310 | 2930331 | 2.71 |
| PhastCons | 132672359 | 2961683 | 12.26 |

| b) | Genic sites (bp) | Genic sites (%) | CDS sites (bp) | CDS sites (%) | 5' UTRs sites (b | 5' UTRs sites (% | 3' UTRs sites (b | 3' UTRs sites (% | Intronic sites (b | Intronic sites (% | Intergenic sites | Intergenic sites (%) |
|---|---|---|---|---|---|---|---|---|---|---|---|---|
| PhyloP acc. (FDR) | 5343488 | 51.56 | 93756 | 0.90 | 65325 | 0.63 | 142070 | 1.37 | 5010136 | 48.34 | 5020642 | 48.44 |
| PhyloP acc. (FDR + Bonferroni) | 32488 | 50.99 | 892 | 1.40 | 673 | 1.06 | 1182 | 1.86 | 29421 | 46.18 | 31222 | 49.01 |
| PhyloP cons. (FDR) | 18317624 | 62.51 | 5011640 | 17.10 | 226695 | 0.77 | 1674976 | 5.72 | 11340733 | 38.70 | 10985686 | 37.49 |
| PhastCons | 80706942 | 60.83 | 18053504 | 13.61 | 911705 | 0.69 | 5520325 | 4.16 | 55894466 | 42.13 | 51965405 | 39.17 |

| c) | Genes (bp) | Genes covered | CDS (bp) | CDS sites (%) | 5' UTRs (bp) | 5' UTRs (%) | 3' UTRs (bp) | 3' UTRs (%) | Introns (bp) | Introns (%) | Intergenes (bp) | Intergenes (%) |
|---|---|---|---|---|---|---|---|---|---|---|---|---|
| PhyloP acc. (FDR) | 567449897 | 0.02 | 27338408 | 0.34 | 3857953 | 1.69 | 19510415 | 0.73 | 519872436 | 0.96 | 515086303 | 0.97 |
| PhyloP acc. (FDR + Bonferroni) | 567449897 | 0.000 | 27338408 | 0.003 | 3857953 | 0.017 | 19510415 | 0.006 | 519872436 | 0.006 | 515086303 | 0.006 |
| PhyloP cons. (FDR) | 567449897 | 0.88 | 27338408 | 18.33 | 3857953 | 5.88 | 19510415 | 8.59 | 519872436 | 2.18 | 515086303 | 2.13 |
| PhastCons | 567449897 | 3.18 | 27338408 | 66.04 | 3857953 | 23.63 | 19510415 | 28.29 | 519872436 | 10.75 | 515086303 | 10.09 |

**Table S13 - PhyloP accelerated genes.** The top 5% (606) of the genes with more accelerated bases overlapped with the entire gene sequence are reported. "LOC" genes were discarded.

| Gene | Chr | Start | End | Strand | Gene length | Overlaps | % |
|---|---|---|---|---|---|---|---|
| HOXC12 | 30 | 652603 | 657799 | - | 5196 | 962 | 18.51 |
| MKRN2OS | 12 | 8363413 | 8366570 | - | 3157 | 530 | 16.79 |
| ZNHIT3 | 19 | 1770586 | 1771733 | - | 1147 | 191 | 16.65 |
| BRF2 | 28 | 4660464 | 4663514 | - | 3050 | 500 | 16.39 |
| IL23A | 30 | 1056051 | 1057597 | + | 1546 | 250 | 16.17 |
| RBM8A | 29 | 1268128 | 1271664 | + | 3536 | 540 | 15.27 |
| RPS23 | Z | 81569892 | 81571688 | + | 1796 | 270 | 15.03 |
| MRPL55 | 1 | 1.54E+08 | 1.54E+08 | + | 1532 | 230 | 15.01 |
| SNRPE | 24 | 6580379 | 6582075 | - | 1696 | 250 | 14.74 |
| COQ10A | 30 | 1023387 | 1027464 | + | 4077 | 590 | 14.47 |
| ZNF653 | 36 | 107018 | 110799 | - | 3781 | 540 | 14.28 |
| HOXC11 | 30 | 633020 | 636806 | - | 3786 | 540 | 14.26 |
| GNGT2 | 27 | 4925604 | 4928161 | + | 2557 | 364 | 14.24 |
| ACVRL1 | 30 | 1580311 | 1591724 | - | 11413 | 1610 | 14.11 |
| POLR3GL | 29 | 1276029 | 1279290 | + | 3261 | 460 | 14.11 |
| SCLY | 10 | 6502205 | 6533568 | - | 31363 | 4369 | 13.93 |
| KCNE3 | 2 | 211817 | 214342 | - | 2525 | 350 | 13.86 |
| KCNIP3 | 28 | 1230952 | 1234593 | + | 3641 | 500 | 13.73 |
| MINDY1 | 29 | 1498912 | 1505272 | - | 6360 | 870 | 13.68 |
| ARMC6 | 26 | 142709 | 150904 | - | 8195 | 1120 | 13.67 |
| GRHPR | Z | 82290592 | 82295341 | - | 4749 | 634 | 13.35 |
| BTG2 | 24 | 6723302 | 6725181 | - | 1879 | 250 | 13.30 |
| RPS11 | 2 | 1.19E+08 | 1.19E+08 | + | 2492 | 330 | 13.24 |
| OST4 | 3 | 1.16E+08 | 1.16E+08 | + | 605 | 80 | 13.22 |
| C5H2orf81 | 5 | 1227940 | 1231138 | - | 3198 | 420 | 13.13 |
| ILF2 | 29 | 1672736 | 1676346 | + | 3610 | 470 | 13.02 |

| | | | | | | | |
|---|---|---|---|---|---|---|---|
| **DBNL** | 28 | 1299954 | 1311296 | + | 11342 | 1430 | 12.61 |
| **FRMPD1** | Z | 82208908 | 82243416 | + | 34508 | 4340 | 12.58 |
| **AK6** | Z | 82175574 | 82178446 | - | 2872 | 360 | 12.53 |
| **NKX3-1** | 28 | 3304503 | 3308635 | + | 4132 | 510 | 12.34 |
| **EXOSC5** | Z | 82197668 | 82199752 | + | 2084 | 250 | 12.00 |
| **GUCA1B** | 24 | 4956023 | 4960086 | - | 4063 | 472 | 11.62 |
| **B9D1** | 15 | 14410476 | 14413494 | - | 3018 | 350 | 11.60 |
| **HSPA8** | 23 | 3864728 | 3870026 | + | 5298 | 610 | 11.51 |
| **TXNL1** | Z | 161294 | 164744 | - | 3450 | 390 | 11.30 |
| **DUSP11** | 28 | 516154 | 522181 | - | 6027 | 679 | 11.27 |
| **PRUNE1** | 29 | 1505351 | 1509438 | + | 4087 | 460 | 11.26 |
| **IQCC** | 25 | 3278811 | 3281138 | - | 2327 | 260 | 11.17 |
| **POLR3D** | 28 | 3794854 | 3798864 | - | 4010 | 440 | 10.97 |
| **AQP8** | 15 | 15645643 | 15650215 | + | 4572 | 500 | 10.94 |
| **S100A1** | 29 | 1690770 | 1692999 | - | 2229 | 242 | 10.86 |
| **TRMT10B** | Z | 82244435 | 82248076 | - | 3641 | 390 | 10.71 |
| **IL23R** | 9 | 1193232 | 1203961 | - | 10729 | 1140 | 10.63 |
| **PERM1** | 22 | 6283562 | 6290728 | + | 7166 | 750 | 10.47 |
| **DDX49** | 26 | 194888 | 204377 | - | 9489 | 990 | 10.43 |
| **ABI3** | 27 | 4917653 | 4923809 | - | 6156 | 640 | 10.40 |
| **HOXB3** | 27 | 5078229 | 5138619 | + | 60390 | 6180 | 10.23 |
| **SLC5A11** | 15 | 15566538 | 15578019 | + | 11481 | 1170 | 10.19 |
| **LSM10** | 25 | 1933480 | 1934756 | + | 1276 | 130 | 10.19 |
| **HOXB13** | 27 | 5019965 | 5022332 | + | 2367 | 240 | 10.14 |
| **HOXB7** | 27 | 5069606 | 5075035 | + | 5429 | 550 | 10.13 |
| **HMGCL** | 25 | 339287 | 343054 | - | 3767 | 380 | 10.09 |
| **FOSB** | 39 | 43138 | 49838 | - | 6700 | 670 | 10.00 |
| **TESC** | 17 | 12047447 | 12054193 | - | 6746 | 670 | 9.93 |
| **PHB** | 27 | 4878788 | 4884743 | + | 5955 | 590 | 9.91 |

| | | | | | | | |
|---|---|---|---|---|---|---|---|
| **RNF222** | 18 | 959835 | 960492 | + | 657 | 65 | 9.89 |
| **RPL38** | 18 | 10933936 | 10935458 | + | 1522 | 150 | 9.86 |
| **RAB29** | 24 | 5996945 | 6006735 | + | 9790 | 960 | 9.81 |
| **FIGLA** | 28 | 1156202 | 1159137 | + | 2935 | 286 | 9.74 |
| **DNM2** | 36 | 47601 | 6.70E+04 | + | 19410 | 1890 | 9.74 |
| **HOXB9** | 27 | 5053405 | 5058890 | + | 5485 | 530 | 9.66 |
| **FAM167B** | 25 | 3259096 | 3260145 | - | 1049 | 100 | 9.53 |
| **MPV17L2** | 26 | 491749 | 494804 | - | 3055 | 290 | 9.49 |
| **MRPL10** | 27 | 4162147 | 4165414 | - | 3267 | 310 | 9.49 |
| **QTRT1** | 36 | 44332 | 46757 | - | 2425 | 230 | 9.48 |
| **TOMM5** | Z | 82205769 | 82206719 | - | 950 | 90 | 9.47 |
| **SUGP2** | 26 | 151246 | 166247 | + | 15001 | 1420 | 9.47 |
| **CAV3** | 12 | 20293907 | 20298945 | + | 5038 | 475 | 9.43 |
| **ESPL1** | 30 | 20449 | 34326 | - | 13877 | 1300 | 9.37 |
| **SLC30A2** | 25 | 889304 | 892209 | - | 2905 | 270 | 9.29 |
| **OXTR** | 12 | 20300659 | 20306507 | - | 5848 | 540 | 9.23 |
| **CHMP1A** | 11 | 1091301 | 1095216 | + | 3915 | 360 | 9.20 |
| **PA2G4** | 30 | 974003 | 983647 | + | 9644 | 880 | 9.12 |
| **DNAJC14** | 30 | 535371 | 538769 | + | 3398 | 310 | 9.12 |
| **SLC25A37** | 28 | 3321922 | 3325112 | - | 3190 | 290 | 9.09 |
| **GALNT5** | 7 | 521717 | 534301 | - | 12584 | 1144 | 9.09 |
| **NDUFA13** | 26 | 21720 | 23597 | - | 1877 | 170 | 9.06 |
| **TOMM6** | 3 | 2148 | 3821 | + | 1673 | 150 | 8.97 |
| **RPL19** | 27 | 3759726 | 3762299 | - | 2573 | 230 | 8.94 |
| **FARSA** | 36 | 152505 | 156316 | + | 3811 | 340 | 8.92 |
| **SYCE2** | 36 | 156423 | 158369 | + | 1946 | 173 | 8.89 |
| **NR0B2** | 25 | 3947181 | 3950338 | + | 3157 | 280 | 8.87 |
| **PRIM1** | 30 | 1145674 | 1156405 | - | 10731 | 950 | 8.85 |
| **CCND3** | 24 | 6780399 | 6792056 | + | 11657 | 1030 | 8.84 |

| | | | | | | | |
|---|---|---|---|---|---|---|---|
| **NUDT17** | 29 | 1534235 | 1.54E+06 | + | 2044 | 180 | 8.81 |
| **TAF11** | 24 | 3456194 | 3458486 | + | 2292 | 200 | 8.73 |
| **LRRC46** | 27 | 4163705 | 4168524 | + | 4819 | 420 | 8.72 |
| **PGC** | 24 | 4175332 | 4180764 | - | 5432 | 470 | 8.65 |
| **PEX11G** | 26 | 2997673 | 3000340 | - | 2667 | 230 | 8.62 |
| **RBBP5** | 24 | 6221000 | 6239206 | + | 18206 | 1570 | 8.62 |
| **CS** | 30 | 1028384 | 1034778 | - | 6394 | 550 | 8.60 |
| **CD164L2** | 25 | 3807011 | 3814338 | + | 7327 | 630 | 8.60 |
| **SYF2** | 25 | 687257 | 690062 | + | 2805 | 240 | 8.56 |
| **RSRP1** | 25 | 690542 | 695375 | - | 4833 | 410 | 8.48 |
| **HOXB4** | 27 | 5109351 | 5111945 | + | 2594 | 220 | 8.48 |
| **MSRB1** | 15 | 15106599 | 15108983 | - | 2384 | 200 | 8.39 |
| **RAD23A** | 36 | 144083 | 148263 | - | 4180 | 350 | 8.37 |
| **MKRN2** | 12 | 8366618 | 8376063 | + | 9445 | 780 | 8.26 |
| **ACVR1C** | 7 | 482966 | 4.98E+05 | + | 15413 | 1270 | 8.24 |
| **S100A13** | 29 | 1693116 | 1696272 | + | 3156 | 260 | 8.24 |
| **HSBP1** | 11 | 3458861 | 3462507 | + | 3646 | 300 | 8.23 |
| **TSPAN10** | 18 | 10972173 | 10974485 | - | 2312 | 190 | 8.22 |
| **CALML4** | 13 | 19241987 | 19249655 | - | 7668 | 630 | 8.22 |
| **PTX4** | 15 | 2689459 | 2692049 | + | 2590 | 210 | 8.11 |
| **CDC37** | 36 | 189267 | 193601 | + | 4334 | 350 | 8.08 |
| **TBKBP1** | 27 | 4116214 | 4127017 | + | 10803 | 870 | 8.05 |
| **TCF7L1** | 28 | 1280639 | 1295739 | - | 15100 | 1214 | 8.04 |
| **PTGER2** | 6 | 2504246 | 2507856 | + | 3610 | 290 | 8.03 |
| **INO80B** | 5 | 1270247 | 1272501 | + | 2254 | 180 | 7.99 |
| **WDR7** | Z | 165573 | 204202 | + | 38629 | 3080 | 7.97 |
| **ZPR1** | 23 | 6276574 | 6280117 | + | 3543 | 281 | 7.93 |
| **TAF8** | 24 | 6754781 | 6761464 | - | 6683 | 530 | 7.93 |
| **XPO7** | 28 | 3891207 | 3920871 | - | 29664 | 2350 | 7.92 |

| | | | | | | | |
|---|---|---|---|---|---|---|---|
| **CCDC103** | 27 | 1418947 | 1422617 | - | 3670 | 290 | 7.90 |
| **FDPS** | 29 | 852347 | 855150 | + | 2803 | 220 | 7.85 |
| **CCDC28B** | 25 | 3281171 | 3283604 | - | 2433 | 190 | 7.81 |
| **ACTG2** | 28 | 491364 | 499200 | - | 7836 | 610 | 7.78 |
| **TNFRSF18** | 22 | 6026616 | 6031929 | + | 5313 | 410 | 7.72 |
| **RCC1** | 25 | 3566346 | 3576222 | - | 9876 | 760 | 7.70 |
| **PCNA** | 28 | 650305 | 654727 | - | 4422 | 340 | 7.69 |
| **EPS8L3** | 24 | 1943341 | 1947634 | - | 4293 | 330 | 7.69 |
| **KXD1** | 26 | 369556 | 371901 | - | 2345 | 180 | 7.68 |
| **ANKRD33** | 30 | 1594933 | 1599508 | - | 4575 | 350 | 7.65 |
| **DNAJB8** | 12 | 9175223 | 9175877 | + | 654 | 50 | 7.65 |
| **COPS9** | 10 | 1751730 | 1753575 | - | 1845 | 141 | 7.64 |
| **TNFRSF13B** | 15 | 13741483 | 1.37E+07 | - | 3012 | 230 | 7.64 |
| **GDF1** | 26 | 215581 | 220185 | + | 4604 | 350 | 7.60 |
| **SNRPG** | 28 | 184713 | 186824 | - | 2111 | 160 | 7.58 |
| **STMN1** | 25 | 858955 | 864519 | - | 5564 | 420 | 7.55 |
| **PPP1R15B** | 24 | 6447243 | 6452951 | + | 5708 | 430 | 7.53 |
| **S100A14** | 29 | 1697169 | 1698901 | + | 1732 | 130 | 7.51 |
| **CNPY2** | 30 | 1036339 | 1039138 | - | 2799 | 210 | 7.50 |
| **FRMD5** | 13 | 19747629 | 19789895 | + | 42266 | 3170 | 7.50 |
| **SNAPC5** | 13 | 18517543 | 18520879 | - | 3336 | 250 | 7.49 |
| **GC** | 5 | 62236695 | 62241248 | + | 4553 | 340 | 7.47 |
| **SPACA9** | 20 | 3953582 | 3957206 | - | 3624 | 270 | 7.45 |
| **SP2** | 27 | 4978820 | 4.99E+06 | + | 11870 | 884 | 7.45 |
| **CIAO2B** | 11 | 21243687 | 21245973 | + | 2286 | 170 | 7.44 |
| **YJEFN3** | 26 | 15001 | 20770 | - | 5769 | 428 | 7.42 |
| **MRPS23** | 19 | 1082646 | 1084805 | - | 2159 | 160 | 7.41 |
| **IL17C** | 11 | 1523543 | 1524220 | - | 677 | 50 | 7.39 |
| **HOXB2** | 27 | 5144571 | 5148101 | + | 3530 | 260 | 7.37 |

| | | | | | | | |
|---|---|---|---|---|---|---|---|
| **RPS2** | 15 | 15129650 | 15132376 | - | 2726 | 200 | 7.34 |
| **RPS27** | 29 | 1576055 | 1577430 | - | 1375 | 100 | 7.27 |
| **ATP5PB** | 24 | 4912542 | 4915436 | - | 2894 | 210 | 7.26 |
| **ACKR3** | 7 | 36565536 | 36573675 | + | 8139 | 590 | 7.25 |
| **TMEM50A** | 25 | 695468 | 698508 | + | 3040 | 220 | 7.24 |
| **INTS3** | 29 | 1630750 | 1657442 | - | 26692 | 1920 | 7.19 |
| **UPF1** | 26 | 221946 | 243657 | - | 21711 | 1560 | 7.19 |
| **PEX11B** | 29 | 1258193 | 1260977 | - | 2784 | 200 | 7.18 |
| **GNRH1** | 28 | 2982442 | 2986065 | + | 3623 | 260 | 7.18 |
| **TRNP1** | 25 | 3922719 | 3927181 | - | 4462 | 320 | 7.17 |
| **SNF8** | 27 | 4947283 | 4950358 | - | 3075 | 220 | 7.15 |
| **SLC35B1** | 27 | 4819934 | 4825106 | + | 5172 | 370 | 7.15 |
| **PIRT** | 18 | 980067 | 987756 | - | 7689 | 550 | 7.15 |
| **TMEM222** | 25 | 3824547 | 3831134 | - | 6587 | 470 | 7.14 |
| **MRPL45** | 27 | 4020014 | 4022958 | - | 2944 | 210 | 7.13 |
| **SH3RF2** | 14 | 426391 | 445915 | - | 19524 | 1390 | 7.12 |
| **RBM7** | 23 | 6687024 | 6689133 | - | 2109 | 150 | 7.11 |
| **NUCKS1** | 24 | 6009371 | 6025542 | + | 16171 | 1150 | 7.11 |
| **TTC12** | 23 | 5801374 | 5814031 | + | 12657 | 900 | 7.11 |
| **PGAP3** | 27 | 3614230 | 3619012 | + | 4782 | 340 | 7.11 |
| **EMC4** | Z | 82200636 | 82202212 | + | 1576 | 112 | 7.11 |
| **TMEM209** | 4 | 789559 | 801336 | - | 11777 | 834 | 7.08 |
| **CLPS** | 24 | 3178375 | 3179932 | + | 1557 | 110 | 7.06 |
| **ATP6V1B2** | 28 | 4105355 | 4113707 | - | 8352 | 590 | 7.06 |
| **C28H2orf68** | 28 | 1192994 | 1194556 | + | 1562 | 110 | 7.04 |
| **GNG13** | 15 | 14921519 | 14923792 | + | 2273 | 160 | 7.04 |
| **C22H1orf159** | 22 | 6063043 | 6074977 | - | 11934 | 840 | 7.04 |
| **MAST1** | 36 | 162978 | 1.74E+05 | - | 10540 | 740 | 7.02 |
| **PSMB3** | 27 | 3895410 | 3899265 | - | 3855 | 270 | 7.00 |

| | | | | | | | |
|---|---|---|---|---|---|---|---|
| **C28H8orf58** | 28 | 3710573 | 3714822 | - | 4249 | 297 | 6.99 |
| **NDUFS5** | 25 | 20216 | 21504 | + | 1288 | 90 | 6.99 |
| **PHOSPHO1** | 27 | 4915810 | 4917815 | + | 2005 | 140 | 6.98 |
| **C8H10orf143** | 8 | 34949227 | 34954389 | - | 5162 | 360 | 6.97 |
| **CTSS** | 29 | 1571351 | 1575427 | - | 4076 | 283 | 6.94 |
| **GBX2** | 7 | 36401821 | 36403983 | - | 2162 | 150 | 6.94 |
| **S100A16** | 29 | 1703170 | 1704613 | + | 1443 | 100 | 6.93 |
| **TRIM63** | 25 | 893136 | 8.97E+05 | - | 3465 | 240 | 6.93 |
| **YKT6** | 28 | 1346355 | 1354591 | + | 8236 | 570 | 6.92 |
| **PRKCB** | 15 | 15665340 | 15758417 | + | 93077 | 6440 | 6.92 |
| **BYSL** | 24 | 6792934 | 6795830 | - | 2896 | 200 | 6.91 |
| **CELF3** | 29 | 1240569 | 1252602 | - | 12033 | 830 | 6.90 |
| **ZBTB5** | Z | 82281543 | 82288217 | + | 6674 | 460 | 6.89 |
| **LSM4** | 26 | 456717 | 4.64E+05 | + | 6834 | 470 | 6.88 |
| **PTP4A2** | 25 | 3157901 | 3177216 | - | 19315 | 1320 | 6.83 |
| **PPDPF** | 16 | 6189005 | 6191642 | + | 2637 | 180 | 6.83 |
| **APOF** | 30 | 1074837 | 1075716 | - | 879 | 60 | 6.83 |
| **KHDRBS1** | 25 | 3180227 | 3198251 | + | 18024 | 1230 | 6.82 |
| **TMEM186** | 15 | 5192399 | 5194744 | - | 2345 | 160 | 6.82 |
| **RIPOR3** | 16 | 925320 | 966369 | - | 41049 | 2800 | 6.82 |
| **FEZ1** | 23 | 1968484 | 1982895 | - | 14411 | 980 | 6.80 |
| **HOXC10** | 30 | 614758 | 620811 | - | 6053 | 410 | 6.77 |
| **CILP2** | 26 | 6443 | 12987 | - | 6544 | 440 | 6.72 |
| **EGR4** | 5 | 1015997 | 1017931 | - | 1934 | 130 | 6.72 |
| **DMTN** | 28 | 3884001 | 3891292 | - | 7291 | 490 | 6.72 |
| **GDF15** | 26 | 428912 | 430858 | - | 1946 | 130 | 6.68 |
| **MARCKSL1** | 25 | 3230894 | 3232990 | + | 2096 | 140 | 6.68 |
| **R3HDM2** | 30 | 1291192 | 1327485 | - | 36293 | 2420 | 6.67 |
| **CLN6** | 13 | 19249614 | 19258419 | - | 8805 | 583 | 6.62 |

| | | | | | | | |
|---|---|---|---|---|---|---|---|
| **CPLANE2** | 22 | 167287 | 170308 | + | 3021 | 200 | 6.62 |
| **ATF7** | 30 | 715521 | 750903 | + | 35382 | 2330 | 6.59 |
| **CARD8** | 10 | 24187131 | 24198077 | + | 10946 | 720 | 6.58 |
| **VPS33B** | 13 | 19970272 | 2.00E+07 | + | 7457 | 490 | 6.57 |
| **RIMBP2** | 17 | 473682 | 558016 | + | 84334 | 5540 | 6.57 |
| **TUBAL3** | 4 | 1089071 | 1098827 | + | 9756 | 640 | 6.56 |
| **INKA2** | 24 | 4831868 | 4844521 | + | 12653 | 830 | 6.56 |
| **ZC3H11A** | 24 | 6582814 | 6595641 | - | 12827 | 840 | 6.55 |
| **ATCAY** | 26 | 582373 | 594925 | + | 12552 | 820 | 6.53 |
| **HOXB6** | 27 | 5085748 | 5089271 | + | 3523 | 230 | 6.53 |
| **TMEM54** | 25 | 3443036 | 3444876 | - | 1840 | 120 | 6.52 |
| **KCTD16** | 14 | 613793 | 674225 | - | 60432 | 3930 | 6.50 |
| **HMGN2** | 25 | 4122604 | 4126143 | - | 3539 | 230 | 6.50 |
| **MTHFD2** | 28 | 394651 | 405480 | - | 10829 | 703 | 6.49 |
| **LSM1** | 28 | 4723501 | 4725507 | - | 2006 | 130 | 6.48 |
| **PGPEP1** | 26 | 442220 | 456579 | - | 14359 | 930 | 6.48 |
| **MRPL1** | 5 | 61968757 | 61973962 | + | 5205 | 337 | 6.47 |
| **AADAC** | 10 | 25402893 | 25410003 | - | 7110 | 460 | 6.47 |
| **CARM1** | 36 | 70683 | 77661 | + | 6978 | 450 | 6.45 |
| **KCNE1** | 2 | 1.18E+08 | 1.18E+08 | - | 4816 | 310 | 6.44 |
| **IL34** | 11 | 7672735 | 7678177 | - | 5442 | 350 | 6.43 |
| **SAMD11** | 22 | 6373174 | 6455166 | - | 81992 | 5250 | 6.40 |
| **DDX20** | 24 | 4819946 | 4824930 | - | 4984 | 318 | 6.38 |
| **PPP6C** | 20 | 827262 | 830554 | + | 3292 | 210 | 6.38 |
| **EIF3I** | 25 | 3267290 | 3274212 | - | 6922 | 440 | 6.36 |
| **CDK5RAP3** | 27 | 4998752 | 5004277 | + | 5525 | 350 | 6.33 |
| **RPL12** | 20 | 9724399 | 9727242 | - | 2843 | 180 | 6.33 |
| **FGF5** | 5 | 60459220 | 60470643 | - | 11423 | 723 | 6.33 |
| **POLR1E** | Z | 82272975 | 82281349 | - | 8374 | 530 | 6.33 |

| | | | | | | | |
|---|---|---|---|---|---|---|---|
| **TRIM41** | 36 | 330010 | 334615 | + | 4605 | 290 | 6.30 |
| **BTG4** | 23 | 5371642 | 5375140 | - | 3498 | 220 | 6.29 |
| **CDC42SE1** | 29 | 1509439 | 1514370 | - | 4931 | 310 | 6.29 |
| **CELA1** | 35 | 506786 | 508695 | + | 1909 | 120 | 6.29 |
| **REN** | 24 | 6539161 | 6545225 | + | 6064 | 380 | 6.27 |
| **HOXB1** | 27 | 5155679 | 5157444 | + | 1765 | 110 | 6.23 |
| **ARID5A** | 28 | 264046 | 269829 | - | 5783 | 360 | 6.23 |
| **EIF4EBP1** | 28 | 4696568 | 4701872 | + | 5304 | 330 | 6.22 |
| **DAD1** | 27 | 2235854 | 2237950 | - | 2096 | 130 | 6.20 |
| **TFEB** | 24 | 4147384 | 4164737 | - | 17353 | 1070 | 6.17 |
| **NPR1** | 29 | 1658418 | 1671592 | - | 13174 | 810 | 6.15 |
| **NDUFB10** | 15 | 15127799 | 15129430 | + | 1631 | 100 | 6.13 |
| **DCTN6** | 5 | 62229419 | 62231543 | - | 2124 | 130 | 6.12 |
| **LDLRAP1** | 25 | 739949 | 754981 | + | 15032 | 920 | 6.12 |
| **RAN** | 17 | 435915 | 4.41E+05 | - | 5230 | 320 | 6.12 |
| **MSTO1** | 29 | 954486 | 958423 | + | 3937 | 240 | 6.10 |
| **STX2** | 17 | 444707 | 461934 | + | 17227 | 1050 | 6.10 |
| **ADD2** | 28 | 1159269 | 1171101 | + | 11832 | 720 | 6.09 |
| **ENDOU** | 30 | 467106 | 476319 | + | 9213 | 560 | 6.08 |
| **GLT1D1** | 17 | 929321 | 969806 | - | 40485 | 2450 | 6.05 |
| **GRB7** | 27 | 3601367 | 3607654 | - | 6287 | 380 | 6.04 |
| **HS3ST2** | 15 | 16074969 | 16085262 | - | 10293 | 620 | 6.02 |
| **CAMK1G** | 24 | 5034270 | 5047926 | - | 13656 | 820 | 6.00 |
| **ATP5MC1** | 27 | 4936413 | 4939255 | + | 2842 | 170 | 5.98 |
| **SLC25A42** | 26 | 121863 | 141605 | - | 19742 | 1180 | 5.98 |
| **UBC** | 19 | 3402724 | 3405570 | - | 2846 | 170 | 5.97 |
| **TBX21** | 27 | 4143962 | 4152678 | + | 8716 | 520 | 5.97 |
| **GPAT2** | 28 | 1237178 | 1245231 | - | 8053 | 480 | 5.96 |
| **GRM7** | 12 | 19949205 | 20132481 | + | 183276 | 10910 | 5.95 |

| | | | | | | | |
|---|---|---|---|---|---|---|---|
| **RASGRF1** | 13 | 19822877 | 19852469 | - | 29592 | 1760 | 5.95 |
| **RASSF2** | 28 | 559111 | 576280 | - | 17169 | 1020 | 5.94 |
| **ANKRD39** | 28 | 112771 | 114960 | + | 2189 | 130 | 5.94 |
| **MDM4** | 24 | 6393488 | 6411198 | - | 17710 | 1050 | 5.93 |
| **FCRL6** | 29 | 1203958 | 1208177 | + | 4219 | 250 | 5.93 |
| **HDAC1** | 25 | 3234735 | 3247224 | - | 12489 | 740 | 5.93 |
| **POLD2** | 28 | 1327005 | 1333419 | - | 6414 | 380 | 5.92 |
| **GLCE** | 13 | 19321852 | 19328610 | + | 6758 | 400 | 5.92 |
| **BSDC1** | 25 | 3313040 | 3319466 | - | 6426 | 380 | 5.91 |
| **KPNB1** | 27 | 4076714 | 4107579 | + | 30865 | 1820 | 5.90 |
| **LZTS1** | 28 | 4078840 | 4098187 | + | 19347 | 1140 | 5.89 |
| **CCDC125** | Z | 82164714 | 82175140 | - | 10426 | 614 | 5.89 |
| **DPM3** | 29 | 763034 | 763646 | - | 612 | 36 | 5.88 |
| **MMADHC** | 7 | 1764336 | 1775557 | + | 11221 | 660 | 5.88 |
| **CTSK** | 29 | 1449428 | 1453342 | - | 3914 | 230 | 5.88 |
| **RCAN1** | 2 | 1.18E+08 | 1.18E+08 | - | 14498 | 850 | 5.86 |
| **SKAP1** | 27 | 5170080 | 5236450 | + | 66370 | 3890 | 5.86 |
| **CDCA8** | 25 | 1430642 | 1434712 | - | 4070 | 238 | 5.85 |
| **DGUOK** | 28 | 486825 | 490418 | - | 3593 | 210 | 5.84 |
| **FZD3** | 3 | 1.16E+08 | 1.16E+08 | - | 45353 | 2650 | 5.84 |
| **C23H11orf52** | 23 | 5474954 | 5481458 | + | 6504 | 380 | 5.84 |
| **CD72** | Z | 20853257 | 20855486 | - | 2229 | 130 | 5.83 |
| **ACBD4** | 27 | 1332567 | 1339289 | - | 6722 | 390 | 5.80 |
| **CENPX** | 18 | 11898022 | 11899401 | - | 1379 | 80 | 5.80 |
| **ZNF593** | 25 | 914928 | 9.16E+05 | + | 1209 | 70 | 5.79 |
| **R3HDM4** | 26 | 2221913 | 2.23E+06 | + | 6914 | 400 | 5.79 |
| **GCK** | 28 | 1337732 | 1345191 | - | 7459 | 430 | 5.76 |
| **RIT1** | 29 | 1010930 | 1017355 | - | 6425 | 370 | 5.76 |
| **PMEL** | 30 | 916784 | 924259 | - | 7475 | 430 | 5.75 |

| | | | | | | | |
|---|---|---|---|---|---|---|---|
| **G0S2** | 24 | 5001046 | 5002090 | - | 1044 | 60 | 5.75 |
| **CACNG3** | 15 | 15767670 | 15791542 | + | 23872 | 1370 | 5.74 |
| **SMIM29** | 24 | 3584022 | 3589077 | + | 5055 | 290 | 5.74 |
| **C25H1orf232** | 25 | 913449 | 914321 | - | 872 | 50 | 5.73 |
| **LIX1L** | 29 | 1261439 | 1267898 | + | 6459 | 370 | 5.73 |
| **RPL35** | 20 | 889374 | 891122 | + | 1748 | 100 | 5.72 |
| **SCRN2** | 27 | 4167855 | 4171358 | - | 3503 | 200 | 5.71 |
| **CTSH** | 13 | 19815031 | 19822670 | - | 7639 | 436 | 5.71 |
| **CD3E** | 23 | 5917268 | 5920423 | - | 3155 | 180 | 5.71 |
| **BAZ2A** | 30 | 1102416 | 1121348 | - | 18932 | 1080 | 5.70 |
| **ABHD12B** | 6 | 2202951 | 2208045 | + | 5094 | 290 | 5.69 |
| **POLR2J** | 19 | 6128316 | 6130073 | + | 1757 | 100 | 5.69 |
| **HOOK2** | 36 | 180951 | 187638 | + | 6687 | 380 | 5.68 |
| **CD34** | 24 | 5475745 | 5489827 | + | 14082 | 800 | 5.68 |
| **ECM1** | 29 | 1445696 | 1448866 | + | 3170 | 180 | 5.68 |
| **GRHL3** | 25 | 457025 | 478024 | + | 20999 | 1190 | 5.67 |
| **MYDGF** | 26 | 982768 | 986298 | - | 3530 | 200 | 5.67 |
| **FCHO1** | 26 | 1681988 | 1693586 | - | 11598 | 657 | 5.66 |
| **KCNAB1** | 10 | 24822897 | 24889470 | - | 66573 | 3770 | 5.66 |
| **CDK10** | 11 | 1087862 | 1091041 | - | 3179 | 180 | 5.66 |
| **PYM1** | 30 | 913711 | 914948 | - | 1237 | 70 | 5.66 |
| **BAG4** | 28 | 4725647 | 4729716 | + | 4069 | 230 | 5.65 |
| **TTC3** | 2 | 1.17E+08 | 1.17E+08 | - | 39174 | 2207 | 5.63 |
| **ITPA** | 9 | 28012648 | 28016200 | - | 3552 | 200 | 5.63 |
| **TCIM** | 28 | 4953377 | 4954622 | + | 1245 | 70 | 5.62 |
| **GATAD2A** | 26 | 24541 | 50358 | - | 25817 | 1450 | 5.62 |
| **MRPS26** | 5 | 2004226 | 2007254 | - | 3028 | 170 | 5.61 |
| **CIAO2A** | 13 | 19590197 | 19592869 | - | 2672 | 150 | 5.61 |
| **PAQR7** | 25 | 854092 | 857306 | - | 3214 | 180 | 5.60 |

| | | | | | | | |
|---|---|---|---|---|---|---|---|
| **CNN1** | 36 | 115801 | 119735 | + | 3934 | 220 | 5.59 |
| **ADORA3** | 24 | 4903645 | 4906693 | + | 3048 | 170 | 5.58 |
| **ATP5MG** | 23 | 6815140 | 6816397 | + | 1257 | 70 | 5.57 |
| **MFSD1** | 10 | 24311178 | 24325748 | - | 14570 | 810 | 5.56 |
| **HOXB8** | 27 | 5066889 | 5068508 | + | 1619 | 90 | 5.56 |
| **OXLD1** | 18 | 10969019 | 10970821 | + | 1802 | 100 | 5.55 |
| **R3HCC1** | 28 | 3493408 | 3504084 | - | 10676 | 590 | 5.53 |
| **RAP1A** | 24 | 4859472 | 4.89E+06 | - | 30220 | 1670 | 5.53 |
| **UBXN11** | 25 | 4176299 | 4186112 | + | 9813 | 540 | 5.50 |
| **LCMT1** | 15 | 15620431 | 15638607 | + | 18176 | 1000 | 5.50 |
| **TMPRSS12** | 30 | 128066 | 132977 | + | 4911 | 270 | 5.50 |
| **GFRA2** | 28 | 3948119 | 3976686 | + | 28567 | 1570 | 5.50 |
| **BUD13** | 23 | 6281100 | 6286571 | + | 5471 | 300 | 5.48 |
| **TAS1R3** | 22 | 5816387 | 5822770 | - | 6383 | 350 | 5.48 |
| **NODAL** | 28 | 4706092 | 4708464 | - | 2372 | 130 | 5.48 |
| **PCYOX1** | 28 | 178600 | 184568 | + | 5968 | 327 | 5.48 |
| **IFT20** | 19 | 535931 | 537940 | + | 2009 | 110 | 5.48 |
| **KIF18B** | 27 | 1398757 | 1402594 | + | 3837 | 210 | 5.47 |
| **SYTL1** | 25 | 3814804 | 3819192 | - | 4388 | 240 | 5.47 |
| **TMEM214** | 3 | 1.16E+08 | 1.16E+08 | - | 7862 | 430 | 5.47 |
| **DCAF10** | Z | 82250708 | 82261869 | + | 11161 | 610 | 5.47 |
| **HOMER3** | 26 | 170814 | 191533 | + | 20719 | 1130 | 5.45 |
| **KCND3** | 24 | 4703001 | 4813410 | + | 110409 | 6020 | 5.45 |
| **HPCA** | 25 | 3439576 | 3442694 | + | 3118 | 170 | 5.45 |
| **KCNQ4** | 25 | 2991802 | 3009603 | + | 17801 | 970 | 5.45 |
| **BMP1** | 28 | 3805886 | 3826840 | - | 20954 | 1140 | 5.44 |
| **GPR157** | 22 | 6738709 | 6750475 | - | 11766 | 640 | 5.44 |
| **SUCNR1** | 10 | 25395385 | 25400349 | - | 4964 | 270 | 5.44 |
| **ATG101** | 30 | 1538912 | 1541486 | - | 2574 | 140 | 5.44 |

| | | | | | | | |
|---|---|---|---|---|---|---|---|
| **GOLGA7** | 28 | 5083384 | 5088291 | + | 4907 | 266 | 5.42 |
| **SF3B4** | 29 | 1314454 | 1318555 | - | 4101 | 222 | 5.41 |
| **SPG11** | 13 | 19593593 | 1.96E+07 | + | 31035 | 1680 | 5.41 |
| **SC5D** | 23 | 4363722 | 4367055 | - | 3333 | 180 | 5.40 |
| **CERS1** | 26 | 209428 | 215553 | + | 6125 | 330 | 5.39 |
| **SS18L2** | 11 | 20915652 | 20917137 | + | 1485 | 80 | 5.39 |
| **CEP164** | 23 | 6116492 | 6145246 | - | 28754 | 1548 | 5.38 |
| **PITHD1** | 25 | 324958 | 327187 | + | 2229 | 120 | 5.38 |
| **LTO1** | 6 | 61172881 | 61178825 | + | 5944 | 320 | 5.38 |
| **IL10RA** | 23 | 5975812 | 5979904 | - | 4092 | 220 | 5.38 |
| **IL12B** | 14 | 1366188 | 1371959 | + | 5771 | 310 | 5.37 |
| **SYPL2** | 24 | 1860647 | 1866793 | + | 6146 | 330 | 5.37 |
| **APRT** | 11 | 1467059 | 1470598 | + | 3539 | 190 | 5.37 |
| **ZNF710** | 13 | 19917338 | 19939696 | + | 22358 | 1200 | 5.37 |
| **C20H9orf116** | 20 | 2799401 | 2800335 | + | 934 | 50 | 5.35 |
| **RPS6** | Z | 34626134 | 34630993 | - | 4859 | 260 | 5.35 |
| **DNAJC30** | 19 | 7807354 | 7808850 | - | 1496 | 80 | 5.35 |
| **GPR3** | 25 | 3805292 | 3807914 | - | 2622 | 140 | 5.34 |
| **GZMM** | 26 | 2544675 | 2548424 | - | 3749 | 200 | 5.33 |
| **PCSK4** | 26 | 1801823 | 1808778 | - | 6955 | 370 | 5.32 |
| **SLC45A3** | 24 | 6038652 | 6059537 | + | 20885 | 1110 | 5.31 |
| **MYO1A** | 30 | 1180156 | 1189392 | - | 9236 | 490 | 5.31 |
| **SNRNP27** | 28 | 1171652 | 1174103 | + | 2451 | 130 | 5.30 |
| **FBXO41** | 5 | 983782 | 1005387 | - | 21605 | 1145 | 5.30 |
| **MYG1** | 30 | 805466 | 807732 | - | 2266 | 120 | 5.30 |
| **LIN28A** | 25 | 4140909 | 4153564 | - | 12655 | 670 | 5.29 |
| **LIPT2** | 2 | 214934 | 216634 | - | 1700 | 90 | 5.29 |
| **GLOD4** | 19 | 3058644 | 3062989 | - | 4345 | 230 | 5.29 |
| **TRMT12** | Z | 89759155 | 89762181 | - | 3026 | 160 | 5.29 |

| | | | | | | | |
|---|---|---|---|---|---|---|---|
| **LGI3** | 28 | 3836575 | 3840553 | + | 3978 | 210 | 5.28 |
| **ATP5IF1** | 25 | 935566 | 936325 | + | 759 | 40 | 5.27 |
| **PIK3R6** | 18 | 536841 | 566325 | - | 29484 | 1550 | 5.26 |
| **NKX6-3** | 28 | 5108560 | 5111226 | - | 2666 | 140 | 5.25 |
| **CIAO3** | 15 | 14989430 | 14999905 | + | 10475 | 550 | 5.25 |
| **CRTC1** | 26 | 277164 | 310717 | - | 33553 | 1760 | 5.25 |
| **SLC41A1** | 24 | 5978630 | 5995641 | + | 17011 | 890 | 5.23 |
| **KCNJ11** | 6 | 50169613 | 50172866 | + | 3253 | 170 | 5.23 |
| **DPM2** | 20 | 5958838 | 5.96E+06 | - | 2297 | 120 | 5.22 |
| **PTGES3** | 30 | 1127211 | 1135107 | - | 7896 | 410 | 5.19 |
| **CNKSR1** | 25 | 918790 | 925734 | + | 6944 | 360 | 5.18 |
| **MPZL3** | 23 | 5928321 | 5934691 | + | 6370 | 330 | 5.18 |
| **SNIP1** | 25 | 1534647 | 1538509 | + | 3862 | 200 | 5.18 |
| **TMPRSS4** | 23 | 5961822 | 5965689 | - | 3867 | 200 | 5.17 |
| **THOP1** | 26 | 1693821 | 1698661 | - | 4840 | 250 | 5.17 |
| **GNG12** | 9 | 1105776 | 1123404 | + | 17628 | 910 | 5.16 |
| **CLK3** | 13 | 1963060 | 1973716 | - | 10656 | 550 | 5.16 |
| **ARPC5L** | 20 | 830636 | 833736 | + | 3100 | 160 | 5.16 |
| **CRTAM** | 23 | 3913510 | 3925150 | - | 11640 | 600 | 5.15 |
| **ZCCHC7** | Z | 82308260 | 82395129 | - | 86869 | 4470 | 5.15 |
| **SNX27** | 29 | 1209611 | 1237410 | + | 27799 | 1430 | 5.14 |
| **LAMTOR5** | 24 | 2180612 | 2182752 | - | 2140 | 110 | 5.14 |
| **DENND4B** | 29 | 1598074 | 1615004 | + | 16930 | 870 | 5.14 |
| **LMCD1** | 12 | 20244672 | 20267062 | + | 22390 | 1150 | 5.14 |
| **TLX2** | 5 | 1313393 | 1316904 | + | 3511 | 180 | 5.13 |
| **TMEM81** | 24 | 6240637 | 6242393 | + | 1756 | 90 | 5.13 |
| **PIK3R5** | 18 | 569212 | 609009 | - | 39797 | 2039 | 5.12 |
| **DKK1** | 8 | 4278845 | 4280797 | + | 1952 | 100 | 5.12 |
| **DDHD2** | 28 | 4731743 | 4740920 | + | 9177 | 470 | 5.12 |

| | | | | | | | |
|---|---|---|---|---|---|---|---|
| **AAAS** | 30 | 797141 | 803975 | + | 6834 | 350 | 5.12 |
| **MED20** | 24 | 6795933 | 6799253 | + | 3320 | 170 | 5.12 |
| **MRPL58** | 18 | 4968670 | 4970820 | - | 2150 | 110 | 5.12 |
| **ARHGAP39** | 1 | 6 | 122728 | - | 122722 | 6270 | 5.11 |
| **BIN2** | 35 | 515549 | 523203 | + | 7654 | 390 | 5.10 |
| **TAGLN** | 23 | 6175947 | 6179095 | - | 3148 | 160 | 5.08 |
| **DHX37** | 17 | 1710328 | 1726874 | + | 16546 | 840 | 5.08 |
| **C18H17orf80** | 18 | 10823619 | 10826968 | + | 3349 | 170 | 5.08 |
| **PPT1** | 25 | 298700 | 302450 | - | 3750 | 190 | 5.07 |
| **FBXL20** | 27 | 3715371 | 3.75E+06 | + | 37703 | 1910 | 5.07 |
| **TTC31** | 5 | 1283863 | 1288011 | + | 4148 | 210 | 5.06 |
| **RPS15A** | 15 | 8335583 | 8339930 | - | 4347 | 220 | 5.06 |
| **PALM** | 26 | 2306556 | 2321377 | - | 14821 | 750 | 5.06 |
| **NEGR1** | 9 | 488344 | 676955 | + | 188611 | 9530 | 5.05 |
| **ZBTB39** | 30 | 1168523 | 1175255 | - | 6732 | 340 | 5.05 |
| **ATF4** | 4 | 52911706 | 52914684 | - | 2978 | 150 | 5.04 |
| **PPIE** | 25 | 177285 | 180264 | - | 2979 | 150 | 5.04 |
| **ELP3** | 3 | 1.16E+08 | 1.16E+08 | + | 105280 | 5300 | 5.03 |
| **NDUFAF8** | 18 | 11083418 | 11084014 | - | 596 | 30 | 5.03 |
| **PPP1R27** | 18 | 11782590 | 11784380 | - | 1790 | 90 | 5.03 |
| **SEMA4A** | 29 | 397249 | 404821 | - | 7572 | 380 | 5.02 |
| **KANSL2** | 30 | 1496779 | 1510936 | + | 14157 | 710 | 5.02 |
| **TMEM250** | 20 | 2371594 | 2375585 | - | 3991 | 200 | 5.01 |
| **CLP1** | 6 | 58528779 | 58530975 | + | 2196 | 110 | 5.01 |
| **LYPD6B** | 7 | 1831264 | 1836065 | - | 4801 | 240 | 5.00 |
| **RPS29** | 6 | 1809526 | 1810127 | - | 601 | 30 | 4.99 |
| **NFE2L1** | 27 | 4954116 | 4959325 | + | 5209 | 260 | 4.99 |
| **LGALS3** | 6 | 3508254 | 3512871 | + | 4617 | 230 | 4.98 |
| **YARS1** | 25 | 3396757 | 3401578 | - | 4821 | 240 | 4.98 |

| | | | | | | | |
|---|---|---|---|---|---|---|---|
| **PLAAT1** | 10 | 14109402 | 14113822 | - | 4420 | 220 | 4.98 |
| **WDR54** | 5 | 1231132 | 1236558 | + | 5426 | 270 | 4.98 |
| **TAC3** | 30 | 1176763 | 1178172 | - | 1409 | 70 | 4.97 |
| **ACP5** | 36 | 121069 | 123284 | - | 2215 | 110 | 4.97 |
| **JTB** | 29 | 1580288 | 1582303 | + | 2015 | 100 | 4.96 |
| **TPP1** | 2 | 1.19E+08 | 1.19E+08 | + | 5643 | 280 | 4.96 |
| **RPL37** | Z | 16810482 | 16813102 | + | 2620 | 130 | 4.96 |
| **DCTN1** | 5 | 1159192 | 1219275 | - | 60083 | 2970 | 4.94 |
| **ZC3HC1** | 4 | 740547 | 7.55E+05 | - | 14770 | 730 | 4.94 |
| **NOP56** | 5 | 1153943 | 1158400 | - | 4457 | 220 | 4.94 |
| **PIK3C2B** | 24 | 6414043 | 6445346 | + | 31303 | 1540 | 4.92 |
| **SGTA** | 26 | 1698949 | 1710135 | + | 11186 | 550 | 4.92 |
| **ARHGAP25** | 28 | 688315 | 708674 | + | 20359 | 1000 | 4.91 |
| **LASP1** | 27 | 3810482 | 3837788 | - | 27306 | 1340 | 4.91 |
| **IL12RB1** | 26 | 3023554 | 3027640 | - | 4086 | 200 | 4.89 |
| **MADCAM1** | 26 | 2405204 | 2407452 | - | 2248 | 110 | 4.89 |
| **GLI1** | 30 | 1365151 | 1374349 | + | 9198 | 450 | 4.89 |
| **CRTC2** | 29 | 1588082 | 1596668 | + | 8586 | 420 | 4.89 |
| **MRTO4** | 22 | 1292836 | 1295290 | + | 2454 | 120 | 4.89 |
| **COL16A1** | 25 | 3026996 | 3052869 | - | 25873 | 1260 | 4.87 |
| **SNAPIN** | 29 | 1676323 | 1677557 | - | 1234 | 60 | 4.86 |
| **RABEPK** | 20 | 824719 | 827188 | - | 2469 | 120 | 4.86 |
| **YPEL4** | 6 | 58525965 | 58528848 | - | 2883 | 140 | 4.86 |
| **GMPS** | 10 | 24896710 | 24924134 | - | 27424 | 1330 | 4.85 |
| **ENTR1** | 20 | 2596077 | 2598758 | - | 2681 | 130 | 4.85 |
| **MXD1** | 28 | 1175126 | 1182966 | + | 7840 | 380 | 4.85 |
| **ABLIM3** | 14 | 1701623 | 1747878 | - | 46255 | 2240 | 4.84 |
| **PIWIL2** | 28 | 3785463 | 3794139 | - | 8676 | 420 | 4.84 |
| **IGSF10** | 10 | 25432918 | 25443900 | + | 10982 | 530 | 4.83 |

| | | | | | | | |
|---|---|---|---|---|---|---|---|
| **YIPF5** | 14 | 681060 | 688320 | + | 7260 | 350 | 4.82 |
| **NCAM1** | 23 | 5710063 | 5797100 | + | 87037 | 4190 | 4.81 |
| **KRTCAP2** | 29 | 764311 | 766185 | - | 1874 | 90 | 4.80 |
| **FKBP8** | 26 | 372187 | 377604 | + | 5417 | 260 | 4.80 |
| **NDUFA8** | 20 | 1702037 | 1704540 | + | 2503 | 120 | 4.79 |
| **MVD** | 11 | 1514994 | 1518958 | + | 3964 | 190 | 4.79 |
| **RBM43** | 7 | 1597824 | 1602414 | + | 4590 | 220 | 4.79 |
| **RPL23** | 27 | 3838706 | 3841632 | + | 2926 | 140 | 4.78 |
| **SUGP1** | 26 | 70721 | 88496 | + | 17775 | 850 | 4.78 |
| **ZNF652** | 27 | 4887914 | 4907787 | + | 19873 | 950 | 4.78 |
| **FIGNL2** | 30 | 49901 | 64546 | + | 14645 | 700 | 4.78 |
| **CCT7** | 5 | 957945 | 975780 | + | 17835 | 852 | 4.78 |
| **FADD** | 6 | 60928130 | 60931489 | - | 3359 | 160 | 4.76 |
| **MRPS21** | 29 | 1413785 | 1414835 | + | 1050 | 50 | 4.76 |
| **RSPO1** | 25 | 1500046 | 1512656 | + | 12610 | 600 | 4.76 |
| **EFNA4** | 29 | 725960 | 730584 | + | 4624 | 220 | 4.76 |
| **RHCE** | 25 | 698538 | 706528 | - | 7990 | 380 | 4.76 |
| **SAV1** | 6 | 2121671 | 2137893 | - | 16222 | 770 | 4.75 |
| **STRIT1** | 10 | 25021539 | 25025122 | + | 3583 | 170 | 4.74 |
| **TRIM54** | 3 | 1.17E+08 | 1.17E+08 | - | 3374 | 160 | 4.74 |
| **FMNL3** | 30 | 1523173 | 1537120 | + | 13947 | 660 | 4.73 |
| **SMIM4** | 12 | 4672453 | 4674355 | + | 1902 | 90 | 4.73 |
| **PTPN23** | 1 | 1.56E+08 | 1.56E+08 | - | 12757 | 600 | 4.70 |
| **ATP5F1E** | 16 | 3567247 | 3568524 | - | 1277 | 60 | 4.70 |
| **CAP1** | 25 | 286419 | 298560 | + | 12141 | 570 | 4.69 |
| **FAM110D** | 25 | 911633 | 913124 | + | 1491 | 70 | 4.69 |
| **POLR2E** | 26 | 2131662 | 2134007 | + | 2345 | 110 | 4.69 |
| **MRPL12** | 18 | 11753351 | 11757361 | + | 4010 | 188 | 4.69 |
| **SRPX2** | Z | 89745631 | 89750968 | + | 5337 | 250 | 4.68 |

| | | | | | | | |
|---|---|---|---|---|---|---|---|
| **CERS2** | 29 | 1493457 | 1498158 | - | 4701 | 220 | 4.68 |
| **ADGRD1** | 17 | 301758 | 425324 | - | 123566 | 5780 | 4.68 |
| **DAP3** | 29 | 937889 | 954368 | + | 16479 | 770 | 4.67 |
| **PIGV** | 25 | 3981991 | 3987774 | - | 5783 | 270 | 4.67 |
| **ELOF1** | 36 | 119806 | 120877 | - | 1071 | 50 | 4.67 |
| **SLC50A1** | 29 | 758173 | 760680 | + | 2507 | 117 | 4.67 |
| **LCK** | 25 | 3247813 | 3254652 | - | 6839 | 319 | 4.66 |
| **BAIAP2** | 18 | 11124371 | 11165115 | - | 40744 | 1900 | 4.66 |
| **ZDHHC4** | 15 | 15508235 | 15515424 | - | 7189 | 335 | 4.66 |
| **PSMD5** | 20 | 2127627 | 2134714 | + | 7087 | 330 | 4.66 |
| **DONSON** | 2 | 118000000 | 118000000 | - | 4940 | 230 | 4.66 |
| **OTUD7B** | 29 | 1322137 | 1343451 | - | 21314 | 990 | 4.64 |
| **FMNL1** | 27 | 1288341 | 1311379 | - | 23038 | 1070 | 4.64 |
| **PMF1** | 29 | 381905 | 385787 | - | 3882 | 180 | 4.64 |
| **AICDA** | 2 | 30396714 | 30398225 | - | 1511 | 70 | 4.63 |
| **TENT5B** | 25 | 3896553 | 3902820 | + | 6267 | 290 | 4.63 |
| **PHLDA2** | 6 | 48666753 | 48668266 | - | 1513 | 70 | 4.63 |
| **STX8** | 18 | 694670 | 800132 | - | 105462 | 4870 | 4.62 |
| **HJV** | 29 | 1292494 | 1295745 | - | 3251 | 150 | 4.61 |
| **CCDC142** | 5 | 1278789 | 1283558 | - | 4769 | 220 | 4.61 |
| **TEX12** | 23 | 5534062 | 5538834 | + | 4772 | 220 | 4.61 |
| **TACC1** | 28 | 4861454 | 4881425 | + | 19971 | 920 | 4.61 |
| **PRPF40A** | 7 | 1155401 | 1174513 | + | 19112 | 880 | 4.60 |
| **TMEM39B** | 25 | 3200510 | 3215286 | + | 14776 | 680 | 4.60 |
| **ATOH7** | 8 | 1483199 | 1483851 | - | 652 | 30 | 4.60 |
| **AGRN** | 22 | 6091494 | 6195984 | - | 104490 | 4800 | 4.59 |
| **ARF3** | 30 | 1463402 | 1465797 | + | 2395 | 110 | 4.59 |
| **LRRC3** | 10 | 4475824 | 4478222 | - | 2398 | 110 | 4.59 |
| **JAKMIP3** | 8 | 35405608 | 35472158 | + | 66550 | 3050 | 4.58 |
| **PMP22** | 18 | 355592 | 372412 | - | 16820 | 770 | 4.58 |

| | | | | | | | |
|---|---|---|---|---|---|---|---|
| **CETN2** | 21 | 1054395 | 1058117 | + | 3722 | 170 | 4.57 |
| **TCERG1L** | 8 | 35133939 | 35184784 | - | 50845 | 2320 | 4.56 |
| **LYPLA2** | 25 | 327437 | 334896 | + | 7459 | 340 | 4.56 |
| **CHURC1** | 6 | 59155520 | 59157497 | - | 1977 | 90 | 4.55 |
| **MRPL28** | 15 | 2867308 | 2870166 | - | 2858 | 130 | 4.55 |
| **PAQR5** | 13 | 19329874 | 19344392 | + | 14518 | 660 | 4.55 |
| **DSTYK** | 24 | 6193469 | 6218553 | + | 25084 | 1140 | 4.54 |
| **CALU** | 4 | 206396 | 220929 | - | 14533 | 660 | 4.54 |
| **RBFOX3** | 18 | 11470652 | 11602397 | + | 131745 | 5980 | 4.54 |
| **UBB** | 17 | 1731525 | 1735718 | + | 4193 | 190 | 4.53 |
| **PEBP4** | 28 | 3563702 | 3640723 | + | 77021 | 3490 | 4.53 |
| **GPATCH3** | 25 | 3951407 | 3955821 | + | 4414 | 200 | 4.53 |
| **CALR** | 36 | 148704 | 151796 | - | 3092 | 140 | 4.53 |
| **TNFSF4** | 9 | 27650863 | 27657496 | + | 6633 | 300 | 4.52 |
| **NDEL1** | 18 | 925278 | 948970 | - | 23692 | 1070 | 4.52 |
| **FECH** | Z | 262421 | 274821 | - | 12400 | 560 | 4.52 |
| **SMIM32** | 14 | 8755666 | 8758548 | + | 2882 | 130 | 4.51 |
| **CADM3** | 29 | 1124868 | 1134181 | + | 9313 | 420 | 4.51 |
| **CHCHD10** | 17 | 11931594 | 11932926 | + | 1332 | 60 | 4.50 |
| **DEF8** | 11 | 1001036 | 1006592 | - | 5556 | 250 | 4.50 |
| **GALE** | 25 | 334800 | 338582 | - | 3782 | 170 | 4.49 |
| **PPCS** | 22 | 794476 | 797908 | + | 3432 | 154 | 4.49 |
| **SDF4** | 22 | 5999657 | 6013285 | + | 13628 | 610 | 4.48 |
| **C1H1orf35** | 1 | 154000000 | 154000000 | + | 3355 | 150 | 4.47 |
| **RAB34** | 19 | 4314957 | 4317869 | + | 2912 | 130 | 4.46 |
| **TMCC2** | 24 | 6182712 | 6192568 | - | 9856 | 440 | 4.46 |
| **MACO1** | 25 | 706309 | 731656 | + | 25347 | 1130 | 4.46 |
| **FAM72A** | 24 | 5883769 | 5887810 | + | 4041 | 180 | 4.45 |
| **HPCAL4** | 25 | 231018 | 234611 | - | 3593 | 160 | 4.45 |
| **VDR** | 30 | 808863 | 827054 | - | 18191 | 810 | 4.45 |

| | | | | | | | |
|---|---|---|---|---|---|---|---|
| **SV2A** | 29 | 1300610 | 1312296 | - | 11686 | 520 | 4.45 |
| **CLDN25** | 23 | 6797600 | 6798275 | - | 675 | 30 | 4.44 |
| **SSR3** | 10 | 24816384 | 24819766 | + | 3382 | 150 | 4.44 |
| **IRF8** | 11 | 2383815 | 2393294 | + | 9479 | 420 | 4.43 |
| **PIFO** | 24 | 6738875 | 6752870 | + | 13995 | 620 | 4.43 |
| **ARID3B** | 13 | 1981365 | 2011860 | - | 30495 | 1350 | 4.43 |
| **GCDH** | 36 | 158474 | 162092 | - | 3618 | 160 | 4.42 |
| **TNFRSF4** | 22 | 6013728 | 6022341 | + | 8613 | 380 | 4.41 |
| **CENPH** | Z | 7720810 | 7727847 | - | 7037 | 310 | 4.41 |
| **TMEM161A** | 26 | 111149 | 118871 | + | 7722 | 340 | 4.40 |
| **VGLL1** | W | 9423871 | 9434560 | + | 10689 | 470 | 4.40 |
| **GINS2** | 11 | 2337087 | 2341184 | - | 4097 | 180 | 4.39 |
| **GPN2** | 25 | 3957296 | 3962076 | + | 4780 | 210 | 4.39 |
| **DPYSL4** | 8 | 35473451 | 35487565 | + | 14114 | 620 | 4.39 |
| **TRAF3IP3** | 24 | 4974144 | 4985760 | - | 11616 | 510 | 4.39 |
| **SLC27A3** | 29 | 1626696 | 1629889 | - | 3193 | 140 | 4.38 |
| **ELMOD3** | 28 | 1252216 | 1257760 | - | 5544 | 243 | 4.38 |
| **NPEPPS** | 27 | 4028955 | 4072771 | + | 43816 | 1920 | 4.38 |
| **ALKBH2** | 17 | 4105861 | 4109289 | - | 3428 | 150 | 4.38 |
| **SPCS2** | 2 | 312689 | 316346 | + | 3657 | 160 | 4.38 |
| **RAPGEF3** | 30 | 452349 | 466977 | + | 14628 | 640 | 4.38 |
| **ARHGAP17** | 15 | 15578141 | 15619983 | - | 41842 | 1830 | 4.37 |
| **ELAVL3** | 36 | 101149 | 106408 | - | 5259 | 230 | 4.37 |
| **TMEM167B** | 24 | 204969 | 206801 | - | 1832 | 80 | 4.37 |
| **YRDC** | 25 | 1372091 | 1373925 | + | 1834 | 80 | 4.36 |
| **PLEKHN1** | 22 | 6298202 | 6319525 | - | 21323 | 930 | 4.36 |
| **TM6SF2** | 26 | 90338 | 94698 | + | 4360 | 190 | 4.36 |
| **TTC36** | 23 | 6860027 | 6861175 | + | 1148 | 50 | 4.36 |
| **LHX5** | 17 | 9794943 | 9802301 | + | 7358 | 320 | 4.35 |
| **AGBL5** | 3 | 116000000 | 116000000 | - | 10587 | 460 | 4.34 |

| | | | | | | | |
|---|---|---|---|---|---|---|---|
| **KCNE2** | 2 | 118000000 | 118000000 | + | 2533 | 110 | 4.34 |
| **FHIP1B** | 2 | 119000000 | 119000000 | + | 8064 | 350 | 4.34 |
| **GAS2L1** | 17 | 12326789 | 12336484 | + | 9695 | 420 | 4.33 |
| **STAM2** | 7 | 1282855 | 1306411 | + | 23556 | 1020 | 4.33 |
| **RAB13** | 29 | 1577842 | 1580156 | + | 2314 | 100 | 4.32 |
| **RPL4** | 13 | 18521244 | 18526349 | - | 5105 | 220 | 4.31 |
| **SMAD4** | W | 10296387 | 10334517 | - | 38130 | 1640 | 4.30 |
| **TMEM106C** | 30 | 830025 | 834213 | + | 4188 | 180 | 4.30 |
| **CDK18** | 24 | 6128360 | 6154910 | - | 26550 | 1140 | 4.29 |
| **GFPT1** | 28 | 952012 | 989745 | - | 37733 | 1620 | 4.29 |
| **ANAPC11** | 18 | 11809891 | 11810357 | + | 466 | 20 | 4.29 |
| **GDF11** | 30 | 571302 | 576431 | + | 5129 | 220 | 4.29 |
| **NUDT19** | 11 | 16003080 | 16008679 | + | 5599 | 240 | 4.29 |
| **SPATA16** | 28 | 1588843 | 1601210 | - | 12367 | 530 | 4.29 |
| **STK32C** | 8 | 35490881 | 35562792 | - | 71911 | 3080 | 4.28 |
| **SPSB1** | 22 | 6792209 | 6819769 | + | 27560 | 1180 | 4.28 |
| **BSX** | 23 | 3886960 | 3890230 | + | 3270 | 140 | 4.28 |
| **RETSAT** | 28 | 1258143 | 1264693 | + | 6550 | 280 | 4.27 |
| **BLOC1S3** | 18 | 2005768 | 2006939 | + | 1171 | 50 | 4.27 |
| **CKMT2** | Z | 81864039 | 81886763 | - | 22724 | 970 | 4.27 |
| **NKAPD1** | 23 | 5520703 | 5524922 | + | 4219 | 180 | 4.27 |
| **SPDEF** | 24 | 3533145 | 3535021 | + | 1876 | 80 | 4.26 |
| **SLC35E4** | 17 | 12745151 | 12749144 | + | 3993 | 170 | 4.26 |
| **GOLPH3L** | 29 | 1045501 | 1054597 | + | 9096 | 387 | 4.25 |
| **MTMR11** | 29 | 1318857 | 1323572 | - | 4715 | 200 | 4.24 |
| **PTGDR** | 6 | 2493980 | 2498224 | + | 4244 | 180 | 4.24 |

**Table S14 - Gene ontology analysis on PhyloP accelerated genes.** The top 5% (606) genes with more accelerated bases overlapped with the entire gene sequence ("LOC" genes excluded) were tested. The first 100 lines are showed.

| GO term | p.geomean | stat.mean | p.val | q.val | set.size | exp1 | GO term name |
|---|---|---|---|---|---|---|---|
| GO:0005840 | 0.002716583 | 2.810582782 | 0.002716583 | 0.911119217 | 131 | 0.002716583 | ribosome |
| GO:0003735 | 0.003101465 | 2.774255948 | 0.003101465 | 0.911119217 | 105 | 0.003101465 | structural constituent of ribosome |
| GO:0006614 | 0.005429905 | 2.601595637 | 0.005429905 | 0.911119217 | 63 | 0.005429905 | SRP-dependent cotranslational protein targeting to membrane |
| GO:0000184 | 0.007528512 | 2.464413196 | 0.007528512 | 0.911119217 | 85 | 0.007528512 | nuclear-transcribed mRNA catabolic process, nonsense-mediated decay |
| GO:0002181 | 0.008235137 | 2.447373983 | 0.008235137 | 0.911119217 | 59 | 0.008235137 | cytoplasmic translation |
| GO:0032729 | 0.008797316 | 2.462019923 | 0.008797316 | 0.911119217 | 35 | 0.008797316 | positive regulation of interferon-gamma production |
| GO:0006397 | 0.012830402 | 2.23817656 | 0.012830402 | 0.911119217 | 277 | 0.012830402 | mRNA processing |
| GO:0022626 | 0.013062291 | 2.266066889 | 0.013062291 | 0.911119217 | 56 | 0.013062291 | cytosolic ribosome |
| GO:0008380 | 0.013458016 | 2.221247385 | 0.013458016 | 0.911119217 | 223 | 0.013458016 | RNA splicing |
| GO:0002250 | 0.013871074 | 2.225163114 | 0.013871074 | 0.911119217 | 87 | 0.013871074 | adaptive immune response |
| GO:0006413 | 0.014907866 | 2.191351528 | 0.014907866 | 0.911119217 | 105 | 0.014907866 | translational initiation |
| GO:0042104 | 0.016806747 | 2.336522582 | 0.016806747 | 0.911119217 | 13 | 0.016806747 | positive regulation of activated T cell proliferation |
| GO:0019083 | 0.017319288 | 2.136969473 | 0.017319288 | 0.911119217 | 78 | 0.017319288 | viral transcription |
| GO:0032418 | 0.017955646 | 2.194029443 | 0.017955646 | 0.911119217 | 24 | 0.017955646 | lysosome localization |
| GO:0022625 | 0.021557571 | 2.069554285 | 0.021557571 | 0.911119217 | 40 | 0.021557571 | cytosolic large ribosomal subunit |
| GO:0001772 | 0.023825023 | 2.045293123 | 0.023825023 | 0.911119217 | 29 | 0.023825023 | immunological synapse |
| GO:0050821 | 0.024176257 | 1.984683903 | 0.024176257 | 0.911119217 | 139 | 0.024176257 | protein stabilization |
| GO:0006412 | 0.029402233 | 1.89515675 | 0.029402233 | 0.911119217 | 222 | 0.029402233 | translation |
| GO:0008047 | 0.034906035 | 1.850843197 | 0.034906035 | 0.911119217 | 37 | 0.034906035 | enzyme activator activity |
| GO:0005681 | 0.037000102 | 1.795931298 | 0.037000102 | 0.911119217 | 120 | 0.037000102 | spliceosomal complex |
| GO:0000398 | 0.039942598 | 1.757022581 | 0.039942598 | 0.911119217 | 179 | 0.039942598 | mRNA splicing, via spliceosome |
| GO:0032740 | 0.040221655 | 1.859315227 | 0.040221655 | 0.911119217 | 14 | 0.040221655 | positive regulation of interleukin-17 production |
| GO:0015935 | 0.040221655 | 1.859315227 | 0.040221655 | 0.911119217 | 14 | 0.040221655 | small ribosomal subunit |
| GO:0006364 | 0.040870628 | 1.746937147 | 0.040870628 | 0.911119217 | 161 | 0.040870628 | rRNA processing |
| GO:0044183 | 0.043481786 | 1.756423925 | 0.043481786 | 0.911119217 | 28 | 0.043481786 | protein folding chaperone |
| GO:0070125 | 0.045566911 | 1.70738492 | 0.045566911 | 0.911119217 | 59 | 0.045566911 | mitochondrial translational elongation |
| GO:0048704 | 0.046525732 | 1.704361261 | 0.046525732 | 0.911119217 | 44 | 0.046525732 | embryonic skeletal system morphogenesis |
| GO:0007040 | 0.049862505 | 1.681791989 | 0.049862505 | 0.911119217 | 31 | 0.049862505 | lysosome organization |
| GO:0070126 | 0.050380048 | 1.65683338 | 0.050380048 | 0.911119217 | 62 | 0.050380048 | mitochondrial translational termination |
| GO:0005249 | 0.054023965 | 1.626924905 | 0.054023965 | 0.911119217 | 48 | 0.054023965 | voltage-gated potassium channel activity |
| GO:1904724 | 0.054357756 | 1.63477738 | 0.054357756 | 0.911119217 | 33 | 0.054357756 | tertiary granule lumen |
| GO:0009952 | 0.055394197 | 1.605187417 | 0.055394197 | 0.911119217 | 82 | 0.055394197 | anterior/posterior pattern specification |
| GO:0043236 | 0.056815343 | 1.636389905 | 0.056815343 | 0.911119217 | 20 | 0.056815343 | laminin binding |
| GO:0051015 | 0.057901133 | 1.577533247 | 0.057901133 | 0.911119217 | 158 | 0.057901133 | actin filament binding |
| GO:0002230 | 0.05974924 | 1.60586527 | 0.05974924 | 0.911119217 | 21 | 0.05974924 | positive regulation of defense response to virus by host |

| | | | | | | | |
|---|---|---|---|---|---|---|---|
| **GO:0002376** | 0.062681674 | 1.535279046 | 0.062681674 | 0.911119217 | 264 | 0.062681674 | immune system process |
| **GO:0006369** | 0.065791962 | 1.548118599 | 0.065791962 | 0.911119217 | 23 | 0.065791962 | termination of RNA polymerase II transcription |
| **GO:0005685** | 0.066238124 | 1.618428593 | 0.066238124 | 0.911119217 | 10 | 0.066238124 | U1 snRNP |
| **GO:0006898** | 0.066598337 | 1.512193848 | 0.066598337 | 0.911119217 | 71 | 0.066598337 | receptor-mediated endocytosis |
| **GO:1990904** | 0.068195131 | 1.495834485 | 0.068195131 | 0.911119217 | 108 | 0.068195131 | ribonucleoprotein complex |
| **GO:0018279** | 0.06890387 | 1.520620908 | 0.06890387 | 0.911119217 | 24 | 0.06890387 | protein N-linked glycosylation via asparagine |
| **GO:0022627** | 0.06890387 | 1.520620908 | 0.06890387 | 0.911119217 | 24 | 0.06890387 | cytosolic small ribosomal subunit |
| **GO:0003729** | 0.069208769 | 1.486254831 | 0.069208769 | 0.911119217 | 147 | 0.069208769 | mRNA binding |
| **GO:0031083** | 0.070451031 | 1.567562579 | 0.070451031 | 0.911119217 | 11 | 0.070451031 | BLOC-1 complex |
| **GO:0001833** | 0.070451031 | 1.567562579 | 0.070451031 | 0.911119217 | 11 | 0.070451031 | inner cell mass cell proliferation |
| **GO:0097623** | 0.070451031 | 1.567562579 | 0.070451031 | 0.911119217 | 11 | 0.070451031 | potassium ion export across plasma membrane |
| **GO:0010614** | 0.070451031 | 1.567562579 | 0.070451031 | 0.911119217 | 11 | 0.070451031 | negative regulation of cardiac muscle hypertrophy |
| **GO:0008250** | 0.070451031 | 1.567562579 | 0.070451031 | 0.911119217 | 11 | 0.070451031 | oligosaccharyltransferase complex |
| **GO:0007625** | 0.070451031 | 1.567562579 | 0.070451031 | 0.911119217 | 11 | 0.070451031 | grooming behavior |
| **GO:0045667** | 0.070451031 | 1.567562579 | 0.070451031 | 0.911119217 | 11 | 0.070451031 | regulation of osteoblast differentiation |
| **GO:0071005** | 0.071657646 | 1.482747007 | 0.071657646 | 0.911119217 | 40 | 0.071657646 | U2-type precatalytic spliceosome |
| **GO:0032735** | 0.072077533 | 1.49389815 | 0.072077533 | 0.911119217 | 25 | 0.072077533 | positive regulation of interleukin-12 production |
| **GO:1904813** | 0.074457845 | 1.450419271 | 0.074457845 | 0.911119217 | 93 | 0.074457845 | ficolin-1-rich granule lumen |
| **GO:0099524** | 0.074680058 | 1.522230301 | 0.074680058 | 0.911119217 | 12 | 0.074680058 | postsynaptic cytosol |
| **GO:0043202** | 0.07677699 | 1.435676585 | 0.07677699 | 0.911119217 | 76 | 0.07677699 | lysosomal lumen |
| **GO:0007166** | 0.07820989 | 1.421107298 | 0.07820989 | 0.911119217 | 153 | 0.07820989 | cell surface receptor signaling pathway |
| **GO:0006506** | 0.078612514 | 1.442456555 | 0.078612514 | 0.911119217 | 27 | 0.078612514 | GPI anchor biosynthetic process |
| **GO:0071007** | 0.078612514 | 1.442456555 | 0.078612514 | 0.911119217 | 27 | 0.078612514 | U2-type catalytic step 2 spliceosome |
| **GO:0086005** | 0.078951363 | 1.480921145 | 0.078951363 | 0.911119217 | 13 | 0.078951363 | ventricular cardiac muscle cell action potential |
| **GO:1900029** | 0.078951363 | 1.480921145 | 0.078951363 | 0.911119217 | 13 | 0.078951363 | positive regulation of ruffle assembly |
| **GO:0071157** | 0.078951363 | 1.480921145 | 0.078951363 | 0.911119217 | 13 | 0.078951363 | negative regulation of cell cycle arrest |
| **GO:0030425** | 0.080269371 | 1.404754898 | 0.080269371 | 0.911119217 | 368 | 0.080269371 | dendrite |
| **GO:0043025** | 0.082882634 | 1.38758491 | 0.082882634 | 0.911119217 | 326 | 0.082882634 | neuronal cell body |
| **GO:0030030** | 0.082969798 | 1.388971188 | 0.082969798 | 0.911119217 | 156 | 0.082969798 | cell projection organization |
| **GO:0101031** | 0.083280707 | 1.442650418 | 0.083280707 | 0.911119217 | 14 | 0.083280707 | chaperone complex |
| **GO:0060307** | 0.083280707 | 1.442650418 | 0.083280707 | 0.911119217 | 14 | 0.083280707 | regulation of ventricular cardiac muscle cell membrane repolarization |
| **GO:0048489** | 0.083280707 | 1.442650418 | 0.083280707 | 0.911119217 | 14 | 0.083280707 | synaptic vesicle transport |
| **GO:0097067** | 0.083280707 | 1.442650418 | 0.083280707 | 0.911119217 | 14 | 0.083280707 | cellular response to thyroid hormone stimulus |
| **GO:0001736** | 0.083280707 | 1.442650418 | 0.083280707 | 0.911119217 | 14 | 0.083280707 | establishment of planar polarity |
| **GO:0048026** | 0.087677779 | 1.406747869 | 0.087677779 | 0.911119217 | 15 | 0.087677779 | positive regulation of mRNA splicing, via spliceosome |
| **GO:1901379** | 0.087677779 | 1.406747869 | 0.087677779 | 0.911119217 | 15 | 0.087677779 | regulation of potassium ion transmembrane transport |
| **GO:0050776** | 0.088442541 | 1.363651355 | 0.088442541 | 0.911119217 | 46 | 0.088442541 | regulation of immune response |

| | | | | | | | |
|---|---|---|---|---|---|---|---|
| **GO:1901224** | 0.094434506 | 1.325698095 | 0.094434506 | 0.911119217 | 48 | 0.094434506 | positive regulation of NIK/NF-kappaB signaling |
| **GO:0008076** | 0.095743893 | 1.314250204 | 0.095743893 | 0.911119217 | 66 | 0.095743893 | voltage-gated potassium channel complex |
| **GO:0043198** | 0.096051501 | 1.322940357 | 0.096051501 | 0.911119217 | 32 | 0.096051501 | dendritic shaft |
| **GO:0035145** | 0.096696519 | 1.340285679 | 0.096696519 | 0.911119217 | 17 | 0.096696519 | exon-exon junction complex |
| **GO:0019882** | 0.096696519 | 1.340285679 | 0.096696519 | 0.911119217 | 17 | 0.096696519 | antigen processing and presentation |
| **GO:0044325** | 0.096948063 | 1.303930327 | 0.096948063 | 0.911119217 | 104 | 0.096948063 | ion channel binding |
| **GO:0007052** | 0.099161635 | 1.290971981 | 0.099161635 | 0.911119217 | 105 | 0.099161635 | mitotic spindle organization |
| **GO:0015459** | 0.099726675 | 1.30027915 | 0.099726675 | 0.911119217 | 33 | 0.099726675 | potassium channel regulator activity |
| **GO:0009925** | 0.099726675 | 1.30027915 | 0.099726675 | 0.911119217 | 33 | 0.099726675 | basal plasma membrane |
| **GO:0042102** | 0.099726675 | 1.30027915 | 0.099726675 | 0.911119217 | 33 | 0.099726675 | positive regulation of T cell proliferation |
| **GO:0051603** | 0.099726675 | 1.30027915 | 0.099726675 | 0.911119217 | 33 | 0.099726675 | proteolysis involved in cellular protein catabolic process |
| **GO:0016032** | 0.100603432 | 1.279193712 | 0.100603432 | 0.911119217 | 409 | 0.100603432 | viral process |
| **GO:0034599** | 0.101180025 | 1.282197682 | 0.101180025 | 0.911119217 | 68 | 0.101180025 | cellular response to oxidative stress |
| **GO:0008202** | 0.101180025 | 1.282197682 | 0.101180025 | 0.911119217 | 68 | 0.101180025 | steroid metabolic process |
| **GO:0045672** | 0.101323697 | 1.309126549 | 0.101323697 | 0.911119217 | 18 | 0.101323697 | positive regulation of osteoclast differentiation |
| **GO:0044306** | 0.101323697 | 1.309126549 | 0.101323697 | 0.911119217 | 18 | 0.101323697 | neuron projection terminus |
| **GO:0005244** | 0.1036723 | 1.265222163 | 0.1036723 | 0.911119217 | 107 | 0.1036723 | voltage-gated ion channel activity |
| **GO:0008198** | 0.10603092 | 1.279068349 | 0.10603092 | 0.911119217 | 19 | 0.10603092 | ferrous iron binding |
| **GO:0043113** | 0.10603092 | 1.279068349 | 0.10603092 | 0.911119217 | 19 | 0.10603092 | receptor clustering |
| **GO:0031369** | 0.10603092 | 1.279068349 | 0.10603092 | 0.911119217 | 19 | 0.10603092 | translation initiation factor binding |
| **GO:0048306** | 0.107000414 | 1.251988767 | 0.107000414 | 0.911119217 | 52 | 0.107000414 | calcium-dependent protein binding |
| **GO:0032438** | 0.11081826 | 1.249959581 | 0.11081826 | 0.911119217 | 20 | 0.11081826 | melanosome organization |
| **GO:0051602** | 0.11081826 | 1.249959581 | 0.11081826 | 0.911119217 | 20 | 0.11081826 | response to electrical stimulus |
| **GO:0005743** | 0.112167985 | 1.216380118 | 0.112167985 | 0.911119217 | 304 | 0.112167985 | mitochondrial inner membrane |
| **GO:0001669** | 0.113568155 | 1.216127821 | 0.113568155 | 0.911119217 | 54 | 0.113568155 | acrosomal vesicle |
| **GO:0042277** | 0.113568155 | 1.216127821 | 0.113568155 | 0.911119217 | 54 | 0.113568155 | peptide binding |
| **GO:0004896** | 0.115035512 | 1.212904195 | 0.115035512 | 0.911119217 | 37 | 0.115035512 | cytokine receptor activity |
| **GO:0034765** | 0.115431147 | 1.201785174 | 0.115431147 | 0.911119217 | 112 | 0.115431147 | regulation of ion transmembrane transport |

**Table S15 - PhyloP conserved genes.** The top 5% (606) genes with most conserved bases overlapped with CDS are reported. "LOC" genes were discarded.

| Gene | Chr | Start | End | Strand | Gene length | CDS length | Overlaps | % |
|---|---|---|---|---|---|---|---|---|
| CAMK2N2 | 10 | 17272192 | 17276215 | - | 4023 | 240 | 214 | 89.2 |
| INHBB | 7 | 12006466 | 12012818 | - | 6352 | 1179 | 1019 | 86.4 |
| SUMO2 | 18 | 4890590 | 4897641 | + | 7051 | 288 | 242 | 84.0 |
| NFIA | 9 | 2692957 | 2937661 | - | 244704 | 2112 | 1721 | 81.5 |
| SOX2 | 10 | 18396452 | 18484266 | - | 87814 | 948 | 728 | 76.8 |
| CNOT2 | 4 | 36037862 | 36122470 | + | 84608 | 1611 | 1222 | 75.9 |
| UBE2D3 | 5 | 48896664 | 48919750 | + | 23086 | 494 | 372 | 75.3 |
| UBE2D2 | 14 | 15056594 | 15081979 | + | 25385 | 444 | 333 | 75.0 |
| GJD2 | 6 | 35675902 | 35680651 | + | 4749 | 915 | 685 | 74.9 |
| KDM6A | 2 | 1.13E+08 | 1.13E+08 | + | 150459 | 4416 | 3293 | 74.6 |
| FOXP2 | 4 | 25889103 | 26292811 | - | 403708 | 2202 | 1621 | 73.6 |
| AMD1 | 3 | 69717400 | 69735354 | - | 17954 | 1026 | 755 | 73.6 |
| FGF12 | 10 | 14298120 | 14515264 | + | 217144 | 833 | 612 | 73.5 |
| MARCHF4 | 7 | 14264113 | 14359677 | - | 95564 | 1173 | 850 | 72.5 |
| HMGN3 | 3 | 83390205 | 83414146 | + | 23941 | 384 | 278 | 72.4 |
| POU2F1 | 2 | 4578587 | 4681092 | - | 102505 | 2630 | 1899 | 72.2 |
| UBE2N | 4 | 46253511 | 46273858 | - | 20347 | 459 | 331 | 72.1 |
| LMO4 | 9 | 15037963 | 15053202 | + | 15239 | 498 | 359 | 72.1 |
| MOSMO | 15 | 9113188 | 9144197 | + | 31009 | 504 | 363 | 72.0 |
| NR2F2 | 13 | 15687702 | 15703513 | + | 15811 | 1355 | 972 | 71.7 |
| TCF7L2 | 8 | 27339850 | 27513391 | + | 173541 | 2234 | 1602 | 71.7 |
| TRIB2 | 3 | 1.03E+08 | 1.03E+08 | + | 19999 | 1032 | 736 | 71.3 |
| LHFPL3 | 4 | 12615089 | 12859349 | + | 244260 | 752 | 534 | 71.0 |
| HOXD10 | 7 | 21864399 | 21867442 | + | 3043 | 1034 | 734 | 71.0 |
| RAB6A | 2 | 1062742 | 1105266 | + | 42524 | 733 | 520 | 70.9 |
| HOXA7 | 1 | 1.22E+08 | 1.22E+08 | + | 3423 | 660 | 466 | 70.6 |
| USP22 | 15 | 13645229 | 13736892 | + | 91663 | 1578 | 1113 | 70.5 |
| ZBTB18 | 3 | 33832044 | 33838694 | + | 6650 | 3165 | 2232 | 70.5 |

| | | | | | | | | |
|---|---|---|---|---|---|---|---|---|
| **UBE2G1** | 19 | 7293527 | 7318497 | - | 24970 | 513 | 361 | 70.4 |
| **CSNK1A1** | 14 | 1594299 | 1626748 | + | 32449 | 1365 | 959 | 70.3 |
| **ELOC** | 1 | 32226747 | 32240793 | + | 14046 | 339 | 237 | 69.9 |
| **CCNC** | 3 | 74990868 | 75007126 | + | 16258 | 891 | 622 | 69.8 |
| **LHX9** | 9 | 2.23E+07 | 2.23E+07 | - | 19700 | 1721 | 1201 | 69.8 |
| **XKR6** | 3 | 114310763 | 114479066 | + | 168303 | 1851 | 1290 | 69.7 |
| **UBE2L3** | 17 | 8182192 | 8199804 | - | 17612 | 507 | 352 | 69.4 |
| **CXXC4** | 5 | 35806393 | 35967416 | - | 161023 | 1324 | 919 | 69.4 |
| **YWHAE** | 19 | 4976934 | 4997696 | - | 20762 | 776 | 538 | 69.3 |
| **MAF** | 11 | 5524146 | 5586702 | + | 62556 | 1113 | 769 | 69.1 |
| **NUTF2** | 11 | 6807941 | 6830320 | + | 22379 | 495 | 342 | 69.1 |
| **OAZ1** | 26 | 3642018 | 3645915 | - | 3897 | 651 | 447 | 68.7 |
| **RAP2B** | 10 | 25254870 | 25255666 | - | 796 | 552 | 378 | 68.5 |
| **PPP2R5E** | 6 | 6663369 | 6734427 | - | 71058 | 1530 | 1043 | 68.2 |
| **CELF2** | 4 | 5085531 | 5460683 | + | 375152 | 2257 | 1533 | 67.9 |
| **CACNG2** | 4 | 54100575 | 54152607 | + | 52032 | 1114 | 755 | 67.8 |
| **ELAVL4** | 9 | 6621993 | 6702986 | - | 80993 | 1645 | 1114 | 67.7 |
| **CDK8** | 2 | 4.72E+07 | 4.73E+07 | + | 74200 | 1678 | 1132 | 67.5 |
| **PTCHD1** | 2 | 105434425 | 105471431 | + | 37006 | 2667 | 1799 | 67.5 |
| **CXXC5** | 14 | 1.51E+07 | 1.52E+07 | + | 36588 | 831 | 559 | 67.3 |
| **RBMS3** | 1 | 115196684 | 115902929 | - | 706245 | 2150 | 1445 | 67.2 |
| **ZNF706** | 1 | 21093172 | 21100268 | + | 7096 | 231 | 155 | 67.1 |
| **FGF14** | 2 | 80195381 | 80588549 | - | 393168 | 995 | 667 | 67.0 |
| **QKI** | 3 | 3.92E+07 | 3.94E+07 | + | 152427 | 1695 | 1136 | 67.0 |
| **SATB1** | 1 | 119612547 | 119705454 | + | 92907 | 2358 | 1579 | 67.0 |
| **PLAG1** | 1 | 3.92E+07 | 3.93E+07 | + | 53175 | 2903 | 1943 | 66.9 |
| **TMEM196** | 1 | 125368294 | 125387666 | + | 19372 | 927 | 620 | 66.9 |
| **MYT1L** | 3 | 98124354 | 98443495 | - | 319141 | 4721 | 3149 | 66.7 |
| **FGF7** | 13 | 10019075 | 10045245 | + | 26170 | 820 | 545 | 66.5 |
| **FGF10** | Z | 15616301 | 15676178 | + | 59877 | 639 | 424 | 66.4 |

| | | | | | | | | |
|---|---|---|---|---|---|---|---|---|
| **PTEN** | 8 | 3591945 | 3640725 | - | 48780 | 1229 | 811 | 66.0 |
| **SPOPL** | 7 | 5466612 | 5499770 | - | 33158 | 1179 | 777 | 65.9 |
| **PURA** | 14 | 15545390 | 15546352 | + | 962 | 951 | 626 | 65.8 |
| **PBX1** | 9 | 28591098 | 28717104 | - | 126006 | 1340 | 880 | 65.7 |
| **GSK3B** | 2 | 26324625 | 26474255 | - | 149630 | 1302 | 854 | 65.6 |
| **LMO3** | 4 | 72850450 | 72909594 | + | 59144 | 677 | 444 | 65.6 |
| **CYRIB** | 1 | 8926375 | 9023591 | + | 97216 | 1097 | 714 | 65.1 |
| **AKIRIN2** | 3 | 80184991 | 80201419 | + | 16428 | 582 | 378 | 64.9 |
| **ZEB2** | 7 | 3344325 | 3459500 | + | 115175 | 3645 | 2360 | 64.7 |
| **AMMECR1L** | 10 | 3876509 | 3891460 | + | 14951 | 933 | 602 | 64.5 |
| **RHOQ** | 3 | 28538252 | 28554734 | + | 16482 | 910 | 586 | 64.4 |
| **RBFOX2** | 4 | 54300853 | 54472392 | + | 171539 | 1655 | 1065 | 64.4 |
| **RUNX1T1** | 1 | 2.49E+07 | 2.50E+07 | + | 114009 | 1965 | 1264 | 64.3 |
| **UBE2E1** | 1 | 117688052 | 117733245 | - | 45193 | 1173 | 752 | 64.1 |
| **COPS2** | 13 | 9916997 | 9938782 | - | 21785 | 1443 | 923 | 64.0 |
| **PPP3CB** | 8 | 15872014 | 15918470 | + | 46456 | 1879 | 1200 | 63.9 |
| **BRSK2** | 6 | 47008531 | 47318605 | + | 310074 | 3035 | 1937 | 63.8 |
| **HNRNPR** | 25 | 940564 | 960997 | + | 20433 | 2393 | 1526 | 63.8 |
| **YPEL1** | 17 | 7951831 | 7973817 | - | 21986 | 360 | 229 | 63.6 |
| **PAX2** | 8 | 16860524 | 16917636 | - | 57112 | 1170 | 743 | 63.5 |
| **LDB2** | 5 | 14892081 | 15266783 | - | 374702 | 1833 | 1160 | 63.3 |
| **MEIS1** | 3 | 3093446 | 3202736 | + | 109290 | 1604 | 1015 | 63.3 |
| **RAB12** | 1 | 5.19E+07 | 5.20E+07 | + | 23950 | 753 | 476 | 63.2 |
| **CTDSPL** | 1 | 152146633 | 152212327 | - | 65694 | 486 | 307 | 63.2 |
| **HDGFL3** | 13 | 10744841 | 10782483 | - | 37642 | 612 | 386 | 63.1 |
| **MAFK** | 15 | 11391344 | 11405724 | + | 14380 | 471 | 297 | 63.1 |
| **ZIC1** | 10 | 12735229 | 12739550 | + | 4321 | 1338 | 841 | 62.9 |
| **MEIS2** | 6 | 34504387 | 34679190 | + | 174803 | 1407 | 882 | 62.7 |
| **FOXP1** | 12 | 16752338 | 17133161 | - | 380823 | 2351 | 1473 | 62.7 |
| **C1H6orf62** | 1 | 60787498 | 60795214 | - | 7716 | 690 | 432 | 62.6 |

| | | | | | | | | |
|---|---|---|---|---|---|---|---|---|
| **MBNL2** | 2 | 78039887 | 78147165 | + | 107278 | 1334 | 834 | 62.5 |
| **SNAP25** | 3 | 6648711 | 6710223 | - | 61512 | 621 | 388 | 62.5 |
| **GRIA4** | 2 | 4.26E+07 | 4.28E+07 | + | 221338 | 3259 | 2035 | 62.4 |
| **TFAP2B** | 3 | 113055613 | 113089530 | + | 33917 | 1711 | 1068 | 62.4 |
| **GPR85** | 4 | 26664618 | 26669132 | + | 4514 | 1113 | 693 | 62.3 |
| **PHLDA2** | 6 | 48666753 | 48668266 | - | 1513 | 399 | 248 | 62.2 |
| **CSNK2A2** | 11 | 6258118 | 6286909 | + | 28791 | 1053 | 653 | 62.0 |
| **MXD4** | 5 | 7946730 | 7987861 | - | 41131 | 884 | 547 | 61.9 |
| **TRPS1** | 1 | 14825916 | 15035345 | + | 209429 | 4684 | 2898 | 61.9 |
| **EIF5** | 6 | 10104065 | 10111639 | - | 7574 | 1293 | 799 | 61.8 |
| **YTHDF3** | 1 | 36523306 | 36546688 | - | 23382 | 3327 | 2053 | 61.7 |
| **RAP1B** | 4 | 35500432 | 35532192 | + | 31760 | 555 | 342 | 61.6 |
| **ERI3** | 9 | 1.01E+07 | 1.02E+07 | + | 131207 | 816 | 501 | 61.4 |
| **HOXA13** | 1 | 122346283 | 122349051 | + | 2768 | 918 | 563 | 61.3 |
| **TIAL1** | 8 | 30684506 | 30705020 | - | 20514 | 1268 | 777 | 61.3 |
| **SIX6** | 6 | 5538951 | 5541086 | + | 2135 | 741 | 454 | 61.3 |
| **COPS3** | 15 | 13849170 | 13862796 | - | 13626 | 1323 | 810 | 61.2 |
| **SUMO1** | 7 | 26551612 | 26563972 | + | 12360 | 345 | 211 | 61.2 |
| **JARID2** | 1 | 91469937 | 91680696 | - | 210759 | 3862 | 2359 | 61.1 |
| **GRIA2** | 5 | 42856280 | 42943070 | + | 86790 | 3557 | 2167 | 60.9 |
| **CALM1** | 6 | 16973890 | 16985007 | - | 11117 | 450 | 274 | 60.9 |
| **CALM2** | 3 | 28995618 | 29009091 | - | 13473 | 450 | 274 | 60.9 |
| **CELF4** | Z | 23019373 | 23734598 | - | 715225 | 1981 | 1205 | 60.8 |
| **KCNMA1** | 8 | 13076497 | 13511360 | + | 434863 | 4967 | 3021 | 60.8 |
| **CACUL1** | 8 | 30232196 | 30274974 | - | 42778 | 1005 | 610 | 60.7 |
| **EBF3** | 8 | 3.48E+07 | 3.49E+07 | - | 122625 | 1854 | 1117 | 60.2 |
| **GPM6B** | 2 | 100919593 | 101011665 | - | 92072 | 1244 | 748 | 60.1 |
| **ACTB** | 15 | 1.32E+07 | 1.32E+07 | - | 4839 | 1128 | 677 | 60.0 |
| **TMSB4X** | 2 | 100515905 | 100517440 | + | 1535 | 135 | 81 | 60.0 |
| **EIF4E** | 5 | 49946872 | 49971271 | + | 24399 | 654 | 392 | 59.9 |

| | | | | | | | | |
|---|---|---|---|---|---|---|---|---|
| **SLC6A6** | 12 | 11098783 | 11150200 | + | 51417 | 2348 | 1406 | 59.9 |
| **TMEM211** | 19 | 3223942 | 3282917 | + | 58975 | 654 | 391 | 59.8 |
| **RBFOX1** | 15 | 3232387 | 4383962 | + | 1151575 | 2125 | 1266 | 59.6 |
| **NOVA1** | 6 | 37217122 | 37361386 | + | 144264 | 1524 | 907 | 59.5 |
| **TAOK1** | 19 | 4.00E+06 | 4.07E+06 | - | 66845 | 3006 | 1789 | 59.5 |
| **PTCHD4** | 3 | 110914722 | 110995645 | + | 80923 | 3141 | 1867 | 59.4 |
| **PHF6** | Z | 84394549 | 84421116 | + | 26567 | 1262 | 749 | 59.4 |
| **SAMD13** | 9 | 14126449 | 14138055 | + | 11606 | 309 | 183 | 59.2 |
| **LEF1** | 5 | 37059532 | 37129982 | - | 70450 | 1310 | 775 | 59.2 |
| **ATXN7L1** | 4 | 13166288 | 13278247 | - | 111959 | 2754 | 1627 | 59.1 |
| **TWIST2** | 7 | 35422777 | 35449339 | - | 26562 | 483 | 285 | 59.0 |
| **SIX3** | 3 | 27603583 | 27607513 | + | 3930 | 951 | 561 | 59.0 |
| **TLK1** | 7 | 19892769 | 19960106 | - | 67337 | 2415 | 1424 | 59.0 |
| **ESRRG** | 3 | 21314675 | 21687282 | + | 372607 | 2061 | 1215 | 59.0 |
| **C15H16orf72** | 15 | 5388789 | 5405607 | + | 16818 | 822 | 484 | 58.9 |
| **CPEB3** | 8 | 20539803 | 20619778 | + | 79975 | 3173 | 1864 | 58.7 |
| **CXCL12** | 8 | 1.88E+07 | 1.88E+07 | - | 18228 | 412 | 242 | 58.7 |
| **POU6F2** | 1 | 101426073 | 101738362 | + | 312289 | 2076 | 1216 | 58.6 |
| **PBX3** | 20 | 593351 | 696857 | - | 103506 | 1424 | 834 | 58.6 |
| **HAND2** | 5 | 48074331 | 48076611 | - | 2280 | 654 | 383 | 58.6 |
| **EIF4G2** | 6 | 5.94E+07 | 5.94E+07 | + | 11951 | 2868 | 1679 | 58.5 |
| **WAC** | 1 | 140712412 | 140738364 | + | 25952 | 1361 | 795 | 58.4 |
| **ZNF384** | 2 | 28505778 | 28528340 | - | 22562 | 1671 | 974 | 58.3 |
| **PFN2** | 10 | 2.57E+07 | 2.57E+07 | + | 5342 | 521 | 303 | 58.2 |
| **HOXA9** | 1 | 122384117 | 122387221 | + | 3104 | 786 | 457 | 58.1 |
| **GNAS** | 16 | 3.41E+06 | 3.54E+06 | + | 131093 | 1429 | 829 | 58.0 |
| **NXPH1** | 1 | 130117260 | 130256787 | - | 139527 | 816 | 473 | 58.0 |
| **CNOT4** | 4 | 49627896 | 49706308 | - | 78412 | 2699 | 1563 | 57.9 |
| **TMEM170B** | 1 | 89671091 | 89684767 | + | 13676 | 399 | 231 | 57.9 |
| **CSNK2A1** | 16 | 7266508 | 7289154 | + | 22646 | 1176 | 680 | 57.8 |

| | | | | | | | | | |
|---|---|---|---|---|---|---|---|---|---|
| **FBRSL1** | 17 | 6318565 | 6822354 | + | | 503789 | 4839 | 2797 | 57.8 |
| **ZNRF1** | 11 | 7959806 | 7979996 | + | | 20190 | 705 | 407 | 57.7 |
| **PHF21A** | 6 | 39042140 | 39176524 | - | | 134384 | 2150 | 1241 | 57.7 |
| **ATOH8** | 5 | 5408840 | 5433202 | - | | 24362 | 618 | 356 | 57.6 |
| **IKZF2** | 7 | 37005874 | 37121282 | - | | 115408 | 1658 | 955 | 57.6 |
| **CCND1** | 6 | 61180511 | 61204407 | - | | 23896 | 897 | 516 | 57.5 |
| **SF3B5** | 3 | 50519706 | 50522031 | + | | 2325 | 261 | 150 | 57.5 |
| **PAX6** | 6 | 55386409 | 55407723 | - | | 21314 | 1539 | 884 | 57.4 |
| **YWHAG** | 19 | 6022850 | 6043890 | - | | 21040 | 744 | 427 | 57.4 |
| **LRRTM3** | 8 | 1126743 | 1206494 | + | | 79751 | 3333 | 1911 | 57.3 |
| **INO80D** | 7 | 25636807 | 25682837 | - | | 46030 | 3090 | 1771 | 57.3 |
| **NAA30** | 6 | 4323288 | 4344284 | + | | 20996 | 1239 | 709 | 57.2 |
| **PDE6D** | 10 | 16616442 | 16650828 | - | | 34386 | 453 | 259 | 57.2 |
| **WDR20** | 6 | 10857724 | 10903466 | - | | 45742 | 1911 | 1092 | 57.1 |
| **EMX2** | 8 | 29698778 | 29704916 | + | | 6138 | 744 | 425 | 57.1 |
| **CBFB** | 11 | 8482221 | 8519991 | + | | 37770 | 714 | 407 | 57.0 |
| **CACNG3** | 15 | 15767670 | 15791542 | + | | 23872 | 1337 | 761 | 56.9 |
| **LINGO1** | 13 | 1478621 | 1617609 | - | | 138988 | 3729 | 2120 | 56.9 |
| **PDGFA** | 15 | 1.07E+07 | 1.07E+07 | - | | 36288 | 781 | 444 | 56.9 |
| **LRATD1** | 3 | 104301310 | 104303211 | + | | 1901 | 855 | 486 | 56.8 |
| **PROX1** | 3 | 22653286 | 22703112 | - | | 49826 | 2400 | 1364 | 56.8 |
| **ATXN7** | 12 | 14053628 | 14121828 | + | | 68200 | 2677 | 1521 | 56.8 |
| **MAP2K4** | 18 | 1647624 | 1712591 | - | | 64967 | 1269 | 720 | 56.7 |
| **TAFA1** | 12 | 15667223 | 15890843 | + | | 223620 | 402 | 228 | 56.7 |
| **BCL7A** | 17 | 3107470 | 3127432 | + | | 19962 | 633 | 359 | 56.7 |
| **SIX2** | 3 | 27668945 | 27673400 | - | | 4455 | 864 | 490 | 56.7 |
| **TNRC6C** | 18 | 3449883 | 3547994 | + | | 98111 | 6368 | 3609 | 56.7 |
| **RBMS1** | 7 | 16257666 | 16400182 | - | | 142516 | 1540 | 871 | 56.6 |
| **UBL3** | 2 | 48568361 | 48622698 | - | | 54337 | 354 | 200 | 56.5 |
| **GID8** | 16 | 5489358 | 5496778 | + | | 7420 | 687 | 388 | 56.5 |

| | | | | | | | | |
|---|---|---|---|---|---|---|---|---|
| **FGF13** | W | 28534725 | 28896347 | + | 361622 | 955 | 539 | 56.4 |
| **ARIH2** | 12 | 11923393 | 11958666 | - | 35273 | 1473 | 830 | 56.3 |
| **TPM1** | 13 | 3685993 | 3705908 | - | 19915 | 1426 | 802 | 56.2 |
| **RAB2A** | 1 | 37577304 | 37620080 | - | 42776 | 639 | 359 | 56.2 |
| **PCMT1** | 3 | 5.81E+07 | 5.81E+07 | - | 50788 | 940 | 528 | 56.2 |
| **ABI1** | 1 | 139922640 | 140001230 | - | 78590 | 1521 | 854 | 56.1 |
| **LGI1** | 8 | 19999221 | 20028733 | - | 29512 | 1658 | 930 | 56.1 |
| **ATXN2** | 17 | 3548864 | 3598436 | - | 49572 | 3806 | 2134 | 56.1 |
| **EBF1** | 14 | 3896619 | 4162966 | + | 266347 | 2040 | 1143 | 56.0 |
| **PAPOLA** | 6 | 14063180 | 14108479 | - | 45299 | 2381 | 1333 | 56.0 |
| **ZFAND3** | 3 | 31682187 | 31817158 | - | 134971 | 739 | 413 | 55.9 |
| **PKIA** | 1 | 29972708 | 30017046 | - | 44338 | 231 | 129 | 55.8 |
| **POU3F3** | 2 | 90105036 | 90106371 | + | 1335 | 1323 | 738 | 55.8 |
| **MBNL3** | W | 5105655 | 5231697 | - | 126042 | 1162 | 648 | 55.8 |
| **CFL2** | 6 | 25796132 | 25801812 | + | 5680 | 501 | 279 | 55.7 |
| **BHLHE22** | 1 | 36007133 | 36008034 | - | 901 | 900 | 501 | 55.7 |
| **CTNND2** | 1 | 73667315 | 74309029 | - | 641714 | 4077 | 2267 | 55.6 |
| **BTG1** | 4 | 45765041 | 45767873 | - | 2832 | 513 | 285 | 55.6 |
| **SGCD** | 14 | 4744922 | 5060488 | - | 315566 | 924 | 513 | 55.5 |
| **PHF12** | 19 | 4179233 | 4210742 | + | 31509 | 3030 | 1681 | 55.5 |
| **CHD7** | 1 | 37425703 | 37558908 | - | 133205 | 9335 | 5176 | 55.4 |
| **NFIB** | Z | 32240896 | 32392771 | - | 151875 | 2451 | 1358 | 55.4 |
| **VKORC1L1** | 19 | 5.27E+06 | 5.29E+06 | - | 19522 | 531 | 294 | 55.4 |
| **SEPTIN7** | 1 | 108806546 | 108887445 | - | 80899 | 1290 | 714 | 55.3 |
| **CRYBB2** | 17 | 4511001 | 4513714 | + | 2713 | 850 | 470 | 55.3 |
| **BMP4** | 6 | 3028195 | 3031297 | - | 3102 | 1212 | 670 | 55.3 |
| **FAM155A** | 2 | 82768203 | 83217028 | - | 448825 | 1287 | 711 | 55.2 |
| **CBLN2** | 1 | 58332604 | 58337598 | - | 4994 | 639 | 353 | 55.2 |
| **PITX1** | 14 | 9335995 | 9342664 | + | 6669 | 909 | 501 | 55.1 |
| **TPT1** | 2 | 55992797 | 55997941 | + | 5144 | 519 | 286 | 55.1 |

| | | | | | | | | | |
|---|---|---|---|---|---|---|---|---|---|
| **MSI1** | 17 | 10455059 | 10485402 | - | | 30343 | 1170 | 644 | 55.0 |
| **RAP2C** | W | 5024533 | 5034298 | - | | 9765 | 552 | 303 | 54.9 |
| **HOXB5** | 27 | 5091002 | 5093053 | + | | 2051 | 798 | 438 | 54.9 |
| **NPAS3** | 6 | 26097760 | 26694190 | - | | 596430 | 2849 | 1562 | 54.8 |
| **EFNB2** | 2 | 82464578 | 82510134 | - | | 45556 | 1002 | 549 | 54.8 |
| **FAM120A** | 12 | 439649 | 486370 | + | | 46721 | 3449 | 1889 | 54.8 |
| **CNOT6** | 14 | 7382440 | 7415325 | - | | 32885 | 1764 | 965 | 54.7 |
| **BCL11B** | 6 | 12658130 | 12749907 | + | | 91777 | 3021 | 1651 | 54.7 |
| **SPTSSA** | 6 | 25863945 | 25868825 | + | | 4880 | 207 | 113 | 54.6 |
| **ARIH1** | 13 | 2620973 | 2671071 | - | | 50098 | 1935 | 1056 | 54.6 |
| **SOX18** | 16 | 6583283 | 6587237 | + | | 3954 | 1254 | 683 | 54.5 |
| **VEZF1** | 19 | 1113495 | 1126918 | - | | 13423 | 1674 | 910 | 54.4 |
| **KIAA0232** | 5 | 11125037 | 11190859 | - | | 65822 | 4176 | 2270 | 54.4 |
| **TAFA2** | 4 | 32877891 | 33069515 | - | | 191624 | 396 | 215 | 54.3 |
| **THRA** | 27 | 3463037 | 3470847 | - | | 7810 | 1227 | 666 | 54.3 |
| **FGF8** | 8 | 22654828 | 22663721 | + | | 8893 | 815 | 442 | 54.2 |
| **PSMD14** | 7 | 1.66E+07 | 1.67E+07 | + | | 46470 | 933 | 505 | 54.1 |
| **HOXA4** | 1 | 122420460 | 122422834 | + | | 2374 | 909 | 492 | 54.1 |
| **TLNRD1** | 13 | 11763402 | 11765462 | - | | 2060 | 1068 | 578 | 54.1 |
| **SET** | 20 | 5.59E+06 | 5.60E+06 | - | | 8180 | 880 | 476 | 54.1 |
| **YWHAQ** | 3 | 101633523 | 101656084 | - | | 22561 | 738 | 399 | 54.1 |
| **RASD1** | 15 | 13902010 | 13903985 | - | | 1975 | 831 | 449 | 54.0 |
| **ASF1A** | 3 | 66102983 | 66113750 | - | | 10767 | 615 | 332 | 54.0 |
| **CAMK2G** | 8 | 15308603 | 15428031 | + | | 119428 | 1841 | 993 | 53.9 |
| **YWHAZ** | 1 | 2.12E+07 | 2.12E+07 | + | | 26041 | 738 | 398 | 53.9 |
| **AP1S2** | 2 | 101961912 | 101992868 | - | | 30956 | 564 | 304 | 53.9 |
| **ISL1** | Z | 14482062 | 14493337 | - | | 11275 | 1334 | 718 | 53.8 |
| **UBE3A** | 2 | 86971945 | 87024720 | + | | 52775 | 2645 | 1423 | 53.8 |
| **ZNF326** | 9 | 15836752 | 15859530 | + | | 22778 | 1692 | 910 | 53.8 |
| **SERP1** | 10 | 25563246 | 25565156 | + | | 1910 | 201 | 108 | 53.7 |

| | | | | | | | | |
|---|---|---|---|---|---|---|---|---|
| **AKT3** | 3 | 33601943 | 33755956 | - | 154013 | 1541 | 827 | 53.7 |
| **EIF4E2** | 10 | 16927486 | 16946690 | - | 19204 | 751 | 403 | 53.7 |
| **CDK5R2** | 7 | 15481140 | 15482526 | + | 1386 | 960 | 515 | 53.6 |
| **GPR27** | 12 | 17192317 | 17194535 | + | 2218 | 1143 | 613 | 53.6 |
| **MCRIP1** | 18 | 11780409 | 11782253 | - | 1844 | 291 | 156 | 53.6 |
| **SST** | 10 | 1.59E+07 | 1.59E+07 | + | 1555 | 351 | 188 | 53.6 |
| **ETV1** | 1 | 127905574 | 127971796 | + | 66222 | 1572 | 841 | 53.5 |
| **FBXO45** | 10 | 4650597 | 4655757 | + | 5160 | 819 | 438 | 53.5 |
| **ZNF148** | 7 | 9279356 | 9317060 | + | 37704 | 2370 | 1265 | 53.4 |
| **HAS2** | 1 | 12218128 | 12237662 | + | 19534 | 1662 | 887 | 53.4 |
| **PPP2R5C** | 6 | 10991327 | 11068563 | - | 77236 | 2243 | 1197 | 53.4 |
| **UBE2K** | 5 | 22872226 | 22916944 | + | 44718 | 666 | 355 | 53.3 |
| **SRSF1** | 19 | 1137703 | 1141215 | - | 3512 | 747 | 398 | 53.3 |
| **SOCS6** | 1 | 57335895 | 57360863 | + | 24968 | 1605 | 855 | 53.3 |
| **SSBP3** | 9 | 4895113 | 4950763 | + | 55650 | 1167 | 621 | 53.2 |
| **HOXD8** | 7 | 21877147 | 21879115 | + | 1968 | 1404 | 746 | 53.1 |
| **HMGB1** | 2 | 48823835 | 48832592 | - | 8757 | 648 | 344 | 53.1 |
| **MAPRE2** | 1 | 42961763 | 43061218 | - | 99455 | 1187 | 630 | 53.1 |
| **LRRC1** | 3 | 92562446 | 92642482 | - | 80036 | 1575 | 835 | 53.0 |
| **TBL1X** | 2 | 98609295 | 98748896 | + | 139601 | 1572 | 833 | 53.0 |
| **UBE2QL1** | 1 | 71789811 | 71807902 | + | 18091 | 921 | 488 | 53.0 |
| **STMN2** | 1 | 29576326 | 29618188 | - | 41862 | 540 | 286 | 53.0 |
| **UHMK1** | 9 | 26967115 | 26981777 | - | 14662 | 1284 | 680 | 53.0 |
| **KCNK9** | 1 | 4155990 | 4244525 | + | 88535 | 1425 | 754 | 52.9 |
| **EPHA7** | 3 | 77603723 | 77770686 | + | 166963 | 4471 | 2361 | 52.8 |
| **IER5L** | 20 | 5329716 | 5331210 | + | 1494 | 966 | 510 | 52.8 |
| **CNNM2** | 8 | 23509459 | 23622954 | + | 113495 | 1836 | 969 | 52.8 |
| **CEBPA** | 11 | 20213662 | 20216548 | - | 2886 | 978 | 516 | 52.8 |
| **RAB7A** | 12 | 8946439 | 8968479 | - | 22040 | 624 | 329 | 52.7 |
| **OAZ2** | 13 | 3257476 | 3270019 | - | 12543 | 573 | 302 | 52.7 |

| | | | | | | | | |
|---|---|---|---|---|---|---|---|---|
| **TBL1XR1** | 10 | 19672996 | 19781615 | + | 108619 | 1542 | 812 | 52.7 |
| **FNDC5** | 25 | 3414957 | 3430042 | - | 15085 | 847 | 446 | 52.7 |
| **ARF6** | 6 | 1919183 | 1923055 | + | 3872 | 528 | 278 | 52.7 |
| **MSI2** | 19 | 844647 | 1052710 | + | 208063 | 1691 | 890 | 52.6 |
| **UBE2W** | 1 | 32250073 | 32282181 | + | 32108 | 462 | 243 | 52.6 |
| **MAPK8** | 8 | 17931309 | 18076014 | + | 144705 | 1373 | 722 | 52.6 |
| **OTX2** | 6 | 4049693 | 4053752 | - | 4059 | 1043 | 548 | 52.5 |
| **VEGFA** | 3 | 32831568 | 32854853 | - | 23285 | 773 | 406 | 52.5 |
| **KICS2** | 4 | 33807907 | 33817479 | - | 9572 | 1552 | 815 | 52.5 |
| **RAB1A** | 3 | 2548630 | 2562126 | - | 13496 | 981 | 515 | 52.5 |
| **PRKACB** | 9 | 14048610 | 14116310 | + | 67700 | 1322 | 694 | 52.5 |
| **SUB1** | Z | 1.99E+07 | 1.99E+07 | - | 12957 | 381 | 200 | 52.5 |
| **ZMYND11** | 1 | 146410221 | 146515049 | - | 104828 | 1854 | 972 | 52.4 |
| **CDC42** | 22 | 2296099 | 2323239 | - | 27140 | 666 | 349 | 52.4 |
| **ARID2** | 4 | 30834592 | 30929427 | + | 94835 | 5496 | 2880 | 52.4 |
| **GNAL** | 1 | 53339841 | 53519094 | + | 179253 | 1630 | 854 | 52.4 |
| **GULP1** | 7 | 31204823 | 31373162 | - | 168339 | 1173 | 614 | 52.3 |
| **KCTD15** | 11 | 20694871 | 20739079 | + | 44208 | 968 | 506 | 52.3 |
| **DESI2** | 3 | 34012029 | 34026850 | + | 14821 | 582 | 304 | 52.2 |
| **UBE2Q2** | 13 | 711991 | 757546 | + | 45555 | 1221 | 637 | 52.2 |
| **MSANTD1** | 5 | 8.78E+06 | 8.83E+06 | + | 43638 | 1235 | 644 | 52.1 |
| **CASK** | 2 | 111927719 | 112118466 | - | 190747 | 2850 | 1485 | 52.1 |
| **ARPC2** | 7 | 15205950 | 15225791 | + | 19841 | 960 | 499 | 52.0 |
| **RORA** | 13 | 4.28E+06 | 4.54E+06 | + | 261678 | 1706 | 886 | 51.9 |
| **GLRA2** | 2 | 101303990 | 101430299 | + | 126309 | 1507 | 781 | 51.8 |
| **ADGRB3** | 3 | 87362886 | 87828744 | - | 465858 | 4707 | 2437 | 51.8 |
| **YAF2** | 4 | 2.95E+07 | 2.95E+07 | - | 24684 | 543 | 281 | 51.7 |
| **CUL2** | 1 | 143161980 | 143203469 | - | 41489 | 2238 | 1158 | 51.7 |
| **NKX2-1** | 6 | 25028226 | 25030187 | + | 1961 | 1038 | 537 | 51.7 |
| **TEAD1** | 6 | 57679607 | 57833779 | + | 154172 | 1305 | 675 | 51.7 |

| | | | | | | | | |
|---|---|---|---|---|---|---|---|---|
| **TOMM20** | 3 | 4.56E+07 | 4.56E+07 | - | 9357 | 441 | 228 | 51.7 |
| **HOXA3** | 1 | 122426251 | 122446095 | + | 19844 | 1242 | 642 | 51.7 |
| **HOXD4** | 7 | 21894191 | 21896485 | + | 2294 | 714 | 369 | 51.7 |
| **TBR1** | 7 | 16655748 | 16661953 | + | 6205 | 2034 | 1051 | 51.7 |
| **MAB21L2** | 5 | 59173501 | 59176147 | + | 2646 | 1080 | 558 | 51.7 |
| **DDX6** | 23 | 6999766 | 7020934 | + | 21168 | 1452 | 750 | 51.7 |
| **GPBP1L1** | 9 | 9.31E+06 | 9.34E+06 | + | 32100 | 1416 | 731 | 51.6 |
| **IGF2BP3** | 1 | 123917269 | 124027671 | + | 110402 | 1958 | 1007 | 51.4 |
| **HMGB3** | 21 | 7098181 | 7104576 | + | 6395 | 609 | 313 | 51.4 |
| **TOX** | 1 | 38067625 | 38286089 | + | 218464 | 1578 | 811 | 51.4 |
| **RFK** | Z | 35501516 | 35510037 | - | 8521 | 468 | 240 | 51.3 |
| **RFX3** | Z | 2677050 | 2777360 | + | 100310 | 2250 | 1153 | 51.2 |
| **FUBP3** | 20 | 4817760 | 4856892 | - | 39132 | 1847 | 946 | 51.2 |
| **HOXB8** | 27 | 5.07E+06 | 5.07E+06 | + | 1619 | 1034 | 529 | 51.2 |
| **FKBP1B** | 3 | 110115241 | 110161017 | + | 45776 | 327 | 167 | 51.1 |
| **TRAF3** | 6 | 10487903 | 10557852 | - | 69949 | 1704 | 870 | 51.1 |
| **LDB1** | 8 | 22393781 | 22418371 | + | 24590 | 1379 | 704 | 51.1 |
| **DACH1** | 2 | 66278987 | 66631894 | - | 352907 | 2312 | 1180 | 51.0 |
| **MAP4K4** | 2 | 8.87E+07 | 8.89E+07 | + | 161371 | 4662 | 2379 | 51.0 |
| **FZD1** | 1 | 133362178 | 133365977 | - | 3799 | 1758 | 897 | 51.0 |
| **TMEFF2** | 7 | 29856064 | 29982395 | + | 126331 | 1221 | 623 | 51.0 |
| **SCRT2** | 16 | 7244070 | 7247162 | + | 3092 | 831 | 424 | 51.0 |
| **CELF5** | 26 | 2582693 | 2619818 | + | 37125 | 1733 | 884 | 51.0 |
| **NIPBL** | Z | 1.82E+07 | 1.84E+07 | - | 140908 | 8568 | 4367 | 51.0 |
| **OST4** | 3 | 116475871 | 116476476 | + | 605 | 114 | 58 | 50.9 |
| **PPP2CA** | 14 | 9658948 | 9674149 | + | 15201 | 930 | 473 | 50.9 |
| **HNRNPAB** | 14 | 7873902 | 7899618 | - | 25716 | 1029 | 523 | 50.8 |
| **SPIN1** | Z | 42701541 | 42761825 | + | 60284 | 789 | 401 | 50.8 |
| **EFNA5** | Z | 4.85E+07 | 4.88E+07 | + | 214331 | 687 | 349 | 50.8 |
| **RARB** | 1 | 117006921 | 117327600 | - | 320679 | 2000 | 1016 | 50.8 |

| | | | | | | | | |
|---|---|---|---|---|---|---|---|---|
| **HNRNPH3** | 8 | 1514362 | 1521327 | + | 6965 | 1314 | 667 | 50.8 |
| **TBPL1** | 3 | 59409023 | 59423199 | - | 14176 | 552 | 280 | 50.7 |
| **CEBPB** | 16 | 7.68E+05 | 7.69E+05 | + | 1797 | 978 | 496 | 50.7 |
| **CBX3** | 1 | 122869397 | 122883985 | - | 14588 | 734 | 372 | 50.7 |
| **DAB1** | 9 | 3773607 | 4180190 | + | 406583 | 1737 | 880 | 50.7 |
| **TAB2** | 3 | 48231649 | 48294278 | - | 62629 | 3490 | 1767 | 50.6 |
| **FOXN3** | 6 | 17314932 | 17531563 | + | 216631 | 1557 | 788 | 50.6 |
| **ELAVL1** | 26 | 4718876 | 4760442 | + | 41566 | 1095 | 553 | 50.5 |
| **RAB14** | 20 | 2016710 | 2030936 | + | 14226 | 648 | 327 | 50.5 |
| **ACTG1** | 18 | 11016056 | 11019084 | + | 3028 | 1128 | 569 | 50.4 |
| **DPF3** | 6 | 29322067 | 29491277 | + | 169210 | 1442 | 725 | 50.3 |
| **AMMECR1** | 21 | 7600333 | 7651887 | + | 51554 | 1542 | 773 | 50.1 |
| **UBE2B** | 14 | 9620420 | 9632664 | - | 12244 | 459 | 230 | 50.1 |
| **CDK19** | 3 | 69764653 | 69886363 | + | 121710 | 1595 | 799 | 50.1 |
| **RNF7** | 10 | 11010595 | 11017107 | + | 6512 | 573 | 287 | 50.1 |
| **NLGN1** | 10 | 20532912 | 20955907 | - | 422995 | 2875 | 1440 | 50.1 |
| **YAP1** | 2 | 41290171 | 41384995 | + | 94824 | 1706 | 854 | 50.1 |
| **TMSB15B** | 21 | 3.89E+06 | 3.89E+06 | + | 1459 | 138 | 69 | 50.0 |
| **VSNL1** | 3 | 105893252 | 105979663 | + | 86411 | 576 | 288 | 50.0 |
| **BRD1** | 4 | 18218038 | 18286153 | - | 68115 | 5161 | 2576 | 49.9 |
| **TENT4B** | 11 | 12508580 | 12547946 | - | 39366 | 2058 | 1025 | 49.8 |
| **MTF2** | 9 | 16861833 | 16886543 | + | 24710 | 1864 | 928 | 49.8 |
| **RABIF** | 24 | 1338481 | 1339908 | - | 1427 | 432 | 215 | 49.8 |
| **PAN3** | 2 | 47930085 | 48010029 | + | 79944 | 2772 | 1379 | 49.7 |
| **DCLK1** | 2 | 50955544 | 51184895 | - | 229351 | 2512 | 1248 | 49.7 |
| **UBE2A** | W | 23662542 | 23671877 | - | 9335 | 459 | 228 | 49.7 |
| **FGF9** | 2 | 45559481 | 45587511 | + | 28030 | 733 | 364 | 49.7 |
| **DAPK3** | 26 | 615305 | 620121 | - | 4816 | 1371 | 680 | 49.6 |
| **C6H11orf96** | 6 | 4.16E+07 | 4.16E+07 | - | 860 | 363 | 180 | 49.6 |
| **KCNJ6** | 2 | 116535295 | 116694563 | + | 159268 | 2170 | 1076 | 49.6 |

| | | | | | | | | |
|---|---|---|---|---|---|---|---|---|
| **ARPP19** | 13 | 8188977 | 8200667 | + | 11690 | 339 | 168 | 49.6 |
| **GAP43** | 2 | 24018603 | 24076746 | + | 58143 | 747 | 370 | 49.5 |
| **RGS7** | 3 | 47755625 | 47954383 | - | 198758 | 1329 | 658 | 49.5 |
| **CUX1** | 19 | 6.22E+06 | 6.48E+06 | - | 267044 | 5843 | 2884 | 49.4 |
| **PIP4K2A** | 1 | 138178878 | 138284738 | - | 105860 | 1218 | 601 | 49.3 |
| **INPP5A** | 8 | 35687508 | 35869602 | + | 182094 | 1326 | 654 | 49.3 |
| **PGRMC2** | 5 | 67636487 | 67649176 | - | 12689 | 582 | 287 | 49.3 |
| **DICER1** | 6 | 14783119 | 14847103 | + | 63984 | 5766 | 2843 | 49.3 |
| **C6H14orf132** | 6 | 1.43E+07 | 1.44E+07 | - | 40901 | 282 | 139 | 49.3 |
| **CYRIA** | 3 | 105374129 | 105428239 | - | 54110 | 1096 | 540 | 49.3 |
| **OLFM1** | 20 | 2907092 | 2922297 | - | 15205 | 1257 | 619 | 49.2 |
| **TFDP1** | 2 | 92474959 | 92515276 | - | 40317 | 1383 | 681 | 49.2 |
| **MMP16** | 1 | 26249743 | 26419954 | + | 170211 | 1934 | 952 | 49.2 |
| **ING3** | 4 | 2.30E+07 | 2.30E+07 | - | 16593 | 1611 | 792 | 49.2 |
| **THRB** | 1 | 117435103 | 117592421 | + | 157318 | 1886 | 927 | 49.2 |
| **PXDC1** | 1 | 86515482 | 86539852 | - | 24370 | 696 | 342 | 49.1 |
| **DDX17** | 4 | 53256295 | 53276351 | + | 20056 | 2020 | 992 | 49.1 |
| **PPARGC1A** | 5 | 17284222 | 17652417 | - | 368195 | 2565 | 1259 | 49.1 |
| **PAFAH1B1** | 19 | 391382 | 417362 | - | 25980 | 1233 | 605 | 49.1 |
| **NFIC** | 26 | 4453151 | 4492250 | + | 39099 | 1590 | 780 | 49.1 |
| **PMEPA1** | 16 | 2900423 | 2940124 | - | 39701 | 1578 | 774 | 49.0 |
| **AKT1** | 6 | 8.53E+06 | 8.61E+06 | + | 77160 | 1544 | 757 | 49.0 |
| **RHEB** | 1 | 150270924 | 150309524 | + | 38600 | 818 | 401 | 49.0 |
| **ARID1B** | 3 | 54799932 | 55124950 | - | 325018 | 7268 | 3562 | 49.0 |
| **IRF2BP2** | 3 | 45384971 | 45387636 | - | 2665 | 1707 | 836 | 49.0 |
| **XPO4** | 2 | 45146824 | 45220906 | - | 74082 | 3510 | 1718 | 48.9 |
| **YTHDF1** | 16 | 5818232 | 5833220 | - | 14988 | 3261 | 1596 | 48.9 |
| **PPP3R1** | 3 | 3919244 | 3957069 | - | 37825 | 513 | 251 | 48.9 |
| **KCNIP2** | 8 | 22569404 | 22612570 | + | 43166 | 1202 | 588 | 48.9 |
| **UBE2V2** | 1 | 42549641 | 42577191 | - | 27550 | 595 | 291 | 48.9 |

| | | | | | | | | |
|---|---|---|---|---|---|---|---|---|
| **TAOK3** | 17 | 9887864 | 9962480 | - | 74616 | 2697 | 1319 | 48.9 |
| **CREB1** | 7 | 26087462 | 26127004 | + | 39542 | 1125 | 550 | 48.9 |
| **BARX1** | 12 | 661408 | 664565 | - | 3157 | 741 | 362 | 48.9 |
| **PPP1R14C** | 3 | 57974369 | 58026240 | - | 51871 | 477 | 233 | 48.8 |
| **CTXN2** | 13 | 9574525 | 9579695 | + | 5170 | 246 | 120 | 48.8 |
| **ARGLU1** | 2 | 82516031 | 82526652 | - | 10621 | 831 | 405 | 48.7 |
| **PSMA6** | 6 | 25560958 | 25572608 | - | 11650 | 741 | 361 | 48.7 |
| **TASP1** | 3 | 5611597 | 5689047 | + | 77450 | 1363 | 664 | 48.7 |
| **TCF21** | 3 | 59434057 | 59436921 | - | 2864 | 540 | 263 | 48.7 |
| **RIMS1** | 3 | 8.60E+07 | 8.63E+07 | - | 299254 | 5192 | 2528 | 48.7 |
| **HOXA11** | 1 | 122362756 | 122365185 | + | 2429 | 894 | 435 | 48.7 |
| **EYA4** | 3 | 59556600 | 59698777 | - | 142177 | 2527 | 1228 | 48.6 |
| **CHIC2** | 5 | 2.70E+07 | 2.70E+07 | - | 30406 | 552 | 268 | 48.6 |
| **CCNY** | 1 | 143275134 | 143395289 | + | 120155 | 1057 | 513 | 48.5 |
| **SP3** | 7 | 20978900 | 21013491 | - | 34591 | 3577 | 1736 | 48.5 |
| **MAPK1** | 17 | 7991427 | 8025421 | - | 33994 | 1107 | 537 | 48.5 |
| **IRX2** | 1 | 64500684 | 64506548 | + | 5864 | 1437 | 697 | 48.5 |
| **CADPS** | 12 | 1.36E+07 | 1.38E+07 | - | 207938 | 4089 | 1982 | 48.5 |
| **PRKAG2** | 1 | 150054832 | 150263556 | + | 208724 | 2039 | 988 | 48.5 |
| **ATF2** | 7 | 21441758 | 21489562 | - | 47804 | 1464 | 709 | 48.4 |
| **PRKRIP1** | 19 | 6.17E+06 | 6.18E+06 | - | 10669 | 570 | 276 | 48.4 |
| **SP8** | 1 | 124975955 | 124978644 | + | 2689 | 1500 | 726 | 48.4 |
| **DOLPP1** | 20 | 5383237 | 5400038 | - | 16801 | 717 | 347 | 48.4 |
| **CBLN1** | 11 | 13008843 | 13011974 | + | 3131 | 585 | 283 | 48.4 |
| **MEMO1** | 3 | 35085186 | 35112559 | - | 27373 | 982 | 475 | 48.4 |
| **VAPB** | 16 | 3227338 | 3264700 | + | 37362 | 825 | 399 | 48.4 |
| **MAP1LC3A** | 16 | 13269950 | 13287199 | + | 17249 | 366 | 177 | 48.4 |
| **THAP11** | 11 | 6692310 | 6695203 | - | 2893 | 732 | 354 | 48.4 |
| **AMBRA1** | 6 | 39365899 | 39491713 | - | 125814 | 5629 | 2722 | 48.4 |
| **FIGN** | 7 | 17453979 | 17559249 | - | 105270 | 4549 | 2199 | 48.3 |

| | | | | | | | | |
|---|---|---|---|---|---|---|---|---|
| **CHIC1** | 21 | 1863025 | 1882544 | + | 19519 | 549 | 265 | 48.3 |
| **CAPZA2** | 4 | 24804818 | 24836876 | - | 32058 | 970 | 468 | 48.2 |
| **MTA1** | 9 | 27267133 | 27342296 | - | 75163 | 2329 | 1123 | 48.2 |
| **FEZF2** | 12 | 13549436 | 13552515 | - | 3079 | 1311 | 632 | 48.2 |
| **FBXO33** | 6 | 23936058 | 23952910 | + | 16852 | 1572 | 757 | 48.2 |
| **RBPJ** | 5 | 18186504 | 18245144 | + | 58640 | 1523 | 733 | 48.1 |
| **PPM1A** | 6 | 5409623 | 5444659 | + | 35036 | 1269 | 610 | 48.1 |
| **E2F3** | 1 | 93262444 | 93304202 | + | 41758 | 1480 | 711 | 48.0 |
| **NSG1** | 5 | 12516550 | 12539395 | - | 22845 | 558 | 268 | 48.0 |
| **SFRP2** | 5 | 41438636 | 41443023 | - | 4387 | 879 | 422 | 48.0 |
| **PPP3CA** | 5 | 49345601 | 49516211 | + | 170610 | 1754 | 842 | 48.0 |
| **MXI1** | 8 | 26050681 | 26105803 | + | 55122 | 871 | 418 | 48.0 |
| **YBX1** | 22 | 2351563 | 2360876 | - | 9313 | 1210 | 580 | 47.9 |
| **PAIP2** | 14 | 16917602 | 16925874 | - | 8272 | 384 | 184 | 47.9 |
| **PCBP2** | 30 | 753013 | 761202 | - | 8189 | 789 | 378 | 47.9 |
| **KDELR2** | 15 | 9364746 | 9377403 | + | 12657 | 639 | 306 | 47.9 |
| **ACTN1** | 6 | 31303100 | 31389950 | + | 86850 | 2831 | 1354 | 47.8 |
| **ZNF423** | 11 | 12631704 | 12857942 | + | 226238 | 7216 | 3448 | 47.8 |
| **BARHL1** | 20 | 4093179 | 4099201 | - | 6022 | 969 | 463 | 47.8 |
| **ZSWIM6** | Z | 10527782 | 10637784 | - | 110002 | 3805 | 1817 | 47.8 |
| **BTF3L4** | 9 | 5928811 | 5937507 | - | 8696 | 633 | 302 | 47.7 |
| **KPNA3** | 2 | 54387994 | 54435451 | + | 47457 | 1617 | 770 | 47.6 |
| **MECOM** | 10 | 21980364 | 22307526 | + | 327162 | 4396 | 2091 | 47.6 |
| **HOXD12** | 7 | 21841951 | 21849313 | + | 7362 | 801 | 381 | 47.6 |
| **GRIN1** | 20 | 10288667 | 10327727 | + | 39060 | 3621 | 1722 | 47.6 |
| **ILRUN** | 24 | 3489994 | 3518649 | + | 28655 | 876 | 416 | 47.5 |
| **HOXD11** | 7 | 2.19E+07 | 2.19E+07 | + | 1552 | 843 | 400 | 47.4 |
| **CDK14** | 1 | 133367144 | 133715637 | - | 348493 | 1716 | 813 | 47.4 |
| **RAB35** | 17 | 10327025 | 10341906 | - | 14881 | 606 | 287 | 47.4 |
| **MAGI2** | 4 | 10867120 | 11581079 | + | 713959 | 5028 | 2381 | 47.4 |

| | | | | | | | | |
|---|---|---|---|---|---|---|---|---|
| **SPRED1** | 6 | 33989619 | 34051546 | - | 61927 | 1242 | 588 | 47.3 |
| **MTPN** | 4 | 65623757 | 65655159 | + | 31402 | 357 | 169 | 47.3 |
| **RNF11** | 9 | 6214755 | 6245225 | - | 30470 | 465 | 220 | 47.3 |
| **BRINP1** | 20 | 6832399 | 6920105 | + | 87706 | 2286 | 1081 | 47.3 |
| **UBA52** | 26 | 3.68E+05 | 3.69E+05 | - | 1086 | 387 | 183 | 47.3 |
| **NPY** | 1 | 123694226 | 123703124 | - | 8898 | 294 | 139 | 47.3 |
| **MARK3** | 6 | 10012881 | 10074767 | - | 61886 | 2640 | 1248 | 47.3 |
| **MBNL1** | 10 | 25318972 | 25379132 | - | 60160 | 1301 | 615 | 47.3 |
| **NOL4** | 1 | 43398132 | 43588859 | + | 190727 | 2450 | 1158 | 47.3 |
| **RNF208** | 20 | 1.04E+07 | 1.04E+07 | + | 4668 | 798 | 377 | 47.2 |
| **KLF6** | 1 | 144458879 | 144468072 | + | 9193 | 852 | 402 | 47.2 |
| **RAP1GDS1** | 5 | 50080381 | 50170452 | - | 90071 | 1935 | 912 | 47.1 |
| **DAZAP1** | 26 | 1838156 | 1862611 | - | 24455 | 1365 | 643 | 47.1 |
| **USP7** | 15 | 5274392 | 5344934 | - | 70542 | 3312 | 1560 | 47.1 |
| **RAB8B** | 13 | 3621181 | 3650380 | - | 29199 | 845 | 398 | 47.1 |
| **ZCCHC24** | 8 | 11823767 | 11931501 | + | 107734 | 720 | 339 | 47.1 |
| **FOXF2** | 1 | 8.57E+07 | 8.57E+07 | - | 5878 | 1179 | 555 | 47.1 |
| **ARF1** | 1 | 154133543 | 154146916 | - | 13373 | 546 | 257 | 47.1 |
| **AGFG1** | 10 | 1.01E+07 | 1.02E+07 | + | 35953 | 1968 | 926 | 47.1 |
| **MEOX2** | 1 | 127118509 | 127172590 | + | 54081 | 909 | 427 | 47.0 |
| **POU4F2** | 5 | 57748556 | 57750373 | + | 1817 | 2217 | 1041 | 47.0 |
| **CITED2** | 3 | 52194530 | 52197469 | - | 2939 | 720 | 338 | 46.9 |
| **PTP4A1** | 3 | 90050456 | 90064436 | - | 13980 | 522 | 245 | 46.9 |
| **KRAS** | 4 | 67495741 | 67521352 | + | 25611 | 567 | 266 | 46.9 |
| **SAMD12** | 1 | 13417718 | 13594557 | + | 176839 | 712 | 334 | 46.9 |
| **SOX11** | 3 | 99835902 | 99838360 | + | 2458 | 1212 | 568 | 46.9 |
| **SYT7** | 6 | 60189435 | 60212459 | - | 23024 | 2055 | 963 | 46.9 |
| **DYNLL2** | 19 | 1155131 | 1162074 | + | 6943 | 423 | 198 | 46.8 |
| **EIF1** | 27 | 3097436 | 3098586 | - | 1150 | 342 | 160 | 46.8 |
| **GNPDA2** | 5 | 24633745 | 24641475 | - | 7730 | 1281 | 599 | 46.8 |

| | | | | | | | | |
|---|---|---|---|---|---|---|---|---|
| **IGF1** | 4 | 57710520 | 57757738 | + | 47218 | 462 | 216 | 46.8 |
| **UBE2I** | 15 | 16484909 | 16497500 | + | 12591 | 477 | 223 | 46.8 |
| **SNTG1** | 1 | 41464408 | 41791769 | - | 327361 | 1566 | 731 | 46.7 |
| **MPPED2** | 6 | 54763924 | 54867237 | - | 103313 | 885 | 413 | 46.7 |
| **PRDM12** | 20 | 4799758 | 4808174 | - | 8416 | 1349 | 629 | 46.6 |
| **SPCS3** | 5 | 35328745 | 35336625 | - | 7880 | 543 | 253 | 46.6 |
| **MAPK6** | 13 | 8.41E+06 | 8.44E+06 | - | 28153 | 2166 | 1009 | 46.6 |
| **CUL1** | 1 | 1.06E+08 | 1.06E+08 | + | 53721 | 2331 | 1085 | 46.5 |
| **RALA** | 1 | 101852617 | 101887996 | + | 35379 | 621 | 289 | 46.5 |
| **ARL8B** | 12 | 19694954 | 19711612 | + | 16658 | 561 | 261 | 46.5 |
| **DR1** | 9 | 16934471 | 16945634 | + | 11163 | 531 | 247 | 46.5 |
| **ZNF503** | 8 | 14261308 | 14265440 | + | 4132 | 1830 | 851 | 46.5 |
| **RHOBTB2** | 28 | 3540094 | 3555684 | - | 15590 | 2475 | 1150 | 46.5 |
| **GAN** | 11 | 4650660 | 4678774 | - | 28114 | 1794 | 833 | 46.4 |
| **CDYL2** | 11 | 4849446 | 4909249 | - | 59803 | 1524 | 707 | 46.4 |
| **ZFHX4** | 1 | 30749745 | 30927765 | - | 178020 | 9318 | 4319 | 46.4 |
| **ADGRL2** | 9 | 13205306 | 13381793 | + | 176487 | 4700 | 2175 | 46.3 |
| **TSPAN5** | 5 | 49994391 | 50077240 | + | 82849 | 807 | 373 | 46.2 |
| **TECR** | 9 | 17133975 | 17157130 | + | 23155 | 939 | 434 | 46.2 |
| **CUL3** | 10 | 9025737 | 9083740 | - | 58003 | 2307 | 1066 | 46.2 |
| **LHX1** | 19 | 1463779 | 1470230 | - | 6451 | 1221 | 564 | 46.2 |
| **GRIN2B** | 4 | 48721345 | 48936564 | + | 215219 | 4521 | 2087 | 46.2 |
| **SSU72** | 22 | 5596105 | 5614864 | + | 18759 | 585 | 270 | 46.2 |
| **HACD1** | 1 | 135923638 | 135941747 | - | 18109 | 824 | 380 | 46.1 |
| **NELFA** | 5 | 7618831 | 7642804 | - | 23973 | 1620 | 747 | 46.1 |
| **WDR26** | 3 | 18650430 | 18678013 | + | 27583 | 1872 | 863 | 46.1 |
| **NOL4L** | 16 | 7524028 | 7583562 | - | 59534 | 2113 | 973 | 46.0 |
| **EIF3H** | 1 | 14338770 | 14430033 | + | 91263 | 1047 | 482 | 46.0 |
| **SRSF10** | 25 | 365874 | 376284 | - | 10410 | 1154 | 531 | 46.0 |
| **DVL3** | 10 | 17166346 | 17200237 | + | 33891 | 2190 | 1007 | 46.0 |

| | | | | | | | | | |
|---|---|---|---|---|---|---|---|---|---|
| **BCLAF1** | 3 | 50865020 | 50890779 | - | | 25759 | 3022 | 1389 | 46.0 |
| **TRAM1** | 1 | 33554194 | 33569233 | + | | 15039 | 1125 | 517 | 46.0 |
| **ADAMTS6** | Z | 9171102 | 9296645 | + | | 125543 | 3210 | 1474 | 45.9 |
| **SFN** | 25 | 3966025 | 3968080 | - | | 2055 | 741 | 340 | 45.9 |
| **KCNC1** | 6 | 49642928 | 49768496 | - | | 125568 | 1972 | 904 | 45.8 |
| **CABP7** | 17 | 12446233 | 12454619 | + | | 8386 | 648 | 297 | 45.8 |
| **POU3F1** | 25 | 1267143 | 1268700 | + | | 1557 | 1113 | 510 | 45.8 |
| **HOXA6** | 1 | 122402066 | 122404264 | + | | 2198 | 821 | 376 | 45.8 |
| **HOXD13** | 7 | 21840464 | 21847759 | + | | 7295 | 963 | 441 | 45.8 |
| **STRN3** | 6 | 27469969 | 27532237 | + | | 62268 | 2521 | 1154 | 45.8 |
| **FGF16** | 21 | 2570241 | 2581393 | + | | 11152 | 627 | 287 | 45.8 |
| **BMP2** | 3 | 8593445 | 8662147 | - | | 68702 | 1576 | 721 | 45.7 |
| **FOXD2** | 9 | 8370309 | 8372779 | - | | 2470 | 1329 | 608 | 45.7 |
| **DUSP14** | 19 | 1270560 | 1281498 | - | | 10938 | 597 | 273 | 45.7 |
| **MLLT10** | 1 | 137731666 | 137857321 | + | | 125655 | 3333 | 1524 | 45.7 |
| **YWHAB** | 16 | 9029975 | 9043876 | + | | 13901 | 735 | 336 | 45.7 |
| **PTBP2** | 9 | 17925863 | 17980605 | + | | 54742 | 2111 | 964 | 45.7 |
| **TFAP2A** | 1 | 89289055 | 89302567 | - | | 13512 | 1452 | 663 | 45.7 |
| **PHF10** | 3 | 42454992 | 42472207 | - | | 17215 | 1610 | 735 | 45.7 |
| **ARL8A** | 24 | 1632147 | 1645927 | + | | 13780 | 561 | 256 | 45.6 |
| **FZD10** | 17 | 639171 | 640997 | - | | 1826 | 1749 | 798 | 45.6 |
| **BACH2** | 3 | 79038779 | 79228944 | + | | 190165 | 2475 | 1129 | 45.6 |
| **YY1** | 6 | 11938783 | 11964238 | - | | 25455 | 1263 | 576 | 45.6 |
| **TGFB2** | 3 | 20666019 | 20725694 | - | | 59675 | 1239 | 565 | 45.6 |
| **GPR137B** | 3 | 45993162 | 46018391 | + | | 25229 | 1158 | 528 | 45.6 |
| **PTP4A3** | 1 | 2987401 | 3026028 | + | | 38627 | 522 | 238 | 45.6 |
| **CAMKMT** | 3 | 27264266 | 27476257 | + | | 211991 | 933 | 425 | 45.6 |
| **BAMBI** | 1 | 140799152 | 140803843 | + | | 4691 | 786 | 358 | 45.5 |
| **RCOR3** | 3 | 23914950 | 23941429 | - | | 26479 | 2173 | 989 | 45.5 |
| **SATB2** | 7 | 27374401 | 27502661 | + | | 128260 | 2721 | 1238 | 45.5 |
| **CABP1** | 17 | 10606541 | 10633170 | + | | 26629 | 1284 | 584 | 45.5 |

| | | | | | | | | |
|---|---|---|---|---|---|---|---|---|
| **AIG1** | 3 | 53580050 | 53705717 | + | 125667 | 717 | 326 | 45.5 |
| **PCDH17** | 2 | 60154561 | 60244753 | + | 90192 | 4054 | 1843 | 45.5 |
| **KLHL15** | 2 | 105659305 | 105687286 | - | 27981 | 1815 | 825 | 45.5 |
| **AIDA** | 3 | 19170897 | 19199893 | + | 28996 | 933 | 424 | 45.4 |
| **RGS17** | 3 | 56614450 | 56687256 | + | 72806 | 711 | 323 | 45.4 |
| **MSANTD3** | 1 | 62010357 | 62022694 | + | 12337 | 1147 | 521 | 45.4 |
| **PCGF1** | 5 | 1303827 | 1306870 | - | 3043 | 786 | 357 | 45.4 |
| **AEBP2** | 4 | 71770431 | 71812910 | - | 42479 | 1512 | 686 | 45.4 |
| **PPM1L** | 10 | 24003770 | 24065179 | + | 61409 | 582 | 264 | 45.4 |
| **GTF2A1** | 6 | 20522466 | 20549065 | + | 26599 | 1193 | 541 | 45.3 |
| **ARL4C** | 7 | 35800768 | 35803662 | + | 2894 | 651 | 295 | 45.3 |
| **HPCAL4** | 25 | 231018 | 234611 | - | 3593 | 576 | 261 | 45.3 |
| **MID1IP1** | 2 | 111021012 | 111022806 | + | 1794 | 495 | 224 | 45.3 |
| **NLK** | 19 | 542050 | 578054 | - | 36004 | 2009 | 909 | 45.2 |
| **CNEP1R1** | 11 | 12571111 | 12577208 | - | 6097 | 378 | 171 | 45.2 |
| **BTRC** | 8 | 22798470 | 22918544 | - | 120074 | 1887 | 853 | 45.2 |
| **VAPA** | 1 | 52502404 | 52530530 | + | 28126 | 750 | 339 | 45.2 |
| **EPC1** | 1 | 142062909 | 142126331 | - | 63422 | 2713 | 1226 | 45.2 |
| **VSX2** | 6 | 23744366 | 23767149 | - | 22783 | 1556 | 702 | 45.1 |
| **EPHX4** | 9 | 16578409 | 16595192 | + | 16783 | 1104 | 498 | 45.1 |
| **EVL** | 6 | 11998889 | 12144843 | - | 145954 | 1455 | 656 | 45.1 |
| **USP15** | 4 | 33087676 | 33147305 | + | 59629 | 3058 | 1378 | 45.1 |
| **CRYBA1** | 19 | 4112670 | 4117099 | - | 4429 | 648 | 292 | 45.1 |
| **NETO1** | 1 | 58422057 | 58484903 | - | 62846 | 1677 | 755 | 45.0 |
| **BNC2** | Z | 33250304 | 33382837 | - | 132533 | 471 | 212 | 45.0 |
| **PSMD12** | 18 | 8709989 | 8718577 | + | 8588 | 1371 | 617 | 45.0 |
| **SLC35F3** | 3 | 45102781 | 45273229 | + | 170448 | 1558 | 701 | 45.0 |
| **PITPNA** | 19 | 4882642 | 4907328 | + | 24686 | 816 | 367 | 45.0 |
| **CREB5** | 1 | 121494012 | 121749269 | - | 255257 | 1628 | 732 | 45.0 |
| **NTMT1** | 20 | 5201203 | 5204397 | - | 3194 | 672 | 302 | 44.9 |
| **SHOC2** | 8 | 26439998 | 26499904 | + | 59906 | 1749 | 786 | 44.9 |

| | | | | | | | | |
|---|---|---|---|---|---|---|---|---|
| **LRRTM1** | 5 | 3182470 | 3186246 | - | 3776 | 1566 | 703 | 44.9 |
| **UBE2D1** | 8 | 5762792 | 5778675 | + | 15883 | 468 | 210 | 44.9 |
| **FNBP1** | 20 | 5005824 | 5098852 | + | 93028 | 2691 | 1207 | 44.9 |
| **MOSPD1** | Z | 84546692 | 84560738 | - | 14046 | 881 | 395 | 44.8 |
| **ELFN1** | 15 | 11462162 | 11565714 | + | 103552 | 2391 | 1071 | 44.8 |
| **PDS5A** | 5 | 22924394 | 23002958 | - | 78564 | 4222 | 1891 | 44.8 |
| **SESN3** | 2 | 38675373 | 38720084 | - | 44711 | 1302 | 583 | 44.8 |
| **NTNG1** | 9 | 20738969 | 20889445 | + | 150476 | 1749 | 783 | 44.8 |
| **STT3B** | 1 | 114624245 | 114675050 | - | 50805 | 2400 | 1074 | 44.8 |
| **ISL2** | 13 | 922599 | 925814 | + | 3215 | 1080 | 483 | 44.7 |
| **GPHN** | 6 | 32215750 | 32491497 | - | 275747 | 2859 | 1277 | 44.7 |
| **MAP3K7** | 3 | 78899462 | 78948694 | + | 49232 | 2092 | 934 | 44.6 |
| **PPP1R12A** | 4 | 40053463 | 40177546 | - | 124083 | 3108 | 1386 | 44.6 |
| **DERL3** | 17 | 11882626 | 11887667 | + | 5041 | 729 | 325 | 44.6 |
| **MYL10** | 19 | 6573210 | 6595943 | + | 22733 | 498 | 222 | 44.6 |
| **ABLIM2** | 5 | 10189171 | 10315615 | + | 126444 | 2046 | 912 | 44.6 |
| **ZBTB20** | 2 | 23394065 | 23869013 | - | 474948 | 3814 | 1700 | 44.6 |
| **NMT2** | 1 | 134950491 | 134983007 | - | 32516 | 1497 | 667 | 44.6 |
| **COTL1** | 11 | 3219648 | 3239010 | + | 19362 | 429 | 191 | 44.5 |
| **SHOX** | 2 | 84713441 | 84724422 | - | 10981 | 876 | 390 | 44.5 |
| **KCNQ2** | 16 | 6076997 | 6152378 | - | 75381 | 2997 | 1334 | 44.5 |
| **RASL10B** | 19 | 5852731 | 5862843 | + | 10112 | 1101 | 490 | 44.5 |
| **PTDSS1** | 1 | 23042622 | 23070764 | - | 28142 | 1628 | 724 | 44.5 |
| **KLHL14** | 1 | 44016040 | 44076745 | + | 60705 | 1878 | 835 | 44.5 |
| **ZNF521** | 1 | 46871469 | 47103379 | + | 231910 | 7451 | 3312 | 44.5 |
| **PCGF3** | Z | 72282181 | 72336436 | - | 54255 | 862 | 383 | 44.4 |
| **BIN1** | 7 | 13931784 | 14023211 | - | 91427 | 1830 | 813 | 44.4 |
| **PPARA** | 4 | 68379700 | 68412278 | - | 32578 | 1513 | 672 | 44.4 |
| **RGS8** | 9 | 29032316 | 29058544 | + | 26228 | 635 | 282 | 44.4 |
| **DCUN1D4** | 5 | 26189176 | 26229014 | + | 39838 | 1131 | 502 | 44.4 |
| **NUDT4** | 4 | 46222232 | 46249319 | + | 27087 | 631 | 280 | 44.4 |

| | | | | | | | | |
|---|---|---|---|---|---|---|---|---|
| **CSNK1E** | 4 | 53320495 | 53342562 | + | 22067 | 1251 | 555 | 44.4 |
| **NAA50** | 2 | 27213028 | 27233673 | - | 20645 | 819 | 363 | 44.3 |
| **CADM2** | 2 | 6218343 | 6795080 | - | 576737 | 1375 | 609 | 44.3 |
| **NKX2-3** | 8 | 21859467 | 21861707 | - | 2240 | 1974 | 874 | 44.3 |
| **ABHD17C** | 13 | 11907179 | 11935978 | - | 28799 | 933 | 413 | 44.3 |

**Table S16 -Gene ontology analysis on PhyloP conserved genes.** The top 5% (606) genes with more accelerated bases overlapped with CDS ("LOC" genes excluded) were tested. The first 100 lines are showed.

| GO term | p.geomean | stat.mean | p.val | q.val | set.size | exp1 | GO term name |
|---|---|---|---|---|---|---|---|
| GO:1990837 | 1.24E-08 | 5.646982295 | 1.24E-08 | 4.22E-05 | 394 | 1.24E-08 | sequence-specific double-stranded DNA binding |
| GO:0045892 | 3.61E-08 | 5.448719491 | 3.61E-08 | 6.12E-05 | 407 | 3.61E-08 | negative regulation of transcription, DNA-templated |
| GO:0003700 | 6.63E-08 | 5.325497347 | 6.63E-08 | 7.50E-05 | 469 | 6.63E-08 | DNA-binding transcription factor activity |
| GO:0043565 | 5.22E-07 | 4.939109527 | 5.22E-07 | 0.000442879 | 343 | 5.22E-07 | sequence-specific DNA binding |
| GO:0001228 | 1.66E-06 | 4.701596018 | 1.66E-06 | 0.001123946 | 324 | 1.66E-06 | DNA-binding transcription activator activity, RNA polymerase II-specific |
| GO:0008134 | 1.73E-05 | 4.191052732 | 1.73E-05 | 0.009768698 | 240 | 1.73E-05 | transcription factor binding |
| GO:0007399 | 2.81E-05 | 4.051681629 | 2.81E-05 | 0.01360715 | 440 | 2.81E-05 | nervous system development |
| GO:0005667 | 6.00E-05 | 3.906384172 | 6.00E-05 | 0.025428411 | 166 | 6.00E-05 | transcription regulator complex |
| GO:0000977 | 0.000126182 | 3.695609073 | 0.000126182 | 0.047570723 | 229 | 0.000126182 | RNA polymerase II transcription regulatory region sequence-specific DNA binding |
| GO:0000165 | 0.000335715 | 3.429373373 | 0.000335715 | 0.113908255 | 230 | 0.000335715 | MAPK cascade |
| GO:0000987 | 0.000484272 | 3.417081778 | 0.000484272 | 0.14364186 | 61 | 0.000484272 | cis-regulatory region sequence-specific DNA binding |
| GO:0009887 | 0.000508017 | 3.338363757 | 0.000508017 | 0.14364186 | 122 | 0.000508017 | animal organ morphogenesis |
| GO:0009952 | 0.000560403 | 3.338674072 | 0.000560403 | 0.146265234 | 82 | 0.000560403 | anterior/posterior pattern specification |
| GO:0070936 | 0.000673827 | 3.347672489 | 0.000673827 | 0.163306678 | 48 | 0.000673827 | protein K48-linked ubiquitination |
| GO:0001227 | 0.000942588 | 3.13560358 | 0.000942588 | 0.213213356 | 183 | 0.000942588 | DNA-binding transcription repressor activity, RNA polymerase II-specific |
| GO:0061631 | 0.001020932 | 3.382977443 | 0.001020932 | 0.216501408 | 23 | 0.001020932 | ubiquitin conjugating enzyme activity |
| GO:0008284 | 0.001413664 | 2.997248451 | 0.001413664 | 0.282150649 | 370 | 0.001413664 | positive regulation of cell population proliferation |
| GO:0006511 | 0.001586101 | 2.969629434 | 0.001586101 | 0.298980004 | 223 | 0.001586101 | ubiquitin-dependent protein catabolic process |
| GO:0098978 | 0.0018269 | 2.92000323 | 0.0018269 | 0.326245811 | 295 | 0.0018269 | glutamatergic synapse |
| GO:0031625 | 0.001944592 | 2.905542961 | 0.001944592 | 0.329422708 | 216 | 0.001944592 | ubiquitin protein ligase binding |
| GO:1900740 | 0.002038867 | 3.244409302 | 0.002038867 | 0.329422708 | 16 | 0.002038867 | positive regulation of protein insertion into mitochondrial membrane involved in apoptotic signaling pathway |
| GO:0010468 | 0.002367351 | 2.841389276 | 0.002367351 | 0.36511014 | 222 | 0.002367351 | regulation of gene expression |
| GO:0003682 | 0.002736452 | 2.787901134 | 0.002736452 | 0.401281403 | 338 | 0.002736452 | chromatin binding |
| GO:0019003 | 0.002838418 | 2.837093262 | 0.002838418 | 0.401281403 | 59 | 0.002838418 | GDP binding |
| GO:0010628 | 0.002995891 | 2.757488391 | 0.002995891 | 0.406602358 | 356 | 0.002995891 | positive regulation of gene expression |
| GO:0003151 | 0.003492259 | 2.79302391 | 0.003492259 | 0.45573976 | 43 | 0.003492259 | outflow tract morphogenesis |
| GO:0048704 | 0.003750725 | 2.764175965 | 0.003750725 | 0.471341059 | 44 | 0.003750725 | embryonic skeletal system morphogenesis |
| GO:0000976 | 0.004417561 | 2.634543109 | 0.004417561 | 0.535313765 | 188 | 0.004417561 | transcription regulatory region sequence-specific DNA binding |
| GO:0030182 | 0.004580254 | 2.626522148 | 0.004580254 | 0.535889695 | 150 | 0.004580254 | neuron differentiation |
| GO:0010629 | 0.004955134 | 2.592999359 | 0.004955134 | 0.560425688 | 205 | 0.004955134 | negative regulation of gene expression |
| GO:0051549 | 0.005752803 | 2.931801726 | 0.005752803 | 0.59802306 | 11 | 0.005752803 | positive regulation of keratinocyte migration |
| GO:0008083 | 0.006198829 | 2.52690836 | 0.006198829 | 0.59802306 | 108 | 0.006198829 | growth factor activity |
| GO:0001654 | 0.006369607 | 2.593571039 | 0.006369607 | 0.59802306 | 33 | 0.006369607 | eye development |
| GO:0030334 | 0.006495688 | 2.523690533 | 0.006495688 | 0.59802306 | 74 | 0.006495688 | regulation of cell migration |
| GO:0003729 | 0.006528657 | 2.50038294 | 0.006528657 | 0.59802306 | 147 | 0.006528657 | mRNA binding |
| GO:0021983 | 0.006628593 | 2.617049945 | 0.006628593 | 0.59802306 | 25 | 0.006628593 | pituitary gland development |
| GO:0003925 | 0.006628593 | 2.617049945 | 0.006628593 | 0.59802306 | 25 | 0.006628593 | obsolete small monomeric GTPase activity |
| GO:0043484 | 0.006874707 | 2.559594307 | 0.006874707 | 0.59802306 | 34 | 0.006874707 | regulation of RNA splicing |
| GO:0010975 | 0.006874707 | 2.559594307 | 0.006874707 | 0.59802306 | 34 | 0.006874707 | regulation of neuron projection development |
| GO:0003924 | 0.007084287 | 2.465237564 | 0.007084287 | 0.59802306 | 203 | 0.007084287 | GTPase activity |
| GO:0003690 | 0.007355656 | 2.466071095 | 0.007355656 | 0.59802306 | 100 | 0.007355656 | double-stranded DNA binding |

| | | | | | | | |
|---|---|---|---|---|---|---|---|
| GO:0001658 | 0.007402584 | 2.526790881 | 0.007402584 | 0.59802306 | 35 | 0.007402584 | branching involved in ureteric bud morphogenesis |
| GO:0030324 | 0.009479826 | 2.376130804 | 0.009479826 | 0.740119143 | 82 | 0.009479826 | lung development |
| GO:0090575 | 0.009840225 | 2.374329876 | 0.009840225 | 0.740119143 | 60 | 0.009840225 | RNA polymerase II transcription regulator complex |
| GO:0001656 | 0.010067244 | 2.417325816 | 0.010067244 | 0.740119143 | 30 | 0.010067244 | metanephros development |
| GO:0038095 | 0.010362212 | 2.340774231 | 0.010362212 | 0.740119143 | 84 | 0.010362212 | Fc-epsilon receptor signaling pathway |
| GO:0000209 | 0.010667119 | 2.312241225 | 0.010667119 | 0.740119143 | 203 | 0.010667119 | protein polyubiquitination |
| GO:0007223 | 0.010845416 | 2.382500879 | 0.010845416 | 0.740119143 | 31 | 0.010845416 | Wnt signaling pathway, calcium modulating pathway |
| GO:0048856 | 0.010851519 | 2.343853392 | 0.010851519 | 0.740119143 | 51 | 0.010851519 | anatomical structure development |
| GO:0008543 | 0.01090656 | 2.332048968 | 0.01090656 | 0.740119143 | 62 | 0.01090656 | fibroblast growth factor receptor signaling pathway |
| GO:0004842 | 0.011990904 | 2.2676406 | 0.011990904 | 0.797747797 | 193 | 0.011990904 | ubiquitin-protein transferase activity |
| GO:0006417 | 0.012317927 | 2.265337942 | 0.012317927 | 0.802021374 | 113 | 0.012317927 | regulation of translation |
| GO:0006325 | 0.013018603 | 2.233432465 | 0.013018603 | 0.802021374 | 239 | 0.013018603 | chromatin organization |
| GO:0021762 | 0.013377186 | 2.285110766 | 0.013377186 | 0.802021374 | 34 | 0.013377186 | substantia nigra development |
| GO:0003730 | 0.013467378 | 2.252202474 | 0.013467378 | 0.802021374 | 55 | 0.013467378 | mRNA 3'-UTR binding |
| GO:0060076 | 0.013711241 | 2.305151624 | 0.013711241 | 0.802021374 | 25 | 0.013711241 | excitatory synapse |
| GO:0048536 | 0.013711241 | 2.305151624 | 0.013711241 | 0.802021374 | 25 | 0.013711241 | spleen development |
| GO:0043488 | 0.013933744 | 2.221069906 | 0.013933744 | 0.802021374 | 91 | 0.013933744 | regulation of mRNA stability |
| GO:0031954 | 0.01412248 | 2.350517468 | 0.01412248 | 0.802021374 | 17 | 0.01412248 | positive regulation of protein autophosphorylation |
| GO:0042752 | 0.014182518 | 2.230090501 | 0.014182518 | 0.802021374 | 56 | 0.014182518 | regulation of circadian rhythm |
| GO:0001501 | 0.014782176 | 2.191746247 | 0.014782176 | 0.822228253 | 118 | 0.014782176 | skeletal system development |
| GO:0007420 | 0.015682963 | 2.160259043 | 0.015682963 | 0.822530709 | 217 | 0.015682963 | brain development |
| GO:0030900 | 0.015689729 | 2.186716873 | 0.015689729 | 0.822530709 | 58 | 0.015689729 | forebrain development |
| GO:0030326 | 0.015751253 | 2.194599639 | 0.015751253 | 0.822530709 | 47 | 0.015751253 | embryonic limb morphogenesis |
| GO:0048935 | 0.015999713 | 2.396877951 | 0.015999713 | 0.822530709 | 11 | 0.015999713 | peripheral nervous system neuron development |
| GO:0001823 | 0.015999713 | 2.396877951 | 0.015999713 | 0.822530709 | 11 | 0.015999713 | mesonephros development |
| GO:0010494 | 0.016482794 | 2.165428424 | 0.016482794 | 0.834718206 | 59 | 0.016482794 | cytoplasmic stress granule |
| GO:0009653 | 0.016987461 | 2.138951967 | 0.016987461 | 0.838640287 | 96 | 0.016987461 | anatomical structure morphogenesis |
| GO:0060425 | 0.017124574 | 2.240496875 | 0.017124574 | 0.838640287 | 19 | 0.017124574 | lung morphogenesis |
| GO:0016055 | 0.017545139 | 2.11650851 | 0.017545139 | 0.838640287 | 178 | 0.017545139 | Wnt signaling pathway |
| GO:0050679 | 0.017548913 | 2.14667746 | 0.017548913 | 0.838640287 | 49 | 0.017548913 | positive regulation of epithelial cell proliferation |
| GO:0007254 | 0.018305307 | 2.140394571 | 0.018305307 | 0.852251414 | 39 | 0.018305307 | JNK cascade |
| GO:0003712 | 0.01833609 | 2.106804594 | 0.01833609 | 0.852251414 | 98 | 0.01833609 | transcription coregulator activity |
| GO:0042056 | 0.018692344 | 2.191918015 | 0.018692344 | 0.857069234 | 20 | 0.018692344 | chemoattractant activity |
| GO:0005525 | 0.019347222 | 2.073323422 | 0.019347222 | 0.866307018 | 255 | 0.019347222 | GTP binding |
| GO:0000381 | 0.01940446 | 2.113455478 | 0.01940446 | 0.866307018 | 40 | 0.01940446 | regulation of alternative mRNA splicing, via spliceosome |
| GO:0051403 | 0.020309554 | 2.146599881 | 0.020309554 | 0.877360576 | 21 | 0.020309554 | stress-activated MAPK cascade |
| GO:0019904 | 0.0213354 | 2.034826342 | 0.0213354 | 0.877360576 | 185 | 0.0213354 | protein domain specific binding |
| GO:0106310 | 0.021619568 | 2.026508624 | 0.021619568 | 0.877360576 | 275 | 0.021619568 | protein serine kinase activity |
| GO:0106311 | 0.021619568 | 2.026508624 | 0.021619568 | 0.877360576 | 275 | 0.021619568 | protein threonine kinase activity |
| GO:0007409 | 0.021651726 | 2.041571986 | 0.021651726 | 0.877360576 | 77 | 0.021651726 | axonogenesis |
| GO:0017053 | 0.021722346 | 2.061188641 | 0.021722346 | 0.877360576 | 42 | 0.021722346 | transcription repressor complex |
| GO:0048538 | 0.021722346 | 2.061188641 | 0.021722346 | 0.877360576 | 42 | 0.021722346 | thymus development |
| GO:0021537 | 0.021979266 | 2.103997674 | 0.021979266 | 0.877360576 | 22 | 0.021979266 | telencephalon development |

| | | | | | | | |
|---|---|---|---|---|---|---|---|
| **GO:0046329** | 0.021979266 | 2.103997674 | 0.021979266 | 0.877360576 | 22 | 0.021979266 | negative regulation of JNK cascade |
| **GO:0003007** | 0.022942506 | 2.035786411 | 0.022942506 | 0.880049865 | 43 | 0.022942506 | heart morphogenesis |
| **GO:0045778** | 0.023084126 | 2.14526608 | 0.023084126 | 0.880049865 | 14 | 0.023084126 | positive regulation of ossification |
| **GO:0006376** | 0.023084126 | 2.14526608 | 0.023084126 | 0.880049865 | 14 | 0.023084126 | mRNA splice site selection |
| **GO:0046827** | 0.023084126 | 2.14526608 | 0.023084126 | 0.880049865 | 14 | 0.023084126 | positive regulation of protein export from nucleus |
| **GO:0043161** | 0.023727107 | 1.99185778 | 0.023727107 | 0.884990003 | 146 | 0.023727107 | proteasome-mediated ubiquitin-dependent protein catabolic process |
| **GO:0045665** | 0.023735364 | 2.011027281 | 0.023735364 | 0.884990003 | 55 | 0.023735364 | negative regulation of neuron differentiation |
| **GO:0003714** | 0.024405902 | 1.980730889 | 0.024405902 | 0.88660381 | 133 | 0.024405902 | transcription corepressor activity |
| **GO:0001934** | 0.025186892 | 1.96593398 | 0.025186892 | 0.88660381 | 148 | 0.025186892 | positive regulation of protein phosphorylation |
| **GO:1990090** | 0.025347677 | 2.002810835 | 0.025347677 | 0.88660381 | 34 | 0.025347677 | cellular response to nerve growth factor stimulus |
| **GO:0048557** | 0.025466995 | 2.082873982 | 0.025466995 | 0.88660381 | 15 | 0.025466995 | embryonic digestive tract morphogenesis |
| **GO:0048663** | 0.0254865 | 2.02533112 | 0.0254865 | 0.88660381 | 24 | 0.0254865 | neuron fate commitment |
| **GO:0032436** | 0.026076936 | 1.968076542 | 0.026076936 | 0.88660381 | 57 | 0.026076936 | positive regulation of proteasomal ubiquitin-dependent protein catabolic process |
| **GO:1990904** | 0.026269246 | 1.951343376 | 0.026269246 | 0.88660381 | 108 | 0.026269246 | ribonucleoprotein complex |
| **GO:0048156** | 0.026883645 | 1.974110394 | 0.026883645 | 0.88660381 | 35 | 0.026883645 | tau protein binding |
| **GO:0006605** | 0.026883645 | 1.974110394 | 0.026883645 | 0.88660381 | 35 | 0.026883645 | protein targeting |
| **GO:0098685** | 0.030581211 | 1.890973513 | 0.030581211 | 0.882742861 | 72 | 0.030581211 | Schaffer collateral - CA1 synapse |
| **GO:0019901** | 0.03073677 | 1.8733243 | 0.03073677 | 0.882742861 | 350 | 0.03073677 | protein kinase binding |
| **GO:0001759** | 0.030913453 | 2.068909427 | 0.030913453 | 0.882742861 | 10 | 0.030913453 | organ induction |
| **GO:0051865** | 0.031021778 | 1.889690565 | 0.031021778 | 0.882742861 | 58 | 0.031021778 | protein autoubiquitination |
| **GO:0010800** | 0.031298032 | 1.9455994 | 0.031298032 | 0.882742861 | 20 | 0.031298032 | positive regulation of peptidyl-threonine phosphorylation |
| **GO:0070534** | 0.031822692 | 1.901519256 | 0.031822692 | 0.882742861 | 32 | 0.031822692 | protein K63-linked ubiquitination |
| **GO:0048013** | 0.033978332 | 1.841876511 | 0.033978332 | 0.882742861 | 75 | 0.033978332 | ephrin receptor signaling pathway |
| **GO:0048935** | 0.03403641 | 1.991312897 | 0.03403641 | 0.882742861 | 11 | 0.03403641 | peripheral nervous system neuron development |
| **GO:0004672** | 0.034795782 | 1.817192367 | 0.034795782 | 0.882742861 | 396 | 0.034795782 | protein kinase activity |
| **GO:0030326** | 0.034856697 | 1.84172909 | 0.034856697 | 0.882742861 | 47 | 0.034856697 | embryonic limb morphogenesis |
| **GO:0021537** | 0.035418604 | 1.874831543 | 0.035418604 | 0.882742861 | 22 | 0.035418604 | telencephalon development |
| **GO:0001944** | 0.035418604 | 1.874831543 | 0.035418604 | 0.882742861 | 22 | 0.035418604 | vasculature development |
| **GO:0046329** | 0.035418604 | 1.874831543 | 0.035418604 | 0.882742861 | 22 | 0.035418604 | negative regulation of JNK cascade |
| **GO:0051965** | 0.036351995 | 1.821171156 | 0.036351995 | 0.882742861 | 48 | 0.036351995 | positive regulation of synapse assembly |
| **GO:0045444** | 0.036351995 | 1.821171156 | 0.036351995 | 0.882742861 | 48 | 0.036351995 | fat cell differentiation |
| **GO:0007409** | 0.036376008 | 1.809731214 | 0.036376008 | 0.882742861 | 77 | 0.036376008 | axonogenesis |
| **GO:0048156** | 0.036908922 | 1.82573449 | 0.036908922 | 0.882742861 | 35 | 0.036908922 | tau protein binding |
| **GO:0006605** | 0.036908922 | 1.82573449 | 0.036908922 | 0.882742861 | 35 | 0.036908922 | protein targeting |
| **GO:0046983** | 0.037059851 | 1.793498598 | 0.037059851 | 0.882742861 | 142 | 0.037059851 | protein dimerization activity |
| **GO:0021895** | 0.037086859 | 1.926023518 | 0.037086859 | 0.882742861 | 12 | 0.037086859 | cerebral cortex neuron differentiation |
| **GO:0002089** | 0.037086859 | 1.926023518 | 0.037086859 | 0.882742861 | 12 | 0.037086859 | lens morphogenesis in camera-type eye |
| **GO:0046622** | 0.037086859 | 1.926023518 | 0.037086859 | 0.882742861 | 12 | 0.037086859 | positive regulation of organ growth |
| **GO:0099054** | 0.037086859 | 1.926023518 | 0.037086859 | 0.882742861 | 12 | 0.037086859 | presynapse assembly |
| **GO:0032330** | 0.037086859 | 1.926023518 | 0.037086859 | 0.882742861 | 12 | 0.037086859 | regulation of chondrocyte differentiation |
| **GO:0006367** | 0.037131995 | 1.795200286 | 0.037131995 | 0.882742861 | 109 | 0.037131995 | transcription initiation from RNA polymerase II promoter |
| **GO:0051893** | 0.037553039 | 1.8418953 | 0.037553039 | 0.882742861 | 23 | 0.037553039 | regulation of focal adhesion assembly |
| **GO:0071276** | 0.037553039 | 1.8418953 | 0.037553039 | 0.882742861 | 23 | 0.037553039 | cellular response to cadmium ion |

| | | | | | | | |
|---|---|---|---|---|---|---|---|
| **GO:0018105** | 0.038797733 | 1.773166175 | 0.038797733 | 0.882742861 | 127 | 0.038797733 | peptidyl-serine phosphorylation |
| **GO:0031016** | 0.039739941 | 1.810290659 | 0.039739941 | 0.882742861 | 24 | 0.039739941 | pancreas development |
| **GO:0008016** | 0.039739941 | 1.810290659 | 0.039739941 | 0.882742861 | 24 | 0.039739941 | regulation of heart contraction |
| **GO:0001829** | 0.040112393 | 1.869275373 | 0.040112393 | 0.882742861 | 13 | 0.040112393 | trophectodermal cell differentiation |
| **GO:0031462** | 0.040112393 | 1.869275373 | 0.040112393 | 0.882742861 | 13 | 0.040112393 | Cul2-RING ubiquitin ligase complex |
| **GO:0030014** | 0.040112393 | 1.869275373 | 0.040112393 | 0.882742861 | 13 | 0.040112393 | CCR4-NOT complex |
| **GO:0021542** | 0.040112393 | 1.869275373 | 0.040112393 | 0.882742861 | 13 | 0.040112393 | dentate gyrus development |
| **GO:0031175** | 0.040417015 | 1.755198982 | 0.040417015 | 0.882742861 | 112 | 0.040417015 | neuron projection development |
| **GO:0008283** | 0.040890312 | 1.748355386 | 0.040890312 | 0.882742861 | 129 | 0.040890312 | cell population proliferation |
| **GO:0045727** | 0.041067641 | 1.761021258 | 0.041067641 | 0.882742861 | 51 | 0.041067641 | positive regulation of translation |
| **GO:0043388** | 0.041981109 | 1.779845877 | 0.041981109 | 0.882742861 | 25 | 0.041981109 | positive regulation of DNA binding |
| **GO:0051090** | 0.041981109 | 1.779845877 | 0.041981109 | 0.882742861 | 25 | 0.041981109 | regulation of DNA-binding transcription factor activity |
| **GO:0048469** | 0.041981109 | 1.779845877 | 0.041981109 | 0.882742861 | 25 | 0.041981109 | cell maturation |
| **GO:0009953** | 0.042409965 | 1.754508605 | 0.042409965 | 0.882742861 | 38 | 0.042409965 | dorsal/ventral pattern formation |
| **GO:0019899** | 0.042461046 | 1.726209768 | 0.042461046 | 0.882742861 | 274 | 0.042461046 | enzyme binding |
| **GO:0001934** | 0.042997181 | 1.7234701 | 0.042997181 | 0.882742861 | 148 | 0.042997181 | positive regulation of protein phosphorylation |
| **GO:0046827** | 0.043143784 | 1.818722748 | 0.043143784 | 0.882742861 | 14 | 0.043143784 | positive regulation of protein export from nucleus |
| **GO:0048511** | 0.043893389 | 1.715719519 | 0.043893389 | 0.882742861 | 115 | 0.043893389 | rhythmic process |
| **GO:0090263** | 0.043893389 | 1.715719519 | 0.043893389 | 0.882742861 | 115 | 0.043893389 | positive regulation of canonical Wnt signaling pathway |
| **GO:0035116** | 0.044278043 | 1.750420217 | 0.044278043 | 0.882742861 | 26 | 0.044278043 | embryonic hindlimb morphogenesis |
| **GO:0051781** | 0.044338199 | 1.73160894 | 0.044338199 | 0.882742861 | 39 | 0.044338199 | positive regulation of cell division |
| **GO:0009948** | 0.046201719 | 1.772826323 | 0.046201719 | 0.882742861 | 15 | 0.046201719 | anterior/posterior axis specification |
| **GO:0036477** | 0.046201719 | 1.772826323 | 0.046201719 | 0.882742861 | 15 | 0.046201719 | somatodendritic compartment |
| **GO:0010880** | 0.046201719 | 1.772826323 | 0.046201719 | 0.882742861 | 15 | 0.046201719 | regulation of release of sequestered calcium ion into cytosol by sarcoplasmic reticulum |
| **GO:0008380** | 0.046487833 | 1.683868791 | 0.046487833 | 0.882742861 | 223 | 0.046487833 | RNA splicing |
| **GO:0045931** | 0.046631997 | 1.721897183 | 0.046631997 | 0.882742861 | 27 | 0.046631997 | positive regulation of mitotic cell cycle |
| **GO:0001501** | 0.047565084 | 1.676737294 | 0.047565084 | 0.882742861 | 118 | 0.047565084 | skeletal system development |
| **GO:0001889** | 0.047944763 | 1.679063384 | 0.047944763 | 0.882742861 | 70 | 0.047944763 | liver development |
| **GO:0006469** | 0.047944763 | 1.679063384 | 0.047944763 | 0.882742861 | 70 | 0.047944763 | negative regulation of protein kinase activity |
| **GO:0000086** | 0.048316773 | 1.670463034 | 0.048316773 | 0.882742861 | 102 | 0.048316773 | G2/M transition of mitotic cell cycle |
| **GO:0030336** | 0.048532857 | 1.670188281 | 0.048532857 | 0.882742861 | 86 | 0.048532857 | negative regulation of cell migration |
| **GO:0050321** | 0.049300549 | 1.730532944 | 0.049300549 | 0.882742861 | 16 | 0.049300549 | tau-protein kinase activity |
| **GO:0030851** | 0.049300549 | 1.730532944 | 0.049300549 | 0.882742861 | 16 | 0.049300549 | granulocyte differentiation |
| **GO:0045182** | 0.049300549 | 1.730532944 | 0.049300549 | 0.882742861 | 16 | 0.049300549 | translation regulator activity |
| **GO:0035019** | 0.05041208 | 1.665058492 | 0.05041208 | 0.882742861 | 42 | 0.05041208 | somatic stem cell population maintenance |
| **GO:0004879** | 0.05041208 | 1.665058492 | 0.05041208 | 0.882742861 | 42 | 0.05041208 | nuclear receptor activity |
| **GO:0001102** | 0.05041208 | 1.665058492 | 0.05041208 | 0.882742861 | 42 | 0.05041208 | RNA polymerase II activating transcription factor binding |
| **GO:0044325** | 0.051091109 | 1.642856815 | 0.051091109 | 0.882742861 | 104 | 0.051091109 | ion channel binding |
| **GO:0030335** | 0.052119034 | 1.629393659 | 0.052119034 | 0.882742861 | 174 | 0.052119034 | positive regulation of cell migration |
| **GO:0000289** | 0.05245047 | 1.691097017 | 0.05245047 | 0.882742861 | 17 | 0.05245047 | nuclear-transcribed mRNA poly(A) tail shortening |
| **GO:0004970** | 0.05245047 | 1.691097017 | 0.05245047 | 0.882742861 | 17 | 0.05245047 | ionotropic glutamate receptor activity |
| **GO:0060999** | 0.05245047 | 1.691097017 | 0.05245047 | 0.882742861 | 17 | 0.05245047 | positive regulation of dendritic spine development |
| **GO:0008135** | 0.05245047 | 1.691097017 | 0.05245047 | 0.882742861 | 17 | 0.05245047 | translation factor activity, RNA binding |

| | | | | | | | |
|---|---|---|---|---|---|---|---|
| **GO:0031252** | 0.052534142 | 1.643519311 | 0.052534142 | 0.882742861 | 43 | 0.052534142 | cell leading edge |
| **GO:0003713** | 0.054407284 | 1.608077173 | 0.054407284 | 0.882742861 | 176 | 0.054407284 | transcription coactivator activity |
| **GO:0009792** | 0.055658864 | 1.653975382 | 0.055658864 | 0.882742861 | 18 | 0.055658864 | embryo development ending in birth or egg hatching |
| **GO:0010613** | 0.055658864 | 1.653975382 | 0.055658864 | 0.882742861 | 18 | 0.055658864 | positive regulation of cardiac muscle hypertrophy |
| **GO:0030225** | 0.055658864 | 1.653975382 | 0.055658864 | 0.882742861 | 18 | 0.055658864 | macrophage differentiation |
| **GO:0016605** | 0.05640274 | 1.597022277 | 0.05640274 | 0.882742861 | 75 | 0.05640274 | PML body |
| **GO:0047485** | 0.05640274 | 1.597022277 | 0.05640274 | 0.882742861 | 75 | 0.05640274 | protein N-terminus binding |
| **GO:0016032** | 0.056632923 | 1.585474999 | 0.056632923 | 0.882742861 | 409 | 0.056632923 | viral process |
| **GO:0001568** | 0.056925897 | 1.601299583 | 0.056925897 | 0.882742861 | 45 | 0.056925897 | blood vessel development |
| **GO:0060079** | 0.056925897 | 1.601299583 | 0.056925897 | 0.882742861 | 45 | 0.056925897 | excitatory postsynaptic potential |
| **GO:0006366** | 0.057722152 | 1.579602921 | 0.057722152 | 0.882742861 | 143 | 0.057722152 | transcription by RNA polymerase II |
| **GO:0035925** | 0.058931135 | 1.618762609 | 0.058931135 | 0.882742861 | 19 | 0.058931135 | mRNA 3'-UTR AU-rich region binding |
| **GO:2001237** | 0.059288015 | 1.589894255 | 0.059288015 | 0.882742861 | 32 | 0.059288015 | negative regulation of extrinsic apoptotic signaling pathway |
| **GO:0006606** | 0.061410898 | 1.556106527 | 0.061410898 | 0.882742861 | 62 | 0.061410898 | protein import into nucleus |
| **GO:0021987** | 0.061410898 | 1.556106527 | 0.061410898 | 0.882742861 | 62 | 0.061410898 | cerebral cortex development |
| **GO:0005844** | 0.062000632 | 1.565189263 | 0.062000632 | 0.882742861 | 33 | 0.062000632 | polysome |
| **GO:0035235** | 0.062271268 | 1.585149662 | 0.062271268 | 0.882742861 | 20 | 0.062271268 | ionotropic glutamate receptor signaling pathway |
| **GO:0032281** | 0.062271268 | 1.585149662 | 0.062271268 | 0.882742861 | 20 | 0.062271268 | AMPA glutamate receptor complex |
| **GO:0043401** | 0.062271268 | 1.585149662 | 0.062271268 | 0.882742861 | 20 | 0.062271268 | steroid hormone mediated signaling pathway |
| **GO:0034063** | 0.062271268 | 1.585149662 | 0.062271268 | 0.882742861 | 20 | 0.062271268 | stress granule assembly |
| **GO:0030878** | 0.062271268 | 1.585149662 | 0.062271268 | 0.882742861 | 20 | 0.062271268 | thyroid gland development |
| **GO:0009954** | 0.062271268 | 1.585149662 | 0.062271268 | 0.882742861 | 20 | 0.062271268 | proximal/distal pattern formation |
| **GO:0006397** | 0.062323482 | 1.538055651 | 0.062323482 | 0.882742861 | 277 | 0.062323482 | mRNA processing |
| **GO:0030666** | 0.063885089 | 1.539900431 | 0.063885089 | 0.882742861 | 48 | 0.063885089 | endocytic vesicle membrane |
| **GO:0008285** | 0.064263651 | 1.52204015 | 0.064263651 | 0.882742861 | 318 | 0.064263651 | negative regulation of cell population proliferation |
| **GO:0031982** | 0.064271817 | 1.525361642 | 0.064271817 | 0.882742861 | 130 | 0.064271817 | vesicle |
| **GO:0019904** | 0.065540975 | 1.513531618 | 0.065540975 | 0.882742861 | 185 | 0.065540975 | protein domain specific binding |
| **GO:0055038** | 0.065638177 | 1.520976774 | 0.065638177 | 0.882742861 | 64 | 0.065638177 | recycling endosome membrane |
| **GO:0005104** | 0.065682196 | 1.552896633 | 0.065682196 | 0.882742861 | 21 | 0.065682196 | fibroblast growth factor receptor binding |
| **GO:0009880** | 0.065682196 | 1.552896633 | 0.065682196 | 0.882742861 | 21 | 0.065682196 | embryonic pattern specification |

**Table S17 - 38 species with CAMK2N2 transcripts downloaded from NCBI.** The last column regards the level of association with humans.
W = wild; D = domestic; S = synanthropic.

| Gene symbol | Gene ID | Description | Scientific name | Common name | RefSeq Transcri | RefSeq Protein | Association |
|---|---|---|---|---|---|---|---|
| **CAMK2N2** | 101747380 | calcium/calmodu | *Gallus gallus* | Chicken | XM_015291551.: | XP_015147037.1 | D |
| **CAMK2N2** | 102107607 | calcium/calmodu | *Pseudopodoces* | Tibetan ground-ti | XM_005525032. | XP_005525089.1 | W |
| **CAMK2N2** | 103528144 | calcium/calmodu | *Calypte anna* | Anna's humming | XM_030456215. | XP_030312075.1 | W |
| **CAMK2N2** | 104057870 | calcium/calmodu | *Cuculus canorus* | Common cuckoo | XM_009559309. | XP_009557604.1 | W |
| **CAMK2N2** | 105759523 | calcium/calmodu | *Taeniopygia gutta* | Zebra finch | XM_041718084. | XP_041574018.1 | D |
| **CAMK2N2** | 106850415 | calcium/calmodu | *Sturnus vulgaris* | Common starling | XM_014871707. | XP_014727193.1 | S |
| **CAMK2N2** | 107208613 | calcium/calmodu | *Parus major* | Great Tit | XM_015637111.2 | XP_015492597.1 | S |
| **CAMK2N2** | 107318230 | calcium/calmodu | *Coturnix japonica* | Japanese quail | XM_015871828.: | XP_015727314.1 | D |
| **CAMK2N2** | 108495276 | calcium/calmodu | *Lepidothrix coror* | Blue-crowned ma | XM_017810817. | XP_017666306.1 | W |
| **CAMK2N2** | 110397805 | calcium/calmodu | *Numida meleagri* | Helmeted guinea | XM_021394645. | XP_021250320.1 | D |
| **CAMK2N2** | 110477863 | calcium/calmodu | *Lonchura striata* | Bengalese finch | XM_021544020.: | XP_021399695.1 | D |
| **CAMK2N2** | 113844458 | calcium/calmodu | *Anas platyrhynch* | Mallard | XM_027464230.: | XP_027320031.1 | D |
| **CAMK2N2** | 113949204 | calcium/calmodu | *Corapipo altera* | White-ruffed mar | XM_027647498. | XP_027503299.1 | W |
| **CAMK2N2** | 113968780 | calcium/calmodu | *Neopelma chryso* | Saffron-crested t | XM_027685656. | XP_027541457.1 | W |
| **CAMK2N2** | 114003775 | calcium/calmodu | *Pipra filicauda* | Wire-tailed mana | XM_027751109.2 | XP_027606910.1 | W |
| **CAMK2N2** | 114060775 | calcium/calmodu | *Empidonax traillii* | Willow flycatcher | XM_027890551. | XP_027746352.1 | W |
| **CAMK2N2** | 115347171 | calcium/calmodu | *Aquila chrysaeto* | Goled eagle | XM_030028241.: | XP_029884101.1 | W |
| **CAMK2N2** | 115611213 | calcium/calmodu | *Strigops habropti* | Kakapo | XM_030493772. | XP_030349632.1 | W |
| **CAMK2N2** | 115907087 | calcium/calmodu | *Camarhynchus p* | Small tree finch | XM_030954740. | XP_030810600.1 | W |
| **CAMK2N2** | 116235402 | calcium/calmodu | *Phasianus colchi* | Ring-necked phe | XM_031603550. | XP_031459410.1 | D |
| **CAMK2N2** | 116448661 | calcium/calmodu | *Corvus monedulo* | New Caledonian | XM_032119428.1 | XP_031975319.1 | W |
| **CAMK2N2** | 116492425 | calcium/calmodu | *Aythya fuligula* | Tufted duck | XM_032193158. | XP_032049049.1 | W |
| **CAMK2N2** | 116791811 | calcium/calmodu | *Chiroxiphia lance* | Lance-tailed mar | XM_032698173. | XP_032554064.1 | W |
| **CAMK2N2** | 117000567 | calcium/calmodu | *Catharus ustulatu* | Swainson's thrus | XM_033068263. | XP_032924154.1 | W |
| **LOC117436255** | 117436255 | calcium/calmodu | *Melopsittacus un* | Budgerigar | XM_034063749. | XP_033919640.1 | D |
| **CAMK2N2** | 118157648 | calcium/calmodu | *Oxyura jamaicen* | Ruddy duck | XM_035312059. | XP_035167950.1 | W |
| **CAMK2N2** | 118247211 | calcium/calmodu | *Cygnus atratus* | Black swan | XM_035545010. | XP_035400903.1 | S |
| **CAMK2N2** | 118690048 | calcium/calmodu | *Molothrus ater* | Brown-headed co | XM_036388723. | XP_036244616.1 | W |

| | | | | | | | |
|---|---|---|---|---|---|---|---|
| **CAMK2N2** | 119157061 | calcium/calmodu | *Falco rusticolus* | Gyrfalcon | XM_037407595.1 | XP_037263492.1 | W |
| **CAMK2N2** | 119704357 | calcium/calmodu | *Motacilla alba alb* | White wagtail | XM_038145455.1 | XP_038001383.1 | S |
| **CAMK2N2** | 120410472 | calcium/calmodu | *Corvus cornix co* | Hooded crow | XM_039557048.1 | XP_039412982.1 | S |
| **CAMK2N2** | 120504284 | calcium/calmodu | *Passer montanus* | Eurasian tree spa | XM_039713643.1 | XP_039569577.1 | S |
| **CAMK2N2** | 120757412 | calcium/calmodu | *Hirundo rustica* | Barn swallow | XM_040074724.1 | XP_039930658.1 | S |
| **CAMK2N2** | 121075074 | calcium/calmodu | *Cygnus olor* | Mute swan | XM_040567937.1 | XP_040423871.1 | S |
| **CAMK2N2** | 121096835 | calcium/calmodu | *Falco naumanni* | Lesser kestrel | XM_040613367.1 | XP_040469301.1 | S |
| **CAMK2N2** | 121336579 | calcium/calmodu | *Onychostruthus t* | White-rumped sn | XM_041405897.1 | XP_041261831.1 | W |
| **CAMK2N2** | 121353747 | calcium/calmodu | *Pyrgilauda rufico* | Rufous-necked s | XM_041467857.1 | XP_041323791.1 | W |
| **CAMK2N2** | 121671265 | calcium/calmodu | *Corvus kubaryi* | Mariana crow | XM_042042637.1 | XP_041898571.1 | W |

**Table S18 - PhastCons conserved genes.** The top 5% (606) genes with more PhastCons conserved bases overlapped with CDS are reported. "LOC" genes were discarded.

| Gene | Chr | Start | End | Strand | Gene length | CDS length | Overlaps | % |
|---|---|---|---|---|---|---|---|---|
| AKIRIN2 | 3 | 80184991 | 80201419 | + | 16428 | 582 | 582 | 100 |
| AMMECR1L | 10 | 3876509 | 3891460 | + | 14951 | 933 | 933 | 100 |
| ARPP19 | 13 | 8188977 | 8200667 | + | 11690 | 339 | 339 | 100 |
| ASF1A | 3 | 66102983 | 66113750 | - | 10767 | 615 | 615 | 100 |
| ATP6V1G1 | 20 | 8540747 | 8544261 | - | 3514 | 357 | 357 | 100 |
| BARX1 | 12 | 661408 | 664565 | - | 3157 | 741 | 741 | 100 |
| BRK1 | 12 | 3530071 | 3532504 | - | 2433 | 228 | 228 | 100 |
| BTG1 | 4 | 45765041 | 45767873 | - | 2832 | 513 | 513 | 100 |
| C15H16orf72 | 15 | 5388789 | 5405607 | + | 16818 | 822 | 822 | 100 |
| C6H14orf132 | 6 | 14320525 | 14361426 | - | 40901 | 282 | 282 | 100 |
| CALM1 | 6 | 16973890 | 16985007 | - | 11117 | 450 | 450 | 100 |
| CALM2 | 3 | 28995618 | 29009091 | - | 13473 | 450 | 450 | 100 |
| CAMK2N2 | 10 | 17272192 | 17276215 | - | 4023 | 240 | 240 | 100 |
| CBLN1 | 11 | 13008843 | 13011974 | + | 3131 | 585 | 585 | 100 |
| CBX3 | 1 | 1.23E+08 | 1.23E+08 | - | 14588 | 734 | 734 | 100 |
| CEBPA | 11 | 20213662 | 20216548 | - | 2886 | 978 | 978 | 100 |
| CITED2 | 3 | 52194530 | 52197469 | - | 2939 | 720 | 720 | 100 |
| CLTB | 14 | 3584132 | 3590564 | + | 6432 | 684 | 684 | 100 |
| CNEP1R1 | 11 | 12571111 | 12577208 | - | 6097 | 378 | 378 | 100 |
| CSNK2A2 | 11 | 6258118 | 6286909 | + | 28791 | 1053 | 1053 | 100 |
| CTXN2 | 13 | 9574525 | 9579695 | + | 5170 | 246 | 246 | 100 |
| CXCL12 | 8 | 18803990 | 18822218 | - | 18228 | 412 | 412 | 100 |
| CXXC5 | 14 | 15120058 | 15156646 | + | 36588 | 831 | 831 | 100 |
| DEXI | 15 | 6197355 | 6205811 | - | 8456 | 282 | 282 | 100 |
| DIO3 | 6 | 11134171 | 11135728 | - | 1557 | 825 | 825 | 100 |
| DLX6 | 1 | 1.31E+08 | 1.31E+08 | - | 3740 | 807 | 807 | 100 |

| | | | | | | | | |
|---|---|---|---|---|---|---|---|---|
| **DNAJC5** | 16 | 6736324 | 6767203 | - | 30879 | 597 | 597 | 100 |
| **DR1** | 9 | 16934471 | 16945634 | + | 11163 | 531 | 531 | 100 |
| **DYNLL2** | 19 | 1155131 | 1162074 | + | 6943 | 423 | 423 | 100 |
| **EIF1AX** | 2 | 1.04E+08 | 1.04E+08 | - | 8909 | 435 | 435 | 100 |
| **EIF1B** | 1 | 1.11E+08 | 1.11E+08 | - | 8314 | 342 | 342 | 100 |
| **EIF4E** | 5 | 49946872 | 49971271 | + | 24399 | 654 | 654 | 100 |
| **EIF5A2** | 10 | 21670550 | 21675396 | + | 4846 | 462 | 462 | 100 |
| **ELOC** | 1 | 32226747 | 32240793 | + | 14046 | 339 | 339 | 100 |
| **EMC6** | 19 | 3356712 | 3357521 | + | 809 | 333 | 333 | 100 |
| **ERH** | 6 | 3.11E+07 | 3.11E+07 | + | 6749 | 315 | 315 | 100 |
| **FKBP1B** | 3 | 110115241 | 110161017 | + | 45776 | 327 | 327 | 100 |
| **FOXN3** | 6 | 17314932 | 17531563 | + | 216631 | 1557 | 1557 | 100 |
| **GABARAPL2** | 11 | 16938 | 21659 | + | 4721 | 354 | 354 | 100 |
| **GDNF** | Z | 17977614 | 17995206 | + | 17592 | 730 | 730 | 100 |
| **GET4** | 15 | 10886392 | 10898338 | + | 11946 | 1062 | 1062 | 100 |
| **GID8** | 16 | 5489358 | 5496778 | + | 7420 | 687 | 687 | 100 |
| **GJD2** | 6 | 35675902 | 35680651 | + | 4749 | 915 | 915 | 100 |
| **GNG10** | Z | 65353743 | 65359998 | - | 6255 | 207 | 207 | 100 |
| **GPR27** | 12 | 17192317 | 17194535 | + | 2218 | 1143 | 1143 | 100 |
| **GSK3B** | 2 | 26324625 | 26474255 | - | 149630 | 1302 | 1302 | 100 |
| **HDGFL3** | 13 | 10744841 | 10782483 | - | 37642 | 612 | 612 | 100 |
| **HMGB3** | 21 | 7098181 | 7104576 | + | 6395 | 609 | 609 | 100 |
| **HOMER1** | Z | 7.32E+06 | 7.41E+06 | - | 87746 | 1068 | 1068 | 100 |
| **HOXA13** | 1 | 1.22E+08 | 1.22E+08 | + | 2768 | 918 | 918 | 100 |
| **HOXA3** | 1 | 1.22E+08 | 1.22E+08 | + | 19844 | 1242 | 1242 | 100 |
| **HOXA5** | 1 | 1.22E+08 | 1.22E+08 | + | 5466 | 1199 | 1199 | 100 |
| **HOXA7** | 1 | 122393042 | 122396465 | + | 3423 | 660 | 660 | 100 |
| **INHBB** | 7 | 12006466 | 12012818 | - | 6352 | 1179 | 1179 | 100 |
| **INSIG2** | 7 | 8377879 | 8391216 | + | 13337 | 777 | 777 | 100 |

| | | | | | | | | |
|---|---|---|---|---|---|---|---|---|
| **ISL1** | Z | 14482062 | 14493337 | - | 11275 | 1334 | 1334 | 100 |
| **KRAS** | 4 | 67495741 | 67521352 | + | 25611 | 567 | 567 | 100 |
| **LDB2** | 5 | 14892081 | 15266783 | - | 374702 | 1833 | 1833 | 100 |
| **LGI1** | 8 | 19999221 | 20028733 | - | 29512 | 1658 | 1658 | 100 |
| **LHFPL3** | 4 | 12615089 | 12859349 | + | 244260 | 752 | 752 | 100 |
| **LHX9** | 9 | 22281366 | 22301066 | - | 19700 | 1721 | 1721 | 100 |
| **LMO1** | 6 | 51979391 | 52053222 | - | 73831 | 699 | 699 | 100 |
| **LMO3** | 4 | 72850450 | 72909594 | + | 59144 | 677 | 677 | 100 |
| **LMO4** | 9 | 1.50E+07 | 1.51E+07 | + | 15239 | 498 | 498 | 100 |
| **LRATD1** | 3 | 104301310 | 104303211 | + | 1901 | 855 | 855 | 100 |
| **LYSMD2** | 13 | 8583775 | 8595154 | + | 11379 | 588 | 588 | 100 |
| **MAB21L2** | 5 | 59173501 | 59176147 | + | 2646 | 1080 | 1080 | 100 |
| **MAPRE2** | 1 | 42961763 | 43061218 | - | 99455 | 1187 | 1187 | 100 |
| **MARCHF4** | 7 | 14264113 | 14359677 | - | 95564 | 1173 | 1173 | 100 |
| **MEIS2** | 6 | 34504387 | 34679190 | + | 174803 | 1407 | 1407 | 100 |
| **MOSMO** | 15 | 9113188 | 9144197 | + | 31009 | 504 | 504 | 100 |
| **MTPN** | 4 | 65623757 | 65655159 | + | 31402 | 357 | 357 | 100 |
| **NKX2-1** | 6 | 25028226 | 25030187 | + | 1961 | 1038 | 1038 | 100 |
| **NOG** | 18 | 7599566 | 7601149 | + | 1583 | 672 | 672 | 100 |
| **NOVA1** | 6 | 3.72E+07 | 3.74E+07 | + | 144264 | 1524 | 1524 | 100 |
| **NPY** | 1 | 123694226 | 123703124 | - | 8898 | 294 | 294 | 100 |
| **NR2F2** | 13 | 15687702 | 15703513 | + | 15811 | 1355 | 1355 | 100 |
| **NSG1** | 5 | 12516550 | 12539395 | - | 22845 | 558 | 558 | 100 |
| **NTMT1** | 20 | 5.20E+06 | 5.20E+06 | - | 3194 | 672 | 672 | 100 |
| **NXPH1** | 1 | 130117260 | 130256787 | - | 139527 | 816 | 816 | 100 |
| **OSR2** | 1 | 2.21E+07 | 2.21E+07 | - | 5723 | 1014 | 1014 | 100 |
| **OST4** | 3 | 1.16E+08 | 1.16E+08 | + | 605 | 114 | 114 | 100 |
| **PCP4** | 2 | 115700657 | 115748854 | - | 48197 | 189 | 189 | 100 |
| **PDCD10** | 10 | 22673331 | 22688081 | + | 14750 | 639 | 639 | 100 |

| | | | | | | | | |
|---|---|---|---|---|---|---|---|---|
| **PDE6D** | 10 | 16616442 | 16650828 | - | 34386 | 453 | 453 | 100 |
| **PHLDA2** | 6 | 48666753 | 48668266 | - | 1513 | 399 | 399 | 100 |
| **PLN** | 3 | 66253922 | 66262968 | - | 9046 | 159 | 159 | 100 |
| **POLR2K** | 1 | 21467464 | 21471174 | - | 3710 | 177 | 177 | 100 |
| **PPP1R14C** | 3 | 57974369 | 58026240 | - | 51871 | 477 | 477 | 100 |
| **PPP2R5E** | 6 | 6663369 | 6734427 | - | 71058 | 1530 | 1530 | 100 |
| **PPP3R1** | 3 | 3919244 | 3957069 | - | 37825 | 513 | 513 | 100 |
| **PTP4A1** | 3 | 90050456 | 90064436 | - | 13980 | 522 | 522 | 100 |
| **PURA** | 14 | 15545390 | 15546352 | + | 962 | 951 | 951 | 100 |
| **RAB14** | 20 | 2016710 | 2030936 | + | 14226 | 648 | 648 | 100 |
| **RAB21** | 4 | 36767672 | 36782135 | + | 14463 | 663 | 663 | 100 |
| **RAB4A** | 3 | 43669319 | 43688097 | - | 18778 | 659 | 659 | 100 |
| **RAC3** | 18 | 11911414 | 11916654 | + | 5240 | 579 | 579 | 100 |
| **RGS17** | 3 | 56614450 | 56687256 | + | 72806 | 711 | 711 | 100 |
| **RHOQ** | 3 | 28538252 | 28554734 | + | 16482 | 910 | 910 | 100 |
| **RNF7** | 10 | 11010595 | 11017107 | + | 6512 | 573 | 573 | 100 |
| **RPL39** | W | 23521796 | 23524725 | + | 2929 | 156 | 156 | 100 |
| **RPS28** | 26 | 4836071 | 4836885 | + | 814 | 210 | 210 | 100 |
| **SFRP2** | 5 | 41438636 | 41443023 | - | 4387 | 879 | 879 | 100 |
| **SIX6** | 6 | 5538951 | 5541086 | + | 2135 | 741 | 741 | 100 |
| **SNAP25** | 3 | 6648711 | 6710223 | - | 61512 | 621 | 621 | 100 |
| **SNX6** | 6 | 25824345 | 25853005 | + | 28660 | 1272 | 1272 | 100 |
| **SOX2** | 10 | 18396452 | 18484266 | - | 87814 | 948 | 948 | 100 |
| **SPTSSA** | 6 | 25863945 | 25868825 | + | 4880 | 207 | 207 | 100 |
| **SRSF1** | 19 | 1137703 | 1141215 | - | 3512 | 747 | 747 | 100 |
| **SST** | 10 | 15940068 | 15941623 | + | 1555 | 351 | 351 | 100 |
| **STMN2** | 1 | 29576326 | 29618188 | - | 41862 | 540 | 540 | 100 |
| **SUMO2** | 18 | 4890590 | 4897641 | + | 7051 | 288 | 288 | 100 |
| **TAFA2** | 4 | 32877891 | 33069515 | - | 191624 | 396 | 396 | 100 |

| | | | | | | | | |
|---|---|---|---|---|---|---|---|---|
| **TBR1** | 7 | 16655748 | 16661953 | + | 6205 | 2034 | 2034 | 100 |
| **TBX15** | 2 | 27669565 | 27723652 | - | 54087 | 420 | 420 | 100 |
| **TENT5A** | 3 | 82352397 | 82357478 | + | 5081 | 1203 | 1203 | 100 |
| **THY1** | 23 | 5132406 | 5137789 | + | 5383 | 486 | 486 | 100 |
| **TIAL1** | 8 | 30684506 | 30705020 | - | 20514 | 1268 | 1268 | 100 |
| **TIMM8A** | 21 | 3966100 | 3967313 | + | 1213 | 285 | 285 | 100 |
| **TLNRD1** | 13 | 1.18E+07 | 1.18E+07 | - | 2060 | 1068 | 1068 | 100 |
| **TMSB4X** | 2 | 100515905 | 100517440 | + | 1535 | 135 | 135 | 100 |
| **TPT1** | 2 | 55992797 | 55997941 | + | 5144 | 519 | 519 | 100 |
| **TRAPPC6B** | 6 | 24009123 | 24014535 | + | 5412 | 477 | 477 | 100 |
| **TRIQK** | 1 | 24495251 | 24555837 | + | 60586 | 273 | 273 | 100 |
| **UBE2A** | W | 23662542 | 23671877 | - | 9335 | 459 | 459 | 100 |
| **UBE2D3** | 5 | 48896664 | 48919750 | + | 23086 | 494 | 494 | 100 |
| **UBE2G1** | 19 | 7293527 | 7318497 | - | 24970 | 513 | 513 | 100 |
| **UBE2I** | 15 | 16484909 | 16497500 | + | 12591 | 477 | 477 | 100 |
| **UBE2N** | 4 | 46253511 | 46273858 | - | 20347 | 459 | 459 | 100 |
| **UBE2Q2** | 13 | 711991 | 757546 | + | 45555 | 1221 | 1221 | 100 |
| **VKORC1L1** | 19 | 5273680 | 5293202 | - | 19522 | 531 | 531 | 100 |
| **YBX1** | 22 | 2351563 | 2360876 | - | 9313 | 1210 | 1210 | 100 |
| **YPEL1** | 17 | 7951831 | 7973817 | - | 21986 | 360 | 360 | 100 |
| **YWHAB** | 16 | 9029975 | 9043876 | + | 13901 | 735 | 735 | 100 |
| **YWHAE** | 19 | 4976934 | 4997696 | - | 20762 | 776 | 776 | 100 |
| **YWHAZ** | 1 | 21171799 | 21197840 | + | 26041 | 738 | 738 | 100 |
| **YY1** | 6 | 11938783 | 11964238 | - | 25455 | 1263 | 1263 | 100 |
| **ZBTB18** | 3 | 33832044 | 33838694 | + | 6650 | 3165 | 3165 | 100 |
| **ZNF706** | 1 | 2.11E+07 | 2.11E+07 | + | 7096 | 231 | 231 | 100 |
| **TFAP2B** | 3 | 113055613 | 113089530 | + | 33917 | 1711 | 1710 | 99.94155465 |
| **YTHDF3** | 1 | 36523306 | 36546688 | - | 23382 | 3327 | 3325 | 99.93988578 |
| **COPS2** | 13 | 9916997 | 9938782 | - | 21785 | 1443 | 1442 | 99.93069993 |

| | | | | | | | | |
|---|---|---|---|---|---|---|---|---|
| **FEZF2** | 12 | 13549436 | 13552515 | - | 3079 | 1311 | 1310 | 99.92372235 |
| **MAB21L1** | 2 | 5.08E+07 | 5.08E+07 | - | 1843 | 1080 | 1079 | 99.90740741 |
| **TRIB2** | 3 | 103472213 | 103492212 | + | 19999 | 1032 | 1031 | 99.90310078 |
| **FGF12** | 10 | 14298120 | 14515264 | + | 217144 | 833 | 832 | 99.87995198 |
| **SRD5A2** | 3 | 35017764 | 35043358 | - | 25594 | 768 | 767 | 99.86979167 |
| **ZFAND3** | 3 | 31682187 | 31817158 | - | 134971 | 739 | 738 | 99.864682 |
| **ZNRF1** | 11 | 7959806 | 7979996 | + | 20190 | 705 | 704 | 99.85815603 |
| **RGS7** | 3 | 4.78E+07 | 4.80E+07 | - | 198758 | 1329 | 1327 | 99.84951091 |
| **SEPTIN7** | 1 | 108806546 | 108887445 | - | 80899 | 1290 | 1288 | 99.84496124 |
| **BMP4** | 6 | 3028195 | 3031297 | - | 3102 | 1212 | 1210 | 99.8349835 |
| **MYT1L** | 3 | 98124354 | 98443495 | - | 319141 | 4721 | 4713 | 99.83054438 |
| **SPOPL** | 7 | 5.47E+06 | 5.50E+06 | - | 33158 | 1179 | 1177 | 99.83036472 |
| **KDM6A** | 2 | 112861854 | 113012313 | + | 150459 | 4416 | 4408 | 99.81884058 |
| **SNX3** | 3 | 70888833 | 70906704 | + | 17871 | 549 | 548 | 99.81785064 |
| **UBE2B** | 14 | 9620420 | 9632664 | - | 12244 | 459 | 458 | 99.78213508 |
| **CELF2** | 4 | 5085531 | 5460683 | + | 375152 | 2257 | 2252 | 99.77846699 |
| **TLX1** | 8 | 23042589 | 23049596 | - | 7007 | 894 | 892 | 99.77628635 |
| **MXD4** | 5 | 7946730 | 7987861 | - | 41131 | 884 | 882 | 99.77375566 |
| **COPS3** | 15 | 13849170 | 13862796 | - | 13626 | 1323 | 1320 | 99.77324263 |
| **TOMM20** | 3 | 45572312 | 45581669 | - | 9357 | 441 | 440 | 99.77324263 |
| **HNRNPA3** | 7 | 22335551 | 22352379 | + | 16828 | 1290 | 1287 | 99.76744186 |
| **MARCKS** | 3 | 68243346 | 68248426 | - | 5080 | 855 | 853 | 99.76608187 |
| **CDK8** | 2 | 47191284 | 47265484 | + | 74200 | 1678 | 1674 | 99.76162098 |
| **CISD2** | 5 | 48875145 | 48882631 | - | 7486 | 405 | 404 | 99.75308642 |
| **USP22** | 15 | 13645229 | 13736892 | + | 91663 | 1578 | 1574 | 99.74651458 |
| **RAB12** | 1 | 5.19E+07 | 5.20E+07 | + | 23950 | 753 | 751 | 99.73439575 |
| **SP8** | 1 | 124975955 | 124978644 | + | 2689 | 1500 | 1496 | 99.73333333 |
| **PLAG1** | 1 | 39228236 | 39281411 | + | 53175 | 2903 | 2895 | 99.72442301 |
| **PSMD14** | 7 | 16605102 | 16651572 | + | 46470 | 933 | 930 | 99.67845659 |

| | | | | | | | | |
|---|---|---|---|---|---|---|---|---|
| **NFIA** | 9 | 2692957 | 2937661 | - | 244704 | 2112 | 2105 | 99.66856061 |
| **ACTB** | 15 | 13177735 | 13182574 | - | 4839 | 1128 | 1124 | 99.64539007 |
| **CACNG2** | 4 | 54100575 | 54152607 | + | 52032 | 1114 | 1110 | 99.64093357 |
| **EPHX4** | 9 | 1.66E+07 | 1.66E+07 | + | 16783 | 1104 | 1100 | 99.63768116 |
| **PTF1A** | 1 | 1.39E+08 | 1.39E+08 | + | 1195 | 828 | 825 | 99.63768116 |
| **HACD1** | 1 | 135923638 | 135941747 | - | 18109 | 824 | 821 | 99.63592233 |
| **ADSS1** | 6 | 8636556 | 8661775 | - | 25219 | 1368 | 1363 | 99.63450292 |
| **ZIC1** | 10 | 12735229 | 12739550 | + | 4321 | 1338 | 1333 | 99.62630792 |
| **EBF3** | 8 | 34778243 | 34900868 | - | 122625 | 1854 | 1847 | 99.62243797 |
| **POU3F3** | 2 | 90105036 | 90106371 | + | 1335 | 1323 | 1318 | 99.62207105 |
| **RUNX1T1** | 1 | 2.49E+07 | 2.50E+07 | + | 114009 | 1965 | 1957 | 99.59287532 |
| **JAZF1** | 1 | 121831475 | 122030926 | + | 199451 | 732 | 729 | 99.59016393 |
| **CZIB** | 9 | 5522137 | 5527465 | + | 5328 | 483 | 481 | 99.58592133 |
| **RPRD1A** | 1 | 67664023 | 67708596 | - | 44573 | 948 | 944 | 99.57805907 |
| **PCMT1** | 3 | 58098778 | 58149566 | - | 50788 | 940 | 936 | 99.57446809 |
| **ING3** | 4 | 23013765 | 23030358 | - | 16593 | 1611 | 1604 | 99.56548727 |
| **ZNF503** | 8 | 14261308 | 14265440 | + | 4132 | 1830 | 1822 | 99.56284153 |
| **PITX1** | 14 | 9335995 | 9342664 | + | 6669 | 909 | 905 | 99.559956 |
| **TCF7L2** | 8 | 27339850 | 27513391 | + | 173541 | 2234 | 2224 | 99.55237243 |
| **CRYBB2** | 17 | 4511001 | 4513714 | + | 2713 | 850 | 846 | 99.52941176 |
| **MTA1** | 9 | 27267133 | 27342296 | - | 75163 | 2329 | 2318 | 99.52769429 |
| **FOXP2** | 4 | 25889103 | 26292811 | - | 403708 | 2202 | 2191 | 99.50045413 |
| **EMC3** | 12 | 3.57E+06 | 3.57E+06 | + | 5164 | 786 | 782 | 99.49109415 |
| **AP1S2** | 2 | 101961912 | 101992868 | - | 30956 | 564 | 561 | 99.46808511 |
| **PSMA6** | 6 | 25560958 | 25572608 | - | 11650 | 741 | 737 | 99.46018893 |
| **TBPL1** | 3 | 59409023 | 59423199 | - | 14176 | 552 | 549 | 99.45652174 |
| **RAB6A** | 2 | 1062742 | 1105266 | + | 42524 | 733 | 729 | 99.45429741 |
| **CPSF4** | 15 | 13434492 | 13440670 | + | 6178 | 732 | 728 | 99.45355191 |
| **SPCS3** | 5 | 35328745 | 35336625 | - | 7880 | 543 | 540 | 99.44751381 |

| | | | | | | | | |
|---|---|---|---|---|---|---|---|---|
| **PHF6** | Z | 84394549 | 84421116 | + | 26567 | 1262 | 1255 | 99.44532488 |
| **UBL3** | 2 | 4.86E+07 | 4.86E+07 | - | 54337 | 354 | 352 | 99.43502825 |
| **DYNLT3** | 2 | 110600432 | 110607724 | - | 7292 | 351 | 349 | 99.43019943 |
| **FBXO33** | 6 | 23936058 | 23952910 | + | 16852 | 1572 | 1563 | 99.42748092 |
| **CDC42** | 22 | 2296099 | 2323239 | - | 27140 | 666 | 662 | 99.3993994 |
| **FBXO45** | 10 | 4650597 | 4655757 | + | 5160 | 819 | 814 | 99.38949939 |
| **PDGFA** | 15 | 10669433 | 10705721 | - | 36288 | 781 | 776 | 99.35979513 |
| **HNRNPA0** | 14 | 8082636 | 8085513 | + | 2877 | 936 | 930 | 99.35897436 |
| **GNA11** | 26 | 2525800 | 2540522 | + | 14722 | 1080 | 1073 | 99.35185185 |
| **PHF21A** | 6 | 39042140 | 39176524 | - | 134384 | 2150 | 2136 | 99.34883721 |
| **UST** | 3 | 48362068 | 48516345 | - | 154277 | 1224 | 1216 | 99.34640523 |
| **CSNK1A1** | 14 | 1594299 | 1626748 | + | 32449 | 1365 | 1356 | 99.34065934 |
| **SOX11** | 3 | 99835902 | 99838360 | + | 2458 | 1212 | 1204 | 99.33993399 |
| **HAS2** | 1 | 12218128 | 12237662 | + | 19534 | 1662 | 1651 | 99.33814681 |
| **CHP1** | 6 | 38261112 | 38281337 | - | 20225 | 588 | 584 | 99.31972789 |
| **AMD1** | 3 | 69717400 | 69735354 | - | 17954 | 1026 | 1019 | 99.31773879 |
| **HLF** | 18 | 7.25E+06 | 7.28E+06 | + | 32385 | 879 | 873 | 99.31740614 |
| **VSNL1** | 3 | 105893252 | 105979663 | + | 86411 | 576 | 572 | 99.30555556 |
| **RFX3** | Z | 2677050 | 2777360 | + | 100310 | 2250 | 2234 | 99.28888889 |
| **DCUN1D5** | 2 | 41629451 | 41642470 | - | 13019 | 702 | 697 | 99.28774929 |
| **GPM6A** | 5 | 35493859 | 35609478 | + | 115619 | 837 | 831 | 99.28315412 |
| **PRR7** | 14 | 3304060 | 3305298 | + | 1238 | 1109 | 1101 | 99.2786294 |
| **RAP2B** | 10 | 25254870 | 25255666 | - | 796 | 552 | 548 | 99.27536232 |
| **TNNI2** | 6 | 4.76E+07 | 4.76E+07 | + | 5648 | 552 | 548 | 99.27536232 |
| **CUL1** | 1 | 106097884 | 106151605 | + | 53721 | 2331 | 2314 | 99.27069927 |
| **FGF13** | W | 28534725 | 28896347 | + | 361622 | 955 | 948 | 99.26701571 |
| **PDAP1** | 15 | 13418921 | 13427094 | - | 8173 | 543 | 539 | 99.26335175 |
| **CNOT6** | 14 | 7382440 | 7415325 | - | 32885 | 1764 | 1751 | 99.26303855 |
| **SOCS6** | 1 | 57335895 | 57360863 | + | 24968 | 1605 | 1593 | 99.25233645 |

| | | | | | | | | |
|---|---|---|---|---|---|---|---|---|
| **TPM1** | 13 | 3.69E+06 | 3.71E+06 | - | 19915 | 1426 | 1415 | 99.2286115 |
| **RAB5A** | 1 | 119067394 | 119084065 | - | 16671 | 648 | 643 | 99.22839506 |
| **TXNDC12** | 9 | 5937550 | 5947337 | + | 9787 | 513 | 509 | 99.2202729 |
| **WIPI2** | 15 | 13019440 | 13046353 | + | 26913 | 1409 | 1398 | 99.21930447 |
| **TOP1** | 16 | 9523803 | 9590792 | - | 66989 | 2301 | 2283 | 99.21773142 |
| **CTXN1** | 26 | 822548 | 842278 | - | 19730 | 255 | 253 | 99.21568627 |
| **CCNC** | 3 | 74990868 | 75007126 | + | 16258 | 891 | 884 | 99.21436588 |
| **ACTR3** | 7 | 7324738 | 7353436 | + | 28698 | 1257 | 1247 | 99.20445505 |
| **SS18** | 1 | 46574377 | 46618262 | + | 43885 | 1257 | 1247 | 99.20445505 |
| **FGF14** | 2 | 80195381 | 80588549 | - | 393168 | 995 | 987 | 99.1959799 |
| **MAPK1** | 17 | 7991427 | 8025421 | - | 33994 | 1107 | 1098 | 99.18699187 |
| **LINGO1** | 13 | 1478621 | 1617609 | - | 138988 | 3729 | 3698 | 99.16867793 |
| **MAFK** | 15 | 11391344 | 11405724 | + | 14380 | 471 | 467 | 99.1507431 |
| **JARID2** | 1 | 91469937 | 91680696 | - | 210759 | 3862 | 3829 | 99.14552046 |
| **KIAA0232** | 5 | 11125037 | 11190859 | - | 65822 | 4176 | 4140 | 99.13793103 |
| **SGCD** | 14 | 4744922 | 5060488 | - | 315566 | 924 | 916 | 99.13419913 |
| **SNAI2** | 1 | 42179073 | 42182758 | + | 3685 | 807 | 800 | 99.13258984 |
| **C1H6orf62** | 1 | 6.08E+07 | 6.08E+07 | - | 7716 | 690 | 684 | 99.13043478 |
| **SATB1** | 1 | 119612547 | 119705454 | + | 92907 | 2358 | 2337 | 99.10941476 |
| **LHX5** | 17 | 9794943 | 9802301 | + | 7358 | 1449 | 1436 | 99.10282954 |
| **SHOX** | 2 | 84713441 | 84724422 | - | 10981 | 876 | 868 | 99.08675799 |
| **CNNM2** | 8 | 23509459 | 23622954 | + | 113495 | 1836 | 1819 | 99.07407407 |
| **GRIA2** | 5 | 42856280 | 42943070 | + | 86790 | 3557 | 3524 | 99.0722519 |
| **GATA2** | 12 | 9.10E+06 | 9.12E+06 | + | 16486 | 1401 | 1388 | 99.07209136 |
| **PRPS2** | 2 | 100414318 | 100439515 | + | 25197 | 1070 | 1060 | 99.06542056 |
| **CSNK2A1** | 16 | 7266508 | 7289154 | + | 22646 | 1176 | 1165 | 99.06462585 |
| **EMX2** | 8 | 29698778 | 29704916 | + | 6138 | 744 | 737 | 99.05913978 |
| **RPL18A** | 4 | 33818880 | 33822443 | - | 3563 | 531 | 526 | 99.05838041 |
| **HDAC2** | 3 | 68174850 | 68199019 | + | 24169 | 1467 | 1453 | 99.04567144 |

| | | | | | | | | |
|---|---|---|---|---|---|---|---|---|
| **BRI3** | 15 | 9706672 | 9717345 | + | 10673 | 417 | 413 | 99.04076739 |
| **ZEB2** | 7 | 3.34E+06 | 3.46E+06 | + | 115175 | 3645 | 3610 | 99.03978052 |
| **RAB18** | 1 | 140288044 | 140302980 | + | 14936 | 624 | 618 | 99.03846154 |
| **DDX6** | 23 | 6999766 | 7020934 | + | 21168 | 1452 | 1438 | 99.03581267 |
| **CRABP1** | 13 | 502516 | 517550 | + | 15034 | 414 | 410 | 99.03381643 |
| **ZCCHC24** | 8 | 11823767 | 11931501 | + | 107734 | 720 | 713 | 99.02777778 |
| **CPLX2** | 14 | 3.62E+06 | 3.63E+06 | - | 15507 | 405 | 401 | 99.01234568 |
| **HOXA4** | 1 | 1.22E+08 | 1.22E+08 | + | 2374 | 909 | 900 | 99.00990099 |
| **PSMA2** | 1 | 103216268 | 103222508 | - | 6240 | 705 | 698 | 99.0070922 |
| **YAP1** | 2 | 41290171 | 41384995 | + | 94824 | 1706 | 1689 | 99.003517 |
| **MAP3K7** | 3 | 78899462 | 78948694 | + | 49232 | 2092 | 2071 | 98.99617591 |
| **RSL24D1** | 13 | 7447767 | 7454564 | + | 6797 | 492 | 487 | 98.98373984 |
| **SLC32A1** | 16 | 10793659 | 10797393 | + | 3734 | 1560 | 1544 | 98.97435897 |
| **SP3** | 7 | 20978900 | 21013491 | - | 34591 | 3577 | 3540 | 98.96561364 |
| **UBLCP1** | 14 | 3814716 | 3827054 | - | 12338 | 957 | 947 | 98.95506792 |
| **LHFPL4** | 12 | 1.16E+07 | 1.16E+07 | - | 12282 | 669 | 662 | 98.95366218 |
| **PTCHD1** | 2 | 105434425 | 105471431 | + | 37006 | 2667 | 2639 | 98.95013123 |
| **DSTN** | 3 | 13416947 | 13428638 | + | 11691 | 758 | 750 | 98.94459103 |
| **TPM2** | Z | 20549190 | 20562659 | - | 13469 | 1226 | 1213 | 98.93964111 |
| **CLDN5** | 17 | 7792227 | 7795970 | + | 3743 | 651 | 644 | 98.92473118 |
| **RAP2C** | W | 5024533 | 5034298 | - | 9765 | 552 | 546 | 98.91304348 |
| **TRPS1** | 1 | 1.48E+07 | 1.50E+07 | + | 209429 | 4684 | 4633 | 98.91118702 |
| **DYNC1LI1** | 1 | 114327592 | 114353318 | + | 25726 | 1645 | 1627 | 98.90577508 |
| **XPOT** | 4 | 33823561 | 33853160 | + | 29599 | 2889 | 2857 | 98.89235029 |
| **ISL2** | 13 | 922599 | 925814 | + | 3215 | 1080 | 1068 | 98.88888889 |
| **EHF** | 6 | 44757840 | 44788782 | - | 30942 | 897 | 887 | 98.8851728 |
| **PPP2R5A** | 3 | 23361139 | 23403739 | - | 42600 | 1431 | 1415 | 98.88190077 |
| **CEBPB** | 16 | 767615 | 769412 | + | 1797 | 978 | 967 | 98.87525562 |
| **ZFAND6** | 13 | 12146453 | 12182845 | - | 36392 | 799 | 790 | 98.87359199 |

| | | | | | | | | |
|---|---|---|---|---|---|---|---|---|
| **AP2M1** | 10 | 1.72E+07 | 1.72E+07 | + | 26699 | 1326 | 1311 | 98.86877828 |
| **STARD3NL** | 1 | 107881559 | 107905707 | - | 24148 | 705 | 697 | 98.86524823 |
| **VSTM2A** | 1 | 69125784 | 69153457 | - | 27673 | 878 | 868 | 98.86104784 |
| **ARIH2** | 12 | 11923393 | 11958666 | - | 35273 | 1473 | 1456 | 98.84589274 |
| **TLK1** | 7 | 19892769 | 19960106 | - | 67337 | 2415 | 2387 | 98.84057971 |
| **NAPG** | 1 | 52803563 | 52816096 | + | 12533 | 945 | 934 | 98.83597884 |
| **TAB2** | 3 | 4.82E+07 | 4.83E+07 | - | 62629 | 3490 | 3449 | 98.8252149 |
| **RHOB** | 3 | 107581626 | 107583994 | + | 2368 | 591 | 584 | 98.81556684 |
| **HOXD11** | 7 | 21855007 | 21856559 | + | 1552 | 843 | 833 | 98.81376038 |
| **GPBP1L1** | 9 | 9306394 | 9338494 | + | 32100 | 1416 | 1399 | 98.79943503 |
| **HNRNPR** | 25 | 940564 | 960997 | + | 20433 | 2393 | 2364 | 98.78813205 |
| **HAND2** | 5 | 48074331 | 48076611 | - | 2280 | 654 | 646 | 98.77675841 |
| **FOXG1** | 6 | 36548378 | 36550318 | - | 1940 | 1374 | 1357 | 98.76273654 |
| **BARHL1** | 20 | 4093179 | 4099201 | - | 6022 | 969 | 957 | 98.76160991 |
| **FRS2** | 4 | 35775715 | 35794363 | + | 18648 | 1527 | 1508 | 98.75573019 |
| **FGF10** | Z | 15616301 | 15676178 | + | 59877 | 639 | 631 | 98.74804382 |
| **TIMP3** | 4 | 55595165 | 55628812 | - | 33647 | 639 | 631 | 98.74804382 |
| **MXI1** | 8 | 2.61E+07 | 2.61E+07 | + | 55122 | 871 | 860 | 98.73708381 |
| **TMEM47** | 2 | 109861692 | 109881257 | - | 19565 | 546 | 539 | 98.71794872 |
| **PXDC1** | 1 | 86515482 | 86539852 | - | 24370 | 696 | 687 | 98.70689655 |
| **TCF21** | 3 | 59434057 | 59436921 | - | 2864 | 540 | 533 | 98.7037037 |
| **RAB8B** | 13 | 3621181 | 3650380 | - | 29199 | 845 | 834 | 98.69822485 |
| **PPP1CC** | 17 | 3396700 | 3411566 | - | 14866 | 972 | 959 | 98.66255144 |
| **PFN2** | 10 | 25652139 | 25657481 | + | 5342 | 521 | 514 | 98.65642994 |
| **FOXB1** | 13 | 4741025 | 4746439 | - | 5414 | 963 | 950 | 98.65005192 |
| **HOXD10** | 7 | 21864399 | 21867442 | + | 3043 | 1034 | 1020 | 98.64603482 |
| **MAP3K9** | 6 | 30246628 | 30308679 | + | 62051 | 3150 | 3107 | 98.63492063 |
| **NKIRAS2** | 27 | 2965405 | 2967551 | - | 2146 | 576 | 568 | 98.61111111 |
| **ATXN2** | 17 | 3548864 | 3598436 | - | 49572 | 3806 | 3753 | 98.6074619 |

| | | | | | | | | |
|---|---|---|---|---|---|---|---|---|
| **CHST2** | 10 | 1.14E+07 | 1.14E+07 | + | 4509 | 1290 | 1272 | 98.60465116 |
| **NMT2** | 1 | 134950491 | 134983007 | - | 32516 | 1497 | 1476 | 98.59719439 |
| **S100A1** | 29 | 1690770 | 1692999 | - | 2229 | 285 | 281 | 98.59649123 |
| **MED9** | 15 | 13896323 | 13900628 | + | 4305 | 354 | 349 | 98.58757062 |
| **INPP5A** | 8 | 3.57E+07 | 3.59E+07 | + | 182094 | 1326 | 1307 | 98.56711916 |
| **ASAP2** | 3 | 101454591 | 101537870 | + | 83279 | 3209 | 3163 | 98.56653163 |
| **SOX18** | 16 | 6583283 | 6587237 | + | 3954 | 1254 | 1236 | 98.5645933 |
| **BCL9** | 2 | 5312524 | 5370141 | + | 57617 | 5351 | 5274 | 98.56101663 |
| **BRINP1** | 20 | 6832399 | 6920105 | + | 87706 | 2286 | 2253 | 98.55643045 |
| **CNOT4** | 4 | 49627896 | 49706308 | - | 78412 | 2699 | 2660 | 98.55502038 |
| **RASA3** | 2 | 92138990 | 92270497 | + | 131507 | 2559 | 2522 | 98.5541227 |
| **NFIC** | 26 | 4453151 | 4492250 | + | 39099 | 1590 | 1567 | 98.55345912 |
| **PRLHR** | 8 | 30167891 | 30175005 | - | 7114 | 1035 | 1020 | 98.55072464 |
| **ATG13** | 6 | 39497039 | 39521028 | + | 23989 | 1647 | 1623 | 98.5428051 |
| **GTF2H1** | 6 | 49373893 | 49397950 | - | 24057 | 1647 | 1623 | 98.5428051 |
| **ATP6V1E1** | 4 | 7.43E+07 | 7.43E+07 | + | 11136 | 681 | 671 | 98.53157122 |
| **RNF144A** | 3 | 100352872 | 100413587 | + | 60715 | 879 | 866 | 98.52104664 |
| **TSPAN5** | 5 | 49994391 | 50077240 | + | 82849 | 807 | 795 | 98.51301115 |
| **NATD1** | 15 | 13593795 | 13605061 | + | 11266 | 333 | 328 | 98.4984985 |
| **CSDE1** | 24 | 2762774 | 2783824 | + | 21050 | 2391 | 2355 | 98.49435383 |
| **ACTG1** | 18 | 11016056 | 11019084 | + | 3028 | 1128 | 1111 | 98.4929078 |
| **GTF2A1** | 6 | 20522466 | 20549065 | + | 26599 | 1193 | 1175 | 98.49119866 |
| **EGLN3** | 6 | 26043813 | 26071078 | + | 27265 | 726 | 715 | 98.48484848 |
| **UBE2W** | 1 | 32250073 | 32282181 | + | 32108 | 462 | 455 | 98.48484848 |
| **UBE2QL1** | 1 | 71789811 | 71807902 | + | 18091 | 921 | 907 | 98.47991314 |
| **DPF3** | 6 | 29322067 | 29491277 | + | 169210 | 1442 | 1420 | 98.47434119 |
| **SF3B5** | 3 | 50519706 | 50522031 | + | 2325 | 261 | 257 | 98.46743295 |
| **HS3ST5** | 3 | 67938199 | 68132244 | + | 194045 | 1038 | 1022 | 98.45857418 |
| **FBXO11** | 3 | 16111105 | 16181090 | + | 69985 | 2658 | 2617 | 98.45748683 |

| | | | | | | | | |
|---|---|---|---|---|---|---|---|---|
| **CABP7** | 17 | 12446233 | 12454619 | + | 8386 | 648 | 638 | 98.45679012 |
| **NPAS3** | 6 | 26097760 | 26694190 | - | 596430 | 2849 | 2805 | 98.45559846 |
| **KICS2** | 4 | 33807907 | 33817479 | - | 9572 | 1552 | 1528 | 98.45360825 |
| **RAP1GDS1** | 5 | 50080381 | 50170452 | - | 90071 | 1935 | 1905 | 98.4496124 |
| **PAQR9** | 10 | 11363298 | 11370766 | - | 7468 | 1215 | 1196 | 98.43621399 |
| **KDELR2** | 15 | 9364746 | 9377403 | + | 12657 | 639 | 629 | 98.43505477 |
| **CFAP20** | 11 | 6288711 | 6295239 | + | 6528 | 702 | 691 | 98.43304843 |
| **LZIC** | 22 | 6994478 | 7003092 | - | 8614 | 573 | 564 | 98.42931937 |
| **NUDT4** | 4 | 46222232 | 46249319 | + | 27087 | 631 | 621 | 98.41521395 |
| **CXXC4** | 5 | 3.58E+07 | 3.60E+07 | - | 161023 | 1324 | 1303 | 98.41389728 |
| **HIBADH** | 1 | 122102915 | 122186981 | + | 84066 | 1131 | 1113 | 98.40848806 |
| **PCMTD1** | 1 | 41017981 | 41059472 | + | 41491 | 1131 | 1113 | 98.40848806 |
| **YTHDF1** | 16 | 5818232 | 5833220 | - | 14988 | 3261 | 3209 | 98.40539712 |
| **DYRK2** | 4 | 3.52E+07 | 3.52E+07 | + | 11728 | 3375 | 3321 | 98.4 |
| **CCNY** | 1 | 143275134 | 143395289 | + | 120155 | 1057 | 1040 | 98.39167455 |
| **CCZ1** | 15 | 9548101 | 9559440 | - | 11339 | 1425 | 1402 | 98.38596491 |
| **AKT3** | 3 | 3.36E+07 | 3.38E+07 | - | 154013 | 1541 | 1516 | 98.37767683 |
| **POU6F2** | 1 | 101426073 | 101738362 | + | 312289 | 2076 | 2042 | 98.36223507 |
| **MCTS1** | Z | 82746188 | 82753388 | - | 7200 | 546 | 537 | 98.35164835 |
| **ESRRG** | 3 | 21314675 | 21687282 | + | 372607 | 2061 | 2027 | 98.35031538 |
| **SRSF2** | 18 | 3808936 | 3813630 | + | 4694 | 666 | 655 | 98.34834835 |
| **DUSP7** | 12 | 4220129 | 4228182 | + | 8053 | 1149 | 1130 | 98.34638816 |
| **MMP16** | 1 | 26249743 | 26419954 | + | 170211 | 1934 | 1902 | 98.34539814 |
| **MIER3** | Z | 12194726 | 12216893 | + | 22167 | 1748 | 1719 | 98.3409611 |
| **HPRT1** | Z | 84432464 | 84449346 | + | 16882 | 657 | 646 | 98.32572298 |
| **GRB2** | 18 | 4781203 | 4832428 | + | 51225 | 654 | 643 | 98.31804281 |
| **TRIM8** | 8 | 23353282 | 23383215 | + | 29933 | 1656 | 1628 | 98.30917874 |
| **WDR5** | 20 | 3390751 | 3402253 | - | 11502 | 1005 | 988 | 98.30845771 |
| **OPRL1** | 16 | 6442121 | 6456706 | - | 14585 | 1107 | 1088 | 98.2836495 |

| | | | | | | | | |
|---|---|---|---|---|---|---|---|---|
| **PPP2CA** | 14 | 9658948 | 9674149 | + | 15201 | 930 | 914 | 98.27956989 |
| **RAB2A** | 1 | 37577304 | 37620080 | - | 42776 | 639 | 628 | 98.27856025 |
| **CWC15** | 2 | 38631514 | 38646314 | - | 14800 | 693 | 681 | 98.26839827 |
| **HRAS** | 6 | 45494404 | 45534274 | - | 39870 | 570 | 560 | 98.24561404 |
| **RAB7A** | 12 | 8946439 | 8968479 | - | 22040 | 624 | 613 | 98.23717949 |
| **DICER1** | 6 | 14783119 | 14847103 | + | 63984 | 5766 | 5664 | 98.23100937 |
| **DACH1** | 2 | 66278987 | 66631894 | - | 352907 | 2312 | 2271 | 98.2266436 |
| **NAP1L4** | 6 | 48685261 | 48713730 | - | 28469 | 1352 | 1328 | 98.22485207 |
| **TRAM1** | 1 | 33554194 | 33569233 | + | 15039 | 1125 | 1105 | 98.22222222 |
| **EIF5** | 6 | 10104065 | 10111639 | - | 7574 | 1293 | 1270 | 98.22119103 |
| **DAB1** | 9 | 3773607 | 4180190 | + | 406583 | 1737 | 1706 | 98.21531376 |
| **SH3GL3** | 13 | 10926333 | 10968624 | + | 42291 | 1062 | 1043 | 98.21092279 |
| **PRDM1** | 3 | 71833026 | 71865908 | - | 32882 | 2904 | 2852 | 98.20936639 |
| **PVALB** | 4 | 54057176 | 54066989 | + | 9813 | 333 | 327 | 98.1981982 |
| **RAP1B** | 4 | 35500432 | 35532192 | + | 31760 | 555 | 545 | 98.1981982 |
| **ELOVL4** | 3 | 83157116 | 83197422 | + | 40306 | 1107 | 1087 | 98.19331527 |
| **PSMC3** | 6 | 40140298 | 40147225 | - | 6927 | 1269 | 1246 | 98.18754925 |
| **STK11** | 26 | 2041887 | 2073960 | - | 32073 | 1323 | 1299 | 98.18594104 |
| **HOXD4** | 7 | 21894191 | 21896485 | + | 2294 | 714 | 701 | 98.17927171 |
| **HMGN3** | 3 | 83390205 | 83414146 | + | 23941 | 384 | 377 | 98.17708333 |
| **PAIP2** | 14 | 1.69E+07 | 1.69E+07 | - | 8272 | 384 | 377 | 98.17708333 |
| **HOXA6** | 1 | 1.22E+08 | 1.22E+08 | + | 2198 | 821 | 806 | 98.17295981 |
| **ARF1** | 1 | 154133543 | 154146916 | - | 13373 | 546 | 536 | 98.16849817 |
| **TM9SF3** | 8 | 16396561 | 16441906 | - | 45345 | 1746 | 1714 | 98.1672394 |
| **RGS19** | 16 | 6539422 | 6554555 | + | 15133 | 600 | 589 | 98.16666667 |
| **ST3GAL3** | 9 | 10399083 | 10573797 | - | 174714 | 1308 | 1284 | 98.16513761 |
| **GMPPB** | 12 | 6672270 | 6674500 | + | 2230 | 1138 | 1117 | 98.15465729 |
| **SPRED2** | 3 | 2631827 | 2689507 | - | 57680 | 1407 | 1381 | 98.15209666 |
| **ADGRB3** | 3 | 87362886 | 87828744 | - | 465858 | 4707 | 4620 | 98.15168897 |

| | | | | | | | | |
|---|---|---|---|---|---|---|---|---|
| **DOK6** | 1 | 56914697 | 57159534 | + | 244837 | 1081 | 1061 | 98.14986124 |
| **NEK7** | 9 | 22170348 | 22237467 | - | 67119 | 909 | 892 | 98.12981298 |
| **GNAZ** | 17 | 11098040 | 11147808 | + | 49768 | 1068 | 1048 | 98.12734082 |
| **ZFP36L1** | 6 | 31442203 | 31445762 | + | 3559 | 1008 | 989 | 98.11507937 |
| **GPR85** | 4 | 26664618 | 26669132 | + | 4514 | 1113 | 1092 | 98.11320755 |
| **DESI2** | 3 | 34012029 | 34026850 | + | 14821 | 582 | 571 | 98.10996564 |
| **GPR12** | 2 | 47394176 | 47405109 | - | 10933 | 1005 | 986 | 98.10945274 |
| **EFNA5** | Z | 48535929 | 48750260 | + | 214331 | 687 | 674 | 98.1077147 |
| **NAP1L1** | 4 | 3.83E+07 | 3.83E+07 | - | 28300 | 1308 | 1283 | 98.08868502 |
| **EPC1** | 1 | 142062909 | 142126331 | - | 63422 | 2713 | 2661 | 98.08330262 |
| **SLC35F3** | 3 | 45102781 | 45273229 | + | 170448 | 1558 | 1528 | 98.07445443 |
| **BCL11B** | 6 | 12658130 | 12749907 | + | 91777 | 3021 | 2962 | 98.0470043 |
| **VASH1** | 6 | 22575530 | 22593194 | - | 17664 | 870 | 853 | 98.04597701 |
| **PITPNA** | 19 | 4882642 | 4907328 | + | 24686 | 816 | 800 | 98.03921569 |
| **STIM2** | 5 | 18357239 | 18424676 | + | 67437 | 2140 | 2098 | 98.03738318 |
| **MLC1** | 4 | 19599988 | 19614077 | - | 14089 | 1170 | 1147 | 98.03418803 |
| **ELOVL6** | 5 | 3.80E+07 | 3.81E+07 | - | 80164 | 812 | 796 | 98.02955665 |
| **PTCHD4** | 3 | 110914722 | 110995645 | + | 80923 | 3141 | 3079 | 98.02610634 |
| **SUFU** | 8 | 23252776 | 23341749 | + | 88973 | 1612 | 1580 | 98.01488834 |
| **PICALM** | 2 | 33883447 | 33949257 | + | 65810 | 2166 | 2123 | 98.01477378 |
| **SERP1** | 10 | 25563246 | 25565156 | + | 1910 | 201 | 197 | 98.00995025 |
| **TENT4B** | 11 | 12508580 | 12547946 | - | 39366 | 2058 | 2017 | 98.00777454 |
| **ZBTB20** | 2 | 23394065 | 23869013 | - | 474948 | 3814 | 3738 | 98.00734137 |
| **FOXP1** | 12 | 16752338 | 17133161 | - | 380823 | 2351 | 2304 | 98.0008507 |
| **DPY30** | 3 | 35119083 | 35124715 | - | 5632 | 300 | 294 | 98 |
| **RPS12** | 3 | 59909503 | 59913486 | - | 3983 | 399 | 391 | 97.99498747 |
| **CCND1** | 6 | 6.12E+07 | 6.12E+07 | - | 23896 | 897 | 879 | 97.99331104 |
| **HOXA11** | 1 | 122362756 | 122365185 | + | 2429 | 894 | 876 | 97.98657718 |
| **ARF4** | 12 | 1670331 | 1680683 | + | 10352 | 543 | 532 | 97.97421731 |

| | | | | | | | | |
|---|---|---|---|---|---|---|---|---|
| **USP15** | 4 | 33087676 | 33147305 | + | 59629 | 3058 | 2996 | 97.97253107 |
| **RER1** | 22 | 4997575 | 5006678 | - | 9103 | 591 | 579 | 97.96954315 |
| **CAND1** | 4 | 35064281 | 35091176 | + | 26895 | 3693 | 3618 | 97.96913079 |
| **SKP1** | 14 | 9684480 | 9693744 | + | 9264 | 492 | 482 | 97.96747967 |
| **RASD1** | 15 | 13902010 | 13903985 | - | 1975 | 831 | 814 | 97.95427196 |
| **SRP54** | 6 | 25650978 | 25666542 | - | 15564 | 1515 | 1484 | 97.95379538 |
| **EIF1** | 27 | 3097436 | 3098586 | - | 1150 | 342 | 335 | 97.95321637 |
| **DVL3** | 10 | 17166346 | 17200237 | + | 33891 | 2190 | 2145 | 97.94520548 |
| **NTS** | 4 | 43113117 | 43128483 | + | 15366 | 486 | 476 | 97.94238683 |
| **ID4** | 1 | 93062315 | 93065578 | + | 3263 | 864 | 846 | 97.91666667 |
| **SIX2** | 3 | 27668945 | 27673400 | - | 4455 | 864 | 846 | 97.91666667 |
| **CASTOR2** | 19 | 8144378 | 8274668 | - | 130290 | 1004 | 983 | 97.90836653 |
| **RBM18** | 20 | 1657486 | 1668955 | + | 11469 | 573 | 561 | 97.90575916 |
| **RUVBL1** | 12 | 9466741 | 9484962 | + | 18221 | 1371 | 1342 | 97.88475565 |
| **NOL4** | 1 | 43398132 | 43588859 | + | 190727 | 2450 | 2398 | 97.87755102 |
| **CLTCL1** | 17 | 8485489 | 8520427 | + | 34938 | 5055 | 4947 | 97.86350148 |
| **HM13** | 16 | 7316857 | 7331739 | + | 14882 | 1351 | 1322 | 97.85344189 |
| **RNF44** | 14 | 3550420 | 3566932 | + | 16512 | 1392 | 1362 | 97.84482759 |
| **ING1** | 2 | 93983467 | 93990191 | - | 6724 | 1479 | 1447 | 97.83637593 |
| **HMX3** | 8 | 31963427 | 31964747 | + | 1320 | 924 | 904 | 97.83549784 |
| **KPNA3** | 2 | 54387994 | 54435451 | + | 47457 | 1617 | 1582 | 97.83549784 |
| **TXNDC17** | 19 | 98130 | 100337 | + | 2207 | 369 | 361 | 97.83197832 |
| **PEX7** | 3 | 51145281 | 51188815 | + | 43534 | 966 | 945 | 97.82608696 |
| **LDB1** | 8 | 22393781 | 22418371 | + | 24590 | 1379 | 1349 | 97.82451051 |
| **MAP1LC3A** | 16 | 13269950 | 13287199 | + | 17249 | 366 | 358 | 97.81420765 |
| **CRYBA2** | 7 | 15498504 | 15505232 | - | 6728 | 594 | 581 | 97.81144781 |
| **DCLK1** | 2 | 5.10E+07 | 5.12E+07 | - | 229351 | 2512 | 2457 | 97.81050955 |
| **GLRA2** | 2 | 101303990 | 101430299 | + | 126309 | 1507 | 1474 | 97.81021898 |
| **ENOX1** | 2 | 56600348 | 56957842 | + | 357494 | 2008 | 1964 | 97.80876494 |

| | | | | | | | | |
|---|---|---|---|---|---|---|---|---|
| **NAA50** | 2 | 27213028 | 27233673 | - | 20645 | 819 | 801 | 97.8021978 |
| **AKT1** | 6 | 8528373 | 8605533 | + | 77160 | 1544 | 1510 | 97.79792746 |
| **FOXF2** | 1 | 85652936 | 85658814 | - | 5878 | 1179 | 1153 | 97.79474131 |
| **TTC9** | 6 | 3.03E+07 | 3.04E+07 | - | 27790 | 633 | 619 | 97.78830964 |
| **CDK14** | 1 | 1.33E+08 | 1.34E+08 | - | 348493 | 1716 | 1678 | 97.78554779 |
| **UBE2E1** | 1 | 117688052 | 117733245 | - | 45193 | 1173 | 1147 | 97.78346121 |
| **GLYR1** | 15 | 24872 | 51703 | - | 26831 | 1848 | 1807 | 97.78138528 |
| **GRK4** | 5 | 8650420 | 8689072 | + | 38652 | 1710 | 1672 | 97.77777778 |
| **TMED10** | 6 | 2.33E+07 | 2.33E+07 | + | 15880 | 672 | 657 | 97.76785714 |
| **CUL2** | 1 | 143161980 | 143203469 | - | 41489 | 2238 | 2188 | 97.76586238 |
| **FHL1** | Z | 84678798 | 84708956 | + | 30158 | 1208 | 1181 | 97.76490066 |
| **TECR** | 9 | 17133975 | 17157130 | + | 23155 | 939 | 918 | 97.76357827 |
| **BRSK2** | 6 | 47008531 | 47318605 | + | 310074 | 3035 | 2967 | 97.75947282 |
| **AIDA** | 3 | 19170897 | 19199893 | + | 28996 | 933 | 912 | 97.74919614 |
| **TOB1** | 18 | 5650929 | 5655371 | - | 4442 | 972 | 950 | 97.73662551 |
| **BARHL2** | 9 | 16116772 | 16120745 | - | 3973 | 1501 | 1467 | 97.73484344 |
| **PPARD** | 24 | 3288348 | 3306724 | - | 18376 | 1847 | 1805 | 97.72604223 |
| **PAX6** | 6 | 5.54E+07 | 5.54E+07 | - | 21314 | 1539 | 1504 | 97.72579597 |
| **RSU1** | 1 | 135518765 | 135621778 | - | 103013 | 834 | 815 | 97.72182254 |
| **EIF2B3** | 9 | 9669515 | 9769163 | + | 99648 | 1359 | 1328 | 97.71891096 |
| **CABLES1** | 1 | 47775504 | 47845196 | - | 69692 | 1578 | 1542 | 97.71863118 |
| **CHD7** | 1 | 37425703 | 37558908 | - | 133205 | 9335 | 9122 | 97.7182646 |
| **ILRUN** | 24 | 3489994 | 3518649 | + | 28655 | 876 | 856 | 97.71689498 |
| **RPS15** | 26 | 1833943 | 1836564 | - | 2621 | 438 | 428 | 97.71689498 |
| **TMEFF2** | 7 | 2.99E+07 | 3.00E+07 | + | 126331 | 1221 | 1193 | 97.70679771 |
| **MBOAT2** | 3 | 1.01E+08 | 1.01E+08 | - | 92108 | 1609 | 1572 | 97.70043505 |
| **ITGB1BP1** | 3 | 101540503 | 101549935 | - | 9432 | 651 | 636 | 97.69585253 |
| **MARK3** | 6 | 10012881 | 10074767 | - | 61886 | 2640 | 2579 | 97.68939394 |
| **CACNG3** | 15 | 15767670 | 15791542 | + | 23872 | 1337 | 1306 | 97.68137622 |

| | | | | | | | | |
|---|---|---|---|---|---|---|---|---|
| **PCGF3** | Z | 72282181 | 72336436 | - | 54255 | 862 | 842 | 97.67981439 |
| **HNRNPK** | Z | 38415865 | 38436111 | - | 20246 | 1551 | 1515 | 97.67891683 |
| **NKX2-3** | 8 | 21859467 | 21861707 | - | 2240 | 1974 | 1928 | 97.66970618 |
| **HNRNPAB** | 14 | 7873902 | 7899618 | - | 25716 | 1029 | 1005 | 97.66763848 |
| **HS2ST1** | 9 | 14861701 | 14941239 | + | 79538 | 1071 | 1046 | 97.66573296 |
| **EGR3** | 28 | 3653788 | 3656189 | + | 2401 | 1194 | 1166 | 97.65494137 |
| **FOXJ2** | 2 | 30239126 | 30270292 | - | 31166 | 1825 | 1782 | 97.64383562 |
| **ABHD17C** | 13 | 11907179 | 11935978 | - | 28799 | 933 | 911 | 97.64201501 |
| **SNTG1** | 1 | 4.15E+07 | 4.18E+07 | - | 327361 | 1566 | 1529 | 97.63729246 |
| **TMEM196** | 1 | 125368294 | 125387666 | + | 19372 | 927 | 905 | 97.62675297 |
| **FOXL2** | 10 | 1105601 | 1118532 | + | 12931 | 921 | 899 | 97.61129207 |
| **FGF16** | 21 | 2570241 | 2581393 | + | 11152 | 627 | 612 | 97.6076555 |
| **YAF2** | 4 | 29512723 | 29537407 | - | 24684 | 543 | 530 | 97.60589319 |
| **NELFA** | 5 | 7618831 | 7642804 | - | 23973 | 1620 | 1581 | 97.59259259 |
| **CABP1** | 17 | 10606541 | 10633170 | + | 26629 | 1284 | 1253 | 97.58566978 |
| **THOC7** | 12 | 14026834 | 14035091 | - | 8257 | 615 | 600 | 97.56097561 |
| **ATP11B** | 10 | 17956386 | 18020285 | - | 63899 | 4121 | 4020 | 97.54913856 |
| **OAZ1** | 26 | 3642018 | 3645915 | - | 3897 | 651 | 635 | 97.5422427 |
| **YPEL5** | 3 | 16920125 | 16923995 | + | 3870 | 366 | 357 | 97.54098361 |
| **SEC61G** | 1 | 104196451 | 104200805 | + | 4354 | 325 | 317 | 97.53846154 |
| **ARF6** | 6 | 1919183 | 1923055 | + | 3872 | 528 | 515 | 97.53787879 |
| **RBMS3** | 1 | 115196684 | 115902929 | - | 706245 | 2150 | 2097 | 97.53488372 |
| **VAT1L** | 11 | 16224736 | 16289523 | - | 64787 | 1257 | 1226 | 97.53381066 |
| **AGPS** | 7 | 22378563 | 22456769 | + | 78206 | 1986 | 1937 | 97.5327291 |
| **UBE2D2** | 14 | 15056594 | 15081979 | + | 25385 | 444 | 433 | 97.52252252 |
| **CSNK1E** | 4 | 53320495 | 53342562 | + | 22067 | 1251 | 1220 | 97.52198241 |
| **ARL5B** | 1 | 136570052 | 136586146 | + | 16094 | 684 | 667 | 97.51461988 |
| **HOXD12** | 7 | 21841951 | 21849313 | + | 7362 | 801 | 781 | 97.5031211 |
| **PID1** | 10 | 10544266 | 10630147 | - | 85881 | 801 | 781 | 97.5031211 |
| **MOSPD1** | Z | 84546692 | 84560738 | - | 14046 | 881 | 859 | 97.50283768 |

| | | | | | | | | |
|---|---|---|---|---|---|---|---|---|
| **CAMK2G** | 8 | 15308603 | 15428031 | + | 119428 | 1841 | 1795 | 97.50135796 |
| **GPR137B** | 3 | 45993162 | 46018391 | + | 25229 | 1158 | 1129 | 97.49568221 |
| **FBXW2** | 20 | 50763 | 59029 | + | 8266 | 1468 | 1431 | 97.47956403 |
| **THRA** | 27 | 3463037 | 3470847 | - | 7810 | 1227 | 1196 | 97.47351263 |
| **CALB1** | 1 | 25670557 | 25688838 | + | 18281 | 789 | 769 | 97.46514575 |
| **AZIN1** | 1 | 20487094 | 20514217 | + | 27123 | 1455 | 1418 | 97.45704467 |
| **GAP43** | 2 | 24018603 | 24076746 | + | 58143 | 747 | 728 | 97.45649264 |
| **GULP1** | 7 | 31204823 | 31373162 | - | 168339 | 1173 | 1143 | 97.44245524 |
| **NHLH2** | 2 | 25968928 | 25972992 | - | 4064 | 429 | 418 | 97.43589744 |
| **RALB** | 7 | 12031962 | 12077078 | - | 45116 | 621 | 605 | 97.42351047 |
| **ZNF423** | 11 | 12631704 | 12857942 | + | 226238 | 7216 | 7030 | 97.42239468 |
| **ATXN7L1** | 4 | 13166288 | 13278247 | - | 111959 | 2754 | 2683 | 97.42193174 |
| **SYNPR** | 12 | 13872255 | 13979560 | + | 107305 | 891 | 868 | 97.41863075 |
| **GRHL2** | 1 | 20931626 | 20998042 | - | 66416 | 2243 | 2185 | 97.41417744 |
| **PACSIN2** | 4 | 60454961 | 60507986 | - | 53025 | 1578 | 1537 | 97.4017744 |
| **PBX3** | 20 | 593351 | 696857 | - | 103506 | 1424 | 1387 | 97.40168539 |
| **NT5DC3** | 4 | 57202062 | 57221928 | + | 19866 | 1500 | 1461 | 97.4 |
| **ATP6V0B** | 9 | 10347462 | 10354071 | - | 6609 | 615 | 599 | 97.39837398 |
| **HOXD9** | 7 | 21869495 | 21873346 | + | 3851 | 1994 | 1942 | 97.39217653 |
| **FAM120A** | 12 | 439649 | 486370 | + | 46721 | 3449 | 3359 | 97.39054798 |
| **SLC35B3** | 1 | 88420680 | 88440236 | - | 19556 | 1110 | 1081 | 97.38738739 |
| **GJA1** | 3 | 65073653 | 65083090 | - | 9437 | 1146 | 1116 | 97.38219895 |
| **MAPK8** | 8 | 17931309 | 18076014 | + | 144705 | 1373 | 1337 | 97.37800437 |
| **QRICH1** | 12 | 11847357 | 11874015 | + | 26658 | 2319 | 2258 | 97.36955584 |
| **DDX5** | 18 | 8437163 | 8444520 | - | 7357 | 1823 | 1775 | 97.36697751 |
| **ACTN2** | 3 | 46148662 | 46217724 | + | 69062 | 2694 | 2623 | 97.36451373 |
| **FIGN** | 7 | 17453979 | 17559249 | - | 105270 | 4549 | 4429 | 97.3620576 |
| **MED31** | 19 | 101328 | 103697 | - | 2369 | 378 | 368 | 97.35449735 |
| **BMI1** | 1 | 138065401 | 138075650 | + | 10249 | 981 | 955 | 97.34964322 |
| **GRK6** | 14 | 3284178 | 3299465 | + | 15287 | 1734 | 1688 | 97.34717416 |

| | | | | | | | | |
|---|---|---|---|---|---|---|---|---|
| **TUSC2** | 12 | 5349150 | 5354552 | - | 5402 | 339 | 330 | 97.34513274 |
| **POU2F1** | 2 | 4578587 | 4681092 | - | 102505 | 2630 | 2560 | 97.33840304 |
| **RPRD1B** | 16 | 7808606 | 7833442 | - | 24836 | 1014 | 987 | 97.33727811 |
| **S100A14** | 29 | 1697169 | 1698901 | + | 1732 | 300 | 292 | 97.33333333 |
| **PAN3** | 2 | 47930085 | 48010029 | + | 79944 | 2772 | 2698 | 97.33044733 |
| **ELAVL4** | 9 | 6621993 | 6702986 | - | 80993 | 1645 | 1601 | 97.32522796 |
| **ARL15** | Z | 13255176 | 13481616 | + | 226440 | 633 | 616 | 97.31437599 |
| **LGALSL** | 3 | 2291730 | 2299400 | + | 7670 | 516 | 502 | 97.28682171 |
| **CPLX1** | Z | 71921044 | 72026709 | + | 105665 | 405 | 394 | 97.28395062 |
| **PCGF5** | 8 | 19451214 | 19551675 | + | 100461 | 771 | 750 | 97.27626459 |
| **GPS1** | 18 | 11928498 | 11941552 | + | 13054 | 1760 | 1712 | 97.27272727 |
| **ACTC1** | 6 | 35650043 | 35655341 | + | 5298 | 1134 | 1103 | 97.26631393 |
| **RPS7** | 3 | 98939046 | 98946152 | + | 7106 | 585 | 569 | 97.26495726 |
| **RALA** | 1 | 101852617 | 101887996 | + | 35379 | 621 | 604 | 97.26247987 |
| **NETO1** | 1 | 58422057 | 58484903 | - | 62846 | 1677 | 1631 | 97.25700656 |
| **KLHL41** | 7 | 19344277 | 19351982 | + | 7705 | 1821 | 1771 | 97.2542559 |
| **MPPED1** | 4 | 60845355 | 60908649 | + | 63294 | 909 | 884 | 97.24972497 |
| **AP3S1** | Z | 64270583 | 64306898 | - | 36315 | 618 | 601 | 97.24919094 |
| **CGA** | 3 | 80426386 | 80432647 | + | 6261 | 363 | 353 | 97.24517906 |
| **GUCY1A1** | 5 | 42184574 | 42219383 | + | 34809 | 2067 | 2010 | 97.24238026 |
| **PITX2** | 5 | 38201401 | 38215848 | - | 14447 | 1048 | 1019 | 97.23282443 |
| **RNF152** | 1 | 83894858 | 83940207 | - | 45349 | 612 | 595 | 97.22222222 |
| **ASIC4** | 7 | 15850163 | 15878718 | + | 28555 | 1617 | 1572 | 97.21706865 |
| **ERP44** | 1 | 61804459 | 61850181 | - | 45722 | 1221 | 1187 | 97.21539722 |
| **RHOJ** | 6 | 6577125 | 6637787 | + | 60662 | 645 | 627 | 97.20930233 |
| **CRY1** | 4 | 56090822 | 56130373 | - | 39551 | 1863 | 1811 | 97.20880301 |
| **ZNF410** | 6 | 23913901 | 23936638 | - | 22737 | 1354 | 1316 | 97.19350074 |
| **OSR1** | 3 | 106875170 | 106883135 | - | 7965 | 783 | 761 | 97.19029374 |
| **BCLAF1** | 3 | 50865020 | 50890779 | - | 25759 | 3022 | 2937 | 97.18729318 |
| **PAPOLA** | 6 | 14063180 | 14108479 | - | 45299 | 2381 | 2314 | 97.18605628 |

| | | | | | | | | | |
|---|---|---|---|---|---|---|---|---|---|
| **TFAP2A** | 1 | 89289055 | 89302567 | - | | 13512 | 1452 | 1411 | 97.17630854 |
| **PACSIN3** | 6 | 39815621 | 39840251 | - | | 24630 | 1374 | 1335 | 97.16157205 |
| **PAFAH1B1** | 19 | 391382 | 417362 | - | | 25980 | 1233 | 1198 | 97.16139497 |
| **KCNG3** | 3 | 25606264 | 25618532 | - | | 12268 | 1901 | 1847 | 97.15938979 |
| **ELFN1** | 15 | 11462162 | 11565714 | + | | 103552 | 2391 | 2323 | 97.15600167 |
| **WDR1** | 5 | 12796528 | 12815318 | - | | 18790 | 1824 | 1772 | 97.14912281 |
| **MEMO1** | 3 | 35085186 | 35112559 | - | | 27373 | 982 | 954 | 97.14867617 |
| **USP25** | 2 | 12332109 | 12426068 | - | | 93959 | 3471 | 3372 | 97.14779602 |
| **CUL5** | 2 | 43535728 | 43566323 | + | | 30595 | 2343 | 2276 | 97.14041827 |
| **KBTBD4** | 6 | 40225721 | 40245414 | - | | 19693 | 2200 | 2137 | 97.13636364 |
| **GABRB3** | 2 | 86403455 | 86519840 | + | | 116385 | 1431 | 1390 | 97.13487072 |
| **PIP4K2A** | 1 | 138178878 | 138284738 | - | | 105860 | 1218 | 1183 | 97.12643678 |
| **PHACTR1** | 1 | 90122257 | 90427337 | + | | 305080 | 2037 | 1978 | 97.1035837 |
| **FST** | Z | 13607237 | 13613955 | - | | 6718 | 1032 | 1002 | 97.09302326 |
| **RIMKLB** | 2 | 30435811 | 30478246 | + | | 42435 | 1161 | 1127 | 97.07149009 |
| **PPP1CB** | 3 | 29078363 | 29107774 | - | | 29411 | 1023 | 993 | 97.06744868 |
| **GSX1** | 2 | 47808034 | 47809769 | + | | 1735 | 744 | 722 | 97.04301075 |
| **SCAMP5** | 13 | 1843846 | 1851123 | - | | 7277 | 708 | 687 | 97.03389831 |
| **IRF2BPL** | 6 | 22436447 | 22440015 | + | | 3568 | 2217 | 2151 | 97.02300406 |
| **YWHAQ** | 3 | 101633523 | 101656084 | - | | 22561 | 738 | 716 | 97.01897019 |
| **PHF10** | 3 | 42454992 | 42472207 | - | | 17215 | 1610 | 1562 | 97.01863354 |
| **AMMECR1** | 21 | 7600333 | 7651887 | + | | 51554 | 1542 | 1496 | 97.01686122 |
| **KCTD12** | 2 | 68558272 | 68562753 | - | | 4481 | 999 | 969 | 96.996997 |
| **TMEM272** | 2 | 53469503 | 53495111 | + | | 25608 | 564 | 547 | 96.9858156 |
| **CTNND2** | 1 | 73667315 | 74309029 | - | | 641714 | 4077 | 3954 | 96.98307579 |
| **CDK5R2** | 7 | 15481140 | 15482526 | + | | 1386 | 960 | 931 | 96.97916667 |

**Table S19 - Barn swallow individuals sequenced with Hifi technology**

| Sample name | Location | Latitude | Longitude | sex | Sequencing coverage |
|---|---|---|---|---|---|
| **2** | Parco Adda (CR) | 45°26′N | 9°30′E | F | 25x |
| **3** | Brescia | 45°22′N | 10°15'E | F | 19.5x |
| **4** | Oleggio (NO) | 45°34′N | 8°39'E | F | 20x |
| **A1** | Matera | 40°40′N | 16°36′E | F | 33x |
| **A2** | Matera | 40°40′N | 16°36′E | F | 15x |

**Table S20 - HiFi sequencing statistics.** Sequencing coverage was estimated based on barn swallow genome size

| | Sample | | | | |
|---|---|---|---|---|---|
| | **A1** | **A2** | **2** | **3** | **4** |
| **>= Q20 Reads** | 2,493,595 | 940,660 | 1,856,659 | 1,530,055 | 1,430,811 |
| **>= Q20 Yield (bp)** | 36,551,797,585 | 17,101,319,656 | 27,772,551,635 | 21,646,429,144 | 22,740,719,734 |
| **>= Q20 Read Lenght (mean, bp)** | 14657.5 | 18,180 | 14,928 | 14,147 | 16,136 |
| **>= Q20 Read Quality (median)** | Q32 | Q29 | Q32 | Q32 | Q31 |
| **Sequencing coverage** | 33 | 15 | 25 | 19 | 20 |

**Table S21- Hifi assembly statistics.**

| | Primary before purging | | Primary after purging | | | Alternate before purging | | Alternate after purging | | |
|---|---|---|---|---|---|---|---|---|---|---|
| **Sample 2** | **Contigs** | **Gaps** | **Contigs** | **Gaps** | **Purged (bp)** | **Contigs** | **Gaps** | **Contigs** | **Gaps** | **Purged (bp)** |
| **Total bp** | 1,307,545,121 | 0 | 1,163,058,884 | 1,288 | 144,486,237 | 1,106,170,783 | 0 | 1,011,189,907 | 138 | 94,980,876 |
| **Number** | 1,959 | 0 | 1,106 | 56 | - | 4,940 | 0 | 2,840 | 6 | - |
| **Max length (bp)** | 40,208,769 | 0 | 40,208,769 | 23 | - | 10,650,792 | 0 | 10,650,792 | 23 | - |
| **N50 (bp)** | 5,371,840 | 0 | 7,140,827 | 23 | - | 1,099,778 | 0 | 1,225,108 | 23 | - |
| **N90 (bp)** | 295,307 | 0 | 564,576 | 23 | - | 89,329 | 0 | 198,244 | 23 | - |
| **NG50 (bp)** | 8,189,401 | 0 | 8,189,401 | - | - | 1,165,674 | 0 | 1,114,799 | - | - |
| **NG90 (bp)** | 1,339,570 | 0 | 1,156,149 | - | - | 166,867 | 0 | 51,878 | - | - |
| **Sample 3** | **Contigs** | **Gaps** | **Contigs** | **Gaps** | **Purged (bp)** | **Contigs** | **Gaps** | **Contigs** | **Gaps** | **Purged (bp)** |
| **Total bp** | 1,273,430,777 | 0 | 1,135,657,041 | 1,081 | 137,773,736 | 1,072,637,793 | 0 | 991,522,024 | 138 | 81,115,769 |
| **Number** | 2,368 | 0 | 1,434 | 47 | - | 6,324 | 0 | 3,941 | 6 | - |
| **Max length (bp)** | 38,082,319 | 0 | 38,082,319 | 23 | - | 4,654,537 | 0 | 4,654,537 | 23 | - |
| **N50 (bp)** | 3,477,328 | 0 | 4,230,173 | 23 | - | 579,024 | 0 | 636,015 | 23 | - |
| **N90 (bp)** | 216,449 | 0 | 405,189 | 23 | - | 66,297 | 0 | 117,718 | 23 | - |
| **NG50 (bp)** | 4,473,894 | 0 | 4,473,894 | - | - | 567,346 | 0 | 585,425 | - | - |
| **NG90 (bp)** | 887,578 | 0 | 753,224 | - | - | 55,473 | 0 | 43,696 | - | - |
| **Sample 4** | **Contigs** | **Gaps** | **Contigs** | **Gaps** | **Purged (bp)** | **Contigs** | **Gaps** | **Contigs** | **Gaps** | **Purged (bp)** |
| **Total bp** | 1,301,377,510 | 0 | 1,139,278,982 | 1,219 | 162,098,528 | 1,072,102,039 | 0 | 984,724,034 | 69 | 87,378,005 |
| **Number** | 2,199 | 0 | 1,267 | 53 | - | 5,743 | 0 | 3,549 | 3 | - |
| **Max length (bp)** | 21,040,034 | 0 | 21,040,034 | 23 | - | 5,303,267 | 0 | 5,303,267 | 23 | - |
| **N50 (bp)** | 3,177,341 | 0 | 3,906,945 | 23 | - | 670,370 | 0 | 748,892 | 23 | - |
| **N90 (bp)** | 247,663 | 0 | 483,079 | 23 | - | 70,261 | 0 | 129,142 | 23 | - |
| **NG50 (bp)** | 4,168,408 | 0 | 4,097,292 | - | - | 664,257 | 0 | 652,964 | - | - |
| **NG90 (bp)** | 965,081 | 0 | 799,088 | - | - | 62,351 | 0 | 29,319 | - | - |
| **Sample A1** | **Contigs** | **Gaps** | **Contigs** | **Gaps** | **Purged (bp)** | **Contigs** | **Gaps** | **Contigs** | **Gaps** | **Purged (bp)** |
| **Total bp** | 1,249,218,340 | 0 | 1,143,554,647 | 851 | 105,663,693 | 1,115,757,570 | 0 | 1,014,043,497 | 115 | 101,714,073 |
| **Number** | 1,301 | 0 | 742 | 37 | - | 4,220 | 0 | 2,144 | 5 | - |
| **Max length (bp)** | 45,188,444 | 0 | 45,188,444 | 23 | - | 10,892,277 | 0 | 10,892,277 | 23 | - |
| **N50 (bp)** | 6,816,398 | 0 | 7,650,723 | 23 | - | 1,619,225 | 0 | 1,847,324 | 23 | - |
| **N90 (bp)** | 544,953 | 0 | 1,017,178 | 23 | - | 121,942 | 0 | 308,732 | 23 | - |
| **NG50 (bp)** | 8,633,890 | 0 | 8,633,890 | - | - | 1,713,272 | 0 | 1,713,272 | - | - |
| **NG90 (bp)** | 2,097,283 | 0 | 1,911,453 | - | - | 246,935 | 0 | 147,958 | - | - |
| **Sample A2** | **Contigs** | **Gaps** | **Contigs** | **Gaps** | **Purged (bp)** | **Contigs** | **Gaps** | **Contigs** | **Gaps** | **Purged (bp)** |
| **Total bp** | 1,276,591,930 | 0 | 1,123,142,668 | 759 | 153,449,262 | 1,051,556,461 | 0 | 954,145,800 | 46 | 97,410,661 |

| | | | | | | | | | | |
|---|---|---|---|---|---|---|---|---|---|---|
| **Number** | 3,111 | 0 | 1,861 | 33 | - | 8,500 | 0 | 5,550 | 2 | - |
| **Max length (bp)** | 12,728,721 | 0 | 12,728,721 | 23 | - | 1,850,512 | 0 | 1,850,512 | 23 | - |
| **N50 (bp)** | 1,547,009 | 0 | 1,860,711 | 23 | - | 259,207 | 0 | 291,680 | 23 | - |
| **N90 (bp)** | 155,020 | 0 | 265,534 | 23 | - | 49,549 | 0 | 80,915 | 23 | - |
| **NG50 (bp)** | 1,944,253 | 0 | 1,930,674 | - | - | 250,570 | 0 | 246,485 | - | - |
| **NG90 (bp)** | 451,290 | 0 | 348,627 | - | - | 39,495 | 0 | - | - | - |

**Table S22 - Pangenome ortholog analysis.** The individual, or combination of individuals, are reported with the corresponding number of "private" or shared bHirRus1 genes.

| Individuals | n. genes | Tot genes searc | % |
|---|---|---|---|
| **all** | 16801 | 18136 | 92.64 |
| **bHirRus1** | 236 | 18136 | 1.30 |
| **HiFi raw-reads** | 79 | 18136 | 0.44 |
| **2,3,4,A1** | 151 | 18136 | 0.83 |
| **2,4,A1,a2** | 103 | 18136 | 0.57 |
| **3,4,A1,A2** | 89 | 18136 | 0.49 |
| **2,3,A1,A2** | 85 | 18136 | 0.47 |
| **2,3,4,A2** | 66 | 18136 | 0.36 |
| **2,4,A1** | 48 | 18136 | 0.26 |
| **A1** | 46 | 18136 | 0.25 |
| **2,3,A1** | 41 | 18136 | 0.23 |
| **2,A1** | 41 | 18136 | 0.23 |
| **2,A1,A2** | 40 | 18136 | 0.22 |
| **3,4,A2** | 39 | 18136 | 0.22 |
| **3** | 32 | 18136 | 0.18 |
| **2** | 25 | 18136 | 0.14 |
| **4** | 24 | 18136 | 0.13 |
| **A2** | 23 | 18136 | 0.13 |
| **A1,A2** | 23 | 18136 | 0.13 |
| **2,3,4** | 21 | 18136 | 0.12 |
| **3,4,A1** | 21 | 18136 | 0.12 |
| **2,4** | 21 | 18136 | 0.12 |
| **2,3,A2** | 20 | 18136 | 0.11 |
| **2,4,A2** | 18 | 18136 | 0.10 |
| **4,A1** | 18 | 18136 | 0.10 |
| **3,A1,A2** | 16 | 18136 | 0.09 |
| **3,A1** | 16 | 18136 | 0.09 |
| **2,3** | 15 | 18136 | 0.08 |

| | | | |
|---|---|---|---|
| **2,A2** | 13 | 18136 | 0.07 |
| **4,A2** | 13 | 18136 | 0.07 |
| **4,A1,A2** | 12 | 18136 | 0.07 |
| **3,4** | 11 | 18136 | 0.06 |
| **3,A2** | 8 | 18136 | 0.04 |

**Table S23 - 236 genes found only in bHirRus1.** The 236 genes thet were not found in the HiFi assembly are listed with the corresponding coordinates in bHirRus1.
Yes or No in the last 5 columns indicates whether the gene was found with BLAST in the HiFi raw reads. Green = found in the HiFi reads. Red = Not found in the HiFi reads.
The 157 missing genes also from raw-reads are marked in yellow.

| Gene | chr | start | end | Sample 2 raw reads | Sample 3 raw reads | Sample 4 raw re | Sample A1 raw | Sample A2 raw reads |
|---|---|---|---|---|---|---|---|---|
| **LOC120748843** | Chr1 | 88901742 | 88903345 | Yes | Yes | Yes | Yes | Yes |
| **LOC120757517** | Chr10 | 6522574 | 6522693 | Yes | Yes | Yes | Yes | Yes |
| **LOC120761903** | Chr20 | 10335743 | 10336250 | Yes | Yes | Yes | Yes | Yes |
| **LOC120762994** | Chr25 | 490006 | 498600 | Yes | Yes | Yes | Yes | Yes |
| **LOC120763337** | Chr26 | 1789240 | 1790822 | Yes | Yes | Yes | Yes | Yes |
| **LOC120748378** | unplaced_BUSCO_68766at7742 | 56862 | 57053 | Yes | Yes | Yes | Yes | Yes |
| **LOC120751923** | Chr4 | 17855658 | 17858155 | Yes | Yes | Yes | Yes | Yes |
| **LOC120752095** | Chr4 | 61191185 | 61192875 | Yes | Yes | Yes | Yes | Yes |
| **LOC120756742** | Chr9 | 3622228 | 3624220 | Yes | Yes | Yes | Yes | Yes |
| **LOC120764913** | ChrW | 10017182 | 10018707 | Yes | Yes | Yes | Yes | Yes |
| **LOC120747603** | ChrW_unloc_2 | 18652 | 32480 | Yes | Yes | Yes | Yes | Yes |
| **CUL4B** | ChrZ | 82756612 | 82781215 | Yes | Yes | Yes | Yes | Yes |
| **LAMP2** | ChrZ | 82790897 | 82806511 | Yes | Yes | Yes | Yes | Yes |
| **ATP1B4** | ChrZ | 82805379 | 82816088 | Yes | Yes | Yes | Yes | Yes |
| **STEEP1** | ChrZ | 82847048 | 82849849 | Yes | Yes | Yes | Yes | Yes |
| **LOC120747646** | scaffold_126_arrow_ctg1 | 85758 | 87812 | Yes | Yes | Yes | Yes | Yes |
| **LOC120747659** | scaffold_137_arrow_ctg1 | 76806 | 78660 | Yes | Yes | Yes | Yes | Yes |
| **LOC120747763** | scaffold_193_arrow_ctg1 | 57304 | 58832 | Yes | Yes | Yes | Yes | Yes |
| **LOC120747871** | scaffold_249_arrow_ctg1 | 34771 | 36805 | Yes | Yes | Yes | Yes | Yes |
| **LOC120747884** | scaffold_266_arrow_ctg1 | 697 | 2883 | Yes | Yes | Yes | Yes | Yes |
| **LOC120748023** | scaffold_441_arrow_ctg1 | 10477 | 22650 | Yes | Yes | Yes | Yes | Yes |
| **RAD9A** | scaffold_504_arrow_ctg1 | 11981 | 14132 | Yes | Yes | Yes | Yes | Yes |
| **LOC120748109** | scaffold_540_arrow_ctg1 | 6282 | 14202 | Yes | Yes | Yes | Yes | Yes |
| **LOC120748114** | scaffold_548_arrow_ctg1 | 238 | 4447 | Yes | Yes | Yes | Yes | Yes |
| **LOC120748129** | scaffold_575_arrow_ctg1 | 7268 | 8198 | Yes | Yes | Yes | Yes | Yes |
| **LOC120748179** | scaffold_640_arrow_ctg1 | 61 | 1058 | Yes | Yes | Yes | Yes | Yes |
| **LOC120748181** | scaffold_646_arrow_ctg1 | 529 | 1235 | Yes | Yes | Yes | Yes | Yes |
| **LOC120748185** | scaffold_654_arrow_ctg1 | 23 | 948 | Yes | Yes | Yes | Yes | Yes |
| **LOC120748194** | scaffold_86_arrow_ctg1 | 98225 | 102432 | Yes | Yes | Yes | Yes | Yes |
| **LOC120748335** | unplaced_BUSCO_395792at7742 | 1361 | 1925 | Yes | Yes | Yes | Yes | Yes |
| **CD58** | Chr2_unloc_BUSCO_191412at7742 | 25839 | 41921 | Yes | Yes | Yes | Yes | No |

| | | | | | | | | |
|---|---|---|---|---|---|---|---|---|
| **LOC120764642** | Chr33 | 434530 | 435463 | Yes | Yes | Yes | Yes | No |
| **RAD1** | unplaced_BUSCO_309129at7742_3 | 11501 | 16689 | Yes | Yes | Yes | Yes | No |
| **LOC120747688** | scaffold_163_arrow_ctg1 | 36300 | 49803 | Yes | Yes | Yes | No | Yes |
| **LOC120750678** | Chr3 | 111000000 | 111000000 | Yes | Yes | Yes | No | No |
| **LOC120747908** | scaffold_289_arrow_ctg1 | 756 | 1773 | Yes | Yes | Yes | No | No |
| **PLA1A** | Chr2_unloc_BUSCO_191412at7742 | 4679 | 20955 | Yes | Yes | No | Yes | Yes |
| **SLC13A3** | Chr16 | 8615365 | 8641520 | Yes | Yes | No | No | No |
| **LOC120747689** | scaffold_163_arrow_ctg1 | 15233 | 22553 | Yes | No | Yes | Yes | Yes |
| **LOC120748197** | scaffold_86_arrow_ctg1 | 285076 | 286016 | Yes | No | Yes | Yes | Yes |
| **TWIST1** | scaffold_430_arrow_ctg1 | 15454 | 17775 | Yes | No | Yes | Yes | No |
| **LOC120759927** | Chr16 | 13743962 | 13744586 | Yes | No | Yes | No | Yes |
| **LOC120747690** | scaffold_163_arrow_ctg1 | 69344 | 78692 | Yes | No | Yes | No | Yes |
| **PALB2** | Chr15_unloc_BUSCO_378949at774 | 3022 | 11951 | Yes | No | No | Yes | No |
| **PPP1CA** | scaffold_504_arrow_ctg1 | 14147 | 16808 | Yes | No | No | Yes | No |
| **SH3TC1** | Chr5 | 10140681 | 10181183 | Yes | No | No | No | No |
| **LOC120766120** | Chr15_unloc_BUSCO_378949at774 | 24071 | 34797 | Yes | No | No | No | No |
| **LOC120747932** | scaffold_311_arrow_ctg1 | 12923 | 14999 | Yes | No | No | No | No |
| **LOC120765838** | ChrZ | 77352642 | 77357666 | No | Yes | Yes | Yes | Yes |
| **LOC120748132** | scaffold_583_arrow_ctg1 | 750 | 9292 | No | Yes | Yes | Yes | Yes |
| **LOC120748183** | scaffold_649_arrow_ctg1 | 3677 | 5481 | No | Yes | Yes | No | Yes |
| **LOC120748176** | scaffold_628_arrow_ctg1 | 1164 | 2097 | No | Yes | Yes | No | No |
| **SLC25A43** | ChrZ | 82859309 | 82875201 | No | Yes | No | Yes | Yes |
| **BRIX1** | unplaced_BUSCO_309129at7742_3 | 16820 | 21438 | No | Yes | No | Yes | Yes |
| **LOC120748261** | unplaced_BUSCO_320379at7742 | 28 | 3547 | No | Yes | No | Yes | Yes |
| **LOC120761627** | Chr1 | 13949271 | 13970981 | No | Yes | No | Yes | No |
| **NOX1** | ChrZ | 89770143 | 89774759 | No | Yes | No | Yes | No |
| **LOC120747675** | scaffold_152_arrow_ctg1 | 54322 | 55255 | No | Yes | No | Yes | No |
| **LOC120747947** | scaffold_329_arrow_ctg1 | 563 | 4814 | No | Yes | No | Yes | No |
| **GADD45B** | Chr26 | 1785190 | 1786920 | No | Yes | No | No | Yes |
| **LOC120748057** | scaffold_474_arrow_ctg1 | 3637 | 20235 | No | Yes | No | No | Yes |
| **LOC120764643** | Chr33 | 28663 | 34941 | No | Yes | No | No | No |
| **LOC120747790** | scaffold_203_arrow_ctg1 | 56621 | 59203 | No | Yes | No | No | No |
| **LOC120766103** | Chr5_unloc_1 | 170250 | 170565 | No | Yes | No | No | No |
| **LOC120766109** | Chr5_unloc_BUSCO_144578at7742 | 2692 | 9570 | No | Yes | No | No | No |
| **LOC120747677** | scaffold_155_arrow_ctg1 | 43854 | 46597 | No | Yes | No | No | No |

| | | | | | | | | |
|---|---|---|---|---|---|---|---|---|
| **PARVG** | Chr4 | 61158264 | 61179876 | No | No | Yes | Yes | No |
| **LOC120748047** | scaffold_464_arrow_ctg1 | 17163 | 18245 | No | No | Yes | No | No |
| **NIPAL1** | Chr5 | 25791932 | 25806624 | No | No | Yes | No | No |
| **LOC120755382** | Chr7 | 36755753 | 36759790 | No | No | Yes | No | No |
| **LOC120747845** | scaffold_224_arrow_ctg1 | 45278 | 46635 | No | No | Yes | No | No |
| **CDT1** | Chr11_unloc_BUSCO_236895at7742 | 31818 | 36075 | No | No | No | Yes | No |
| **DIRAS1** | Chr26 | 1717951 | 1721194 | No | No | No | Yes | No |
| **SLC13A1** | Chr4 | 21965236 | 21996564 | No | No | No | Yes | No |
| **STXBP3** | Chr9 | 21205266 | 21227560 | No | No | No | Yes | No |
| **LOC120765639** | ChrZ | 20672592 | 20674931 | No | No | No | Yes | No |
| **LOC120747954** | scaffold_335_arrow_ctg1 | 16733 | 18873 | No | No | No | Yes | No |
| **LOC120748072** | scaffold_490_arrow_ctg1 | 314 | 3965 | No | No | No | Yes | No |
| **FAM104A** | Chr18 | 10813503 | 10823454 | No | No | No | No | Yes |
| **LOC120759993** | Chr1 | 154406 | 154745 | No | No | No | No | No |
| **LOC120749544** | Chr1 | 253157 | 253484 | No | No | No | No | No |
| **LRATD2** | Chr1 | 10223237 | 10230055 | No | No | No | No | No |
| **CTU2** | Chr11_unloc_BUSCO_236895at7742 | 27 | 3835 | No | No | No | No | No |
| **PPP1R15B** | Chr24 | 6447243 | 6452951 | No | No | No | No | No |
| **SHD** | Chr26 | 891828 | 894720 | No | No | No | No | No |
| **LOC120749898** | Chr3 | 7448254 | 7466220 | No | No | No | No | No |
| **LOC120751209** | Chr3 | 7467450 | 7468480 | No | No | No | No | No |
| **HOXC9** | Chr30 | 598175 | 601649 | No | No | No | No | No |
| **LOC120752184** | Chr4 | 36712810 | 36726812 | No | No | No | No | No |
| **LOC120752257** | Chr4 | 63434696 | 63436520 | No | No | No | No | No |
| **RELL1** | Chr5 | 22137235 | 22168752 | No | No | No | No | No |
| **LOC120766108** | Chr5_unloc_BUSCO_144578at7742 | 9855 | 41381 | No | No | No | No | No |
| **LOC120755806** | Chr8 | 16249639 | 16311058 | No | No | No | No | No |
| **NDUFB8** | Chr8 | 17219407 | 17229691 | No | No | No | No | No |
| **LOC120756511** | Chr9 | 31172209 | 31183693 | No | No | No | No | No |
| **LOC120764849** | ChrW | 6933601 | 7016241 | No | No | No | No | No |
| **ZC3H12B** | ChrW | 11998365 | 12080752 | No | No | No | No | No |
| **GIN1** | ChrZ | 50717682 | 50738540 | No | No | No | No | No |
| **LOC120765297** | ChrZ | 85641149 | 85658756 | No | No | No | No | No |
| **LOC120747762** | scaffold_193_arrow_ctg1 | 2743 | 8737 | No | No | No | No | No |
| **RFC5** | scaffold_640_arrow_ctg1 | 1446 | 3854 | No | No | No | No | No |

| | | | | | | | | |
|---|---|---|---|---|---|---|---|---|
| **LOC120748189** | scaffold_86_arrow_ctg1 | 171379 | 185177 | No | No | No | No | No |
| **LOC120748264** | unplaced_BUSCO_320379at7742 | 20480 | 27730 | No | No | No | No | No |
| **DMAC2L** | unplaced_BUSCO_363461at7742 | 5121 | 9947 | No | No | No | No | No |
| **LOC120748352** | unplaced_BUSCO_419118at7742 | 10391 | 14070 | No | No | No | No | No |
| **LOC120756062** | Chr1 | 80185770 | 80187089 | No | No | No | No | No |
| **LOC120760288** | Chr1 | 84856619 | 84873628 | No | No | No | No | No |
| **PIEZO1** | Chr11_unloc_BUSCO_236895at774[illegible] | 3841 | 29418 | No | No | No | No | No |
| **LOC120759332** | Chr14 | 16142177 | 16145937 | No | No | No | No | No |
| **LOC120759274** | Chr14 | 16731517 | 16736498 | No | No | No | No | No |
| **DCTN5** | Chr15_unloc_BUSCO_378949at774[illegible] | 6 | 2987 | No | No | No | No | No |
| **NDUFAB1** | Chr15_unloc_BUSCO_378949at774[illegible] | 12951 | 15779 | No | No | No | No | No |
| **LOC120766116** | Chr15_unloc_BUSCO_378949at774[illegible] | 17479 | 24094 | No | No | No | No | No |
| **COG1** | Chr18 | 10807018 | 10814892 | No | No | No | No | No |
| **LOC120766062** | Chr1_unloc_2 | 274247 | 275441 | No | No | No | No | No |
| **LOC120766122** | Chr21_unloc_BUSCO_410670at774[illegible] | 19199 | 29475 | No | No | No | No | No |
| **LOC120766123** | Chr21_unloc_BUSCO_410670at774[illegible] | 29468 | 34379 | No | No | No | No | No |
| **LOC120763441** | Chr26 | 795285 | 803716 | No | No | No | No | No |
| **FSD1** | Chr26 | 895400 | 898584 | No | No | No | No | No |
| **SGTA** | Chr26 | 1698949 | 1710135 | No | No | No | No | No |
| **SLC39A3** | Chr26 | 1711148 | 1716084 | No | No | No | No | No |
| **GNG7** | Chr26 | 1725313 | 1778391 | No | No | No | No | No |
| **LOC120763340** | Chr26 | 1731296 | 1742325 | No | No | No | No | No |
| **LOC120763566** | Chr26 | 1882953 | 1899810 | No | No | No | No | No |
| **COPA** | Chr29_unloc_BUSCO_136144at774[illegible] | 17099 | 44382 | No | No | No | No | No |
| **LOC120747567** | Chr29_unloc_BUSCO_136144at774[illegible] | 69323 | 72932 | No | No | No | No | No |
| **LOC120747575** | Chr29_unloc_BUSCO_136144at774[illegible] | 79308 | 79908 | No | No | No | No | No |
| **LOC120747568** | Chr29_unloc_BUSCO_136144at774[illegible] | 87405 | 88611 | No | No | No | No | No |
| **LOC120747569** | Chr29_unloc_BUSCO_136144at774[illegible] | 103113 | 105397 | No | No | No | No | No |
| **LOC120747570** | Chr29_unloc_BUSCO_136144at774[illegible] | 126647 | 127286 | No | No | No | No | No |
| **LOC120747571** | Chr29_unloc_BUSCO_136144at774[illegible] | 128395 | 131700 | No | No | No | No | No |
| **FKBP11** | Chr30_unloc_BUSCO_126329at774[illegible] | 1090 | 4073 | No | No | No | No | No |
| **CCDC65** | Chr30_unloc_BUSCO_126329at774[illegible] | 3426 | 7315 | No | No | No | No | No |
| **RND1** | Chr30_unloc_BUSCO_126329at774[illegible] | 7532 | 14149 | No | No | No | No | No |
| **DDX23** | Chr30_unloc_BUSCO_126329at774[illegible] | 14718 | 22950 | No | No | No | No | No |
| **CACNB3** | Chr30_unloc_BUSCO_126329at774[illegible] | 23273 | 27131 | No | No | No | No | No |

| | | | | | | | | |
|---|---|---|---|---|---|---|---|---|
| **LOC120764785** | Chr37 | 234642 | 242188 | No | No | No | No | No |
| **DPF2** | scaffold_121_arrow_ctg1 | 76635 | 86997 | No | No | No | No | No |
| **HNRNPUL2** | unplaced_BUSCO_68766at7742 | 43949 | 55624 | No | No | No | No | No |
| **LOC120748246** | unplaced_BUSCO_267345at7742 | 23939 | 27539 | No | No | No | No | No |
| **LOC120748243** | unplaced_BUSCO_267345at7742 | 30856 | 35115 | No | No | No | No | No |
| **LOC120748244** | unplaced_BUSCO_267345at7742 | 36156 | 41999 | No | No | No | No | No |
| **LOC120748245** | unplaced_BUSCO_267345at7742 | 45388 | 52535 | No | No | No | No | No |
| **LOC120748052** | scaffold_473_arrow_ctg1 | 774 | 2043 | No | No | No | No | No |
| **LOC120748265** | unplaced_BUSCO_330367at7742 | 14174 | 28933 | No | No | No | No | No |
| **LOC120748093** | scaffold_510_arrow_ctg1 | 142 | 6656 | No | No | No | No | No |
| **LOC120748094** | scaffold_510_arrow_ctg1 | 6781 | 17270 | No | No | No | No | No |
| **SAE1** | scaffold_529_arrow_ctg1 | 5 | 7681 | No | No | No | No | No |
| **LOC120748102** | scaffold_529_arrow_ctg1 | 8843 | 15877 | No | No | No | No | No |
| **PSMD8** | scaffold_618_arrow_ctg1 | 1824 | 6772 | No | No | No | No | No |
| **PRDM13** | Chr3_unloc_BUSCO_162664at7742 | 1286 | 8318 | No | No | No | No | No |
| **TFAP2D** | Chr3_unloc_BUSCO_208931at7742 | 4397 | 31177 | No | No | No | No | No |
| **LMF2** | Chr4 | 501875 | 516741 | No | No | No | No | No |
| **CEP41** | Chr4 | 826514 | 849887 | No | No | No | No | No |
| **NET1** | Chr4 | 975770 | 1075324 | No | No | No | No | No |
| **UCN3** | Chr4 | 1105680 | 1114236 | No | No | No | No | No |
| **ALMS1** | Chr5 | 1063355 | 1123879 | No | No | No | No | No |
| **IDH3B** | Chr5 | 1126090 | 1153697 | No | No | No | No | No |
| **RTKN** | Chr5 | 1253253 | 1267376 | No | No | No | No | No |
| **LOC120766107** | Chr5_unloc_3 | 88470 | 116564 | No | No | No | No | No |
| **LOC120753593** | Chr6 | 666260 | 690013 | No | No | No | No | No |
| **CARNS1** | Chr6 | 60024326 | 60047047 | No | No | No | No | No |
| **KLHL28** | Chr6_unloc_BUSCO_122064at7742 | 6242 | 17679 | No | No | No | No | No |
| **NMI** | Chr7 | 1587277 | 1596707 | No | No | No | No | No |
| **RBM43** | Chr7 | 1597824 | 1602414 | No | No | No | No | No |
| **PTGER3** | Chr9 | 739111 | 747574 | No | No | No | No | No |
| **IL12RB2** | Chr9 | 1163880 | 1181555 | No | No | No | No | No |
| **GLRX2** | Chr9 | 23624059 | 23630882 | No | No | No | No | No |
| **LOC120764966** | ChrW | 3918774 | 4055914 | No | No | No | No | No |
| **LOC120765517** | ChrZ | 20668559 | 20669970 | No | No | No | No | No |
| **NRG1** | ChrZ | 73649768 | 74054191 | No | No | No | No | No |

| | | | | | | | | |
|---|---|---|---|---|---|---|---|---|
| **SLC25A5** | ChrZ | 82855098 | 82859166 | No | No | No | No | No |
| **LOC120765349** | ChrZ | 85638035 | 85640860 | No | No | No | No | No |
| **LOC120747613** | scaffold_106_arrow_ctg1 | 93554 | 134290 | No | No | No | No | No |
| **LOC120747674** | scaffold_151_arrow_ctg1 | 88299 | 91869 | No | No | No | No | No |
| **LOC120747680** | scaffold_156_arrow_ctg1 | 89695 | 92689 | No | No | No | No | No |
| **LOC120747703** | scaffold_176_arrow_ctg1 | 75382 | 77551 | No | No | No | No | No |
| **LOC120747776** | scaffold_201_arrow_ctg1 | 52 | 1445 | No | No | No | No | No |
| **LOC120747781** | scaffold_201_arrow_ctg1 | 52334 | 62252 | No | No | No | No | No |
| **LOC120747779** | scaffold_201_arrow_ctg1 | 62378 | 63398 | No | No | No | No | No |
| **LOC120747785** | scaffold_201_arrow_ctg1 | 65084 | 66983 | No | No | No | No | No |
| **LOC120747789** | scaffold_203_arrow_ctg1 | 2128 | 10630 | No | No | No | No | No |
| **RABGGTA** | scaffold_207_arrow_ctg1 | 27361 | 49048 | No | No | No | No | No |
| **LOC120747847** | scaffold_224_arrow_ctg1 | 52068 | 5.33E+04 | No | No | No | No | No |
| **LOC120747874** | scaffold_252_arrow_ctg1 | 16013 | 30908 | No | No | No | No | No |
| **LOC120747873** | scaffold_252_arrow_ctg1 | 37758 | 40446 | No | No | No | No | No |
| **LOC120747872** | scaffold_252_arrow_ctg1 | 40526 | 49953 | No | No | No | No | No |
| **LOC120747890** | scaffold_275_arrow_ctg1 | 1546 | 43133 | No | No | No | No | No |
| **LOC120747913** | scaffold_292_arrow_ctg1 | 616 | 945 | No | No | No | No | No |
| **LOC120747926** | scaffold_306_arrow_ctg1 | 3 | 33728 | No | No | No | No | No |
| **LOC120747933** | scaffold_311_arrow_ctg1 | 9560 | 11633 | No | No | No | No | No |
| **BRD4** | scaffold_330_arrow_ctg1 | 752 | 37042 | No | No | No | No | No |
| **LOC120747950** | scaffold_331_arrow_ctg1 | 1584 | 33915 | No | No | No | No | No |
| **SELENBP1** | scaffold_360_arrow_ctg1 | 12113 | 31728 | No | No | No | No | No |
| **LOC120747976** | scaffold_362_arrow_ctg1 | 19525 | 30691 | No | No | No | No | No |
| **LOC120747993** | scaffold_382_arrow_ctg1 | 12686 | 13620 | No | No | No | No | No |
| **COL9A2** | scaffold_392_arrow_ctg1 | 70 | 8570 | No | No | No | No | No |
| **LOC120747998** | scaffold_392_arrow_ctg1 | 16925 | 27585 | No | No | No | No | No |
| **LOC120748012** | scaffold_433_arrow_ctg1 | 194 | 10667 | No | No | No | No | No |
| **POLR1G** | scaffold_433_arrow_ctg1 | 11814 | 13678 | No | No | No | No | No |
| **ERCC1** | scaffold_433_arrow_ctg1 | 13569 | 19064 | No | No | No | No | No |
| **LOC120748013** | scaffold_433_arrow_ctg1 | 19036 | 20838 | No | No | No | No | No |
| **PPP1R37** | scaffold_447_arrow_ctg1 | 256 | 22788 | No | No | No | No | No |
| **LOC120748039** | scaffold_454_arrow_ctg1 | 4172 | 8949 | No | No | No | No | No |
| **LOC120748040** | scaffold_456_arrow_ctg1 | 10181 | 11973 | No | No | No | No | No |
| **G6PC3** | scaffold_460_arrow_ctg1 | 11687 | 15553 | No | No | No | No | No |

| | | | | | | | | |
|---|---|---|---|---|---|---|---|---|
| **LOC120748046** | scaffold_461_arrow_ctg1 | 1888 | 20466 | No | No | No | No | No |
| **LOC120748051** | scaffold_469_arrow_ctg1 | 936 | 1869 | No | No | No | No | No |
| **LOC120748063** | scaffold_479_arrow_ctg1 | 2355 | 14876 | No | No | No | No | No |
| **ERCC2** | scaffold_485_arrow_ctg1 | 336 | 12132 | No | No | No | No | No |
| **LOC120748068** | scaffold_485_arrow_ctg1 | 13632 | 19102 | No | No | No | No | No |
| **PHF23** | scaffold_492_arrow_ctg1 | 525 | 7547 | No | No | No | No | No |
| **LOC120748076** | scaffold_492_arrow_ctg1 | 8177 | 12484 | No | No | No | No | No |
| **SNAPC2** | scaffold_492_arrow_ctg1 | 13626 | 18339 | No | No | No | No | No |
| **LOC120748086** | scaffold_504_arrow_ctg1 | 6 | 7473 | No | No | No | No | No |
| **EML3** | scaffold_532_arrow_ctg1 | 6637 | 15771 | No | No | No | No | No |
| **LOC120748131** | scaffold_580_arrow_ctg1 | 2140 | 9849 | No | No | No | No | No |
| **LOC120748137** | scaffold_592_arrow_ctg1 | 4454 | 7197 | No | No | No | No | No |
| **LOC120748146** | scaffold_613_arrow_ctg1 | 1988 | 5611 | No | No | No | No | No |
| **LOC120748173** | scaffold_620_arrow_ctg1 | 251 | 3909 | No | No | No | No | No |
| **COPE** | scaffold_620_arrow_ctg1 | 4070 | 7654 | No | No | No | No | No |
| **LOC120748182** | scaffold_648_arrow_ctg1 | 968 | 3146 | No | No | No | No | No |
| **LOC120748184** | scaffold_652_arrow_ctg1 | 386 | 5106 | No | No | No | No | No |
| **LOC120748260** | unplaced_BUSCO_309958at7742 | 847 | 3918 | No | No | No | No | No |
| **IBA57** | unplaced_BUSCO_309958at7742 | 11061 | 15010 | No | No | No | No | No |
| **LOC120748259** | unplaced_BUSCO_309958at7742 | 28841 | 38924 | No | No | No | No | No |
| **LOC120748276** | unplaced_BUSCO_339203at7742 | 93306 | 97495 | No | No | No | No | No |
| **CDKL1** | unplaced_BUSCO_363461at7742 | 1013 | 4342 | No | No | No | No | No |
| **L2HGDH** | unplaced_BUSCO_363461at7742 | 9947 | 21774 | No | No | No | No | No |
| **LOC120748297** | unplaced_BUSCO_363461at7742 | 24158 | 64209 | No | No | No | No | No |
| **LOC120748348** | unplaced_BUSCO_407723at7742 | 346 | 2449 | No | No | No | No | No |
| **PEA15** | unplaced_BUSCO_407723at7742 | 3766 | 11884 | No | No | No | No | No |
| **LOC120748353** | unplaced_BUSCO_419118at7742 | 6 | 4341 | No | No | No | No | No |
| **MRPL20** | unplaced_BUSCO_419118at7742 | 5053 | 8268 | No | No | No | No | No |
| **LOC120748359** | unplaced_BUSCO_420085at7742 | 1250 | 14384 | No | No | No | No | No |
| **VPS16** | unplaced_BUSCO_63198at7742 | 1350 | 39112 | No | No | No | No | No |

**Table S24 - Geographical origin, type of genomic data, population sizes and sequencing coverage (based on genome size) for all genomic data analysed.** ds: dataset

| Identifier | Publication | Data type | N individuals | Location(s) | Subspecies | Average sequencing coverage |
|---|---|---|---|---|---|---|
| **ds1** | This publication | Pacbio HiFi WGS | 5 | Italy | *H. r. rustica* | ~20x |
| **ds2.1** | Schield et al., 2021 | Illumina WGS | 34 | China, Japan, Russia | *H. r. gutturalis* | ~6.68x |
| **ds2.2** | Schield et al., 2021 | Illumina WGS | 25 | China, Mongolia, Morocco, Russia | *H. r. rustica* | ~6.08x |
| **ds2.3** | Schield et al., 2021 | Illumina WGS | 8 | Israel | *H. r. transitiva* | ~7.92x |
| **ds2.4** | Schield et al., 2021 | Illumina WGS | 10 | Russia | *H. r. tytleri* | ~6.35x |
| **ds2.5** | Schield et al., 2021 | Illumina WGS | 29 | Mongolia, Russia (hybrid zone) | *H. r. gutturalis x H. r. tytleri* | ~6.99x |
| **ds2.6** | Schield et al., 2021 | Illumina WGS | 21 | China (hybrid zone) | *H. r. rustica x H. r. gutturalis* | ~7.27x |
| **ds2.7** | Schield et al., 2021 | Illumina WGS | 16 | Russia (hybrid zone) | *H. r. rustica x H. r. tytleri* | ~5.79x |
| **ds3.1.1** | Smith et al., 2018 | Illumina WGS | 8 | Egypt | *H. r. savignii* | ~6.1x |
| **ds3.1.2** | Smith et al., 2018 | Illumina WGS | 8 | Colorado, USA | *H. r. erythrogaster* | ~7.83x |
| **ds3.2.1** | Smith et al., 2018 | ddRAD | 36 | Egypt | *H. r. savignii* | ~0.12x |
| **ds3.2.2** | Smith et al., 2018 | ddRAD | 26 | Colorado, USA | *H. r. erythrogaster* | ~0.08x |
| **ds4** | Scordato et al., 2017 | ddRAD | 533 | Russia (west-to-east transect) | *H. r. rustica; H. r. tytleri; H. r. gutturalis* | ~0.15x |
| **ds5** | Von Ronn et al., 201( | ddRAD | 216 | Transect from Sweden to Switzerland | *H. r. rustica* | ~0.33x |
| **ds6.1** | Safran et al., 2016 | ddRAD | 145 | Colorado, USA | *H. r. erythrogaster* | ~0.02x |
| **ds6.2** | Safran et al., 2016 | ddRAD | 27 | Ithaca, New York ,USA | *H. r. erythrogaster* | ~0.03x |
| **ds6.3** | Safran et al., 2016 | ddRAD | 24 | Czech Republic | *H. r. rustica* | ~0.02x |
| **ds6.4** | Safran et al., 2016 | ddRAD | 16 | Romania | *H. r. rustica* | ~0.02x |
| **ds6.5** | Safran et al., 2016 | ddRAD | 50 | Turkey | *H. r. rustica* | ~0.02x |
| **ds6.6** | Safran et al., 2016 | ddRAD | 26 | United Kingdom | *H. r. rustica* | ~0.01x |
| **ds6.7** | Safran et al., 2016 | ddRAD | 45 | Israel | *H. r. transitiva* | ~0.02x |
| **ds6.8** | Safran et al., 2016 | ddRAD | 18 | Taiwan | *H. r. gutturalis* | ~0.03x |

**Table S25 - SNP counts and density across chromosomes for all datasets analysed.** HiFi 20x:  ds1 at 20x sequencing coverage (except for A1 sample whose initial coverage was 15x); Hifi 5x: ds1 downsampled to 5x; HiFi intersection 20x_5x: after downsampling to 5x, 20x ds was considered as truth set and used for intersection; all Illumina WGS data: ds2 and ds3.1; all ddRAD data: ds3.2 through ds6. All samples from the same sequencing technology were considered together. No SNPs were detected on chr39.

| chr | type | chr size (bp) | total number of SNPs (Hifi 20x) | weighted average SNPS density/10kb (Hifi 20x) | total number of SNPs (Hifi 5x) | weighted average SNPS density/10kb (Hifi 5x) | total number of SNPs (HiFi intersection 20x_5x) | weighted avg SNPS density/10kb (HiFi intersection 20x_5x) | total number of SNPs (all Illumina WGS data) | weighted average SNPS density/10kb (all Illumina WGS data) | total number of SNPs (all ddRAD data) | weighted average SNPS density/10kb (all ddRAD data) |
|---|---|---|---|---|---|---|---|---|---|---|---|---|
| **chr1** | macrochromosome | 156,035,725 | 2,235,865 | 143.29 | 1,712,439 | 109.75 | 1,552,209 | 99.48 | 2,378,916 | 152.46 | 12,046 | 0.77 |
| **chr2** | macrochromosome | 119,023,421 | 1,874,228 | 157.47 | 1,430,648 | 120.20 | 1,302,118 | 109.40 | 2,033,905 | 170.88 | 9,613 | 0.81 |
| **chr3** | macrochromosome | 116,801,625 | 1,713,055 | 146.66 | 1,327,408 | 113.65 | 1,200,944 | 102.82 | 1,915,698 | 164.01 | 9,641 | 0.83 |
| **chrZ** | macrochromosome | 90,132,487 | 286,586 | 31.80 | 24,286 | 2.69 | 21,094 | 2.34 | 480,430 | 53.30 | 2,452 | 0.27 |
| **chr4** | macrochromosome | 76,187,387 | 1,002,509 | 131.58 | 765,664 | 100.50 | 692,717 | 90.92 | 1,144,428 | 150.21 | 5,704 | 0.75 |
| **chr5** | macrochromosome | 73,257,097 | 1,166,751 | 159.26 | 903,652 | 123.35 | 818,574 | 111.74 | 1,232,652 | 168.26 | 6,145 | 0.84 |
| **chr6** | macrochromosome | 63,258,489 | 877,095 | 138.65 | 674,193 | 106.58 | 609,627 | 96.37 | 1,011,284 | 159.87 | 5,015 | 0.79 |
| **chr7** | intermediate | 38,459,648 | 659,436 | 171.46 | 511,137 | 132.90 | 460,616 | 119.77 | 752,364 | 195.62 | 3,793 | 0.99 |
| **chr8** | intermediate | 36,085,389 | 638,569 | 176.96 | 493,525 | 136.77 | 445,824 | 123.55 | 739,420 | 204.91 | 3,555 | 0.99 |
| **chrW** | intermediate | 31,704,074 | 10,022 | 3.16 | 3,727 | 1.18 | 3,215 | 1.01 | 4,364 | 1.38 | 24 | 0.008 |
| **chr9** | intermediate | 31,262,510 | 535,687 | 171.35 | 413,729 | 132.34 | 372,503 | 119.15 | 609,915 | 195.09 | 3,320 | 1.06 |
| **chr10** | intermediate | 25,880,253 | 488,472 | 188.74 | 373,477 | 144.31 | 339,236 | 131.08 | 565,813 | 218.63 | 3,049 | 1.18 |
| **chr11** | intermediate | 21,491,857 | 329,758 | 153.43 | 249,183 | 115.94 | 222,277 | 103.42 | 372,113 | 173.14 | 2,085 | 0.97 |
| **chr12** | intermediate | 20,890,524 | 382,823 | 183.25 | 291,496 | 139.54 | 265,602 | 127.14 | 444,832 | 212.93 | 2,199 | 1.05 |
| **chr13** | intermediate | 20,272,128 | 355,942 | 175.58 | 273,164 | 134.75 | 247,645 | 122.16 | 421,690 | 208.01 | 2,106 | 1.04 |
| **chr14** | microchromosome | 18,810,845 | 348,540 | 185.29 | 259,966 | 138.20 | 237,131 | 126.06 | 394,972 | 209.97 | 1,769 | 0.94 |
| **chr15** | microchromosome | 16,541,138 | 311,453 | 188.29 | 236,045 | 142.70 | 213,204 | 128.89 | 365,592 | 221.02 | 1,866 | 1.13 |
| **chr16** | microchromosome | 15,277,844 | 276,785 | 181.17 | 206,184 | 134.95 | 190,293 | 124.55 | 325,289 | 212.91 | 1,460 | 0.96 |
| **chr17** | microchromosome | 13,985,943 | 254,428 | 181.90 | 191,298 | 136.77 | 173,657 | 124.15 | 313,512 | 224.14 | 1,559 | 1.11 |
| **chr18** | microchromosome | 12,073,725 | 201,040 | 166.51 | 142,265 | 117.83 | 130,382 | 107.99 | 232,814 | 192.82 | 1,169 | 0.97 |
| **chr19** | microchromosome | 11,382,101 | 208,474 | 183.16 | 154,764 | 135.97 | 142,984 | 125.62 | 244,586 | 214.89 | 1,327 | 1.17 |
| **chr20** | microchromosome | 11,194,341 | 210,406 | 187.94 | 152,143 | 135.91 | 139,941 | 125.01 | 250,938 | 224.15 | 1,183 | 1.06 |
| **chr21** | microchromosome | 9,617,204 | 193,476 | 201.16 | 143,517 | 149.22 | 132,068 | 137.31 | 213,087 | 221.56 | 1,094 | 1.14 |
| **chr22** | microchromosome | 7,507,825 | 124,436 | 165.72 | 88,747 | 118.20 | 79,904 | 106.42 | 150,522 | 200.49 | 733 | 0.98 |
| **chr23** | microchromosome | 7,098,401 | 145,259 | 204.64 | 97,972 | 138.02 | 91,185 | 128.46 | 164,593 | 231.87 | 762 | 1.07 |
| **chr24** | microchromosome | 6,843,954 | 123,102 | 179.86 | 82,037 | 119.86 | 74,420 | 108.73 | 130,777 | 191.07 | 606 | 0.89 |
| **chr25** | microchromosome | 6,778,862 | 120,232 | 177.35 | 80,810 | 119.20 | 71,673 | 105.72 | 134,800 | 198.84 | 708 | 1.04 |
| **chr26** | microchromosome | 5,553,549 | 90,808 | 163.51 | 60,915 | 109.69 | 55,033 | 99.10 | 96,966 | 174.59 | 463 | 0.83 |
| **chr28** | microchromosome | 5,297,670 | 79,828 | 150.68 | 50,934 | 96.14 | 44,796 | 84.56 | 77,700 | 146.67 | 485 | 0.92 |
| **chr27** | microchromosome | 5,236,451 | 87,168 | 166.46 | 61,632 | 117.65 | 54,924 | 104.89 | 87,208 | 166.53 | 441 | 0.84 |
| **chr29** | microchromosome | 2,102,120 | 34,416 | 163.72 | 21,680 | 103.13 | 18,928 | 90.04 | 31,055 | 147.69 | 105 | 0.50 |
| **chr30** | microchromosome | 1,648,998 | 16,907 | 102.48 | 9,750 | 59.11 | 7,934 | 48.10 | 15,060 | 91.33 | 62 | 0.37 |
| **chr31** | microchromosome | 1,590,086 | 6,559 | 41.25 | 5,174 | 32.54 | 3,958 | 24.89 | 2,527 | 15.89 | 3 | 0.02 |
| **chr32** | microchromosome | 784,579 | 5,733 | 73.07 | 4,173 | 53.18 | 3,175 | 40.47 | 2,104 | 26.82 | 9 | 0.11 |
| **chr34** | microchromosome | 606,149 | 1,925 | 31.76 | 1,260 | 20.72 | 958 | 15.80 | 1,104 | 18.21 | 3 | 0.05 |

| | | | | | | | | | | | | |
|---|---|---|---|---|---|---|---|---|---|---|---|---|
| **chr35** | microchromosome | 523,230 | 7,090 | 134.46 | 4,775 | 90.85 | 3,817 | 72.61 | 8,220 | 157.10 | 44 | 0.84 |
| **chr33** | microchromosome | 437,724 | 720 | 16.45 | 888 | 20.29 | 288 | 6.58 | 104 | 2.37 | 0 | 0.00 |
| **chr36** | microchromosome | 338,027 | 1,741 | 51.34 | 1,255 | 36.91 | 726 | 21.40 | 1,812 | 53.47 | 6 | 0.18 |
| **chr37** | microchromosome | 276,370 | 4,050 | 144.36 | 2,229 | 79.06 | 1,896 | 67.37 | 2,675 | 94.43 | 9 | 0.33 |
| **chr38** | microchromosome | 220,485 | 265 | 12.02 | 157 | 7.12 | 42 | 1.9 | 408 | 18.5 | 11 | 0.5 |
| **chr39** | microchromosome | 65,965 | / | / | / | / | / | / | / | | / | / |

**Table S26 - Whole genome LD values in Illumina WGS data from dataset2 and dataset3.1.** Average r^2 values calculated according to physical distance (kbps) between SNPs are reported, as well as the total number of computed marker pairs per interval.

| Distance (kbps) | average R^2 (*H. r. erythrogaster*) | SNP pairs number (*H. r. erythrogaster*) | average R^2 (*H. r. savignii*) | SNP pairs number (*H. r. savignii*) | average R^2 (*H. r. gutturalis*) | SNP pairs number (*H. r. gutturalis*) | average R^2 (*H. r. rustica*) | SNP pairs number (*H. r. rustica*) | average R^2 (*H. r. transitiva*) | SNP pairs number (*H. r. transitiva*) | average R^2 (*H. r. tytleri*) | SNP pairs number (*H. r. tytleri*) | average R^2 (*H. r. gutturalis x tytleri*) | SNP pairs number (*H. r. gutturalis x tytleri*) | average R^2 (*H. r. rustica x gutturalis*) | SNP pairs number (*H. r. rustica x gutturalis*) | average R^2 (*H. r. rustica x tytleri*) | SNP pairs number (*H. r. rustica x tytleri*) |
|---|---|---|---|---|---|---|---|---|---|---|---|---|---|---|---|---|---|---|
| **0-1** | 0.133933316 | 36201043 | 0.137887383 | 15146532 | 0.068242657 | 26465560 | 0.064058402 | 9175544 | 0.131788753 | 46247086 | 0.112951759 | 75120088 | 0.058990866 | 21619950 | 0.06985821 | 56990532 | 0.082580313 | 9870545 |
| **1-2** | 0.120482536 | 30323625 | 0.12277292 | 11843527 | 0.047493452 | 21612283 | 0.046960604 | 6966282 | 0.12263391 | 39206127 | 0.105412774 | 63643430 | 0.042645131 | 17146557 | 0.057651246 | 49133054 | 0.066642206 | 7352148 |
| **2-3** | 0.117555281 | 30150507 | 0.120147712 | 11777665 | 0.042476921 | 21518830 | 0.043967217 | 6930137 | 0.120875302 | 39002584 | 0.103772147 | 63330664 | 0.039268547 | 17050844 | 0.05484585 | 48901104 | 0.063732561 | 7295972 |
| **3-4** | 0.115820389 | 30059766 | 0.118335219 | 11720790 | 0.039567761 | 21427963 | 0.042433322 | 6895407 | 0.119986327 | 38904478 | 0.102911017 | 63111137 | 0.037436743 | 16985421 | 0.053290424 | 48739086 | 0.062125758 | 7269437 |
| **4-5** | 0.114649637 | 30021085 | 0.11705706 | 11704810 | 0.037663073 | 21368053 | 0.041337234 | 6894311 | 0.119403089 | 38816407 | 0.102380456 | 62976090 | 0.036349977 | 16944793 | 0.052262699 | 48614231 | 0.061054312 | 7258929 |
| **5-6** | 0.113678417 | 29947333 | 0.116120769 | 11717452 | 0.036355291 | 21348540 | 0.04072646 | 6875591 | 0.118926589 | 38757262 | 0.101967239 | 62851874 | 0.035527579 | 16930186 | 0.051563494 | 48574136 | 0.060321146 | 7237932 |
| **6-7** | 0.112944089 | 29865409 | 0.115232036 | 11661500 | 0.035309818 | 21238491 | 0.040282745 | 6842902 | 0.118607684 | 38627984 | 0.101710051 | 62720385 | 0.034921744 | 16891625 | 0.050989223 | 48411169 | 0.059874146 | 7213697 |
| **7-8** | 0.112364537 | 29814122 | 0.114544539 | 11623595 | 0.034524775 | 21198088 | 0.039881319 | 6830459 | 0.118376254 | 38559048 | 0.101474003 | 62526804 | 0.034498186 | 16823343 | 0.050525292 | 48331347 | 0.05941701 | 7192768 |
| **8-9** | 0.111918728 | 29798972 | 0.113985165 | 11636240 | 0.033878352 | 21198459 | 0.039579057 | 6813282 | 0.118159241 | 38531878 | 0.101260249 | 62525537 | 0.034073363 | 16821510 | 0.050184477 | 48265553 | 0.059095024 | 7189622 |
| **9-10** | 0.111533081 | 29710522 | 0.113392116 | 11593510 | 0.03332682 | 21114409 | 0.039311418 | 6802627 | 0.11803996 | 38439784 | 0.101168552 | 62388202 | 0.033815623 | 16755443 | 0.049828232 | 48140181 | 0.058853611 | 7175265 |
| **10-11** | 0.11116807 | 29679543 | 0.112823189 | 11568991 | 0.032858036 | 21098156 | 0.039117817 | 6792599 | 0.117824832 | 38398410 | 0.101043504 | 62288844 | 0.033601888 | 16738537 | 0.049568527 | 48116608 | 0.05862874 | 7163965 |
| **11-12** | 0.110901683 | 29629762 | 0.112490212 | 11557396 | 0.032509601 | 21059418 | 0.038960231 | 6781093 | 0.117701697 | 38318876 | 0.100905232 | 62247211 | 0.033390209 | 16700936 | 0.049317463 | 48071252 | 0.058440487 | 7151871 |
| **12-13** | 0.110591165 | 29594947 | 0.1121869 | 11548471 | 0.032152987 | 21028268 | 0.038799741 | 6763606 | 0.117504252 | 38255148 | 0.100885453 | 62161774 | 0.033207437 | 16705183 | 0.04911653 | 47997122 | 0.058345637 | 7152549 |
| **13-14** | 0.110317789 | 29571499 | 0.111621311 | 11537812 | 0.031843845 | 21002444 | 0.03867448 | 6760538 | 0.11741959 | 38253238 | 0.100738119 | 62050943 | 0.03303232 | 16690438 | 0.048934801 | 47912356 | 0.058125549 | 7138956 |
| **14-15** | 0.110070627 | 29550367 | 0.111457621 | 11517256 | 0.031619749 | 20978534 | 0.038604937 | 6754509 | 0.117301225 | 38236497 | 0.100628133 | 62039835 | 0.032910831 | 16659139 | 0.048774451 | 47912321 | 0.058024721 | 7124406 |
| **15-16** | 0.10992551 | 29499398 | 0.111142708 | 11522540 | 0.031345422 | 20959512 | 0.038481697 | 6752609 | 0.117245786 | 38201120 | 0.100574699 | 61955970 | 0.032813443 | 16658273 | 0.04862034 | 47876807 | 0.057896835 | 7121544 |
| **16-17** | 0.10981851 | 29442931 | 0.110792252 | 11502581 | 0.031167566 | 20920028 | 0.03841121 | 6756101 | 0.117136821 | 38181067 | 0.100516896 | 61881312 | 0.032721981 | 16626983 | 0.048457128 | 47779297 | 0.057760652 | 7112215 |
| **17-18** | 0.109647563 | 29446052 | 0.110618056 | 11486440 | 0.030969479 | 20901375 | 0.038352339 | 6743327 | 0.117032807 | 38113822 | 0.100487302 | 61831286 | 0.032616439 | 16618174 | 0.04830451 | 47811710 | 0.057598462 | 7112663 |
| **18-19** | 0.109394614 | 29405414 | 0.110311034 | 11476980 | 0.030790092 | 20926487 | 0.038264566 | 6748103 | 0.117010408 | 38102603 | 0.100416216 | 61848098 | 0.032504161 | 16628166 | 0.048168582 | 47796008 | 0.057463406 | 7101682 |
| **19-20** | 0.109220215 | 29380069 | 0.110122097 | 11475184 | 0.030627794 | 20878917 | 0.038161736 | 6733172 | 0.116893145 | 38046632 | 0.100374973 | 61735557 | 0.032434659 | 16592881 | 0.048071314 | 47709553 | 0.05752697 | 7108911 |
| **20-21** | 0.10924986 | 29373104 | 0.109951402 | 11450989 | 0.030528 | 20868483 | 0.038122954 | 6733595 | 0.116803549 | 38065434 | 0.100363214 | 61653586 | 0.032389121 | 16578287 | 0.047963588 | 47686622 | 0.057499666 | 7089954 |
| **21-22** | 0.10904451 | 29338848 | 0.109817516 | 11428555 | 0.030345881 | 20834439 | 0.038136782 | 6712316 | 0.116818083 | 37959380 | 0.100310825 | 61580091 | 0.032312439 | 16548114 | 0.047829784 | 47625715 | 0.057434792 | 7091599 |
| **22-23** | 0.108984287 | 29278558 | 0.109518368 | 11419247 | 0.030240465 | 20800421 | 0.038042329 | 6704800 | 0.116747837 | 37939283 | 0.100292844 | 61544376 | 0.032231534 | 16529404 | 0.047762303 | 47566967 | 0.057350852 | 7064845 |
| **23-24** | 0.108848597 | 29264195 | 0.109307981 | 11414546 | 0.030119279 | 20778283 | 0.0379488 | 6697244 | 0.116615908 | 37917495 | 0.100240088 | 61501459 | 0.03217056 | 16514502 | 0.047684198 | 47534664 | 0.057238893 | 7069970 |
| **24-25** | 0.108754729 | 29252411 | 0.109186803 | 11412317 | 0.030003408 | 20774629 | 0.037931926 | 6688254 | 0.116625469 | 37916634 | 0.100243639 | 61444587 | 0.032142418 | 16494687 | 0.047566526 | 47494485 | 0.057236904 | 7072002 |
| **25-26** | 0.108748319 | 29245767 | 0.109045092 | 11405667 | 0.029931824 | 20765750 | 0.037883436 | 6688440 | 0.116552172 | 37895930 | 0.100201563 | 61425294 | 0.032044823 | 16508929 | 0.047483823 | 47471031 | 0.057204578 | 7064674 |
| **26-27** | 0.108558632 | 29210643 | 0.108904552 | 11405512 | 0.029822541 | 20734114 | 0.037835978 | 6693375 | 0.116551163 | 37879405 | 0.100160495 | 61396454 | 0.031998162 | 16480867 | 0.047427468 | 47451725 | 0.057115963 | 7044597 |
| **27-28** | 0.108473439 | 29219039 | 0.108757116 | 11375178 | 0.02974434 | 20720467 | 0.0378175 | 6674835 | 0.116493498 | 37821363 | 0.100128835 | 61346467 | 0.031981821 | 16463279 | 0.047348387 | 47420235 | 0.057123467 | 7040969 |
| **28-29** | 0.108480611 | 29169778 | 0.108669692 | 11367502 | 0.029666955 | 20704834 | 0.037832617 | 6683054 | 0.116476316 | 37773890 | 0.100184552 | 61278565 | 0.031955039 | 16442085 | 0.047272414 | 47345646 | 0.057015283 | 7049736 |
| **29-30** | 0.108418942 | 29175953 | 0.108549079 | 11369388 | 0.029624633 | 20719096 | 0.037760155 | 6673472 | 0.116409188 | 37782016 | 0.100094492 | 61278944 | 0.031925238 | 16445004 | 0.047221389 | 47350806 | 0.05701871 | 7043599 |
| **30-31** | 0.108353107 | 29116416 | 0.108350393 | 11359956 | 0.029488976 | 20684846 | 0.037742123 | 6667074 | 0.116411925 | 37744650 | 0.100082835 | 61225430 | 0.031887345 | 16443887 | 0.047163609 | 47303278 | 0.057036949 | 7032287 |
| **31-32** | 0.10827471 | 29111588 | 0.108355756 | 11351359 | 0.02943607 | 20660982 | 0.03770363 | 6661831 | 0.116395117 | 37721580 | 0.100066702 | 61208661 | 0.031781238 | 16431168 | 0.047043517 | 47306427 | 0.056965827 | 7030487 |
| **32-33** | 0.108214298 | 29068195 | 0.108100453 | 11330919 | 0.029392877 | 20629983 | 0.037698265 | 6647047 | 0.116375865 | 37694360 | 0.10000229 | 61115001 | 0.031777787 | 16373990 | 0.047010883 | 47230214 | 0.056922092 | 7011659 |
| **33-34** | 0.108152319 | 29058642 | 0.108080955 | 11340012 | 0.0293381 | 20626847 | 0.037738092 | 6650780 | 0.116237954 | 37646547 | 0.100009162 | 61102175 | 0.031733679 | 16375612 | 0.046956061 | 47216362 | 0.056953078 | 7026367 |
| **34-35** | 0.108095472 | 29050394 | 0.10798607 | 11324208 | 0.029299672 | 20609973 | 0.037662008 | 6636767 | 0.116236537 | 37658616 | 0.099980002 | 61037357 | 0.03173451 | 16384402 | 0.046913833 | 47174571 | 0.056891655 | 7007844 |
| **35-36** | 0.108100714 | 28984602 | 0.107760275 | 11321963 | 0.029229728 | 20591173 | 0.037611951 | 6637626 | 0.116185905 | 37606630 | 0.099941546 | 60960490 | 0.031750361 | 16362340 | 0.046874765 | 47133191 | 0.056757074 | 7001518 |
| **36-37** | 0.107975503 | 28986544 | 0.107741148 | 11323448 | 0.029169601 | 20561116 | 0.037626295 | 6625147 | 0.116119278 | 37583741 | 0.09995853 | 60900737 | 0.031679229 | 16335612 | 0.046773932 | 47127736 | 0.056780344 | 6996454 |
| **37-38** | 0.107992453 | 28996976 | 0.107696089 | 11299564 | 0.029143654 | 20571537 | 0.0376001 | 6631293 | 0.116156425 | 37580228 | 0.099958029 | 60947288 | 0.031665872 | 16342498 | 0.046741666 | 47109211 | 0.056703912 | 6994313 |
| **38-39** | 0.107909866 | 28983509 | 0.107554243 | 11302254 | 0.029076938 | 20581801 | 0.037592811 | 6628296 | 0.116153848 | 37557027 | 0.0999435 | 60966124 | 0.031671393 | 16343480 | 0.046697684 | 47114147 | 0.056722517 | 6989713 |
| **39-40** | 0.107858101 | 28974988 | 0.107375315 | 11297868 | 0.029051717 | 20568885 | 0.037531206 | 6631780 | 0.116064232 | 37559603 | 0.099932427 | 60945641 | 0.031619715 | 16334225 | 0.046633357 | 47134654 | 0.056781732 | 7002387 |
| **40-41** | 0.107966395 | 28947894 | 0.107529087 | 11301407 | 0.029044109 | 20537521 | 0.037510477 | 6624755 | 0.116061152 | 37545794 | 0.099922428 | 60884690 | 0.031585334 | 16326658 | 0.046592284 | 47060752 | 0.056671438 | 7003881 |
| **41-42** | 0.107853449 | 28930495 | 0.107243915 | 11288913 | 0.028987895 | 20551231 | 0.037532833 | 6631386 | 0.116047645 | 37611889 | 0.099894395 | 60861483 | 0.031580776 | 16325048 | 0.04658071 | 47080923 | 0.056699642 | 6988587 |
| **42-43** | 0.107752222 | 28926840 | 0.107303828 | 11289281 | 0.028928947 | 20509720 | 0.037519228 | 6620170 | 0.11601948 | 37526835 | 0.099866499 | 60821460 | 0.031584496 | 16313339 | 0.046527326 | 47042872 | 0.056648236 | 6971390 |
| **43-44** | 0.107736783 | 28946865 | 0.107164255 | 11283151 | 0.028924798 | 20523460 | 0.037466781 | 6602016 | 0.116029893 | 37534986 | 0.099904081 | 60818396 | 0.031538004 | 16307528 | 0.046488833 | 47040276 | 0.056633201 | 6985747 |
| **44-45** | 0.107757025 | 28903514 | 0.1070592 | 11282268 | 0.028884488 | 20518894 | 0.037478568 | 6621060 | 0.116012194 | 37528169 | 0.099907826 | 60771112 | 0.031547431 | 16299378 | 0.04641669 | 47039116 | 0.05658803 | 6977830 |
| **45-46** | 0.107730167 | 28921430 | 0.107047293 | 11274257 | 0.028830541 | 20504046 | 0.0374975 | 6610527 | 0.115976461 | 37493106 | 0.099833547 | 60780869 | 0.031496108 | 16310339 | 0.046384196 | 47000921 | 0.056572612 | 6981941 |
| **46-47** | 0.107602876 | 28881981 | 0.106797133 | 11263000 | 0.028849355 | 20498500 | 0.037497273 | 6613364 | 0.115950097 | 37459136 | 0.099863744 | 60771626 | 0.031477734 | 16292653 | 0.04632649 | 46967786 | 0.056486609 | 6976573 |
| **47-48** | 0.107671914 | 28866590 | 0.106951091 | 11270117 | 0.028787754 | 20477402 | 0.037450735 | 6601541 | 0.11595294 | 37451003 | 0.099848139 | 60714477 | 0.031497978 | 16295781 | 0.046312588 | 46954885 | 0.056592412 | 6972784 |
| **48-49** | 0.107570658 | 28895952 | 0.106882214 | 11261074 | 0.028759201 | 20485488 | 0.037431472 | 6598639 | 0.115906174 | 37476151 | 0.099822123 | 60678658 | 0.031457095 | 16283471 | 0.046286468 | 46967812 | 0.05651113 | 6971222 |
| **49-50** | 0.107590856 | 28896368 | 0.106844202 | 11251811 | 0.028734666 | 20485091 | 0.037424629 | 6596524 | 0.115916783 | 37465668 | 0.099796201 | 60647503 | 0.031443106 | 16271071 | 0.04625893 | 46936570 | 0.056550207 | 6965449 |
| **50-51** | 0.107502833 | 28837920 | 0.10673613 | 11255468 | 0.028710302 | 20449476 | 0.037414869 | 6595401 | 0.115853011 | 37435987 | 0.099805502 | 60621916 | 0.031445186 | 16245286 | 0.046194704 | 46912751 | 0.056493206 | 6960951 |
| **51-52** | 0.10751818 | 28877391 | 0.106654457 | 11241778 | 0.028670794 | 20455390 | 0.037377912 | 6595666 | 0.115832086 | 37389229 | 0.099763449 | 60647393 | 0.031414535 | 16267370 | 0.046144941 | 46884998 | 0.056495258 | 6967797 |
| **52-53** | 0.107468387 | 28843181 | 0.106612118 | 11234455 | 0.028662689 | 20432071 | 0.037367386 | 6587873 | 0.115834954 | 37378527 | 0.099744634 | 60604206 | 0.031398277 | 16260709 | 0.046121267 | 46879003 | 0.056484496 | 6958828 |
| **53-54** | 0.107478056 | 28824212 | 0.106553168 | 11236353 | 0.028648439 | 20419100 | 0.03744837 | 6589599 | 0.115777494 | 37377144 | 0.099766616 | 60546904 | 0.031393912 | 16244064 | 0.046107926 | 46861949 | 0.0564824 | 6951502 |
| **54-55** | 0.107532441 | 28790978 | 0.106573131 | 11224565 | 0.028609154 | 20406666 | 0.037358681 | 6573639 | 0.115763707 | 37317350 | 0.099740569 | 60533278 | 0.031365212 | 16228409 | 0.046061717 | 46803500 | 0.056518298 | 6937280 |
| **55-56** | 0.10743102 | 28769287 | 0.106500244 | 11238200 | 0.02858381 | 20412816 | 0.037401906 | 6587845 | 0.115755926 | 37370340 | 0.099757966 | 60528458 | 0.031393779 | 16239643 | 0.046045863 | 46816093 | 0.056463954 | 6951806 |

**Table S27 - LD values according to chromosome type in Illumina WGS data from dataset2 and dataset3.1.** Average r^2 values were calculated for each chromosome group in all populations available according to physical distance (kbps) between SNP pairs

| chr. type | distance (kbps) | average R^2 *(H. r. erythrogaster)* | average R^2 *(H. r. savignii)* | average R^2 *(H. r. gutturalis)* | average R^2 *(H. r. rustica)* | average R^2 *(H. r. transitiva)* | average R^2 *(H. r. tytleri)* | average R^2 *(H. r. gutturalis x tytleri)* | average R^2 *(H. r. rustica x gutturalis)* | average R^2 *(H. r. rustica x tytleri)* | average R^2 (across all populations) | standard deviation R^2 (across all populations) |
|---|---|---|---|---|---|---|---|---|---|---|---|---|
| **macro** | 0-1 | 0.138680452 | 0.142938143 | 0.074640565 | 0.068435983 | 0.135445038 | 0.116408895 | 0.064674051 | 0.074510423 | 0.086739047 | 0.10027 | 0.03274459294 |
| **macro** | 1-2 | 0.123330396 | 0.125640208 | 0.051706172 | 0.049863327 | 0.124818872 | 0.107601706 | 0.046164418 | 0.060474703 | 0.069462620 | 0.08434 | 0.03519139777 |
| **macro** | 2-3 | 0.119822384 | 0.122582372 | 0.045829232 | 0.046272879 | 0.122568030 | 0.105533681 | 0.041969367 | 0.057007275 | 0.066036207 | 0.08085 | 0.03593705389 |
| **macro** | 3-4 | 0.117914442 | 0.120539225 | 0.042308204 | 0.044435604 | 0.121353662 | 0.10444157 | 0.039642818 | 0.055105077 | 0.064134357 | 0.07887 | 0.03633650776 |
| **macro** | 4-5 | 0.116534557 | 0.119204701 | 0.040022092 | 0.04313101 | 0.120596215 | 0.103771754 | 0.038299138 | 0.053856673 | 0.062863402 | 0.07759 | 0.03658238231 |
| **macro** | 5-7 | 0.114948033 | 0.117722917 | 0.037791926 | 0.042049466 | 0.119822483 | 0.10305002 | 0.036805482 | 0.052648175 | 0.061753819 | 0.07629 | 0.03675269602 |
| **macro** | 7-9 | 0.113534616 | 0.116196947 | 0.035797205 | 0.041123021 | 0.11923819 | 0.102403836 | 0.035581273 | 0.051557211 | 0.060726678 | 0.07513 | 0.03688085625 |
| **macro** | 9-11 | 0.112572017 | 0.114906977 | 0.034394086 | 0.040499638 | 0.118813207 | 0.102046688 | 0.034821028 | 0.050771922 | 0.060110362 | 0.07433 | 0.03693478024 |
| **macro** | 11-13 | 0.111785282 | 0.114052945 | 0.033447385 | 0.040061648 | 0.118373347 | 0.101790999 | 0.034264122 | 0.050192909 | 0.059626209 | 0.07373 | 0.03694811651 |
| **macro** | 13-15 | 0.111109736 | 0.113238594 | 0.032728317 | 0.039776683 | 0.118079385 | 0.101550768 | 0.033848736 | 0.049790053 | 0.059237797 | 0.07326 | 0.03692000379 |
| **macro** | 15-20 | 0.11038349 | 0.112112072 | 0.031778291 | 0.039406908 | 0.117772564 | 0.101296631 | 0.033419605 | 0.049187799 | 0.058791518 | 0.07268 | 0.03690340901 |
| **macro** | 20-25 | 0.109612935 | 0.110923202 | 0.030850564 | 0.039027159 | 0.117356118 | 0.101053406 | 0.032940549 | 0.048576299 | 0.058433019 | 0.07209 | 0.03686091717 |
| **macro** | 25-30 | 0.109021068 | 0.110052589 | 0.030228266 | 0.03879043 | 0.11713477 | 0.100883014 | 0.032599039 | 0.048135489 | 0.058132795 | 0.07166 | 0.03682886635 |
| **macro** | 30-40 | 0.108452293 | 0.10889136 | 0.029581992 | 0.038571387 | 0.11684806 | 0.100698165 | 0.032296373 | 0.047625002 | 0.057833839 | 0.07120 | 0.03675403215 |
| **macro** | 40-50 | 0.10793539 | 0.107904921 | 0.029051489 | 0.038381982 | 0.116574262 | 0.100549767 | 0.032030304 | 0.047140715 | 0.057567825 | 0.07079 | 0.03668778955 |
| **macro** | 50-60 | 0.10758775 | 0.107121375 | 0.028690491 | 0.038263723 | 0.116358261 | 0.100397932 | 0.031865763 | 0.046743966 | 0.057422506 | 0.07049 | 0.03662082631 |
| **macro** | 60-70 | 0.107340888 | 0.106645838 | 0.028452392 | 0.03818188 | 0.1161493 | 0.100303034 | 0.031729112 | 0.046432683 | 0.057295304 | 0.07028 | 0.03657981813 |
| **macro** | 70-80 | 0.107125353 | 0.106171697 | 0.028266017 | 0.038091779 | 0.116020309 | 0.100266277 | 0.031614937 | 0.046200466 | 0.057151313 | 0.07010 | 0.03654669576 |
| **macro** | 80-90 | 0.107021421 | 0.105914787 | 0.028101033 | 0.038051945 | 0.115854807 | 0.100190706 | 0.031541206 | 0.045975074 | 0.057058391 | 0.06997 | 0.0365283332 |
| **macro** | 90-100 | 0.106916557 | 0.105643658 | 0.028010902 | 0.037999456 | 0.115792341 | 0.100111153 | 0.031473023 | 0.045777448 | 0.056969067 | 0.06985 | 0.03651150041 |
| **intermediate** | 0-1 | 0.130881978 | 0.134349507 | 0.065348018 | 0.05981625 | 0.128446072 | 0.110103797 | 0.054895384 | 0.067149335 | 0.077797509 | 0.09209 | 0.0333605762 |
| **intermediate** | 1-2 | 0.118922003 | 0.121118587 | 0.045870435 | 0.043403104 | 0.12035583 | 0.103405823 | 0.040006949 | 0.056003337 | 0.062935493 | 0.07911 | 0.035961185 |
| **intermediate** | 2-3 | 0.11635561 | 0.118855711 | 0.041273069 | 0.040990965 | 0.118918764 | 0.102013092 | 0.037086625 | 0.053629448 | 0.060517665 | 0.07663 | 0.03651427839 |
| **intermediate** | 3-4 | 0.11466082 | 0.117450278 | 0.03872637 | 0.039781373 | 0.118402364 | 0.101366449 | 0.035652061 | 0.052311453 | 0.059254016 | 0.07529 | 0.03677686411 |
| **intermediate** | 4-5 | 0.113605335 | 0.11605392 | 0.037083428 | 0.03886741 | 0.117869877 | 0.100961173 | 0.034667803 | 0.051415198 | 0.058420609 | 0.07433 | 0.03688591668 |
| **intermediate** | 5-7 | 0.112388865 | 0.11494266 | 0.035483564 | 0.038303423 | 0.117383153 | 0.100550445 | 0.033841974 | 0.050611079 | 0.057647949 | 0.07346 | 0.03694468822 |
| **intermediate** | 7-9 | 0.111483032 | 0.113680491 | 0.034175678 | 0.037704205 | 0.11698561 | 0.10019985 | 0.033182058 | 0.049836338 | 0.057029012 | 0.07270 | 0.03697939435 |
| **intermediate** | 9-11 | 0.110768816 | 0.11262876 | 0.033270816 | 0.037351636 | 0.116750795 | 0.099942115 | 0.032731203 | 0.049320907 | 0.056672277 | 0.07216 | 0.03696055452 |
| **intermediate** | 11-13 | 0.110264791 | 0.111855162 | 0.032641171 | 0.03716795 | 0.116517302 | 0.099870421 | 0.032465465 | 0.048935072 | 0.056533127 | 0.07181 | 0.03692462299 |
| **intermediate** | 13-15 | 0.109772731 | 0.110894419 | 0.032117475 | 0.037023663 | 0.116281398 | 0.099702485 | 0.032208769 | 0.048621546 | 0.056380545 | 0.07144 | 0.03683217345 |
| **intermediate** | 15-20 | 0.109211394 | 0.110095047 | 0.031503221 | 0.036813278 | 0.116050675 | 0.099524997 | 0.031959029 | 0.048177532 | 0.055868403 | 0.07102 | 0.03680102359 |
| **intermediate** | 20-25 | 0.108605529 | 0.109099119 | 0.030881071 | 0.036647023 | 0.115803229 | 0.099412912 | 0.031675304 | 0.047625823 | 0.055691892 | 0.07060 | 0.03673505212 |
| **intermediate** | 25-30 | 0.108307019 | 0.108343127 | 0.030488296 | 0.03643694 | 0.115638681 | 0.099323668 | 0.03142949 | 0.047226263 | 0.055543277 | 0.07030 | 0.03671262644 |
| **intermediate** | 30-40 | 0.107977415 | 0.107639075 | 0.030070027 | 0.036324716 | 0.115399442 | 0.099158041 | 0.031232519 | 0.046798476 | 0.055345909 | 0.06999 | 0.03666517857 |
| **intermediate** | 40-50 | 0.107704556 | 0.10687156 | 0.029814785 | 0.03616285 | 0.115230096 | 0.099097507 | 0.031096817 | 0.046368113 | 0.055113272 | 0.06972 | 0.03661659035 |
| **intermediate** | 50-60 | 0.107503126 | 0.106454833 | 0.029599992 | 0.036062166 | 0.115010462 | 0.099005108 | 0.030955211 | 0.046007775 | 0.05495053 | 0.06951 | 0.03659067914 |

| | | | | | | | | | | | | |
|---|---|---|---|---|---|---|---|---|---|---|---|---|
| **intermediate** | 60-70 | 0.10731965 | 0.106109564 | 0.02939966 | 0.036000592 | 0.114777121 | 0.098958314 | 0.030813605 | 0.045714316 | 0.05475042 | 0.06932 | 0.0365689156 |
| **intermediate** | 70-80 | 0.10723162 | 0.105873661 | 0.029270079 | 0.035968696 | 0.1146108 | 0.098924137 | 0.03075637 | 0.045489501 | 0.054698776 | 0.06920 | 0.03654813718 |
| **intermediate** | 80-90 | 0.107156446 | 0.105728797 | 0.029213058 | 0.03593053 | 0.114538384 | 0.098863589 | 0.03073721 | 0.045293516 | 0.054626311 | 0.06912 | 0.03653696381 |
| **intermediate** | 90-100 | 0.107140071 | 0.1054276 | 0.029155342 | 0.03587557 | 0.114444776 | 0.098807891 | 0.030697544 | 0.045146348 | 0.054628668 | 0.06904 | 0.03650821159 |
| **micro** | 0-1 | 0.123417656 | 0.127892618 | 0.053285237 | 0.051571119 | 0.123948597 | 0.105804261 | 0.046633248 | 0.059903985 | 0.070629785 | 0.08479 | 0.03481510737 |
| **micro** | 1-2 | 0.114206493 | 0.11718481 | 0.038154989 | 0.039898357 | 0.118279682 | 0.101253719 | 0.035768917 | 0.051938908 | 0.059526735 | 0.07513 | 0.03670648187 |
| **micro** | 2-3 | 0.112287612 | 0.11516655 | 0.034901713 | 0.038540716 | 0.117430628 | 0.100394625 | 0.03407948 | 0.050241161 | 0.057999097 | 0.07345 | 0.03698139075 |
| **micro** | 3-4 | 0.110963268 | 0.113487379 | 0.033063141 | 0.037764417 | 0.116950413 | 0.099946393 | 0.033153218 | 0.049287178 | 0.057219907 | 0.07243 | 0.03703173472 |
| **micro** | 4-5 | 0.110251065 | 0.112473194 | 0.03187929 | 0.037287723 | 0.116736399 | 0.099648901 | 0.032584818 | 0.048667704 | 0.056710793 | 0.07180 | 0.0371029182 |
| **micro** | 5-7 | 0.109382819 | 0.111047141 | 0.030826925 | 0.036984472 | 0.116326662 | 0.099385591 | 0.032116525 | 0.048033967 | 0.056182634 | 0.07114 | 0.03702520785 |
| **micro** | 7-9 | 0.108579372 | 0.109777252 | 0.029825937 | 0.036612342 | 0.11598702 | 0.099236009 | 0.031669804 | 0.047386797 | 0.055819956 | 0.07054 | 0.03699292456 |
| **micro** | 9-11 | 0.108141439 | 0.108856473 | 0.029237801 | 0.036387616 | 0.115794531 | 0.099232514 | 0.031475435 | 0.046912896 | 0.055553831 | 0.07018 | 0.03696821188 |
| **micro** | 11-13 | 0.107866663 | 0.108289479 | 0.0288125 | 0.03623801 | 0.115664948 | 0.099018512 | 0.031292368 | 0.046578916 | 0.055489396 | 0.06992 | 0.03694748845 |
| **micro** | 13-15 | 0.107574429 | 0.107738884 | 0.028449255 | 0.036043383 | 0.1155588 | 0.098839234 | 0.031147623 | 0.046266831 | 0.055280124 | 0.06966 | 0.03693111746 |
| **micro** | 15-20 | 0.107306489 | 0.107125044 | 0.028100499 | 0.035939649 | 0.115286063 | 0.098783421 | 0.030934296 | 0.0458577 | 0.055084899 | 0.06938 | 0.03690021959 |
| **micro** | 20-25 | 0.107058751 | 0.106418307 | 0.027747166 | 0.035800574 | 0.115051288 | 0.098711689 | 0.030758639 | 0.045420528 | 0.054892257 | 0.06910 | 0.03686706117 |
| **micro** | 25-30 | 0.106847291 | 0.105882628 | 0.027526184 | 0.035730676 | 0.114807744 | 0.098618269 | 0.030696311 | 0.045101121 | 0.054705304 | 0.06888 | 0.03680658731 |
| **micro** | 30-40 | 0.106605668 | 0.105450096 | 0.027323375 | 0.035638254 | 0.11459794 | 0.098539725 | 0.030537711 | 0.044689396 | 0.054593738 | 0.06866 | 0.03677870956 |
| **micro** | 40-50 | 0.106510189 | 0.105022009 | 0.027178722 | 0.035552756 | 0.114384102 | 0.098456955 | 0.03042283 | 0.044280693 | 0.054387496 | 0.06847 | 0.03675929929 |
| **micro** | 50-60 | 0.106410231 | 0.104713874 | 0.02710226 | 0.035439254 | 0.114219743 | 0.098407392 | 0.030363079 | 0.043994076 | 0.054302501 | 0.06833 | 0.03673627465 |
| **micro** | 60-70 | 0.106350647 | 0.104660303 | 0.027026602 | 0.035402676 | 0.114046843 | 0.098345266 | 0.030296342 | 0.043775941 | 0.05419456 | 0.06823 | 0.03673515543 |
| **micro** | 70-80 | 0.106358813 | 0.10445845 | 0.02696623 | 0.035345318 | 0.113836653 | 0.098343603 | 0.030257941 | 0.043576382 | 0.054110703 | 0.06814 | 0.03671880071 |
| **micro** | 80-90 | 0.106319732 | 0.104383367 | 0.026945539 | 0.035335935 | 0.113727922 | 0.098296206 | 0.030210457 | 0.043444797 | 0.054055175 | 0.06808 | 0.03670638969 |
| **micro** | 90-100 | 0.106290554 | 0.104283672 | 0.026916709 | 0.035261326 | 0.11356346 | 0.09825523 | 0.030161998 | 0.043313548 | 0.053978239 | 0.06800 | 0.03669380364 |

**Table S28 - ROIs with highest average LD values ( > 0.3) computed over 5 kbps windows using Illumina WGS data from ds3.1.**
A minimum of 100 markers per window was required to perform this analysis.

| ROI n | chr | start | end | average r^2 | marker pairs number | genes within the interval | Interval excluded for proximity to ambigous regions of the assembly | Interval excluded for not containing annotated genes |
|---|---|---|---|---|---|---|---|---|
| 1 | 1 | 4,525,000 | 4,529,999 | 0.323718501 | 104 | COL22A1 | | |
| 2 | 1 | 26,525,000 | 26,529,999 | 0.326156141 | 190 | / | | X |
| 3 | 1 | 96,335,000 | 96,339,999 | 0.485467547 | 105 | CTDP1 | | |
| 3 | 1 | 96,400,000 | 96,404,999 | 0.303938321 | 120 | CTDP1 | | |
| 3 | 1 | 96,450,000 | 96,454,999 | 0.447613366 | 120 | CTDP1 | | |
| 4 | 1 | 101,030,000 | 101,034,999 | 0.451958407 | 230 | / | | X |
| 4 | 1 | 101,075,000 | 101,079,999 | 0.306657997 | 120 | / | | X |
| 5 | 1 | 101,190,000 | 101,194,999 | 0.306817598 | 152 | AMPH | | |
| 6 | 1 | 101,600,000 | 101,604,999 | 0.303188657 | 136 | POU6F2 | | |
| 7 | 1 | 103,225,000 | 103,229,999 | 0.426162424 | 105 | MRPL32 | | |
| 8 | 1 | 104,275,000 | 104,279,999 | 0.304933051 | 105 | TPK1 | | |
| 9 | 1 | 134,855,000 | 134,859,999 | 0.374239894 | 105 | ABCB1 | | |
| 9 | 1 | 134,860,000 | 134,864,999 | 0.327953806 | 120 | ABCB1 | | |
| 10 | 2 | 7,745,000 | 7,749,999 | 0.331257264 | 171 | / | | X |
| 11 | 3 | 45,000 | 49,999 | 0.548404774 | 136 | ACSS1 | | |
| 11 | 3 | 80,000 | 84,999 | 0.331551673 | 231 | TTBK1 | | |
| 11 | 3 | 85,000 | 89,999 | 0.376619967 | 120 | TTBK1 | | |
| 11 | 3 | 90,000 | 94,999 | 0.410552098 | 120 | TTBK1 | | |
| 11 | 3 | 95,000 | 99,999 | 0.309422723 | 231 | TTBK1 | | |
| 11 | 3 | 105,000 | 109,999 | 0.417221146 | 120 | TTBK1 | | |
| 11 | 3 | 115,000 | 119,999 | 0.352963042 | 120 | TTBK1 | | |
| 11 | 3 | 160,000 | 164,999 | 0.329566118 | 105 | SLC22A7 | | |
| 11 | 3 | 195,000 | 199,999 | 0.391623229 | 190 | POLR1B | | |
| 12 | 3 | 330,000 | 334,999 | 0.370706919 | 153 | PPP2R5D | | |
| 12 | 3 | 355,000 | 359,999 | 0.363481778 | 253 | MEA1, KLHDC3 | | |
| 13 | 3 | 12,510,000 | 12,514,999 | 0.368875797 | 105 | LOC120750516 | | |
| 14 | 3 | 16,445,000 | 16,449,999 | 0.501605677 | 325 | PCARE | | |
| 14 | 3 | 16,480,000 | 16,484,999 | 0.598724083 | 171 | CLIP4 | | |
| 14 | 3 | 16,485,000 | 16,489,999 | 0.462030338 | 171 | CLIP4 | | |

| | | | | | | | | |
|---|---|---|---|---|---|---|---|---|
| **15** | 3 | 16,650,000 | 16,654,999 | 0.47011107 | 134 | ALK | | |
| **15** | 3 | 16,715,000 | 16,719,999 | 0.538193851 | 276 | ALK | | |
| **15** | 3 | 16,730,000 | 16,734,999 | 0.352739412 | 136 | ALK | | |
| **16** | 3 | 16,850,000 | 16,854,999 | 0.553628128 | 171 | LOC120750002 | | |
| **16** | 3 | 16,880,000 | 16,884,999 | 0.387476236 | 105 | / | | X |
| **16** | 3 | 16,890,000 | 16,894,999 | 0.460067064 | 171 | / | | X |
| **16** | 3 | 16,895,000 | 16,899,999 | 0.330815482 | 105 | / | | X |
| **16** | 3 | 16,905,000 | 16,909,999 | 0.310238401 | 210 | / | | X |
| **16** | 3 | 16,910,000 | 16,914,999 | 0.350068784 | 105 | / | | X |
| **16** | 3 | 16,920,000 | 16,924,999 | 0.598184708 | 105 | YPEL5 | | |
| **17** | 3 | 17,050,000 | 17,054,999 | 0.315353521 | 153 | / | | X |
| **17** | 3 | 17,055,000 | 17,059,999 | 0.349880946 | 171 | / | | X |
| **17** | 3 | 17,060,000 | 17,064,999 | 0.512528177 | 153 | / | | X |
| **17** | 3 | 17,105,000 | 17,109,999 | 0.450686702 | 120 | LCLAT1 | | |
| **17** | 3 | 17,145,000 | 17,149,999 | 0.303347154 | 136 | LCLAT1 | | |
| **18** | 3 | 98,945,000 | 98,949,999 | 0.361068511 | 105 | RPS7 | | |
| **19** | 3 | 111,460,000 | 111,464,999 | 0.325996763 | 276 | PLA2G7 | | |
| **20** | 4 | 40,145,000 | 40,149,999 | 0.392746 | 103 | PPP1R12A | | |
| **21** | 4 | 45,985,000 | 45,989,999 | 0.327447748 | 153 | PLEKHG7 | | |
| **22** | 5 | 21,565,000 | 21,569,999 | 0.434107134 | 105 | / | | X |
| **23** | 5 | 36,835,000 | 36,839,999 | 0.313502134 | 105 | LOC120753169 | | |
| **24** | 5 | 68,095,000 | 68,099,999 | 0.302952662 | 136 | / | | X |
| **25** | 6 | 27,755,000 | 27,759,999 | 0.334276961 | 120 | MGA | | |
| **25** | 6 | 27,810,000 | 27,814,999 | 0.447479505 | 105 | MAPKBP1 | | |
| **25** | 6 | 27,845,000 | 27,849,999 | 0.452836549 | 120 | MAPKBP1 | | |
| **25** | 6 | 27,920,000 | 27,924,999 | 0.385319023 | 351 | LOC120753633 | | |
| **25** | 6 | 27,930,000 | 27,934,999 | 0.460366858 | 369 | LOC120753633 | | |
| **25** | 6 | 27,965,000 | 27,969,999 | 0.42517482 | 210 | LOC120753633 | | |
| **26** | 6 | 28,495,000 | 28,499,999 | 0.301443445 | 120 | ZNF106 | | |
| **27** | 6 | 28,825,000 | 28,829,999 | 0.353217755 | 153 | UBR1 | | |
| **28** | 6 | 29,215,000 | 29,219,999 | 0.332188077 | 105 | RBM25 | | |
| **29** | 6 | 29,825,000 | 29,829,999 | 0.312092979 | 210 | / | | X |
| **29** | 6 | 29,835,000 | 29,839,999 | 0.365393453 | 595 | / | | X |
| **29** | 6 | 29,940,000 | 29,944,999 | 0.503766325 | 104 | SIPA1L1 | | |

| | | | | | | | | |
|---|---|---|---|---|---|---|---|---|
| **30** | 6 | 30,255,000 | 30,259,999 | 0.328386684 | 171 | MAP3K9 | | |
| **30** | 6 | 30,260,000 | 30,264,999 | 0.352607586 | 276 | MAP3K9 | | |
| **31** | 6 | 31,050,000 | 31,054,999 | 0.330805973 | 104 | SLC39A9 | | |
| **32** | 6 | 31,315,000 | 31,319,999 | 0.306403516 | 231 | ACTN1 | | |
| **33** | 6 | 32,305,000 | 32,309,999 | 0.328642463 | 300 | GPHN | | |
| **34** | 6 | 32,430,000 | 32,434,999 | 0.300470827 | 119 | GPHN | | |
| **34** | 6 | 32,455,000 | 32,459,999 | 0.362579067 | 105 | GPHN | | |
| **35** | 6 | 36,325,000 | 36,329,999 | 0.329783717 | 119 | / | | X |
| **36** | 6 | 36,960,000 | 36,964,999 | 0.41218069 | 105 | / | | X |
| **37** | 6 | 37,280,000 | 37,284,999 | 0.469569797 | 253 | NOVA1 | | |
| **38** | 6 | 37,805,000 | 37,809,999 | 0.307179829 | 325 | STXBP6 | | |
| **38** | 6 | 37,810,000 | 37,814,999 | 0.578396897 | 153 | STXBP6 | | |
| **38** | 6 | 37,845,000 | 37,849,999 | 0.355639329 | 119 | STXBP6 | | |
| **38** | 6 | 37,855,000 | 37,859,999 | 0.354314546 | 171 | STXBP6 | | |
| **39** | 6 | 46,515,000 | 46,519,999 | 0.579355153 | 105 | AP2A2 | | |
| **39** | 6 | 46,535,000 | 46,539,999 | 0.503726853 | 135 | AP2A2 | | |
| **40** | 6 | 50,465,000 | 50,469,999 | 0.346252081 | 105 | PLEKHA7 | | |
| **41** | 6 | 50,760,000 | 50,764,999 | 0.339192987 | 120 | SOX6 | | |
| **42** | 6 | 51,280,000 | 51,284,999 | 0.386203126 | 171 | / | | X |
| **43** | 6 | 52,020,000 | 52,024,999 | 0.323671792 | 406 | LMO1 | | |
| **44** | 6 | 52,165,000 | 52,169,999 | 0.312987042 | 153 | LOC120754649 | | |
| **44** | 6 | 52,260,000 | 52,264,999 | 0.314201757 | 105 | LOC120754649 | | |
| **44** | 6 | 52,270,000 | 52,274,999 | 0.372920521 | 120 | LOC120754649 | | |
| **44** | 6 | 52,275,000 | 52,279,999 | 0.372192095 | 135 | LOC120754649 | | |
| **44** | 6 | 52,315,000 | 52,319,999 | 0.337098639 | 105 | / | | X |
| **44** | 6 | 52,400,000 | 52,404,999 | 0.315211782 | 741 | TRIM66 | | |
| **44** | 6 | 52,440,000 | 52,444,999 | 0.414994298 | 210 | DENND2B | | |
| **44** | 6 | 52,455,000 | 52,459,999 | 0.312397507 | 120 | DENND2B | | |
| **44** | 6 | 52,495,000 | 52,499,999 | 0.464522242 | 120 | DENND2B | | |
| **45** | 6 | 53,680,000 | 53,684,999 | 0.771114866 | 136 | / | | X |
| **45** | 6 | 53,695,000 | 53,699,999 | 0.587150949 | 153 | / | | X |
| **45** | 6 | 53,705,000 | 53,709,999 | 0.705205103 | 105 | / | | X |
| **45** | 6 | 53,725,000 | 53,729,999 | 0.872786758 | 120 | CCDC34 | | |
| **45** | 6 | 53,730,000 | 53,734,999 | 0.705027024 | 104 | CCDC34 | | |

| | | | | | | | | |
|---|---|---|---|---|---|---|---|---|
| **45** | 6 | 53,735,000 | 53,739,999 | 0.740417802 | 105 | CCDC34 | | |
| **45** | 6 | 53,755,000 | 53,759,999 | 0.552005477 | 120 | LGR4 | | |
| **45** | 6 | 53,765,000 | 53,769,999 | 0.668553403 | 136 | LGR4 | | |
| **45** | 6 | 53,775,000 | 53,779,999 | 0.666817446 | 153 | LGR4 | | |
| **45** | 6 | 53,780,000 | 53,784,999 | 0.604254879 | 300 | LGR4 | | |
| **45** | 6 | 53,790,000 | 53,794,999 | 0.608621875 | 105 | LGR4 | | |
| **45** | 6 | 53,810,000 | 53,814,999 | 0.547712705 | 105 | LGR4 | | |
| **45** | 6 | 53,815,000 | 53,819,999 | 0.625098194 | 252 | LGR4 | | |
| **45** | 6 | 53,845,000 | 53,849,999 | 0.507142341 | 120 | LIN7C | | |
| **45** | 6 | 53,860,000 | 53,864,999 | 0.30950362 | 190 | / | | X |
| **45** | 6 | 53,870,000 | 53,874,999 | 0.349581369 | 153 | / | | X |
| **45** | 6 | 53,875,000 | 53,879,999 | 0.377250405 | 378 | / | | X |
| **45** | 6 | 53,885,000 | 53,889,999 | 0.432021741 | 190 | BDNF | | |
| **45** | 6 | 53,895,000 | 53,899,999 | 0.496976461 | 120 | BDNF | | |
| **45** | 6 | 53,900,000 | 53,904,999 | 0.329262907 | 120 | BDNF | | |
| **45** | 6 | 53,910,000 | 53,914,999 | 0.34897478 | 120 | BDNF | | |
| **45** | 6 | 53,945,000 | 53,949,999 | 0.37879887 | 324 | / | | X |
| **45** | 6 | 53,950,000 | 53,954,999 | 0.419861929 | 153 | / | | X |
| **46** | 8 | 525,000 | 529,999 | 0.756105245 | 120 | LOC120755920 | | |
| **47** | 8 | 2,910,000 | 2,914,999 | 0.323768569 | 136 | MCU | | |
| **48** | 8 | 3,210,000 | 3,214,999 | 0.327505282 | 136 | LOC120756098 | | |
| **49** | 10 | 1,300,000 | 1,304,999 | 0.312055501 | 153 | VAMP2 | | |
| **50** | 11 | 16,830,000 | 16,834,999 | 0.430271256 | 276 | / | X | |
| **50** | 11 | 16,845,000 | 16,849,999 | 0.460269469 | 210 | / | | X |
| **50** | 11 | 16,855,000 | 16,859,999 | 0.428974743 | 231 | / | | X |
| **51** | 11 | 18,220,000 | 18,224,999 | 0.546459528 | 171 | / | | X |
| **51** | 11 | 18,235,000 | 18,239,999 | 0.551595553 | 120 | / | | X |
| **51** | 11 | 18,260,000 | 18,264,999 | 0.773465513 | 136 | / | | X |
| **51** | 11 | 18,270,000 | 18,274,999 | 0.9384316 | 190 | / | | X |
| **51** | 11 | 18,275,000 | 18,279,999 | 0.694954171 | 105 | / | | X |
| **51** | 11 | 18,295,000 | 18,299,999 | 0.318043332 | 153 | / | | X |
| **51** | 11 | 18,365,000 | 18,369,999 | 0.525914905 | 120 | / | | X |
| **51** | 11 | 18,410,000 | 18,414,999 | 0.336137998 | 561 | / | | X |
| **51** | 11 | 18,515,000 | 18,519,999 | 0.463150477 | 105 | / | | X |

| | | | | | | | | |
|---|---|---|---|---|---|---|---|---|
| **51** | 11 | 18,595,000 | 18,599,999 | 0.32001957 | 120 | / | | X |
| **52** | 11 | 19,455,000 | 19,459,999 | 0.331963055 | 595 | BEAN1 | | |
| **52** | 11 | 19,500,000 | 19,504,999 | 0.300486863 | 435 | LOC120757948 | | |
| **52** | 11 | 19,510,000 | 19,514,999 | 0.333400554 | 190 | TK2 | | |
| **52** | 11 | 19,535,000 | 19,539,999 | 0.382415971 | 136 | LOC120757604,CMTM3 | X | |
| **52** | 11 | 19,550,000 | 19,554,999 | 0.393890105 | 171 | / | | X |
| **52** | 11 | 19,560,000 | 19,564,999 | 0.390796904 | 300 | CMTM4 | | |
| **52** | 11 | 19,565,000 | 19,569,999 | 0.381078004 | 190 | CMTM4 | | |
| **52** | 11 | 19,575,000 | 19,579,999 | 0.346412034 | 300 | CMTM4 | | |
| **52** | 11 | 19,580,000 | 19,584,999 | 0.495238444 | 351 | CMTM4 | | |
| **52** | 11 | 19,585,000 | 19,589,999 | 0.509732531 | 151 | CMTM4 | | |
| **52** | 11 | 19,590,000 | 19,594,999 | 0.463295287 | 187 | / | | X |
| **52** | 11 | 19,595,000 | 19,599,999 | 0.504887215 | 120 | DYNC1LI2 | | |
| **52** | 11 | 19,605,000 | 19,609,999 | 0.713151153 | 171 | DYNC1LI2 | | |
| **52** | 11 | 19,610,000 | 19,614,999 | 0.705062508 | 190 | DYNC1LI2 | | |
| **52** | 11 | 19,615,000 | 19,619,999 | 0.560123309 | 190 | DYNC1LI2 | | |
| **53** | 11 | 19,800,000 | 19,804,999 | 0.947497399 | 153 | LOC120757950 | X | |
| **53** | 11 | 19,815,000 | 19,819,999 | 0.696842914 | 168 | LOC120757727 | | |
| **53** | 11 | 19,915,000 | 19,919,999 | 0.831808 | 253 | LOC120757950 | | |
| **53** | 11 | 19,925,000 | 19,929,999 | 0.753129214 | 231 | LRP3 | | |
| **53** | 11 | 19,930,000 | 19,934,999 | 0.611347163 | 190 | LRP3 | | |
| **53** | 11 | 19,940,000 | 19,944,999 | 0.403813532 | 595 | LRP3 | | |
| **53** | 11 | 19,945,000 | 19,949,999 | 0.596263267 | 253 | LRP3 | | |
| **53** | 11 | 19,965,000 | 19,969,999 | 0.696903778 | 171 | SLC7A10 | | |
| **53** | 11 | 19,970,000 | 19,974,999 | 0.531826285 | 136 | SLC7A10 | | |
| **53** | 11 | 19,980,000 | 19,984,999 | 0.317947238 | 351 | SLC7A10 | | |
| **53** | 11 | 19,990,000 | 19,994,999 | 0.336530293 | 231 | SLC7A10 | | |
| **53** | 11 | 19,995,000 | 19,999,999 | 0.308402149 | 698 | SLC7A10 | | |
| **53** | 11 | 20,000,000 | 20,004,999 | 0.477269596 | 561 | SLC7A10 | | |
| **53** | 11 | 20,005,000 | 20,009,999 | 0.308389392 | 136 | SLC7A10 | | |
| **53** | 11 | 20,020,000 | 20,024,999 | 0.385645733 | 210 | / | | X |
| **53** | 11 | 20,025,000 | 20,029,999 | 0.321230946 | 231 | / | | X |
| **53** | 11 | 20,035,000 | 20,039,999 | 0.413027229 | 171 | / | | X |
| **53** | 11 | 20,040,000 | 20,044,999 | 0.376046977 | 190 | / | | X |

| | | | | | | | | |
|---|---|---|---|---|---|---|---|---|
| **53** | 11 | 20,045,000 | 20,049,999 | 0.569983013 | 136 | / | | X |
| **53** | 11 | 20,050,000 | 20,054,999 | 0.443500276 | 190 | / | | X |
| **53** | 11 | 20,055,000 | 20,059,999 | 0.387810415 | 136 | / | | X |
| **54** | 11 | 21,200,000 | 21,204,999 | 0.365741393 | 435 | UBA2 | | |
| **55** | 11 | 21,385,000 | 21,389,999 | 0.496061776 | 153 | PDP2 | | |
| **55** | 11 | 21,390,000 | 21,394,999 | 0.312506656 | 231 | CA7 | | |
| **55** | 11 | 21,400,000 | 21,404,999 | 0.59924966 | 120 | CA7 | X | |
| **55** | 11 | 21,405,000 | 21,409,999 | 0.827014714 | 630 | NAE1 | X | |
| **56** | 12 | 9,430,000 | 9,434,999 | 0.312684787 | 405 | EEFSEC | | |
| **57** | 16 | 14,720,000 | 14,724,999 | 0.344505056 | 105 | NDRG3 | | |
| **57** | 16 | 14,740,000 | 14,744,999 | 0.37214136 | 253 | NDRG3 | | |
| **57** | 16 | 14,755,000 | 14,759,999 | 0.320529563 | 136 | NDRG3 | | |
| **57** | 16 | 14,770,000 | 14,774,999 | 0.354718301 | 136 | NDRG3 | | |
| **58** | 17 | 160,000 | 164,999 | 0.369956025 | 105 | / | | X |
| **59** | 18 | 6,860,000 | 6,864,999 | 0.340840058 | 136 | / | | X |
| **60** | 27 | 20,000 | 24,999 | 0.429871942 | 105 | / | | X |
| **61** | W | 5,120,000 | 5,124,999 | 0.689897033 | 120 | MBNL3 | | |
| **61** | W | 5,125,000 | 5,129,999 | 0.643972838 | 120 | MBNL3 | | |
| **61** | W | 5,130,000 | 5,134,999 | 0.612508051 | 136 | MBNL3 | | |
| **62** | W | 9,325,000 | 9,329,999 | 0.46989773 | 105 | / | | X |
| **63** | W | 9,570,000 | 9,574,999 | 0.697001733 | 105 | RBMX | | |
| **63** | W | 9,605,000 | 9,609,999 | 0.666389717 | 120 | LOC120764854 | | |
| **64** | W | 12,075,000 | 12,079,999 | 0.528482703 | 171 | ZC3H12B | | |
| **65** | W | 12,290,000 | 12,294,999 | 0.558123099 | 171 | LOC120764947 | | |
| **66** | W | 19,310,000 | 19,314,999 | 0.440821923 | 136 | POF1B | | |
| **67** | W | 19,460,000 | 19,464,999 | 0.60091764 | 136 | LOC120764804 | | |
| **68** | W | 20,630,000 | 20,634,999 | 0.467206752 | 136 | / | | X |
| **69** | W | 23,465,000 | 23,469,999 | 0.462534376 | 276 | ZBTB33 | | |
| **69** | W | 23,520,000 | 23,524,999 | 0.580058622 | 231 | UPF3B,RPL39,LOC120765016 | | |
| **69** | W | 23,615,000 | 23,619,999 | 0.533652241 | 153 | SEPTIN6 | | |
| **69** | W | 23,655,000 | 23,659,999 | 0.484432125 | 496 | NKRF | | |
| **70** | W | 28,315,000 | 28,319,999 | 0.837218083 | 120 | ATP11C | | |
| **71** | W | 31,595,000 | 31,599,999 | 0.579872077 | 210 | LOC120764863 | | |
| **72** | Z | 58,080,000 | 58,084,999 | 0.433165801 | 120 | / | | X |

| | | | | | | | | |
|---|---|---|---|---|---|---|---|---|
| **72** | Z | 58,115,000 | 58,119,999 | 0.322064155 | 351 | / | | X |
| **72** | Z | 58,120,000 | 58,124,999 | 0.453374826 | 276 | / | | X |
| **72** | Z | 58,125,000 | 58,129,999 | 0.399757528 | 351 | / | | X |
| **72** | Z | 58,145,000 | 58,149,999 | 0.304713724 | 465 | / | | X |
| **72** | Z | 58,175,000 | 58,179,999 | 0.419082732 | 153 | / | | X |
| **73** | Z | 59,570,000 | 59,574,999 | 0.368467485 | 276 | / | | X |
| **73** | Z | 59,575,000 | 59,579,999 | 0.312210827 | 120 | YTHDC2 | | |
| **74** | Z | 77,225,000 | 77,229,999 | 0.318757035 | 171 | / | | X |
| **75** | Z | 82,775,000 | 82,779,999 | 0.304575576 | 325 | CUL4B | | |
| **76** | Z | 86,270,000 | 86,274,999 | 0.433500256 | 120 | LOC120765675 | | |
| **77** | Z | 86,550,000 | 86,554,999 | 0.370511446 | 104 | DACH2 | | |
| **78** | Z | 89,970,000 | 89,974,999 | 0.569778107 | 104 | / | | X |
| **78** | Z | 89,975,000 | 89,979,999 | 0.63959112 | 300 | / | | X |
| **78** | Z | 90,010,000 | 90,014,999 | 0.698744137 | 153 | / | | X |

**Table S29 - Genotypes of *H. r. savignii* samples (ds3.1.1) within the *BDNF* gene region.** The genotypes of all *H. r. savignii* samples from ds3.1 within *BDNF* region are reported. 0/0: homozygous reference allele; 0/1: heterozygous; 1/1 homozygous alternate allele. In the last column, X indicates whether the SNP affects a CpG site.

| chr | position | reference allele | alternate allele | genotype SRR7167866 | genotype SRR7167867 | genotype SRR7167868 | genotype SRR7167869 | genotype SRR7167871 | genotype SRR7167872 | genotype SRR7167873 | genotype SRR7167877 | % of homozygous genotypes | part of / altering a CpG site |
|---|---|---|---|---|---|---|---|---|---|---|---|---|---|
| chr6 | 53887847 | G | A | 0/0 | 0/0 | 0/0 | 0/0 | 0/0 | 0/0 | 0/0 | 0/0 | 100 | X |
| chr6 | 53887979 | A | G | 0/1 | ./. | 0/0 | 0/0 | 0/0 | 0/0 | 0/0 | 0/0 | 85.7 | X |
| chr6 | 53888330 | T | A | 0/0 | 0/0 | 0/0 | 0/0 | 0/0 | 0/0 | 0/0 | 0/0 | 100 | |
| chr6 | 53888975 | C | T | 0/1 | 0/0 | 0/0 | 0/0 | 0/0 | 0/0 | 0/0 | 0/0 | 87.5 | X |
| chr6 | 53889046 | G | A | 0/0 | 0/0 | 0/0 | 0/0 | 0/0 | 0/0 | 0/0 | 0/0 | 100 | |
| chr6 | 53889327 | C | T | 0/0 | 0/0 | 0/0 | 0/0 | 0/0 | 0/0 | 0/0 | 0/0 | 100 | X |
| chr6 | 53889342 | G | A | 1/1 | 0/0 | 0/0 | 0/0 | 0/0 | 0/0 | 0/0 | 0/0 | 87.5 | |
| chr6 | 53889381 | T | C | 0/0 | 0/0 | ./. | 0/0 | 0/0 | 0/0 | 0/0 | 0/0 | 100 | X |
| chr6 | 53889520 | A | C | 0/0 | 0/0 | 0/0 | 0/0 | 0/0 | 0/0 | 0/0 | 0/0 | 100 | |
| chr6 | 53890246 | T | C | 1/1 | 1/1 | 0/1 | ./. | 0/1 | 0/0 | 1/1 | 0/0 | 28.6 | |
| chr6 | 53890512 | T | A | 0/0 | 0/0 | 0/0 | 0/0 | 0/0 | 0/0 | 0/0 | 0/0 | 100 | |
| chr6 | 53890692 | T | C | 0/0 | 0/0 | 0/0 | 0/0 | 0/0 | 0/1 | 0/0 | 0/1 | 75.0 | |
| chr6 | 53890815 | G | A | 0/0 | 0/0 | 0/0 | 0/0 | 0/0 | 0/0 | 0/0 | 0/0 | 100 | |
| chr6 | 53890943 | C | T | 0/1 | 0/0 | 0/0 | 0/0 | 0/0 | 0/0 | 0/0 | ./. | 85.7 | |
| chr6 | 53890965 | A | G | 0/1 | 0/0 | 0/0 | 0/0 | 0/0 | 0/0 | 0/0 | 0/0 | 87.5 | X |
| chr6 | 53891012 | A | C | 0/0 | 0/0 | 0/0 | 0/0 | 0/0 | 0/0 | 1/1 | 0/0 | 87.5 | |
| chr6 | 53891107 | A | G | ./. | 0/0 | 0/0 | 0/0 | 0/0 | 0/1 | 0/0 | 0/1 | 71.4 | |
| chr6 | 53891199 | G | T | ./. | 0/0 | 0/0 | 0/0 | 0/0 | 0/0 | 0/0 | 0/0 | 100 | X |
| chr6 | 53891451 | G | A | 0/1 | 1/1 | 0/1 | 1/1 | 0/0 | 0/1 | 1/1 | 0/1 | 12.5 | X |
| chr6 | 53891507 | C | T | 0/0 | 0/0 | 0/0 | ./. | 0/0 | 0/0 | 0/0 | 0/0 | 100 | X |
| chr6 | 53893216 | C | A | 0/1 | 0/0 | 0/0 | 0/0 | 0/0 | 0/0 | 0/0 | 0/0 | 87.5 | |
| chr6 | 53895223 | G | T | 0/0 | 0/0 | 0/0 | 0/0 | 0/0 | 0/0 | 0/0 | 0/0 | 100 | |
| chr6 | 53895351 | C | T | 1/1 | 0/0 | 0/0 | 0/0 | 0/0 | 0/0 | 0/0 | 0/0 | 87.5 | X |
| chr6 | 53896510 | C | A | 0/0 | 0/0 | 0/0 | 0/0 | 0/0 | 0/0 | 0/0 | 0/0 | 100 | |
| chr6 | 53897090 | G | A | 0/1 | 0/0 | ./. | 0/0 | 0/0 | 0/0 | 0/0 | 0/0 | 85.7 | X |
| chr6 | 53897167 | C | T | 0/1 | 0/0 | 0/0 | 0/0 | 0/0 | 0/0 | 0/0 | 0/0 | 87.5 | X |
| chr6 | 53897199 | T | A | 0/1 | 0/0 | 0/0 | 0/0 | 0/0 | 0/0 | 0/0 | 0/0 | 87.5 | |
| chr6 | 53897968 | C | T | 0/0 | 0/0 | 0/0 | 0/0 | 0/0 | 0/0 | 0/0 | 0/0 | 100 | |
| chr6 | 53898080 | C | T | 1/1 | 0/0 | 0/0 | 0/0 | 0/0 | 0/0 | 0/0 | 0/0 | 87.5 | X |
| chr6 | 53898218 | T | C | 0/0 | 0/0 | 0/0 | 0/0 | 0/0 | 0/0 | 0/0 | 0/0 | 100 | |
| chr6 | 53898236 | G | A | 0/0 | 0/0 | 0/0 | 0/0 | 0/0 | 0/0 | 0/0 | 0/0 | 100 | |
| chr6 | 53898362 | G | A | 0/0 | 0/0 | 0/0 | 0/0 | 0/0 | 0/0 | 0/0 | 0/0 | 100 | |
| chr6 | 53898429 | G | T | 0/0 | 0/0 | 0/0 | 0/0 | 0/0 | 0/0 | 0/0 | 0/0 | 100 | |
| chr6 | 53898529 | G | T | 0/0 | 0/0 | 0/0 | 0/0 | 0/0 | 0/0 | 0/0 | 0/0 | 100 | |
| chr6 | 53898983 | G | A | 0/0 | 0/0 | 0/0 | 0/0 | 0/0 | 0/0 | 0/0 | 0/0 | 100 | |
| chr6 | 53899620 | A | G | 0/1 | 0/0 | 0/0 | 0/0 | 0/0 | 0/0 | 0/0 | ./. | 85.7 | |
| chr6 | 53899935 | T | C | 0/0 | 0/0 | 0/0 | 0/0 | 0/0 | 0/0 | 0/0 | . | 100 | |
| chr6 | 53900031 | T | C | 0/0 | 0/0 | 0/0 | 0/0 | 0/0 | 0/0 | 0/0 | . | 100 | |
| chr6 | 53900534 | C | T | 0/0 | 0/0 | 0/0 | 0/0 | 0/0 | 0/0 | 0/0 | 0/0 | 100 | |

| | | | | | | | | | | | | | |
|---|---|---|---|---|---|---|---|---|---|---|---|---|---|
| **chr6** | 53901454 | C | T | 0/1 | 0/0 | 0/0 | 0/0 | 0/0 | 0/0 | 0/0 | 0/0 | 87.5 | X |
| **chr6** | 53901462 | T | A | 0/0 | 0/0 | 0/0 | 0/0 | 0/0 | 0/0 | 0/0 | 0/0 | 100 | |
| **chr6** | 53901536 | T | C | 0/1 | 0/0 | 0/0 | 0/0 | 0/0 | 0/0 | 0/0 | 0/0 | 87.5 | |
| **chr6** | 53901605 | C | A | 0/1 | 0/0 | 0/0 | 0/0 | 0/0 | 0/0 | 0/0 | 0/0 | 87.5 | |
| **chr6** | 53902087 | G | T | 0/0 | 0/0 | 0/0 | 0/0 | 0/0 | 0/0 | 0/0 | 0/0 | 100 | |
| **chr6** | 53902174 | G | T | 0/0 | 0/0 | 0/0 | 0/0 | 0/0 | 0/0 | 0/0 | 0/0 | 100 | |
| **chr6** | 53902193 | A | T | 0/1 | 0/0 | 0/0 | 0/0 | 0/0 | 0/0 | 0/0 | 0/0 | 87.5 | |
| **chr6** | 53902321 | G | A | 0/0 | 0/0 | ./. | 0/0 | 0/0 | 0/1 | 0/0 | 0/1 | 71.4 | X |
| **chr6** | 53902829 | T | G | 0/0 | 0/0 | 0/0 | 0/0 | 0/0 | 0/0 | 0/0 | 0/0 | 100 | |
| **chr6** | 53902896 | C | G | 0/0 | 0/0 | 0/0 | 0/0 | 0/0 | 0/0 | 0/0 | 0/0 | 100 | |
| **chr6** | 53903471 | C | T | 1/1 | 0/0 | 0/1 | ./. | 0/0 | 0/0 | 0/0 | 0/0 | 71.4 | |
| **chr6** | 53903663 | A | G | . | 0/0 | 0/0 | 0/0 | 0/0 | 0/0 | 0/0 | 0/0 | 100 | |
| **chr6** | 53904017 | A | C | 0/0 | 0/0 | 0/0 | 0/0 | 0/0 | 0/1 | 0/0 | 0/1 | 75.0 | |
| **chr6** | 53904926 | T | G | 0/1 | 0/0 | 0/0 | 0/0 | 0/0 | ./. | 0/0 | 0/0 | 85.7 | X |
| **chr6** | 53905398 | C | T | 0/0 | 0/0 | 0/0 | 0/0 | 0/0 | 0/0 | 0/0 | 0/0 | 100 | |
| **chr6** | 53905494 | T | C | 0/0 | 0/0 | ./. | 0/0 | 0/0 | 0/1 | 0/0 | 0/1 | 71.4 | |
| **chr6** | 53905674 | G | A | 0/0 | 1/1 | 0/0 | 0/0 | 0/0 | 0/0 | 0/0 | 0/0 | 87.5 | |
| **chr6** | 53905675 | G | C | 0/1 | 0/0 | 0/0 | 0/0 | 0/0 | 0/0 | 0/0 | 0/0 | 87.5 | |
| **chr6** | 53905922 | T | C | 0/0 | 0/0 | 0/0 | 0/0 | 0/0 | 0/1 | 0/0 | 0/1 | 87.5 | X |
| **chr6** | 53905950 | A | G | 0/1 | 0/0 | 0/0 | 0/0 | 0/0 | 0/0 | 0/0 | 0/0 | 87.5 | |
| **chr6** | 53908016 | C | G | 0/0 | ./. | 0/0 | 0/0 | 0/0 | 0/0 | 0/0 | 0/0 | 100 | X |
| **chr6** | 53908027 | A | G | 0/1 | 0/0 | 0/0 | 0/0 | 0/0 | 0/0 | 0/0 | 0/0 | 87.5 | |
| **chr6** | 53908036 | C | G | 0/1 | 0/0 | 0/0 | 0/0 | 0/0 | 0/0 | 0/0 | 0/0 | 87.5 | X |
| **chr6** | 53908225 | T | G | 0/0 | 0/0 | 0/0 | 0/0 | 0/0 | 0/0 | 0/0 | 0/0 | 100 | X |
| **chr6** | 53908236 | C | T | 0/1 | 0/0 | 0/0 | 0/0 | 0/0 | 0/0 | 0/0 | 0/0 | 87.5 | X |
| **chr6** | 53908257 | T | C | 0/0 | 0/0 | 0/0 | 0/0 | 0/0 | 0/0 | 0/0 | 0/0 | 100 | X |
| **chr6** | 53908287 | C | T | 0/0 | 0/0 | 0/0 | 0/1 | 0/0 | 0/0 | 0/0 | 0/1 | 75.0 | |
| **chr6** | 53908403 | A | G | 0/0 | 0/0 | 0/0 | 0/1 | 0/0 | 0/0 | 0/0 | 0/1 | 75.0 | |
| **chr6** | 53908956 | C | T | 0/0 | 0/0 | 0/0 | 0/0 | 0/0 | 0/0 | 0/0 | 0/0 | 100 | |
| **chr6** | 53909489 | T | C | 0/1 | 0/0 | 0/0 | 0/0 | 0/0 | 0/0 | ./. | 0/0 | 85.7 | |
| **chr6** | 53909534 | C | T | 0/0 | ./. | 0/0 | 0/0 | 0/0 | 0/0 | 0/0 | 0/0 | 100 | |
| **chr6** | 53910061 | G | A | 0/0 | 0/0 | 0/0 | 0/0 | 0/0 | 0/0 | 0/0 | 0/0 | 100 | |
| **chr6** | 53910224 | T | C | 0/0 | 0/0 | 0/0 | 0/0 | 0/0 | 0/0 | 0/0 | 0/0 | 100 | |
| **chr6** | 53910274 | A | C | 0/0 | 0/0 | 0/0 | 0/0 | 0/0 | 0/0 | 0/0 | 0/0 | 100 | |
| **chr6** | 53910311 | T | C | 0/0 | 0/0 | 0/0 | 0/0 | 0/0 | 0/0 | 0/0 | 0/0 | 100 | |
| **chr6** | 53911501 | G | C | 0/1 | 0/0 | 0/0 | 0/0 | 0/0 | 0/0 | 0/0 | 0/0 | 87.5 | |
| **chr6** | 53911534 | G | A | 0/0 | 0/1 | 0/0 | 0/0 | 0/0 | 0/0 | 0/0 | 0/0 | 87.5 | |
| **chr6** | 53911766 | A | G | 1/1 | 1/1 | 0/1 | 1/1 | 0/1 | 0/1 | 1/1 | 0/1 | 0.0 | |
| **chr6** | 53912033 | T | A | 0/1 | 0/0 | 0/0 | 0/0 | 0/0 | 0/0 | 0/0 | 0/0 | 87.5 | |
| **chr6** | 53912377 | C | G | 0/0 | 0/0 | 0/0 | 0/0 | 0/0 | 0/0 | 0/0 | 0/0 | 100 | |
| **chr6** | 53912462 | G | T | 0/1 | 0/0 | 0/0 | 0/0 | 0/0 | 0/0 | 0/0 | 0/0 | 87.5 | |
| **chr6** | 53912519 | C | T | 0/0 | 0/0 | 0/0 | 0/1 | 0/1 | 0/0 | 0/0 | 0/0 | 75.0 | |
| **chr6** | 53912566 | C | T | 0/1 | 0/0 | 0/0 | 0/0 | 0/0 | 0/0 | 0/0 | 0/0 | 87.5 | |

| | | | | | | | | | | | | | |
|---|---|---|---|---|---|---|---|---|---|---|---|---|---|
| **chr6** | 53913491 | T | C | 0/1 | 0/0 | ./. | 0/0 | 0/0 | 0/0 | 0/0 | 0/0 | 85.7 | |
| **chr6** | 53913953 | C | T | 0/0 | 0/0 | ./. | 0/0 | 0/0 | 0/0 | 0/0 | 0/0 | 100 | |
| **chr6** | 53914268 | G | A | 0/0 | 0/0 | 0/0 | 0/0 | 0/0 | 0/0 | 0/0 | 0/0 | 100 | |
| **chr6** | 53914473 | T | C | 0/0 | 0/0 | 0/0 | 0/0 | 0/0 | 0/0 | 0/0 | 0/0 | 100 | |
| **chr6** | 53915039 | T | C | 0/1 | 0/0 | 0/0 | 0/0 | 0/0 | 0/0 | . | 0/0 | 85.7 | X |
| **chr6** | 53916446 | G | C | 0/1 | 0/0 | 0/0 | 0/0 | 0/0 | 0/0 | 0/0 | 0/0 | 87.5 | |
| **chr6** | 53916458 | C | T | 0/0 | 0/0 | 0/0 | 0/0 | 0/0 | 0/0 | 0/0 | 0/0 | 100 | X |
| **chr6** | 53917007 | T | C | 0/1 | 0/0 | 0/0 | 0/0 | 0/0 | 0/0 | 0/0 | 0/0 | 87.5 | |
| **chr6** | 53918629 | C | T | 0/0 | 0/0 | ./. | 0/0 | 0/0 | 0/0 | 0/0 | 0/0 | 100 | X |
| **chr6** | 53918982 | A | C | 0/1 | 0/0 | 0/0 | 0/0 | 0/0 | 0/0 | 0/0 | 0/0 | 87.5 | |
| **chr6** | 53919014 | T | C | 0/1 | 0/0 | 0/0 | 0/0 | 0/0 | 0/0 | 0/0 | 0/0 | 87.5 | X |
| **chr6** | 53919040 | C | T | 0/1 | 0/0 | 0/0 | 0/0 | 0/0 | 0/0 | 0/0 | 0/0 | 87.5 | |
| **chr6** | 53919084 | A | G | 0/0 | 0/0 | 0/0 | 0/0 | 0/0 | 0/0 | 0/0 | 0/0 | 100 | |
| **chr6** | 53919477 | C | T | 0/0 | 0/0 | 0/0 | 0/0 | 0/0 | 0/1 | 0/0 | 0/1 | 75.0 | |
| **chr6** | 53919989 | C | T | 1/1 | 0/0 | 0/0 | 0/0 | 0/0 | 0/0 | 0/1 | 0/0 | 75.0 | |
| **chr6** | 53920145 | T | C | 0/1 | 0/0 | 0/0 | 0/0 | 0/0 | 0/0 | 0/0 | 0/0 | 87.5 | X |
| **chr6** | 53921500 | C | T | 0/0 | 0/0 | 0/0 | 0/0 | 0/0 | 0/0 | 0/0 | 0/0 | 100 | X |
| **chr6** | 53921514 | G | A | 0/1 | 0/0 | 0/0 | 0/0 | 0/0 | 0/0 | 0/0 | 0/0 | 87.5 | X |
| **chr6** | 53921715 | G | A | . | 1/1 | 0/1 | 1/1 | 0/1 | 0/1 | 1/1 | 0/1 | 0.0 | |
| **chr6** | 53923055 | T | C | 0/0 | 0/0 | 0/0 | 0/0 | 0/0 | 0/0 | 0/0 | 0/0 | 100 | |
| **chr6** | 53924906 | G | A | ./. | 0/0 | 0/0 | 0/0 | 0/0 | 0/0 | 0/0 | 0/0 | 100 | X |
| **chr6** | 53927474 | C | A | 0/0 | 0/0 | 0/0 | 0/0 | 0/0 | 0/0 | 0/0 | 0/0 | 100 | |
| **average homozygosity** | | | | | | | | | | | | 88.82 | |

Table S30 - Genomescope2.0 predictions based on HiFi data

| | Sample A1 | | | Sample A2 | | | Sample 2 | | | Sample 3 | | | Sample 4 | | |
|---|---|---|---|---|---|---|---|---|---|---|---|---|---|---|---|
| **Property** | **min** | **max** | **mean** | **min** | **max** | **mean** | **min** | **max** | **mean** | **min** | **max** | **mean** | **min** | **max** | **mean** |
| **Homozygous (aa) (%)** | 98.8248 | 98.8393 | 98.83205 | 98.8456 | 98.8602 | 98.8529 | 98.7786 | 98.8248 | 98.8017 | 98.7622 | 98.7858 | 98.774 | 98.8634 | 98.8843 | 98.87385 |
| **Heterozygous (ab) (%)** | 1.16067 | 1.1752 | 1.167935 | 1.13982 | 1.15442 | 1.14712 | 1.17519 | 1.22142 | 1.198305 | 1.21423 | 1.23777 | 1.226 | 1.11572 | 1.13658 | 1.12615 |
| **Genome Haploid Length (bp)** | 1,060,090,691 | 1,060,747,433 | 1,060,419,062 | 1,087,258,723 | 1,088,738,793 | 1,087,998,758 | 1,080,524,880 | 1,081,680,853 | 1,081,102,867 | 1,061,676,406 | 1,062,843,484 | 1,062,259,945 | 1,078,140,346 | 1,079,282,529 | 1,078,711,438 |
| **Genome Repeat Length (bp)** | 148,917,848 | 149,010,104 | 148,963,976 | 185,998,518 | 186,251,716 | 186,125,117 | 165,703,559 | 165,880,832 | 165,792,196 | 156,530,465 | 156,702,535 | 156,616,500 | 172,221,680 | 172,404,131 | 172,312,906 |
| **Genome Unique Length (bp)** | 911,172,844 | 911,737,328 | 911,455,086 | 901,260,204 | 902,487,077 | 901,873,641 | 914,821,322 | 915,800,020 | 915,310,671 | 905,145,941 | 906,140,949 | 905,643,445 | 905,918,667 | 906,878,398 | 906,398,533 |
| **Model Fit (%)** | 85.263 | 98.2188 | 91.7409 | 83.7977 | 99.4162 | 91.60695 | 84.3326 | 98.5606 | 91.4466 | 85.0648 | 98.8014 | 91.9331 | 84.7102 | 99.5474 | 92.1288 |
| **Read Error Rate (%)** | 0.184164 | 0.184164 | 0.184164 | 0.234586 | 0.234586 | 0.234586 | 0.182921 | 0.182921 | 0.182921 | 0.170348 | 0.170348 | 0.170348 | 0.191115 | 0.191115 | 0.191115 |

**Table S31 - Custom purge_dups cutoffs for HiFi.** The k-mer coverage (kcov) was computed with genomescope2.0.

| Sample | kcov | value1 | value2 |
|---|---|---|---|
| **2** | 12.3 | 18.45 | 55.35 |
| **3** | 9.81 | 14.715 | 44.145 |
| **4** | 10.1 | 15.15 | 45.45 |
| **A1** | 16.5 | 24.75 | 74.25 |
| **A2** | 7.47 | 11.205 | 33.615 |

**Table S32 - Haplotig purging on HiFi assemblies**

| | Sample | | | | |
|---|---|---|---|---|---|
| | **A1** | **A2** | **2** | **3** | **4** |
| **HAPLOTIG** | 67 | 243 | 170 | 115 | 129 |
| **HIGHCOV** | 0 | 1 | 0 | 0 | 0 |
| **JUNK** | 56 | 207 | 119 | 156 | 112 |
| **OVLP** | 76 | 142 | 84 | 92 | 112 |
| **REPEAT** | 472 | 831 | 619 | 710 | 742 |
| **TOT** | 671 | 1424 | 992 | 1073 | 1095 |
| **Removed (Mbp)** | 106 | 153 | 144 | 138 | 162 |

**Table S33 - Parameters used to filter vcf files of different publicly avalable datasets analyzed.** INFO/QUAL: quality score present in the INFO field of the vcf file. INFO/DP: sum of the read depth across all samples for a site. FMT/GQ: quality score in the format field for a genotype. FMT/DP: read depth at a particular site for a sample. Missingness fraction: fraction of missing genotypes at a site. MAF: minor allele frequency threshold. Discarded individuals: number of discarded individuals due to high number of missing genotypes (see Methods).

| Identifier | INFO/QUAL | MAX INFO/DP | FMT/GQ | MAX FMT/DP | MIN FMT/DP | MISSINGNESS FRACTION | MAF | DISCARDED INDIVIDUALS |
|---|---|---|---|---|---|---|---|---|
| **ds2.1** | >30 | 1,367 | >30 | 10 | 2 | <0.1 | >0.05 | / |
| **ds2.2** | >30 | 1,369 | >30 | 9 | 2 | <0.1 | >0.05 | / |
| **ds2.3** | >30 | 1,364 | >30 | 11 | 2 | <0.1 | >0.05 | / |
| **ds2.4** | >30 | 1,368 | >30 | 10 | 2 | <0.1 | >0.05 | / |
| **ds2.5** | >30 | 1,368 | >30 | 10 | 2 | <0.1 | >0.05 | / |
| **ds2.6** | >30 | 1,367 | >30 | 10 | 2 | <0.1 | >0.05 | / |
| **ds2.7** | >30 | 1,368 | >30 | 9 | 2 | <0.1 | >0.05 | / |
| **ds3.1.1** | >30 | 180 | >30 | 9 | 2 | <0.1 | >0.05 | / |
| **ds3.1.2** | >30 | 179 | >30 | 11 | 2 | <0.1 | >0.05 | / |
| **ds3.2.1** | >30 | 703 | >20 | 9 | 2 | <0.2 | >0.05 | / |
| **ds3.2.2** | >30 | 727 | >20 | 7 | 2 | <0.2 | >0.05 | 1 |
| **ds4** | >30 | 8,598 | >20 | 13 | 2 | <0.2 | >0.05 | 5 |
| **ds5** | >30 | 12,852 | >30 | 53 | 5 | <0.3 | >0.02 | 22 |
| **ds6.1** | >20 | 770 | >20 | 3 | 1 | <0.4 | >0.05 | 12 |
| **ds6.2** | >20 | 770 | >20 | 3 | 1 | <0.4 | >0.05 | / |
| **ds6.3** | >20 | 770 | >20 | 3 | 1 | <0.4 | >0.05 | 4 |
| **ds6.4** | >20 | 770 | >20 | 3 | 1 | <0.4 | >0.05 | / |
| **ds6.5** | >20 | 770 | >20 | 3 | 1 | <0.4 | >0.05 | 4 |
| **ds6.6** | >20 | 770 | >20 | 3 | 1 | <0.4 | >0.05 | 18 |
| **ds6.7** | >20 | 770 | >20 | 3 | 1 | <0.4 | >0.05 | 8 |
| **ds6.8** | >20 | 770 | >20 | 3 | 1 | <0.4 | >0.05 | / |

**Table S34 - Missing Chelidonia BUSCOs that were found in bHirRus1.** Information about BUSCO genes in bHirRus1 were retrieved from the "full_table.tsv" BUSCO output.

| BUSCO ID | status in Chelidon | status in bHirRus | bHirRus1 chr. | Gene Start | Gene End | Score | Lenght | OrthoDB url | Description |
|---|---|---|---|---|---|---|---|---|---|
| **129214at7742** | Missing | Complete | SUPER_2 | 43763683 | 43823851 | 1055.5 | 540 | https://www.orthc | probable ATP-dependent RNA helicase DDX10 |
| **239747at7742** | Missing | Complete | SUPER_29_unlo | 53146 | 55386 | 462.2 | 352 | https://www.orthc | Ribosomal RNA adenine dimethylase domain containing 1 |
| **290517at7742** | Missing | Complete | scaffold_270_arr | 31546 | 39663 | 364.8 | 236 | https://www.orthc | eukaryotic translation initiation factor 3 subunit G |
| **295275at7742** | Missing | Complete | SUPER_Z | 38438718 | 38440601 | 465.2 | 326 | https://www.orthc | recQ-mediated genome instability protein 1 |
| **323438at7742** | Missing | Complete | SUPER_Z | 53288635 | 53294590 | 419 | 235 | https://www.orthc | metallo-beta-lactamase domain-containing protein 2 |
| **368989at7742** | Missing | Complete | SUPER_Z | 15924176 | 15931677 | 351 | 192 | https://www.orthc | Transmembrane protein 267 |
| **424874at7742** | Missing | Complete | SUPER_Z | 64307048 | 64309376 | 187.5 | 89 | https://www.orthc | Ubiquitin-like protein ATG12 |
| **46657at7742** | Missing | Complete | SUPER_11 | 15250903 | 15268907 | 1840.8 | 917 | https://www.orthc | teashirt homolog 3 |
| **73217at7742** | Missing | Complete | SUPER_15 | 1638202 | 1645187 | 1250.1 | 626 | https://www.orthc | WD repeat domain 24 |
| **77322at7742** | Missing | Complete | SUPER_8 | 16206525 | 16227942 | 1266.6 | 690 | https://www.orthc | lymphoid-specific helicase |
| **80213at7742** | Missing | Complete | SUPER_Z | 46971972 | 47003188 | 1213.8 | 758 | https://www.orthc | WD repeat-containing protein 36 |
| **139811at7742** | Missing | Fragmented | SUPER_17 | 7998088 | 8025201 | 373.8 | 283 | https://www.orthc | Mitogen-activated protein kinase |
| **36859at7742** | Missing | Fragmented | scaffold_622_arr | 908 | 8003 | 558.3 | 276 | https://www.orthc | symplekin |

**Table S35 - Summary of missing BUSCO in bHirRus1 that were found in Chelidonia, alternate assembly and pipeline intermediates.** The table reports the arbitrary gene ID, the BUSCO gene ID and their presence in all the tested assemblies.

**Legend**

BLASTED busco genes in bHirRus1 (see ST32 for genes information and ST33 for the BLAST search)

Recovered BUSCO genes of the alternate assembly (see ST32b for genes information and ST36a for recovery coordinates)

Recovered BUSCO genes of Chelidonia (see ST32a for genes information and ST36b for recovery coordinates)

| Gene ID | BUSCO ID | bHirRus1 befor | Alternate assem | Chelidonia | c1 | p1 | p2 | bHirRus1 after E | Notes |
|---|---|---|---|---|---|---|---|---|---|
| g1 | 263171at7742 | Missing | Complete | Complete | Complete | Missing | Complete | Missing | Found partially with a BLAST search of Chelidonia gene sequence |
| g2 | 25839at7742 | Missing | Complete | Complete | Complete | Complete | Missing | Missing | Found with a BLAST search of Chelidonia and alternate assembly gene sequence |
| g3 | 47275at7742 | Missing | Complete | Complete | Complete | Complete | Missing | Missing | Found with a BLAST search of Chelidonia and alternate assembly gene sequence |
| g4 | 75522at7742 | Missing | Complete | Complete | Complete | Complete | Missing | Missing | Found with a BLAST search of Chelidonia and alternate assembly gene sequence |
| g5 | 106313at7742 | Missing | Complete | Complete | Complete | Complete | Missing | Missing | Found with a BLAST search of Chelidonia and alternate assembly gene sequence |
| g6 | 274411at7742 | Missing | Complete | Complete | Complete | Complete | Missing | Fragmented | Found with a BLAST search of Chelidonia and alternate assembly gene sequence |
| g7 | 68501at7742 | Missing | Missing | Complete | Complete | Complete | Missing | Missing | Found with a BLAST search of Chelidonia gene sequence |
| g8 | 58506at7742 | Missing | Missing | Complete | Missing | Missing | Missing | Missing | Found with a BLAST search of Chelidonia gene sequence |
| g9 | 59090at7742 | Missing | Missing | Complete | Missing | Missing | Missing | Missing | Found with a BLAST search of Chelidonia gene sequence |
| g10 | 205146at7742 | Missing | Missing | Complete | Missing | Missing | Missing | Missing | Found with a BLAST search of Chelidonia gene sequence |
| g11 | 210721at7742 | Missing | Missing | Complete | Missing | Missing | Missing | Missing | Found with a BLAST search of Chelidonia gene sequence |
| g12 | 283910at7742 | Missing | Missing | Complete | Missing | Missing | Missing | Missing | Found with a BLAST search of Chelidonia gene sequence |
| g13 | 265085at7742 | Missing | Complete | Missing | Complete | Complete | Missing | Missing | Found with a BLAST search of alternate assembly gene sequence |
| g14 | 291147at7742 | Missing | Complete | Missing | Complete | Complete | Missing | Missing | Found with a BLAST search of alternate assembly gene sequence |
| g15 | 259004at7742 | Missing | Complete | Missing | Missing | Complete | Missing | Missing | Found with a BLAST search of alternate assembly gene sequence |
| g16 | 131366at7742 | Missing | Complete | Missing | Missing | Missing | Missing | Missing | Found with a BLAST search of alternate assembly gene sequence |
| g17 | 437433at7742 | Missing | Complete | Missing | Missing | Missing | Missing | Missing | Found with a BLAST search of alternate assembly gene sequence |
| g18 | 333504at7742 | Missing | Fragmented | Missing | Missing | Missing | Missing | Missing | Found with a BLAST search of alternate assembly gene sequence |
| g19 | 149121at7742 | Missing | Missing | Missing | Complete | Complete | Missing | Missing | Found with a BLAST search of p1 gene sequence |
| g20 | 360487at7742 | Missing | Complete | Complete | Complete | Missing | Complete | Complete | |
| g21 | 363461at7742 | Missing | Complete | Complete | Complete | Missing | Complete | Complete | |
| g22 | 419118at7742 | Missing | Complete | Complete | Complete | Missing | Complete | Complete | |
| g23 | 293860at7742 | Missing | Complete | Missing | Complete | Missing | Complete | Complete | |
| g24 | 235274at7742 | Missing | Complete | Complete | Fragmented | Missing | Complete | Complete | |
| g25 | 264645at7742 | Missing | Complete | Complete | Fragmented | Missing | Fragmented | Complete | |
| g26 | 407572at7742 | Missing | Complete | Fragmented | Missing | Missing | Missing | Complete | |
| g27 | 218284at7742 | Missing | Complete | Complete | Fragmented | Missing | Complete | Complete | |
| g28 | 144578at7742 | Missing | Fragmented | Complete | Missing | Missing | Missing | Complete | |
| g29 | 63198at7742 | Missing | Missing | Complete | Missing | Missing | Missing | Complete | |
| g30 | 68766at7742 | Missing | Missing | Complete | Missing | Missing | Missing | Complete | |
| g31 | 122064at7742 | Missing | Missing | Complete | Missing | Missing | Missing | Complete | |
| g32 | 126329at7742 | Missing | Missing | Complete | Missing | Missing | Missing | Complete | |
| g33 | 162664at7742 | Missing | Missing | Complete | Missing | Missing | Missing | Missing | This gene was found with BLAST in bHirRus1 after gene recovery, see ST37 |
| g34 | 191412at7742 | Missing | Missing | Complete | Missing | Missing | Missing | Complete | |
| g35 | 208931at7742 | Missing | Missing | Complete | Missing | Missing | Missing | Complete | |

| | | | | | | | | |
|---|---|---|---|---|---|---|---|---|
| **g36** | 236895at7742 | Missing | Missing | Complete | Missing | Missing | Missing | Complete |
| **g37** | 266219at7742 | Missing | Missing | Complete | Missing | Missing | Missing | Complete |
| **g38** | 267345at7742 | Missing | Missing | Complete | Missing | Missing | Missing | Complete |
| **g39** | 293313at7742 | Missing | Missing | Complete | Missing | Missing | Missing | Complete |
| **g40** | 309129at7742 | Missing | Missing | Complete | Missing | Missing | Missing | Complete |
| **g41** | 309958at7742 | Missing | Missing | Complete | Missing | Missing | Missing | Complete |
| **g42** | 310457at7742 | Missing | Missing | Complete | Missing | Missing | Missing | Complete |
| **g43** | 320379at7742 | Missing | Missing | Complete | Missing | Missing | Missing | Complete |
| **g44** | 334667at7742 | Missing | Missing | Complete | Missing | Missing | Missing | Complete |
| **g45** | 341187at7742 | Missing | Missing | Complete | Missing | Missing | Missing | Complete |
| **g46** | 349414at7742 | Missing | Missing | Complete | Missing | Missing | Missing | Complete |
| **g47** | 357329at7742 | Missing | Missing | Complete | Missing | Missing | Missing | Complete |
| **g48** | 368535at7742 | Missing | Missing | Complete | Missing | Missing | Missing | Complete |
| **g49** | 368798at7742 | Missing | Missing | Complete | Missing | Missing | Missing | Complete |
| **g50** | 378120at7742 | Missing | Missing | Complete | Missing | Missing | Missing | Complete |
| **g51** | 378949at7742 | Missing | Missing | Complete | Missing | Missing | Missing | Complete |
| **g52** | 381001at7742 | Missing | Missing | Complete | Missing | Missing | Missing | Complete |
| **g53** | 381971at7742 | Missing | Missing | Complete | Missing | Missing | Missing | Complete |
| **g54** | 395792at7742 | Missing | Missing | Complete | Missing | Missing | Missing | Complete |
| **g55** | 407723at7742 | Missing | Missing | Complete | Missing | Missing | Missing | Complete |
| **g56** | 410670at7742 | Missing | Missing | Complete | Missing | Missing | Missing | Complete |
| **g57** | 420085at7742 | Missing | Missing | Complete | Missing | Missing | Missing | Complete |
| **g58** | 339203at7742 | Missing | Missing | Duplicated | Missing | Missing | Missing | Complete |
| **g59** | 136144at7742 | Missing | Missing | Fragmented | Missing | Missing | Missing | Fragmented |
| **g60** | 281245at7742 | Missing | Missing | Fragmented | Missing | Missing | Missing | Fragmented |
| **g61** | 330367at7742 | Missing | Missing | Fragmented | Missing | Missing | Missing | Complete |

**Table S36 - Information of BUSCO genes missing from bHirRus1, but found in the other assemblies or intermediates.** Information were retrieved from the BUSCO output "full_table.tsv" for a) Chelidonia; b) alternate assembly; c) p1, intermediate of the VGP pipeline.

**a)**

| Gene ID | Busco id | Status in bHirRu | Status in Chelid | Chelidonia sequ | Gene Start | Gene End | Score | Length | OrthoDB url | Description |
|---|---|---|---|---|---|---|---|---|---|---|
| **g1** | 263171at7742 | Missing | Complete | QRBI01000142.1 | 4878372 | 4901008 | 518.7 | 321 | https://www.ortho | MPN domain-containing protein |
| **g2** | 25839at7742 | Missing | Complete | QRBI01000208.1 | 3403968 | 3418251 | 1893.6 | 1032 | https://www.ortho | DNA mismatch repair protein Msh6 |
| **g3** | 47275at7742 | Missing | Complete | QRBI01000208.1 | 3375445 | 3402719 | 353.6 | 698 | https://www.ortho | F-box only protein 10 |
| **g4** | 75522at7742 | Missing | Complete | QRBI01000223.1 | 87332 | 112771 | 1411.3 | 721 | https://www.ortho | mutS protein homolog 4 |
| **g5** | 106313at7742 | Missing | Complete | QRBI01000116.1 | 22666816 | 22668884 | 535.3 | 422 | https://www.ortho | ataxin-1-like |
| **g6** | 274411at7742 | Missing | Complete | QRBI01000112.1 | 21081776 | 21097751 | 414.6 | 280 | https://www.ortho | Megalencephalic leukoencephalopathy with subcortical cysts 1 |
| **g7** | 68501at7742 | Missing | Complete | QRBI01000107.1 | 11758926 | 11818081 | 987.8 | 773 | https://www.ortho | enteropeptidase |
| **g8** | 58506at7742 | Missing | Complete | QRBI01000270.1 | 31205 | 63502 | 1244.9 | 909 | https://www.ortho | probable cation-transporting ATPase 13A2 |
| **g9** | 59090at7742 | Missing | Complete | QRBI01000123.1 | 53896618 | 53988865 | 1558.9 | 761 | https://www.ortho | Vacuolar protein sorting-associated protein 41 homolog |
| **g10** | 205146at7742 | Missing | Complete | QRBI01000195.1 | 865563 | 895356 | 702.8 | 324 | https://www.ortho | ceramide glucosyltransferase |
| **g11** | 210721at7742 | Missing | Complete | QRBI01000102.1 | 5320500 | 5352337 | 615.4 | 391 | https://www.ortho | dnaJ homolog subfamily C member 21 |
| **g12** | 283910at7742 | Missing | Complete | QRBI01000169.1 | 1627623 | 1686060 | 365.1 | 195 | https://www.ortho | DTW domain-containing protein 2 |
| **g20** | 360487at7742 | Missing | Complete | QRBI01000240.1 | 16972 | 21010 | 403.3 | 249 | https://www.ortho | TELO2-interacting protein 2 |
| **g21** | 363461at7742 | Missing | Complete | QRBI01000152.1 | 2262551 | 2266468 | 300.8 | 162 | https://www.ortho | ATP synthase subunit s, mitochondrial |
| **g22** | 419118at7742 | Missing | Complete | QRBI01000235.1 | 1936562 | 1939520 | 215.8 | 123 | https://www.ortho | Mitochondrial ribosomal protein L20 |
| **g24** | 235274at7742 | Missing | Complete | QRBI01000249.1 | 444782 | 456398 | 854.9 | 439 | https://www.ortho | Synoviolin 1 |
| **g25** | 264645at7742 | Missing | Complete | QRBI01000268.1 | 129061 | 135873 | 426.6 | 311 | https://www.ortho | tRNA modification GTPase GTPBP3, mitochondrial |
| **g26** | 407572at7742 | Missing | Fragmented | QRBI01000205.1 | 146406 | 152708 | 119.5 | 112 | https://www.ortho | alpha-ketoglutarate-dependent dioxygenase alkB homolog 7, mitochondrial |
| **g27** | 218284at7742 | Missing | Complete | QRBI01000116.1 | 21324256 | 21327407 | 397.6 | 263 | https://www.ortho | cytoplasmic tRNA 2-thiolation protein 2 |
| **g28** | 144578at7742 | Missing | Complete | QRBI01000105.1 | 1928349 | 1934461 | 579 | 401 | https://www.ortho | adenosine deaminase domain-containing protein 1 |
| **g29** | 63198at7742 | Missing | Complete | QRBI01000105.1 | 66721255 | 66747819 | 687 | 422 | https://www.ortho | VPS16, CORVET/HOPS core subunit |
| **g30** | 68766at7742 | Missing | Complete | QRBI01000249.1 | 129633 | 133452 | 1029.5 | 693 | https://www.ortho | integrator complex subunit 5 |
| **g31** | 122064at7742 | Missing | Complete | QRBI01000152.1 | 1117805 | 1128803 | 978.6 | 494 | https://www.ortho | kelch-like protein 28 |
| **g32** | 126329at7742 | Missing | Complete | QRBI01000248.1 | 204174 | 212108 | 1383.5 | 684 | https://www.ortho | probable ATP-dependent RNA helicase DDX23 |
| **g33** | 162664at7742 | Missing | Complete | QRBI01000106.1 | 63011072 | 63018103 | 704 | 362 | https://www.ortho | PR domain zinc finger protein 13 |
| **g34** | 191412at7742 | Missing | Complete | QRBI01000120.1 | 91897656 | 91913919 | 500 | 324 | https://www.ortho | Phospholipase A1 member A |
| **g35** | 208931at7742 | Missing | Complete | QRBI01000099.1 | 2727389 | 2757173 | 559.3 | 274 | https://www.ortho | transcription factor AP-2-delta |
| **g36** | 236895at7742 | Missing | Complete | QRBI01000116.1 | 21355634 | 21359393 | 433.3 | 315 | https://www.ortho | DNA replication factor Cdt1 |
| **g37** | 266219at7742 | Missing | Complete | QRBI01000199.1 | 344736 | 347910 | 511.5 | 327 | https://www.ortho | prolactin regulatory element-binding protein |
| **g38** | 267345at7742 | Missing | Complete | QRBI01000249.1 | 353310 | 367170 | 320 | 248 | https://www.ortho | WD repeat domain 74 |
| **g39** | 293313at7742 | Missing | Complete | QRBI01000199.1 | 351238 | 353807 | 458.6 | 293 | https://www.ortho | Ketohexokinase |
| **g40** | 309129at7742 | Missing | Complete | QRBI01000102.1 | 5313240 | 5317727 | 443.4 | 247 | https://www.ortho | ribosome biogenesis protein BRX1 homolog |
| **g41** | 309958at7742 | Missing | Complete | QRBI01000123.1 | 946786 | 949272 | 275.1 | 180 | https://www.ortho | IBA57, iron-sulfur cluster assembly |
| **g42** | 310457at7742 | Missing | Complete | QRBI01000102.1 | 5297043 | 5313074 | 454.3 | 239 | https://www.ortho | cell cycle checkpoint protein RAD1 |
| **g43** | 320379at7742 | Missing | Complete | QRBI01000119.1 | 7955301 | 7964874 | 387.6 | 224 | https://www.ortho | lysosome-associated membrane glycoprotein 5 |
| **g44** | 334667at7742 | Missing | Complete | QRBI01000103.1 | 116693 | 136355 | 237 | 190 | https://www.ortho | spindle and kinetochore-associated protein 1 |
| **g45** | 341187at7742 | Missing | Complete | QRBI01000205.1 | 273 | 17157 | 339.1 | 206 | https://www.ortho | Mitochondrial ribosomal protein L4 |
| **g46** | 349414at7742 | Missing | Complete | QRBI01000240.1 | 2760875 | 2769103 | 338.6 | 182 | https://www.ortho | transmembrane protein 127 |
| **g47** | 357329at7742 | Missing | Complete | QRBI01000099.1 | 24804523 | 24814779 | 379.3 | 193 | https://www.ortho | ras-related protein Rab-23 |
| **g48** | 368535at7742 | Missing | Complete | QRBI01000189.1 | 945590 | 950066 | 82 | 89 | https://www.ortho | B-cell antigen receptor complex-associated protein beta chain |
| **g49** | 368798at7742 | Missing | Complete | QRBI01000260.1 | 51393 | 54218 | 360.6 | 216 | https://www.ortho | coiled-coil domain-containing protein 130 |
| **g50** | 378120at7742 | Missing | Complete | QRBI01000194.1 | 810966 | 813762 | 447.8 | 230 | https://www.ortho | Transmembrane protein 101 |
| **g51** | 378949at7742 | Missing | Complete | QRBI01000098.1 | 611589 | 613898 | 302.1 | 147 | https://www.ortho | dynactin subunit 5 |

| | | | | | | | | | | |
|---|---|---|---|---|---|---|---|---|---|---|
| **g52** | 381001at7742 | Missing | Complete | QRBI01000240.1 | 2629493 | 2633917 | 224.6 | 135 | https://www.orthc | Mitochondrial ribosomal protein S24 |
| **g53** | 381971at7742 | Missing | Complete | QRBI01000340.1 | 83384 | 89355 | 145 | 110 | https://www.orthc | succinate dehydrogenase cytochrome b560 subunit, mitochondrial |
| **g54** | 395792at7742 | Missing | Complete | QRBI01000123.1 | 29639863 | 29640426 | 207.6 | 140 | https://www.orthc | fer3-like protein |
| **g55** | 407723at7742 | Missing | Complete | QRBI01000043.1 | 2461 | 10859 | 187.8 | 107 | https://www.orthc | astrocytic phosphoprotein PEA-15 |
| **g56** | 410670at7742 | Missing | Complete | QRBI01000153.1 | 982835 | 987434 | 202.7 | 99 | https://www.orthc | RNA polymerase I subunit D |
| **g57** | 420085at7742 | Missing | Complete | QRBI01000268.1 | 424857 | 432729 | 159.7 | 108 | https://www.orthc | nuclear receptor 2C2-associated protein |
| **g58** | 339203at7742 | Missing | Duplicated | QRBI01000248.1 | 430368 | 433482 | 302.2 | 199 | https://www.orthc | tRNA (guanine-N(7)-)-methyltransferase |
| **g59** | 136144at7742 | Missing | Fragmented | QRBI01000246.1 | 199035 | 207813 | 400.3 | 295 | https://www.orthc | nicastrin |
| **g60** | 281245at7742 | Missing | Fragmented | QRBI01000012.1 | 57802 | 67653 | 199.5 | 130 | https://www.orthc | lymphokine-activated killer T-cell-originated protein kinase |
| **g61** | 330367at7742 | Missing | Fragmented | QRBI01000258.1 | 447908 | 452949 | 269.1 | 151 | https://www.orthc | FosB proto-oncogene, AP-1 transcription factor subunit |

**b)**

| Gene ID | Busco id | Status in bHirRu | Status in alterna | Alternate seque | Gene Start | Gene End | Score | Length | OrthoDB url | Description |
|---|---|---|---|---|---|---|---|---|---|---|
| **g1** | 263171at7742 | Missing | Complete | 001562F\|arrow\|a | 47326 | 51319 | 662 | 358 | https://www.orthc | MPN domain-containing protein |
| **g2** | 25839at7742 | Missing | Complete | 000069F_009\|ar | 60740 | 75022 | 1891.4 | 1032 | https://www.orthc | DNA mismatch repair protein Msh6 |
| **g3** | 47275at7742 | Missing | Complete | 000069F_009\|ar | 33176 | 59491 | 350.1 | 676 | https://www.orthc | F-box only protein 10 |
| **g4** | 75522at7742 | Missing | Complete | 000268F_001\|ar | 580500 | 605941 | 1411.3 | 721 | https://www.orthc | mutS protein homolog 4 |
| **g5** | 106313at7742 | Missing | Complete | 000280F_006\|ar | 158017 | 160086 | 593.9 | 422 | https://www.orthc | ataxin-1-like |
| **g6** | 274411at7742 | Missing | Complete | 000084F_017\|ar | 1144760 | 1160739 | 414.6 | 280 | https://www.orthc | Megalencephalic leukoencephalopathy with subcortical cysts 1 |
| **g13** | 265085at7742 | Missing | Complete | 000160F_001\|ar | 726701 | 812685 | 563.2 | 298 | https://www.orthc | Branched chain keto acid dehydrogenase E1 subunit beta |
| **g14** | 291147at7742 | Missing | Complete | 000235F_006\|ar | 777922 | 819309 | 438.9 | 278 | https://www.orthc | WD repeat-containing protein 25 |
| **g15** | 259004at7742 | Missing | Complete | 000038F_014\|ar | 2534502 | 2545088 | 478.5 | 279 | https://www.orthc | pituitary-specific positive transcription factor 1 |
| **g16** | 131366at7742 | Missing | Complete | 000180F_002\|ar | 685387 | 718596 | 580.9 | 324 | https://www.orthc | Spastin |
| **g17** | 437433at7742 | Missing | Complete | 000031F_006\|ar | 351016 | 353103 | 115.3 | 77 | https://www.orthc | selenoprotein K |
| **g18** | 333504at7742 | Missing | Fragmented | 000018F_013\|ar | 310834 | 322473 | 248.8 | 136 | https://www.orthc | type II iodothyronine deiodinase |
| **g20** | 360487at7742 | Missing | Complete | 000989F\|arrow\|a | 133501 | 141103 | 403.5 | 249 | https://www.orthc | TELO2-interacting protein 2 |
| **g21** | 363461at7742 | Missing | Complete | 000910F\|arrow\|a | 13913 | 17843 | 300.8 | 162 | https://www.orthc | ATP synthase subunit s, mitochondrial |
| **g22** | 419118at7742 | Missing | Complete | 001128F\|arrow\|a | 71973 | 74747 | 215.8 | 123 | https://www.orthc | Mitochondrial ribosomal protein L20 |
| **g23** | 293860at7742 | Missing | Complete | 001400F\|arrow\|a | 64238 | 69827 | 337.2 | 226 | https://www.orthc | probable cytosolic iron-sulfur protein assembly protein CIAO1 |
| **g24** | 235274at7742 | Missing | Complete | 001510F\|arrow\|a | 51595 | 55923 | 698.6 | 316 | https://www.orthc | Synoviolin 1 |
| **g25** | 264645at7742 | Missing | Complete | 001189F\|arrow\|a | 91200 | 97204 | 430.3 | 299 | https://www.orthc | tRNA modification GTPase GTPBP3, mitochondrial |
| **g26** | 407572at7742 | Missing | Complete | 001439F_001\|ar | 7205 | 8567 | 188.8 | 155 | https://www.orthc | alpha-ketoglutarate-dependent dioxygenase alkB homolog 7, mitochondrial |
| **g27** | 218284at7742 | Missing | Complete | 001299F\|arrow\|a | 71975 | 75127 | 398.2 | 268 | https://www.orthc | cytoplasmic tRNA 2-thiolation protein 2 |
| **g28** | 144578at7742 | Missing | Fragmented | 000162F_007\|ar | 1 | 5173 | 447.4 | 321 | https://www.orthc | adenosine deaminase domain-containing protein 1 |

**c)**

| Gene ID | Busco ID | Status in bHirRu | Status in p1 | p1 sequence | Gene Start | Gene End | Score | Length | OrthoDB url | Description |
|---|---|---|---|---|---|---|---|---|---|---|
| **g19** | 149121at7742 | Missing | Complete | 002029F\|arrow | 618 | 24009 | 745.4 | 467 | https://www.orthc | IlvB acetolactate synthase like |

**Table S37 - BLAST search of the missing genes in bHirRus1.** The table reports the output of the BLAST search of the 139 genes missing in bHirRus1, that were found in the other assemblies/intermediates. Sequences were retrieved from the corresponding assembly/intermediates and blasted on bHirRus1.

a)

| Gene ID | BUSCO gene ID | Chelidonia scaf | Gene start in th | Gene end in the | Gene length in t | bHirRus1 chr be | % identity | alignment lengt | mismatches | gap opens | q. start | q. end | s. start | s. end | evalue | bit score |
|---|---|---|---|---|---|---|---|---|---|---|---|---|---|---|---|---|
| **g1** | 263171at7742 | QRBI01000142.1 | 4878372 | 4901008 | 22636 | SUPER_26 | 95.379 | 3571 | 108 | 31 | 1 | 3559 | 917655 | 914130 | 0 | 5627 |
| **g1** | 263171at7742 | QRBI01000142.1 | 4878372 | 4901008 | 22636 | SUPER_26 | 93.81 | 4152 | 111 | 35 | 2828 | 6913 | 914585 | 910514 | 0 | 6109 |
| **g1** | 263171at7742 | QRBI01000142.1 | 4878372 | 4901008 | 22636 | SUPER_26 | 97.057 | 8935 | 133 | 61 | 6756 | 15623 | 910800 | 901929 | 0 | 14923 |
| **g2** | 25839at7742 | QRBI01000208.1 | 3403968 | 3418251 | 14283 | SUPER_3 | 99.484 | 14330 | 5 | 65 | 1 | 14283 | 16181132 | 16195439 | 0 | 25989 |
| **g3** | 47275at7742 | QRBI01000208.1 | 3375445 | 3402719 | 27274 | SUPER_3 | 98.74 | 27462 | 60 | 259 | 1 | 27274 | 16152515 | 16179878 | 0 | 48532 |
| **g4** | 75522at7742 | QRBI01000223.1 | 87332 | 112771 | 25439 | SUPER_8 | 99.076 | 25546 | 54 | 166 | 1 | 25439 | 36058293 | 36032823 | 0 | 45700 |
| **g5** | 106313at7742 | QRBI01000116.1 | 22666816 | 22668884 | 2068 | SUPER_11 | 98.662 | 2092 | 1 | 21 | 1 | 2068 | 165887 | 163799 | 0 | 3683 |
| **g6** | 274411at7742 | QRBI01000112.1 | 21081776 | 21097751 | 15975 | SUPER_4 | 99.444 | 7559 | 24 | 9 | 96 | 7640 | 19616307 | 19608753 | 0 | 13710 |
| **g6** | 274411at7742 | QRBI01000112.1 | 21081776 | 21097751 | 15975 | SUPER_4 | 99.497 | 8356 | 27 | 8 | 7629 | 15975 | 19608817 | 19600468 | 0 | 15184 |
| **g7** | 68501at7742 | QRBI01000107.1 | 11758926 | 11818081 | 59155 | SUPER_2 | 99.412 | 30270 | 46 | 82 | 1 | 30240 | 11417992 | 11387825 | 0 | 54791 |
| **g7** | 68501at7742 | QRBI01000107.1 | 11758926 | 11818081 | 59155 | SUPER_2 | 99.102 | 28950 | 153 | 49 | 30261 | 59155 | 11387851 | 11358954 | 0 | 51921 |
| **g8** | 58506at7742 | QRBI01000270.1 | 31205 | 63502 | 32297 | SUPER_22 | 99.567 | 32343 | 30 | 86 | 1 | 32297 | 56641 | 24363 | 0 | 58850 |
| **g9** | 59090at7742 | QRBI01000123.1 | 53896618 | 53988865 | 92247 | SUPER_1 | 99.568 | 34060 | 75 | 34 | 1 | 34020 | 1.01E+08 | 1.01E+08 | 0 | 62017 |
| **g10** | 205146at7742 | QRBI01000195.1 | 865563 | 895356 | 29793 | SUPER_Z | 99.872 | 25840 | 3 | 27 | 2 | 25820 | 65238702 | 65212872 | 0 | 47508 |
| **g11** | 210721at7742 | QRBI01000102.1 | 5320500 | 5352337 | 31837 | SUPER_Z | 99.854 | 23908 | 0 | 34 | 7956 | 31837 | 19095024 | 19071126 | 0 | 43923 |
| **g12** | 283910at7742 | QRBI01000169.1 | 1627623 | 1686060 | 58437 | SUPER_Z | 99.861 | 28732 | 0 | 37 | 1 | 28697 | 63012691 | 62983965 | 0 | 52800 |

b)

| Gene ID | BUSCO gene ID | Alternate scaffo | Gene start in th | Gene end in the | Gene length in t | bHirRus1 chr be | % identity | alignment lengt | mismatches | gap opens | q. start | q. end | s. start | s. end | evalue | bit score |
|---|---|---|---|---|---|---|---|---|---|---|---|---|---|---|---|---|
| **g2** | 25839at7742 | 000069F_009\|ar | 60740 | 75022 | 14282 | SUPER_3 | 99.442 | 14329 | 12 | 65 | 1 | 14282 | 16181132 | 16195439 | 0 | 2.60E+04 |
| **g3** | 47275at7742 | 000069F_009\|ar | 33176 | 59491 | 26315 | SUPER_3 | 98.683 | 26500 | 68 | 255 | 1 | 26315 | 16153475 | 16179878 | 0 | 46743 |
| **g4** | 75522at7742 | 000268F_001\|ar | 580500 | 605941 | 25441 | SUPER_8 | 99.068 | 25547 | 56 | 166 | 1 | 25441 | 36032824 | 36058294 | 0 | 45690 |
| **g5** | 106313at7742 | 000280F_006\|ar | 158017 | 160086 | 2069 | SUPER_11 | 98.709 | 2092 | 1 | 20 | 1 | 2069 | 163800 | 165888 | 0 | 3690 |
| **g6** | 274411at7742 | 000084F_017\|ar | 1144760 | 1160739 | 15979 | SUPER_4 | 99.39 | 8357 | 34 | 8 | 7633 | 15979 | 19608817 | 19600468 | 0 | 15134 |
| **g6** | 274411at7742 | 000084F_017\|ar | 1144760 | 1160739 | 15979 | SUPER_4 | 99.392 | 7561 | 28 | 10 | 96 | 7644 | 19616307 | 19608753 | 0 | 13692 |
| **g13** | 265085at7742 | 000160F_001\|ar | 726701 | 812685 | 85984 | SUPER_3 | 99.312 | 86154 | 271 | 149 | 1 | 85984 | 83045036 | 82959035 | 0 | 1.56E+05 |
| **g14** | 291147at7742 | 000235F_006\|ar | 777922 | 819309 | 41387 | SUPER_6 | 99.015 | 29650 | 197 | 38 | 1 | 29623 | 11855573 | 11825992 | 0 | 53047 |
| **g14** | 291147at7742 | 000235F_006\|ar | 777922 | 819309 | 41387 | SUPER_6 | 99.201 | 10513 | 55 | 16 | 29656 | 40149 | 11825990 | 11815488 | 0 | 18921 |
| **g15** | 259004at7742 | 000038F_014\|ar | 2534502 | 2545088 | 10586 | SUPER_2 | 98.644 | 10621 | 98 | 17 | 1 | 10586 | 5796576 | 5785967 | 0 | 18774 |
| **g16** | 131366at7742 | 000180F_002\|ar | 685387 | 718596 | 33209 | SUPER_3 | 98.822 | 33281 | 286 | 46 | 1 | 33209 | 35129109 | 35162355 | 0 | 59189 |
| **g17** | 437433at7742 | 000031F_006\|ar | 351016 | 353103 | 2087 | SUPER_12 | 98.132 | 2088 | 30 | 3 | 1 | 2087 | 3108423 | 3110502 | 0 | 3631 |
| **g18** | 333504at7742 | 000018F_013\|ar | 310834 | 322473 | 11639 | SUPER_6 | 99.278 | 11640 | 66 | 6 | 1 | 11639 | 20819295 | 20830917 | 0 | 21014 |

c)

| Gene ID | BUSCO gene ID | p1 contig | Gene start in th | Gene end in the | Gene length in t | bHirRus1 chr be | % identity | alignment lengt | mismatches | gap opens | q. start | q. end | s. start | s. end | evalue | bit score |
|---|---|---|---|---|---|---|---|---|---|---|---|---|---|---|---|---|
| **g19** | 149121at7742 | 002029F\|arrow | 618 | 24009 | 23391 | scaffold_397_arr | 99.838 | 23399 | 12 | 26 | 1 | 23391 | 26797 | 3417 | 0 | 42976 |

**Table S38 - Purged c1 sequences that contained BUSCO genes.** The table reports: the c1 contig; start and end coordinates of the purged sequence by purge_dups; type of the purged sequence; BUSCO ID of the gene that was included in the purged sequence; the status in which the gene was found in c1 by BUSCO; start and end coordinates of the gene in the c1 contig; BUSCO score; the OrthoDB URL; gene description.
Infomation about purged sequences were recovered from purge_dups "dups.bed" output, while BUSCO genes infomation were retrieved from BUSCO "full_table.tsv" output for c1.

| c1 contig | start purged | end purged | type | Gene ID of the c | Busco ID of the | Status in c1 | Gene Start in th | Gene End in the | Score | Length | OrthoDB url | Description |
|---|---|---|---|---|---|---|---|---|---|---|---|---|
| **000989F\|arrow** | 1 | 140961 | HAPLOTIG | g20 | 360487at7742 | Complete | 133540 | 137580 | 403.3 | 249 | https://www.orthc | TELO2-interacting protein 2 |
| **000910F\|arrow** | 1 | 171175 | HAPLOTIG | g21 | 363461at7742 | Complete | 13921 | 17853 | 300.8 | 162 | https://www.orthc | ATP synthase subunit s, mitochonc |
| **001128F\|arrow** | 1 | 107330 | HAPLOTIG | g22 | 419118at7742 | Complete | 98276 | 101051 | 215.8 | 123 | https://www.orthc | Mitochondrial ribosomal protein L2 |
| **001400F\|arrow** | 1 | 70212 | REPEAT | g23 | 293860at7742 | Complete | 64306 | 70086 | 339.6 | 219 | https://www.orthc | probable cytosolic iron-sulfur prote |
| **001510F\|arrow** | 1 | 59168 | HAPLOTIG | g24 | 235274at7742 | Fragmented | 51702 | 56035 | 642.5 | 291 | https://www.orthc | Synoviolin 1 |
| **001189F\|arrow** | 1 | 97110 | HAPLOTIG | g25 | 264645at7742 | Fragmented | 91315 | 97110 | 364 | 272 | https://www.orthc | tRNA modification GTPase GTPBF |
| **001299F\|arrow** | 1 | 81927 | HAPLOTIG | g27 | 218284at7742 | Fragmented | 73759 | 75105 | 267.7 | 173 | https://www.orthc | cytoplasmic tRNA 2-thiolation prote |

**Table S39 - Alignment coordinates for BUSCO genes recovery. a)** Chelidonia scaffolds containing the missing genes in bHirRus1 were aligned to the latter. The matching alignment row was reported if the gene coordinates aligned completely. For the genes that were actually missing in bHirRus1, the alignment rows around the gene coordinates were reported, in order to select the coordinates of the Chelidonia scaffold fragment that has to be recovered. The first 6 columns report information about the gene included in the aligned scaffold. The remaining columns were taken from MUMMer NUCmer alignment coordinates. **b)** Same procedure in alternate assembly. **c)** Same procedure in the p1 intermediate.

**a)**

| Gene ID | BUSCO ID | Gene status in ( | Gene start | Gene end | Gene sequence | Query seq. ID | Q. scaffold leng | Q. start alignme | Q. end alignmer | Q. alignment ler | Reference seq. | R. chr length | R. start alignme | R. end alignmer | R. alignment ler | % identity | % similarity | strand of the qu | Start missing se | End missing se | Notes |
|---|---|---|---|---|---|---|---|---|---|---|---|---|---|---|---|---|---|---|---|---|---|
| **g1** | 263171at7742 | Complete | 4878372 | 4901008 | 22636 | QRBI01000142.1 | 5062783 | 4893537 | 4885421 | 8117 | SUPER_26 | 5553549 | 902419 | 910526 | 8107 | 98.91879 | 98.91879 | Minus | - | - | Part of the query gene sequence aligned |
| **g2** | 25839at7742 | Complete | 3403968 | 3418251 | 14283 | QRBI01000208.1 | 3457278 | 1861356 | 3431559 | 1570204 | SUPER_3 | 1.17E+08 | 14637965 | 16208754 | 1570789 | 99.69991 | 99.69991 | Plus | - | - | Query gene sequence aligned entirely |
| **g3** | 47275at7742 | Complete | 3375445 | 3402719 | 27274 | QRBI01000208.1 | 3457278 | 1861356 | 3431559 | 1570204 | SUPER_3 | 1.17E+08 | 14637965 | 16208754 | 1570789 | 99.69991 | 99.69991 | Plus | - | - | Query gene sequence aligned entirely |
| **g4** | 75522at7742 | Complete | 87332 | 112771 | 25439 | QRBI01000223.1 | 1164200 | 164855 | 75174 | 89682 | SUPER_8 | 36085389 | 35980713 | 36070454 | 89741 | 99.23019 | 99.23019 | Minus | - | - | Query gene sequence aligned entirely |
| **g5** | 106313at7742 | Complete | 22666816 | 22668884 | 2068 | QRBI01000116.1 | 22866040 | 2297053 | 2307661 | 10609 | SUPER_11 | 21491857 | 19997100 | 20007710 | 10610 | 99.28525 | 99.28525 | Plus | - | - | Query gene sequence aligned entirely |
| **g6** | 274411at7742 | Complete | 21081776 | 21097751 | 15975 | QRBI01000112.1 | 40088381 | 21123704 | 21029027 | 94678 | SUPER_4 | 76187387 | 19574515 | 19669115 | 94600 | 99.21317 | 99.21317 | Minus | - | - | Query gene sequence aligned entirely |
| **g7** | 68501at7742 | Complete | 11758926 | 11818081 | 59155 | QRBI01000107.1 | 14885125 | 11902352 | 11457459 | 444894 | SUPER_2 | 1.19E+08 | 11274764 | 11719436 | 444672 | 99.20023 | 99.20023 | Minus | - | - | Query gene sequence aligned entirely |
| **g8** | 58506at7742 | Complete | 31205 | 63502 | 32297 | QRBI01000270.1 | 947175 | 78935 | 3977 | 74959 | SUPER_22 | 7.51E+06 | 8.91E+03 | 8.39E+04 | 7.50E+04 | 99.48607 | 99.48607 | Minus | - | - | Query gene sequence aligned entirely |
| **g9** | 59090at7742 | Complete | 53896618 | 53988865 | 92247 | QRBI01000123.1 | 55481500 | 53930638 | 53678205 | 252434 | SUPER_1 | 1.56E+08 | 1.01E+08 | 1.02E+08 | 1.00E+06 | 99.79887 | 99.79887 | Minus | - | - | Query gene sequence aligned entirely |
| **g10** | 205146at7742 | Complete | 865563 | 895356 | 29793 | QRBI01000195.1 | 1737651 | 891383 | 404472 | 486912 | SUPER_Z | 90132487 | 65212872 | 65699857 | 486985 | 99.9101 | 99.9101 | Minus | - | - | Query gene sequence aligned entirely |
| **g11** | 210721at7742 | Complete | 5320500 | 5352337 | 31837 | QRBI01000102.1 | 9438414 | 5860533 | 5328456 | 532078 | SUPER_Z | 90132487 | 18562787 | 19095024 | 532237 | 99.89013 | 99.89013 | Minus | - | - | Query gene sequence aligned entirely |
| **g12** | 283910at7742 | Complete | 1627623 | 1686060 | 58437 | QRBI01000169.1 | 3685968 | 1656320 | 1380509 | 275812 | SUPER_Z | 9.01E+07 | 62983965 | 63259744 | 275779 | 99.84854 | 99.84854 | Minus | - | - | Query gene sequence aligned entirely |
| **g28** | 144578at7742 | Complete | 1928349 | 1934461 | 6112 | QRBI01000105.1 | 66802864 | 1925492 | 1585559 | 339934 | SUPER_5 | 73257097 | 65575466 | 65915542 | 340076 | 99.57864 | 99.57864 | Minus | 1925492 | 1967238 | Query gene sequence not aligned |
| **g28** | 144578at7742 | Complete | 1928349 | 1934461 | 6112 | QRBI01000105.1 | 66802864 | 2044755 | 1967238 | 77518 | SUPER_5 | 73257097 | 65456638 | 65534086 | 77448 | 99.40066 | 99.40066 | Minus | | | |
| **g29** | 63198at7742 | Complete | 66721255 | 66747819 | 26564 | QRBI01000105.1 | 66802864 | 66708817 | 66495102 | 213716 | SUPER_5 | 7.33E+07 | 1124142 | 1337831 | 213689 | 99.53743 | 99.53743 | Minus | 66708817 | 66802864 | Query gene sequence not aligned |
| **g30** | 68766at7742 | Complete | 129633 | 133452 | 3819 | QRBI01000249.1 | 721187 | 90113 | 108525 | 18413 | scaffold_493_arr | 18466 | 1 | 18466 | 18465 | 98.53259 | 98.53259 | Plus | 108525 | 200600 | Query gene sequence not aligned |
| **g30** | 68766at7742 | Complete | 129633 | 133452 | 3819 | QRBI01000249.1 | 721187 | 210864 | 200600 | 10265 | scaffold_587_arr | 10268 | 1 | 10268 | 10267 | 99.80533 | 99.80533 | Minus | | | |
| **g31** | 122064at7742 | Complete | 1117805 | 1128803 | 10998 | QRBI01000152.1 | 19025920 | 1026263 | 1111216 | 84954 | SUPER_6 | 63258489 | 821150 | 906062 | 84912 | 98.99326 | 98.99326 | Plus | 1111216 | 1128958 | Query gene sequence not aligned |
| **g31** | 122064at7742 | Complete | 1117805 | 1128803 | 10998 | QRBI01000152.1 | 19025920 | 1128958 | 1165874 | 36917 | SUPER_6 | 63258489 | 906163 | 943015 | 36852 | 98.72636 | 98.72636 | Plus | | | |
| **g32** | 126329at7742 | Complete | 204174 | 212108 | 7934 | QRBI01000248.1 | 455452 | 165169 | 189516 | 24348 | SUPER_30 | 1648998 | 1442837 | 1467198 | 24361 | 99.17721 | 99.17721 | Plus | 189516 | 216835 | Query gene sequence not aligned |
| **g32** | 126329at7742 | Complete | 204174 | 212108 | 7934 | QRBI01000248.1 | 455452 | 216835 | 245532 | 28698 | SUPER_30 | 1648998 | 1468328 | 1497091 | 28763 | 99.16363 | 99.16363 | Plus | | | |
| **g33** | 162664at7742 | Complete | 63011072 | 63018103 | 7031 | QRBI01000106.1 | 79407264 | 63007889 | 63009785 | 1897 | SUPER_3 | 1.17E+08 | 74978681 | 74980575 | 1894 | 98.89532 | 98.89532 | Plus | 63009785 | 63027227 | Query gene sequence not aligned |
| **g33** | 162664at7742 | Complete | 63011072 | 63018103 | 7031 | QRBI01000106.1 | 79407264 | 63027227 | 63058508 | 31282 | SUPER_3 | 1.17E+08 | 74981569 | 75012871 | 31302 | 99.384 | 99.384 | Plus | | | |
| **g34** | 191412at7742 | Complete | 91897656 | 91913919 | 16263 | QRBI01000120.1 | 96661620 | 91892976 | 91879579 | 13398 | SUPER_2 | 1.19E+08 | 26257439 | 26270843 | 13404 | 99.42597 | 99.42597 | Minus | 91892976 | 91936698 | Query gene sequence not aligned |
| **g34** | 191412at7742 | Complete | 91897656 | 91913919 | 16263 | QRBI01000120.1 | 96661620 | 92291648 | 91936698 | 354951 | SUPER_2 | 1.19E+08 | 25858818 | 26213694 | 354876 | 99.46492 | 99.46492 | Minus | | | |
| **g35** | 208931at7742 | Complete | 2727389 | 2757173 | 29784 | QRBI01000099.1 | 24819762 | 2725996 | 2650828 | 75169 | SUPER_3 | 1.17E+08 | 1.13E+08 | 1.13E+08 | 75137 | 99.45001 | 99.45001 | Minus | 2725996 | 2758525 | Query gene sequence not aligned |
| **g35** | 208931at7742 | Complete | 2727389 | 2757173 | 29784 | QRBI01000099.1 | 24819762 | 2758525 | 2777756 | 19232 | SUPER_3 | 1.17E+08 | 75377071 | 75396302 | 19231 | 100 | 0.046797 | Plus | | | |
| **g36** | 236895at7742 | Complete | 21355634 | 21359393 | 3759 | QRBI01000116.1 | 22866040 | 21323815 | 21317239 | 6577 | SUPER_11 | 21491857 | 1508826 | 1515424 | 6598 | 98.38149 | 98.38149 | Minus | 21323815 | 21360556 | Query gene sequence not aligned |
| **g36** | 236895at7742 | Complete | 21355634 | 21359393 | 3759 | QRBI01000116.1 | 22866040 | 21390174 | 21360556 | 29619 | SUPER_11 | 21491857 | 1441921 | 1471502 | 29581 | 99.32204 | 99.32204 | Minus | | | |
| **g37** | 266219at7742 | Complete | 344736 | 347910 | 3174 | QRBI01000199.1 | 606848 | 334638 | 309206 | 25433 | SUPER_3 | 1.17E+08 | 1.16E+08 | 1.17E+08 | 25446 | 99.30133 | 99.30133 | Minus | 334638 | 364196 | Query gene sequence not aligned |
| **g37** | 266219at7742 | Complete | 344736 | 347910 | 3174 | QRBI01000199.1 | 606848 | 364196 | 396729 | 32534 | SUPER_3 | 1.17E+08 | 1.16E+08 | 1.17E+08 | 32523 | 99.63458 | 99.63458 | Plus | | | |
| **g38** | 267345at7742 | Complete | 353310 | 367170 | 13860 | QRBI01000249.1 | 721187 | 334945 | 253665 | 81281 | scaffold_121_arr | 129379 | 48071 | 129379 | 81308 | 99.6401 | 99.6401 | Minus | 334945 | 499425 | Query gene sequence not aligned |
| **g38** | 267345at7742 | Complete | 353310 | 367170 | 13860 | QRBI01000249.1 | 721187 | 499425 | 505594 | 6170 | SUPER_1 | 1.56E+08 | 1.54E+08 | 1.54E+08 | 6169 | 100 | 0.081037 | Plus | | | |
| **g39** | 293313at7742 | Complete | 351238 | 353807 | 2569 | QRBI01000199.1 | 606848 | 334638 | 309206 | 25433 | SUPER_3 | 1.17E+08 | 1.16E+08 | 1.17E+08 | 25446 | 99.30133 | 99.30133 | Minus | 334638 | 364196 | Query gene sequence not aligned |
| **g39** | 293313at7742 | Complete | 351238 | 353807 | 2569 | QRBI01000199.1 | 606848 | 364196 | 396729 | 32534 | SUPER_3 | 1.17E+08 | 1.16E+08 | 1.17E+08 | 32523 | 99.63458 | 99.63458 | Plus | | | |
| **g39** | 293313at7742 | Complete | 351238 | 353807 | 2569 | QRBI01000199.1 | 606848 | 334638 | 309206 | 25433 | SUPER_3 | 1.17E+08 | 1.16E+08 | 1.17E+08 | 25446 | 99.30133 | 99.30133 | Minus | 334638 | 364196 | Query gene sequence not aligned |
| **g39** | 293313at7742 | Complete | 351238 | 353807 | 2569 | QRBI01000199.1 | 606848 | 364196 | 396729 | 32534 | SUPER_3 | 1.17E+08 | 1.16E+08 | 1.17E+08 | 32523 | 99.63458 | 99.63458 | Plus | | | |
| **g40** | 309129at7742 | Complete | 5313240 | 5317727 | 4487 | QRBI01000102.1 | 9438414 | 5296419 | 5286423 | 9997 | SUPER_4 | 76187387 | 6759419 | 6769417 | 9998 | 99.81 | 99.81 | Minus | 5296419 | 5328456 | Query gene sequence not aligned |
| **g40** | 309129at7742 | Complete | 5313240 | 5317727 | 4487 | QRBI01000102.1 | 9438414 | 5860533 | 5328456 | 532078 | SUPER_Z | 90132487 | 18562787 | 19095024 | 532237 | 99.89013 | 99.89013 | Minus | | | |
| **g41** | 309958at7742 | Complete | 946786 | 949272 | 2486 | QRBI01000123.1 | 55481500 | 935724 | 934177 | 1548 | SUPER_Z | 90132487 | 62516068 | 62517615 | 1547 | 100 | 2.196382 | Minus | 935724 | 987011 | Query gene sequence not aligned |
| **g41** | 309958at7742 | Complete | 946786 | 949272 | 2486 | QRBI01000123.1 | 55481500 | 1018346 | 987011 | 31336 | SUPER_1 | 1.56E+08 | 1.54E+08 | 1.54E+08 | 31319 | 98.81446 | 98.81446 | Minus | | | |
| **g42** | 310457at7742 | Complete | 5297043 | 5313074 | 16031 | QRBI01000102.1 | 9438414 | 5286421 | 5296418 | 9998 | SUPER_Z | 90132487 | 26815412 | 26825412 | 10000 | 99.69006 | 99.69006 | Plus | 5296418 | 5328456 | Query gene sequence not aligned |
| **g42** | 310457at7742 | Complete | 5297043 | 5313074 | 16031 | QRBI01000102.1 | 9438414 | 5860533 | 5328456 | 532078 | SUPER_Z | 90132487 | 18562787 | 19095024 | 532237 | 99.89013 | 99.89013 | Minus | | | |
| **g43** | 320379at7742 | Complete | 7955301 | 7964874 | 9573 | QRBI01000119.1 | 13528601 | 7928085 | 7948338 | 20254 | scaffold_474_arr | 20248 | 1 | 20248 | 20247 | 98.72212 | 98.72212 | Plus | 7948338 | 7976236 | Query gene sequence not aligned |
| **g43** | 320379at7742 | Complete | 7955301 | 7964874 | 9573 | QRBI01000119.1 | 13528601 | 7976236 | 8000156 | 23921 | SUPER_Z | 90132487 | 7489159 | 7513079 | 23920 | 100 | 0.020902 | Plus | | | |
| **g44** | 334667at7742 | Complete | 116693 | 136355 | 19662 | QRBI01000103.1 | 6171425 | 119895 | 122169 | 2275 | SUPER_Z | 90132487 | 24335965 | 24338239 | 2274 | 99.78031 | 1.010545 | Plus | 116693 | 141973 | Query gene sequence not aligned |
| **g44** | 334667at7742 | Complete | 116693 | 136355 | 19662 | QRBI01000103.1 | 6171425 | 141973 | 373384 | 231412 | SUPER_Z | 90132487 | 597447 | 828833 | 231386 | 99.82511 | 99.82511 | Plus | | | |
| **g45** | 341187at7742 | Complete | 273 | 17157 | 16884 | QRBI01000205.1 | 609271 | 73850 | 121073 | 47224 | SUPER_5 | 73257097 | 33770927 | 33818150 | 47223 | 100 | 0.010588 | Plus | 0 | 73850 | Query gene sequence not aligned |
| **g46** | 349414at7742 | Complete | 2760875 | 2769103 | 8228 | QRBI01000240.1 | 3274615 | 2755799 | 2717001 | 38799 | SUPER_Z | 90132487 | 24335986 | 24374784 | 38798 | 100 | 0.015464 | Minus | 2755799 | 2776954 | Query gene sequence not aligned |
| **g46** | 349414at7742 | Complete | 2760875 | 2769103 | 8228 | QRBI01000240.1 | 3274615 | 2776954 | 2811979 | 35026 | SUPER_28 | 5297670 | 1 | 35032 | 35031 | 99.81174 | 99.81174 | Plus | | | |
| **g47** | 357329at7742 | Complete | 24804523 | 24814779 | 10256 | QRBI01000099.1 | 24819762 | 24804610 | 24553038 | 251573 | SUPER_3 | 1.17E+08 | 91246103 | 91497641 | 251538 | 99.56194 | 99.56194 | Minus | 24804610 | 24819762 | Query gene sequence not aligned |
| **g48** | 368535at7742 | Complete | 945590 | 950066 | 4476 | QRBI01000189.1 | 1056100 | 932633 | 943048 | 10416 | SUPER_27_unlo | 65965 | 55658 | 65965 | 10307 | 98.27388 | 98.27388 | Plus | 943048 | 967982 | Query gene sequence not aligned |
| **g48** | 368535at7742 | Complete | 945590 | 950066 | 4476 | QRBI01000189.1 | 1056100 | 975708 | 967982 | 7727 | SUPER_27 | 5236451 | 859320 | 867064 | 7744 | 98.72324 | 98.72324 | Minus | | | |
| **g49** | 368798at7742 | Complete | 51393 | 54218 | 2825 | QRBI01000260.1 | 275178 | 45533 | 29442 | 16092 | scaffold_201_arr | 67835 | 15798 | 31859 | 16061 | 98.5275 | 98.5275 | Minus | 45533 | 187975 | Query gene sequence not aligned |
| **g49** | 368798at7742 | Complete | 51393 | 54218 | 2825 | QRBI01000260.1 | 275178 | 187975 | 194170 | 6196 | SUPER_W | 31704074 | 29807904 | 29814099 | 6195 | 100 | 0.096837 | Plus | | | |
| **g50** | 378120at7742 | Complete | 810966 | 813762 | 2796 | QRBI01000194.1 | 1532577 | 809235 | 810349 | 1115 | scaffold_460_arr | 22113 | 21003 | 22113 | 1110 | 98.30206 | 98.30206 | Plus | 810349 | 826625 | Query gene sequence not aligned |
| **g50** | 378120at7742 | Complete | 810966 | 813762 | 2796 | QRBI01000194.1 | 1532577 | 826625 | 913675 | 87051 | SUPER_Z | 90132487 | 3242250 | 3329300 | 87050 | 100 | 0.005744 | Plus | | | |
| **g51** | 378949at7742 | Complete | 611589 | 613898 | 2309 | QRBI01000098.1 | 16813092 | 610964 | 286865 | 324100 | SUPER_15 | 16541138 | 15933712 | 16257853 | 324141 | 99.49706 | 99.49706 | Minus | 610964 | 645811 | Query gene sequence not aligned |
| **g51** | 378949at7742 | Complete | 611589 | 613898 | 2309 | QRBI01000098.1 | 16813092 | 671402 | 645811 | 25592 | SUPER_15 | 16541138 | 15873160 | 15898838 | 25678 | 99.23686 | 99.23686 | Minus | | | |
| **g52** | 381001at7742 | Complete | 2629493 | 2633917 | 4424 | QRBI01000240.1 | 3274615 | 2619038 | 2528327 | 90712 | SUPER_28 | 5297670 | 143151 | 233922 | 90771 | 99.77982 | 99.77982 | Minus | 2619038 | 2636565 | Query gene sequence not aligned |
| **g52** | 381001at7742 | Complete | 2629493 | 2633917 | 4424 | QRBI01000240.1 | 3274615 | 2636565 | 2653565 | 17001 | SUPER_Z | 90132487 | 48576984 | 48593984 | 17000 | 100 | 0.02941 | Plus | | | |
| **g53** | 381971at7742 | Complete | 83384 | 89355 | 5971 | QRBI01000340.1 | 373944 | 103234 | 151914 | 48681 | SUPER_28 | 5297670 | 4120146 | 4168826 | 48680 | 100 | 0.008217 | Plus | 0 | 103234 | Query gene sequence not aligned |
| **g54** | 395792at7742 | Complete | 29639863 | 29640426 | 563 | QRBI01000123.1 | 55481500 | 29638501 | 29620823 | 17679 | scaffold_430_arr | 24449 | 1 | 17671 | 17670 | 99.42383 | 99.42383 | Minus | 29638501 | 29662808 | Query gene sequence not aligned |
| **g54** | 395792at7742 | Complete | 29639863 | 29640426 | 563 | QRBI01000123.1 | 55481500 | 29689239 | 29662808 | 26432 | SUPER_1 | 1.56E+08 | 1.26E+08 | 1.26E+08 | 26453 | 99.74313 | 99.74313 | Minus | | | |
| **g55** | 407723at7742 | Complete | 2461 | 10859 | 8398 | QRBI01000043.1 | 41151 | - | - | - | - | - | - | - | | - | - | - | 0 | 41151 | Query gene sequence not aligned |
| **g56** | 410670at7742 | Complete | 982835 | 987434 | 4599 | QRBI01000153.1 | 9804896 | 947182 | 953087 | 5906 | SUPER_21 | 9617204 | 924667 | 930559 | 5892 | 98.6991 | 98.6991 | Plus | 953087 | 988110 | Query gene sequence not aligned |
| **g56** | 410670at7742 | Complete | 982835 | 987434 | 4599 | QRBI01000153.1 | 9804896 | 988110 | 1077662 | 89553 | SUPER_21 | 9617204 | 965483 | 1055073 | 89590 | 99.01981 | 99.01981 | Plus | | | |
| **g57** | 420085at7742 | Complete | 424857 | 432729 | 7872 | QRBI01000268.1 | 1215624 | 236642 | 244573 | 7932 | SUPER_26 | 5553549 | 89949 | 97869 | 7920 | 98.9679 | 98.9679 | Plus | 244573 | 435978 | Query gene sequence not aligned |
| **g57** | 420085at7742 | Complete | 424857 | 432729 | 7872 | QRBI01000268.1 | 1215624 | 435978 | 471274 | 35297 | scaffold_340_arr | 35322 | 1 | 35322 | 35321 | 99.54217 | 99.54217 | Plus | | | |
| **g58** | 339203at7742 | Duplicated | 430368 | 433482 | 3114 | QRBI01000248.1 | 455452 | 348427 | 350644 | 2218 | SUPER_30 | 1648998 | 1599269 | 1601482 | 2213 | 98.24245 | 98.24245 | Plus | 350644 | 455452 | Query gene sequence not aligned |
| **g59** | 136144at7742 | Fragmented | 199035 | 207813 | 8778 | QRBI01000246.1 | 1116789 | 196445 | 194076 | 2370 | SUPER_29 | 2102120 | 2065049 | 2067428 | 2379 | 98.02935 | 98.02935 | Minus | 196445 | 331656 | Query gene sequence not aligned |

| | | | | | | | | | | | | | | | | | | | | | |
|---|---|---|---|---|---|---|---|---|---|---|---|---|---|---|---|---|---|---|---|---|---|
| **g59** | 136144at7742 | Fragmented | 199035 | 207813 | 8778 | QRBI01000246.1 | 1116789 | 333117 | 331656 | 1462 | SUPER_29 | 2102120 | 1954588 | 1956053 | 1465 | 98.5685 | 98.5685 | Minus | | | |
| **g60** | 281245at7742 | Fragmented | 57802 | 67653 | 9851 | QRBI01000012.1 | 72275 | - | - | - | - | - | - | - | | - | - | - | 0 | 72275 | Query gene sequence not aligned |
| **g61** | 330367at7742 | Fragmented | 447908 | 452949 | 5041 | QRBI01000258.1 | 596393 | 403766 | 370397 | 33370 | SUPER_1 | 1.56E+08 | 896895 | 930264 | 33369 | 100 | 0.011987 | Minus | 403766 | 469731 | Query gene sequence not aligned |
| **g61** | 330367at7742 | Fragmented | 447908 | 452949 | 5041 | QRBI01000258.1 | 596393 | 469731 | 479725 | 9995 | scaffold_590_arr | 1.00E+04 | 1 | 10049 | 10048 | 98.85754 | 98.85754 | Plus | | | |

**b)**

| Gene ID | BUSCO ID | Gene status in t | Gene start | Gene end | Gene sequence | Query seq. ID | Q. scaffold leng | Q. start alignme | Q. end alignmer | Q. alignment ler | Reference seq. | R. chr length | R. start alignme | R. end alignmer | R. alignment ler | % identity | % similarity | strand of the qu | Start missing in | End missing in t | Notes |
|---|---|---|---|---|---|---|---|---|---|---|---|---|---|---|---|---|---|---|---|---|---|
| **g1** | 263171at7742 | Complete | 47326 | 51319 | 3993 | 001562F_arrow_arrow | 53617 | 43851 | 35506 | 8346 | SUPER_26 | 5553549 | 902419 | 910748 | 8329 | 98.08978 | 98.08978 | Minus | 43851 | 53617 | Query gene sequence not aligned |
| **g2** | 25839at7742 | Complete | 60740 | 75022 | 14282 | 000069F_009_arrow_arrow | 98347 | 1 | 90671 | 90671 | SUPER_3 | 1.17E+08 | 16120344 | 16211115 | 90771 | 99.18372 | 99.18372 | Plus | - | - | Query gene sequence aligned entirely |
| **g3** | 47275at7742 | Complete | 33176 | 59491 | 26315 | 000069F_009_arrow_arrow | 98347 | 1 | 90671 | 90671 | SUPER_3 | 1.17E+08 | 16120344 | 16211115 | 90771 | 99.18372 | 99.18372 | Plus | - | - | Query gene sequence aligned entirely |
| **g4** | 75522at7742 | Complete | 580500 | 605941 | 25441 | 000268F_001_arrow_arrow | 632736 | 528407 | 618025 | 89619 | SUPER_8 | 36085389 | 35980713 | 36070454 | 89741 | 99.26682 | 99.26682 | Plus | - | - | Query gene sequence aligned entirely |
| **g5** | 106313at7742 | Complete | 158017 | 160086 | 2069 | 000280F_006_arrow_arrow | 652044 | 1 | 276059 | 276059 | SUPER_11 | 21491857 | 5872 | 282023 | 276151 | 99.53653 | 99.53653 | Plus | - | - | Query gene sequence aligned entirely |
| **g6** | 274411at7742 | Complete | 1144760 | 1160739 | 15979 | 000084F_017_arrow_arrow | 2768865 | 1186708 | 1092014 | 94695 | SUPER_4 | 76187387 | 19574515 | 19669115 | 94600 | 98.92482 | 98.92482 | Minus | - | - | Query gene sequence aligned entirely |
| **g13** | 265085at7742 | Complete | 726701 | 812685 | 85984 | 000160F_001_arrow_arrow | 818199 | 797469 | 631592 | 165878 | SUPER_3 | 1.17E+08 | 82974228 | 83140003 | 165775 | 98.92596 | 98.92596 | Minus | - | - | Query gene sequence aligned entirely |
| **g13** | 265085at7742 | Complete | 726701 | 812685 | 85984 | 000160F_001_arrow_arrow | 818199 | 818199 | 777451 | 40749 | SUPER_3 | 1.17E+08 | 82953519 | 82994239 | 40720 | 99.66901 | 99.66901 | Minus | - | - | Query gene sequence aligned entirely |
| **g14** | 291147at7742 | Complete | 777922 | 819309 | 41387 | 000235F_006_arrow_arrow | 1162873 | 859601 | 761484 | 98118 | SUPER_6 | 63258489 | 11773913 | 11872025 | 98112 | 98.83692 | 98.83692 | Minus | - | - | Query gene sequence aligned entirely |
| **g15** | 259004at7742 | Complete | 2534502 | 2545088 | 10586 | 000038F_014_arrow_arrow | 3123829 | 2615445 | 2502316 | 113130 | SUPER_2 | 1.19E+08 | 5715751 | 5828707 | 112956 | 98.50085 | 98.50085 | Minus | - | - | Query gene sequence aligned entirely |
| **g16** | 131366at7742 | Complete | 685387 | 718596 | 33209 | 000180F_002_arrow_arrow | 1444896 | 356393 | 760311 | 403919 | SUPER_3 | 1.17E+08 | 34800136 | 35203971 | 403835 | 98.65837 | 98.65837 | Plus | - | - | Query gene sequence aligned entirely |
| **g17** | 437433at7742 | Complete | 351016 | 353103 | 2087 | 000031F_006_arrow_arrow | 2595384 | 263634 | 498125 | 234492 | SUPER_12 | 20890524 | 3021194 | 3255434 | 234240 | 98.33811 | 98.33811 | Plus | - | - | Query gene sequence aligned entirely |
| **g18** | 333504at7742 | Fragmented | 310834 | 322473 | 11639 | 000018F_013_arrow_arrow | 447620 | 305885 | 363359 | 57475 | SUPER_6 | 6.33E+07 | 20814343 | 20871736 | 57393 | 99.22629 | 99.22629 | Plus | - | - | Query gene sequence aligned entirely |
| **g20** | 360487at7742 | Complete | 133501 | 141103 | 7602 | 000989F_arrow_arrow | 141510 | 98227 | 101379 | 3153 | SUPER_28 | 5.30E+06 | 4095755 | 4098891 | 3136 | 98.00317 | 98.00317 | Plus | 101379 | 141510 | Query gene sequence not aligned |
| **g21** | 363461at7742 | Complete | 13913 | 17843 | 3930 | 000910F_arrow_arrow | 171119 | 8213 | 1 | 8213 | SUPER_5 | 7.33E+07 | 18336086 | 18344319 | 8233 | 98.38553 | 98.38553 | Minus | 8213 | 74224 | Query gene sequence not aligned |
| **g21** | 363461at7742 | Complete | 13913 | 17843 | 3930 | 000910F_arrow_arrow | 171119 | 94949 | 74224 | 20726 | SUPER_6 | 6.33E+07 | 1930357 | 1951094 | 20737 | 98.46457 | 98.46457 | Minus | | | |
| **g22** | 419118at7742 | Complete | 71973 | 74747 | 2774 | 001128F_arrow_arrow | 81027 | 66526 | 62565 | 3962 | SUPER_22 | 7.51E+06 | 5682203 | 5686160 | 3957 | 99.5458 | 99.5458 | Minus | 66526 | 81027 | Query gene sequence not aligned |
| **g23** | 293860at7742 | Complete | 64238 | 69827 | 5589 | 001400F\|arrow\|arrow | 70135 | - | - | - | - | - | - | - | - | - | - | - | 0 | 70135 | Query scaffold sequence not aligned |
| **g24** | 235274at7742 | Complete | 51595 | 55923 | 4328 | 001510F_arrow_arrow | 59052 | 48862 | 47850 | 1013 | scaffold_121_arr | 129379 | 48071 | 49084 | 1013 | 98.13176 | 98.13176 | Minus | 48862 | 59052 | Query gene sequence not aligned |
| **g25** | 264645at7742 | Complete | 91200 | 97204 | 6004 | 001189F_arrow_arrow | 97204 | 38389 | 36996 | 1394 | SUPER_26 | 5553549 | 41015 | 42408 | 1393 | 99.06743 | 99.06743 | Minus | 38389 | 97204 | Query gene sequence not aligned |
| **g26** | 407572at7742 | Complete | 7205 | 8567 | 1362 | 001439F_001\|arrow\|arrow | 61945 | - | - | - | - | - | - | - | - | - | - | - | 0 | 61945 | Query scaffold sequence not aligned |
| **g27** | 218284at7742 | Complete | 71975 | 75127 | 3152 | 001299F_arrow_arrow | 81976 | 71534 | 64955 | 6580 | SUPER_11 | 2.15E+07 | 1508826 | 1515424 | 6598 | 98.41174 | 98.41174 | Minus | 71534 | 81976 | Query gene sequence not aligned |
| **g28** | 144578at7742 | Fragmented | 1 | 5173 | 5172 | 000162F_007_arrow_arrow | 585565 | 8036 | 310996 | 302961 | SUPER_5 | 73257097 | 65575466 | 65878510 | 303044 | 99.14637 | 99.14637 | Plus | 0 | 8036 | Query gene sequence not aligned |

**c)**

| Gene ID | BUSCO ID | Gene status in p | Gene start | Gene end | Gene sequence | Query seq. ID | Q. scaffold leng | Q. start alignme | Q. end alignmer | Q. alignment ler | Reference seq. | R. chr length | R. start alignme | R. end alignmer | R. alignment ler | % identity | % similarity | strand of the qu | Start missing in | End missing in t | Notes |
|---|---|---|---|---|---|---|---|---|---|---|---|---|---|---|---|---|---|---|---|---|---|
| **g19** | 149121at7742 | Complete | 618 | 24009 | 23391 | 002029F_arrow | 27423 | 27423 | 1 | 27424 | scaffold_397_arr | 27415 | 1 | 27415 | 27415 | 99.82867 | 99.82867 | Minus | - | - | Query gene sequence aligned entirely |

**Table S40 - Recovered sequences from Chelidonia and the alternate assembly.** The tables reports the coordinates of the recovered sequences that contained the genes from a) Chelidonia; b) alternate assembly.

a)

| Gene ID | BUSCO ID | Chelidonia scaffold ID | Start | End | Sequence name in bHirRus1 after BUSCO gene recover | Notes |
|---|---|---|---|---|---|---|
| **g28** | 144578at7742 | QRBI01000105.1 | 1925492 | 1967238 | SUPER_5_unloc_BUSCO_144578at7742 | g28 recovered from Chelidonia and not from the alternate because it was found complete in the former and fragmented in the latter |
| **g29** | 63198at7742 | QRBI01000105.1 | 66708817 | 66802864 | unplaced_BUSCO_63198at7742 | |
| **g30** | 68766at7742 | QRBI01000249.1 | 108525 | 166550 | unplaced_BUSCO_68766at7742 | g30 recovery coordinates (108525-200600, see ST13) trimmed to avoid adding duplicates |
| **g31** | 122064at7742 | QRBI01000152.1 | 1111216 | 1128958 | SUPER_6_unloc_BUSCO_122064at7742 | |
| **g32** | 126329at7742 | QRBI01000248.1 | 189516 | 216835 | SUPER_30_unloc_BUSCO_126329at7742 | |
| **g33** | 162664at7742 | QRBI01000106.1 | 63009785 | 63027227 | SUPER_3_unloc_BUSCO_162664at7742 | |
| **g34** | 191412at7742 | QRBI01000120.1 | 91892976 | 91936698 | SUPER_2_unloc_BUSCO_191412at7742 | |
| **g35** | 208931at7742 | QRBI01000099.1 | 2725996 | 2758525 | SUPER_3_unloc_BUSCO_208931at7742 | |
| **g36, g27** | 236895at7742, 218284at7742 | QRBI01000116.1 | 21323815 | 21360556 | SUPER_11_unloc_BUSCO_236895at7742_218284at7742 | g27 recovered from Chelidonia and not from the alternate to avoid adding duplicates. When recovering g36 from Chelidonia, BUSCO found g27 in the same sequence. |
| **g38** | 267345at7742 | QRBI01000249.1 | 349281 | 404975 | unplaced_BUSCO_267345at7742 | g38 recovery coordinates (334945-499425, see ST13) trimmed to avoid adding duplicates |
| **g39, g37** | 293313at7742, 266219at7742 | QRBI01000199.1 | 334638 | 364196 | SUPER_3_unloc_BUSCO_293313at7742_266219at7742 | g37 and 29 recovered together because are in the same Chelidonia scaffold and the same coordinates didn't align (see ST13) |
| **g40, g42** | 309129at7742, 310457at7742 | QRBI01000102.1 | 5296419 | 5320500 | unplaced_BUSCO_309129at7742_310457at7742 | g40 and 42 recovered together because are in the same Chelidonia scaffold and the same coordinates didn't align (see ST13). Recovery coordinates (5296419-5328456, see ST12) trimmed to avoid adding to avoid adding duplicates |
| **g41** | 309958at7742 | QRBI01000123.1 | 935724 | 987011 | unplaced_BUSCO_309958at7742 | |
| **g43** | 320379at7742 | QRBI01000119.1 | 7948338 | 7976236 | unplaced_BUSCO_320379at7742 | |
| **g44** | 334667at7742 | QRBI01000103.1 | 116693 | 141973 | SUPER_Z_unloc_BUSCO_334667at7742 | |
| **g45** | 341187at7742 | QRBI01000205.1 | 0 | 73850 | unplaced_BUSCO_341187at7742 | |
| **g46** | 349414at7742 | QRBI01000240.1 | 2755799 | 2776954 | unplaced_BUSCO_349414at7742 | |
| **g47** | 357329at7742 | QRBI01000099.1 | 24804610 | 24819762 | unplaced_BUSCO_357329at7742 | |
| **g48** | 368535at7742 | QRBI01000189.1 | 943048 | 967982 | SUPER_27_unloc_BUSCO_368535at7742 | |
| **g49** | 368798at7742 | QRBI01000260.1 | 45533 | 187975 | unplaced_BUSCO_368798at7742 | |
| **g50** | 378120at7742 | QRBI01000194.1 | 810349 | 826625 | unplaced_BUSCO_378120at7742 | |
| **g51** | 378949at7742 | QRBI01000098.1 | 610964 | 645811 | SUPER_15_unloc_BUSCO_378949at7742 | |
| **g52** | 381001at7742 | QRBI01000240.1 | 2619038 | 2636565 | unplaced_BUSCO_381001at7742 | |
| **g53** | 381971at7742 | QRBI01000340.1 | 0 | 103234 | unplaced_BUSCO_381971at7742 | |
| **g54** | 395792at7742 | QRBI01000123.1 | 29638501 | 29662808 | unplaced_BUSCO_395792at7742 | |
| **g55** | 407723at7742 | QRBI01000043.1 | 0 | 41151 | unplaced_BUSCO_407723at7742 | |
| **g56** | 410670at7742 | QRBI01000153.1 | 953087 | 988110 | SUPER_21_unloc_BUSCO_410670at7742 | |
| **g57** | 420085at7742 | QRBI01000268.1 | 384075 | 435978 | unplaced_BUSCO_420085at7742 | g38 recovery coordinates (244573-435978, see ST13) trimmed to avoid adding genes that were already found by BUSCO |
| **g58** | 339203at7742 | QRBI01000248.1 | 350644 | 455452 | unplaced_BUSCO_339203at7742 | |
| **g59** | 136144at7742 | QRBI01000246.1 | 196445 | 331656 | SUPER_29_unloc_BUSCO_136144at7742 | |
| **g60** | 281245at7742 | QRBI01000012.1 | 0 | 72275 | unplaced_BUSCO_281245at7742 | |
| **g61** | 330367at7742 | QRBI01000258.1 | 403766 | 469731 | unplaced_BUSCO_330367at7742 | |

b)

| Gene ID | BUSCO ID | Alternate scaffold ID | Start | End | Sequence name in fasta | Notes |
|---|---|---|---|---|---|---|
| **g20** | 360487at7742 | 000989F\|arrow\|arrow | 101379 | 141510 | unplaced_BUSCO_360487at7742 | |
| **g21** | 363461at7742 | 000910F\|arrow\|arrow | 8213 | 74224 | unplaced_BUSCO_363461at7742 | |
| **g22** | 419118at7742 | 001128F\|arrow\|arrow | 66526 | 81027 | unplaced_BUSCO_419118at7742 | |
| **g23** | 293860at7742 | 001400F\|arrow\|arrow | 0 | 70135 | unplaced_BUSCO_293860at7742 | Recovery of the whole scaffold |
| **g24** | 235274at7742 | 001510F\|arrow\|arrow | 48862 | 59052 | unplaced_BUSCO_235274at7742 | |
| **g25** | 264645at7742 | 001189F\|arrow\|arrow | 38389 | 97204 | unplaced_BUSCO_264645at7742 | |
| **g26** | 407572at7742 | 001439F_001\|arrow\|arrow | 0 | 61945 | unplaced_BUSCO_407572at7742 | Recovery of the whole scaffold |

**Table S41 - BLAST search of the missing genes in bHirRus1 after gene recovery.** The table reports the output of the BLAST search of the 139 genes missing in bHirRus1, that were found in the other assemblies/intermediates.
Sequences were retrieved from the corresponding assembly/intermediates and blasted on bHirRus1 after gene recovery.

a)

| Gene ID | BUSCO gene ID | Chelidonia scaf | Gene start in th | Gene end in the | Gene length in t | bHirRus1 chr | % identity | alignment lengt | mismatches | gap opens | q. start | q. end | s. start | s. end | evalue | bit score |
|---|---|---|---|---|---|---|---|---|---|---|---|---|---|---|---|---|
| **g1** | 263171at7742 | QRBI01000142.1 | 4878372 | 4901008 | 22636 | SUPER_26 | 97.057 | 8935 | 133 | 61 | 6756 | 15623 | 9.11E+05 | 9.02E+05 | 0 | 14923 |
| **g1** | 263171at7742 | QRBI01000142.1 | 4878372 | 4901008 | 22636 | SUPER_26 | 93.81 | 4152 | 111 | 35 | 2828 | 6913 | 914585 | 910514 | 0 | 6109 |
| **g1** | 263171at7742 | QRBI01000142.1 | 4878372 | 4901008 | 22636 | SUPER_26 | 95.379 | 3571 | 108 | 31 | 1 | 3559 | 917655 | 914130 | 0 | 5627 |
| **g2** | 25839at7742 | QRBI01000208.1 | 3403968 | 3418251 | 14283 | SUPER_3 | 99.484 | 14330 | 5 | 65 | 1 | 14283 | 16181132 | 16195439 | 0 | 25989 |
| **g3** | 47275at7742 | QRBI01000208.1 | 3375445 | 3402719 | 27274 | SUPER_3 | 98.74 | 27462 | 60 | 259 | 1 | 27274 | 16152515 | 16179878 | 0 | 48532 |
| **g4** | 75522at7742 | QRBI01000223.1 | 87332 | 112771 | 25439 | SUPER_8 | 99.076 | 25546 | 54 | 166 | 1 | 25439 | 36058293 | 36032823 | 0 | 45700 |
| **g5** | 106313at7742 | QRBI01000116.1 | 22666816 | 22668884 | 2068 | SUPER_11 | 98.662 | 2092 | 1 | 21 | 1 | 2068 | 165887 | 163799 | 0 | 3683 |
| **g6** | 274411at7742 | QRBI01000112.1 | 21081776 | 21097751 | 15975 | SUPER_4 | 99.497 | 8356 | 27 | 8 | 7629 | 15975 | 19608817 | 19600468 | 0 | 15184 |
| **g6** | 274411at7742 | QRBI01000112.1 | 21081776 | 21097751 | 15975 | SUPER_4 | 99.444 | 7559 | 24 | 9 | 96 | 7640 | 19616307 | 19608753 | 0 | 13710 |
| **g7** | 68501at7742 | QRBI01000107.1 | 11758926 | 11818081 | 59155 | SUPER_2 | 99.412 | 30270 | 46 | 82 | 1 | 30240 | 11417992 | 11387825 | 0 | 54791 |
| **g7** | 68501at7742 | QRBI01000107.1 | 11758926 | 11818081 | 59155 | SUPER_2 | 99.102 | 28950 | 153 | 49 | 30261 | 59155 | 11387851 | 11358954 | 0 | 51921 |
| **g8** | 58506at7742 | QRBI01000270.1 | 31205 | 63502 | 32297 | SUPER_22 | 99.567 | 32343 | 30 | 86 | 1 | 32297 | 56641 | 24363 | 0 | 58850 |
| **g9** | 59090at7742 | QRBI01000123.1 | 53896618 | 53988865 | 92247 | SUPER_1 | 99.568 | 34060 | 75 | 34 | 1 | 34020 | 1.01E+08 | 1.01E+08 | 0 | 62017 |
| **g10** | 205146at7742 | QRBI01000195.1 | 865563 | 895356 | 29793 | SUPER_Z | 99.872 | 25840 | 3 | 27 | 2 | 25820 | 65238702 | 65212872 | 0 | 47508 |
| **g11** | 210721at7742 | QRBI01000102.1 | 5320500 | 5352337 | 31837 | SUPER_Z | 99.854 | 23908 | 0 | 34 | 7956 | 31837 | 19095024 | 19071126 | 0 | 43923 |
| **g12** | 283910at7742 | QRBI01000169.1 | 1627623 | 1686060 | 58437 | SUPER_Z | 99.861 | 28732 | 0 | 37 | 1 | 28697 | 63012691 | 62983965 | 0 | 52800 |
| **g20** | 360487at7742 | QRBI01000240.1 | 16972 | 21010 | 4038 | unplaced_BUSCO_360487at7742 | 98.349 | 2181 | 18 | 5 | 1872 | 4038 | 34298 | 32122 | 0 | 3812 |
| **g20** | 360487at7742 | QRBI01000240.1 | 16972 | 21010 | 4038 | unplaced_BUSCO_360487at7742 | 97.8 | 2045 | 19 | 6 | 1 | 2022 | 36357 | 34316 | 0 | 3504 |
| **g20** | 360487at7742 | QRBI01000240.1 | 16972 | 21010 | 4038 | unplaced_BUSCO_360487at7742 | 97.8 | 2045 | 19 | 6 | 1 | 2022 | 36357 | 34316 | 0 | 3504 |
| **g21** | 363461at7742 | QRBI01000152.1 | 2262551 | 2266468 | 3917 | unplaced_BUSCO_363461at7742 | 98.805 | 3934 | 26 | 8 | 1 | 3917 | 9629 | 5700 | 0 | 6985 |
| **g22** | 419118at7742 | QRBI01000235.1 | 1936562 | 1939520 | 2958 | unplaced_BUSCO_419118at7742 | 99.771 | 2616 | 4 | 2 | 1 | 2615 | 5448 | 8062 | 0 | 4796 |
| **g24** | 235274at7742 | QRBI01000249.1 | 444782 | 456398 | 11616 | unplaced_BUSCO_235274at7742 | 99.92 | 7459 | 0 | 6 | 1 | 7455 | 2734 | 10190 | 0 | 13736 |
| **g25** | 264645at7742 | QRBI01000268.1 | 129061 | 135873 | 6812 | unplaced_BUSCO_264645at7742 | 98.028 | 5630 | 28 | 24 | 1021 | 6583 | 58815 | 53202 | 0 | 9705 |
| **g26** | 407572at7742 | QRBI01000205.1 | 146406 | 152708 | 6302 | unplaced_BUSCO_407572at7742 | 98.315 | 6349 | 17 | 68 | 1 | 6302 | 4581 | 10886 | 0 | 11049 |
| **g27** | 218284at7742 | QRBI01000116.1 | 21324256 | 21327407 | 3151 | SUPER_11_unloc_BUSCO_236895at7742_218284at | 100 | 3151 | 0 | 0 | 1 | 3151 | 442 | 3592 | 0 | 5819 |
| **g28** | 144578at7742 | QRBI01000105.1 | 1928349 | 1934461 | 6112 | SUPER_5_unloc_BUSCO_144578at7742 | 100 | 6112 | 0 | 0 | 1 | 6112 | 2858 | 8969 | 0 | 11287 |
| **g29** | 63198at7742 | QRBI01000105.1 | 66721255 | 66747819 | 26564 | unplaced_BUSCO_63198at7742 | 100 | 26564 | 0 | 0 | 1 | 26564 | 12439 | 39002 | 0 | 49055 |
| **g30** | 68766at7742 | QRBI01000249.1 | 129633 | 133452 | 3819 | unplaced_BUSCO_68766at7742 | 100 | 3819 | 0 | 0 | 1 | 3819 | 21109 | 24927 | 0 | 7053 |
| **g31** | 122064at7742 | QRBI01000152.1 | 1117805 | 1128803 | 10998 | SUPER_6_unloc_BUSCO_122064at7742 | 100 | 10998 | 0 | 0 | 1 | 10998 | 6590 | 17587 | 0 | 20310 |
| **g32** | 126329at7742 | QRBI01000248.1 | 204174 | 212108 | 7934 | SUPER_30_unloc_BUSCO_126329at7742 | 100 | 7934 | 0 | 0 | 1 | 7934 | 14659 | 22592 | 0 | 14652 |
| **g33** | 162664at7742 | QRBI01000106.1 | 63011072 | 63018103 | 7031 | SUPER_3_unloc_BUSCO_162664at7742 | 100 | 7031 | 0 | 0 | 1 | 7031 | 1288 | 8318 | 0 | 12984 |
| **g34** | 191412at7742 | QRBI01000120.1 | 91897656 | 91913919 | 16263 | SUPER_2_unloc_BUSCO_191412at7742 | 100 | 16263 | 0 | 0 | 1 | 16263 | 4681 | 20943 | 0 | 30033 |
| **g35** | 208931at7742 | QRBI01000099.1 | 2727389 | 2757173 | 29784 | SUPER_3_unloc_BUSCO_208931at7742 | 100 | 29784 | 0 | 0 | 1 | 29784 | 1394 | 31177 | 0 | 55001 |
| **g36** | 236895at7742 | QRBI01000116.1 | 21355634 | 21359393 | 3759 | SUPER_11_unloc_BUSCO_236895at7742_218284at | 100 | 3759 | 0 | 0 | 1 | 3759 | 31820 | 35578 | 0 | 6942 |
| **g37** | 266219at7742 | QRBI01000199.1 | 344736 | 347910 | 3174 | SUPER_3_unloc_BUSCO_293313at7742_266219at7 | 100 | 3174 | 0 | 0 | 1 | 3174 | 10099 | 13272 | 0 | 5862 |
| **g38** | 267345at7742 | QRBI01000249.1 | 353310 | 367170 | 13860 | unplaced_BUSCO_267345at7742 | 100 | 13860 | 0 | 0 | 1 | 13860 | 4030 | 17889 | 0 | 25547 |
| **g39** | 293313at7742 | QRBI01000199.1 | 351238 | 353807 | 2569 | SUPER_3_unloc_BUSCO_293313at7742_266219at7 | 100 | 2569 | 0 | 0 | 1 | 2569 | 16601 | 19169 | 0 | 4745 |
| **g40** | 309129at7742 | QRBI01000102.1 | 5313240 | 5317727 | 4487 | unplaced_BUSCO_309129at7742_310457at7742 | 100 | 4487 | 0 | 0 | 1 | 4487 | 16822 | 21308 | 0 | 8287 |
| **g41** | 309958at7742 | QRBI01000123.1 | 946786 | 949272 | 2486 | unplaced_BUSCO_309958at7742 | 100 | 2486 | 0 | 0 | 1 | 2486 | 11063 | 13548 | 0 | 4591 |
| **g42** | 310457at7742 | QRBI01000102.1 | 5297043 | 5313074 | 16031 | unplaced_BUSCO_309129at7742_310457at7742 | 100 | 16031 | 0 | 0 | 1 | 16031 | 625 | 16655 | 0 | 2.96E+04 |
| **g43** | 320379at7742 | QRBI01000119.1 | 7955301 | 7964874 | 9573 | unplaced_BUSCO_320379at7742 | 100 | 9573 | 0 | 0 | 1 | 9573 | 6964 | 16536 | 0 | 17679 |
| **g44** | 334667at7742 | QRBI01000103.1 | 116693 | 136355 | 19662 | SUPER_Z_unloc_BUSCO_334667at7742 | 100 | 14186 | 0 | 0 | 5477 | 19662 | 5477 | 19662 | 0 | 26197 |
| **g44** | 334667at7742 | QRBI01000103.1 | 116693 | 136355 | 19662 | SUPER_Z_unloc_BUSCO_334667at7742 | 100 | 3228 | 0 | 0 | 1 | 3228 | 1 | 3228 | 0 | 5962 |
| **g45** | 341187at7742 | QRBI01000205.1 | 273 | 17157 | 16884 | unplaced_BUSCO_341187at7742 | 100 | 16884 | 0 | 0 | 1 | 16884 | 274 | 17157 | 0 | 31179 |
| **g46** | 349414at7742 | QRBI01000240.1 | 2760875 | 2769103 | 8228 | unplaced_BUSCO_349414at7742 | 100 | 8228 | 0 | 0 | 1 | 8228 | 5077 | 13304 | 0 | 15195 |
| **g47** | 357329at7742 | QRBI01000099.1 | 24804523 | 24814779 | 10256 | unplaced_BUSCO_357329at7742 | 100 | 10169 | 0 | 0 | 88 | 10256 | 1 | 10169 | 0 | 18779 |
| **g48** | 368535at7742 | QRBI01000189.1 | 945590 | 950066 | 4476 | SUPER_27_unloc_BUSCO_368535at7742 | 100 | 4476 | 0 | 0 | 1 | 4476 | 2543 | 7018 | 0 | 8266 |
| **g49** | 368798at7742 | QRBI01000260.1 | 51393 | 54218 | 2825 | unplaced_BUSCO_368798at7742 | 100 | 2825 | 0 | 0 | 1 | 2825 | 5861 | 8685 | 0 | 5217 |
| **g50** | 378120at7742 | QRBI01000194.1 | 810966 | 813762 | 2796 | unplaced_BUSCO_378120at7742 | 100 | 2796 | 0 | 0 | 1 | 2796 | 618 | 3413 | 0 | 5164 |
| **g51** | 378949at7742 | QRBI01000098.1 | 611589 | 613898 | 2309 | SUPER_15_unloc_BUSCO_378949at7742 | 100 | 2309 | 0 | 0 | 1 | 2309 | 626 | 2934 | 0 | 4265 |
| **g52** | 381001at7742 | QRBI01000240.1 | 2629493 | 2633917 | 4424 | unplaced_BUSCO_381001at7742 | 100 | 4424 | 0 | 0 | 1 | 4424 | 10456 | 14879 | 0 | 8170 |
| **g53** | 381971at7742 | QRBI01000340.1 | 83384 | 89355 | 5971 | unplaced_BUSCO_381971at7742 | 100 | 5971 | 0 | 0 | 1 | 5971 | 83385 | 89355 | 0 | 11027 |
| **g54** | 395792at7742 | QRBI01000123.1 | 29639863 | 29640426 | 563 | unplaced_BUSCO_395792at7742 | 100 | 563 | 0 | 0 | 1 | 563 | 1363 | 1925 | 0 | 1040 |
| **g55** | 407723at7742 | QRBI01000043.1 | 2461 | 10859 | 8398 | unplaced_BUSCO_407723at7742 | 100 | 8398 | 0 | 0 | 1 | 8398 | 2462 | 10859 | 0 | 15509 |
| **g56** | 410670at7742 | QRBI01000153.1 | 982835 | 987434 | 4599 | SUPER_21_unloc_BUSCO_410670at7742 | 100 | 4599 | 0 | 0 | 1 | 4599 | 29749 | 34347 | 0 | 8493 |
| **g57** | 420085at7742 | QRBI01000268.1 | 424857 | 432729 | 7872 | unplaced_BUSCO_420085at7742 | 100 | 7872 | 0 | 0 | 1 | 7872 | 40783 | 48654 | 0 | 14537 |

| | | | | | | | | | | | | | | | | |
|---|---|---|---|---|---|---|---|---|---|---|---|---|---|---|---|---|
| **g58** | 339203at7742 | QRBI01000248.1 | 430368 | 433482 | 3114 | unplaced_BUSCO_339203at7742 | 100 | 3114 | 0 | 0 | 1 | 3114 | 79725 | 82838 | 0 | 5751 |
| **g59** | 136144at7742 | QRBI01000246.1 | 199035 | 207813 | 8778 | SUPER_29_unloc_BUSCO_136144at7742 | 100 | 8778 | 0 | 0 | 1 | 8778 | 2591 | 11368 | 0 | 16211 |
| **g60** | 281245at7742 | QRBI01000012.1 | 57802 | 67653 | 9851 | unplaced_BUSCO_281245at7742 | 100 | 9851 | 0 | 0 | 1 | 9851 | 57803 | 67653 | 0 | 18192 |
| **g61** | 330367at7742 | QRBI01000258.1 | 447908 | 452949 | 5041 | unplaced_BUSCO_330367at7742 | 100 | 5041 | 0 | 0 | 1 | 5041 | 44143 | 49183 | 0 | 9310 |

**b)**

| Gene ID | BUSCO gene ID | Alternate scaffo | Gene start in th | Gene end in the | Gene length in t | bHirRus1 chr | % identity | alignment lengt | mismatches | gap opens | q. start | q. end | s. start | s. end | evalue | bit score |
|---|---|---|---|---|---|---|---|---|---|---|---|---|---|---|---|---|
| **g2** | 25839at7742 | 000069F_009\|arr | 60740 | 75022 | 14282 | SUPER_3 | 99.442 | 14329 | 12 | 65 | 1 | 14282 | 16181132 | 16195439 | 0 | 2.60E+04 |
| **g3** | 47275at7742 | 000069F_009\|arr | 33176 | 59491 | 26315 | SUPER_3 | 98.683 | 26500 | 68 | 255 | 1 | 26315 | 16153475 | 16179878 | 0 | 46743 |
| **g4** | 75522at7742 | 000268F_001\|arr | 580500 | 605941 | 25441 | SUPER_8 | 99.068 | 25547 | 56 | 166 | 1 | 25441 | 36032824 | 36058294 | 0 | 45690 |
| **g5** | 106313at7742 | 000280F_006\|arr | 158017 | 160086 | 2069 | SUPER_11 | 98.709 | 2092 | 1 | 20 | 1 | 2069 | 163800 | 165888 | 0 | 3690 |
| **g6** | 274411at7742 | 000084F_017\|arr | 1144760 | 1160739 | 15979 | SUPER_4 | 99.39 | 8357 | 34 | 8 | 7633 | 15979 | 19608817 | 19600468 | 0 | 15134 |
| **g6** | 274411at7742 | 000084F_017\|arr | 1144760 | 1160739 | 15979 | SUPER_4 | 99.392 | 7561 | 28 | 10 | 96 | 7644 | 19616307 | 19608753 | 0 | 13692 |
| **g13** | 265085at7742 | 000160F_001\|arr | 726701 | 812685 | 85984 | SUPER_3 | 99.312 | 86154 | 271 | 149 | 1 | 85984 | 83045036 | 82959035 | 0 | 1.56E+05 |
| **g14** | 291147at7742 | 000235F_006\|arr | 777922 | 819309 | 41387 | SUPER_6 | 99.015 | 29650 | 197 | 38 | 1 | 29623 | 11855573 | 11825992 | 0 | 53047 |
| **g14** | 291147at7742 | 000235F_006\|arr | 777922 | 819309 | 41387 | SUPER_6 | 99.201 | 10513 | 55 | 16 | 29656 | 40149 | 11825990 | 11815488 | 0 | 18921 |
| **g15** | 259004at7742 | 000038F_014\|arr | 2534502 | 2545088 | 10586 | SUPER_2 | 98.644 | 10621 | 98 | 17 | 1 | 10586 | 5796576 | 5785967 | 0 | 18774 |
| **g16** | 131366at7742 | 000180F_002\|arr | 685387 | 718596 | 33209 | SUPER_3 | 98.822 | 33281 | 286 | 46 | 1 | 33209 | 35129109 | 35162355 | 0 | 59189 |
| **g17** | 437433at7742 | 000031F_006\|arr | 351016 | 353103 | 2087 | SUPER_12 | 98.132 | 2088 | 30 | 3 | 1 | 2087 | 3108423 | 3110502 | 0 | 3631 |
| **g18** | 333504at7742 | 000018F_013\|arr | 310834 | 322473 | 11639 | SUPER_6 | 99.278 | 11640 | 66 | 6 | 1 | 11639 | 20819295 | 20830917 | 0 | 21014 |
| **g20** | 360487at7742 | 000989F\|arrow\|a | 133501 | 141103 | 7602 | unplaced_BUSCO_360487at7742 | 100 | 7602 | 0 | 0 | 1 | 7602 | 32123 | 39724 | 0 | 14039 |
| **g21** | 363461at7742 | 000910F\|arrow\|a | 13913 | 17843 | 3930 | unplaced_BUSCO_363461at7742 | 100 | 3930 | 0 | 0 | 1 | 3930 | 5701 | 9630 | 0 | 7258 |
| **g22** | 419118at7742 | 001128F\|arrow\|a | 71973 | 74747 | 2774 | unplaced_BUSCO_419118at7742 | 100 | 2774 | 0 | 0 | 1 | 2774 | 5448 | 8221 | 0 | 5123 |
| **g23** | 293860at7742 | 001400F\|arrow\|a | 64238 | 69827 | 5589 | unplaced_BUSCO_293860at7742 | 100 | 5589 | 0 | 0 | 1 | 5589 | 64239 | 69827 | 0 | 10322 |
| **g24** | 235274at7742 | 001510F\|arrow\|a | 51595 | 55923 | 4328 | unplaced_BUSCO_235274at7742 | 100 | 4328 | 0 | 0 | 1 | 4328 | 2734 | 7061 | 0 | 7993 |
| **g25** | 264645at7742 | 001189F\|arrow\|a | 91200 | 97204 | 6004 | unplaced_BUSCO_264645at7742 | 100 | 6004 | 0 | 0 | 1 | 6004 | 5.28E+04 | 5.88E+04 | 0 | 11088 |
| **g26** | 407572at7742 | 001439F_001\|arr | 7205 | 8567 | 1362 | unplaced_BUSCO_407572at7742 | 100 | 1362 | 0 | 0 | 1 | 1362 | 7206 | 8567 | 0 | 2516 |
| **g27** | 218284at7742 | 001299F\|arrow\|a | 71975 | 75127 | 3152 | SUPER_11_unloc_BUSCO_236895at7742_218284at | 99.746 | 3155 | 1 | 6 | 1 | 3152 | 442 | 3592 | 0 | 5775 |
| **g28** | 144578at7742 | 000162F_007\|arr | 1 | 5173 | 5172 | SUPER_5_unloc_BUSCO_144578at7742 | 99.633 | 5172 | 17 | 2 | 1 | 5172 | 8026 | 2857 | 0 | 9444 |

| Table S42 - False duplication content. | | | | | | | | | |
|---|---|---|---|---|---|---|---|---|---|
| **Histogram** | **cutoff** | **1** | **2** | **3** | **4** | **>4** | **dup(>1)** | **all** | **dup%** |
| **bHirRus1 before BUSCO gene recov** | 84 | 9.27E+08 | 3935932 | 253910 | 49034 | 27208 | 4266084 | 9.31E+08 | 0.458118 |
| **bHirRus1** | 84 | 9.28E+08 | 4218223 | 267726 | 54013 | 29560 | 4569522 | 9.32E+08 | 0.490137 |
| **c1** | 84 | 9.01E+08 | 36656566 | 707513 | 147460 | 62899 | 37574438 | 9.39E+08 | 4.00275 |
| **p1** | 84 | 9.22E+08 | 4258910 | 266200 | 53209 | 29325 | 4607644 | 9.27E+08 | 0.49727 |
| **Chelidonia** | 84 | 9.38E+08 | 11470565 | 870422 | 152316 | 74823 | 12568126 | 9.50E+08 | 1.32249 |
| **Chelidonia after haplotig purging** | 84 | 9.38E+08 | 4748015 | 323016 | 62305 | 39539 | 5172875 | 9.43E+08 | 0.54871 |

| Table S43 - Mitogenome annotation | | | | |
|---|---|---|---|---|
| **Annotation** | **start** | **end** | **e-value** | **strand** |
| **OH** | 23 | 229 | 1.7E+06 | + |
| **trnF(gaa)** | 248 | 317 | 1.1E-11 | + |
| **rrnS** | 316 | 1290 | 0 | + |
| **trnV(tac)** | 1289 | 1359 | 9.4E-12 | + |
| **rrnL** | 1367 | 2943 | 0 | + |
| **trnL2(taa)** | 2967 | 3042 | 8.7E-13 | + |
| **nad1** | 3069 | 4040 | 8.9E+08 | + |
| **trnI(gat)** | 4054 | 4125 | 1.7E-08 | + |
| **trnQ(ttg)** | 4132 | 4204 | 1.9E-14 | - |
| **trnM(cat)** | 4203 | 4272 | 2.1E-13 | + |
| **nad2** | 4272 | 5313 | 5.9E+08 | + |
| **trnW(tca)** | 5312 | 5383 | 1.8E-14 | + |
| **trnA(tgc)** | 5384 | 5453 | 5.4E-12 | - |
| **trnN(gtt)** | 5462 | 5535 | 4.8E-13 | - |
| **trnC(gca)** | 5535 | 5601 | 4.9E-10 | - |
| **trnY(gta)** | 5600 | 5671 | 1.5E-14 | - |
| **cox1** | 5672 | 7229 | 2.4E+09 | + |
| **trnS2(tga)** | 7220 | 7295 | 4.5E-14 | - |
| **trnD(gtc)** | 7299 | 7368 | 4.5E-12 | + |
| **cox2** | 7378 | 8062 | 5.0E+08 | + |
| **trnK(ttt)** | 8063 | 8134 | 2.5E-13 | + |
| **atp8** | 8135 | 8303 | 5.3E+06 | + |
| **atp6** | 8293 | 8977 | 3.2E+08 | + |
| **cox3** | 8983 | 9768 | 7.3E+08 | + |
| **trnG(tcc)** | 9767 | 9836 | 5.3E-12 | + |
| **nad3** | 9836 | 10187 | 9.2E+07 | + |
| **trnR(tcg)** | 10189 | 10259 | 8.6E-12 | + |
| **nad4l** | 10260 | 10557 | 4.8E+07 | + |

| | | | | |
|---|---|---|---|---|
| **nad4** | 10550 | 11928 | 1.4E+09 | + |
| **trnH(gtg)** | 11928 | 11997 | 1.9E-11 | + |
| **trnS1(gct)** | 11997 | 12064 | 2.9E-09 | + |
| **trnL1(tag)** | 12063 | 12134 | 3.4E-19 | + |
| **nad5** | 12134 | 13952 | 2.0E+09 | + |
| **cob** | 13960 | 15103 | 1.6E+09 | + |
| **trnT(tgt)** | 15107 | 15176 | 3.5E-12 | + |
| **OL** | 15862 | 15896 | 3.6E-02 | - |

Table S44 - Number of SNPs identified in the different datasets

| Identifier | SNP number after filtering |
|---|---|
| **ds2.1** | 4,180,839 |
| **ds2.2** | 2,296,850 |
| **ds2.3** | 5,783,842 |
| **ds2.4** | 7,543,250 |
| **ds2.5** | 3,762,802 |
| **ds2.6** | 6,471,459 |
| **ds2.7** | 2,414,350 |
| **ds3.1.1** | 3,111,728 |
| **ds3.1.2** | 5,022,964 |
| **ds3.2.1** | 42,022 |
| **ds3.2.2** | 16,103 |
| **ds4** | 30,491 |
| **ds5** | 28,840 |
| **ds6.1** | 22,650 |
| **ds6.2** | 36,835 |
| **ds6.3** | 26,291 |
| **ds6.4** | 17,925 |
| **ds6.5** | 15,682 |
| **ds6.6** | 18,446 |
| **ds6.7** | 16,824 |
| **ds6.8** | 31,804 |

**Table S45 - Summary of the values relative to variant depth in the different datasets after our filtering pipeline.** For each dataset, the first value refers to average depth of coverage per site, while the second refers to the mean depth of coverage per individual of the dataset

| Identifier | min | 1st quantile | median | mean | 3rd quantile | max |
|---|---|---|---|---|---|---|
| **ds2.1** | 2.3 ; 3.09 | 4.58 ; 4.84 | 5.21 ; 5.16 | 5.21 ; 5.21 | 5.82 ; 5.71 | 11.85 ; 6.54 |
| **ds2.2** | 2.09 ; 2.04 | 4.25 ; 4.43 | 4.76 ; 4.86 | 4.77 ; 4.75 | 5.28 ; 5.25 | 7.84 ; 6.14 |
| **ds2.3** | 1.87 ; 5.25 | 4.87 ; 5.41 | 5.75 ; 5.66 | 5.71 ; 5.71 | 6.5 ; 5.88 | 10.5 ; 6.55 |
| **ds2.4** | 1.8 ; 2.98 | 4.3 ; 4.95 | 5.1 ; 5.06 | 5.14 ; 5.13 | 5.9 ; 5.29 | 10.2 ; 6.66 |
| **ds2.5** | 2.29 ; 3.27 | 4.64 ; 4.89 | 5.24 ; 5.28 | 5.25 ; 5.25 | 5.86 ; 5.73 | 8.65 ; 7.11 |
| **ds2.6** | 2 ; 4.7 | 4.57 ; 4.93 | 5.38 ; 5.46 | 5.37 ; 5.37 | 6.14; 5.57 | 10.57 ; 6.17 |
| **ds2.7** | 2 ; 2.43 | 4.12 ; 4.51 | 4.67 ; 4.63 | 4.68 ; 4.67 | 5.25 ; 4.74 | 9 ; 6.97 |
| **ds3.1.1** | 2 ; 4.08 | 4.25 ; 4.75 | 4.87 ; 4.91 | 4.87 ; 4.87 | 5.5 ; 5.08 | 8.62 ; 5.28 |
| **ds3.1.2** | 2 ; 4.91 | 5 ; 5.49 | 5.87 ; 5.88 | 5.8 ; 5.8 | 6.62 ; 6.24 | 10.62 ; 6.29 |
| **ds3.2.1** | 2.35 ; 2.78 | 3.77 ; 3.62 | 4.36 ; 4.35 | 4.5 ; 4.49 | 5.11 ; 5.27 | 7.97 ; 6.72 |
| **ds3.2.2** | 2.21 ; 2.41 | 3.3 ; 3.24 | 3.76 ; 4.15 | 3.88 ; 3.87 | 4.36 ; 4.45 | 6.68 ; 4.96 |
| **ds4** | 3.63 ; 1.76 | 4.94 ; 3.89 | 5.63 ; 5.32 | 5.8 ; 5.73 | 6.53 ; 7.13 | 12.78 ; 14.14 |
| **ds5** | 7.17 ; 7.96 | 19.71 ; 21.09 | 26.19 ; 23.95 | 27.25 ; 27.25 | 33.93 ; 28.33 | 64.41 ; 101.38 |
| **ds6.1** | 1.34 ; 1.25 | 1.71 ; 1.53 | 1.85 ; 1.72 | 1.89 ; 1.83 | 2.02 ; 2.09 | 3.09 ; 3.06 |
| **ds6.2** | 1.06 ; 1.45 | 1.71 ; 1.7 | 1.92 ; 1.87 | 1.96 ; 1.95 | 2.17 ; 2.12 | 3.81 ; 2.74 |
| **ds6.3** | 1 ; 1.24 | 1.65 ; 1.49 | 1.86 ; 1.63 | 1.9 ; 1.83 | 2.08 ; 2.07 | 7.59 ; 3.03 |
| **ds6.4** | 1 ; 1.27 | 1.4 ; 1.48 | 1.6 ; 1.59 | 1.66 ; 1.64 | 1.83 ; 1.81 | 3.21 ; 2.12 |
| **ds6.5** | 1.14 ; 1.23 | 1.59 ; 1.47 | 1.76 ; 1.66 | 1.79 ; 1.74 | 1.94 ; 1.88 | 3.13 ; 3.32 |
| **ds6.6** | 1 ; 1.32 | 1.4 ; 1.43 | 1.67 ; 1.57 | 1.71 ; 1.67 | 2 ; 1.64 | 3.86 ; 2.66 |
| **ds6.7** | 1.04 ; 1.24 | 1.56 ; 1.53 | 1.71 ; 1.72 | 1.74 ; 1.71 | 1.89 ; 1.87 | 4.23 ; 2.59 |
| **ds6.8** | 1 ; 1.4 | 1.61 ; 1.62 | 1.85 ; 1.78 | 1.88 ; 1.86 | 2.08 ; 2.07 | 4.64 ; 2.74 |

| Table S46 - SNP counts from dataset1 (HiFi 20x) falling in the different genomic regions | | | | | | | |
|---|---|---|---|---|---|---|---|
| **chr** | **type** | **chr size (bp)** | **total number of SNPs** | **intergenic** | **genic** | **exonic** | **intronic** |
| **chr1** | macrochromosome | 156,035,725 | 2,235,865 | 1,135,091 | 1,100,774 | 50,344 | 1,057,446 |
| **chr2** | macrochromosome | 119,023,421 | 1,874,228 | 936,815 | 937,413 | 50,861 | 894,063 |
| **chr3** | macrochromosome | 116,801,625 | 1,713,055 | 811,706 | 901,349 | 50,140 | 858,949 |
| **chrZ** | macrochromosome | 90,132,487 | 286,586 | 149,838 | 136,748 | 8,500 | 129,272 |
| **chr4** | macrochromosome | 76,187,387 | 1,002,509 | 400,577 | 601,932 | 34,838 | 572,431 |
| **chr5** | macrochromosome | 73,257,097 | 1,166,751 | 548,354 | 618,397 | 34,454 | 589,413 |
| **chr6** | macrochromosome | 63,258,489 | 877,095 | 383,240 | 493,855 | 37,278 | 461,781 |
| **chr7** | intermediate | 38,459,648 | 659,436 | 259,207 | 400,229 | 24,235 | 379,921 |
| **chr8** | intermediate | 36,085,389 | 638,569 | 216,671 | 421,898 | 23,952 | 402,139 |
| **chrW** | intermediate | 31,704,074 | 10,022 | 2,195 | 7,827 | 1,541 | 6,348 |
| **chr9** | intermediate | 31,262,510 | 535,687 | 206,458 | 329,229 | 23,107 | 309,490 |
| **chr10** | intermediate | 25,880,253 | 488,472 | 209,498 | 278,974 | 20,721 | 261,873 |
| **chr11** | intermediate | 21,491,857 | 329,758 | 152,994 | 176,764 | 14,260 | 164,754 |
| **chr12** | intermediate | 20,890,524 | 382,823 | 124,081 | 258,742 | 15,623 | 246,077 |
| **chr13** | intermediate | 20,272,128 | 355,942 | 127,538 | 228,404 | 16,827 | 213,920 |
| **chr14** | microchromosome | 18,810,845 | 348,540 | 155,294 | 193,246 | 14,345 | 180,657 |
| **chr15** | microchromosome | 16,541,138 | 311,453 | 108,092 | 203,361 | 16,731 | 189,190 |
| **chr16** | microchromosome | 15,277,844 | 276,785 | 128,954 | 147,831 | 14,693 | 134,886 |
| **chr17** | microchromosome | 13,985,943 | 254,428 | 80,606 | 173,822 | 16,285 | 159,672 |
| **chr18** | microchromosome | 12,073,725 | 201,040 | 78,479 | 122,561 | 13,365 | 110,793 |
| **chr19** | microchromosome | 11,382,101 | 208,474 | 56,168 | 152,306 | 14,282 | 140,148 |
| **chr20** | microchromosome | 11,194,341 | 210,406 | 74,426 | 135,980 | 13,740 | 124,329 |
| **chr21** | microchromosome | 9,617,204 | 193,476 | 76,039 | 117,437 | 10,188 | 108,455 |
| **chr22** | microchromosome | 7,507,825 | 124,436 | 40,269 | 84,167 | 8,392 | 76,915 |
| **chr23** | microchromosome | 7,098,401 | 145,259 | 53,183 | 92,076 | 7,244 | 85,532 |
| **chr24** | microchromosome | 6,843,954 | 123,102 | 46,309 | 76,793 | 9,274 | 68,635 |
| **chr25** | microchromosome | 6,778,862 | 120,232 | 58,735 | 61,497 | 7,628 | 54,941 |
| **chr26** | microchromosome | 5,553,549 | 90,808 | 30,230 | 60,578 | 8,244 | 53,151 |

| | | | | | | | |
|---|---|---|---|---|---|---|---|
| **chr28** | microchromosome | 5,297,670 | 79,828 | 37,718 | 42,110 | 4,365 | 38,023 |
| **chr27** | microchromosome | 5,236,451 | 87,168 | 23,190 | 63,978 | 8,192 | 56,495 |
| **chr29** | microchromosome | 2,102,120 | 34,416 | 12,750 | 21,666 | 4,421 | 17,717 |
| **chr30** | microchromosome | 1,648,998 | 16,907 | 4,413 | 12,494 | 2,060 | 10,571 |
| **chr31** | microchromosome | 1,590,086 | 6,559 | 2,672 | 3,887 | 359 | 3,672 |
| **chr32** | microchromosome | 784,579 | 5,733 | 4,197 | 1,536 | 508 | 1,259 |
| **chr34** | microchromosome | 606,149 | 1,925 | 1,479 | 446 | 128 | 354 |
| **chr35** | microchromosome | 523,230 | 7,090 | 4,144 | 2,946 | 537 | 2,451 |
| **chr33** | microchromosome | 437,724 | 720 | 625 | 95 | 20 | 75 |
| **chr36** | microchromosome | 338,027 | 1,741 | 438 | 1,303 | 322 | 1,085 |
| **chr37** | microchromosome | 276,370 | 4,050 | 1,840 | 2,210 | 569 | 1,900 |
| **chr38** | microchromosome | 220,485 | 265 | 29 | 236 | 39 | 206 |

Table S47 - SNP counts from downsampled dataset1 (HiFi 5x intersected with HiFi 20x as truth set) falling in the different genomic regions

| chr | type | chr size (bp) | total number of SNPs | intergenic | genic | exonic | intronic |
|---|---|---|---|---|---|---|---|
| **chr1** | macrochromosome | 156,035,725 | 1,552,209 | 773,675 | 778,534 | 35,266 | 748,172 |
| **chr2** | macrochromosome | 119,023,421 | 1,302,118 | 639,492 | 662,626 | 35,475 | 632,499 |
| **chr3** | macrochromosome | 116,801,625 | 1,200,944 | 559,011 | 641,933 | 34,848 | 612,537 |
| **chrZ** | macrochromosome | 90,132,487 | 21,094 | 13,242 | 7,852 | 1,018 | 6,947 |
| **chr4** | macrochromosome | 76,187,387 | 692,717 | 271,944 | 420,773 | 24,236 | 400,267 |
| **chr5** | macrochromosome | 73,257,097 | 818,574 | 381,163 | 437,411 | 24,140 | 417,153 |
| **chr6** | macrochromosome | 63,258,489 | 609,627 | 258,940 | 350,687 | 26,096 | 328,300 |
| **chr7** | intermediate | 38,459,648 | 460,616 | 176,738 | 283,878 | 16,819 | 269,844 |
| **chr8** | intermediate | 36,085,389 | 445,824 | 146,462 | 299,362 | 16,746 | 285,642 |
| **chrW** | intermediate | 31,704,074 | 3,215 | 669 | 2,546 | 714 | 1,872 |
| **chr9** | intermediate | 31,262,510 | 372,503 | 140,733 | 231,770 | 16,062 | 218,060 |
| **chr10** | intermediate | 25,880,253 | 339,236 | 142,956 | 196,280 | 14,125 | 184,645 |
| **chr11** | intermediate | 21,491,857 | 222,277 | 101,044 | 121,233 | 9,344 | 113,318 |
| **chr12** | intermediate | 20,890,524 | 265,602 | 84,366 | 181,236 | 10,724 | 172,519 |
| **chr13** | intermediate | 20,272,128 | 247,645 | 88,791 | 158,854 | 11,390 | 149,102 |
| **chr14** | microchromosome | 18,810,845 | 237,131 | 103,868 | 133,263 | 9,567 | 124,799 |
| **chr15** | microchromosome | 16,541,138 | 213,204 | 72,919 | 140,285 | 11,278 | 130,780 |
| **chr16** | microchromosome | 15,277,844 | 190,293 | 86,633 | 103,660 | 9,928 | 94,950 |
| **chr17** | microchromosome | 13,985,943 | 173,657 | 52,498 | 121,159 | 11,274 | 111,314 |
| **chr18** | microchromosome | 12,073,725 | 130,382 | 48,813 | 81,569 | 8,814 | 73,754 |
| **chr19** | microchromosome | 11,382,101 | 142,984 | 37,160 | 105,824 | 9,662 | 97,627 |
| **chr20** | microchromosome | 11,194,341 | 139,941 | 48,394 | 91,547 | 9,261 | 83,696 |
| **chr21** | microchromosome | 9,617,204 | 132,068 | 50,382 | 81,686 | 6,902 | 75,646 |
| **chr22** | microchromosome | 7,507,825 | 79,904 | 25,827 | 54,077 | 5,479 | 49,324 |
| **chr23** | microchromosome | 7,098,401 | 91,185 | 32,534 | 58,651 | 4,503 | 54,595 |
| **chr24** | microchromosome | 6,843,954 | 74,420 | 27,254 | 47,166 | 5,636 | 42,190 |
| **chr25** | microchromosome | 6,778,862 | 71,673 | 34,921 | 36,752 | 4,738 | 32,729 |
| **chr26** | microchromosome | 5,553,549 | 55,033 | 18,318 | 36,715 | 4,894 | 32,352 |

| | | | | | | | |
|---|---|---|---|---|---|---|---|
| **chr28** | microchromosome | 5,297,670 | 44,796 | 21,490 | 23,306 | 2,663 | 20,792 |
| **chr27** | microchromosome | 5,236,451 | 54,924 | 14,099 | 40,825 | 5,467 | 35,823 |
| **chr29** | microchromosome | 2,102,120 | 18,928 | 6,492 | 12,436 | 2,479 | 10,216 |
| **chr30** | microchromosome | 1,648,998 | 7,934 | 2,211 | 5,723 | 963 | 4,802 |
| **chr31** | microchromosome | 1,590,086 | 3,958 | 1,564 | 2,394 | 225 | 2,260 |
| **chr32** | microchromosome | 784,579 | 3,175 | 2,363 | 812 | 314 | 638 |
| **chr34** | microchromosome | 606,149 | 958 | 711 | 247 | 64 | 196 |
| **chr35** | microchromosome | 523,230 | 3,817 | 2,266 | 1,551 | 302 | 1,257 |
| **chr33** | microchromosome | 437,724 | 288 | 252 | 36 | 4 | 32 |
| **chr36** | microchromosome | 338,027 | 726 | 193 | 533 | 130 | 443 |
| **chr37** | microchromosome | 276,370 | 1,896 | 810 | 1,086 | 229 | 979 |
| **chr38** | microchromosome | 220,485 | 42 | 12 | 30 | 10 | 20 |

| Table S48 - SNP counts from all Illumina WGS datasets (ds2 and ds3.1) combined falling in the different genomic regions | | | | | | | |
|---|---|---|---|---|---|---|---|
| **chr** | **type** | **chr size (bp)** | **total number of SNPs** | **intergenic** | **genic** | **exonic** | **intronic** |
| **chr1** | macrochromosome | 156,035,725 | 2,378,916 | 1,186,927 | 1,191,989 | 62,021 | 1,138,288 |
| **chr2** | macrochromosome | 119,023,421 | 2,033,905 | 973,683 | 1,060,222 | 64,512 | 1,004,527 |
| **chr3** | macrochromosome | 116,801,625 | 1,915,698 | 916,951 | 998,747 | 61,180 | 946,537 |
| **chrZ** | macrochromosome | 90,132,487 | 480,430 | 256,672 | 223,758 | 15,408 | 210,137 |
| **chr4** | macrochromosome | 76,187,387 | 1,144,428 | 449,665 | 694,763 | 42,860 | 658,428 |
| **chr5** | macrochromosome | 73,257,097 | 1,232,652 | 562,448 | 670,204 | 42,488 | 634,380 |
| **chr6** | macrochromosome | 63,258,489 | 1,011,284 | 426,503 | 584,781 | 46,559 | 544,350 |
| **chr7** | intermediate | 38,459,648 | 752,364 | 296,341 | 456,023 | 30,285 | 430,340 |
| **chr8** | intermediate | 36,085,389 | 739,420 | 247,000 | 492,420 | 30,797 | 466,825 |
| **chrW** | intermediate | 31,704,074 | 4,364 | 1,134 | 3,230 | 890 | 2,379 |
| **chr9** | intermediate | 31,262,510 | 609,915 | 223,719 | 386,196 | 28,682 | 361,380 |
| **chr10** | intermediate | 25,880,253 | 565,813 | 234,198 | 331,615 | 26,679 | 309,478 |
| **chr11** | intermediate | 21,491,857 | 372,113 | 168,312 | 203,801 | 17,082 | 189,353 |
| **chr12** | intermediate | 20,890,524 | 444,832 | 139,174 | 305,658 | 19,409 | 289,607 |
| **chr13** | intermediate | 20,272,128 | 421,690 | 149,382 | 272,308 | 21,205 | 253,839 |
| **chr14** | microchromosome | 18,810,845 | 394,972 | 173,218 | 221,754 | 16,943 | 206,668 |
| **chr15** | microchromosome | 16,541,138 | 365,592 | 126,085 | 239,507 | 21,897 | 220,916 |
| **chr16** | microchromosome | 15,277,844 | 325,289 | 149,326 | 175,963 | 18,402 | 159,654 |
| **chr17** | microchromosome | 13,985,943 | 313,512 | 96,889 | 216,623 | 21,737 | 197,550 |
| **chr18** | microchromosome | 12,073,725 | 232,814 | 89,417 | 143,397 | 16,182 | 128,928 |
| **chr19** | microchromosome | 11,382,101 | 244,586 | 60,781 | 183,805 | 18,164 | 168,236 |
| **chr20** | microchromosome | 11,194,341 | 250,938 | 86,332 | 164,606 | 17,734 | 149,391 |
| **chr21** | microchromosome | 9,617,204 | 213,087 | 78,872 | 134,215 | 12,983 | 122,667 |
| **chr22** | microchromosome | 7,507,825 | 150,522 | 47,155 | 103,367 | 11,454 | 93,173 |
| **chr23** | microchromosome | 7,098,401 | 164,593 | 57,758 | 106,835 | 8,735 | 98,858 |
| **chr24** | microchromosome | 6,843,954 | 130,777 | 45,994 | 84,783 | 11,622 | 74,474 |
| **chr25** | microchromosome | 6,778,862 | 134,800 | 64,150 | 70,650 | 9,187 | 62,721 |
| **chr26** | microchromosome | 5,553,549 | 96,966 | 29,694 | 67,272 | 9,749 | 58,378 |

| | | | | | | | |
|---|---|---|---|---|---|---|---|
| **chr28** | microchromosome | 5,297,670 | 77,700 | 36,196 | 41,504 | 4,754 | 37,112 |
| **chr27** | microchromosome | 5,236,451 | 87,208 | 18,864 | 68,344 | 9,261 | 59,882 |
| **chr29** | microchromosome | 2,102,120 | 31,055 | 10,144 | 20,911 | 4,413 | 16,995 |
| **chr30** | microchromosome | 1,648,998 | 15,060 | 3,135 | 11,925 | 2,184 | 9,824 |
| **chr31** | microchromosome | 1,590,086 | 2,527 | 1,142 | 1,385 | 220 | 1,270 |
| **chr32** | microchromosome | 784,579 | 2,104 | 1,443 | 661 | 198 | 533 |
| **chr34** | microchromosome | 606,149 | 1,104 | 738 | 366 | 101 | 290 |
| **chr35** | microchromosome | 523,230 | 8,220 | 4,410 | 3,810 | 891 | 2,970 |
| **chr33** | microchromosome | 437,724 | 104 | 88 | 16 | 8 | 15 |
| **chr36** | microchromosome | 338,027 | 1,812 | 328 | 1,484 | 496 | 1,040 |
| **chr37** | microchromosome | 276,370 | 2,675 | 1,017 | 1,658 | 400 | 1,416 |
| **chr38** | microchromosome | 220,485 | 408 | 26 | 382 | 107 | 292 |

| Table S49 - SNP counts from all ddRAD datasets (ds3.2 through ds6) combined falling in the different genomic regions | | | | | | | |
|---|---|---|---|---|---|---|---|
| **chr** | **type** | **chr size (bp)** | **total number of SNPs** | **intergenic** | **genic** | **exonic** | **intronic** |
| **chr1** | macrochromosome | 156,035,725 | 12,046 | 5,926 | 6,120 | 326 | 5,847 |
| **chr2** | macrochromosome | 119,023,421 | 9,613 | 4,597 | 5,016 | 325 | 4,719 |
| **chr3** | macrochromosome | 116,801,625 | 9,641 | 4,444 | 5,197 | 323 | 4,915 |
| **chrZ** | macrochromosome | 90,132,487 | 2,452 | 1,359 | 1,093 | 82 | 1,015 |
| **chr4** | macrochromosome | 76,187,387 | 5,704 | 2,263 | 3,441 | 202 | 3,264 |
| **chr5** | macrochromosome | 73,257,097 | 6,145 | 2,824 | 3,321 | 189 | 3,157 |
| **chr6** | macrochromosome | 63,258,489 | 5,015 | 2,087 | 2,928 | 223 | 2,732 |
| **chr7** | intermediate | 38,459,648 | 3,793 | 1,412 | 2,381 | 149 | 2,242 |
| **chr8** | intermediate | 36,085,389 | 3,555 | 1,140 | 2,415 | 101 | 2,338 |
| **chrW** | intermediate | 31,704,074 | 24 | 8 | 16 | 4 | 14 |
| **chr9** | intermediate | 31,262,510 | 3,320 | 1,201 | 2,119 | 156 | 1,972 |
| **chr10** | intermediate | 25,880,253 | 3,049 | 1,280 | 1,769 | 127 | 1,649 |
| **chr11** | intermediate | 21,491,857 | 2,085 | 994 | 1,091 | 76 | 1,025 |
| **chr12** | intermediate | 20,890,524 | 2,199 | 731 | 1,468 | 78 | 1,405 |
| **chr13** | intermediate | 20,272,128 | 2,106 | 757 | 1,349 | 65 | 1,296 |
| **chr14** | microchromosome | 18,810,845 | 1,769 | 767 | 1,002 | 60 | 946 |
| **chr15** | microchromosome | 16,541,138 | 1,866 | 620 | 1,246 | 98 | 1,161 |
| **chr16** | microchromosome | 15,277,844 | 1,460 | 685 | 775 | 93 | 690 |
| **chr17** | microchromosome | 13,985,943 | 1,559 | 439 | 1,120 | 91 | 1,038 |
| **chr18** | microchromosome | 12,073,725 | 1,169 | 522 | 647 | 78 | 575 |
| **chr19** | microchromosome | 11,382,101 | 1,327 | 318 | 1,009 | 97 | 944 |
| **chr20** | microchromosome | 11,194,341 | 1,183 | 351 | 832 | 85 | 760 |
| **chr21** | microchromosome | 9,617,204 | 1,094 | 390 | 704 | 65 | 646 |
| **chr22** | microchromosome | 7,507,825 | 733 | 195 | 538 | 50 | 495 |
| **chr23** | microchromosome | 7,098,401 | 762 | 297 | 465 | 26 | 439 |
| **chr24** | microchromosome | 6,843,954 | 606 | 201 | 405 | 51 | 363 |
| **chr25** | microchromosome | 6,778,862 | 708 | 348 | 360 | 23 | 341 |
| **chr26** | microchromosome | 5,553,549 | 463 | 142 | 321 | 38 | 287 |

| | | | | | | | |
|---|---|---|---|---|---|---|---|
| **chr28** | microchromosome | 5,297,670 | 485 | 250 | 235 | 17 | 219 |
| **chr27** | microchromosome | 5,236,451 | 441 | 92 | 349 | 24 | 326 |
| **chr29** | microchromosome | 2,102,120 | 105 | 36 | 69 | 22 | 51 |
| **chr30** | microchromosome | 1,648,998 | 62 | 5 | 57 | 7 | 50 |
| **chr31** | microchromosome | 1,590,086 | 3 | 2 | 1 | 0 | 1 |
| **chr32** | microchromosome | 784,579 | 9 | 9 | 0 | 0 | 0 |
| **chr34** | microchromosome | 606,149 | 3 | 2 | 1 | 0 | 1 |
| **chr35** | microchromosome | 523,230 | 44 | 11 | 33 | 3 | 30 |
| **chr33** | microchromosome | 437,724 | 0 | 0 | 0 | 0 | 0 |
| **chr36** | microchromosome | 338,027 | 6 | 1 | 5 | 2 | 3 |
| **chr37** | microchromosome | 276,370 | 9 | 9 | 0 | 0 | 0 |
| **chr38** | microchromosome | 220,485 | 11 | 11 | 0 | 0 | 0 |

| **Table S50 - HiFi data alignment statistics.** Alignment metrics from pbmm2 output for all HiFi samples (ds1) | | | | | |
|---|---|---|---|---|---|
| | **Sample** | | | | |
| | **A1** | **A2** | **2** | **3** | **4** |
| **Mapped reads** | 2,475,803 | 930,924 | 1,842,954 | 1,519,281 | 1,420,140 |
| **Alignments** | 2,994,011 | 1,215,608 | 2,258,506 | 1,825,799 | 1,777,646 |
| **Mapped bases** | 35,698,278,367 | 16,611,159,086 | 27,070,079,509 | 21,144,417,593 | 22,159,332,354 |
| **Mean mapped concordance** | 96.9233% | 96.5095% | 96.8043% | 96.9713% | 96.6645% |
| **Max mapped read lenght** | 49,467 | 47,385 | 35,156 | 43,871 | 35,559 |
| **Mean mapped read lenght** | 11,923.20 | 13,664.90 | 11,985.80 | 11,580.90 | 12,465.50 |

**Table S51 - SNP recall rate after downsampling for all HiFi samples (ds1).** The first column indicates sample name and sequencing coverage. For truth set 1, the 20x filtered variants (or 15x for sample A2) constitute the truth set. For the other truth sets, the SNP number relative to the truth set are in the column "Variants after intersection" in correspondence of the full coverage datasets line. See Methods section for a complete definition of the different truth sets. SD: standard deviation.

**Truth set 1**

| Sample A1 | Filtered variants | Variants after intersection | Recall | Triplicates average | SD | Precision | Triplicates average | SD | F1 score | Triplicates average | SD |
|---|---|---|---|---|---|---|---|---|---|---|---|
| 20x | 6,899,805 | | 1 | | | 1.000000 | | | | | |
| 15x | 6,579,094 | 6,430,950 | 0.9320 | 0.932221998 | 0.000310 | 0.977483 | 0.977377 | 0.000134 | 0.954225 | 0.954223 | 0.000061 |
| 15x | 6,579,433 | 6,430,883 | 0.9320 | | | 0.977422 | | | 0.954161 | | |
| 15x | 6,584,578 | 6,434,617 | 0.9326 | | | 0.977225 | | | 0.954282 | | |
| 10x | 6,102,026 | 5,852,497 | 0.8482 | 0.849106509 | 0.000872 | 0.959107 | 0.959261 | 0.000139 | 0.891322 | 0.897882 | 0.005682 |
| 10x | 6,107,075 | 5,859,001 | 0.8492 | | | 0.959379 | | | 0.901044 | | |
| 10x | 6,113,345 | 5,864,510 | 0.8500 | | | 0.959296 | | | 0.901279 | | |
| 7,5x | 5,294,594 | 5,036,554 | 0.7300 | 0.730310736 | 0.00031 | 0.951263 | 0.951739 | 0.00041 | 0.822116 | 0.825264 | 0.002732 |
| 7,5x | 5,295,112 | 5,040,616 | 0.7305 | | | 0.951938 | | | 0.827006 | | |
| 7,5x | 5,293,861 | 5,039,835 | 0.7304 | | | 0.952015 | | | 0.826670 | | |
| 5x | 3,834,530 | 3,605,144 | 0.5225 | 0.521775036 | 0.000745 | 0.940179 | 0.940056 | 0.000365 | 0.666311 | 0.669301 | 0.002589 |
| 5x | 3,831,682 | 3,600,420 | 0.5218 | | | 0.939645 | | | 0.670756 | | |
| 5x | 3,822,936 | 3,594,874 | 0.5210 | | | 0.940344 | | | 0.670835 | | |
| **Sample A2** | **Filtered variants** | **Variants after intersection** | **Recall** | **Triplicates average** | **SD** | **Precision** | **Triplicates average** | **SD** | **F1 score** | **Triplicates average** | **SD** |
| 15x | 6660405 | | 1 | | | 1.000000 | | | | | |
| 10x | 6008544 | 5762246 | 0.86515 | 0.866295568 | 0.000992531 | 0.959009 | 0.958957 | | 0.909664 | 0.910261 | 0.000517 |
| 10x | 6020967 | 5773696 | 0.86687 | | | 0.958932 | | 0.000044 | 0.910541 | | |
| 10x | 6020967 | 5773696 | 0.86687 | | | 0.958932 | | | 0.910579 | | |
| 7,5x | 5178718 | 4918934 | 0.73853 | 0.73764 | 0.000819 | 0.949836 | 0.949808 | | 0.826510 | 0.828991 | 0.002205 |
| 7,5x | 5174620 | 4911792 | 0.73746 | | | 0.949208 | | 0.000587 | 0.829734 | | |
| 7,5x | 5164481 | 4908223 | 0.73693 | | | 0.950381 | | | 0.830729 | | |
| 5x | 3691591 | 3460308 | 0.51953 | 0.51767 | 0.001647 | 0.937349 | 0.936976 | | 0.662603 | 0.664783 | 0.001936 |
| 5x | 3675079 | 3443882 | 0.51707 | | | 0.937091 | | 0.000441 | 0.666301 | | |
| 5x | 3672761 | 3439502 | 0.51641 | | | 0.936489 | | | 0.665446 | | |
| **Sample 2** | **Filtered variants** | **Variants after intersection** | **Recall** | **Triplicates average** | **SD** | **Precision** | **Triplicates average** | **SD** | **F1 score** | **Triplicates average** | **SD** |
| 20x | 6,805,561 | | 1 | | | 1.000000 | | | | | |
| 15x | 6,579,792 | 6,377,379 | 0.9370835 | 0.9368892 | 0.0003321 | 0.969237 | 0.969197 | 0.000081 | 0.952889 | 0.952747 | 0.000143 |
| 15x | 6,575,649 | 6,373,447 | 0.9365057 | | | 0.969250 | | | 0.952603 | | |
| 15x | 6,580,659 | 6,377,344 | 0.9370784 | | | 0.969104 | | | 0.952749 | | |
| 10x | 6,062,669 | 5,757,563 | 0.8460086 | 0.8476337 | 0.0663027 | 0.949675 | 0.949413 | 0.000291 | 0.885271 | 0.892413 | 0.006208 |
| 10x | 6,081,141 | 5,771,609 | 0.8480725 | | | 0.949100 | | | 0.895460 | | |
| 10x | 6,084,159 | 5,776,696 | 0.8488200 | | | 0.949465 | | | 0.896508 | | |
| 7,5x | 5,266,017 | 4,948,838 | 0.7271756 | 0.7259847 | 0.0012100 | 0.939769 | 0.939978 | 0.00018 | 0.815174 | 0.817710 | 0.002246 |
| 7,5x | 5,255,958 | 4,940,988 | 0.7260221 | | | 0.940074 | | | 0.819448 | | |
| 7,5x | 5,246,690 | 4,932,374 | 0.7247564 | | | 0.940093 | | | 0.818508 | | |
| 5x | 3,779,998 | 3,498,356 | 0.5140437 | 0.5144114 | 0.0011110 | 0.925491 | 0.925842 | 0.000350 | 0.654331 | 0.659256 | 0.004364 |
| 5x | 3,774,794 | 3,494,866 | 0.5135309 | | | 0.925843 | | | 0.660794 | | |
| 5x | 3,789,013 | 3,509,353 | 0.5156596 | | | 0.926192 | | | 0.662642 | | |
| **Sample 3** | **Filtered variants** | **Variants after intersection** | **Recall** | **Triplicates average** | **SD** | **Precision** | **Triplicates average** | **SD** | **F1 score** | **Triplicates average** | **SD** |
| 20x | 6,789,479 | | 1 | | | 1.000000 | | | | | |
| 15x | 6,550,999 | 6,359,642 | 0.93669 | 0.9364883 | 0.00037090 | 0.970790 | 0.970942 | 0.000134 | 0.953435 | 0.953446 | 0.000139 |
| 15x | 6,545,193 | 6,355,361 | 0.93606 | | | 0.970997 | | | 0.953312 | | |
| 15x | 6,549,470 | 6,359,799 | 0.93671 | | | 0.971040 | | | 0.953590 | | |
| 10x | 6,061,636 | 5,757,149 | 0.84795 | 0.8190156 | 0.05971497 | 0.949768 | 0.949734 | 0.000287 | 0.885499 | 0.892925 | 0.006466 |
| 10x | 6,074,314 | 5,770,610 | 0.84993 | | | 0.950002 | | | 0.897303 | | |
| 10x | 6,069,323 | 5,762,403 | 0.84873 | | | 0.949431 | | | 0.895974 | | |
| 7,5x | 5,266,690 | 4,952,596 | 0.72945 | 0.7303293 | 0.00093788 | 0.940362 | 0.940138 | 0.00044 | 0.817149 | 0.820459 | 0.002951 |
| 7,5x | 5,279,850 | 4,965,265 | 0.73132 | | | 0.940418 | | | 0.822818 | | |
| 7,5x | 5,276,322 | 4,957,806 | 0.73022 | | | 0.939633 | | | 0.821409 | | |
| 5x | 3,813,134 | 3,536,101 | 0.52082 | 0.5202553 | 0.00079984 | 0.927348 | 0.927388 | 0.000127 | 0.661413 | 0.664691 | 0.002912 |
| 5x | 3,810,803 | 3,534,637 | 0.52061 | | | 0.927531 | | | 0.666979 | | |
| 5x | 3,802,549 | 3,526,049 | 0.51934 | | | 0.927286 | | | 0.665680 | | |
| **Sample 4** | **Filtered variants** | **Variants after intersection** | **Recall** | **Triplicates average** | **SD** | **Precision** | **Triplicates average** | **SD** | **F1 score** | **Triplicates average** | **SD** |
| 20x | 6,791,626 | | 1 | | | 1.000000 | | | | | |
| 15x | 6,578,490 | 6,365,722 | 0.9372898331 | 0.9369292911 | 0.000465302746 | 0.967657 | 0.967831 | 0.000156 | 0.952231 | 0.952166 | 0.000084 |
| 15x | 6,570,219 | 6,359,706 | 0.9364040364 | | | 0.967960 | | | 0.952072 | | |
| 15x | 6,575,624 | 6,364,392 | 0.9370940037 | | | 0.967877 | | | 0.952195 | | |
| 10x | 6,081,021 | 5,756,502 | 0.8475881917 | 0.8477179593 | 0.000712578451 | 0.946634 | 0.946618 | 0.000116 | 0.883912 | 0.890969 | 0.006116 |
| 10x | 6,088,362 | 5,762,603 | 0.8484865038 | | | 0.946495 | | | 0.894746 | | |
| 10x | 6,076,789 | 5,753,045 | 0.8470791825 | | | 0.946724 | | | 0.894249 | | |
| 7,5x | 5,278,852 | 4,941,880 | 0.7276431299 | 0.7262630481 | 0.001201027126 | 0.936166 | 0.935965 | 0.00032 | 0.813674 | 0.816159 | 0.002174 |
| 7,5x | 5,267,903 | 4,928,624 | 0.7256913146 | | | 0.935595 | | | 0.817102 | | |
| 7,5x | 5,263,156 | 4,927,017 | 0.7254546997 | | | 0.936134 | | | 0.817703 | | |
| 5x | 3,791,729 | 3,491,936 | 0.5141531645 | 0.514401166 | 0.000798817947 | 0.920935 | 0.921127 | 0.000166 | 0.652977 | 0.657886 | 0.004307 |
| 5x | 3,798,991 | 3,499,688 | 0.5152945701 | | | 0.921215 | | | 0.661032 | | |
| 5x | 3,787,581 | 3,489,237 | 0.5137557633 | | | 0.921231 | | | 0.659648 | | |

**Truth set 2**

| Sample A1 | Variants after intersection | Recall | Triplicates average | SD | Precision | Triplicates average | SD | F1 score | Triplicates average | SD |
|---|---|---|---|---|---|---|---|---|---|---|
| 20x | 1,249,452 | 1 | | | 1.0000000 | | | | | |
| 15x | 1,177,147 | 0.9421306301 | 0.9424886537 | 0.000313166705 | 0.1789223562 | 0.1789375764 | 0.000079396528 | 0.3007319622 | 0.3007716917 | 0.000117979344 |
| 15x | 1,177,873 | 0.9427116848 | | | 0.1790234812 | | | 0.3009044072 | | |
| 15x | 1,177,763 | 0.9426236462 | | | 0.1788668917 | | | 0.3006787056 | | |
| 10x | 1,085,136 | 0.8684895458 | 0.8696265776 | 0.00116719845 | 0.177832084 | 0.1779057865 | 0.000073700947 | 0.2952157376 | 0.2953829266 | 0.00016887706 |
| 10x | 1,086,484 | 0.8695684188 | | | 0.1779057896 | | | 0.2953795996 | | |
| 10x | 1,088,050 | 0.8708217683 | | | 0.1779794859 | | | 0.2955534425 | | |
| 7,5x | 939,337 | 0.7517991888 | 0.752023287 | 0.000269702807 | 0.1774143589 | 0.1774696437 | 0.000053214753 | 0.2870814172 | 0.2871701338 | 0.000088569433 |
| 7,5x | 939,991 | 0.7523226182 | | | 0.1775205133 | | | 0.2872585553 | | |
| 7,5x | 939,523 | 0.751948054 | | | 0.1774740591 | | | 0.2871704288 | | |
| 5x | 677,029 | 0.5418607518 | 0.5409619577 | 0.000824441819 | 0.1765611431 | 0.1764898905 | 0.000128669929 | 0.2663380791 | 0.2661482789 | 0.000190063348 |
| 5x | 675,684 | 0.5407842798 | | | 0.1763413561 | | | 0.2659579535 | | |
| 5x | 675,005 | 0.5402408416 | | | 0.1765671725 | | | 0.2661488041 | | |
| **Sample A2** | **Variants after intersection** | **Recall** | **Triplicates average** | **SD** | **Precision** | **Triplicates average** | **SD** | **F1 score** | **Triplicates average** | **SD** |
| 15x | 1,213,105 | 1 | | | 1.0000000 | | | | | |
| 10x | 1,067,713 | 0.880148874 | 0.881527705 | 0.001194103 | 0.1776991231 | 0.1777325002 | 0.000028905432 | 0.2956978385 | 0.2958217472 | 0.000107308028 |
| 10x | 1,070,222 | 0.882217121 | | | 0.1777491888 | | | 0.2958837015 | | |
| 10x | 1,070,222 | 0.882217121 | | | 0.1777491888 | | | 0.2958837015 | | |
| 7,5x | 918,083 | 0.756804234 | 0.755993642 | 0.001005915 | 0.1772799755 | 0.1772993503 | 0.000017455037 | 0.2872679672 | 0.2872348877 | 0.000053971264 |
| 7,5x | 917,482 | 0.756308811 | | | 0.1773042272 | | | 0.2872640886 | | |
| 7,5x | 915,734 | 0.75486788 | | | 0.1773138482 | | | 0.2871726073 | | |
| 5x | 650,480 | 0.536210798 | 0.534503609 | 0.001484016 | 0.176205869 | 0.1762072022 | 0.000097919914 | 0.265247836 | 0.2650397122 | 0.000220184484 |
| 5x | 647,218 | 0.53352183 | | | 0.1761099557 | | | 0.2648091806 | | |
| 5x | 647,529 | 0.533778197 | | | 0.1763057819 | | | 0.26506212 | | |
| **Sample 2** | **Variants after intersection** | **Recall** | **Triplicates average** | **SD** | **Precision** | **Triplicates average** | **SD** | **F1 score** | **Triplicates average** | **SD** |
| 20x | 1,230,900 | 1 | | | 1.0000000 | | | | | |
| 15x | 1,167,430 | 0.948436104 | 0.948105993 | 0.000375411 | 0.1774265813 | 0.1773942703 | 0.000035605551 | 0.2989312599 | 0.2988689986 | 0.000057221483 |
| 15x | 1,166,521 | 0.94769762 | | | 0.1774001319 | | | 0.2988570238 | | |
| 15x | 1,167,120 | 0.948184255 | | | 0.1773560976 | | | 0.2988187121 | | |
| 10x | 1,068,040 | 0.867690308 | 0.868318033 | 0.000580327 | 0.1761666355 | 0.1759078687 | 0.000224112640 | 0.2928717066 | 0.2925495756 | 0.000279089395 |
| 10x | 1,068,949 | 0.868428792 | | | 0.1757809924 | | | 0.2923804722 | | |
| 10x | 1,069,449 | 0.868834999 | | | 0.1757759782 | | | 0.292396548 | | |
| 7,5x | 922,549 | 0.749491429 | 0.747584694 | 0.001783498 | 0.175189142 | 0.1750689866 | 0.000104117288 | 0.2839959322 | 0.2837009209 | 0.000260972438 |
| 7,5x | 919,858 | 0.747305224 | | | 0.1750124335 | | | 0.2836066398 | | |
| 7,5x | 918,199 | 0.74595743 | | | 0.1750053843 | | | 0.2835001907 | | |
| 5x | 656,711 | 0.533521001 | 0.534207761 | 0.001025954 | 0.1737331607 | 0.1738983936 | 0.000153293548 | 0.2621130983 | 0.2623839077 | 0.000237622661 |
| 5x | 656,950 | 0.533715168 | | | 0.1740359871 | | | 0.2624810866 | | |
| 5x | 659,008 | 0.535387115 | | | 0.173926033 | | | 0.2625575383 | | |
| **Sample 3** | **Variants after intersection** | **Recall** | **Triplicates average** | **SD** | **Precision** | **Triplicates average** | **SD** | **F1 score** | **Triplicates average** | **SD** |
| 20x | 1,232,184 | 1 | | | 1.0000000 | | | | | |
| 15x | 1,166,550 | 0.946762889 | 0.94662797 | 0.000485143 | 0.1780720773 | 0.1781131641 | 0.000046852048 | 0.2997631464 | 0.299814588 | 0.000080294679 |
| 15x | 1,165,720 | 0.946089648 | | | 0.1781032278 | | | 0.2997735064 | | |
| 15x | 1,166,881 | 0.947031374 | | | 0.1781641873 | | | 0.2999071111 | | |
| 10x | 1,068,430 | 0.86717449 | 0.867881798 | 0.000786403 | 0.1762609962 | 0.176207542 | 0.000053960076 | 0.2929726687 | 0.2929391129 | 0.000072469041 |
| 10x | 1,070,346 | 0.868728622 | | | 0.1762085398 | | | 0.2929887223 | | |
| 10x | 1,069,130 | 0.867742283 | | | 0.1761530899 | | | 0.2928559477 | | |
| 7,5x | 922,680 | 0.748951813 | 0.75034534 | 0.001228015 | 0.1751916289 | 0.1752649542 | 0.000063742645 | 0.2839604376 | 0.2841567776 | 0.000170272675 |
| 7,5x | 925,537 | 0.751269221 | | | 0.1752960785 | | | 0.2842639335 | | |
| 7,5x | 924,977 | 0.750814986 | | | 0.1753071552 | | | 0.2842459616 | | |
| 5x | 662,609 | 0.537999567 | 0.537752712 | 0.000823548 | 0.1737701848 | 0.173886721 | 0.000122251225 | 0.26269249 | 0.262796093 | 0.000195609678 |
| 5x | 663,133 | 0.538424601 | | | 0.1740139808 | | | 0.2630217132 | | |
| 5x | 661,172 | 0.536833968 | | | 0.1738759974 | | | 0.2626740757 | | |
| **Sample 4** | **Variants after intersection** | **Recall** | **Triplicates average** | **SD** | **Precision** | **Triplicates average** | **SD** | **F1 score** | **Triplicates average** | **SD** |
| 20x | 1,224,638 | 1 | | | 1.0000000 | | | | | |
| 15x | 1,161,040 | 0.9480679188 | 0.9482677058 | 0.000331301539 | 0.1764903496 | 0.1766272589 | 0.000120243427 | 0.2975832256 | 0.2977876598 | 0.000177637784 |
| 15x | 1,161,061 | 0.9480850668 | | | 0.1767157229 | | | 0.2979043746 | | |
| 15x | 1,161,753 | 0.9486501317 | | | 0.1766757041 | | | 0.2978753791 | | |
| 10x | 1,065,983 | 0.8704474302 | 0.8697223179 | 0.000730060965 | 0.1752967142 | 0.1751209148 | 0.000177649977 | 0.2918239135 | 0.2915395345 | 0.000268100990 |
| 10x | 1,065,107 | 0.8697321168 | | | 0.1749414703 | | | 0.2912913989 | | |
| 10x | 1,064,195 | 0.8689874069 | | | 0.17512456 | | | 0.2915032911 | | |
| 7,5x | 922,070 | 0.7529327034 | 0.7512886801 | 0.00145496311 | 0.1746724477 | 0.1745847612 | 0.000166678836 | 0.2835615954 | 0.283329271 | 0.000295678267 |
| 7,5x | 918,683 | 0.7501669881 | | | 0.1743925429 | | | 0.2829964416 | | |
| 7,5x | 919,417 | 0.7507663489 | | | 0.1746892929 | | | 0.283429776 | | |
| 5x | 656,820 | 0.5363380852 | 0.5374597772 | 0.001221792971 | 0.173224405 | 0.1735391568 | 0.000273469082 | 0.2618707921 | 0.2623640781 | 0.000431838242 |
| 5x | 659,788 | 0.5387616585 | | | 0.1736745362 | | | 0.2626738559 | | |
| 5x | 657,973 | 0.5372795879 | | | 0.1737185291 | | | 0.2625475862 | | |

**Truth set 3**

| Sample A1 | Variants after intersection | Recall | Triplicates average | SD | Precision | Triplicates average | SD | F1 score | Triplicates average | SD |
|---|---|---|---|---|---|---|---|---|---|---|
| 20x | 1697859 | | | | | | | | | |
| 15x | 1578794 | 0.9298734465 | 0.9298877783 | 0.0001216695079 | 0.2399713395 | 0.2399043027 | 0.000137798758 | 0.3814915948 | 0.3814080765 | 0.0001830246106 |
| 15x | 1579036 | 0.9300159789 | | | 0.2399957565 | | | 0.3815344439 | | |
| 15x | 1578625 | 0.9297739094 | | | 0.2397458121 | | | 0.3811981908 | | |
| 10x | 1422972 | 0.8380978633 | 0.8388446862 | 0.001293535225 | 0.2331966465 | 0.2331958738 | 0.0001916365045 | 0.3648699949 | 0.3649397378 | 0.0003459613353 |
| 10x | 1422972 | 0.8380978633 | | | 0.2330038521 | | | 0.364633961 | | |
| 10x | 1426776 | 0.840338332 | | | 0.2333871228 | | | 0.3653152574 | | |
| 7,5x | 1174642 | 0.6918371902 | 0.6923920066 | 0.0008453993778 | 0.22185686 | 0.2220377685 | 0.0002515628708 | 0.3359742282 | 0.3362470909 | 0.0003879741949 |
| 7,5x | 1177236 | 0.6933649967 | | | 0.2223250424 | | | 0.3366912289 | | |
| 7,5x | 1174874 | 0.6919738329 | | | 0.2219314032 | | | 0.3360758154 | | |
| 5x | 783471 | 0.4614464452 | 0.4603680282 | 0.001133806539 | 0.2043199558 | 0.2040985063 | 0.0001987357982 | 0.283230626 | 0.2828145251 | 0.000399051545 |
| 5x | 781816 | 0.4604716882 | | | 0.2040398968 | | | 0.2827779015 | | |
| 5x | 779633 | 0.4591859512 | | | 0.2039356662 | | | 0.2824350479 | | |
| **Sample A2** | **Variants after intersection** | **Recall** | **Triplicates average** | **SD** | **Precision** | **Triplicates average** | **SD** | **F1 score** | **Triplicates average** | **SD** |
| 15x | 1,697,859 | | | | | | | | | |
| 10x | 1,450,944 | 0.8545727295 | 0.8552590841 | 0.0005944005189 | 0.2414801323 | 0.2413415158 | 0.0001200453744 | 0.3765554436 | 0.37645336 | 0.00008840698767 |
| 10x | 1,452,692 | 0.8556022614 | | | 0.2412722076 | | | 0.3764023182 | | |
| 10x | 1,452,692 | 0.8556022614 | | | 0.2412722076 | | | 0.3764023182 | | |
| 7,5x | 1,182,525 | 0.6964800964 | 0.6953233455 | 0.001024830406 | 0.2283431923 | 0.2282333324 | 0.0001799735003 | 0.3439283818 | 0.3436625569 | 0.0002729857857 |
| 7,5x | 1,179,946 | 0.6949611246 | | | 0.2280256328 | | | 0.3433829336 | | |
| 7,5x | 1,179,212 | 0.6945288154 | | | 0.2283311721 | | | 0.3436763553 | | |
| 5x | 762,694 | 0.4492092688 | 0.4477495874 | 0.001464217856 | 0.2066030608 | 0.2065909144 | 0.0002764903943 | 0.2830322204 | 0.2827301468 | 0.0004830685637 |
| 5x | 760,231 | 0.4477586184 | | | 0.2068611314 | | | 0.2829852122 | | |
| 5x | 757,722 | 0.4462808749 | | | 0.206308551 | | | 0.282173008 | | |
| **Sample 2** | **Variants after intersection** | **Recall** | **Triplicates average** | **SD** | **Precision** | **Triplicates average** | **SD** | **F1 score** | **Triplicates average** | **SD** |
| 20x | 1,697,859 | | | | | | | | | |
| 15x | 1,592,638 | 0.9380272449 | 0.9376497106 | 0.0003698396076 | 0.2420499007 | 0.2419926729 | 0.0001454620678 | 0.3848043364 | 0.3847002427 | 0.0002079137465 |
| 15x | 1,591,970 | 0.9376338082 | | | 0.2421008177 | | | 0.3848355498 | | |
| 15x | 1,591,383 | 0.9372880787 | | | 0.2418273003 | | | 0.3844608419 | | |
| 10x | 1,419,331 | 0.8359533978 | 0.8373510403 | 0.001559835105 | 0.2341099275 | 0.2339873299 | 0.000241222811 | 0.3657820705 | 0.3657657706 | 0.0003587468923 |
| 10x | 1,421,220 | 0.8370659754 | | | 0.2337094305 | | | 0.3653991516 | | |
| 10x | 1,424,561 | 0.8390337478 | | | 0.2341426317 | | | 0.3661160897 | | |
| 7,5x | 1,171,749 | 0.6901332796 | 0.6888966241 | 0.001107285695 | 0.2225114351 | 0.2225267137 | 0.0001062716203 | 0.3365220748 | 0.3363921719 | 0.0001421235125 |
| 7,5x | 1,169,077 | 0.6885595329 | | | 0.2224289083 | | | 0.3362403699 | | |
| 7,5x | 1,168,122 | 0.6879970598 | | | 0.2226397977 | | | 0.3364140709 | | |
| 5x | 771,095 | 0.4541572651 | 0.4546559324 | 0.001537293119 | 0.2039934942 | 0.2041484974 | 0.0003091141816 | 0.2815316282 | 0.2817749041 | 0.0005876061971 |
| 5x | 769,860 | 0.4534298785 | | | 0.2039475532 | | | 0.2813480043 | | |
| 5x | 774,870 | 0.4563806535 | | | 0.2045044448 | | | 0.2824450798 | | |
| **Sample 3** | **Variants after intersection** | **Recall** | **Triplicates average** | **SD** | **Precision** | **Triplicates average** | **SD** | **F1 score** | **Triplicates average** | **SD** |
| 20x | 1,697,859 | | | | | | | | | |
| 15x | 1,613,382 | 0.950244985 | 0.950094207 | 0.000348719 | 0.2462803001 | 0.2463331691 | 0.00004591077659 | 0.3911770575 | 0.3912309535 | 0.0000564169939 |
| 15x | 1,612,449 | 0.9496954694 | | | 0.246356219 | | | 0.3912262109 | | |
| 15x | 1,613,547 | 0.9503421662 | | | 0.2463629882 | | | 0.3912895921 | | |
| 10x | 1,436,143 | 0.8458552801 | 0.846348058 | 0.00080255 | 0.2369233322 | 0.23679622 | 0.0001439545428 | 0.3701640377 | 0.370055956 | 0.0002012988786 |
| 10x | 1,438,552 | 0.8472741258 | | | 0.2368254259 | | | 0.3701801285 | | |
| 10x | 1,436,244 | 0.8459147668 | | | 0.236639902 | | | 0.3698237018 | | |
| 7,5x | 1,185,010 | 0.6979437044 | 0.697824338 | 0.00027846 | 0.2250009019 | 0.2246386336 | 0.0003138395694 | 0.3402976991 | 0.3398689779 | 0.0003734290136 |
| 7,5x | 1,185,145 | 0.6980232163 | | | 0.2244656572 | | | 0.3396945903 | | |
| 7,5x | 1,184,267 | 0.6975060944 | | | 0.2244493418 | | | 0.3396146444 | | |
| 5x | 784,609 | 0.4863132228 | 0.461801402 | 0.00050983 | 0.2057648643 | 0.2058569758 | 0.00008565475604 | 0.2891759648 | 0.2862489923 | 0.00253536762 |
| 5x | 784,537 | 0.4620742947 | | | 0.2058718333 | | | 0.2848375885 | | |
| 5x | 783,075 | 0.4612132103 | | | 0.2059342299 | | | 0.2847334234 | | |
| **Sample 4** | **Variants after intersection** | **Recall** | **Triplicates average** | **SD** | **Precision** | **Triplicates average** | **SD** | **F1 score** | **Triplicates average** | **SD** |
| 20x | 1,697,859 | | | | | | | | | |
| 15x | 1,608,992 | 0.947659376 | 0.951407063 | 0.054493768 | 0.2445837875 | 0.2446924715 | 0.0001002030099 | 0.3888168563 | 0.3892671078 | 0.0006385238843 |
| 15x | 1,608,266 | 0.958817035 | | | 0.2447811861 | | | 0.3899978697 | | |
| 15x | 1,609,137 | 0.947744777 | | | 0.244712441 | | | 0.3889865973 | | |
| 10x | 1,431,101 | 0.842885658 | 0.842922567 | 0.000436272 | 0.235338934 | 0.2353092052 | 0.0001055351763 | 0.3679452569 | 0.3679123979 | 0.00008835745888 |
| 10x | 1,431,934 | 0.843376276 | | | 0.2351919942 | | | 0.3678123187 | | |
| 10x | 1,430,456 | 0.842505768 | | | 0.2353966873 | | | 0.3679796179 | | |
| 7,5x | 1,176,767 | 0.693088766 | 0.692080241 | 0.00091336 | 0.2229210063 | 0.2229718272 | 0.00004618902795 | 0.3373414779 | 0.3372799365 | 0.00005561029642 |
| 7,5x | 1,174,654 | 0.691844258 | | | 0.222983225 | | | 0.3372650401 | | |
| 7,5x | 1,173,743 | 0.6913077 | | | 0.2230112503 | | | 0.3372332914 | | |
| 5x | 772,887 | 0.455212712 | 0.45530793 | 0.000184053 | 0.2038349787 | 0.2038220537 | 0.0002331192295 | 0.2815828802 | 0.2815886669 | 0.000190823837 |
| 5x | 773,409 | 0.455520158 | | | 0.2035827408 | | | 0.2814008023 | | |
| 5x | 772,850 | 0.45519092 | | | 0.2040484415 | | | 0.2817823183 | | |

**Table S52 - SVs, SNVs and indels number for all HiFi samples (ds1) after individual variant calling (per-sample calling) compared to joint variant calling.**

| | Full coverage | | | | | | | Coverage downsampled to 5x | | | | | | |
|---|---|---|---|---|---|---|---|---|---|---|---|---|---|---|
| | Mean mapped coverage | SVs | | SNVs | | indels | | Mean mapped coverage | SVs | | SNVs | | indels | |
| | | Per-sample SV calling | Joint SV calling | Per-sample SNV calling | Joint SNV calling | Per-sample InDel calling | Joint InDel calling | | Per-sample SV calling (5x) | Joint SV calling (5x | Per-sample SNV calling (5x) | Joint SNV calling (5x) | Per-sample InDel calling (5x) | Joint InDel calling (5x) |
| **Sample 2** | 24.5 | 106347 | 110487 | 11504129 | 11576741 | 1607890 | 1714966 | 4.9 | 49999 | 72321 | 8449139 | 8502965 | 1040502 | 1116725 |
| **Sample 3** | 19.1 | 104173 | 108893 | 11447507 | 11522595 | 1586883 | 1695578 | 4.9 | 50097 | 72518 | 8463837 | 8519275 | 1048283 | 1123563 |
| **Sample 4** | 20 | 104807 | 108992 | 11456889 | 11528616 | 1586883 | 1691942 | 4.9 | 50982 | 73012 | 8484189 | 8537628 | 1038290 | 1123563 |
| **Sample A1** | 32.3 | 108024 | 112113 | 11557933 | 11631281 | 1617358 | 1724515 | 4.9 | 50898 | 72842 | 8515260 | 8569764 | 1051987 | 1127938 |
| **Sample A2** | 15 | 100080 | 105757 | 11241454 | 11311189 | 1522829 | 1635297 | 4.8 | 50534 | 72763 | 8357696 | 8408522 | 1005652 | 1083174 |
| **Average** | 22.18 | 104686 | 109248 | 11441582 | 11514084 | 1584369 | 1692460 | 4.88 | 50502 | 72691 | 8454024 | 8507631 | 1036943 | 1114993 |

Table S53 - Statistics of phased blocks across chromosomes for all HiFi samples (ds1)

| | Sample A1 | | | | | Sample A2 | | | | | Sample 2 | | | | | Sample 3 | | | | | Sample 4 | | | | |
|---|---|---|---|---|---|---|---|---|---|---|---|---|---|---|---|---|---|---|---|---|---|---|---|---|---|
| Chromosome | Percentage of phased variants | Phased blocks | bp per block median | Percentage of phased chromosome | NG50 | Percentage of phased variants | Phased blocks | bp per block median | Percentage of phased chromosome | NG50 | Percentage of phased variants | Phased blocks | bp per block median | Percentage of phased chromosome | NG50 | Percentage of phased variants | Phased blocks | bp per block median | Percentage of phased chromosome | NG50 | Percentage of phased variants | Phased blocks | bp per block median | Percentage of phased chromosome | NG50 |
| Chr1 | 66% | 311 | 213,563.5 | 77% | 834,960 | 66% | 348 | 205,287 | 76% | 726,281 | 66% | 338 | 193,963 | 79% | 879,834 | 66% | 286 | 263,845 | 79% | 912,214 | 66% | 351 | 160,534 | 80% | 757,281 |
| Chr2 | 66% | 236 | 233,802 | 77% | 951,110 | 66% | 259 | 215,288 | 78% | 641,884 | 66% | 228 | 262,429 | 81% | 779,976 | 66% | 237 | 226,404 | 77% | 790,391 | 66% | 249 | 209,258 | 79% | 835,641 |
| Chr3 | 66% | 216 | 225,433 | 70% | 910,831 | 66% | 227 | 234,265 | 70% | 763,751 | 67% | 206 | 234,999 | 71% | 900,632 | 66% | 177 | 259,506 | 67% | 1,002,443 | 66% | 213 | 230,313 | 73% | 905,571 |
| Chr4 | 66% | 174 | 172,579.5 | 68% | 755,929 | 66% | 168 | 208,416 | 71% | 659,623 | 66% | 170 | 567,891 | 70% | 760,898 | 66% | 193 | 88,373 | 68% | 759,607 | 66% | 186 | 98,614 | 68% | 762,670 |
| Chr5 | 66% | 100 | 359,227 | 75% | 1,129,851 | 66% | 118 | 255,161 | 69% | 1,034,662 | 66% | 118 | 567,891 | 72% | 1,012,043 | 66% | 103 | 348,561 | 74% | 1,132,356 | 66% | 102 | 329,699 | 69% | 1,007,372 |
| Chr6 | 66% | 147 | 109,430.5 | 67% | 823,794 | 66% | 133 | 164,362 | 73% | 731,645 | 66% | 119 | 567,891 | 73% | 1,012,548 | 66% | 129 | 150,505 | 71% | 1,018,918 | 66% | 101 | 299,395.5 | 67% | 907,600 |
| Chr7 | 66% | 60 | 309,834 | 81% | 1,203,398 | 66% | 61 | 328,946 | 79% | 956,099 | 66% | 57 | 567,891 | 84% | 1,302,615 | 67% | 58 | 378,094 | 90% | 1,001,173 | 67% | 65 | 248,802 | 91% | 1,125,131 |
| Chr8 | 66% | 35 | 541,171 | 67% | 1,507,262 | 66% | 42 | 387,913 | 75% | 1,357,220 | 67% | 33 | 411,181 | 73% | 1,724,255 | 66% | 28 | 477,576.5 | 66% | 3,144,292 | 67% | 28 | 516,823.50 | 72% | 1,892,096 |
| Chr9 | 66% | 43 | 420,512.5 | 74% | 1,151,148 | 67% | 34 | 383,116 | 61% | 1,856,639 | 66% | 44 | 377,030 | 79% | 1,090,195 | 67% | 39 | 294,177 | 73% | 1,856,072 | 66% | 40 | 337,680.5 | 68% | 1,535,811 |
| Chr10 | 67% | 27 | 239,246 | 86% | 3,603,366 | 66% | 29 | 448,398 | 93% | 704,232 | 66% | 24 | 363,920.5 | 86% | 2,557,349 | 66% | 30 | 391,963 | 89% | 1,482,761 | 66% | 27 | 354,562 | 93% | 1,723,445 |
| Chr11 | 65% | 28 | 105,361 | 59% | 1,424,791 | 61% | 64 | 32,266 | 49% | 1,376,388 | 61% | 48 | 17,438 | 50% | 2,429,861 | 65% | 36 | 90,822 | 72% | 1,515,607 | 64% | 34 | 103,592 | 56% | 2,752,734 |
| Chr12 | 67% | 15 | 793,697 | 75% | 1,200,103 | 66% | 12 | 875,187 | 65% | 2,825,724 | 67% | 19 | 491,901 | 75% | 1,592,368 | 67% | 17 | 408,881 | 73% | 2,474,732 | 66% | 13 | 491,342 | 54% | 2,759,564 |
| Chr13 | 66% | 27 | 164,238 | 98% | 2,614,645 | 66% | 23 | 220,769 | 98% | 1,673,634 | 67% | 18 | 511,479.5 | 98% | 3,802,422 | 66% | 25 | 360,955 | 98% | 2,845,586 | 67% | 25 | 308,684 | 98% | 1,549,748 |
| Chr14 | 66% | 15 | 349,217 | 72% | 1,526,110 | 66% | 27 | 67,370.5 | 68% | 2,577,482 | 66% | 21 | 163,173 | 78% | 2,422,557 | 66% | 21 | 86,572 | 67% | 1,747,052 | 67% | 19 | 208,276 | 75% | 3,179,446 |
| Chr15 | 67% | 16 | 367,533 | 52% | 1,398,219 | 66% | 18 | 372,144 | 72% | 1,790,394 | 67% | 18 | 567,891 | 77% | 1,430,603 | 67% | 21 | 303,350 | 80% | 2,288,579 | 66% | 15 | 470,793.5 | 45% | 1,821,825 |
| Chr16 | 66% | 17 | 318,092 | 82% | 1,061,623 | 66% | 18 | 421,498 | 73% | 1,057,899 | 66% | 15 | 451,426.5 | 68% | 1,619,977 | 67% | 14 | 308,590 | 76% | 1,744,943 | 66% | 17 | 313,411 | 78% | 1,870,719 |
| Chr17 | 66% | 16 | 455,874 | 82% | 1,049,686 | 66% | 24 | 295,935.5 | 92% | 929,643 | 67% | 20 | 317,479.5 | 91% | 1,211,423 | 67% | 19 | 303,784 | 85% | 1,294,724 | 67% | 16 | 506,413 | 84% | 1,222,207 |
| Chr18 | 66% | 32 | 68,726 | 52% | 1,031,639 | 66% | 29 | 150,597 | 54% | 1,034,040 | 67% | 25 | 147,030 | 49% | 1,794,273 | 66% | 33 | 62,056 | 45% | 1,535,422 | 66% | 33 | 62,379.5 | 61% | 426,488 |
| Chr19 | 66% | 12 | 463,662 | 74% | 1,550,427 | 66% | 9 | 496,575 | 55% | 1,491,144 | 66% | 8 | 471,804 | 28% | 1,549,282 | 66% | 13 | 214,179 | 73% | 1,629,091 | 66% | 12 | 531,682.5 | 77% | 1,543,808 |
| Chr20 | 66% | 12 | 483,477 | 94% | 1,437,644 | 66% | 14 | 273,839 | 95% | 1,243,136 | 66% | 12 | 314,815 | 95% | 2,556,972 | 66% | 12 | 332,500 | 83% | 2,399,158 | 67% | 11 | 647,311.5 | 72% | 1,263,244 |
| Chr21 | 66% | 12 | 392,192 | 67% | 1,100,595 | 67% | 15 | 453,172.5 | 79% | 884,784 | 67% | 17 | 372,330 | 85% | 946,322 | 67% | 12 | 486,739.5 | 66% | 881,760 | 67% | 11 | 319,469 | 66% | 1,097,204 |
| Chr22 | 66% | 25 | 73,456.5 | 79% | 687,221 | 66% | 23 | 74,479 | 77% | 628,955 | 65% | 33 | 26,496.5 | 75% | 656,042 | 66% | 25 | 45,046 | 78% | 823,337 | 66% | 20 | 77,731 | 77% | 866,993 |
| Chr23 | 67% | 15 | 462,603 | 87% | 703,040 | 66% | 15 | 241,122 | 90% | 721,327 | 64% | 15 | 253,555 | 87% | 730,378 | 67% | 17 | 226,115 | 88% | 503,794 | 66% | 13 | 447,870 | 74% | 591,150 |
| Chr24 | 66% | 24 | 92,568 | 73% | 434,358 | 66% | 23 | 96,380 | 67% | 363,109 | 67% | 20 | 121,833 | 66% | 610,210 | 66% | 21 | 110,992 | 83% | 638,703 | 66% | 16 | 103,673 | 70% | 639,056 |
| Chr25 | 67% | 32 | 35,455 | 89% | 503,335 | 66% | 29 | 50,132 | 88% | 778,208 | 67% | 32 | 84,901.5 | 91% | 434,111 | 66% | 33 | 58,916 | 91% | 527,581 | 67% | 25 | 47,709 | 91% | 554,111 |
| Chr26 | 66% | 27 | 62,292.5 | 85% | 456,939 | 66% | 24 | 84,470.5 | 82% | 431,719 | 67% | 20 | 89,047 | 81% | 439,830 | 66% | 30 | 51,933 | 79% | 322,885 | 67% | 22 | 72,261.5 | 82% | 471,779 |
| Chr27 | 66% | 31 | 40,864 | 84% | 448,088 | 66% | 26 | 107,390.5 | 86% | 285,196 | 65% | 37 | 60,096 | 83% | 175,144 | 65% | 41 | 38,831 | 86% | 284,924 | 66% | 29 | 49,396 | 88% | 388,508 |
| Chr28 | 66% | 30 | 44,353 | 58% | 193,684 | 66% | 26 | 46,631 | 58% | 324,935 | 66% | 28 | 91,191 | 79% | 243,899 | 67% | 31 | 44,641 | 77% | 276,298 | 66% | 23 | 58,799.5 | 61% | 374,287 |
| Chr29 | 66% | 15 | 39,613.5 | 77% | 183,376 | 66% | 17 | 28,536 | 72% | 175,997 | 65% | 18 | 16,032 | 79% | 173,055 | 66% | 18 | 26,798 | 77% | 266,996 | 66% | 12 | 32,887.5 | 82% | 398,706 |
| Chr30 | 65% | 16 | 32,192 | 59% | 46,042 | 64% | 14 | 23,773 | 55% | 40,453 | 64% | 16 | 24,437 | 57% | 40,168 | 65% | 16 | 14,561 | 62% | 52,906 | 66% | 14 | 34,811 | 60% | 49,588 |
| Chr31 | 67% | 6 | 33,918 | 10% | 67,821 | 66% | 8 | 40,246.5 | 25% | 81,523 | 66% | 10 | 43,276 | 18% | 48,280 | 70% | 11 | 43,046.5 | 25% | 47,586 | 67% | 4 | 37,876.5 | 9% | 79,904 |
| Chr32 | 65% | 10 | 28,780 | 50% | 25,738 | 68% | 9 | 41,638.5 | 52% | 47,264 | 65% | 9 | 35,745 | 60% | 35,745 | 61% | 10 | 29,064.5 | 55% | 25,244 | 63% | 11 | 33,361 | 67% | 57,976 |
| Chr33 | 53% | 4 | 6,394 | 7% | 0 | 54% | 4 | 9,639 | 3% | 0 | 54% | 7 | 5,432 | 18% | 0 | 55% | 8 | 3,094 | 4% | 0 | 59% | 6 | 6,502.5 | 10% | 0 |
| Chr34 | 84% | 1 | 57,192 | 23% | 106,162 | 67% | 2 | 10,554 | 44% | 44,120 | 80% | 6 | 11,285 | 37% | 8,683 | 81% | 2 | 60,512 | 27% | 103,494 | 54% | 4 | 18,164.5 | 39% | 0 |
| Chr35 | 68% | 4 | 44,775 | 70% | 55,865 | 64% | 4 | 44,621 | 69% | 48,415 | 66% | 8 | 23,473.5 | 60% | 29,019 | 64% | 6 | 31,846 | 63% | 51,734 | 66% | 4 | 61,523.5 | 73% | 87,876 |
| Chr36 | 68% | 5 | 12,578 | 27% | 0 | 62% | 5 | 14,857 | 36% | 0 | 65% | 3 | 27,639 | 19% | 0 | 66% | 4 | 9,376 | 25% | 0 | 66% | 5 | 23,752 | 52% | 10,066 |
| Chr37 | 64% | 3 | 32,251 | 53% | 59,192 | 64% | 3 | 42,191.5 | 58% | 54,473 | 63% | 2 | 41,812.5 | 55% | 67,682 | 64% | 2 | 44,357 | 54% | 55,832 | 62% | 3 | 36,106 | 42% | 32,591 |
| Chr38 | 66% | 2 | 8,086 | 8%% | 0 | 64% | 3 | 632 | 10% | 0 | 70% | 3 | 13,931 | 20% | 0 | 72%% | 1 | 26,481 | 13% | 0 | 0% | 0 | - | - | - |
| Z | 11% | 106 | 4,532 | 2% | 0 | 13% | 89 | 3,663 | 3% | 0 | 11% | 110 | 3,009 | 3% | 0 | 10% | 135 | 1,926 | 2% | 0 | 11% | 109 | 5008.5 | 3% | 0 |
| W | 91% | 160 | 7,150 | 7% | 0 | 87% | 128 | 7,123 | 5% | 0 | 89% | 149 | 6,336.5 | 6% | 0 | 91% | 167 | 6,239 | 6% | 0 | 89% | 151 | 7,190 | 6% | 0 |
